# Genome-targeted enrichment and sequencing of human-infecting *Cryptosporidium* spp.

**DOI:** 10.1101/2024.03.29.586458

**Authors:** NJ Bayona-Vásquez, AH Sullivan, MS Beaudry, A Khan, RP Baptista, KN Petersen, MIU Bhuiyan, B Brunelle, G Robinson, RM Chalmers, EVC Alves-Ferreira, ME Grigg, JC Kissinger, TC Glenn

## Abstract

*Cryptosporidium* spp. are parasites that cause severe illness in vulnerable human populations. Obtaining pure and sufficient *Cryptosporidium* DNA from clinical and environmental samples is a challenging task. Oocysts shed in available fecal samples can be limited in quantity, require purification (biased towards dominant strains), and yield limited DNA (<40 fg/oocyst). Here, we use updated genomic sequences from a broad diversity of *Cryptosporidium* species that have been found to infect humans (*C. cuniculus*, *C. hominis*, *C. meleagridis*, *C. parvum*, *C. tyzzeri*, and *C. viatorum*) to develop and validate a set of 100,000 RNA baits (CryptoCap_100k) with the aim of enriching *Cryptosporidium* DNA from varied samples. Compared to unenriched libraries, CryptoCap_100k increases the percentage of reads mapping to target genome sequences, increases the depth and breadth of genome coverage, and facilitates analyses of genetic variants in many samples, while decreasing overall costs.

## Introduction

Cryptosporidiosis, a disease characterized by mild to severe gastrointestinal symptoms, is caused by protist parasites in the genus *Cryptosporidium*. Infection can lead to adverse health outcomes in vulnerable human populations (infants, the elderly, immunocompromised, and pregnant)^1–6^. An estimated 7.5M global cases of cryptosporidiosis occur in children aged 0–24 months, and ∼133,422 deaths occur annually^7–9^. In children under five years, cryptosporidiosis is the fourth leading cause of diarrheal-related deaths globally^6,10,11^. In the United States, *Cryptosporidium* is the third leading cause of zoonotic gastroenteritis and the leading cause of waterborne disease outbreaks^12^. In low- and middle-income countries, *Cryptosporidium* are the second most common pathogen in infants from 0–11 months^10^.

There are at least 44 recognized species within the genus *Cryptosporidium*^13–15^, with a wide range of host specificity and likelihood of infecting humans. Thus, *Cryptosporidium* spp. vary in public health importance^13^. At the genome sequence level, a cluster of species shares greater than 80% identity across much of their genomes^16–19^, including *C. cuniculus*, *C. hominis*, *C. meleagridis*, *C. parvum*, *C. tyzzeri,* and *C. viatorum*.

Several molecular methods have been developed to gain insight into the genetic diversity of *Cryptosporidium*, often relying on single-marker typing of conserved (e.g., rRNA-SSU) or polymorphic genes (e.g., *gp60* or *hsp70*)^2,20^. Other approaches include multi-locus sequence typing (MLST)^21–24^ and, more recently, whole-genome sequencing (WGS). The disadvantages of MLST include unreliability of markers in different regions due to limited information about the genetic variability of *Cryptosporidium*^24^, and thus, assays are not universal. Challenges with methods such as WGS are constrained by the need to purify oocysts^25^, given the limited amount of target DNA from fecal clinical samples (which also have an abundance of other organisms’ DNA) ^26,27^. These challenges have led to the use of purification methods combined with costly deep sequencing^28,29^.

For example, to obtain sufficient DNA, oocysts may be propagated through animals, which has been shown to change *Cryptosporidium* populations through selection of individual subtypes or recombination^14,30^. Some WGS analyses have relied on whole-genome amplification protocols to overcome low concentrations of target DNA^14,26,27,31^. A helpful alternative that still requires semi-purified oocyst suspensions^4,14,32^, and that may be prone to uneven amplification and the production of chimeric DNA fragments. Other promising protocols for WGS directly from human stool samples do not require whole-genome amplification but rely on a sufficient sample volume and an adequate concentration of oocysts, achieving WGS data for ∼53% of samples^33^, showing medium success rates. Thus, due to the limitations of existing approaches, additional methods that can characterize *Cryptosporidium* genomes from limited amounts of DNA, if possible, directly from fecal samples, which can be implemented easily and reliably, would be beneficial.

Host-derived and environmental samples can also contain more than one species or genotype of *Cryptosporidium* spp.^34–39^. Mixed infections may be limited to specific geographic areas, particularly in endemic regions^40^. Most reported human mixed-species infections contain *C. parvum* and *C. hominis*^35,37,41^; still, mixed infections are more commonly represented by multiple strains of a single species^17^. Their prevalence has been reported to range from 5% in the US^42^ to 12% in Scotland^43^, and up to 33% in Peru^35^. However, their frequency and prevalence by different species or strains are challenging to estimate, given the reliance on limited reference databases and PCR-generated data that may fail to detect low-abundance types/species within mixed infection samples^20^. Thus, methods that enable the detection and characterization of mixed infections are necessary to gain a better understanding of cryptosporidiosis.

Hybridization sequence capture has well-known advantages^44–46^ relevant to studying *Cryptosporidium* spp. and other parasites: it is feasible to design thousands of baits to capture genomes, such as the 9 Mb genome of *Cryptosporidium* spp.; baits can tolerate mismatches (thus regions with < 20% sequence divergence may be captured); and successful enrichment reduces the amount of sequencing needed (thereby reducing sequencing costs in comparison with non-enriched samples). The first *Cryptosporidium* whole-genome bait set (CryptoCap_75k^47^) was designed using the *C. parvum* IOWAII genome assembly^48^ and is reported to perform well in gene-rich regions (∼90%) of the genome. However, in mixed samples comprising other, more distantly related species, the analyses showed a bias towards *C. parvum* enrichment over the other species^47^. Because fecal samples can contain unknown or mixed *Cryptosporidium* species, a second-generation bait set is needed to enrich for a broader spectrum of *Cryptosporidium* species and consider highly polymorphic genomic regions to maximize the potential to capture these parts of the genome.

To ameliorate the current challenges in the genomic study of human-infecting *Cryptosporidium* spp., we developed, validated, and critically assessed CryptoCap_100k. Motivated by CryptoCap_75k^47^, we advance this approach by creating a bait set that also captures highly diverse genomic regions (e.g., including all subtelomeric regions) and a wide diversity of species that have been found to infect humans. CryptoCap_100k was designed (separately from CryptoCap_75k) from the most complete publicly available genome sequences from six *Cryptosporidium* species (Table 1). We validated the sensitivity and specificity of CryptoCap_100k *in silico* and *in vitro* with simulated and mock samples, multiple bait dilutions, and library preparation methods, including enriching pools of sample libraries, rather than single libraries, for time- and cost-efficiency purposes. We tested the reliability of these methods in pure oocysts and in clinical samples and demonstrated that CryptoCap_100k is an efficient and economical method to obtain genomic information for *Cryptosporidium*. These genomic data facilitate inter-sample population genomics analyses and intra-sample genetic variation via deconvolution analyses, opening new opportunities to understand the complex biology of these important pathogens.

**Table 1.**
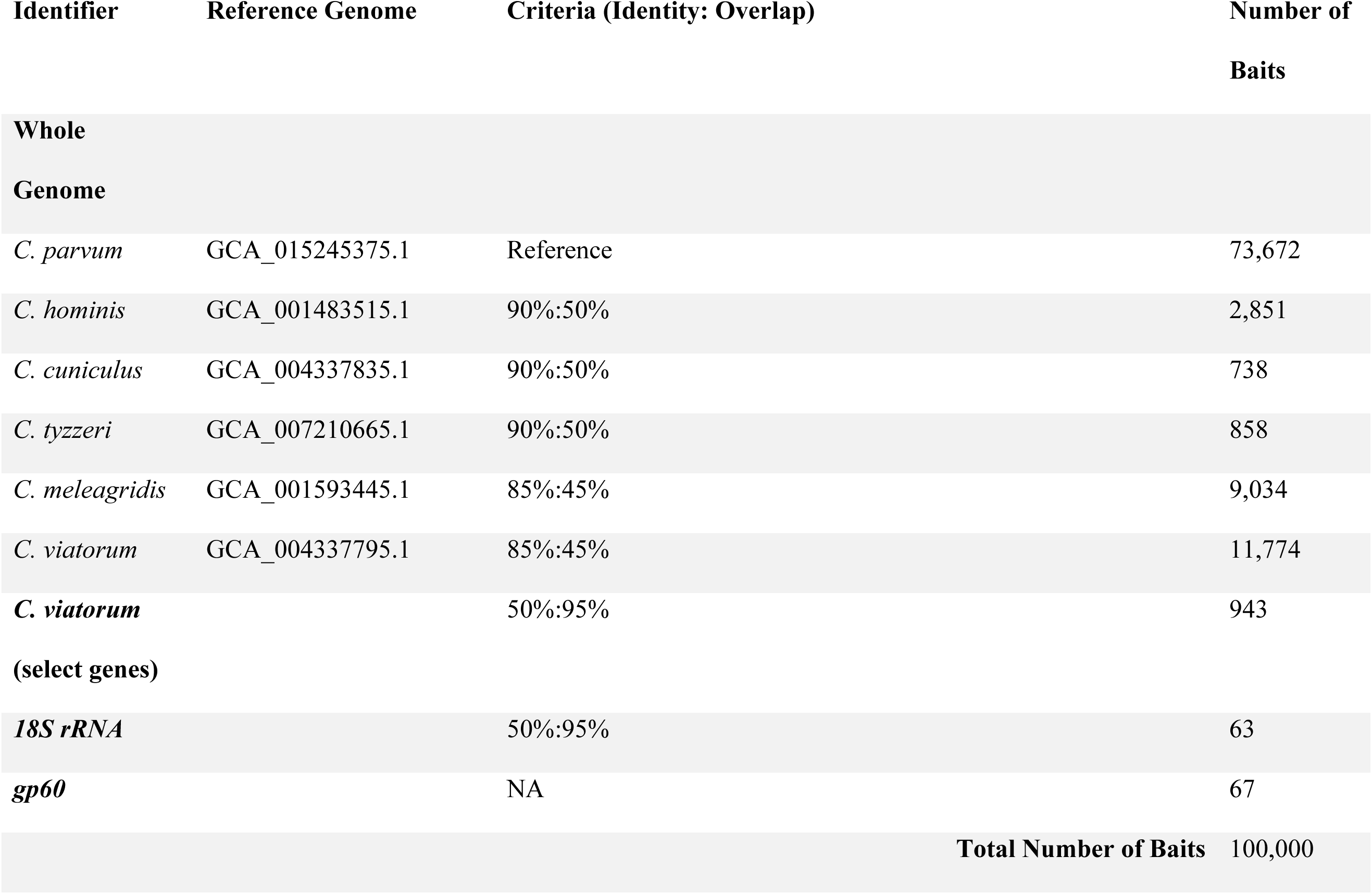
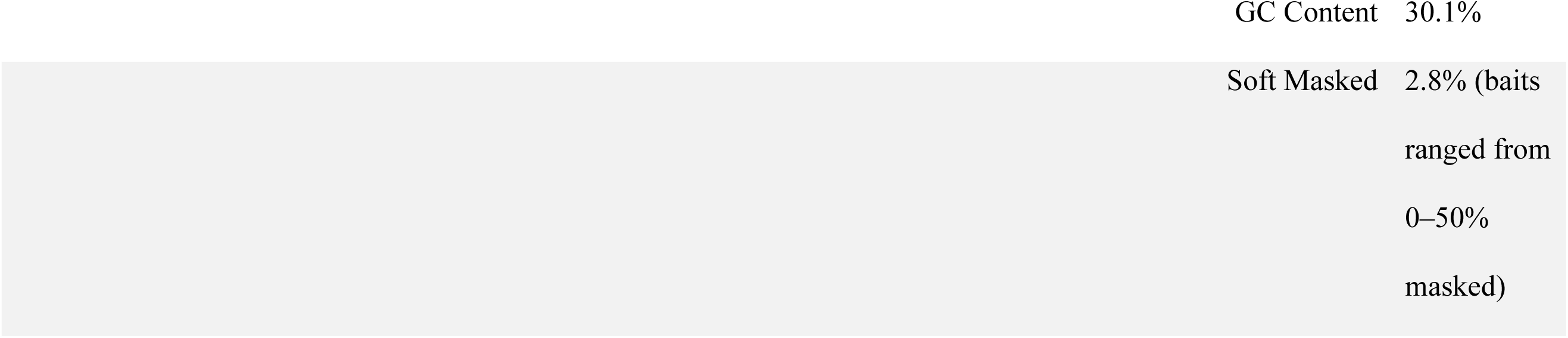
Summary of *Cryptosporidium* species considered, genome version used, and criteria for designing CryptoCap_100k bait set.

**Table 2.**
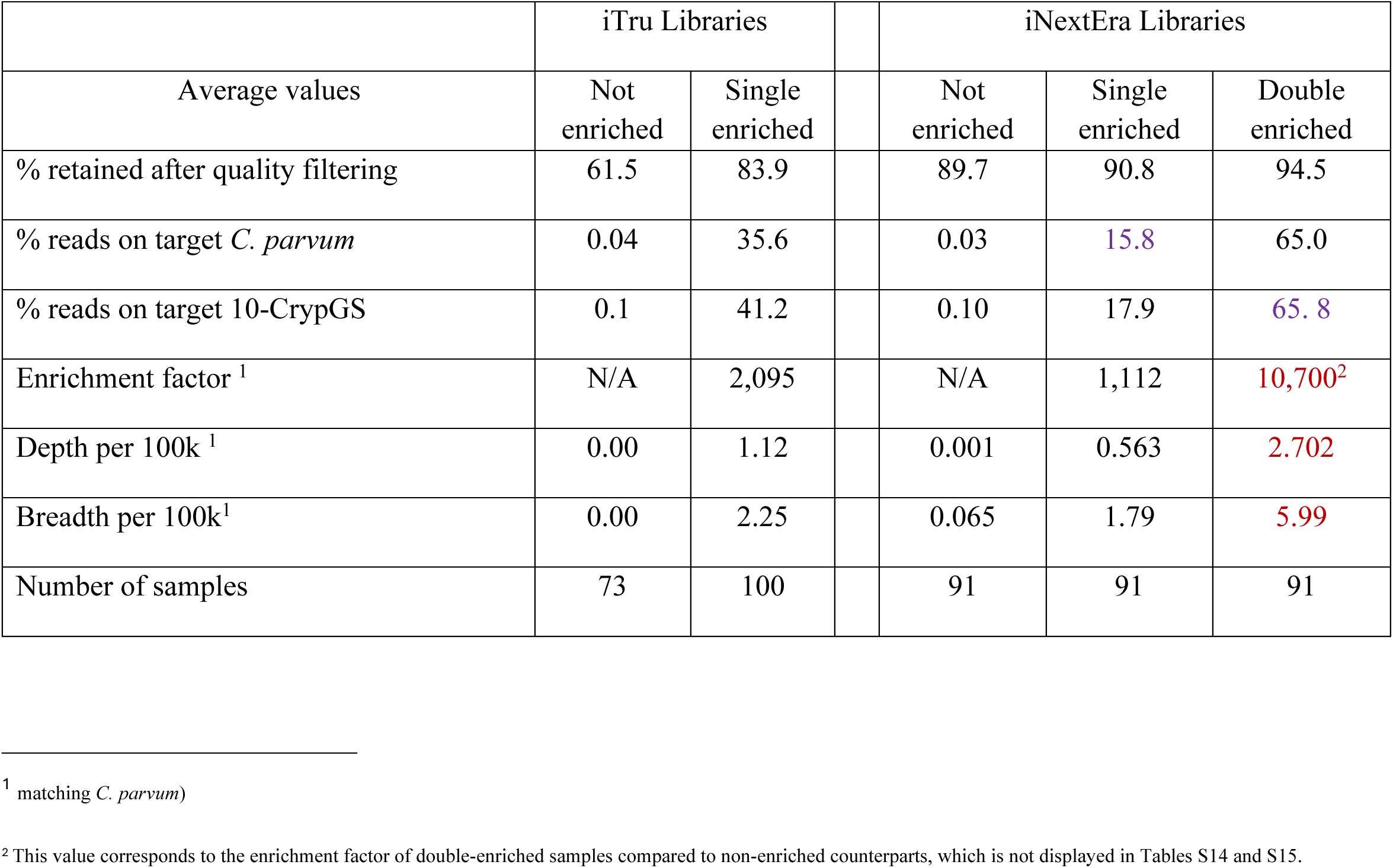
Summary statistics from enriching 100 human fecal DNA samples with two library preparation methods using CryptoCap_100k baits.

|  | iTru Libraries |  |  | iNextEra Libraries |  |  |
| --- | --- | --- | --- | --- | --- | --- |
| Average values | Not enriched | Single enriched |  | Not enriched | Single enriched | Double enriched |
| % retained after quality filtering | 61.5 | 83.9 |  | 89.7 | 90.8 | 94.5 |
| % reads on target *C. parvum* | 0.04 | 35.6 |  | 0.03 | 15.8 | 65.0 |
| % reads on target 10-CrypGS | 0.1 | 41.2 |  | 0.10 | 17.9 | 65.8 |
| Enrichment factor <sup>1</sup> | N/A | 2,095 |  | N/A | 1,112 | 10,700<sup>2</sup> |
| Depth per 100k <sup>1</sup> | 0.00 | 1.12 |  | 0.001 | 0.563 | 2.702 |
| Breadth per 100k<sup>1</sup> | 0.00 | 2.25 |  | 0.065 | 1.79 | 5.99 |
| Number of samples | 73 | 100 |  | 91 | 91 | 91 |

<sup>1</sup> matching *C. parvum*)
<sup>2</sup> This value corresponds to the enrichment factor of double-enriched samples compared to non-enriched counterparts, which is not displayed in Tables S14 and S15.

## Results

### CryptoCap_100k Bait Design & Characteristics

We designed a set of 100,000 120-mer RNA baits targeting six *Cryptosporidium* species, including 73,672 baits for full coverage of *C. parvum* and 4,447 baits to cover divergent regions among *C. hominis, C. cuniculus, and C. tyzzeri*. Additional baits were designed to cover *C. meleagridis* and *C. viatorum*. Finally, we added 130 baits for genes of particular interest (Table 1).

Mapping the baits to twelve *Cryptosporidium* genome sequences (Table S1, Supplementary Notes #1) revealed ∼90.5% breadth of genome coverage for target species, but much lower values for distant species not considered in bait design (7.3% at 0.08X; Table S1; Fig. S1).

When comparing the multi-species design (CryptoCap_100k) to the single-species design (CryptoCap_75k) (Supplementary Notes #2), despite a slightly higher percentage of target reads for CryptoCap_75k than CryptoCap_100k (64.7% vs. 55.6% baits, respectively), both bait sets showed similar depth and breadth of coverage (Table S1; p-values > 0.05; Figs. S2–S3).

### In Silico Simulations of Enrichment and Sequencing

*In silico* simulations exhibited breadth of genome coverage of up to 99.9% and a depth of coverage of 7.6X (for ∼200,000 paired reads), minimal off-target hybridization to more than a thousand non-*Cryptosporidium* genome sequences (including the human genome), and reliable mapping fidelity to the database containing ten genome sequences from ten *Cryptosporidium* species (10-CrypGS), which translated to high accuracy in species assignment for each simulated *Cryptosporidium* species (Supplementary Notes #3–7, Supplementary Tables S2–S7, Supplementary Figures S4–S13).

### In Vitro Simulation of Single Infections and Mixed Infections

DNA from oocysts of *C. parvum* and *C. meleagridis* (Supplementary Notes #8; Table S8) showed improved read mapping (95.8% vs. 68.1%, p-value = 0.094; Fig. S14), similar breadth (85.2% vs. 81.6%, p-value = 0.75), and greater depth of coverage in enriched vs. unenriched libraries (17.3X vs. 5X, respectively; p-value = 0.01; Fig. S15–S16). Species assignment was largely accurate for both library types, with a low proportion of reads from *C. parvum* oocysts assigned to *C. cuniculus* and *C. hominis* (Fig. S17; Table S9).

The proportion of reads obtained for each species (*C. parvum* and *C. meleagridis*) from the unenriched mock mixed infection sample (Supplementary Notes #9; Table S10) showed that initially the mixture was unequal (60.6% *C. parvum* and 39.4% *C. meleagridis*; Fig. 1A) and enrichment slightly increased the bias (64.3% *C. parvum* and 35.7% *C. meleagridis*; Fig. 1A; Table S10). In total, 95.7% and 81.4% reads mapped to 10-CrypGS, for enriched and unenriched libraries, respectively (p-value = 0.50), with very accurate species assignment for both library types (> 97%; Fig. 1B). Species assignment with *gp60* performed well for *C. parvum*, but less accurately for *C. meleagridis* (Fig. 1C; Supplementary Notes #9). Species assignments with *18S rRNA* were ∼70% correct (Fig. 1D; Supplementary Notes #9).

**Figure 1.**
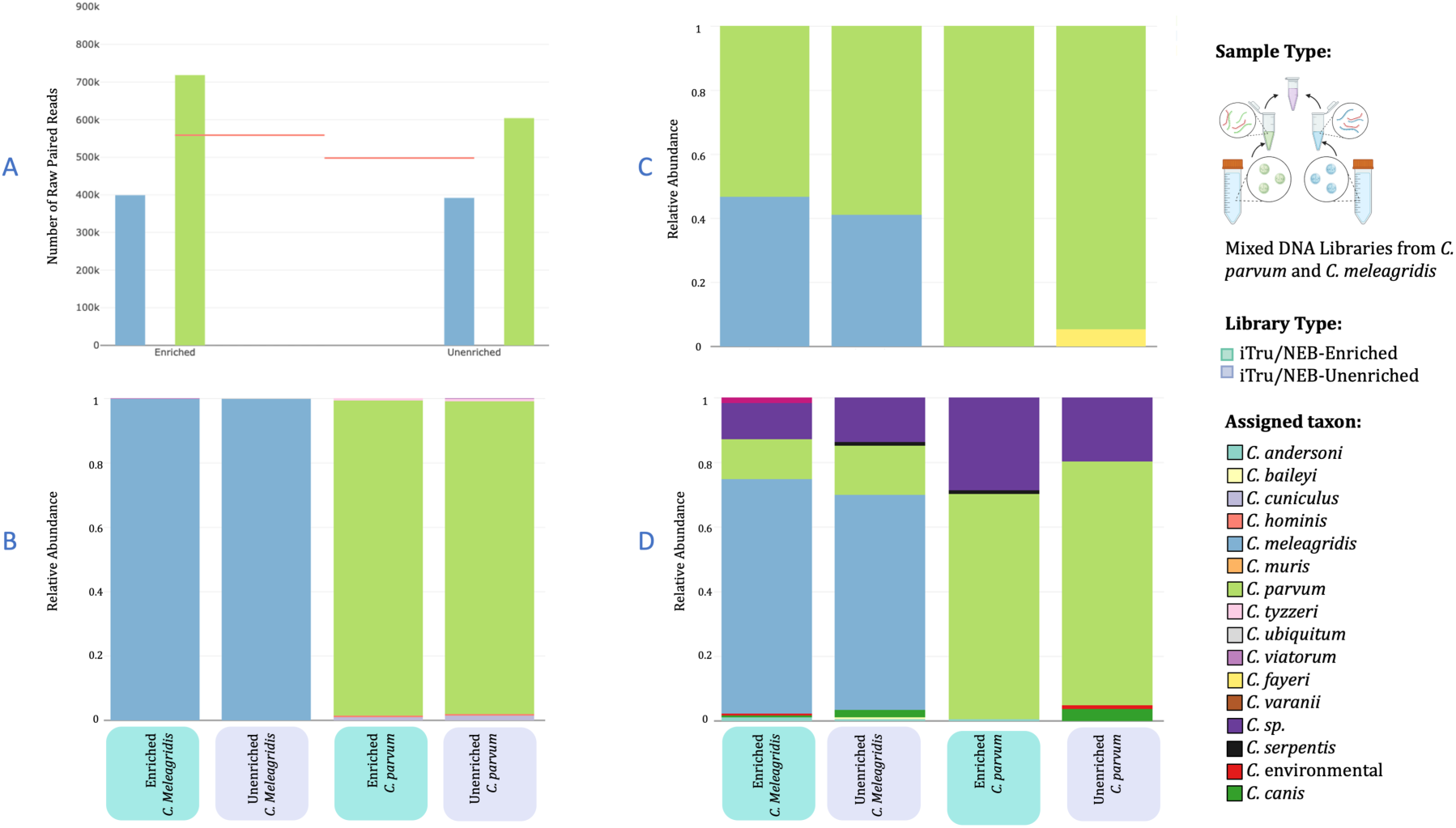
Species assignment plots based on mapping reads from unenriched and enriched libraries for one mock mixed-infection sample. (A) The number of raw paired reads obtained for each species in the mix, *C. parvum* in green and *C. meleagridis* in blue, for both library types. The pink lines indicate the expected number of reads for each sample for a 50-50 proportion. Relative abundance of species assigned from mapping the sequence reads to (B) the ten-genome sequence reference database (10-CrypGS), (C) the *gp60* database, and (D) the *18S rRNA* database.

### LOD: In Vitro Sensitivity Test with Different Inputs of C. parvum DNA

Dilutions of target DNA allowed the determination of the limit of detection (LOD) in a background community. For NEB-iTru libraries (Supplementary Notes #10.1, Table S11), the percentage of reads mapped to *C. parvum* correlated with the initial concentration of target DNA, with inputs below 0.0001 ng having very low percentages, with differences between 1 ng and ≤ 0.01 ng, and 0.1 ng and ≤ 0.01 ng being statistically significant (p-values ≤ 0.040; Fig. S18). The average depth of genome coverage was 4X for ≤ 0.01 ng, 29X for 0.1 ng, and 126X for 1 ng, with significant differences between 1 ng and all other dilutions (p-values < 0.0001; Fig. 2A; normalized values Fig. S19). The breadth of genome coverage was significantly different between comparisons below and above 0.01 ng (p-values ≤ 0.02; Fig. 2A; Supplementary Notes #10.1; normalized values Fig. S19). For unenriched libraries, accurate species assignment is observed only for 1 ng and 0.1 ng inputs (Fig. 2B). For enrichments, *C. parvum* is accurately detected in inputs ≥ 0.001 ng (Fig. 2B).

**Figure 2.**
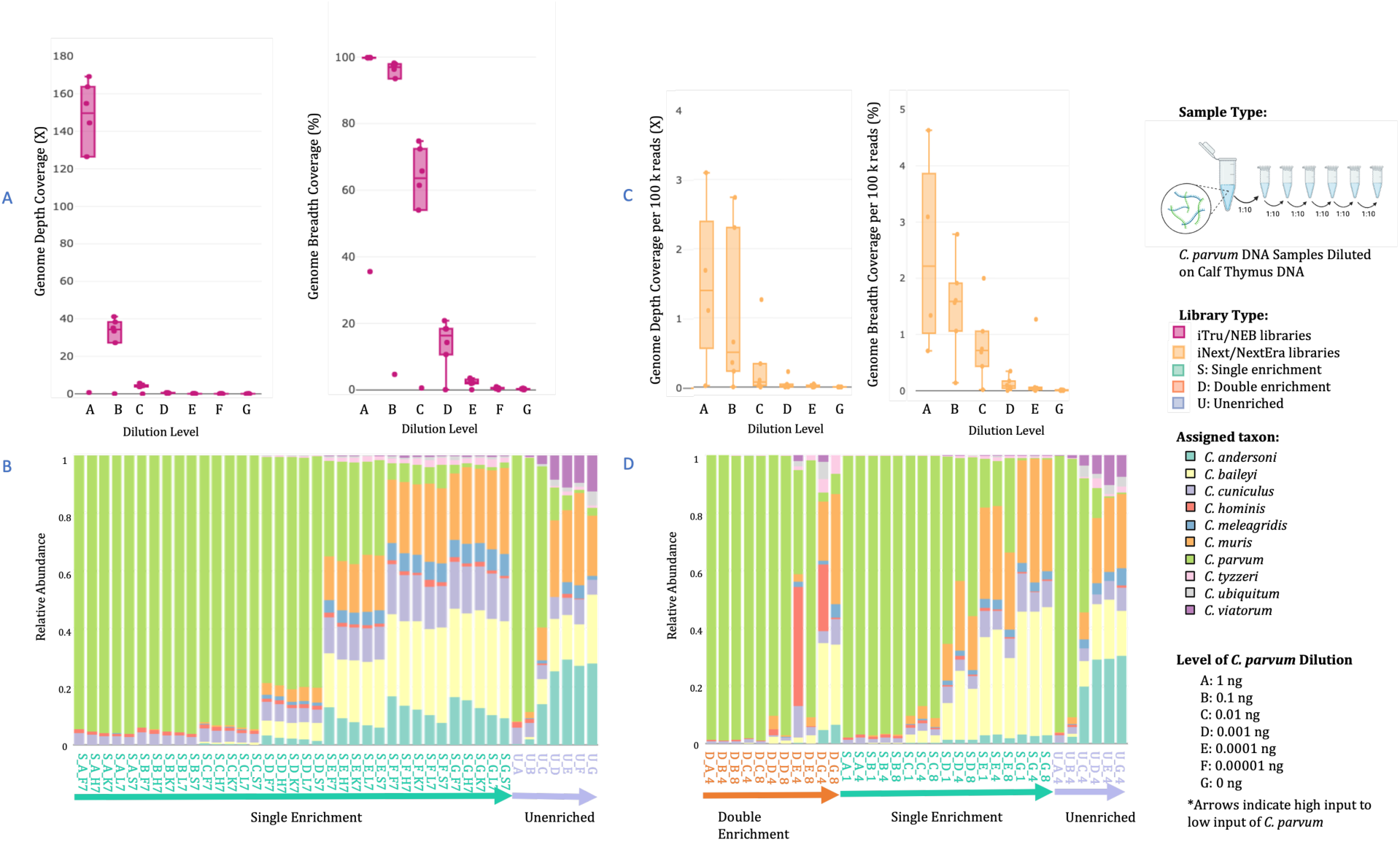
Summary statistics for sequence data from serially diluted *C. parvum* samples to assess detection levels given different input amounts. (A) *C. parvum* genome depth and breadth of coverage for diluted DNA samples prepared with NEB/ iTru libraries (B) species assignment of diluted *C. parvum* samples prepared with NEB-iTru protocol and unenriched (*U*) or single-enriched (*S*) mapped to the ten *Cryptosporidium* genome sequences reference (10-CrypGS); (C) *C. parvum* genome depth and breadth coverage for diluted DNA samples prepared with iNext/NextEra libraries, (D) species assignment of diluted *C. parvum* samples prepared with iNextEra protocol and unenriched (*U*), single-enriched (*S*), or double-enriched (*D*) mapped to the ten *Cryptosporidium* genome sequences reference (10-CrypGS). In the species assignment plots, the first letter in the sample name corresponds to the library type, and the second letter corresponds to the level of input of DNA (A*–*G). The direction of the arrow indicates high to low input (A → G).

For iNextEra libraries (Supplementary Notes #10.2; Table S12, Fig. S19), the average percentage of reads mapped increased with target input in enriched libraries (Fig. S20). The percentage of mapped reads was lower for inputs of ≤ 0.01 ng vs. 1 ng (p-values ≤ 0.015) and for inputs ≤ 0.001 ng vs. 0.1 ng (p-values ≤ 0.05). Depth and breadth of genome coverage tended to increase as target DNA increased (Figs. 2C and S21).

Although depth did not show statistically significant differences among dilutions (normalized values per 100,000 reads did: Supplementary Notes #10.1, Fig. S21) for libraries with ≤ 0.1 ng we recovered depths ≤ 4X whereas for those with 1 ng average depth was 34.8X (Fig. 2C). Breadth of genome coverage between the two highest input samples (i.e., 1 ng and 0.1 ng, mean 57%–58%) differed from all other input samples (p values ≤ 0.017, Fig. 2C).

The average percentage of reads mapped to the *Bos taurus* genome sequence (DNA used as background diluent for *C. parvum*) decreased as *C. parvum* input increased in enriched libraries. With mapped reads only being significantly different for inputs of 1 ng vs. all the other input values (0.004 ≤ p-values ≤ 0.021). In unenriched libraries, mapping percentages to *Bos taurus* reference were high across all dilutions, averaging 90.5% (Table S12, Fig. S22; Supplementary Notes #10.1).

Accurate species assignment was obtained in unenriched samples with inputs ≥ 0.1 ng, in single-enriched samples with ≥ 0.01 ng, and in double-enriched samples consistently with ≥ 0.001 ng, and less consistently in samples of 0.0001 ng (Fig. 2D).

### LOD: In Vitro Sensitivity Test with Different Dilution Levels of Baits

Bait dilutions enabled the determination of the minimum quantity that could be used in a capture reaction. For both library types, no difference was observed among bait dilutions (NEB-iTru: full down to one-sixteenth; iNextEra: one-quarter vs. one-eighth) in the number of reads obtained, the percentage of reads mapped to *C. parvum,* and the depth and breadth of genome coverage (Supplementary Notes #11.1–11.2; p-values ≥ 0.05; Fig. S23–S29; Table S11).

### Enriching Seven Human Fecal DNA Samples with the CryptoCap_100k Bait Set

From seven clinical patients’ samples (Supplementary Notes #12.1, Table S13, Fig. S30), the percentage of reads mapped increased from 0.02–0.15% (unenriched) to 0.4–18.2% (enriched), with a fold change of 9.25–479X (Table S13, Fig. S31, p-value = 0.02). Species assignment revealed that in unenriched libraries, the few mapped reads were assigned to non-human infecting *C. muris* (Fig. S32 right), similar to the background effect observed in the LOD experiment (Supplemental Notes #12.2). In contrast, in enriched libraries, DOCK and UKH101 are predominantly assigned to *C. hominis* and *C. cuniculus*, while EC4 and UKP196 to *C. parvum* (Fig. S32 left). No reads mapped to the *gp60* database, and very few mapped to the *18S rRNA* database with assignment to species not common in human infections (Fig. S33).

### Enriching 100 Human Fecal DNA Samples with the CryptoCap_100k Bait Set

#### NEB-iTru libraries

For fecal DNA obtained from 100 patients when using the NEB-iTru protocol, we observed a significantly higher proportion of enriched vs. unenriched retained reads mapped to the *C. parvum* genome sequence (p-value < 0.0001), increasing enrichment by an average factor of 2,095X, and increasing significantly the breadth and depth of coverage (p-values < 0.0008). In enriched libraries, qPCR C_T_ values were a good predictor of the percentage of mapping reads, but not in unenriched libraries (Supplementary Notes #12.2.1, Table S14; Fig. S34–S37).

Species assignments for unenriched libraries resemble the complex background from LOD experiments (cf. Figs. 2B and S38). Following enrichment, most hits matched *C. parvum*, but many samples have moderate proportions of reads assigned to other species, including those with limited evidence to infect humans (Fig. S38; see Supplementary Notes #12.2.2).

#### iNextEra Libraries

For the 91 clinical iNextEra libraries (Supplementary Notes #12.2.2, Tables 2 and S15; Fig. S39), on average, 0.03% of unenriched reads, 15.8% of single-enriched reads, and 65.0% of double-enriched reads mapped to the *C. parvum* genome sequence (Supplementary Notes #12.2.2, p-values < 0.0001, Fig. 3A, Table S15, Fig. S40), yielding an average enrichment factor of 1,112X from unenriched to single-enriched, and an extra 9.24X from single- to double-enriched (Table S15). The mean depth of genome coverage was significantly higher for double-enriched samples (95.5X) than for single-enriched libraries (32.1X) and unenriched libraries (0.05X) (p-values < 0.005; Fig. 3B; normalized values per 100,000 reads in Supplementary Notes #12.2.2, and Fig. S41, Table S15). The mean breadth of genome coverage was significantly higher for double-and single-enriched samples (48% and 54%, respectively) than for unenriched samples (2.6%, p-values < 0.0001).

**Figure 3.**
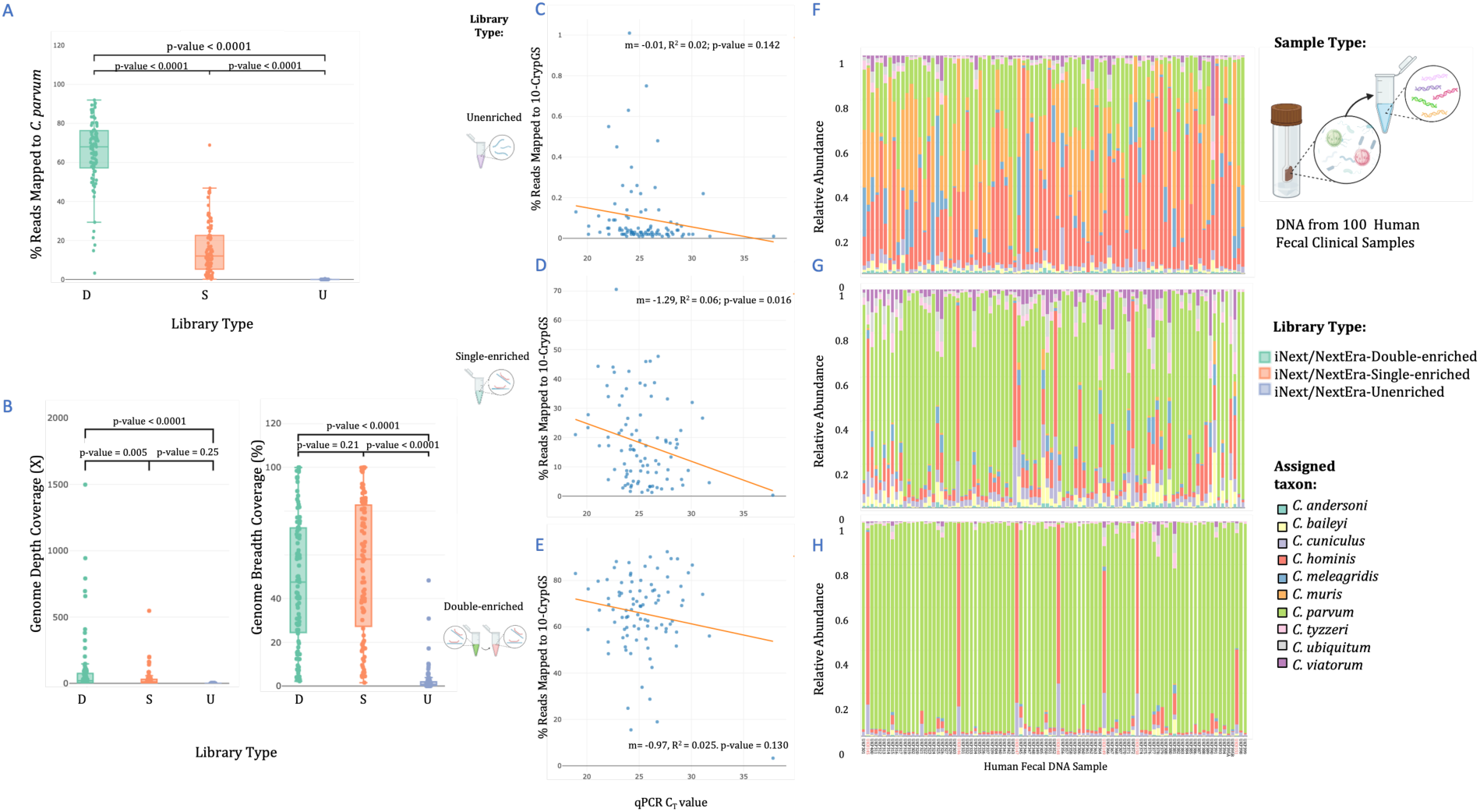
**Summary statistics for sequence data from 91 clinical samples prepared with the iNextEra protocol and iNext primers, comparing libraries enriched with CryptoCap_100k bait set versus unenriched libraries for validation**. (A) Boxplot of percentage of reads mapped to *C. parvum* from double-enriched [D], single-enriched [S], and unenriched libraries[U]. (B) Genome depth and breadth of coverage for each sample in double-enriched, single-enriched, and unenriched libraries. Linear regression model for predicting the percentage of mapped reads to a reference database containing ten *Cryptosporidium* genome sequences (10-CrypGS) given a C_T_ value from qPCR for (C) unenriched libraries and (D) single-enriched libraries with CryptoCap_100k, and (E) double-enriched libraries with CryptoCap_100k. Species assignment of each sample prepared with iNextEra protocol with (F) no further enrichment, (G) single-enriched with CryptoCap_100k, and (H) double-enriched with CryptoCap_100k; when mapped to a reference database containing ten *Cryptosporidium* genome sequences (10-CrypGS). Sample names for plot 4F are not shown.

The percentage of reads mapped was negatively correlated with qPCR C_T_ values for all library types (Figs. 3C–3E). In unenriched and double-enriched libraries, the linear model fails to detect C_T_ values as predictors of percentage of reads mapped (p-value > 0.05; Figs. 3C and 3E), but in single-enriched libraries qPCR C_T_ has a significant but small effect on percentage of reads mapping (R^2^ = 0.006, p-value = 0.016), that is for every increase of one qPCR C_T_ unit, the percentage decreased by 1.3% (Fig 3D). Double-enrichments allowed for the retrieval of a higher percentage of target reads, even at higher C_T_ values, than the other two library types (Figs. 3C–3E).

Species assignments were similar to those from NEB-iTru libraries in unenriched and single-enriched libraries (cf. Figs. S38 vs. 3F and 3G). Double-enriched libraries compared to single- and unenriched libraries exhibit an increased proportion of hits to the most frequently observed human-infecting species (*C. parvum* and *C. hominis*; Fig. 3H). Enriched libraries mapped most frequently along either *C. parvum* chromosomes for samples labeled as UKP or *C. hominis*-*C. cuniculus* chromosomes for samples labeled as UKH (Fig. S42), aligning with the species identified in the multi-locus typing study in the same samples^49^.

#### iNextEra vs. NEB-iTru Libraries

iNextEra and NEB-iTru libraries performed similarly (Tables 2 and S16) but with noteworthy differences. In single-enriched libraries, NEB-iTru samples obtained and retained more reads following quality filtering, and a higher percentage of reads mapped to the *C. parvum* genome sequence than iNextEra samples. The average depth of coverage is similar for both protocols (p-value = 0.7); however, the breadth of coverage was significantly higher for iNextEra libraries (p-value < 0.0001; Supplementary Notes #12.2.3, Table S16; Fig. S43–S47).

### Population Genomics Analyses of Clinical Samples

Enrichment drastically improved SNP recovery: obtaining 94,783–127,102 SNPs for double- and single-enriched vs. 15,430 for unenriched in raw VCF files (Figs. S48–S53), with large variations in read depth (DP; Fig. S54Supplementary Notes #13.4). Filtered VCF files contained: 76 and 77 samples, and 12,498 and 13,578 SNPs for single- and double-enrichments, respectively (Figs. S55–S58). For unenriched libraries, filtering was based on read depth and biallelic variants only, given that missing data filtering would have removed all variants and all samples (except for three), yielding 91 samples and 11,226 SNPs (Figs. S59–S60; Supplementary Notes #13.4).

Population genomics analyses (Supplementary Notes #13.5), such as principal component analysis (PCA), phylogenetic networks, genetic distance trees, and Structure, are highly concordant in enriched datasets. Both datasets suggest hierarchical structure (K = 2; Figs. 4, S61–S70), with distinct clustering of UKP (*C. parvum*) and UKH (*C. hominis*) samples, aligning with the species assignment analyses from above and previous multi-locus typing results^49^ (cf. Figs. 3H and 4A–4B). The sparse variant database obtained from unenriched libraries was insufficient for species assignment (Fig. 4C).

**Figure 4.**
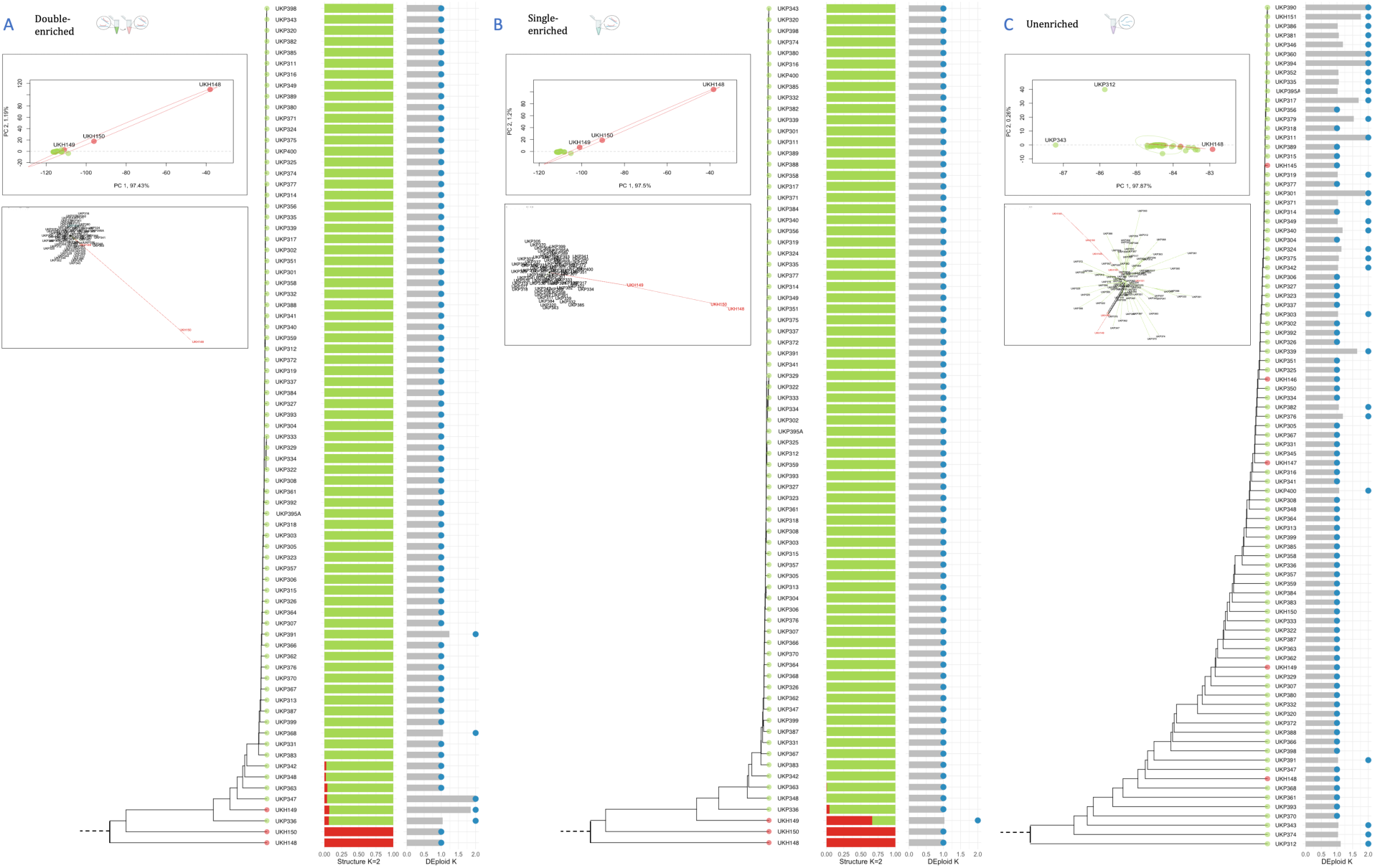
Population genomic analyses from called SNP variants of clinical samples prepared with the iNextEra protocol and iNext primers. Libraries were double-, single-enriched, and unenriched, respectively (A, B, and C). Each panel includes: Principal Components Analyses, phylogenetic distance networks, UPGMA Phylogenetic trees of Nei’s genetic distances, assignment plots from Structure analyses for K=2 (selected based on Evanno’s method—unenriched samples could not be analyzed due to high levels of missing data), and effective (gray bars) and inferred (blue circles) number of strains (K) detected in each sample by the DEploid software.

Minor distinctions between single- and double-enriched libraries are evident in sample

UKH149, which is positioned in the trees as more closely related to *C. parvum* than to *C. hominis*. However, in double-enriched libraries, UKH149 clusters much closer to all other *C. parvum* samples than in single enrichments (Figs. 4A and 4B).

High pairwise *F_ST_* values between UKP and UKH samples were estimated in enriched (0.94 in double vs. 0.98 in single enrichments), but not in unenriched libraries (0.08).

These values are balanced across all chromosomes, showing no locus bias (Figs. S71–S73). Pairwise genetic distances within samples of *C. parvum* reveal two to three clusters in enriched libraries (Fig. S74–S75Supplementary Notes #13.5). For *C. hominis*, enrichments reveal that the three samples are highly distinct from each other, even more than the differences shown between clusters of *C. parvum* (Figs. S74–S75). Genetic distances using unenriched libraries fail to reflect any biologically meaningful pattern (Fig. S76).

Deconvolution analyses (Supplementary Notes #13.6.1) revealed different numbers of inferred strains (inferred K) among library types (Figs. 4A–4C). Sample UKH149 was detected as having two strains by both enrichment methods (Table S17; Fig. 4B), and two more samples were detected as having two strains for double-enriched libraries (UKP347 and UKP391) with effective Ks above 1.2 and a proportion of the least dominant haplotype larger than 10% (Table S17; Fig. 4C, Supplementary Notes #13.6.1). Three samples (UKH149, UKP347, and UKP391) exhibit the most differences between capture methods. When assessing allelic read depth (AD) and genotype quality (GQ), UKH149 and UKP347 had statistically larger AD and GQ values in double-enrichments compared to single-enrichments (p-values < 0.0001; Figs. S77–S80, Table S18–S19). Sample UKP391 had similar AD (p-value = 0.64; Fig. S81), but GQ was statistically larger in double than in single enrichments (p-values < 0.0001; Fig. S81, Table S20; Supplementary Notes #13.6.2–13.6.3).

Genetic diversity, measured when ploidy was set to 2 in GATK (Supplementary Notes #13.6.4) revealed that the nucleotide diversity (π) of polymorphic sites within UKH samples (considering UKH149: single-enriched mean π = 0.066; double-enriched mean π = 0.335; not considering UKH149: single-enriched mean π = 0.005; double-enriched mean π = 0.0204) and within UKP samples (single-enriched mean π = 0.0122; double-enriched mean π = 0.0187) was statistically different between double- and single-enrichments (p-values < 0.0001); and also between UKH and UKP samples (p-values < 0.0002; Fig. S82). High levels of within-sample allele variation were observed in samples with high values of effective K in DEploid (Supplementary Notes #13.6.5). These values were consistent among chromosomes, showing no locus bias (Figs. S83–S88). Double enrichments showed a higher mean of within-sample allele variation compared to single enrichments (0.009 vs. 0.002), whereas unenriched libraries showed a mean within-sample allele variation of 1 for all samples (Fig. S88).

## Discussion

### Development and Assessment of CryptoCap_100k

Obtaining sufficient *Cryptosporidium* DNA for genomic studies has been a challenge in the quest to understand its global diversity and its links to transmission, infection, pathogenesis, and host response. Here, we developed and evaluated a hybridization capture system to obtain *Cryptosporidium* genomic data from limited amounts of input DNA and evaluated its performance in a variety of settings.

With a foundation similar to the previous whole-genome sequencing bait capture for

*C. parvum* (CryptoCap_75k)^47^, CryptoCap_100k used more complete genome sequences that included the 16 subtelomeric regions (i.e., *C. parvum* IOWA-ATCC^50^) and included additional baits to account for variation among *Cryptosporidium* species that commonly infect humans. Performance between CryptoCap_75k^47^ and CryptoCap_100k is similar *in silico* (Table S1, Figs. S1–S3); however, CryptoCap_100k allows pooling ∼10 libraries before enrichment, reducing processing costs and time. To make recommendations about how to use CryptoCap_100k best, we defined two distinct limits of detection (LOD) for: i) minimum target genomic equivalents and ii) minimum bait concentration. Given that a single haploid sporozoite contains ∼9 fg of DNA (with four sporozoites per oocyst) and is typically found in a complex community, we diluted *C. parvum* DNA in a consistent background of calf DNA and determined the data quality threshold. Enriched libraries exhibit a high percentage of target reads, substantial breadth and depth of genome coverage, and accurate species assignment with inputs as low as 0.01 ng of target DNA (∼1,000 sporozoites or ∼250 oocysts equivalents; Figs. 2A–2B), but higher input of target DNA significantly reduces non-target reads (Fig. S22). Non-target reads are likely caused by non-specific interactions of beads and/or biotin with background DNA when target DNA is low^51^. These parameters improved further in double-enriched iNextEra libraries with 0.001 ng of target DNA (∼100 sporozoites or ∼25 oocyst equivalents; Fig. 2D), highlighting that double-enrichments should be prioritized for samples with very low target DNA. These findings validate that CryptoCap_100k can obtain genomic information at low DNA levels without purifying oocysts. The LOD assay with bait dilutions suggests that there are no differences in estimated parameters between bait dilutions, offering a cost-effective strategy for large-scale and effective use.

### Validating CryptoCap_100k

Experiments based on simulations and on DNA from purified oocysts demonstrate that CryptoCap_100k retains nearly the whole genome of the target species with minimal bias (breadth of genome coverage: ∼95% *in silico*, ∼85% from pure oocysts; Figs. 1, S15–S17). However, obtaining pure oocysts requires increased effort and time spent per sample, and via propagation, strains may be altered ^14,30,32^. Thus, we aimed to eliminate the need for purifying oocysts to obtain usable genomic data.

We also successfully tested CryptoCap_100k on a small initial set of seven clinical samples. Two of these had sufficient genome-wide read coverage and were also analyzed with CryptoCap_75k^47^, which identified UK101 as *C. hominis* and UK196 as *C. parvum*^47^, both of which agreed with our analyses (Fig. S32).

When validating the baits on fecal DNA from 100 clinical samples, CryptoCap_100k substantially increased on-target reads relative to unenriched libraries (>437X fold change) for both NEB-iTru and iNextEra protocols. While NEB-iTru libraries still performed better than unenriched libraries, with a high percentage of reads mapping to target genomes, iNextEra libraries achieved greater genome coverage breadth at ∼50%.

Several studies recommend sequencing only samples with C_T_ values < 16^31,52^, however our set of 100 clinical samples had an average C_T_ value of 25.4. Still, we were able to recover significantly more target reads, higher averages of breadth and depth of genome coverage, and better accuracy of species assignment in enriched protocols (Fig. S38; Fig. 3; Tables S14–S15). Double enrichments consistently resulted in significant increases of target reads and depth of genome coverage (Figs. 3A–3B), as well as more accurate species assignment than single-enrichments (Figs. 3F–3H). The presence of some *C. cuniculus* hits in samples characterized by the multi-locus system as *C. hominis* (and similar results from our *in silico* tests) underscores more likely the high genetic identity between these two species^53^, instead of representing mixed infections, particularly given that *C. cuniculus* prevalence in the UK (where our clinical samples come from) has been reported to be very low (1.2%)^54^

### CryptoCap_100k Applicability: Enables the Study of Population Genomics

CryptoCap_100k is a useful tool for SNP calling, enabling robust comparative analyses of genetic variation. CryptoCap_100k enabled the acquisition of genomic data for downstream analyses in 76%–85% of the clinical samples used in this study. Enrichment methods resolved species-level (*C. parvum* and *C. hominis*) clustering, which matched the multi-locus typing results^49^ (Figs. 3H, 4A–4B).

Beyond detecting different species, we were able to assess the diversity within each species. Enrichments show that *C. parvum* haplotypes in this study are highly similar and have low nucleotide diversity. Still, two clusters within the species are detected (Figs. S70 and S75), which may correspond to two circulating but highly similar variants, or samples with some levels of mixed infections; however, the latter was not confirmed by our deconvolution analyses. Genomic studies have revealed two distinct clusters for *C. parvum* in the UK^26^. Our samples may represent two less divergent lineages within just one of these clusters, highlighting the prevalence of clonality and the dominance of some lineages in the region (Figs. 4A and 4B).

We also identified two distinct clusters of *C. hominis* separated by the first PC axis, which explains 56–57% of the variance (UKH148 + [UKH149 and UKH150]; Figs. 4A–4B). Two subspecies of *C. hominis* have been detected in the UK, *C. h. hominis*, and *C. h. aquapotentis*, with some evidence of gene flow and recombination between them in the region^27^. The highly divergent *C. hominis* samples in our study may reflect the two subspecies, given their comparable levels of variation. However, explicit comparative analyses with samples of known lineages for both species are warranted.

Given that more than 90% of cryptosporidiosis cases in humans are caused by *C. parvum* and *C. hominis*^55^, our clinical results were representative. Because ∼74% of the baits in our set match to *C. parvum*, it is relevant to assess CryptoCap_100k performance for other species. In our study, we demonstrate that CryptoCap_100k enables the collection of genomic data for *C. hominis* in clinical samples, from which only ∼3% of the baits had been designed specifically, but we expected a larger set to work, thanks to the > 97% percent similarity with *C. parvum* genome sequences^56^ and the mismatch tolerance of the baits. Our evidence suggests that CryptoCap_100k works appropriately in other species: i) *in silico* analyses reveal proper hybridization of our baits to other target species (Fig. S7); ii) *in vitro* tests using pure oocyst of *C. meleagridis* show proper species assignment and a high percentage of target reads (Fig. 1B); and iii) the number of baits for some species, such as *C. meleagridis* and *C. viatorum* included in CryptoCap_100k is even higher than the number for *C. hominis*, which worked successfully (Table 1).

### CryptoCap_100k Applicability: Potential for the Detection of Mixed Infections

*Cryptosporidium* infections can contain multiple strains, subspecies, or species. Current methods for detecting mixed infections include amplicon, PCR-RFLP, and MLST approaches^35,57,58^. However, these methods have inherent biases due to preferential amplification and can miss mixed infections that are not comprised of *C. parvum* and *C. hominis*^35,36,57^. Regarding mixed-species infections, CryptoCap_100k was able to detect these in *in silico* simulations (Fig. S13); in *in vitro* pools of *C. parvum* and *C. meleagridis*, with a slight bias towards *C. parvum*; and to a lesser extent in clinical samples (i.e., EC4, UKH149, UKP381, and UKH151; Figs. 3H and S32). For instance, deconvolution analyses of sample UKH149 indicate a mixed-species origin, suggesting the potential for the genomic data obtained via CryptoCap_100k to support the deconvolution of mixed-infections within samples. Mixed infections with strains of the same species were detected in UKP347 and UKP391 (Fig. 4A) in double-enriched but not in single-enriched libraries. These discordances were associated with higher allelic depth (AD) and/or improved genotype quality (GQ) in double-enriched libraries (Figs. S77–S81), which likely allowed for greater resolution and ability to detect imbalances in within-host allele frequency by DEploid. However, these mixed-species and mixed-strain results should be interpreted cautiously and primarily as proof-of-concept, given the limited resolution of our dataset. Also, double-enrichments exhibit higher proportions of *C. parvum* compared to single-enrichments. These higher proportions of *C. parvum* can be caused by minor enrichment biases due to the high number of baits designed from *C. parvum*^47^.

### CryptoCap_100k: Conclusion

In summary, CryptoCap_100k results in a cost reduction of approximately 98% relative to deep sequencing of fecal samples. This translates into relatively inexpensive genomic data that can be applied to a large number of samples, including archived or fresh samples, from diverse origins such as stool, environmental sources, oocysts, or existing extracted DNA. However, cost savings will vary depending on the abundance of *Cryptosporidium* DNA in the sample (Supplementary Notes #14 and Table S21).

CryptoCap_100k can recover >80% of the genome at high depth from pure oocyst samples (Figs. S13–S14) and achieve as high as 3,690X fold-change enrichment with the iNextEra protocol in fecal samples, without the need for oocyst purification, allowing high-accuracy species assignment (Table S15). Also, it enables bait dilution and pooling of ∼10 libraries per capture reaction, which increases affordability. The quality of the retrieved genomic data allows for SNPs to be called, utility for population genomics, and provides the potential to assess mixed-infections. Double enrichment improves coverage for low-input DNA samples, but this outcome can be matched with single enrichments and higher sequencing effort. Studies using the CryptoCap_100k will enhance our understanding of the population genetics of human-infecting *Cryptosporidium* species and their relationship to epidemiology. Generating economical and efficient genome-level data enables the growth of reference resources and the refinement of analyses and methods for subtyping.

## Methods

### CryptoCap_100k Bait Design & Characterization

The genome sequences of six *Cryptosporidium* species were chosen for bait design (Table 1), along with 55 *18S rRNA* sequences and eight *gp60* sequences (Data Availability). All sequences were soft-masked for simple and low-complexity repeats using RepeatMasker 4.1.1 (http://www.repeatmasker.org/). We aimed to design 100,000 120-nucleotide baits. Baits that were ≤ 50% soft-masked for repeats were retained. These parameters resulted in 435,323 raw baits. Then, VSEARCH 2.15.0^59^ was used to cluster baits at various levels of sequence overlap and identity (Table 1), allowing for mismatches to reduce the number of baits while maintaining diversity. The genome order was designed to prioritize genome sequences based on their completeness and likelihood of human infection, with *C. parvum* IOWA-ATCC listed first for all clustering procedures. Then, *C. hominis*, *C. cuniculus*, and *C. tyzzeri* were collapsed at 50% overlap, and 90% identity relative to *C. parvum*; and *C. meleagridis* and *C. viatorum* were collapsed at 45% overlap and 85% identity relative to *C. parvum*. From *C. viatorum*, a subset of baits was randomly selected. Raw baits were obtained for the *18S rRNA* targets and collapsed at 50% overlap and 95% identity (Table 1).

We characterized CryptoCap_100k (Supplementary Notes #1) by using twelve genome sequences available for ten species of *Cryptosporidium* (Data Availability). BWA 0.7.17^60^ was used to map the CryptoCap_100k bait sequences to each genome sequence. Alignment files were generated and converted using Samtools 1.10^60,61^. The bamCoverage function from DeepTools 3.3.1^62^ was used to produce bigwig files to visualize coverage tracks calculated as the number of reads per bin in IGV-Web app version 1.10.8^63^. To obtain alignment statistics, the depth option from Samtools v1.1.0 and the *-a* flag were used to get all sites and create a depth file^61^. The mean depth, the proportion of baits mapped to each reference, and the breadth of genome coverage were estimated. The breadth of coverage, as reported throughout this manuscript, was determined by estimating the number of bases covered in the alignment over the reference length, with a minimum coverage depth of 1X.

We also compared the CryptoCap_100k bait set to CryptoCap_75k^47^ (Supplementary Notes #2) by estimating the same parameters as above, and using Welch’s two-sample t-test for comparisons.

### In Silico Simulations of Enrichment with CryptoCap_100k & Sequencing

To validate data quality from CryptoCap_100k before bait synthesis, a simple infection scenario, error-free and bias-free, was simulated. We used twelve available genome sequences from ten *Cryptosporidium* species to simulate the enrichment and sequencing process with CryptoCap_100k, assuming two different ranges of insert size fragments (Supplementary Notes #3). To simulate hybridization, we used the coordinates of the mapped baits in relation to each reference genome, and we extended the position by several bases to “capture” larger fragments than the bait length using Samtools^61^. To simulate sequencing, ART 2016.06.05^64^ was used to reproduce ∼200,000 paired-end 150 bp fastq reads from the extended coordinates in each reference genome. Then, reads were mapped to their respective genome sequences using BWA 0.7.17^60^, and we estimated depth and breadth of coverage. To validate the accuracy of species assignment, we used a reference database containing ten *Cryptosporidium* species genome sequences (10-CrypGS), a *gp60* database^2^, and an *18S rRNA* database^55,65,66^, and used BBMap 38.90^67^ to map simulated fastq reads to each and explore the proportion of hits to each *Cryptosporidium* species.

To assess the sensitivity of the baits to off-target DNA, we assessed the affinity of our baits to 1,008 genome sequences^68^, including the human genome. Because *in vitro* libraries were diluted in calf thymus DNA (see *In Vitro Test of Limits of Detection with Different Levels of C. parvum Dilution*), we also assessed affinity to the *Bos taurus* genome sequence (GCA_002263795.3). For both analyses, we used BWA 0.7.17^60^ to map the bait sequences, and we estimated the number of hits as well as the quality of the mapping.

To explore the ability to discriminate different species within mixed-species infection samples, we created four simulated samples by concatenating the fastq files already simulated for each species above. The four *Cryptosporidium* mixed infection samples contained at equal proportions: i) reads from twelve genome sequences from ten species; ii) reads from ten genome sequences from ten species; iii) reads from five genome sequences and species that share high percent identity; and iv) reads from three genome sequences and species commonly found in human infections. Then, we mapped these mixed reads to 10-CrypGS, the *gp60*, and the *18S rRNA* databases. See Supplementary Notes #3–7 for details.

### In Vitro Enrichment & Validation with CryptoCap_100k Baits with Pure Oocysts’ DNA

Pure oocyst DNA was obtained from three *C. parvum* samples from Dr. Michael Grigg (National Institutes of Health, US) and *C. meleagridis* DNA isolate TU1867 from BEI Resources (BEI Resources NR2521 Manassa, VA). Metagenomic libraries were prepared with the New England BioLabs Ultra FS II library kit (see *Metagenomic Libraries*), and aliquots of these were left unenriched or enriched with one-tenth of the recommended volume of CryptoCap_100k (see *Hybridization Capture Enrichments*). Libraries were sequenced using an Illumina MiSeq with a PE250 kit.

### In Vitro Simulation of Mixed Infections

To test the ability to detect mixed infection samples, after NEB/iTru library preparation (see *In Vitro Enrichment & Validation with CryptoCap_100k Baits with Pure Oocysts’ DNA*) but before bait capture, two indexed libraries were pooled at equimolar ratios to simulate a mixed infection. After pooling, an aliquot was enriched with one-tenth of the recommended volume of CryptoCap_100k, and another remained unenriched. These were sequenced in an Illumina MiSeq using a PE250 kit. We estimated the number of reads per sample after demultiplexing, expecting roughly 50% for each species.

### In Vitro Test of Limits of Detection with Different Levels of C. parvum Dilution

To determine the CryptoCap_100k limits of detection (LOD) of *Cryptosporidium*, we performed two types of library preparations, NEB-iTru and iNextEra^69^, on samples with known and varied input DNA. For NEB-iTru libraries, we diluted *C. parvum* genomic DNA in a background of calf thymus DNA (Sigma-Aldrich, St. Louis, MO). A 45-ng aliquot of calf thymus DNA was pipetted into five tubes. In the first tube, 5 ng of *C. parvum* genomic DNA was added to a final volume of 50 µL. Then, three 1:10 serial dilutions were performed. The fifth tube corresponded to a negative control with just the calf DNA. Metagenomic libraries using the NEB-iTru protocol and 10 µL of each dilution were generated (see *Metagenomic Libraries*). In this experiment, up to 7 libraries were pooled, an aliquot was left unenriched, and independent aliquots were enriched with full, one-half, a quarter, an eighth, or a sixteenth of the recommended volume of CryptoCap_100k (see *Hybridization Capture Enrichments*). Pools were sequenced at Novogene using a HiSeq X with a PE150 kit. The number of reads, percentage of reads mapped to the *C. parvum* genome sequence, and depth and breadth of genome coverage were estimated.

For iNextEra libraries, 90 ng aliquots of calf DNA were pipetted into six tubes. In the first tube, 10 ng of *C. parvum* genomic DNA was added in a final volume of 100 µL. From this, four 1:10 serial dilutions were performed. The sixth tube corresponded to the negative control (calf DNA only). iNextEra library preparation was performed with 10 µL inputs from these dilutions (see *Metagenomic Libraries*). Up to 6 libraries were pooled, and aliquots were left unenriched or enriched with one-quarter or one-eighth of the recommended baits. Aliquots of single-enriched libraries were double-enriched with one-quarter of the recommended baits (see *Hybridization Capture Enrichments*). Pools of unenriched, single-enriched, and double-enriched libraries were sequenced at Novogene using a NovaSeq with a PE150 kit. The number of reads, the percentage of reads mapped to the *C. parvum* genome sequence, the depth and breadth of genome coverage, and species assignment based on the mapping of reads against 10-CrypGS were analyzed among dilutions.

### In Vitro Test of Limits of Detection with Different Levels of Bait Dilution

To determine the sensitivity or limits of detection (LOD) when using a lower concentration of baits than recommended, we modulated the quantity of baits added to the capture reaction in one experiment with NEB-iTru libraries and one experiment with iNextEra libraries.

For both library types, the bait dilution sensitivity assay considered pools of the metagenomic libraries from the *In Vitro Test of Limits of Detection with Different Levels of C. parvum Dilution* experiment. For NEB-iTru libraries, we tested five different bait quantities: full volume (5.5 µL, per the manufacturer’s specifications), and dilutions in water at one-half, a quarter, an eighth, and a sixteenth. For iNextEra libraries, we tested a quarter and an eighth bait dilutions. Sequencing was performed as described in the section above. The number of reads, the percentage of reads mapped to the *C. parvum* genome sequence, and the depth and breadth of genome coverage were compared among bait dilutions.

### In Vitro Enrichment & Validation with CryptoCap_100k Baits with Clinical Samples

Two clinical datasets were used for real-world validation. First, Dr. Michael Grigg (National Institutes of Health, US) and Prof. Rachel Chalmers’s laboratory (Public Health Wales—PHW— Microbiology and Health Protection, UK) provided DNA extracted (Qiagen PowerSoil Kit) from seven de-identified patient fecal samples. These samples were obtained under an NIAID IRB-approved protocol NCT00006150. The NEB-iTru library protocol was used, pools made, aliquots were either left unenriched or enriched with one-tenth of the recommended volume for baits, and libraries were sequenced in an Illumina HiSeq using a PE150 kit.

Second, 100 DNA extractions (Qiagen QIAamp Fast DNA Stool kit) from clinical samples from Prof. Rachel Chalmers’s laboratory (PHW) were used. This work was carried out in accordance with a material transfer agreement between UGA and PHW, which did not require ethical approval. These samples had previously been screened using a multi-locus typing system ^49^ to detect the species present, and these previous results are reflected in the sample names used here (*C. parvum* = UKP and *C. hominis* = UKH). The laboratory workflow used for these clinical samples in this study is depicted in Fig. 5. A real-time qPCR assay was used to quantify *Cryptosporidium*, using primers for the *18S rRNA* gene and slightly modified probes ^70^ as specified in Supplementary Notes #12.2. Two types of genomic libraries were prepared for each sample (NEB-iTru and iNextEra; see *Metagenomic Libraries*). Nine DNA samples were depleted when used for iTru libraries, thus not used for iNextEra libraries. Aliquots of pools of ∼12 libraries were left unenriched or enriched. iTru-NEB libraries were enriched with one-tenth and iNextEra libraries were enriched with one-eighth of the recommended volume of CryptoCap_100k. For iNextEra libraries, we performed additional double enrichments on single-enriched aliquots using one-quarter of the recommended volume of baits (see *Hybridization Capture Enrichments with CryptoCap_100k*). All libraries were sequenced using an Illumina NovaSeq X with a PE150 kit (Fig. 5).

**Figure 5.**
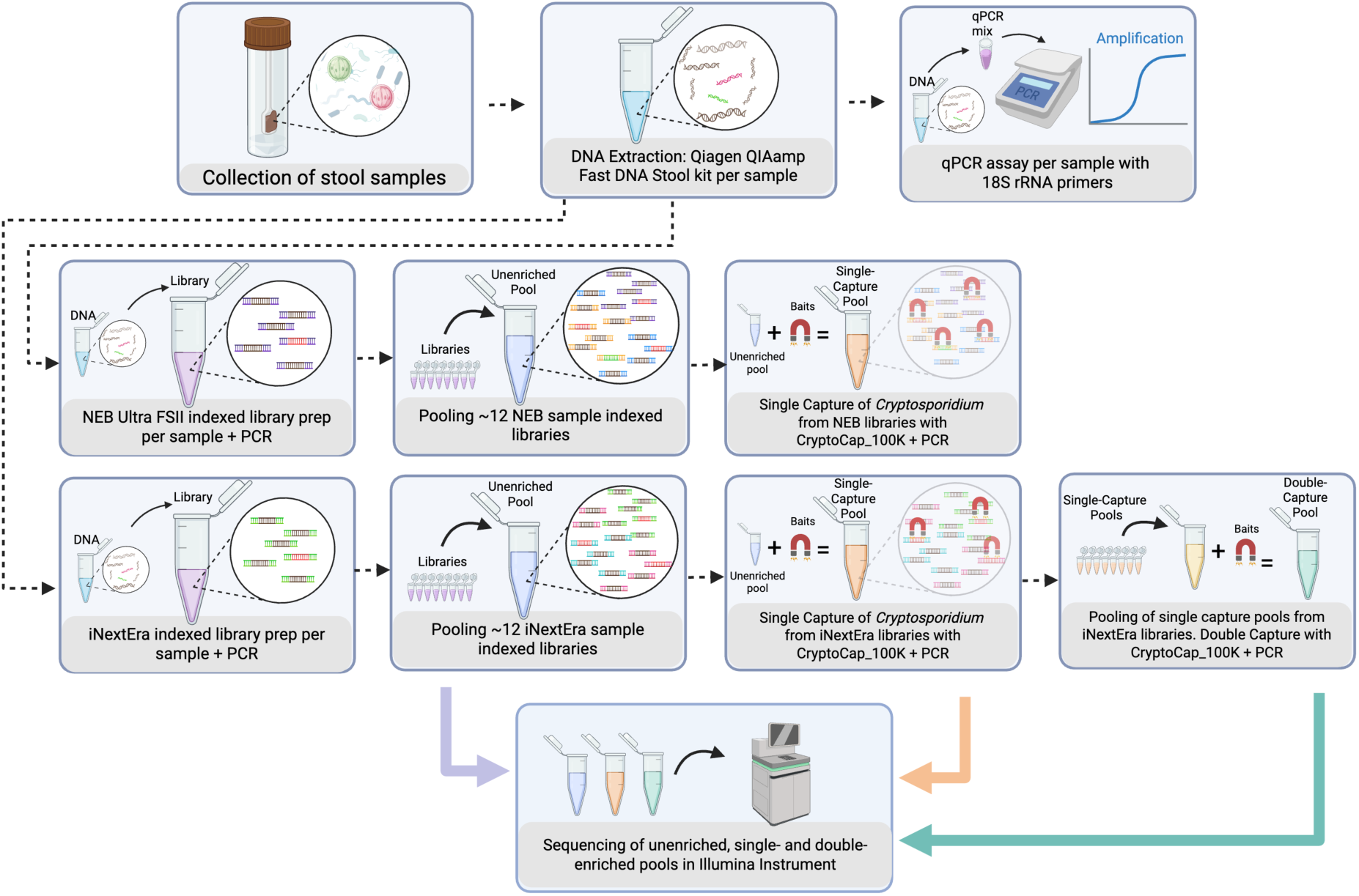
**Laboratory workflow used for the analysis of a set of clinical samples from the UK**. The genomic DNA of each patient’s sample was extracted, and *Cryptosporidium* was validated via qPCR. Each DNA extract was used for two types of library prep, NEB Ultra FSII and iNextEra library prep. Unenriched library preparation aliquots were used for sequencing. Also, aliquots from unenriched libraries were pooled according to library type, and enrichments were performed with CryptoCap_100k on these pools. Aliquots of these products, referred to as single-enriched products, were used for sequencing. Aliquots of single-enriched libraries, which used the iNextEra protocol, were pooled, and enrichments were performed again with CryptoCap_100k. Aliquots of these products, named double-enriched products, were used for sequencing. Colors represent those used throughout the manuscript to differentiate unenriched, single-enriched, and double-enriched products. Created in BioRender. Bayona N (2025) https://BioRender.com/yysbjqh.

### Metagenomic Libraries

For iTru-NEB libraries, the New England BioLabs (NEB) Ultra FS II library kit (New England BioLabs, Ipswich, MA) protocol was used according to the manufacturer, with enzymatic fragmentation lasting 5–11 minutes, based on the integrity of DNA in gel electrophoresis. A 12- or 14-cycle PCR with iTru primers^69^ was used to add unique dual-indexes to each ligation product. Products were cleaned with 1.25X SpeedBeads^71^ and quantified with a Qubit 4.0 Fluorometer (Thermofisher, Waltham, MA).

iNextEra library preparation was used to decrease the time, cost, and amount of input DNA needed. In brief, 2.5 µL of 10 ng/µL (25 ng) of sample DNA was used for a tagmentation reaction with 3 µL of 2X TMP buffer and 0.5 µL of Transposome (BLT; CAT # 20015880, Illumina Inc.), incubated at 53 ℃ for 30 minutes. Then, PCR reactions contained 9.5 µL of Q5 Buffer, 1.25 µL of 10 mM dNTPs, 0.5 µL of Q5 High Fidelity Polymerase (New England Biolabs M0491L), 40.25 µL of nuclease-free water, 2.5 µL of iNextEra5 indexed primer at 5 µM, 2.5 µL of iNextEra7 indexed primer at 5 µM^69^, and 6 µL of tagmented product. Cycling conditions were: 3 minutes at 72 ℃, 3 minutes at 98 ℃, followed by ten cycles of 98 ℃ for 45 seconds, 62 ℃ for 30 seconds, and 72 ℃ for 2 minutes, with a final extension at 72 ℃ for 1 minute.

### Hybridization Capture Enrichments with CryptoCap_100k

Metagenomic libraries were pooled (e.g., Fig. 5) in preparation for enrichment. For different experiments, the input concentration per pool of samples varied (i.e., for clinical samples using NEB/ iTru, the input was between 25–80 ng/µL; for clinical samples using iNextEra, it was 103 ng/µL; and for the LOD experiments, it was 26–59 ng/µL). The enrichment process per pool followed manufacturers’ standard protocols (mybaits v5, Daicel Arbor Biosciences, Ann Arbor, MI), with wash and hybridization temperatures at 63 ℃ and modifications of bait quantities as described above. After enrichments, P5/P7 PCRs with 16–22 cycles were performed, followed by 1–1.25X SpeedBeads clean-ups (Sigma Aldrich, St. Louis, MO).

For the 100-clinical sample set and the LOD experiments, double enrichments were performed on the iNextEra single-captured pools (e.g., Fig. 5). For these, aliquots of the cleaned single-captured pools were pooled and used independently as input for a second capture using one-quarter of the recommended baits. For post-double-enrichments P5/P7 PCRs with 12–16 cycles were performed, followed by 1X SpeedBeads clean-up.

All PCRs were performed using the KAPA HiFi Kit (Roche, Basel, Switzerland). All P5/P7 PCRs consisted of 98°C for 2 minutes.; then, cycles of: 98°C for 20 sec., 60°C for 30 sec., 72°C for 1 min.; followed by 72°C for 5 min.

### In Vitro Bioinformatic Processing and Analyses

Bioinformatic and data analysis code are available in the Supplementary Notes. Sequencing data were processed using Trimmomatic 0.39^72^, and retained reads were mapped to specific genome sequences or databases using BBMap 38.9^67^. Samtools^61^ was used to sort bam files. We determined mapping statistics, including genome depth and breadth of coverage. The results of the mapping to 10-CrypGS were used to assess the proportion of hits to the different *Cryptosporidium* species and produce species assignment plots. For iNextEra libraries from clinical samples, we also generated bigwig files, which were visualized using the IGV-Web app version 2.2.7.

Cleaned reads from pure oocysts libraries and *in vitro* mixed-infection libraries were mapped to the corresponding target references (i.e., *C. parvum* samples to *C. parvum* IOWA-ATCC, *C. meleagridis* samples to chromosome-level assembly^73^, also to 10-CrypGS, to the *gp60* database, and the *18S rRNA* database—used in simulations). The sequencing reads from the LOD dilution experiments (i.e., input DNA and bait dilutions) were mapped to *C. parvum* IOWA-ATCC and 10-CrypGS. iNextEra reads from LOD were also mapped to *Bos taurus* (GCA_002263795.3). The cleaned sequence reads from clinical samples were mapped to *C. parvum* IOWA-ATCC and 10-CrypGS. For the set of 100 clinical samples, qPCR C_T_ values were used in the *parsnip 1.1.1*^74^ to define linear regression models predicting the percentage of reads mapped against 10-CrypGS (Supplementary Notes #12).

### Population Genomics and Mixed Infection Analyses of Clinical Samples

To examine the genetic variation in natural populations of *Cryptosporidium* from the set of 91 clinical samples prepared using the iNextEra protocol, we performed genotype calling (Fig. S89). In short, reads from each treatment (unenriched, single-enriched, and double-enriched) were mapped to the *C. parvum* IOWA-ATCC genome sequence using BWA 0.7.17^60^, and the results were used in GATK 4.6.0.0^75^. Duplicate reads and haplotypes were determined using Haplotypecaller with GVCF mode and merged into databases for each library type to perform joint genotyping using GenotypeGVCFs.

Importantly, HaplotypeCaller was run with --sample-ploidy set to 1. A first filtering approach was performed to obtain SNPs with QD < 2, FS > 60, MQ <40, MQRankSum < -12.5, and ReadPosRankSum < −8.0.

The resulting VCFs were explored for missing data and coverage depth per sample and chromosome using vcfR 1.15.5^76^. Then, filtering using dartR 2.9.7^77^ considered only SNPs with coverage (DP) larger than 5 and lower than 200, that were biallelic in the dataset, and with less than 40% of missing data. Individuals with > 80% missing data were removed.

To explore the diversity and divergence among samples, filtered VCF files for each treatment were used for population genomic analyses. These included Principal Components Analyses performed with SNPRelate 1.40.0^78^ and ade4 1.7-23^79^. We used phangorn 2.12.1^80^and SplitsTree 6.4.11^81^ to compute pairwise genetic distances and visualize phylogenetic networks. We estimated Nei’s genetic distances between samples and visualized them in dendrograms with bootstrap support using poppr 2.9.6^82^ and plotted these with ape 5.8-1^83^. We tested K = 1–5 clusters, each with three replicates, in Structure 2.3.4^84^, run through structure_threader 1.3.10^85^, which utilizes Structure Harvester 0.6.94^86^. We used 10,000 chains for burn-in, 100,000 chains for MCMC, and set ploidy to 1, with other options left as default. We used readr 2.1.5^87^ and starmie 0.1.2^88^ to graph Evanno’s results and to plot Structure assignment plots. We estimated pairwise F_ST_ statistics between UKP and UKH samples using hierfstat 0.5-11^89^ and mapped these values along *C. parvum* chromosomes using qqman 0.1.9^90^. We compared Allelic depth (AD) and genotype quality (GQ) values between VCF files of the different treatments and used ggridges 0.5.6^91^ to plot the frequency of values across samples.^90^

To explore the diversity within each clinical sample, we deconvoluted multiple infecting strains using DEploid 0.7.1^92^with no reference panel. The effective and inferred number of strains (K) was plotted. Additionally, using filtered VCFs from the GATK 4.6.0.0^75^ run with the setting --sample-ploidy set to 2, we estimated nucleotide diversity (π) of SNPs in VCFTools 0.1.16^93^ for double and single enrichments that were subset to contain only UKH (*C. hominis*) or UKP (*C. parvum*) samples; and, we assessed and plotted within-sample allele variation (what in a diploid system would be categorized as heterozygosity) levels per sample. Data analyses were performed in R Studio 2023.06.0. Least-square comparisons of the means between experimental groups (e.g., unenriched and enriched, single-capture vs. double-capture, etc.) were performed to calculate p-values using a confidence interval of 0.95 with the package: *lsmeans*^94^, which provides a t-test output when comparing two variables, or performs a Tukey method adjustment when comparing a family of three or more estimates. Other packages used were *tidyverse 2.0.0*^95^, *ggplot2 3.5.2*^96^, and *plotly 4.10.4*^97^.

## Acknowledgments

T.G. and J.C.K. disclose support from the National Institutes of Health under award number NIH R01AI148667. M.G. discloses support from the Intramural Research Program of the National Institute of Allergy and Infectious Diseases (NIAID) at the National Institutes of Health. We thank the authors of CryptoCap_75k^47^ for sharing unpublished results that facilitated the work and comparisons presented here. We also thank Prisha Sharma and Fiifi Agyabeng-Dadzie for sample processing, Christopher Fitzgerald for sample tracking and processing, and Mustafa Nural for sample database help. This study was supported in part by resources and technical expertise from the Georgia Advanced Computing Resource Center, a partnership between the University of Georgia’s Office of the Vice President for Research and Office of the Vice President for Information Technology.

## Author Contributions

N.J.B-V., A.H.S., M.S.B., A.K., R.P.B., M.E.G., J.C.K., and T.C.G. contributed to conceptualization, methodology, investigation, and writing—review and editing. M.S.B., R.P.B., and B.B. designed and oversaw the production and QC of the baits. M.S.B., K.N.P., M.I.U.B., contributed to investigation and writing—review and editing. N.J.B-V. and A.H.S. performed validation, data curation, formal analyses, and with T.C.G. writing—original draft. G.R., R.M.C., E. A-F, M.E.G., and J.C.K. contributed samples and analyses to the manuscript. N.J.B-V, M.S.B, A.K., R.P.B., M.E.G., J.C.K., and T.C.G contributed to funding acquisition. All authors read and approved the final version of the manuscript.

## Competing Interests

B.B. and M.B. are employed by Diacel Arbor Biosciences, which manufactures and sells the baits described herein. The CryptoCap_100k bait set is available for commercial purchase (Design ID: D10006Crypto, Cat# #308508.V5, #308548.V5, and #308596.V5, Arbor Biosciences, MI, USA).

## Data Availability

The six genome sequences that supported the CryptoCap_100k bait design were from GenBank: GCA_015245375.1, GCA_004337795.1, GCA_001483515.1, GCA_007210665.1, GCA_004337835.1, and GCA_001593445.1. The extra sub-telomeric sequences from *C. parvum* were: MZ892386.1, MZ892388.1, and MZ892387.1.

The *18S rRNA* sequence data that supported the CryptoCap_100k bait design were from GenBank with the EU250845, LC089976, DQ288166, EF641022, GU951714, AF108864, LC012016, GQ121020, HQ397716, KR296812, EF641014, FJ435960, JX416368, KP099082, AY120903, AY120902, JX644908, AY324641, AY324638, AF159113.1, JQ413438, AF108861, GQ121021, KP730304, AY737602, AY731235, JX294358, AY268584, KR819168, AY120904.1, KU608308, AF112576, EU553551, JF710247, KP004205, JQ002555, KT235702, AB909498.1, AF316630, AY504515, AY504516, AF262325, GQ227479, EU827297, AF093500, AF093497, AY954886, HM116385, DQ650344, KY490554, KC305650, EF547155, FJ769050, HM243547, and AY524773 accession codes.

The *gp60* sequence data that supported the CryptoCap_100k bait design were from GenBank with the FJ490060, FJ490087, FJ490058, JX412915, JX412926, JX412925, KC204983, and KC204984 accession codes.

The twelve *Cryptosporidium* genome sequences used for CryptoCap_100k bait set characterization and *in silico* simulations were from GenBank with the GCF_00000165345.1, GCA_004337835.1, GCA_004337795.1, GCA_007210665.1, GCF_001865345.1, GCA_001865355.1, GCA_001593455.1, GCF_000006515.1, GCA_000165345.1, GCA_001483515.1, and GCA_002223825.1 accession codes *C. hominis* UdeA01 and *C. parvum* IOWA-ATCC were removed for those analyses with only ten genome sequences reference database (10-CrypGS).

Generated fastq files from all *in silico* and *in vitro* experiments for mock and clinical samples are available in GenBank BioProject number PRJNA1061798. Clinical samples were scrubbed of human DNA upon submission by NCBI using their Human Read Removal Tool.

## Computer Code

The code and data used for all sections of this paper are described in the Supplementary Notes, also available in Dryad, doi: 10.5061/dryad.gtht76j0h.

## Notes

### Summary of Updates

This version streamlines the work to focus on the method that works best, increases figure quality, clarifies interpretations, reduces overstatements, and aligns results and conclusions.

