## Supplementary Figures for "Genome-targeted enrichment and sequencing of human-infecting *Cryptosporidium* spp."

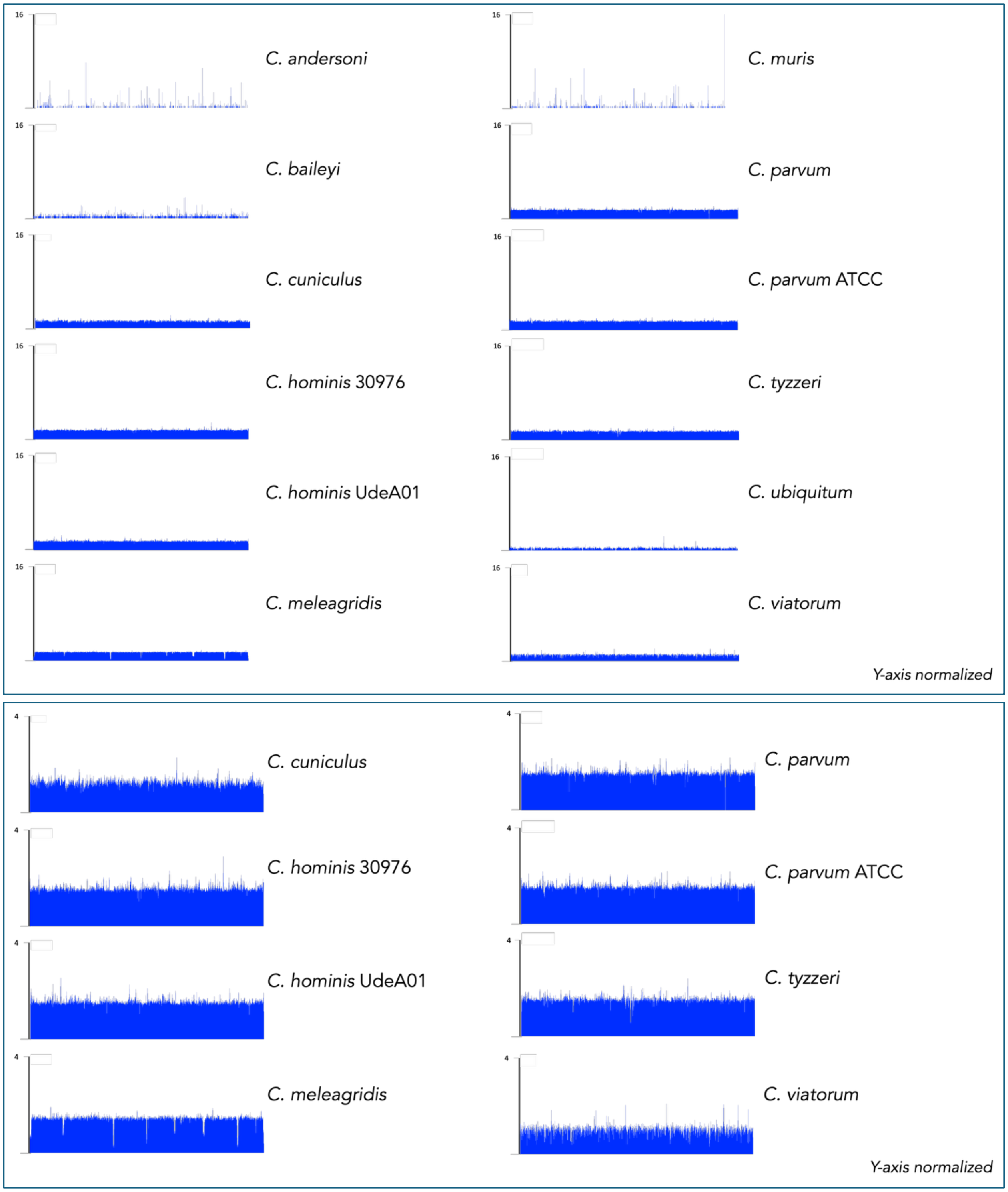


**Supplementary Figure S1.** Mean breadth and depth of coverage per 50-bases bin from the mapping of sequences from the CryptoCap_100k bait set to each *Cryptosporidium* reference genome. Top, all species analyzed. Bottom, species considered in bait design.

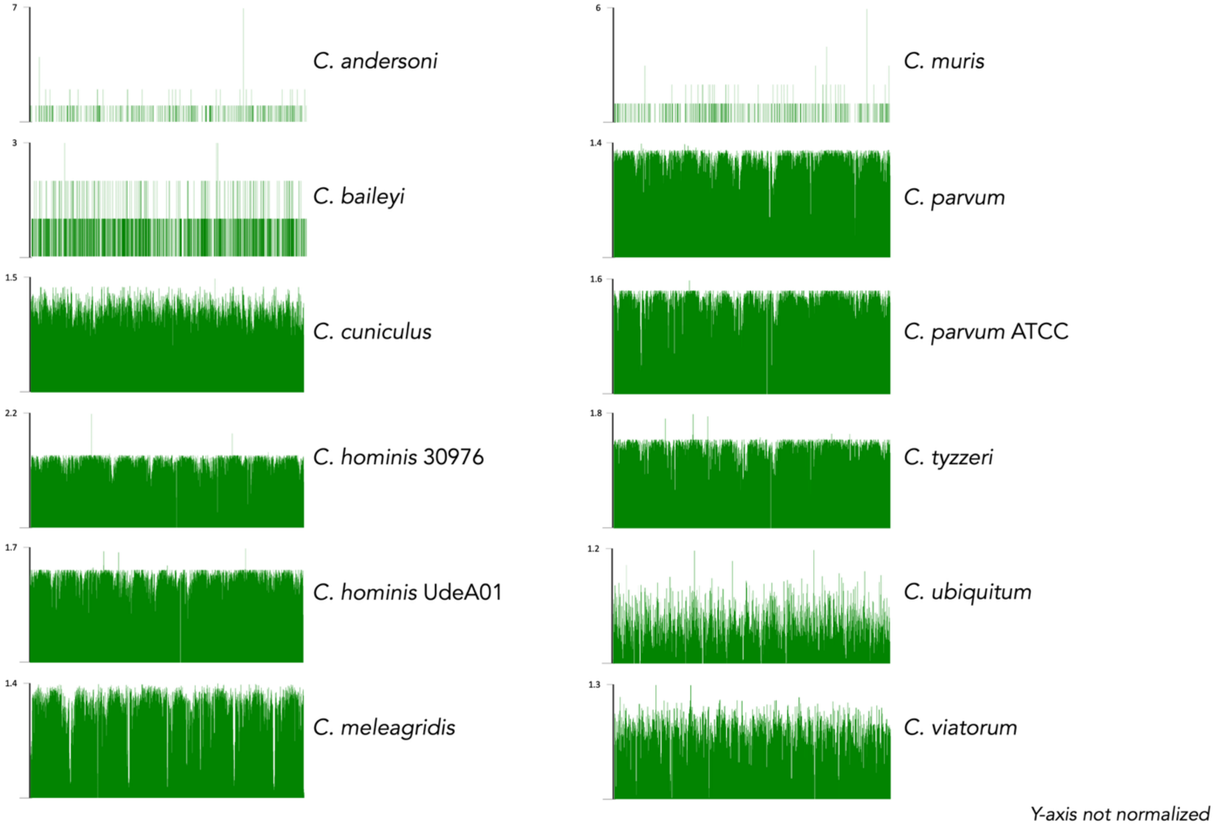


**Suppl. Fig. S2**. Mean coverage per 50-bases bin from the mapping of sequences from the CryptoCap_75k bait set to each *Cryptosporidium* genome.

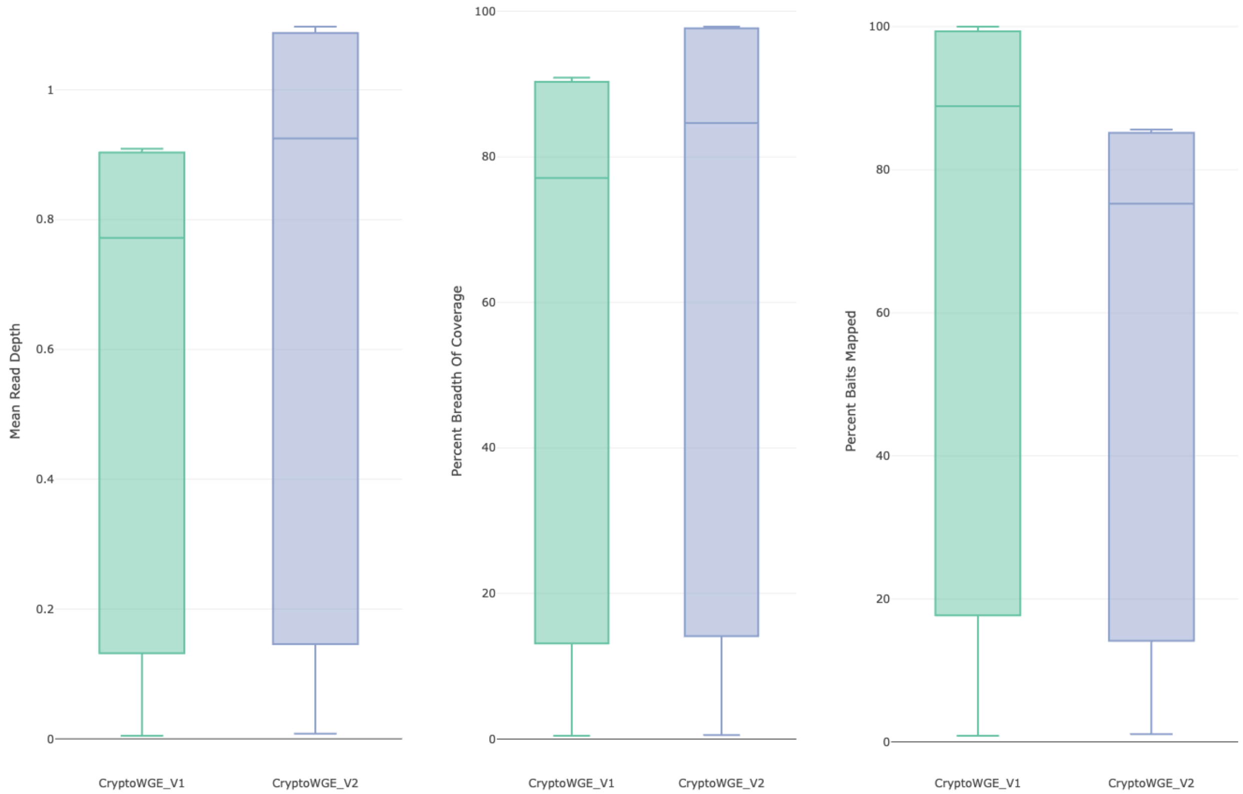


**Suppl. Fig. S3** Boxplot of mean read depth covered by baits, breadth of coverage along a genome of the baits, and percent of baits designed that mapped to each *Cryptosporidium* reference genome. CryptoCap_75k (CryptoWGE_V1, green) and CryptoCap_100k (CryptoWGE_V2, blue).

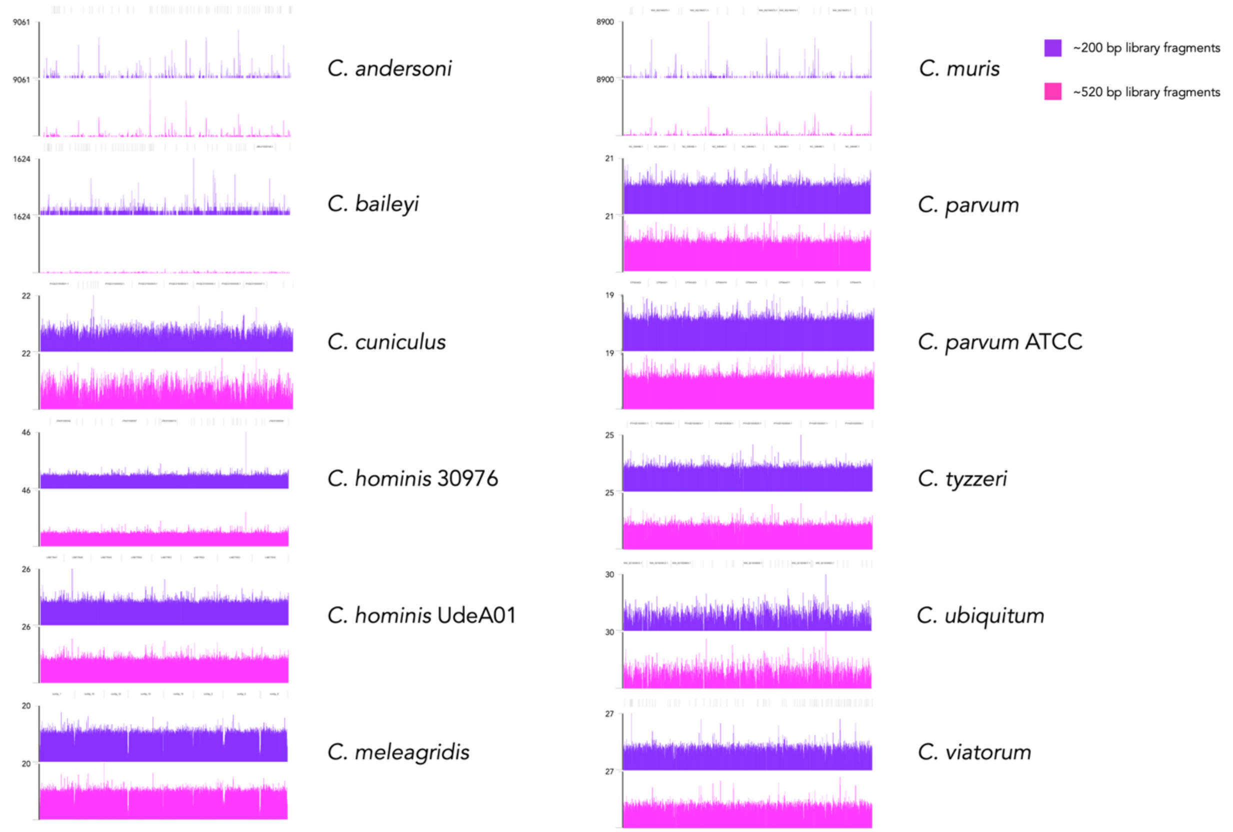


**Suppl. Fig. S4.** Horizontal and vertical coverage of simulated enriched reads from two fragment sizes (~200 bp, purple and ~520 bp, pink) when mapped against twelve *Cryptosporidium* genomes.

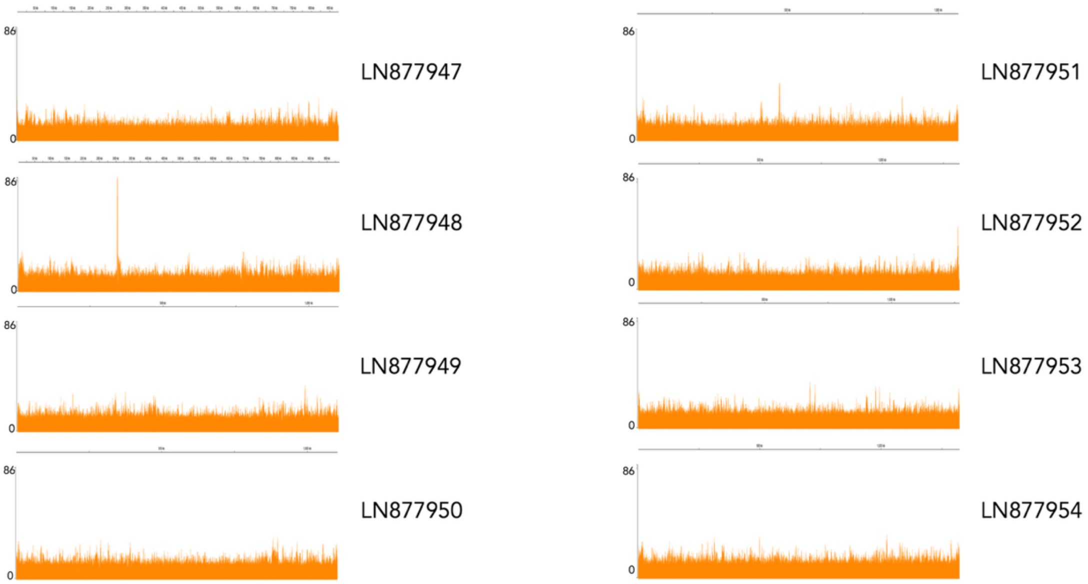


**Suppl. Fig. S5.** Chromosomal horizontal and vertical coverage obtained from the mapping to the *Cryptosporidium hominis* UdeA01 reference genome of simulated reads from fragments of ~520 bp size and enriched with CryptoCap_100k.

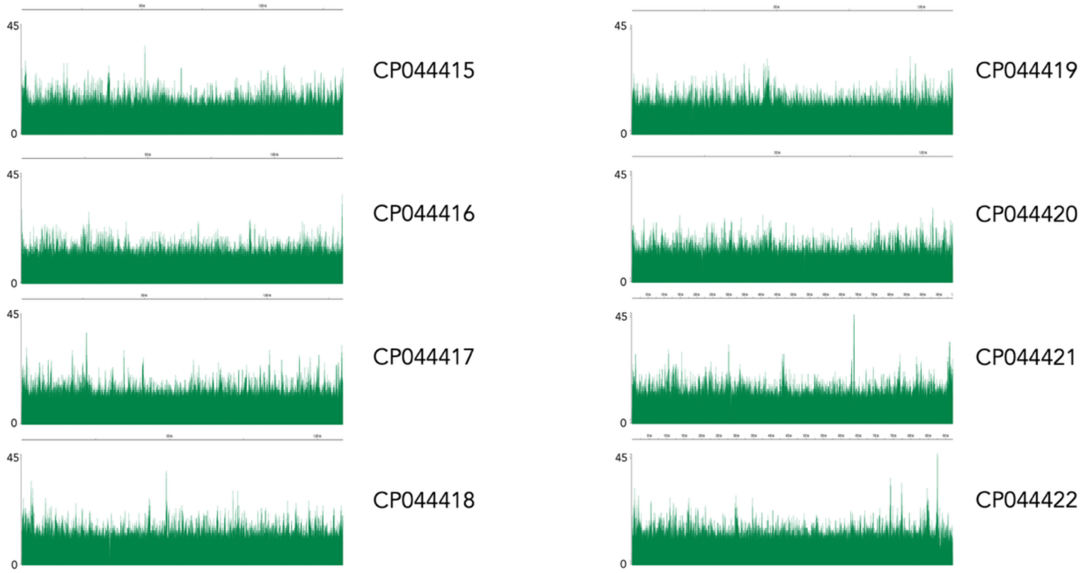


**Suppl. Fig. S6.** Chromosomal horizontal and vertical coverage obtained from the mapping to the *Cryptosporidium parvum* IOWA-ATCC reference genome of simulated reads from fragments of ~520 bp size and enriched with CryptoCap_100k.

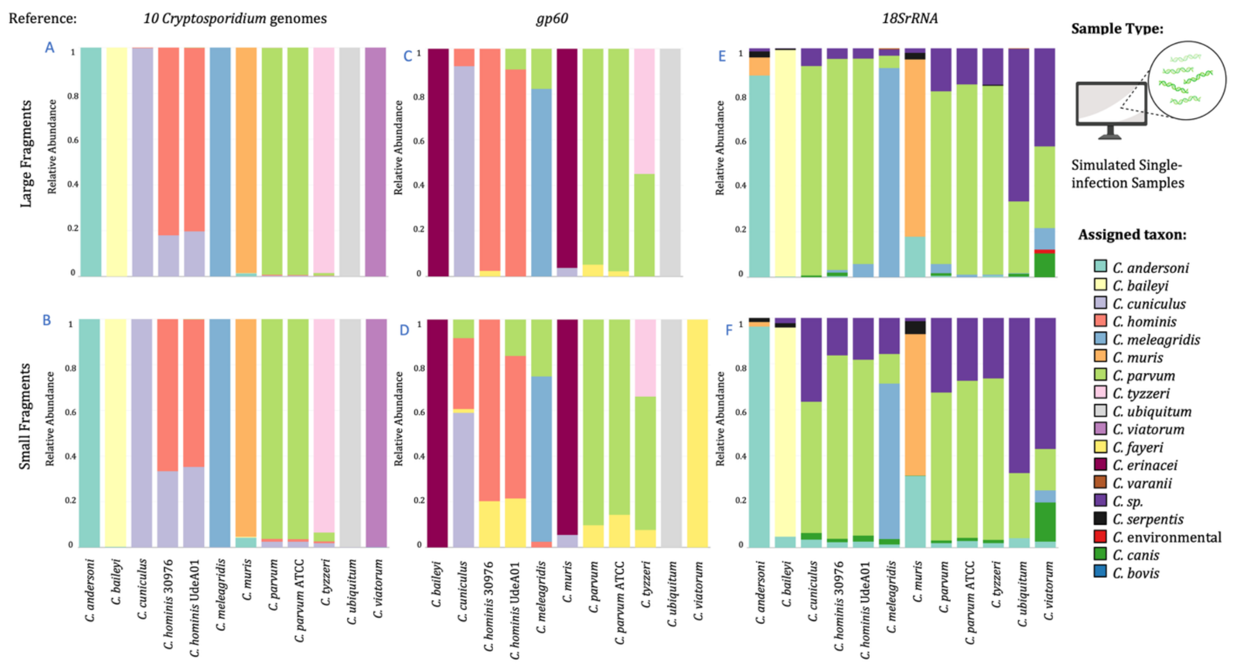


**Suppl. Fig. S7.** Bar plots of relative abundance of species assignment from mapping *in silico* simulated enriched and sequenced reads for each of the twelve *Cryptosporidium* genomes to three reference databases. Simulations were modeled separately for two fragment-insert size classes, one of large-520 bp fragments (above) and one of small-200 bp fragments (below). Three reference databases were used to map simulated reads, one containing ten *Cryptosporidium* genome sequences (10-CrypGS), a *gp60* database (Khan et al., 2018), and an *18S rRNA* database (Xiao et al., 2010).

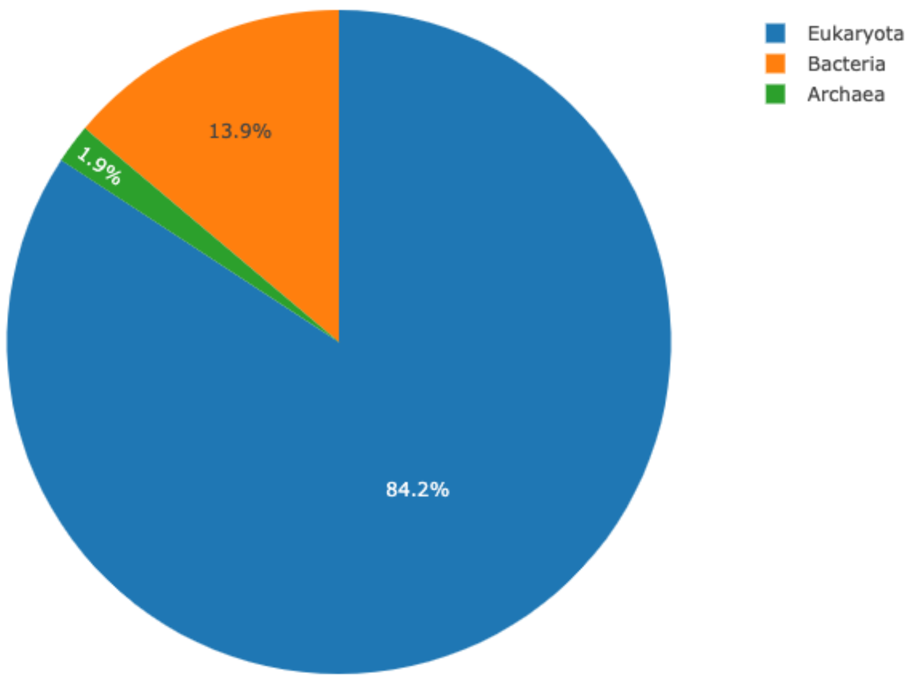


**Suppl. Fig. S8**. Pie chart of the proportion of the domains obtained from the 894 hits recovered from the mapping of the sequence of the baits from the CryptoCap_100k to the Lindgreen database.

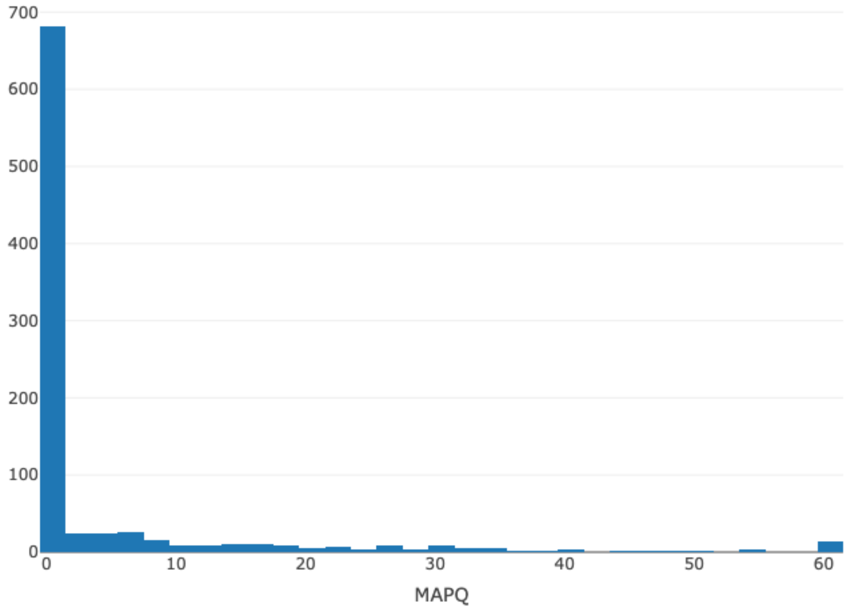


**Suppl. Fig. S9.** Histogram of MAPQ scores for the 894 hits from the mapping of the sequence of the baits from the CryptoCap_100k to the Lindgreen reference genomes database.

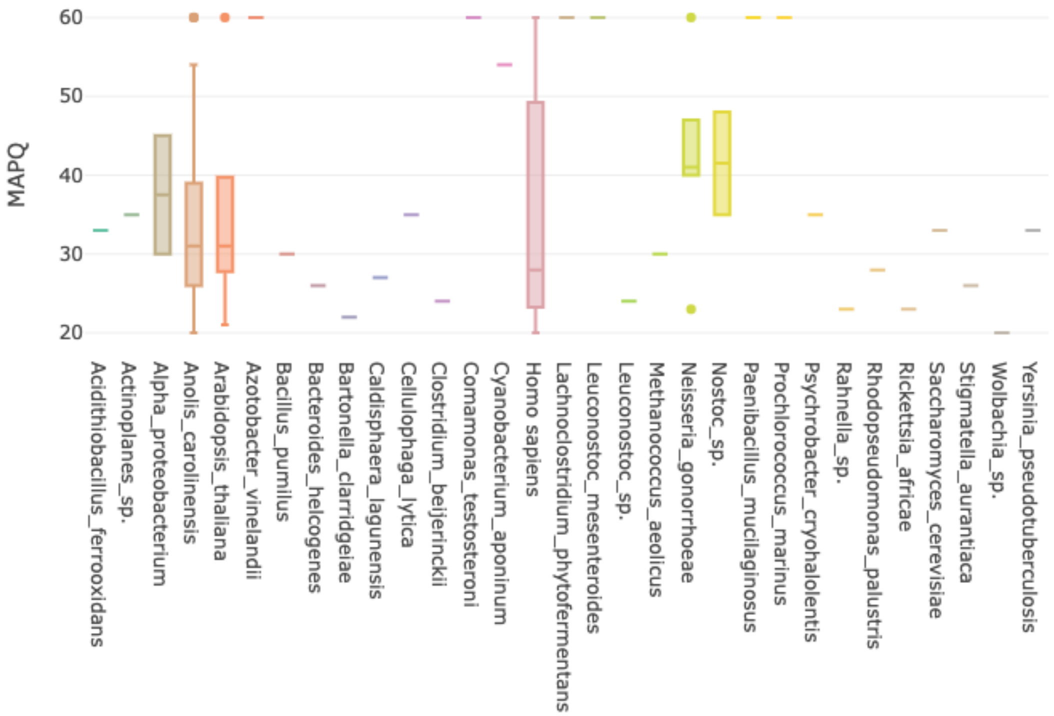


**Suppl. Fig. S10**. Boxplot of species with hits with high MAPQ scores (>20) from the mapping of the sequence of the baits from the CryptoCap_100k to the Lindgreen reference genomes database.

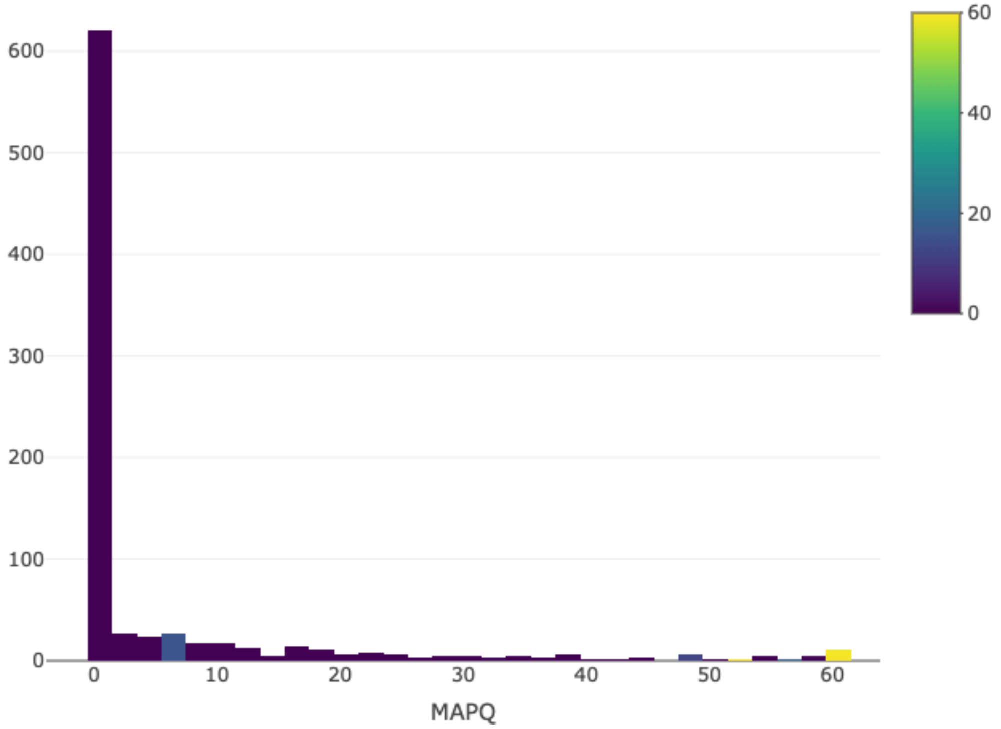


**Suppl. Fig. S11**. Histogram of MAPQ scores from the mapping of the sequence of the baits from the CryptoCap_100k to the *Bos taurus* reference genome (GCA_002263795.3).

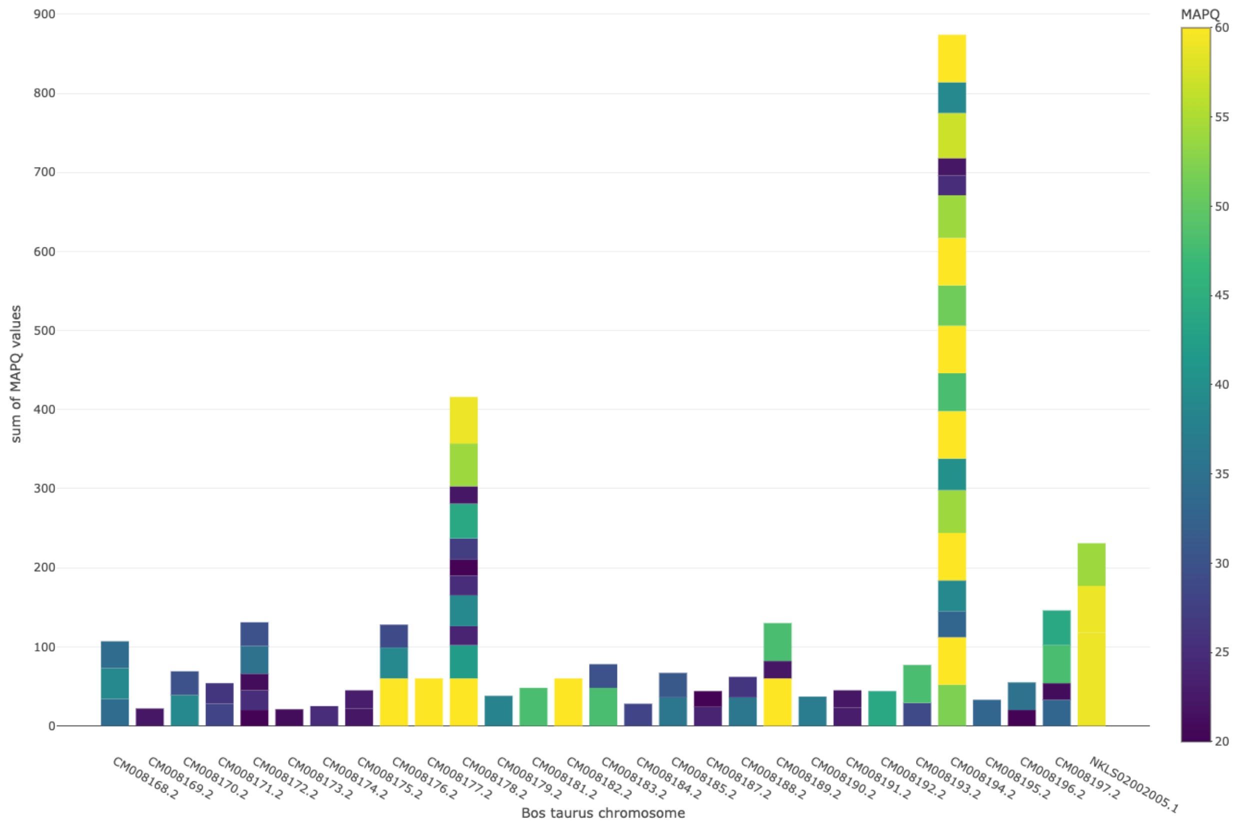


**Suppl. Fig. S12.** Stacked barplot of MAPQ scores (>= 20) from the alignment of CryptoCap_100k baits to each of the *Bos taurus* chromosomes.

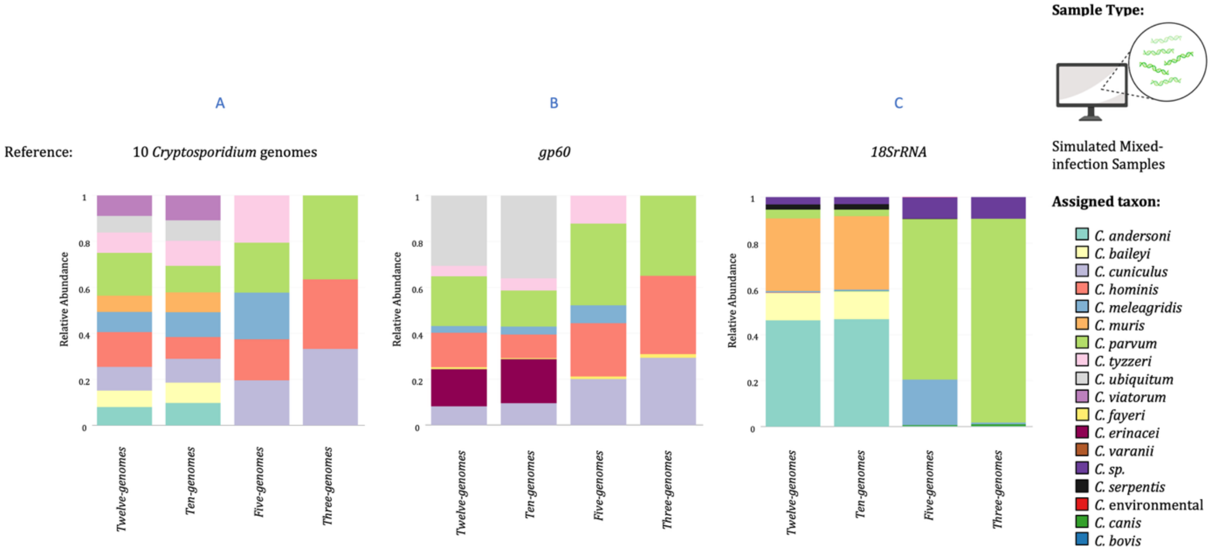


**Suppl. Fig. S13.** Bar plots of the relative abundance of species assignment from mapping four classes of *in silico* simulated enriched and sequenced mixed-infection samples. Three reference databases were used to map mixed infection simulated reads, one containing A) ten *Cryptosporidium* genome sequences (10-CrypGS), B) a *gp60* database (Khan et al., 2018), and C) an *18S rRNA* database (Xiao et al., 2010).

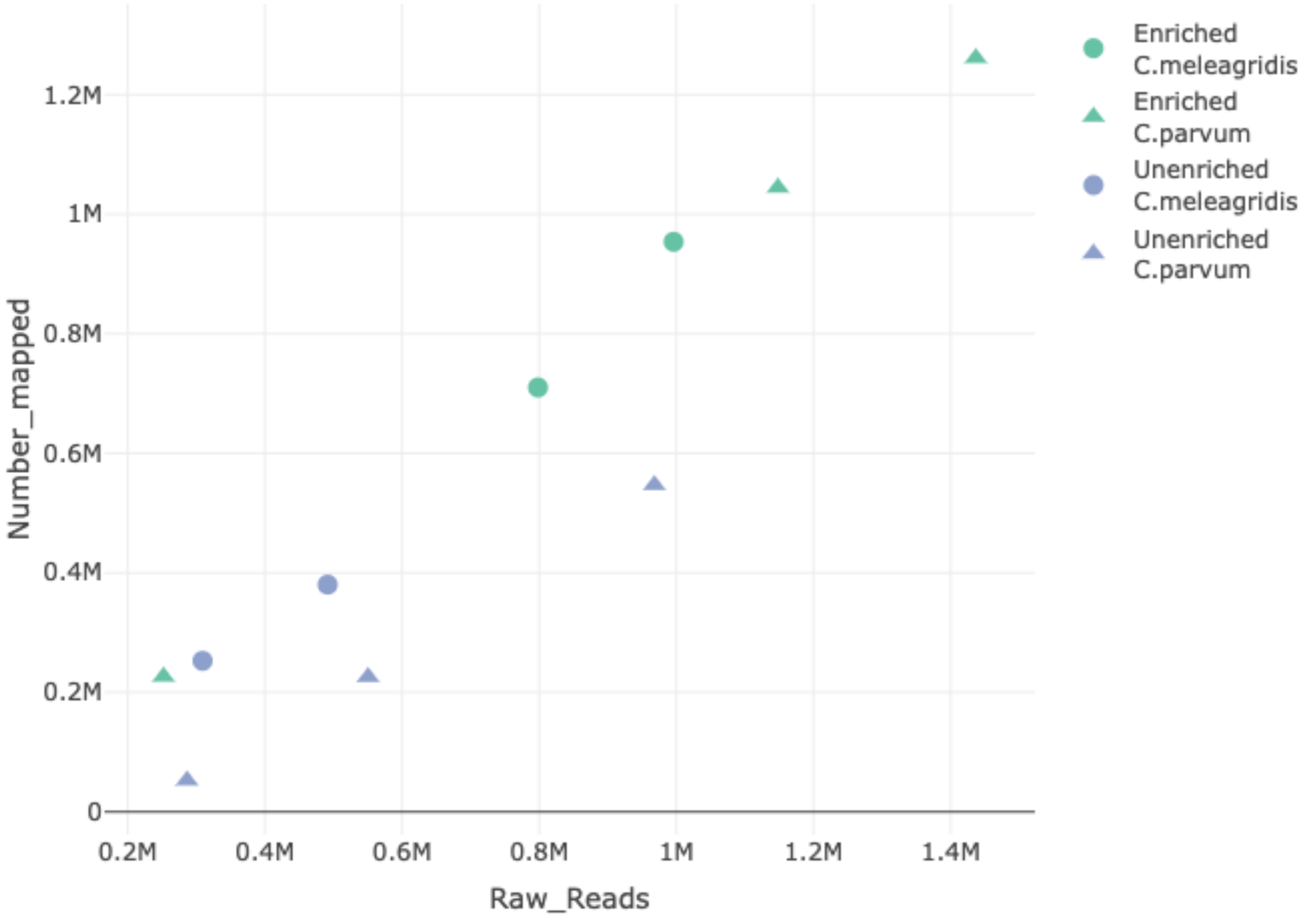


**Suppl. Fig. S14.** Scatterplot of the percentage of reads mapped to the respective *Cryptosporidium* genome given the number of raw reads obtained for a specific sample. In green, enriched libraries; in purple, unenriched libraries. In circles, samples from *C. parvum* pure oocysts, and in triangles, samples from *C. meleagridis* oocysts.

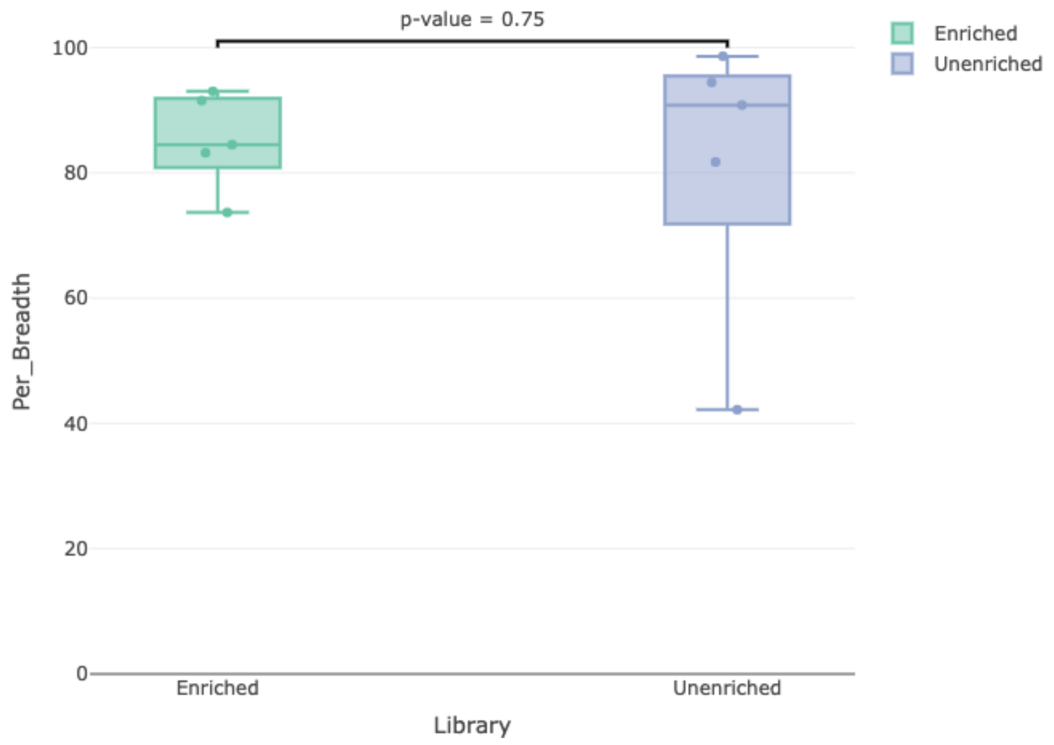


**Suppl. Fig. S15**. Boxplot of the percentage of the breadth of genome coverage obtained from the mapping of sequences from enriched (green) and unenriched (purple) libraries to the corresponding reference genome. These libraries were prepared from pure oocyst DNA, three samples of *C. parvum*, and two samples of *C. meleagridis*.

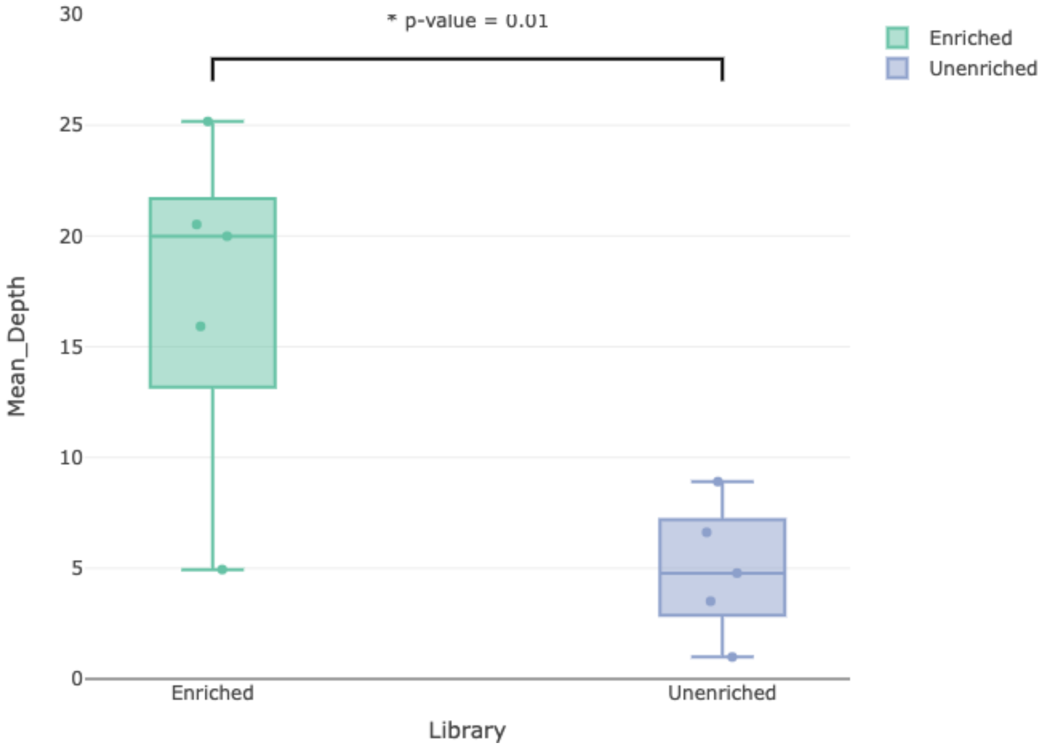


**Suppl. Fig. S16**. Boxplot of the mean depth of genome coverage obtained from the mapping of sequences from enriched (green) and unenriched (purple) libraries to the corresponding reference genome. These libraries were prepared from pure oocyst DNA, three samples of *C. parvum*, and two samples of *C. meleagridis*.

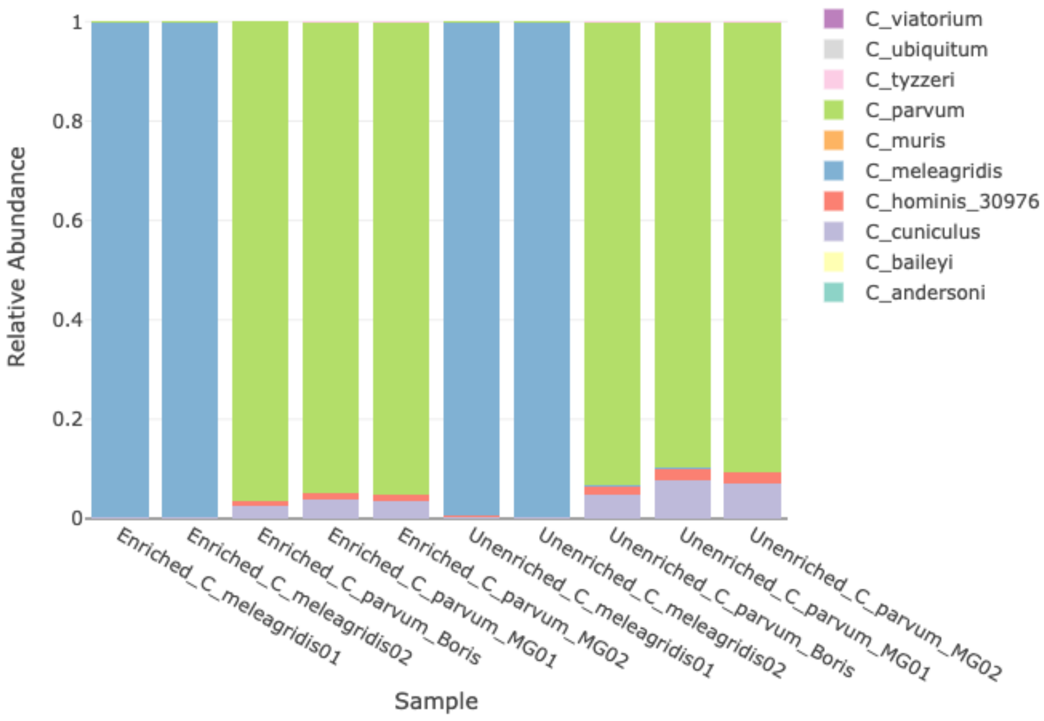


**Suppl. Fig. S17**. Stacked barplot of the relative abundance of hits from pure oocyst DNA samples from two *Cryptosporidium* species and two library types mapped to the ten genomes reference.

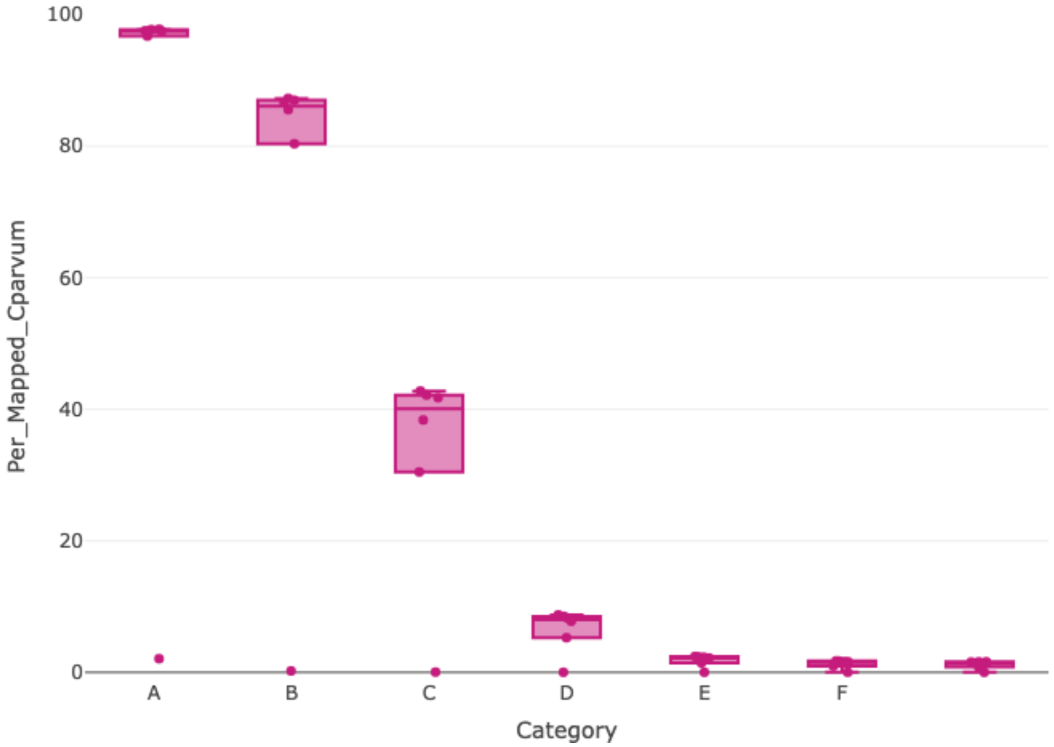


**Suppl. Fig. S18**. Boxplot of the percentage of paired reads that mapped to the *C. parvum* IOWA-ATCC reference genome for the *C. parvum* dilution experiment for six levels of serial 1:10 dilutions (A-F), and a negative control (G). A: 1 ng, B: 0.1 ng, C: 0.01 ng, D: 0.001 ng, E: 0.0001 ng, F: 0.00001 ng, G: 0 ng.

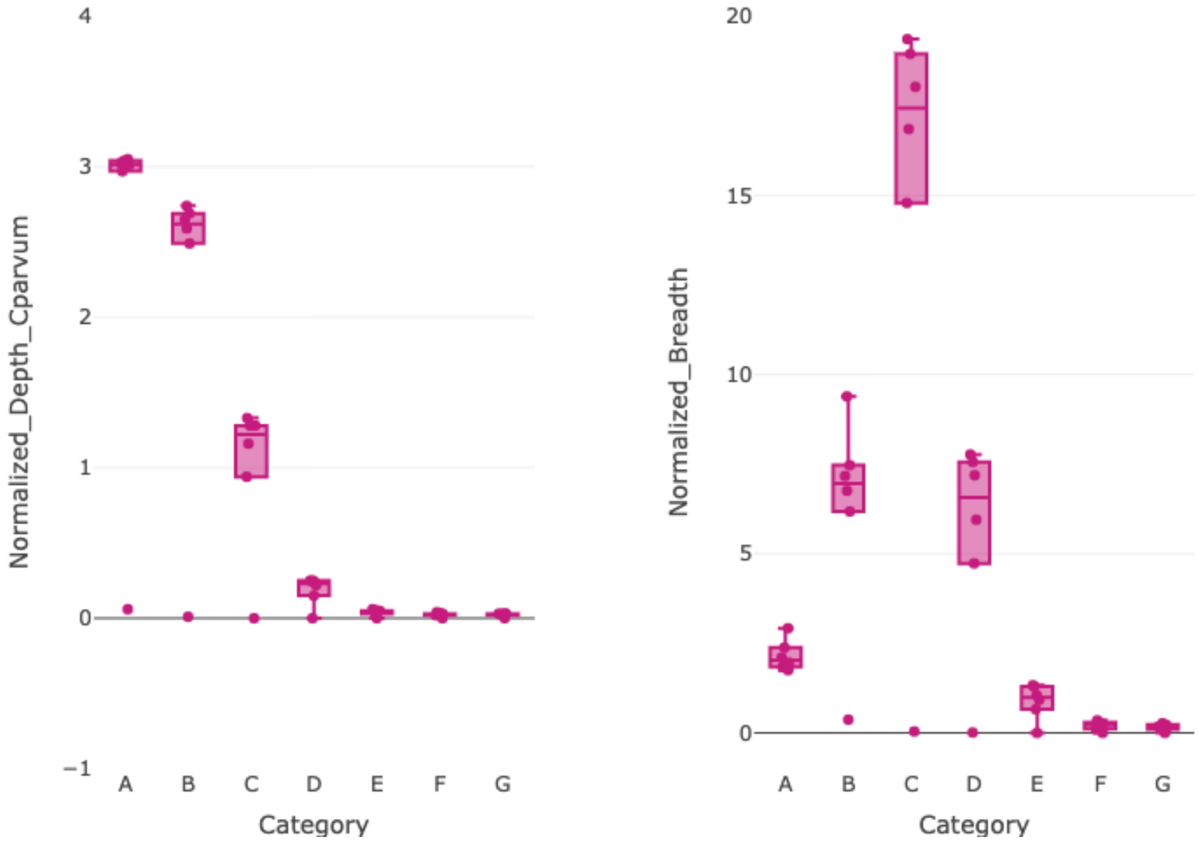


**Suppl. Fig. S19**. Plot of the depth and breadth of genome coverage normalized per 100,000 reads, obtained by the mapping of serial diluted *C. parvum* libraries prepared using the NEB protocol and iTru primers for six levels of serial 1:10 dilutions (A-F), and a negative control (G). A: 1 ng, B: 0.1 ng, C: 0.01 ng, D: 0.001 ng, E: 0.0001 ng, F: 0.00001 ng, G: 0 ng.

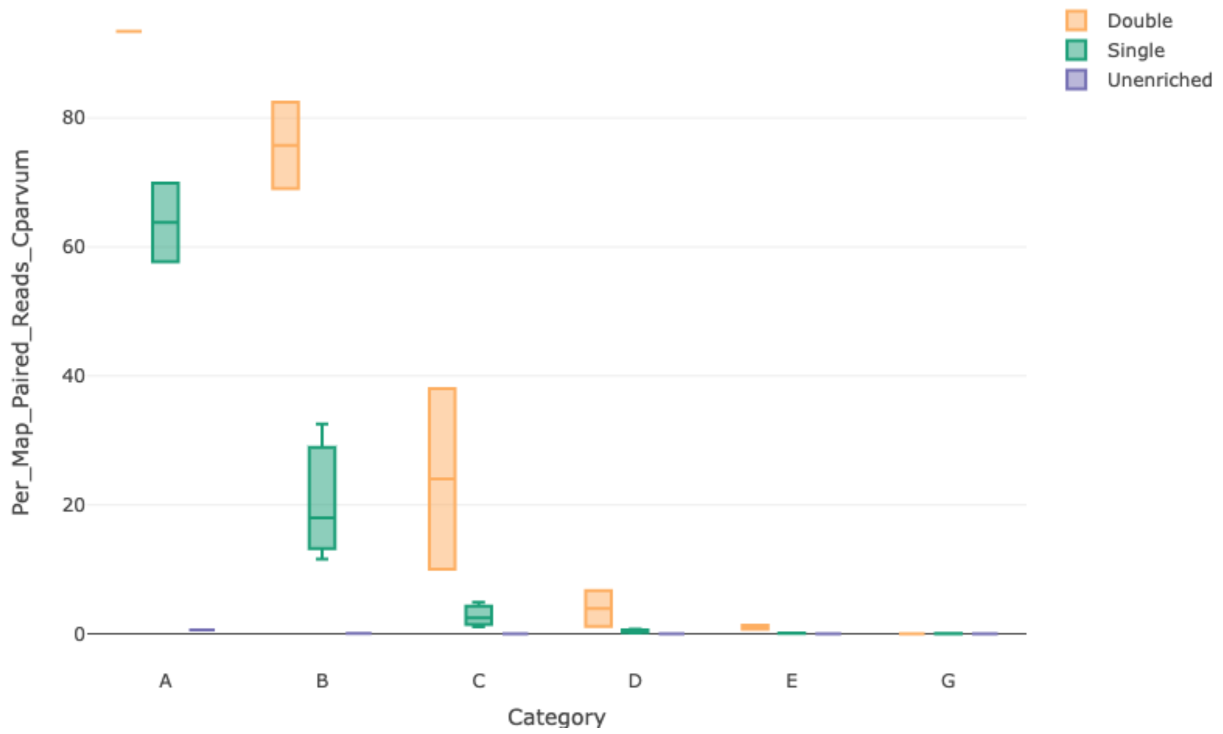


**Suppl. Fig. S20**. Plot of the percentage of raw reads that mapped to the *C. parvum* IOWA-ATCC reference genome for libraries of serially diluted *C. parvum* (A-E) and a negative control (G) prepared using the iNextEra protocol with different enrichment types and diluted baits. A: 1 ng, B: 0.1 ng, C: 0.01 ng, D: 0.001 ng, E: 0.0001 ng, and G: 0 ng.

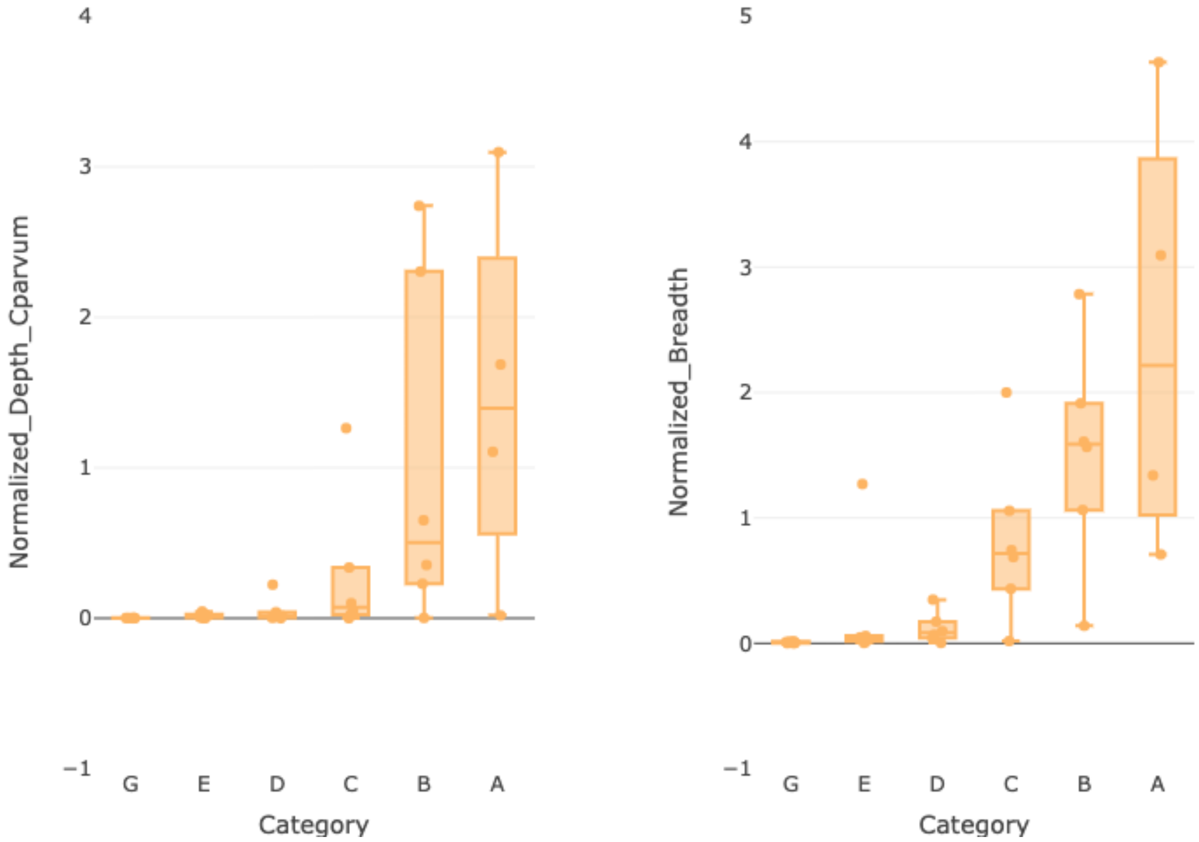


**Suppl. Fig. S21.** Plot of the depth and breadth of *C. parvum* IOWA-ATCC genome coverage, normalized per 100,000 reads, obtained for libraries of serially diluted *C. parvum* (A-E) and a negative control (G) prepared using the iNextEra protocol and different enrichment types. A: 1 ng, B: 0.1 ng, C: 0.01 ng, D: 0.001 ng, E: 0.0001 ng, and G: 0 ng.

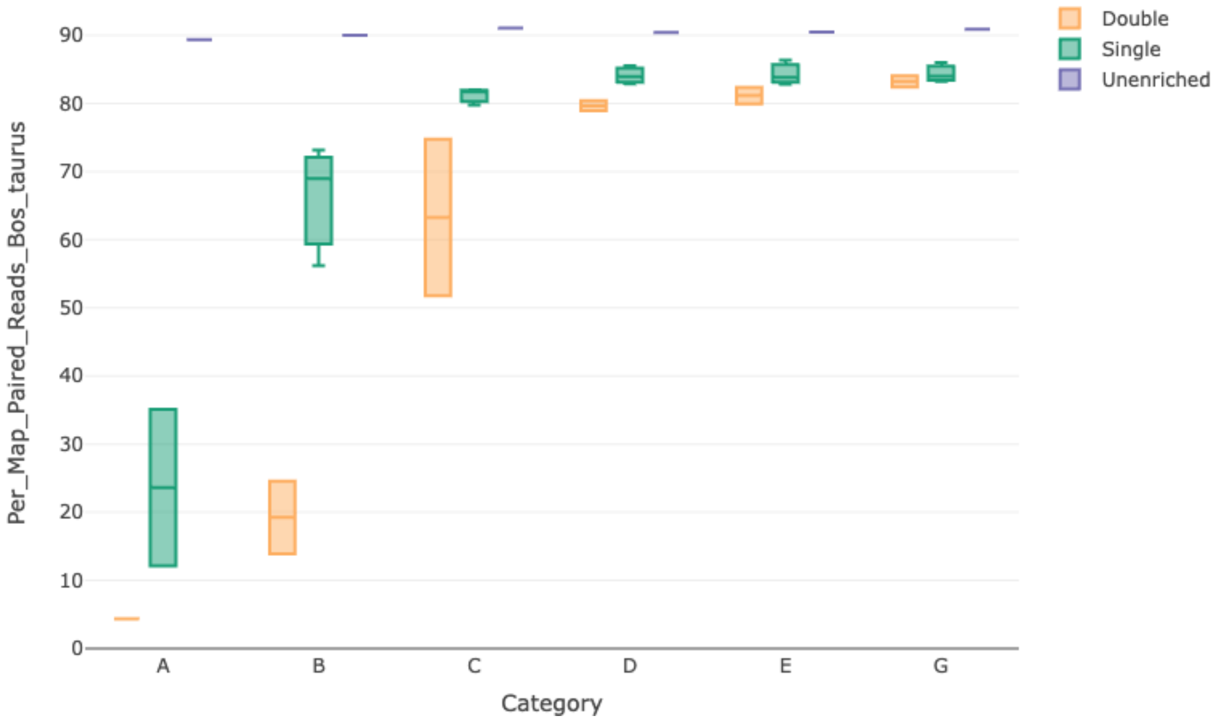


**Suppl. Fig. S22**. Plot of the percentage of raw reads that mapped to the *Bos taurus* reference genome (GCA_002263795.3) for libraries of serially diluted *C. parvum* (A-E) and a negative control (G) prepared using the iNextEra protocol with different enrichment types and diluted baits. A: 1 ng, B: 0.1 ng, C: 0.01 ng, D: 0.001 ng, E: 0.0001 ng, and G: 0 ng.

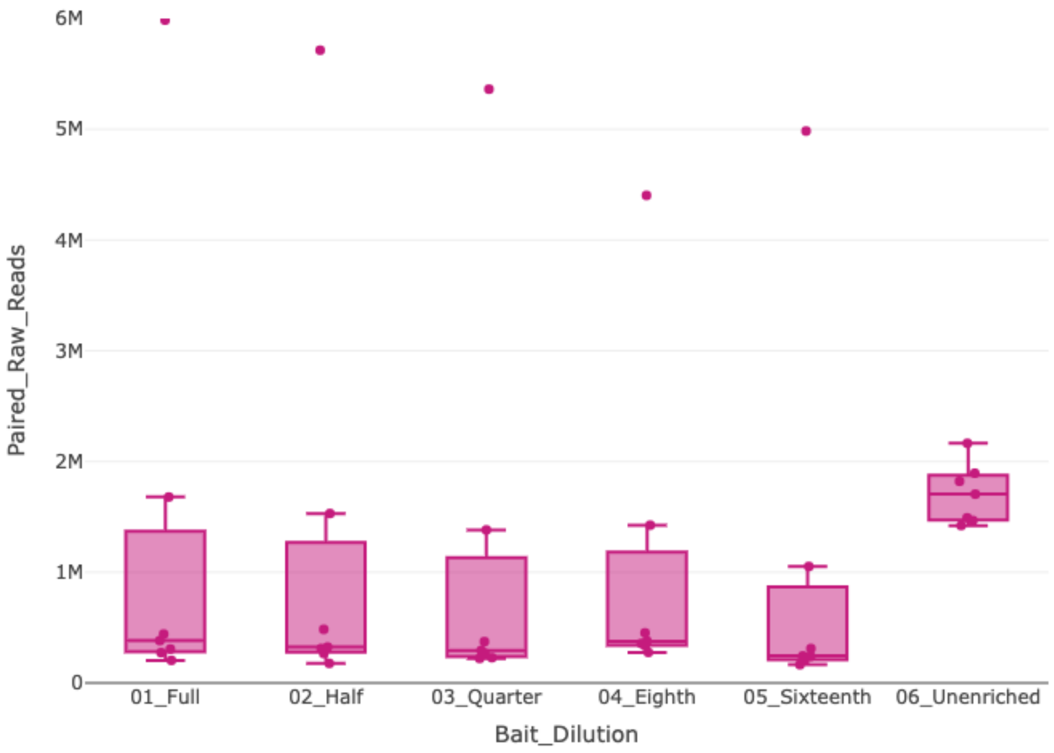


**Suppl. Fig. S23.** Number of paired reads obtained for CryptoCap_100k enriched *C. parvum* libraries with different amounts of baits (with respect to the standard amount from the manufacturer) or left unenriched. These libraries were prepared using the NEB protocol and iTru primers.

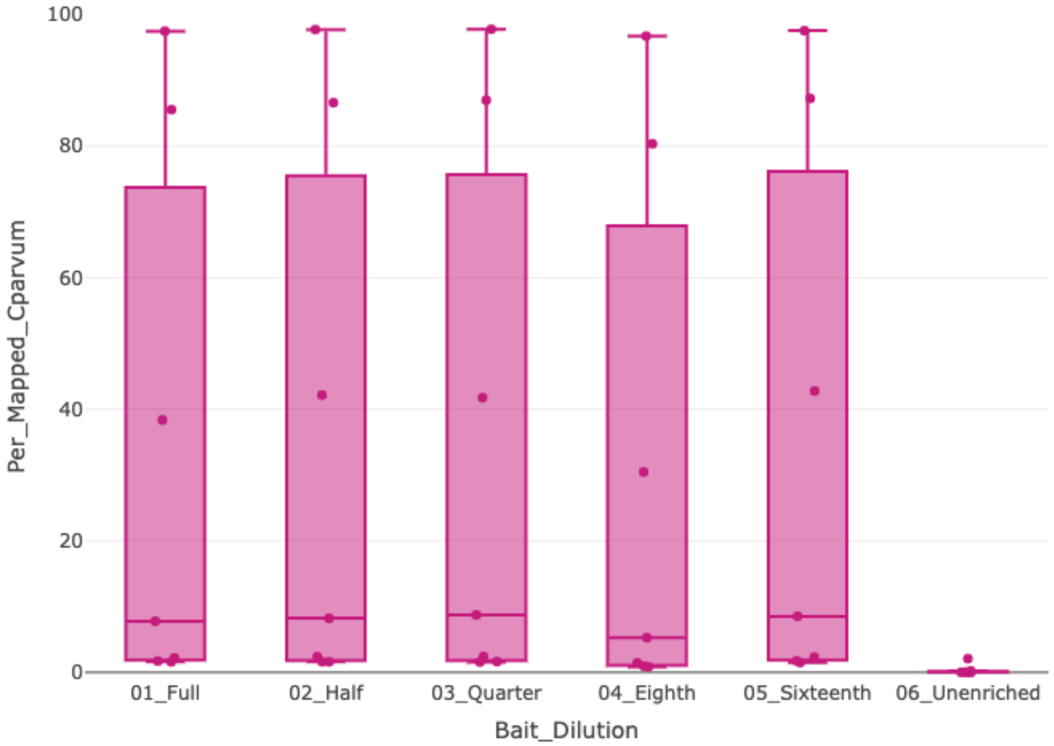


**Suppl. Fig. S24.** Percentage of paired reads that mapped to the *C. parvum* IOWA-ATCC reference genome, obtained for CryptoCap_100k enriched *C. parvum* libraries with different amounts of baits (with respect to the standard amount from the manufacturer) or left unenriched. These libraries were prepared using the NEB protocol and iTru primers.

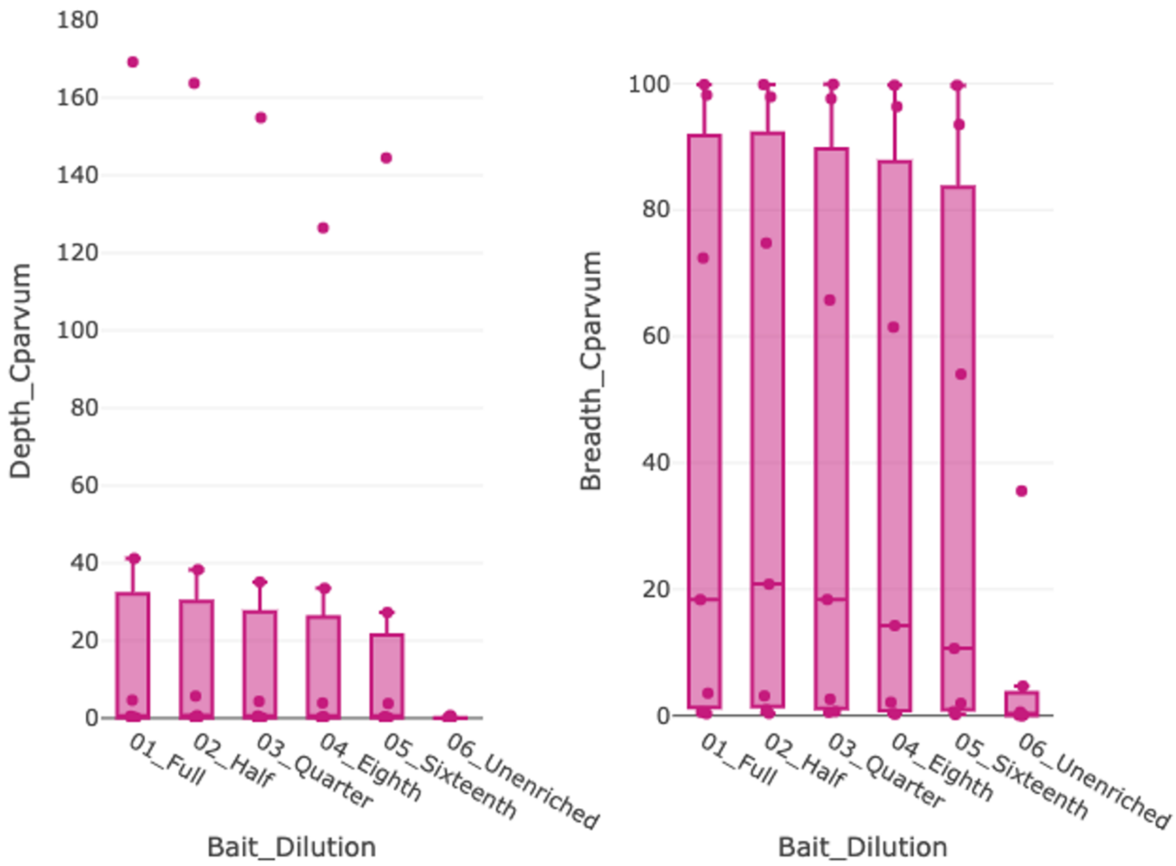


**Suppl. Fig. S25.** Plot of the depth and breadth of *C. parvum* IOWA-ATCC genome coverage obtained by the mapping of *C. parvum* sequence reads from libraries prepared with different amounts of baits (with respect to the standard amount from the manufacturer) or left unenriched. These libraries were prepared using the NEB protocol and iTru primers.

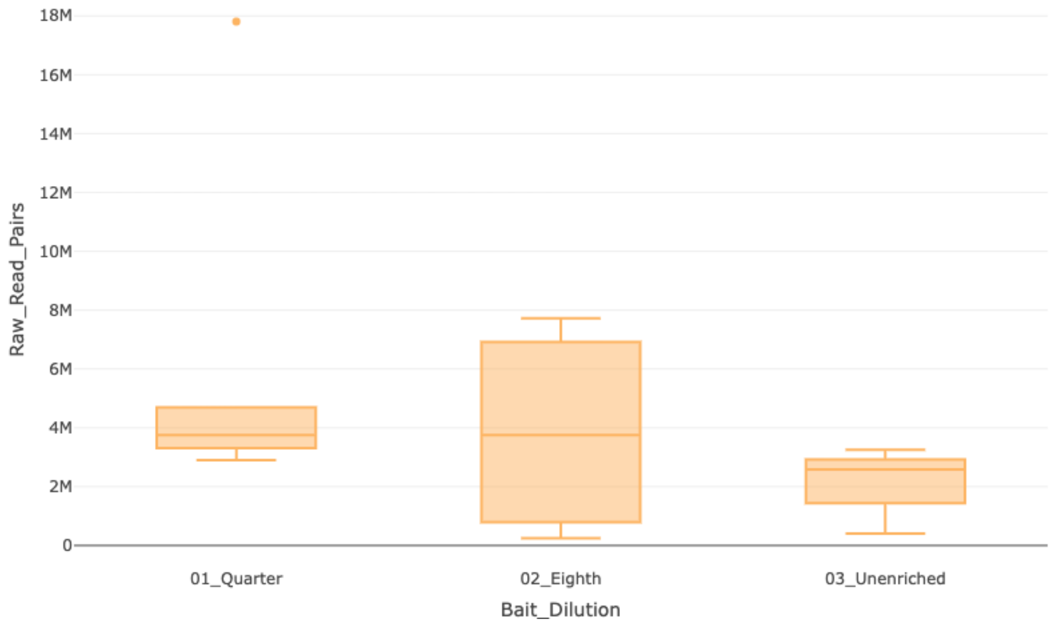


**Suppl. Fig. S26.** Number of paired reads obtained for CryptoCap_100k enriched *C. parvum* libraries with different amounts of baits (with respect to the standard amount from the manufacturer) or left unenriched. These libraries were prepared using the iNextEra protocol and iNext primers.

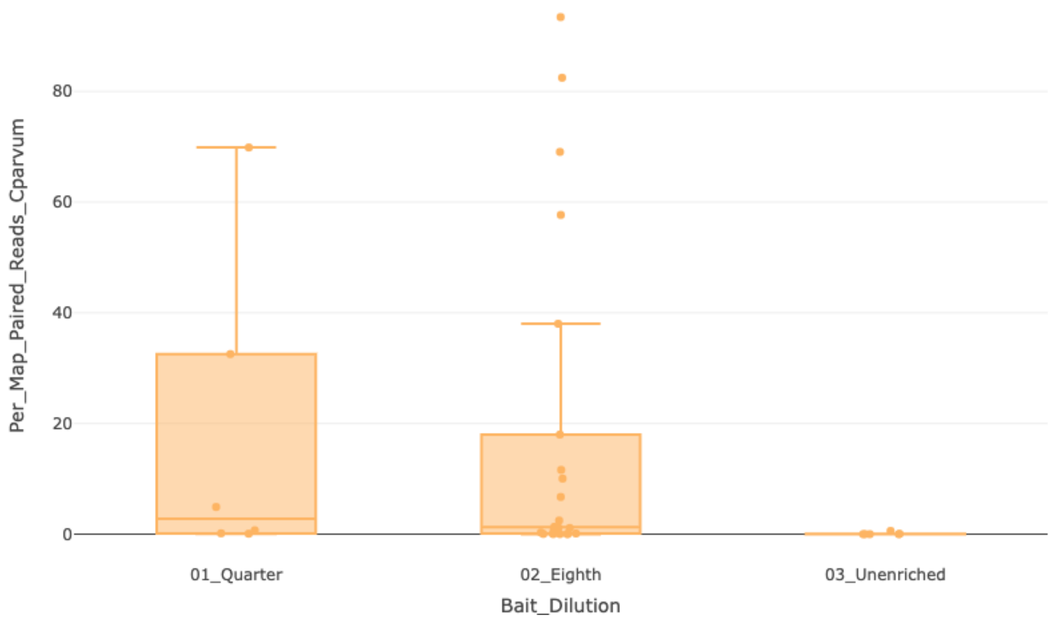


**Suppl. Fig. S27.** Percentage of paired reads that mapped to the *C. parvum* IOWA-ATCC reference genome, obtained for CryptoCap_100k enriched *C. parvum* libraries with different amounts of baits (with respect to the standard amount from the manufacturer) or left unenriched. These libraries were prepared using the iNextEra protocol and iNext primers.

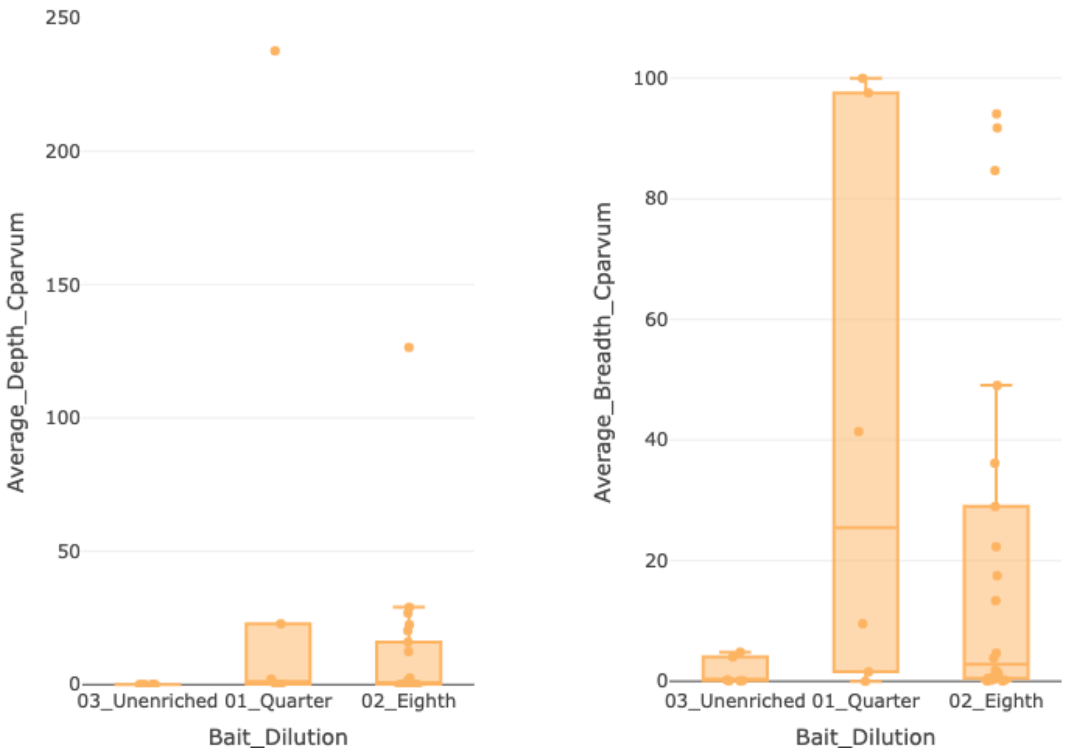


**Suppl. Fig. S28.** Plot of the depth and breadth of *C. parvum* IOWA-ATCC genome coverage obtained by the mapping of *C. parvum* sequence reads from libraries prepared with different amounts of baits (with respect to the standard amount from the manufacturer) or left unenriched. These libraries were prepared using the iNextEra protocol and iNext primers.

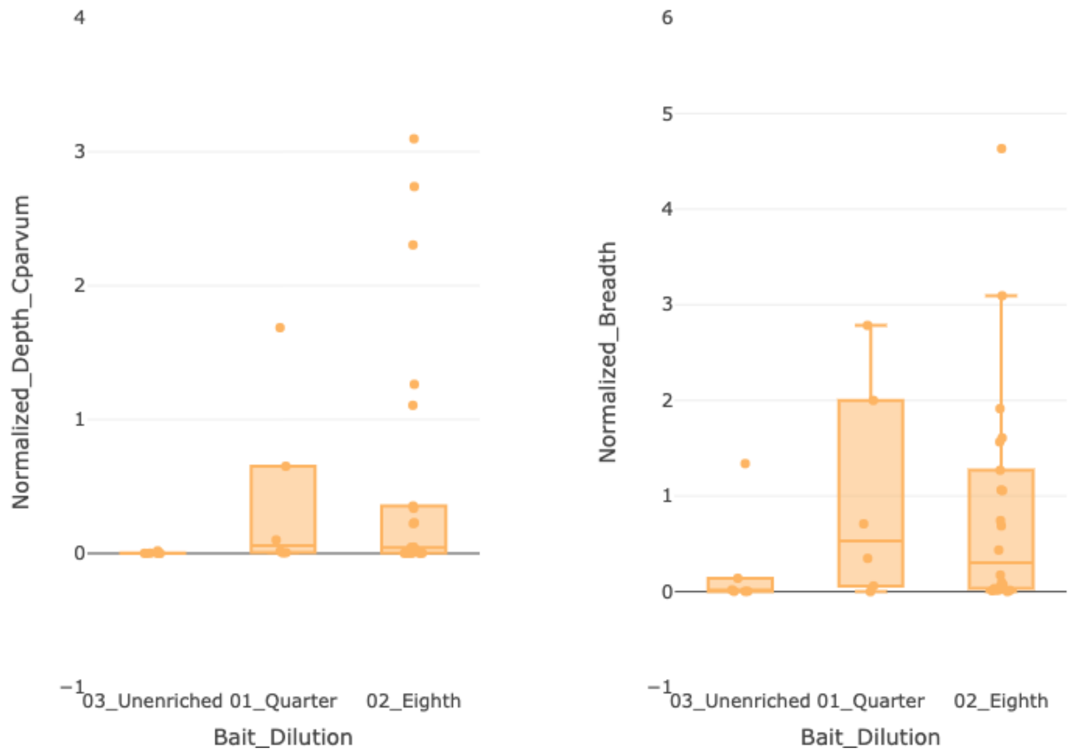


**Suppl. Fig. S29.** Plot of the depth and breadth of *C. parvum* IOWA-ATCC genome coverage normalized per 100,000 reads, obtained by the mapping of *C. parvum* sequence reads from libraries prepared with different amounts of baits (with respect to the standard amount from the manufacturer) or left unenriched. These libraries were prepared using the iNextEra protocol and iNext primers.

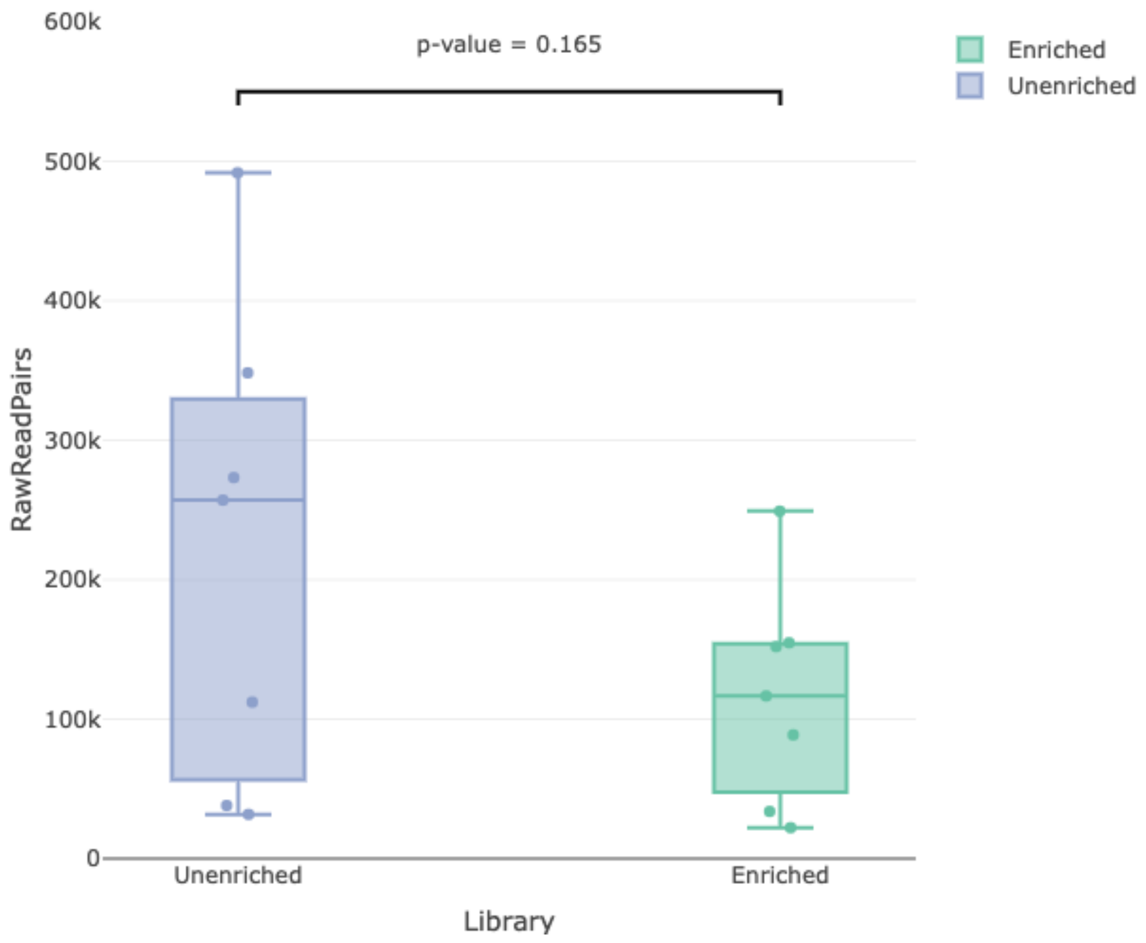


**Suppl. Fig. S30.** Boxplot of the number of raw paired reads per sample obtained for enriched and unenriched libraries from the seven clinical samples set that were prepared with the NEB protocol with iTru primers.

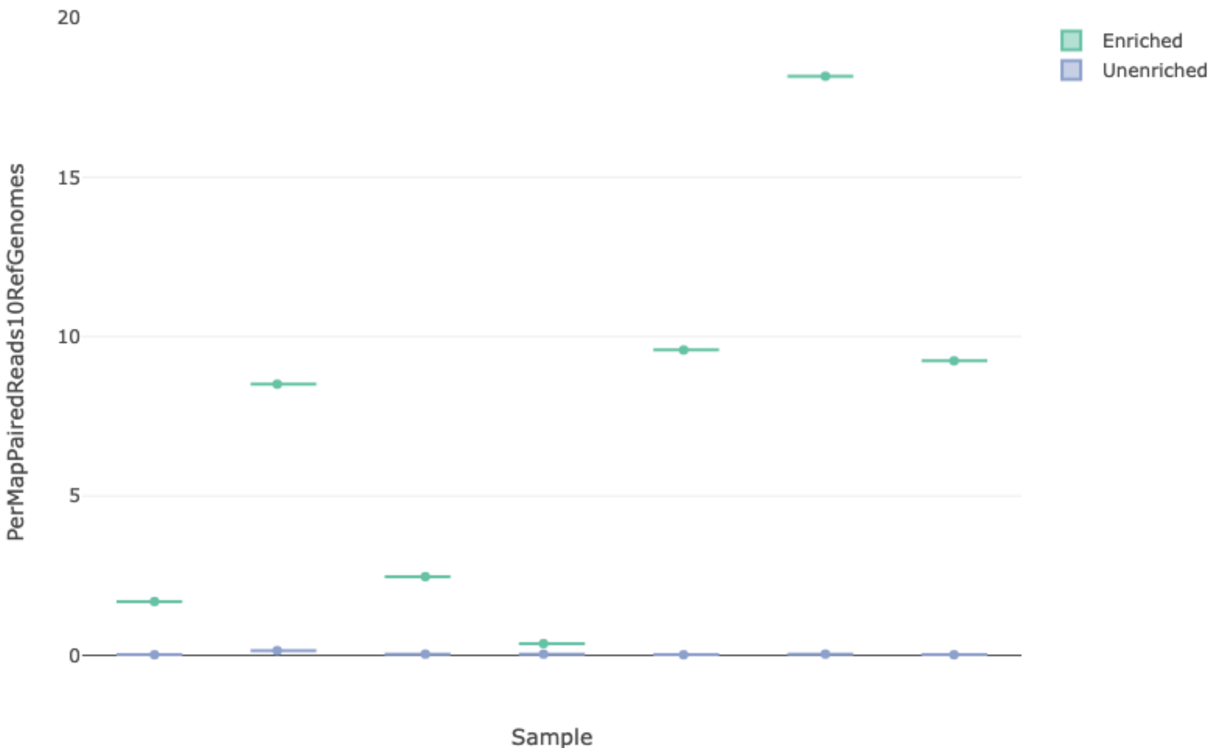


**Suppl. Fig. S31.** Percentage of reads per sample that mapped to the ten-genome reference (10-CrypGS) for enriched and unenriched libraries made from seven clinical samples. These libraries were prepared with the NEB protocol with iTru primers and single-enriched with CryptoCap_100k.

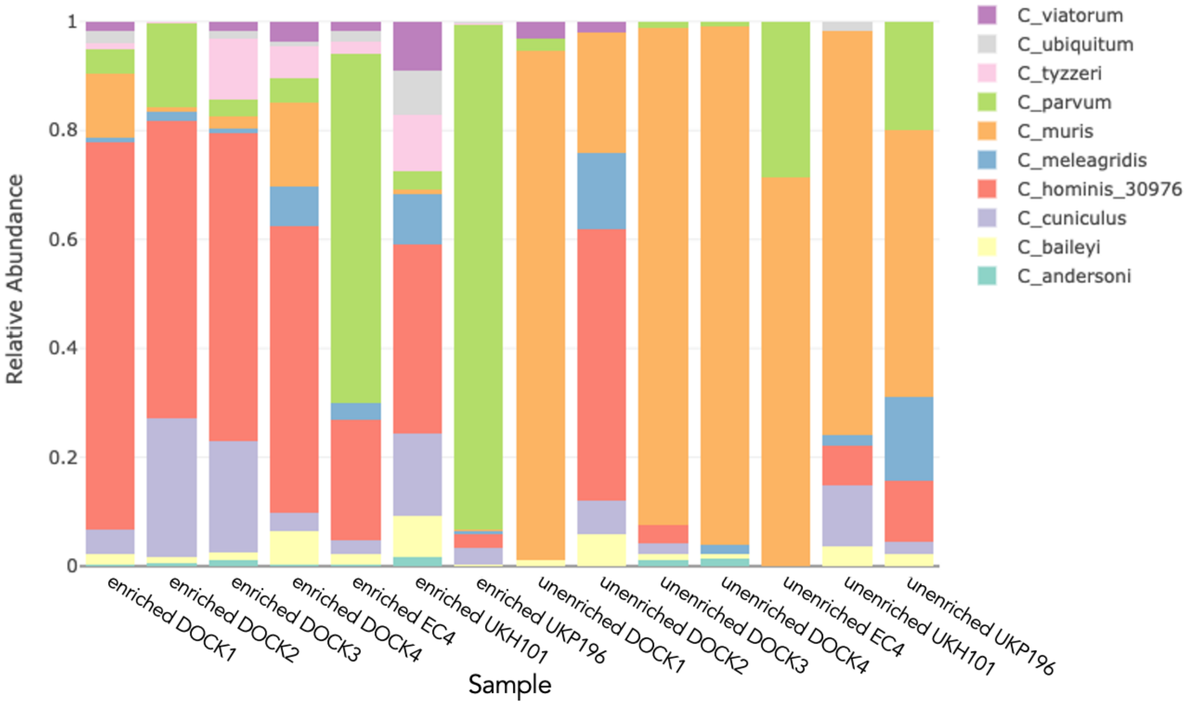


**Suppl. Fig. S32**. Stacked barplot of relative abundance of species detected from the mapping of sequencing reads from seven clinical samples to the reference containing ten *Cryptosporidium* species genome sequences (10-CrypGS). These libraries were prepared with the NEB protocol with iTru primers. Libraries on the left represent those single-enriched with CryptoCap_100k. Libraries on the right represent those being left unenriched.

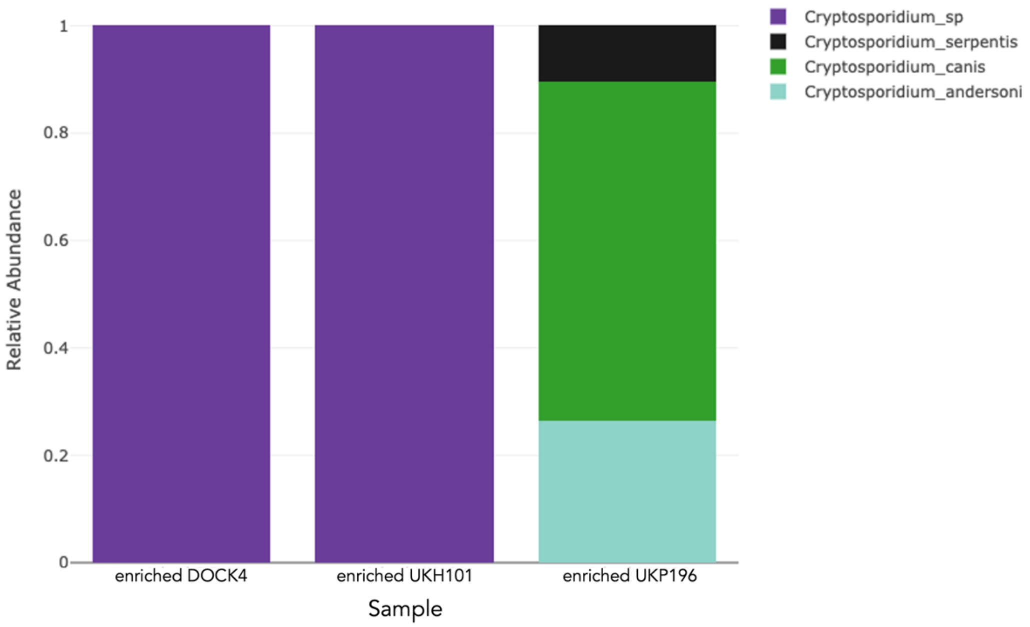


**Suppl. Fig. S33**. Stacked barplot of relative abundance of species detected from the mapping of sequencing reads from seven clinical samples to the 18S rRNA Crypto DB reference database. These libraries were prepared with the NEB protocol with iTru primers and single-enriched with CryptoCap_100k.

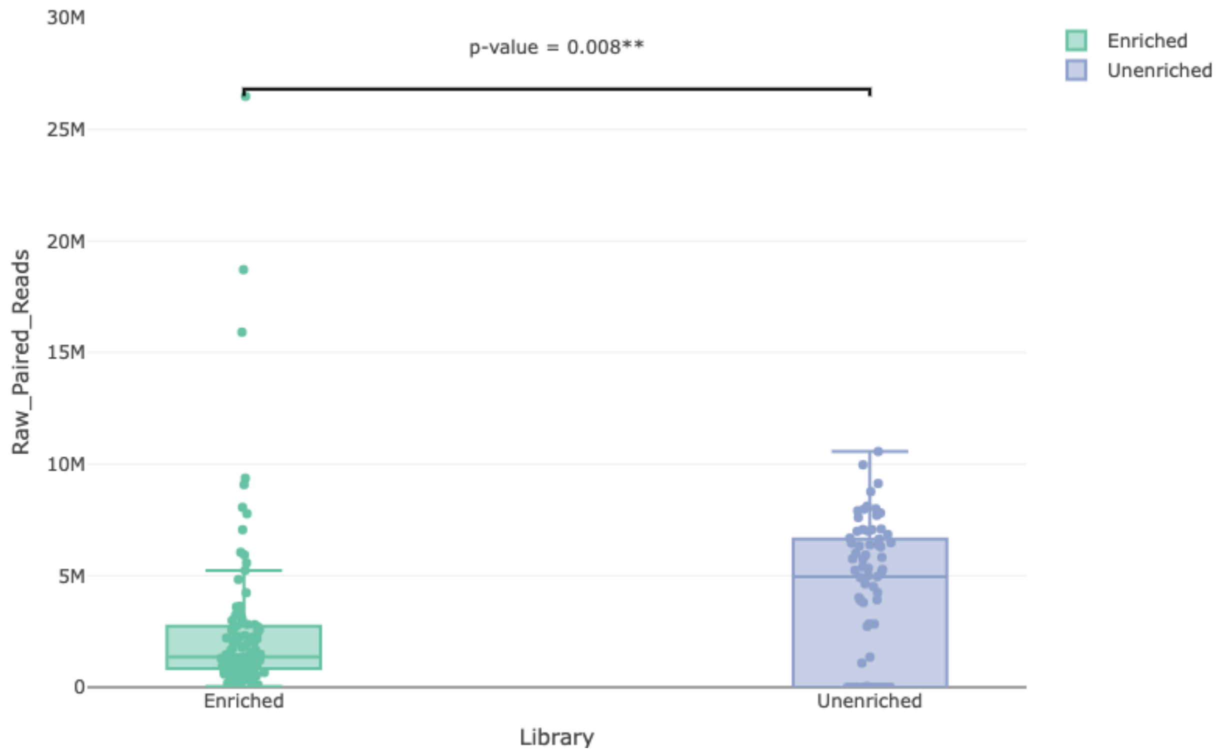


**Suppl. Fig. S34.** Boxplot of the raw number of reads obtained from the sequencing of single-enriched with CryptoCap_100k (left, green) and left unenriched (right, purple) libraries prepared from the set of 100 clinical samples. These libraries were prepared with the NEB protocol with iTru primers.

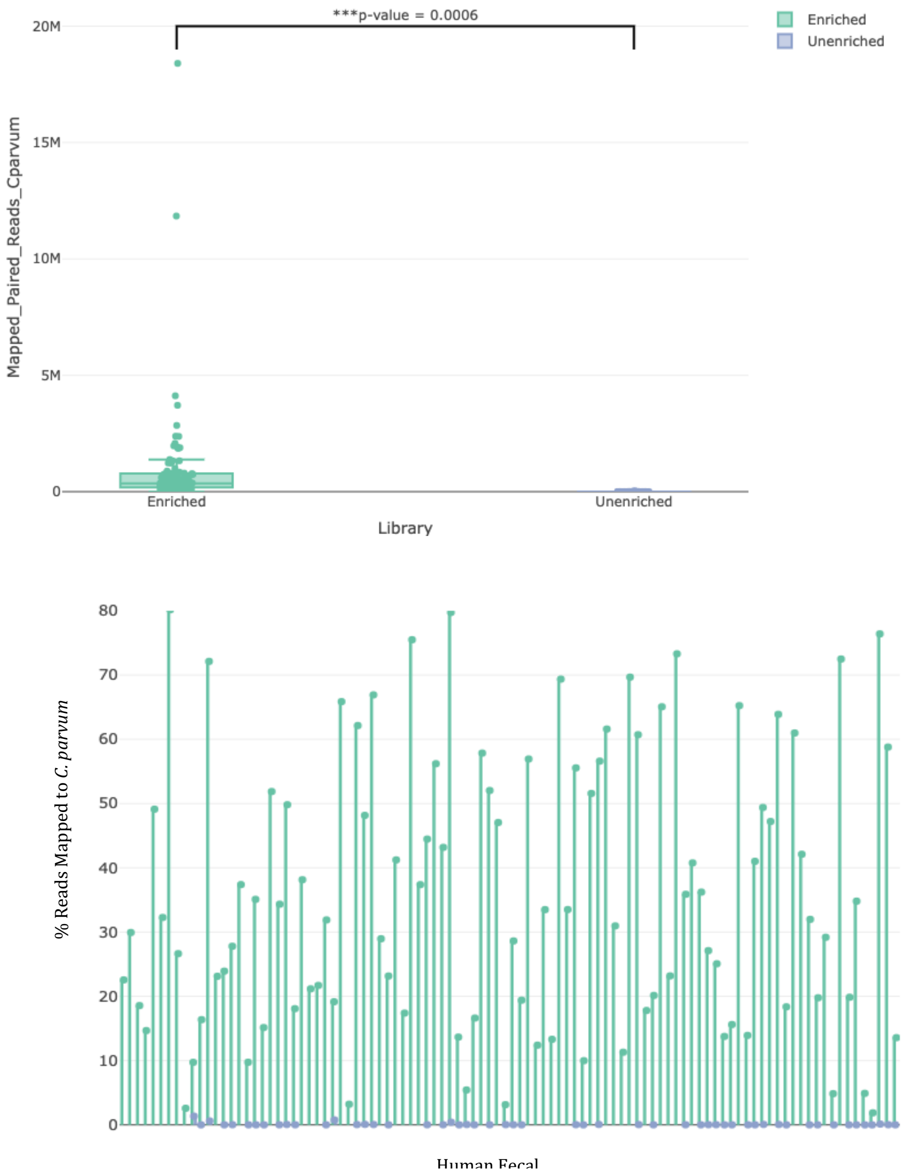


**Suppl. Fig. S35.** Top, boxplot of the number of reads obtained from sequencing. Bottom, percentage of reads mapped per sample (x-axis labels not shown but in order from left to right, sample 1 to sample 100). Green single-enriched with CryptoCap_100k. Purple left unenriched. Libraries were prepared from the set of 100 clinical samples that mapped to the *C. parvum* IOWA-ATCC genome. These libraries were prepared with the NEB protocol with iTru primers.

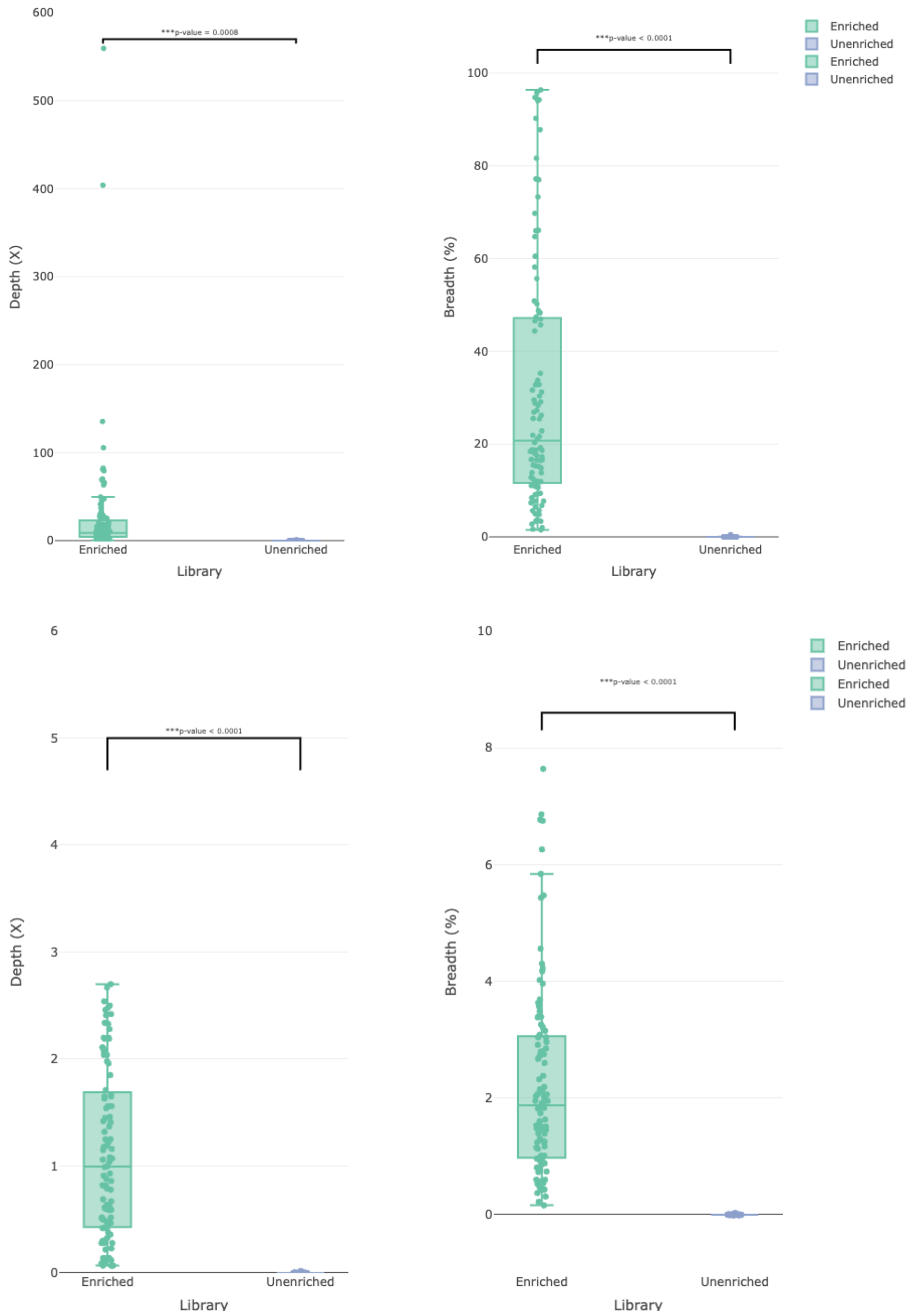


**Suppl. Fig. S36.** Plot of the depth and breadth of genome coverage. Top, raw coverage data. Bottom, normalized by 100,000 reads. Statistics obtained by the mapping of enriched (green) and unenriched (purple) sequence reads to the *C. parvum* IOWA-ATCC reference genome from libraries prepared from 100 clinical samples using the NEB protocol.

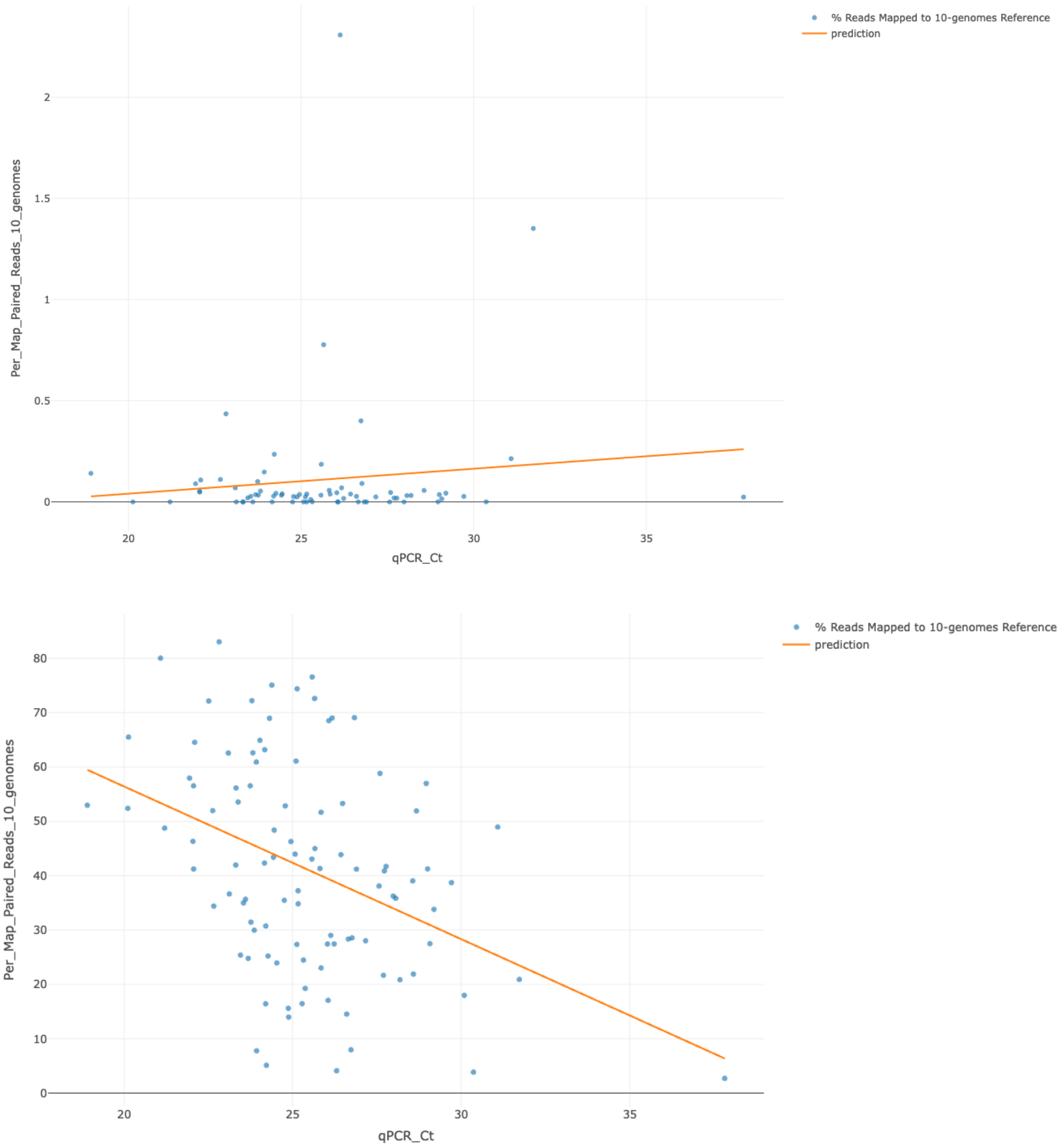


**Suppl. Fig. S37.** Linear regression model for predicting the percentage of mapped reads to a reference database containing ten Cryptosporidium genome sequences (10-CrypGS) given a $C_T$ value from qPCR. Top, unenriched libraries. Bottom, single-enriched libraries with CryptoCap_100k.

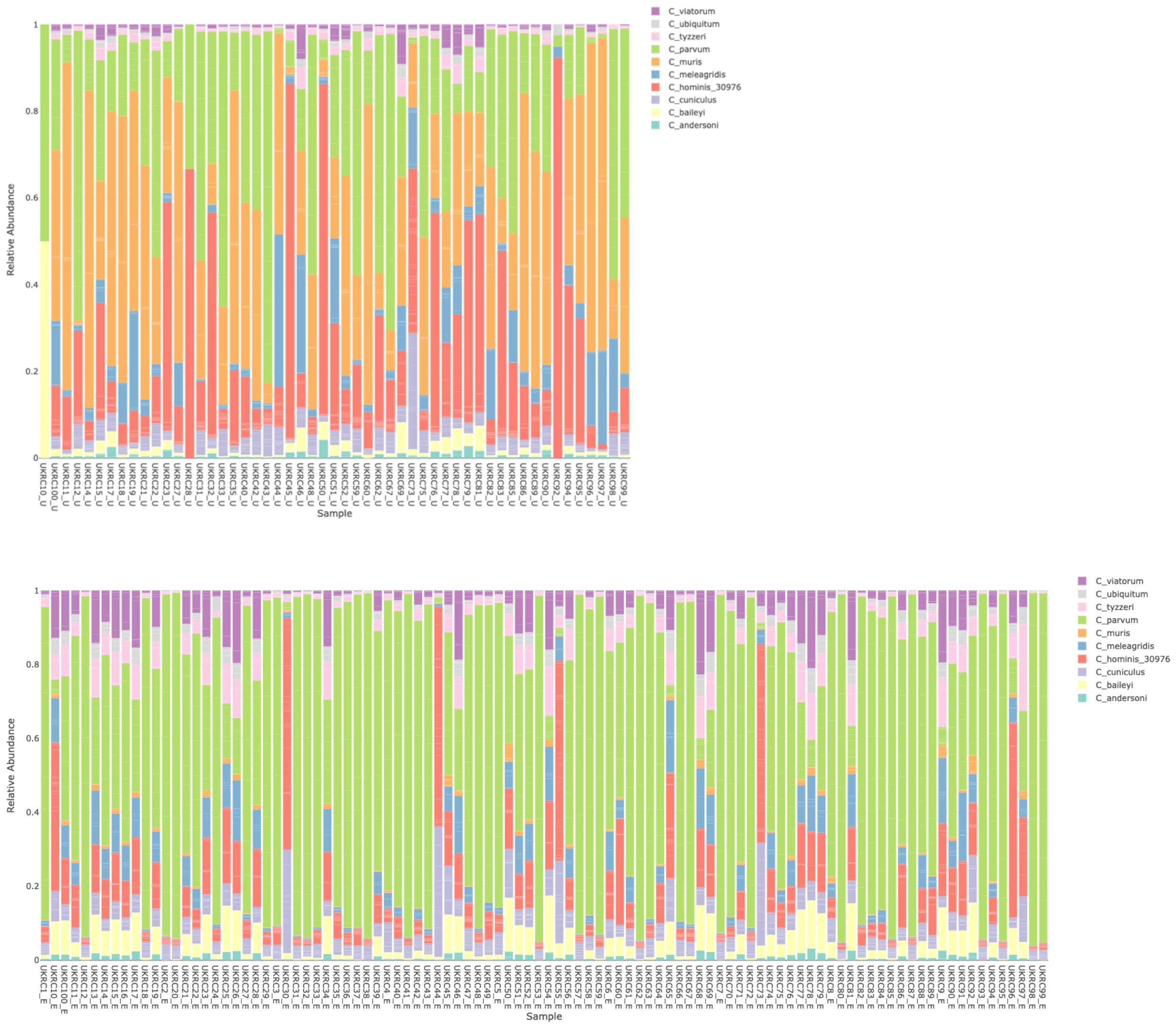


**Suppl. Fig. S38.** Species assignment when mapped to 10-CrypGS for libraries left unenriched (top), and enriched libraries with CryptoCap_100k (bottom).

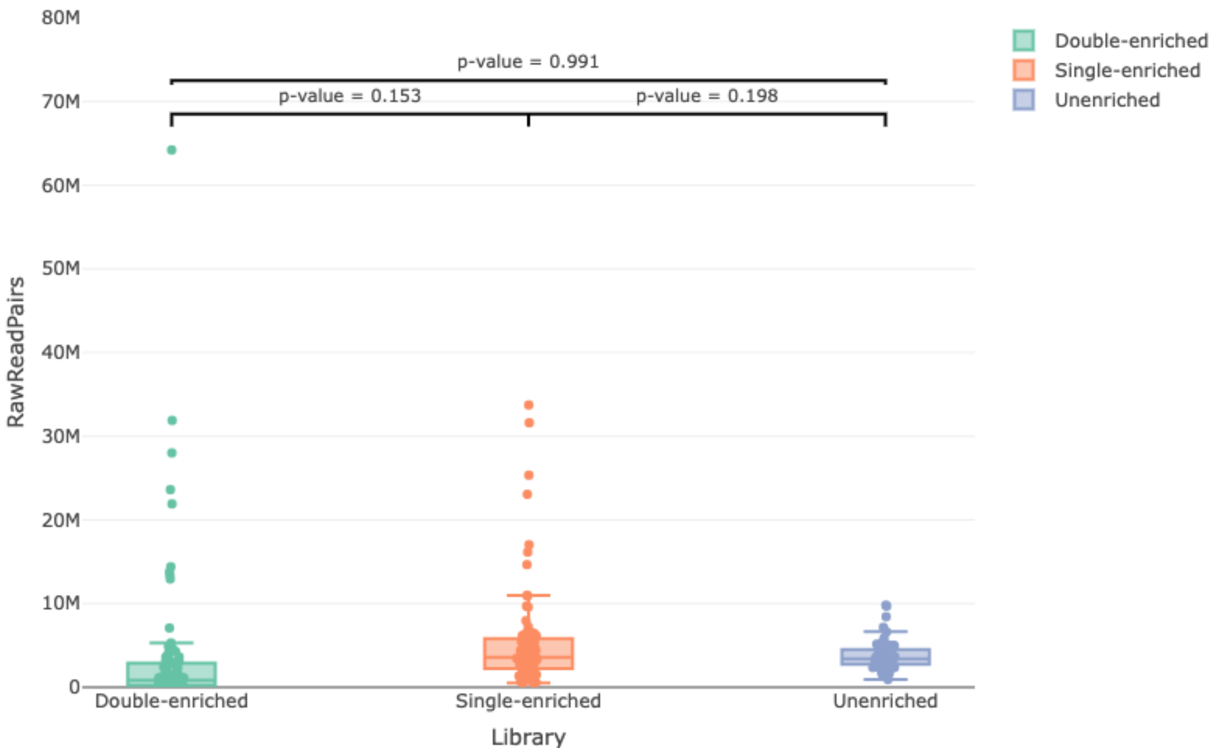


**Suppl. Fig. S39.** Boxplot of the raw number of paired reads from double-enriched (green), single-enriched (orange), and unenriched (purple) libraries prepared from 91 clinical samples set with the iNextEra protocol.

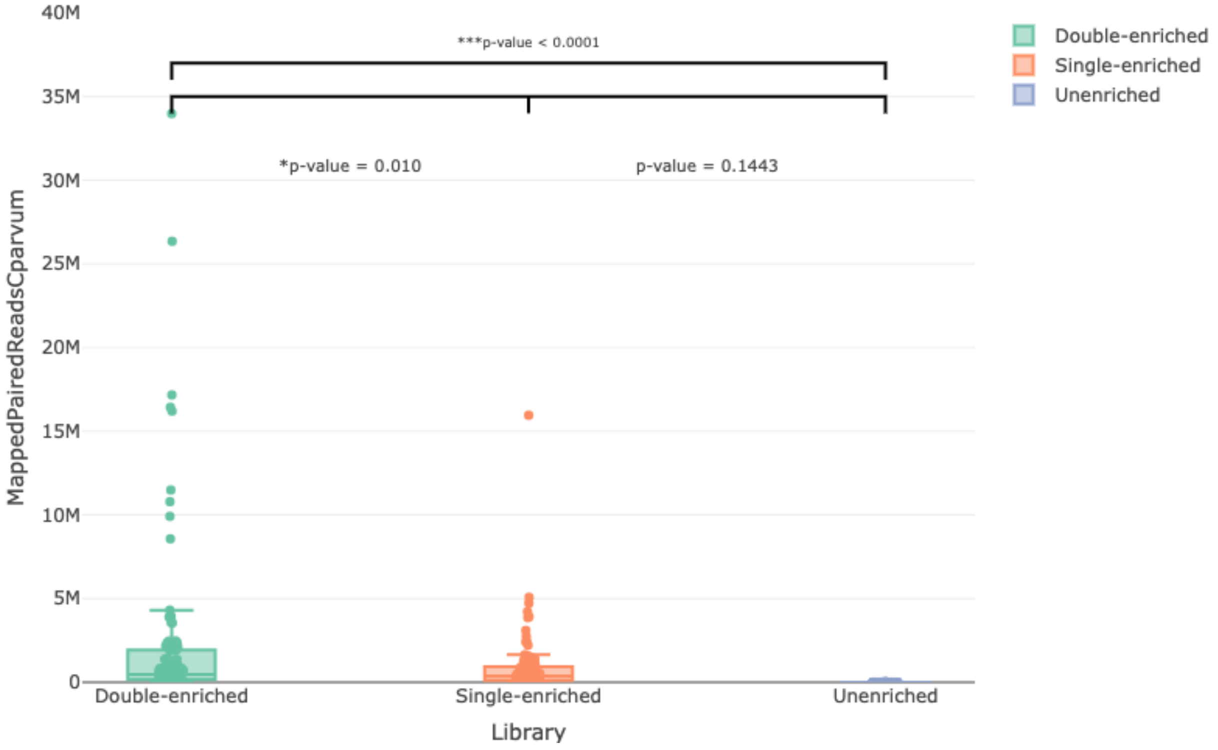


**Suppl. Fig. S40.** Boxplot of the average number of reads from across 91 clinical samples set that mapped to the *C. parvum* IOWA-ATCC reference genome from double-enriched (green), single-enriched (orange), and unenriched (purple) libraries prepared with the iNextEra protocol.

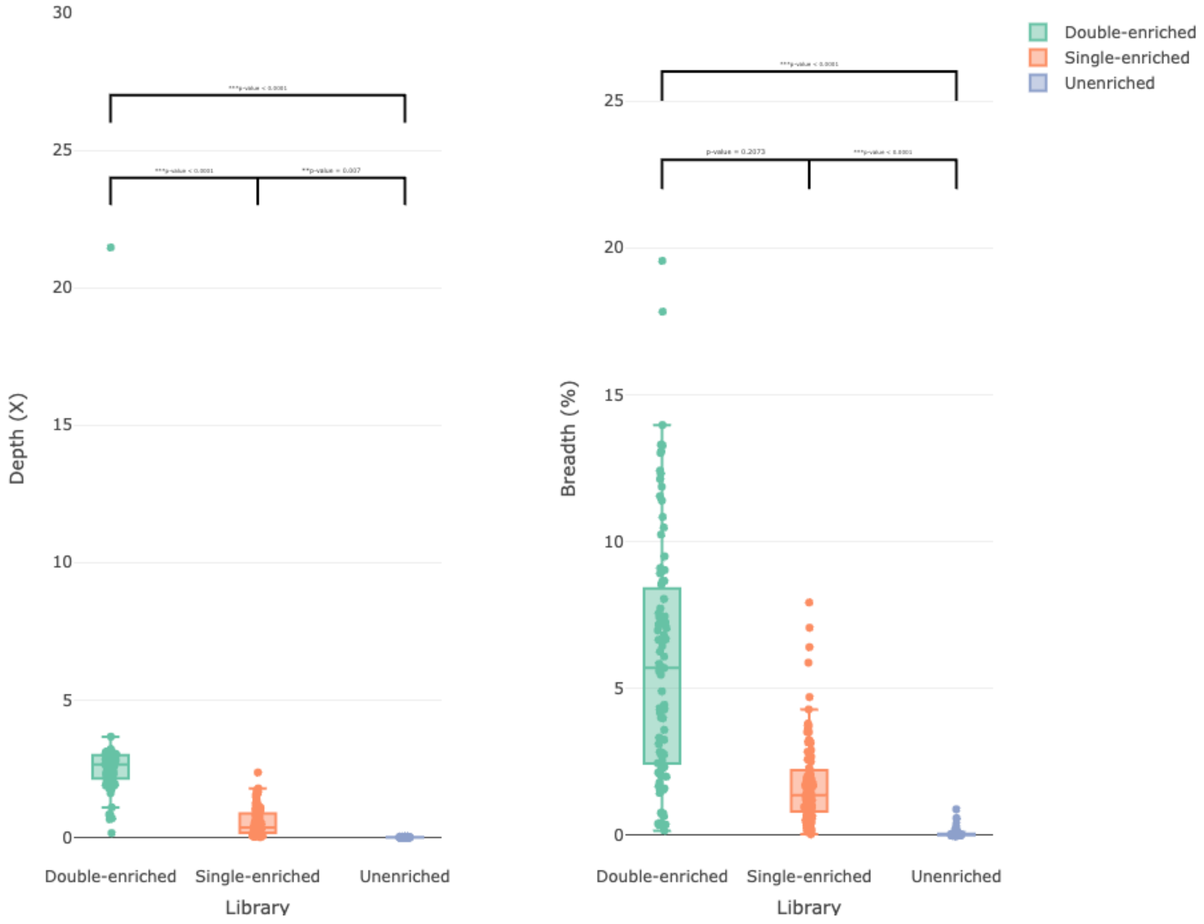


**Suppl. Fig. S41.** Plot of the depth and breadth of genome coverage normalized by 100,000 reads obtained by the mapping to the *C. parvum* IOWA-ATCC reference genome from double-enriched (green), single-enriched (orange), and unenriched (purple) libraries prepared from 91 clinical samples using the iNextEra protocol.

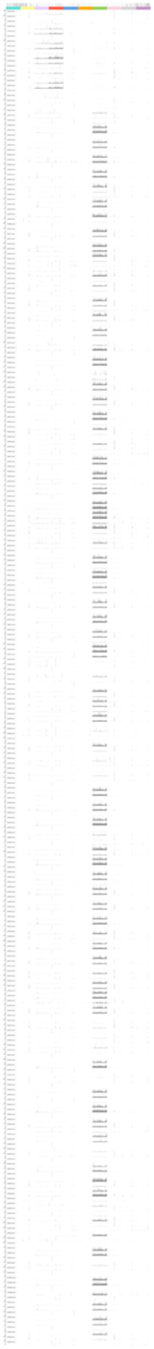

**Suppl. Fig. S42.** IVG plot of the depth and breadth of genome coverage obtained by the mapping to the 10-CrypGS reference file from double-enriched (D), single-enriched (S), and unenriched (U) libraries prepared from 91 clinical samples using the iNextEra protocol. Colors of genome sequences correspond to those used to reference *Cryptosporidium* species in the paper. A full-size resolution image can be found at: https://doi.org/10.6084/m9.figshare.29621024.

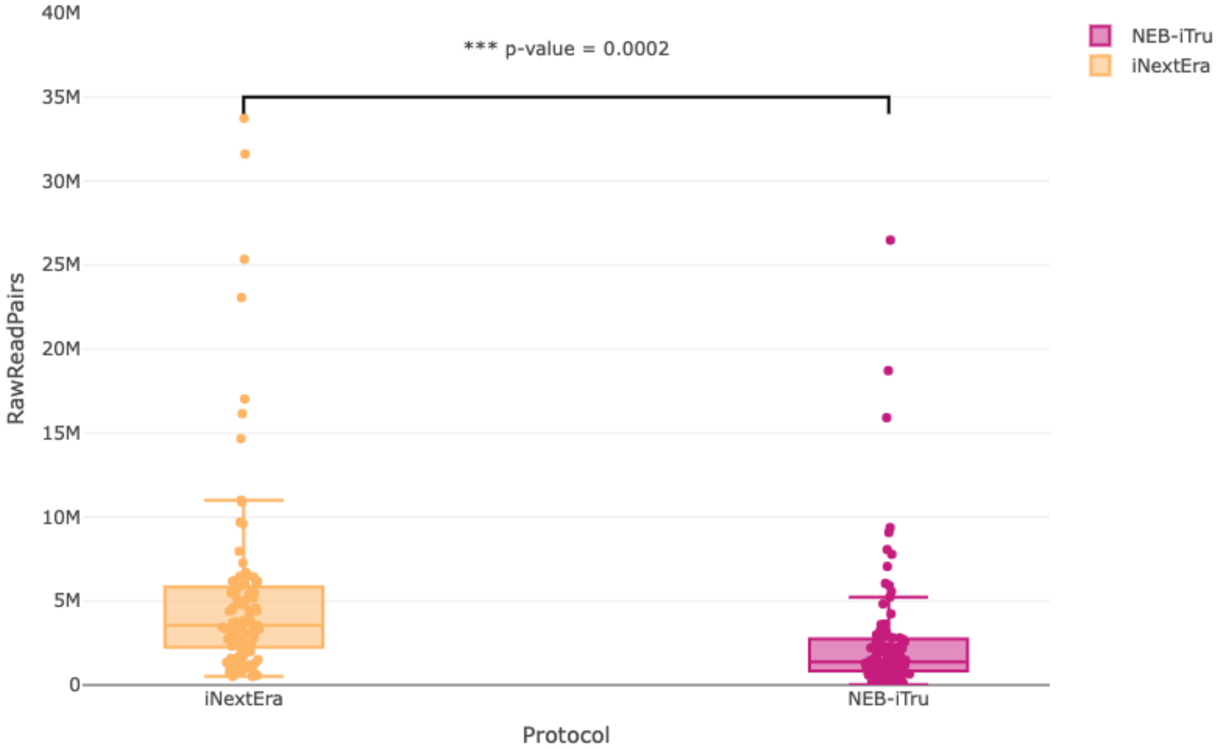


**Suppl. Fig. S43.** Boxplot of the number of raw paired reads per sample obtained for single-enriched libraries from the 100-clinical samples set that were prepared using either the NEB-iTru (pink) protocol or the iNextEra-iNext (orange) protocol.

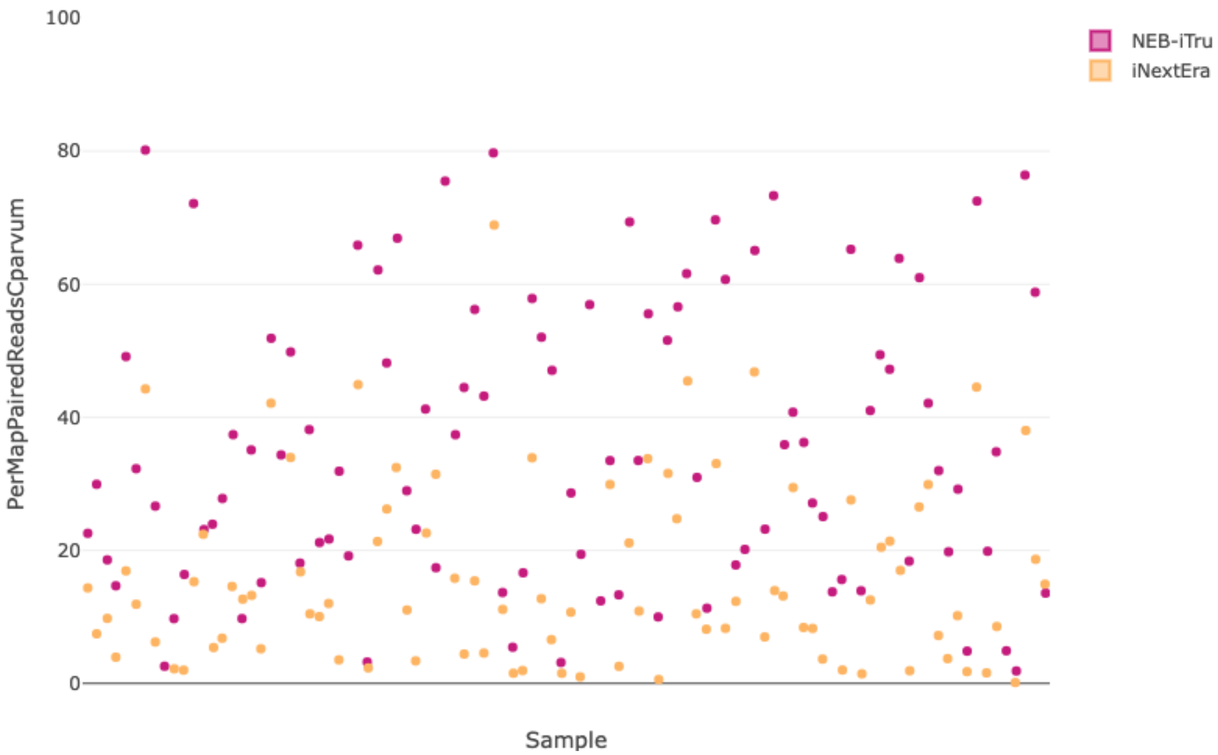


**Suppl. Fig. S44.** Boxplot of the percentage of reads mapped per sample to *C. parvum* obtained for single-enriched libraries from the 100-clinical samples set that were prepared using either the NEB-iTru (pink) protocol or the iNextEra (orange) protocol.

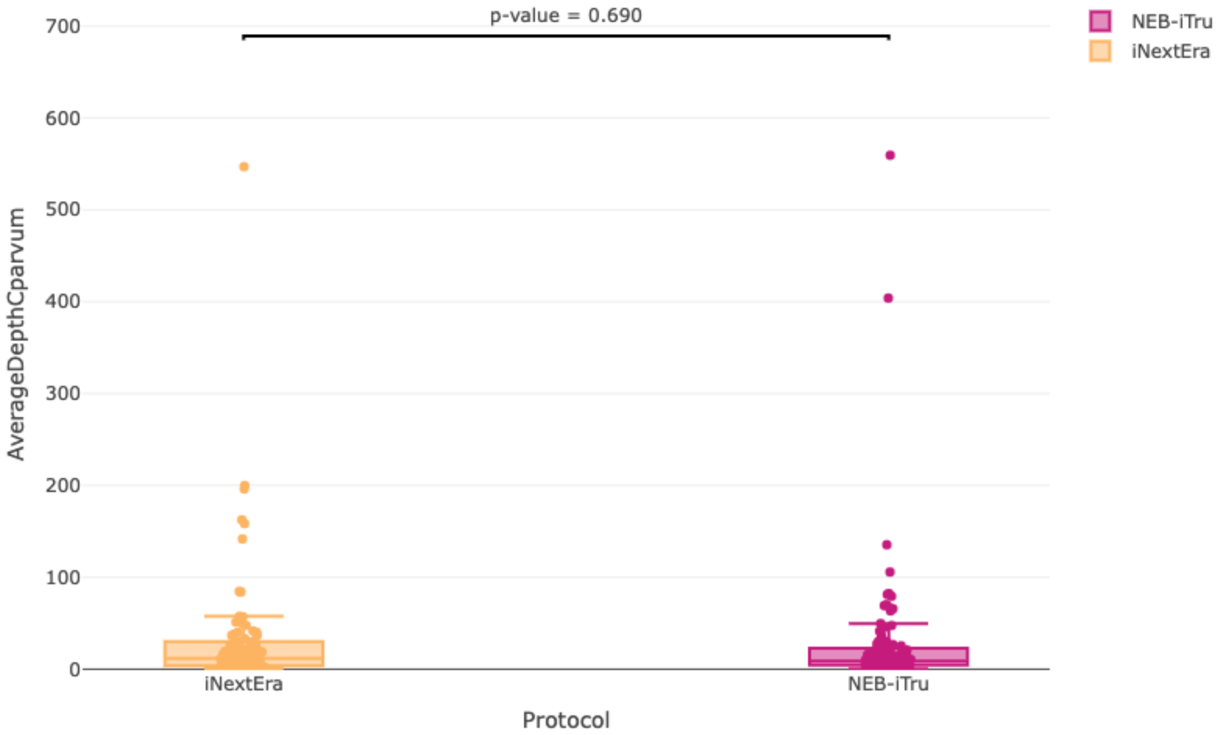


**Suppl. Fig. S45.** Boxplot of the depth of *C. parvum* genome coverage obtained for single-enriched libraries in the 100-clinical samples set that were prepared using either the NEB-iTru (pink) protocol or the iNextEra (orange) protocol.

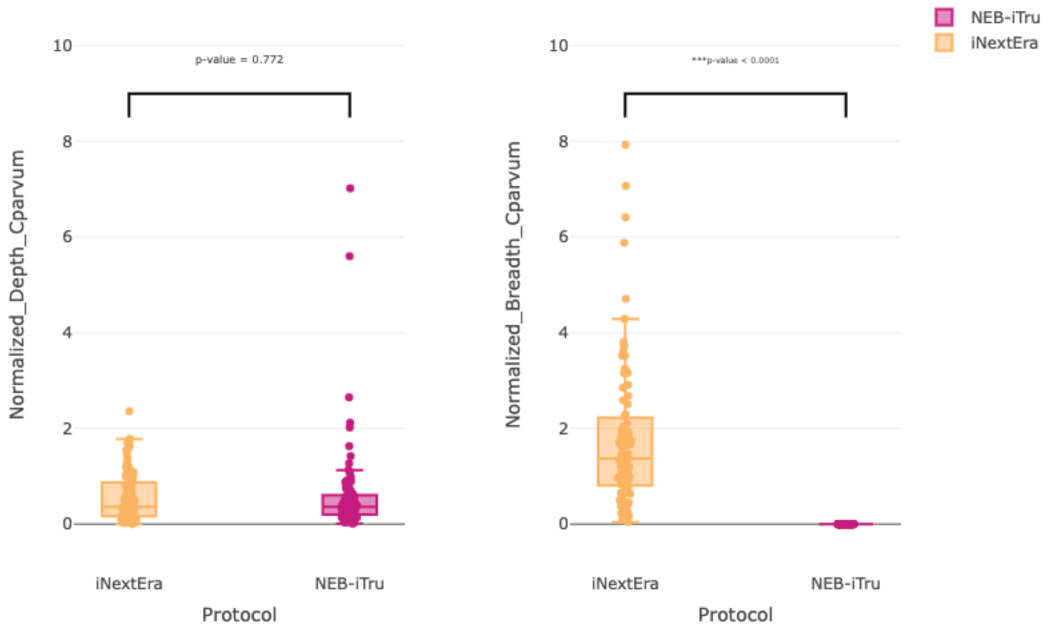


**Suppl. Fig. S46.** Plot of the depth and breadth of genome coverage normalized for 100,000 reads obtained by the mapping of single-enriched sequence reads to *C. parvum* from libraries prepared from the 100 clinical samples set that were prepared using either the NEB-iTru (pink) protocol or the iNextEra-iNext (orange) protocol.

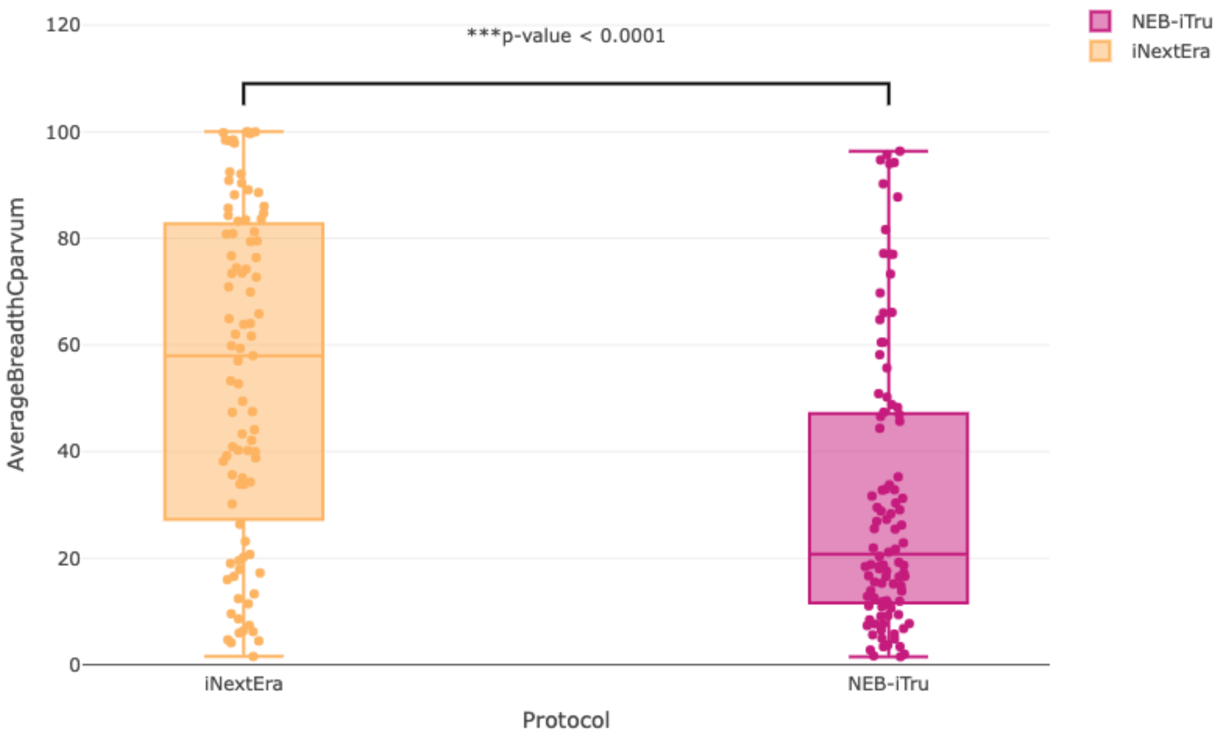


**Suppl. Fig. S47.** Boxplot of the breadth of genome coverage obtained by the mapping of enriched sequence reads to *C. parvum* from libraries prepared from the 100 clinical samples set that were prepared using either the NEB-iTru (pink) protocol or the iNextEra-iNext (orange) protocol.

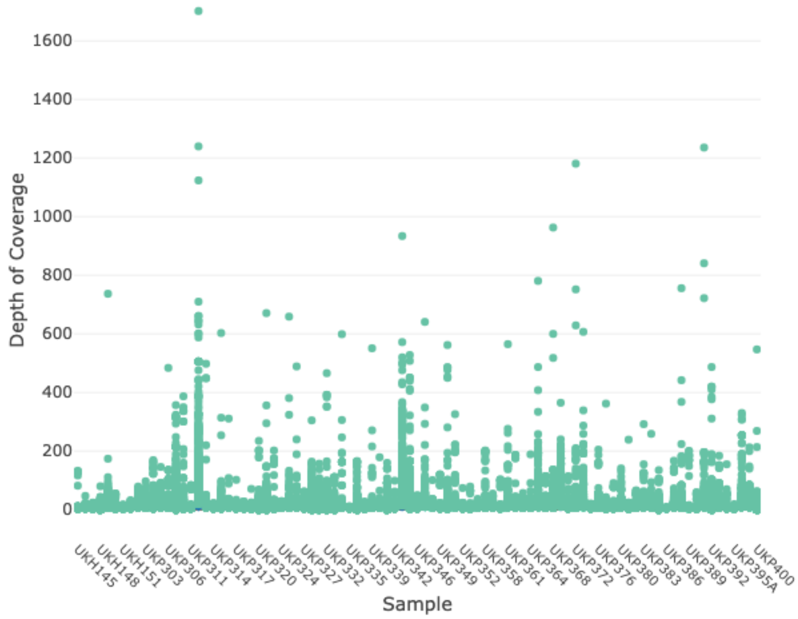


**Suppl. Fig. S48.** Plot of the mean depth of coverage per sample among 94,422 called variants obtained for double-enriched libraries from the clinical samples set that were prepared using the iNextEra protocol.

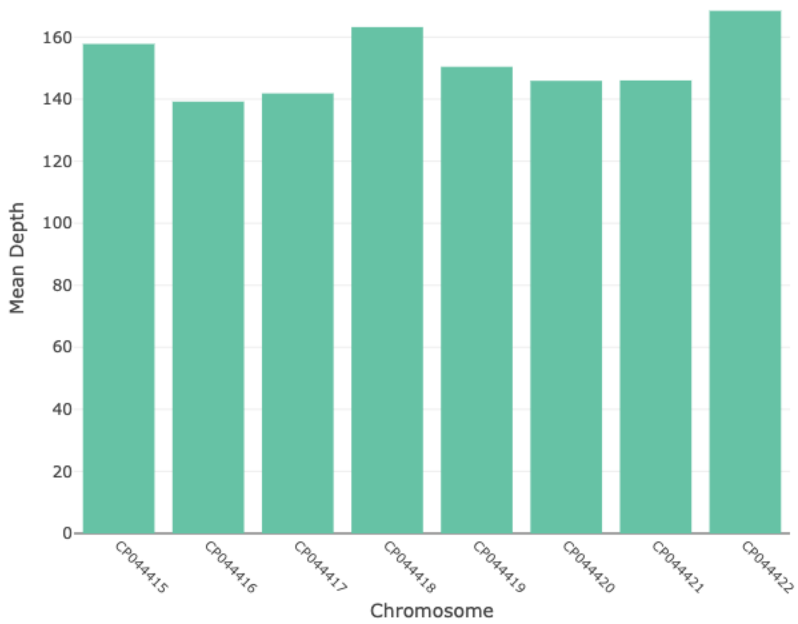


**Suppl. Fig. S49.** Barplot of the mean depth of coverage per *C. parvum* chromosome among 94,422 called variants obtained for double-enriched libraries from the clinical samples set that were prepared used the iNextEra protocol.

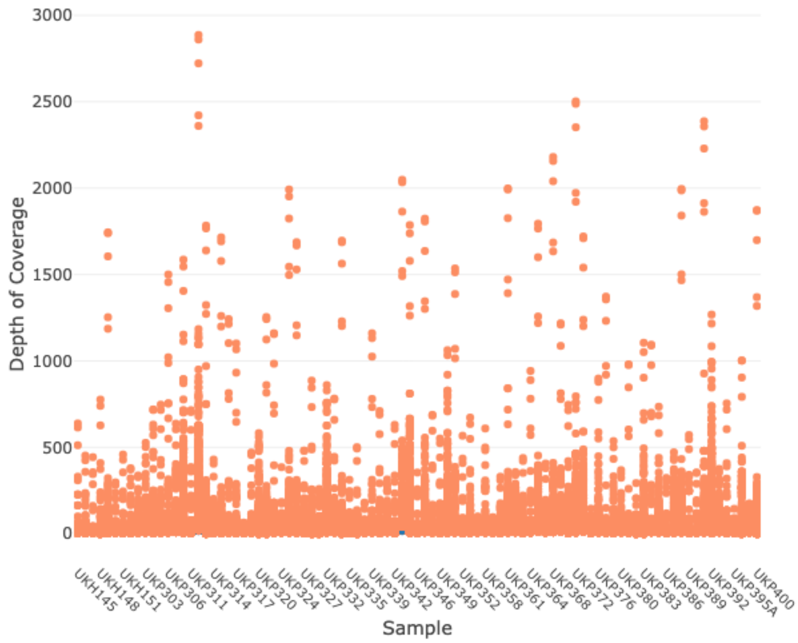


**Suppl. Fig. S50.** Plot of the mean depth of coverage per sample among 126,598 called variants obtained for single-enriched libraries from the clinical samples set that were prepared using the iNextEra protocol.

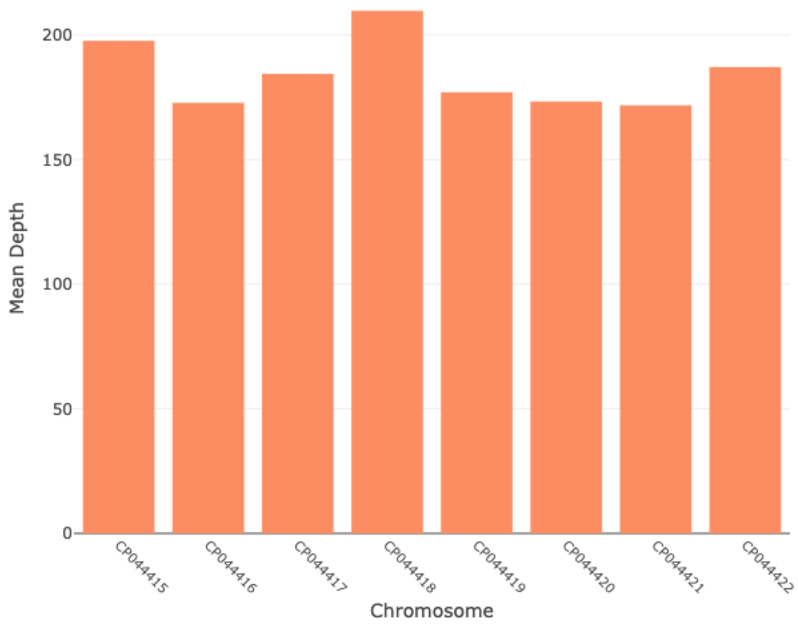


**Suppl. Fig. S51.** Barplot of the mean depth of coverage per *C. parvum* chromosome among 126,598 called variants obtained for single-enriched libraries from the clinical samples set that were prepared using the iNextEra protocol.

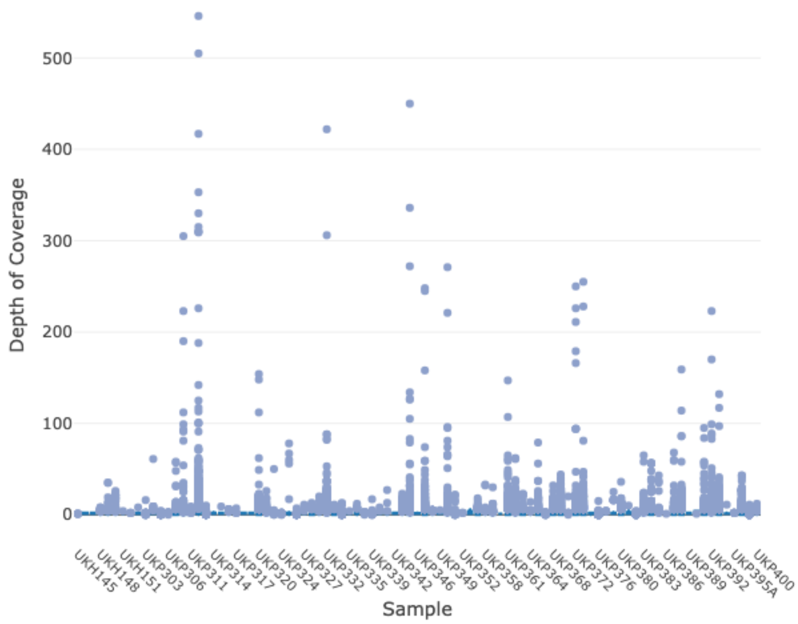


**Suppl. Fig. S52.** Plot of the mean depth of coverage per sample among 13,578 called variants obtained for unenriched libraries from the clinical samples set that were prepared using the iNextEra protocol.

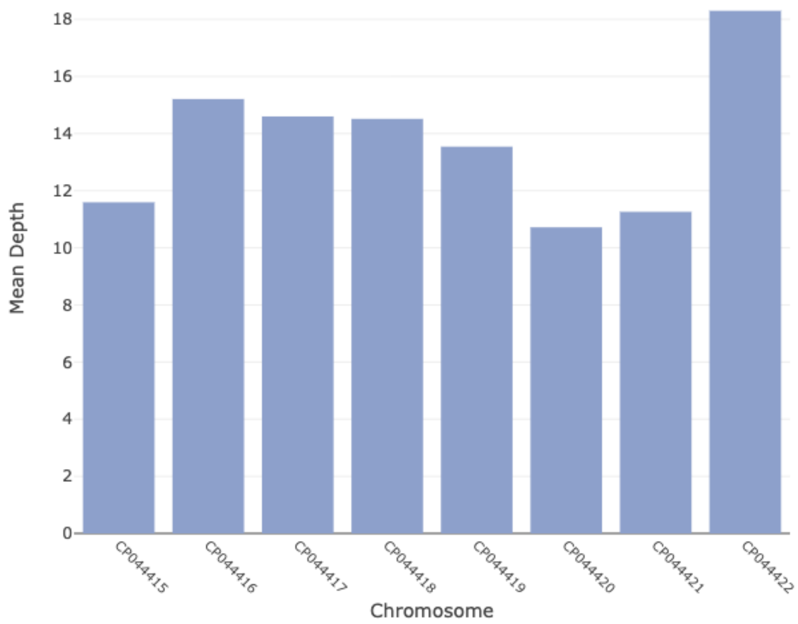


**Suppl. Fig. S53.** Barplot of the mean depth of coverage per *C. parvum* chromosome among 13,578 called variants obtained for single-enriched libraries from the clinical samples set that were prepared using the iNextEra protocol.

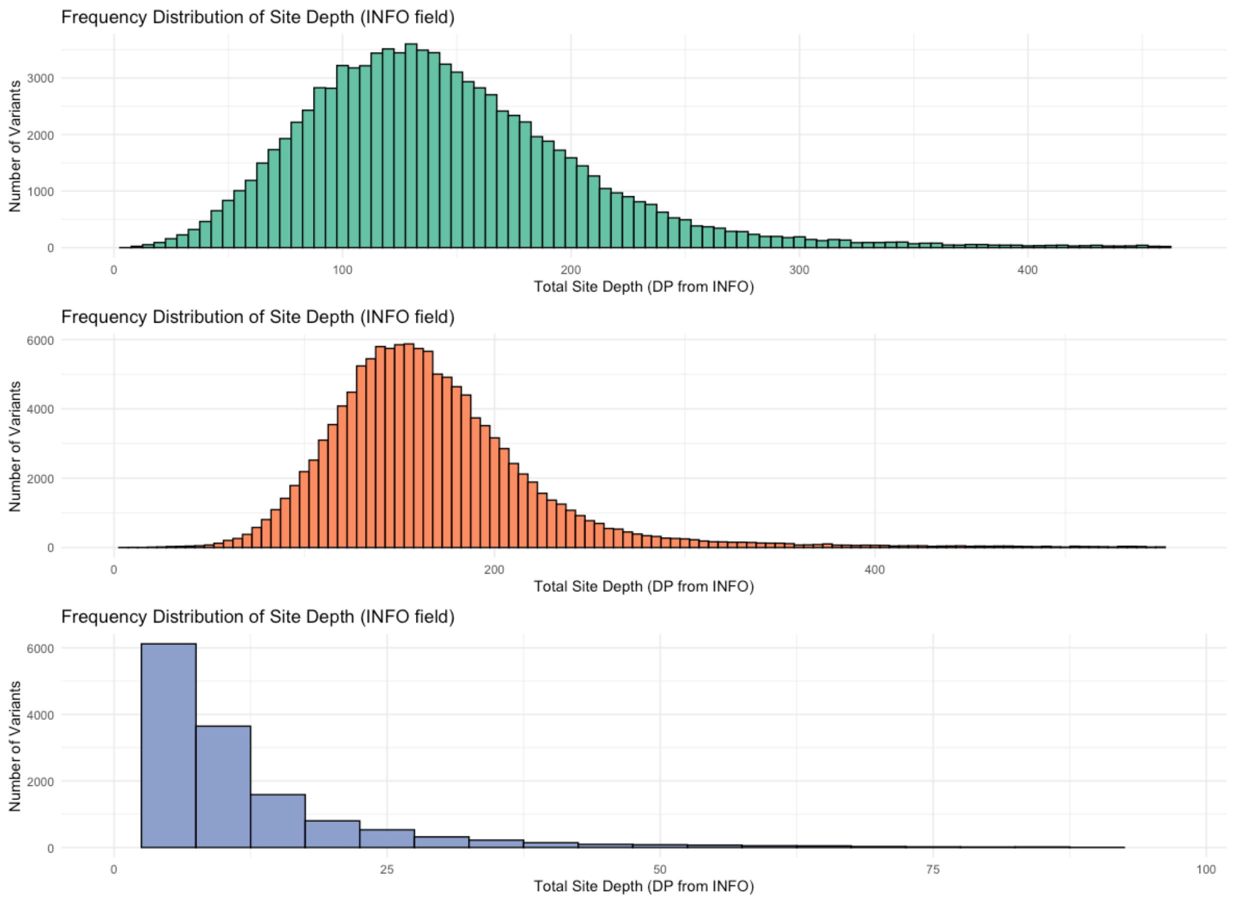


**Suppl. Fig. S54.** Histogram of the distribution of the approximate read depth (DP) from the INFO field across variants for the set of clinical samples from each raw VCF produced by GATK v. 4.6. The Top plot represents double-enriched libraries, containing 94,422 SNPs; the middle plot represents single-enriched libraries, containing 126,598 SNPs; and the bottom plot represents unenriched libraries, containing 15,198 SNPs.

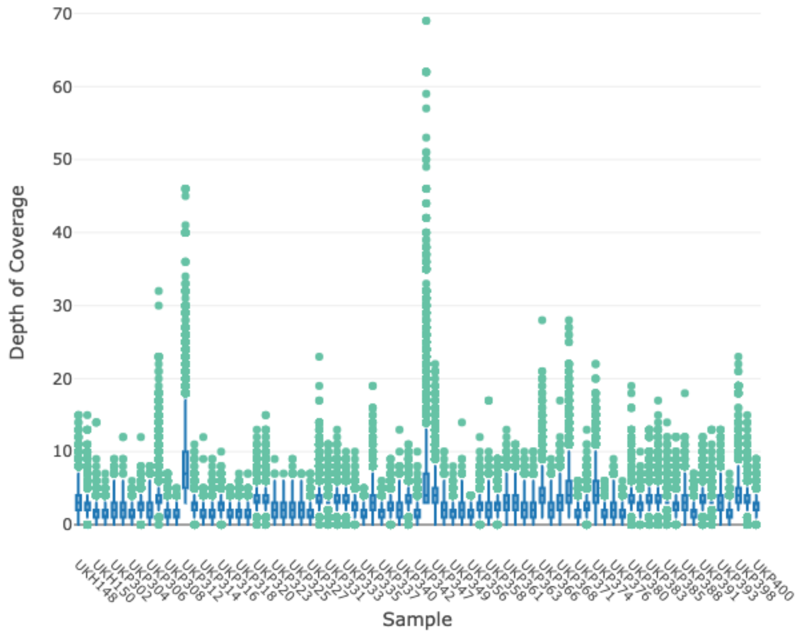


**Suppl. Fig. S55.** Plot of the mean depth of coverage per sample among 13,578 filtered variants obtained for double-enriched libraries from the clinical samples set that were prepared using the iNextEra protocol.

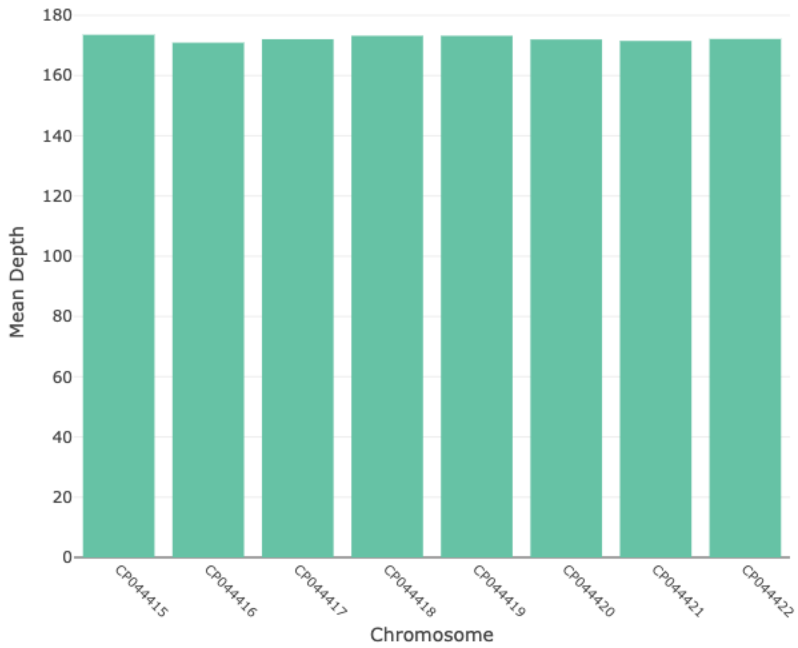


**Suppl. Fig. S56.** Barplot of the mean depth of coverage per *C. parvum* chromosome for 13,578 filtered variants that passed filters obtained for double-enriched libraries from the clinical samples set that were prepared using the iNextEra protocol.

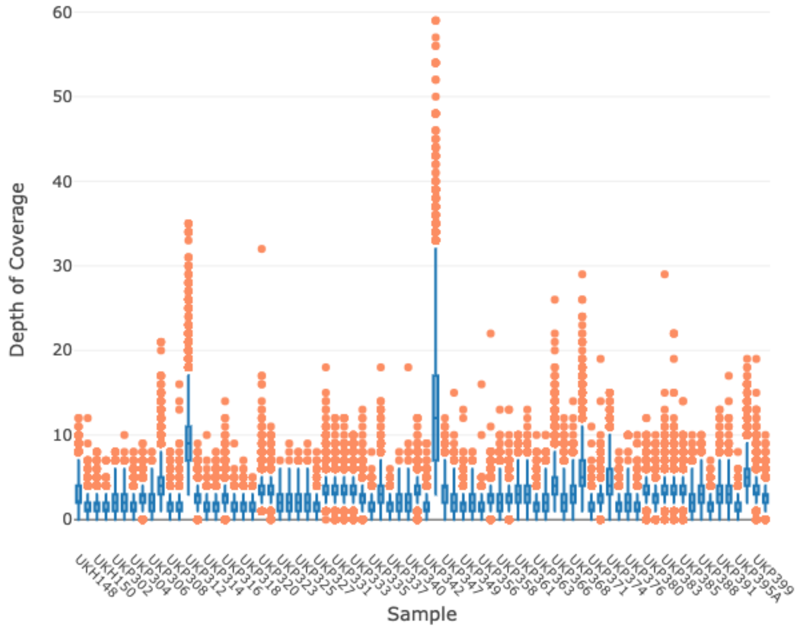


**Suppl. Fig. S57.** Barplot of the mean depth of coverage per sample among 12,498 filtered variants obtained for single-enriched libraries from the clinical samples set that were prepared using the iNextEra protocol.

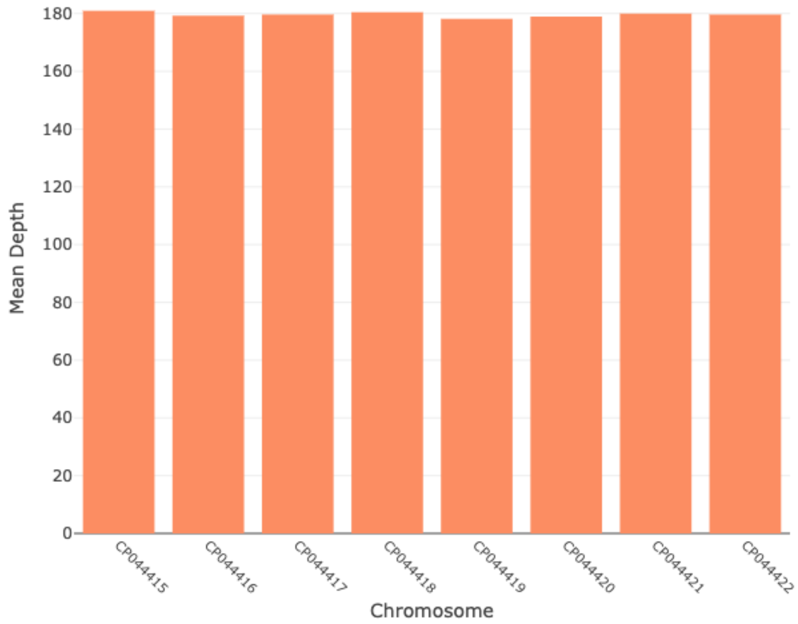


**Suppl. Fig. S58.** Barplot of the mean depth of coverage per *C. parvum* chromosome for 12,498 filtered variants that passed filters obtained for single-enriched libraries from the clinical samples set that were prepared using the iNextEra protocol.

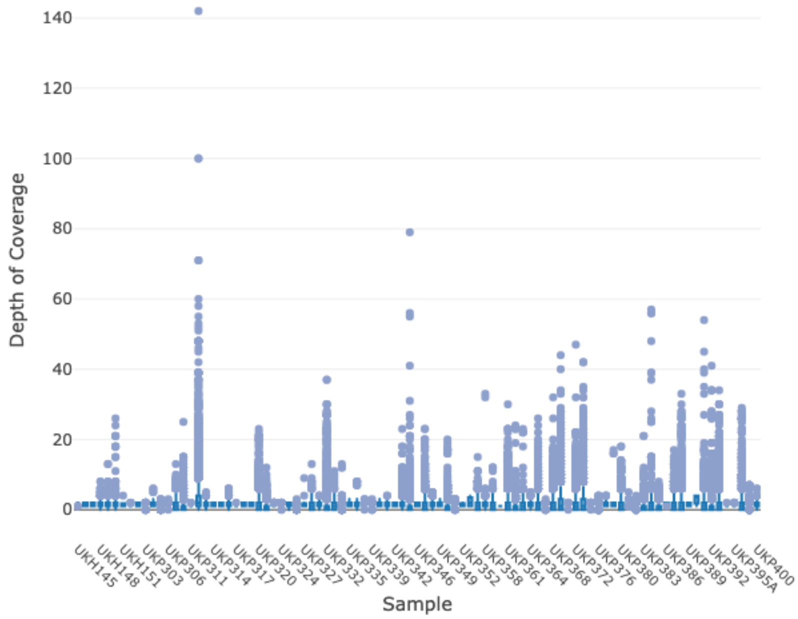


**Suppl. Fig. S59.** Plot of the mean depth of coverage per sample among 11,226 filtered (to some extent) variants obtained for unenriched libraries from the clinical samples set that were prepared using the iNextEra protocol.

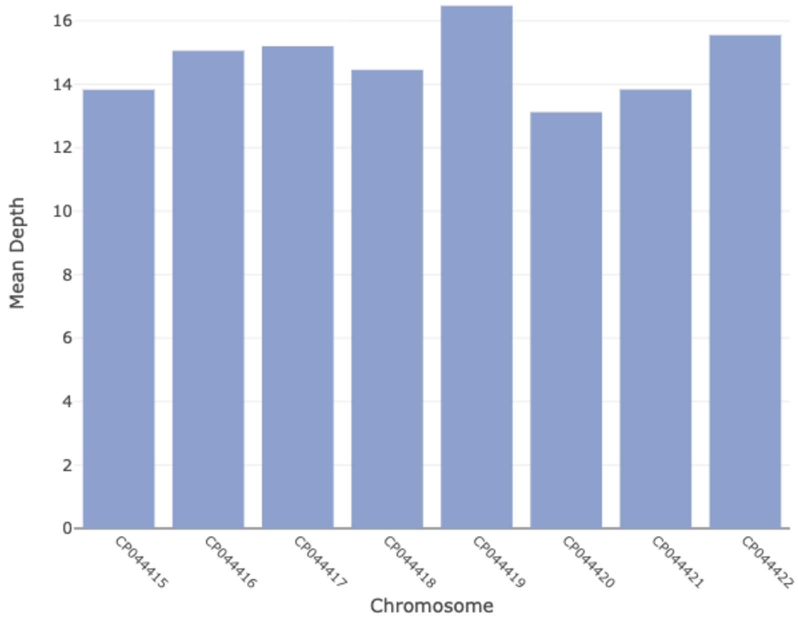


**Suppl. Fig. S60.** Barplot of the mean depth of coverage per *C. parvum* chromosome for 11,226 filtered (to some extent) variants that passed filters obtained for single-enriched libraries from the clinical samples set that were prepared using the iNextEra protocol.

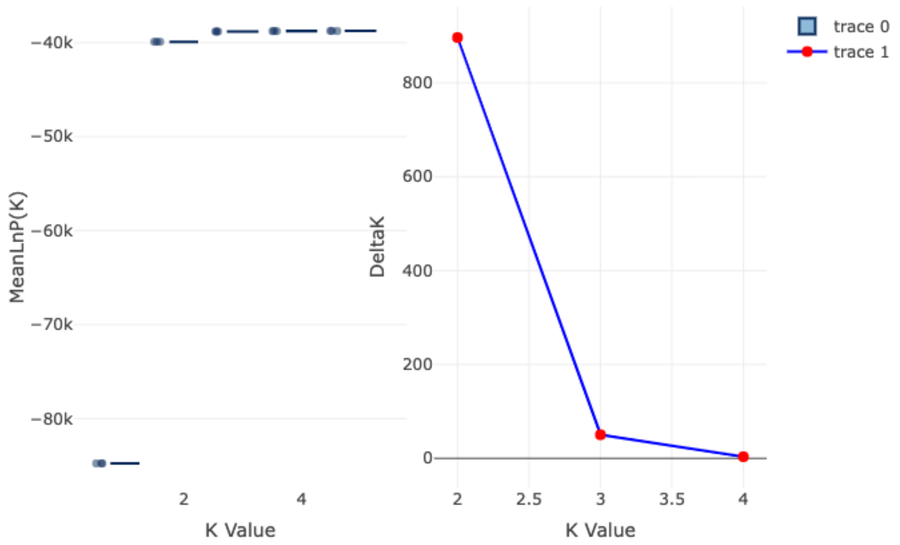


**Suppl. Fig. S61.** Mean LnP for each K value tested (1-5) and Delta K for K 2-4 calculated by the Evanno method using 13,578 SNPs from all the clinical samples set that were prepared using the iNextEra protocol and double-enriched with CryptoCap_100k.

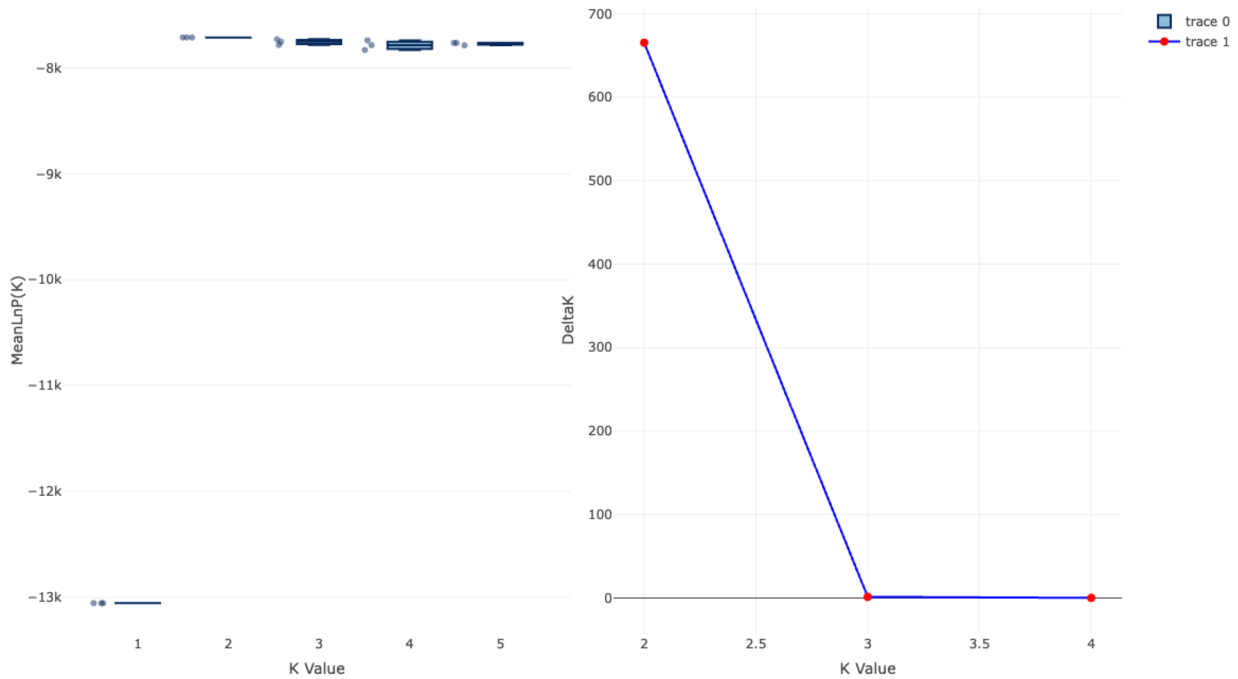


**Suppl. Fig. S62.** Mean LnP for each K value tested (1-5) and Delta K for K 2-4 calculated by the Evanno method using 13,578 SNPs from only clinical samples categorized as UKH (*Cryptosporidium hominis*) set that were prepared using the iNextEra protocol and double-enriched with CryptoCap_100k.

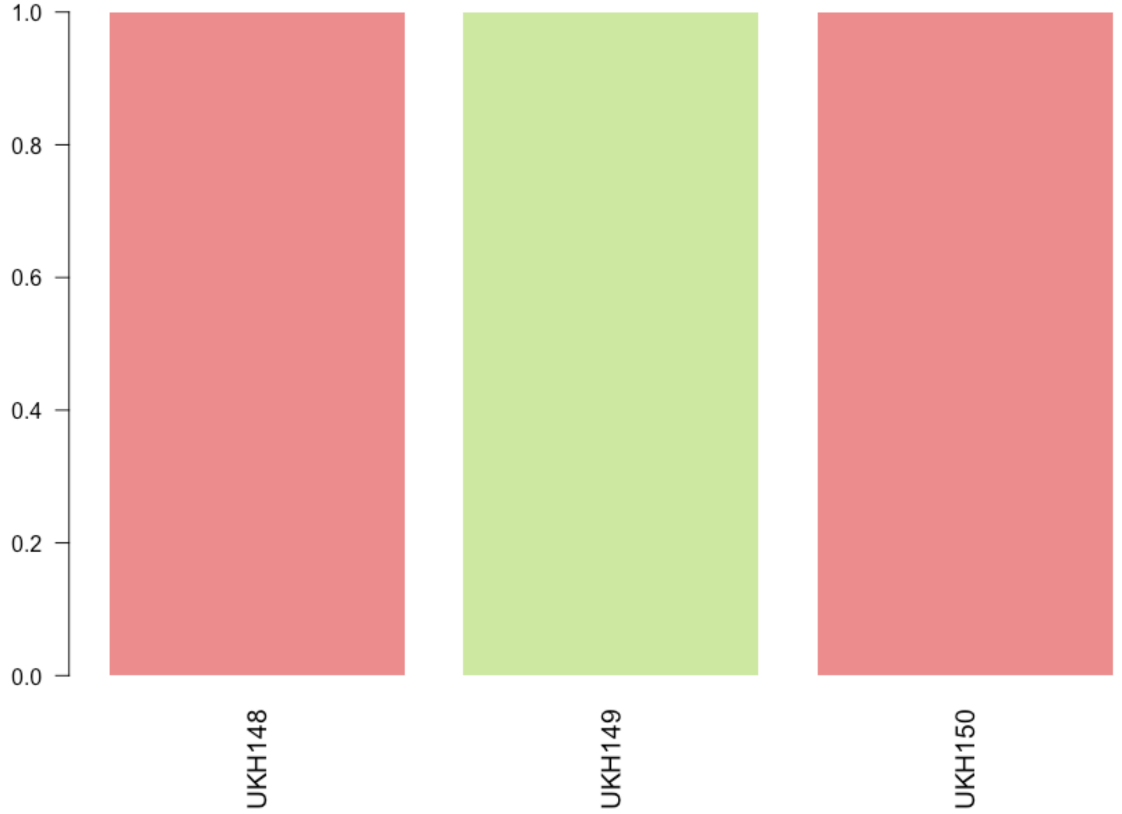


**Suppl. Fig. S63.** Structure Plot for K=2 (best K according to the Evanno method) using 13,578 SNPs from only clinical samples categorized as UKH (*Cryptosporidium hominis*) set that were prepared using the iNextEra protocol and double-enriched with CryptoCap_100k.

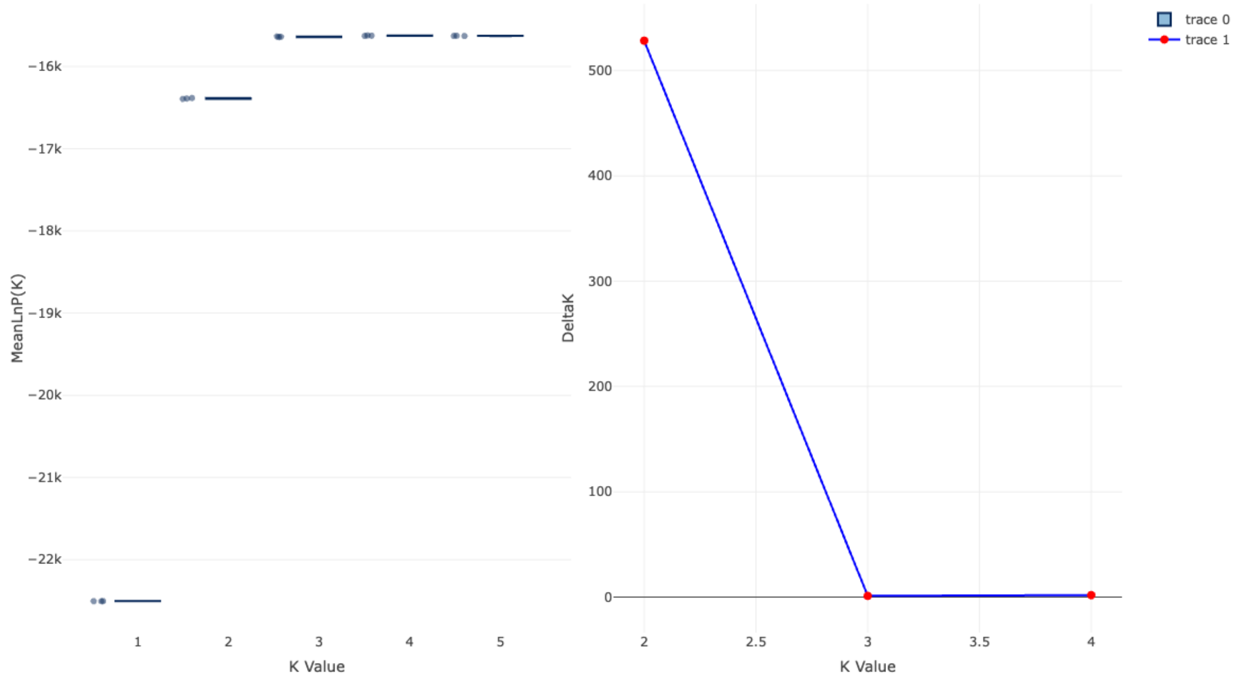


**Suppl. Fig. S64.** Mean LnP for each K value tested (1-5) and Delta K for K 2-4 calculated by the Evanno method using 13,578 SNPs from only clinical samples categorized as UKP (*Cryptosporidium parvum*) set that were prepared using the iNextEra protocol and double-enriched with CryptoCap_100k.

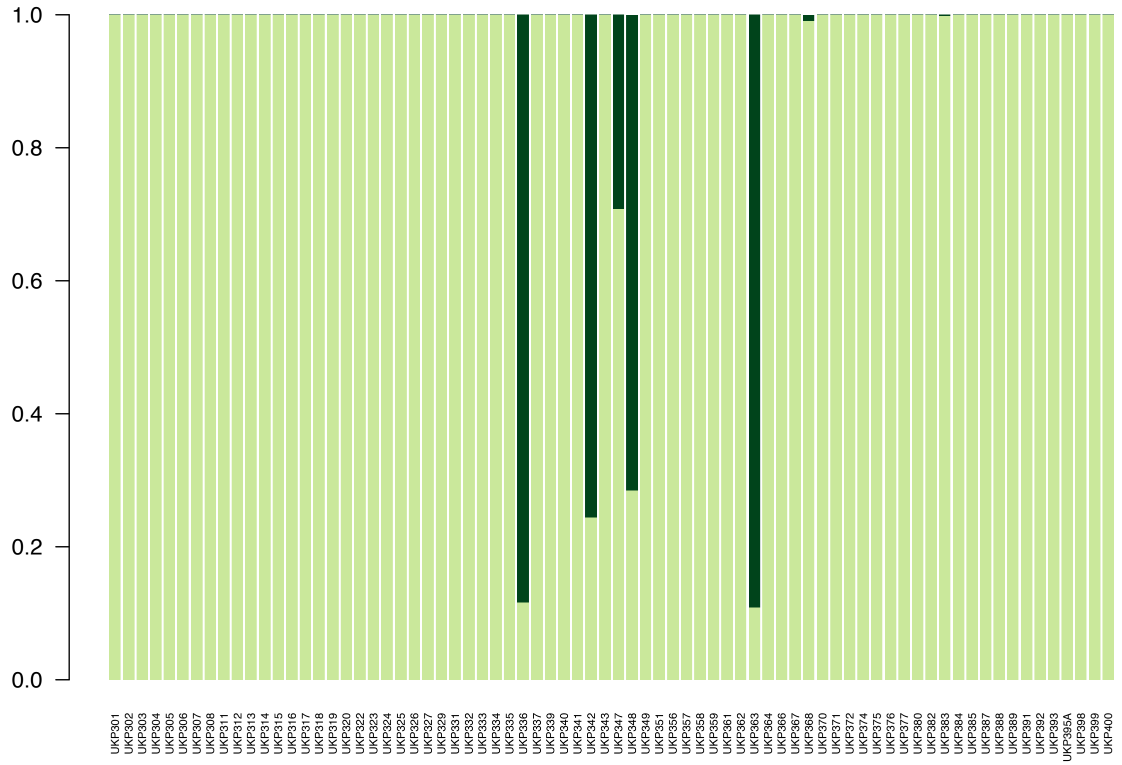


**Suppl. Fig. S65.** Structure Plot for K=2 (best K according to the Evanno method) using 13,578 SNPs from only clinical samples categorized as UKP (*Cryptosporidium parvum*) set that were prepared using the iNextEra protocol and double-enriched with CryptoCap_100k.

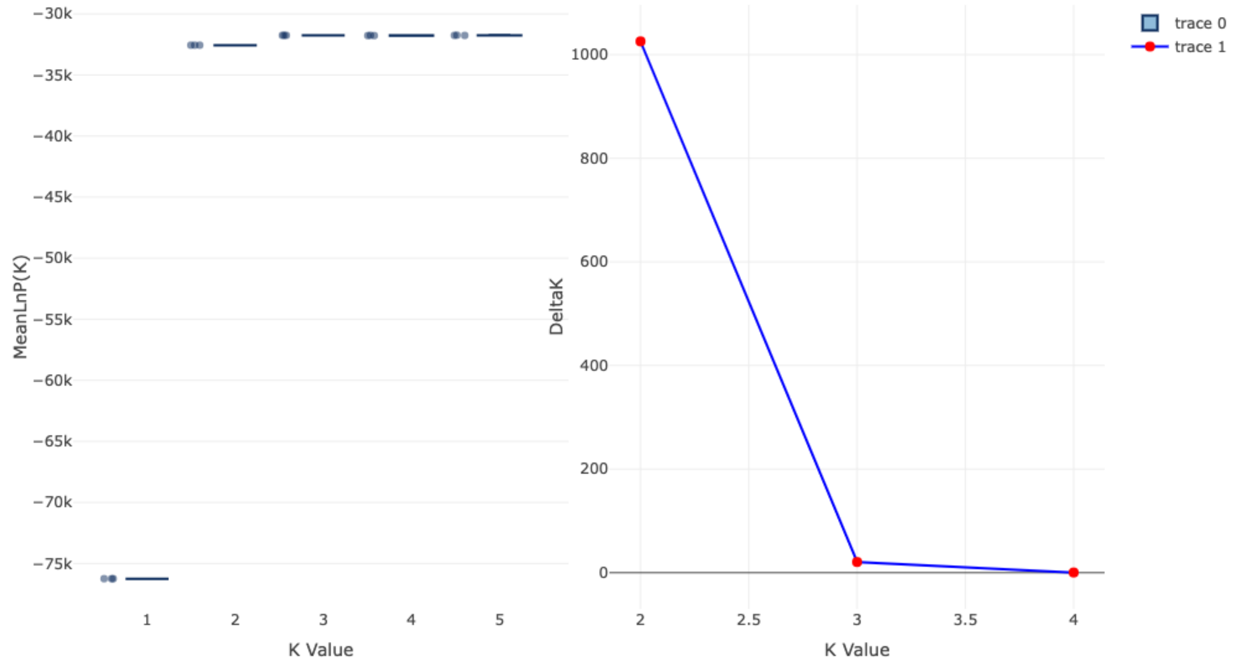


**Suppl. Fig. S66.** Mean LnP for each K value tested (1-5) and Delta K for K 2-4 calculated by the Evanno method using 12,498 SNPs from the clinical samples set that were prepared using the iNextEra protocol and single-enriched with CryptoCap_100k.

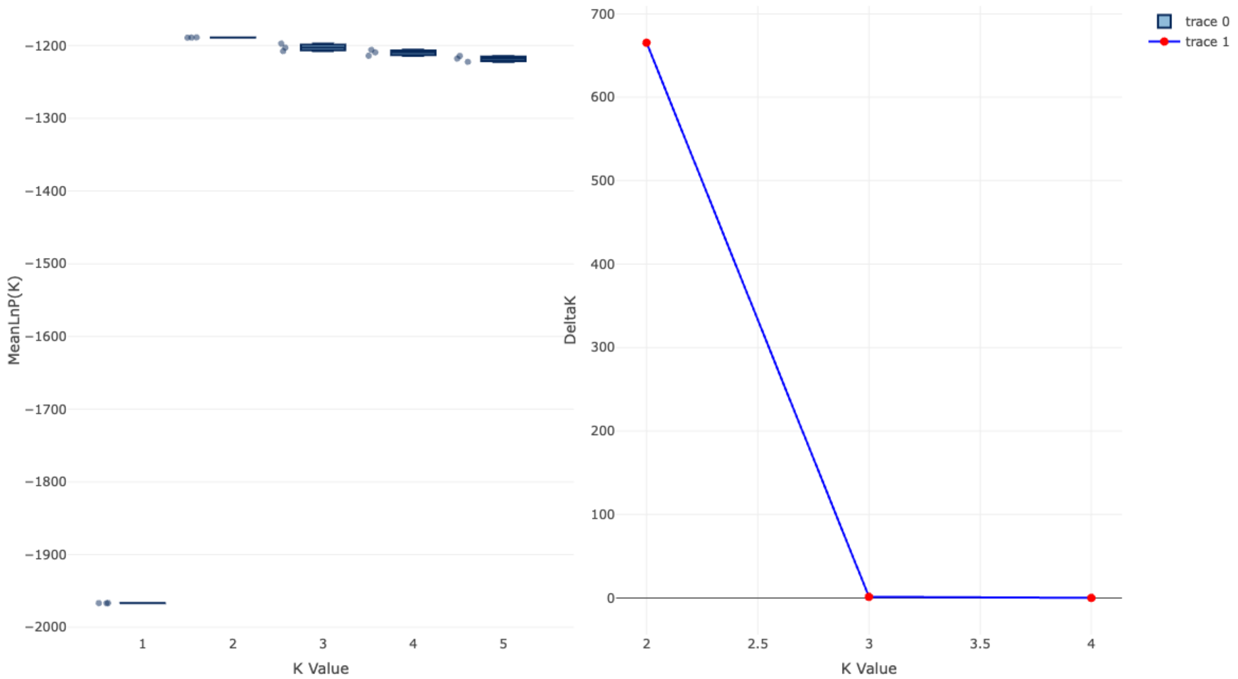


**Suppl. Fig. S67.** Mean LnP for each K value tested (1-5) and Delta K for K 2-4 calculated by the Evanno method using 12,498 SNPs from only clinical samples categorized as UKH (*Cryptosporidium hominis*) set that were prepared using the iNextEra protocol and single-enriched with CryptoCap_100k.

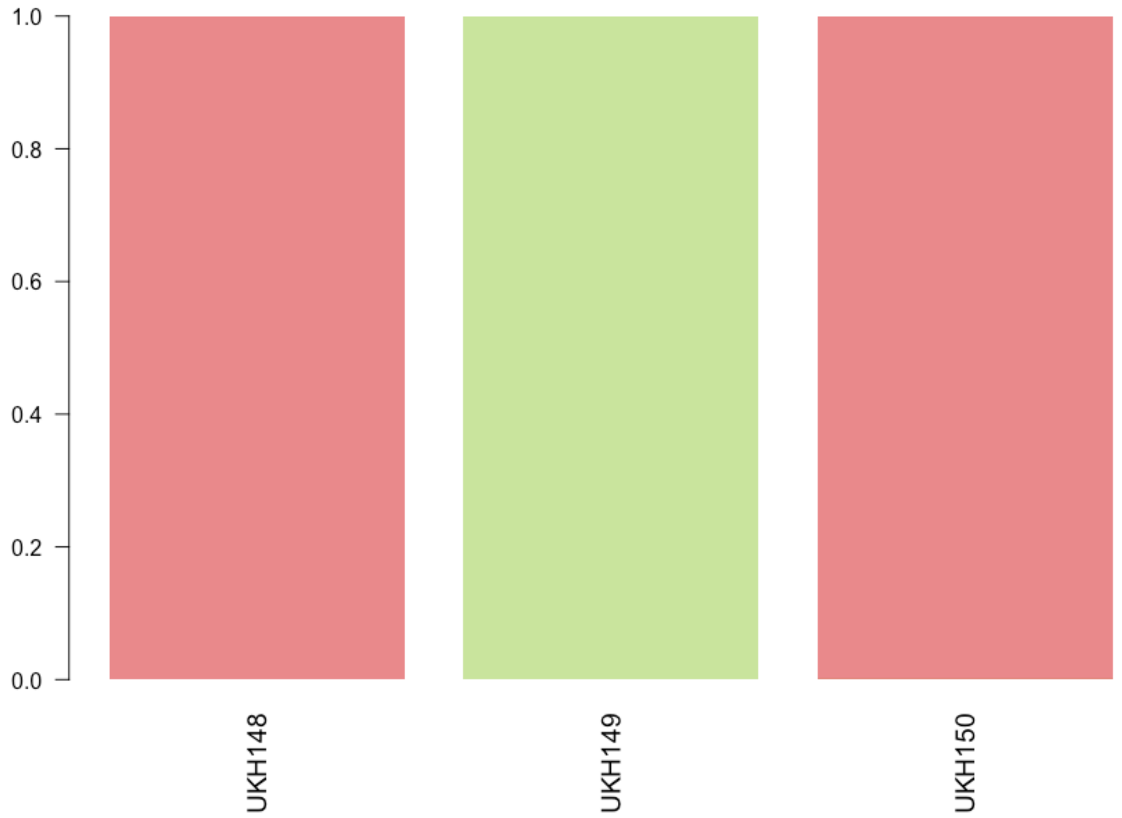


**Suppl. Fig. S68.** Structure Plot for K=2 (best K according to the Evanno method) using 12,498 SNPs from only clinical samples categorized as UKH (*Cryptosporidium hominis*) set that were prepared using the iNextEra protocol and single-enriched with CryptoCap_100k.

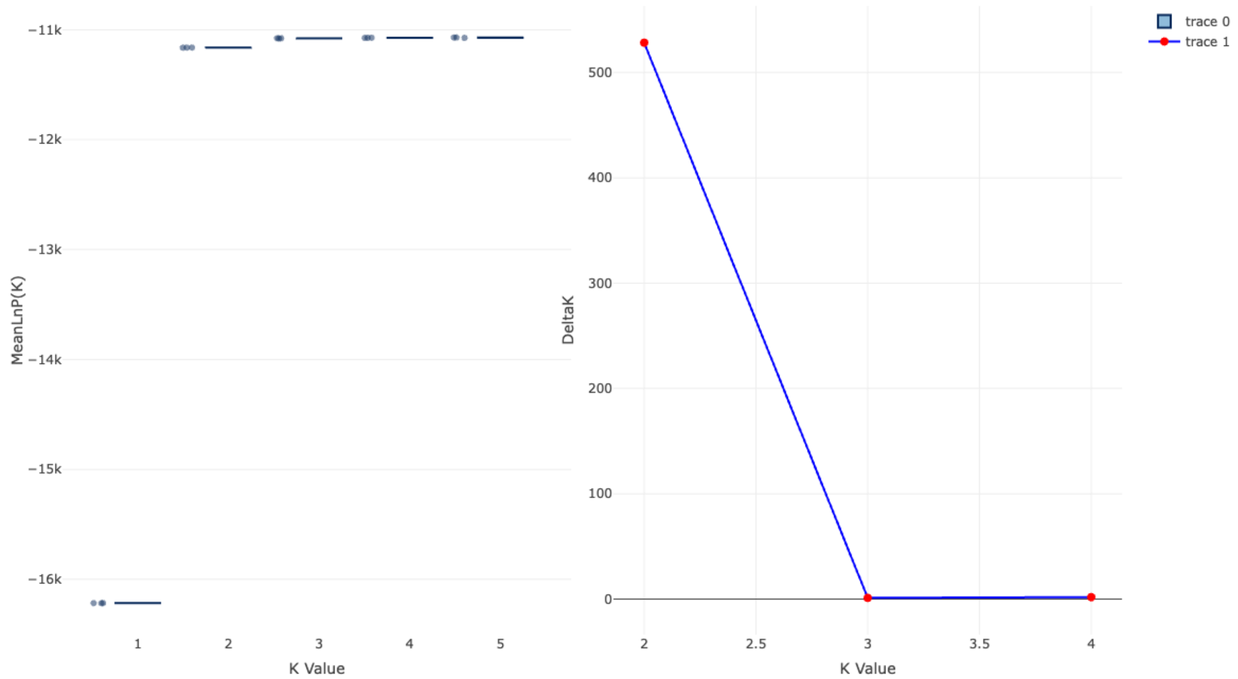


**Suppl. Fig. S69.** Mean LnP for each K value tested (1-5) and Delta K for K 2-4 calculated by the Evanno method using 12,498 SNPs from only clinical samples categorized as UKP (*Cryptosporidium parvum*) set that were prepared using the iNextEra protocol and single-enriched with CryptoCap_100k.

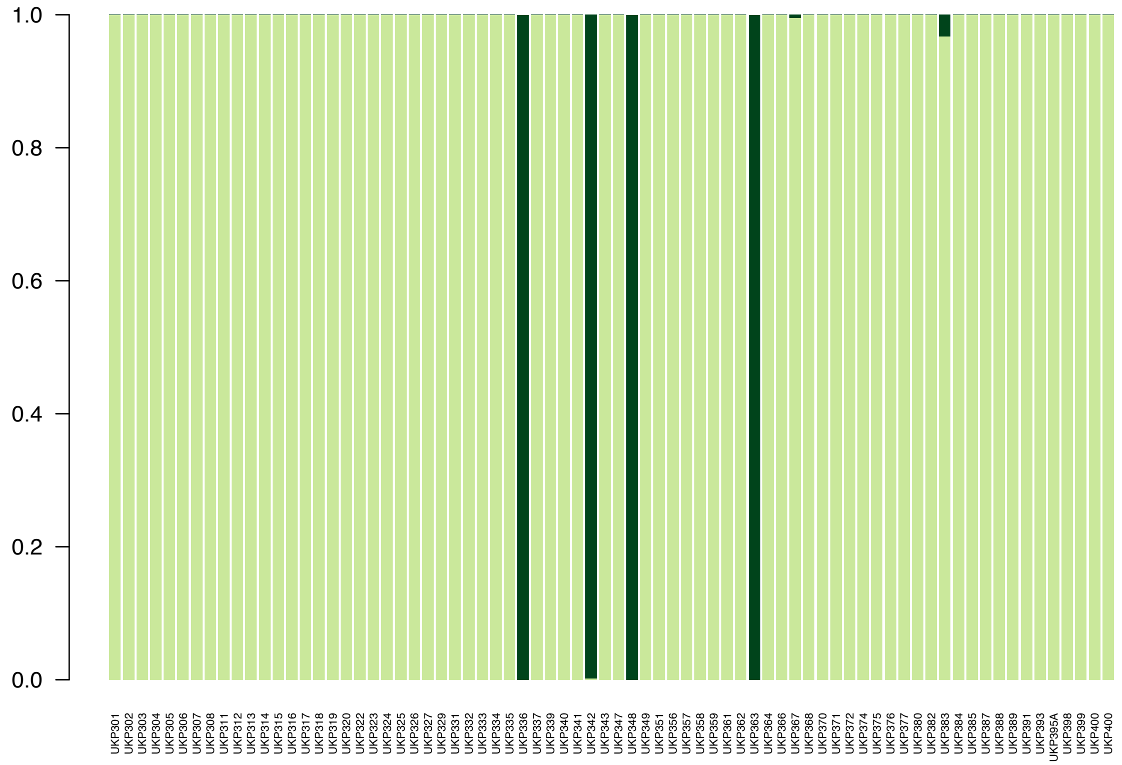

**Suppl. Fig. S70.** Structure Plot for K=2 (best K according to the Evanno method) using 12,498 SNPs from only clinical samples categorized as UKP (*Cryptosporidium parvum*) set that were prepared using the iNextEra protocol and single-enriched with CryptoCap_100k.

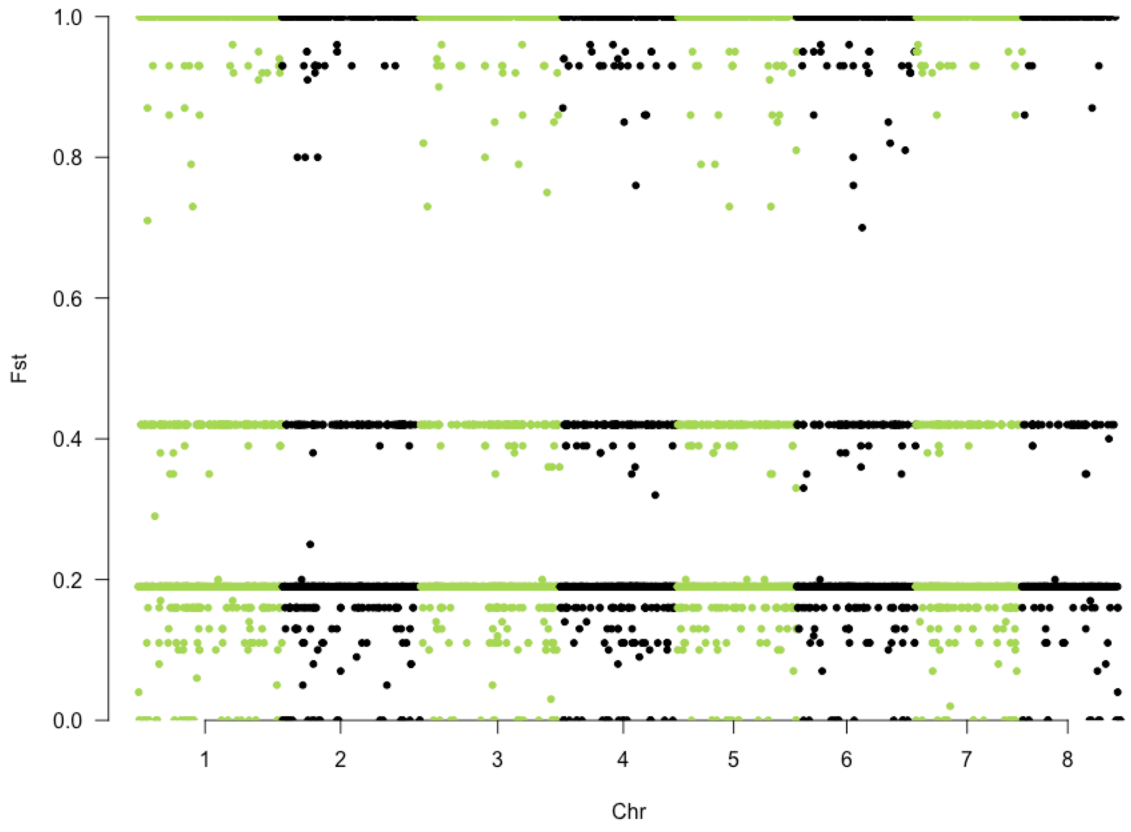


**Suppl. Fig. S71.** Manhattan plot of Fst values across *C. parvum* IOWA-ATCC reference genome chromosomes between UKH and UKP samples for 13,578 variants from the clinical samples set that were prepared using the iNextEra protocol and double-enriched with CryptoCap_100k.

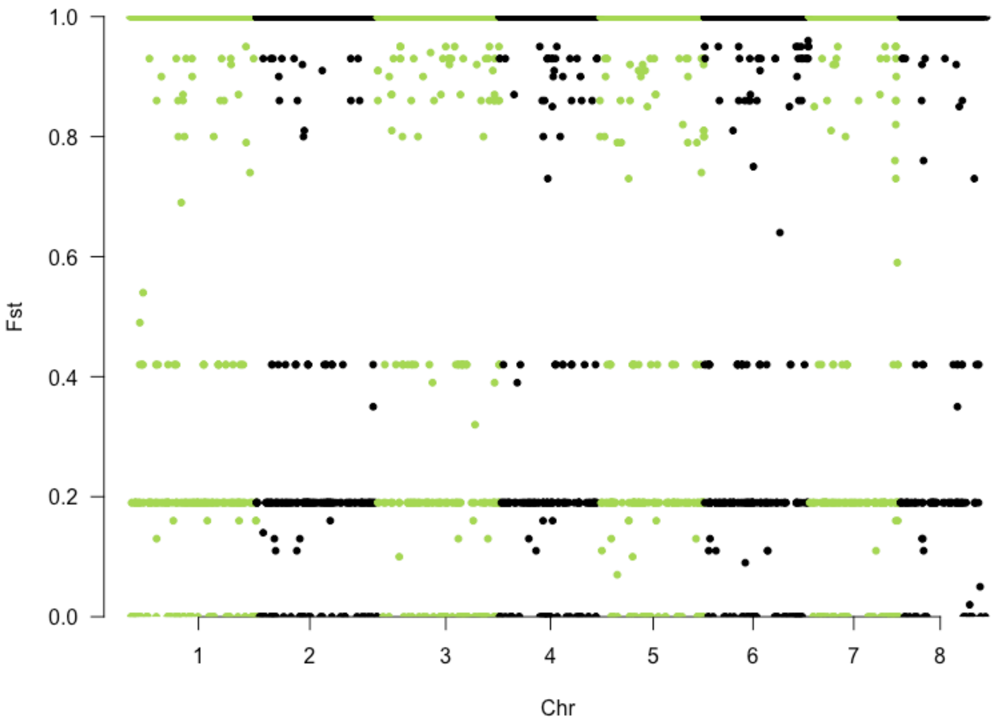


**Suppl. Fig. S72.** Manhattan plot of Fst values across *C. parvum* IOWA-ATCC reference genome chromosomes between UKH and UKP samples for 12,498 variants from the clinical samples set that were prepared using the iNextEra protocol and single-enriched with CryptoCap_100k.

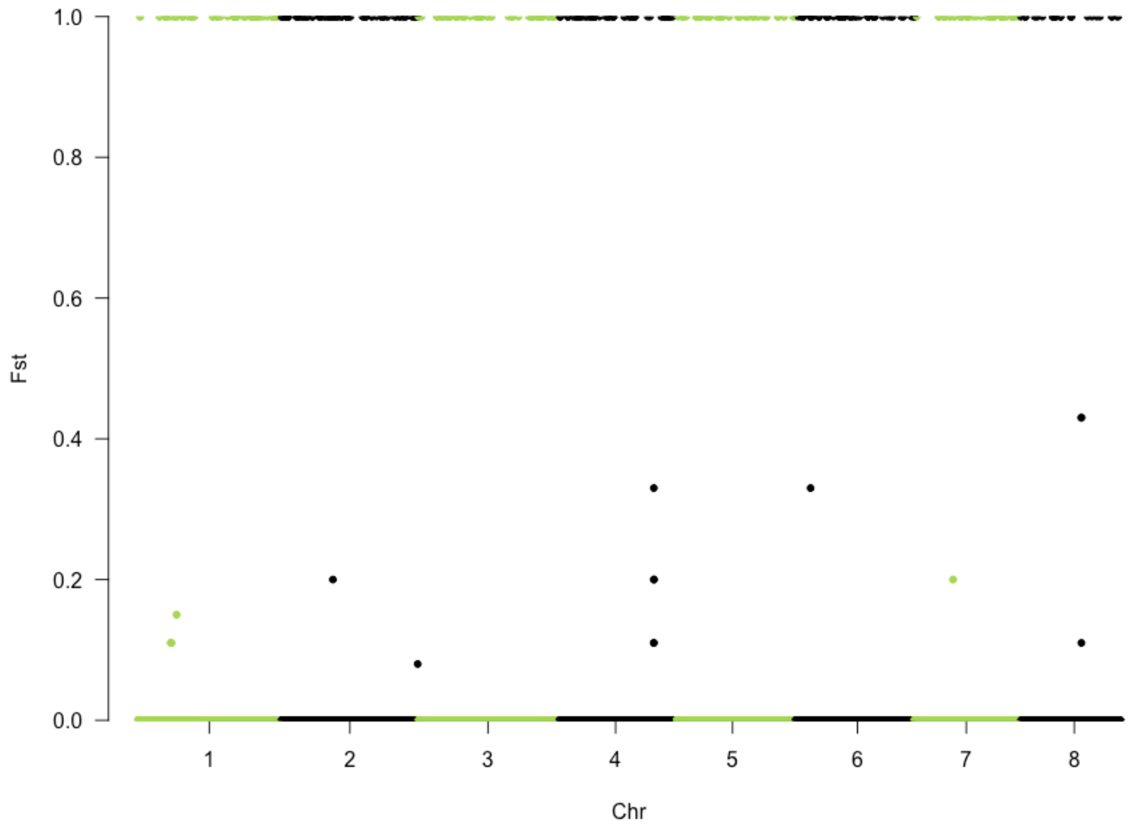


**Suppl. Fig. S73.** Manhattan plot of Fst values across *C. parvum* IOWA-ATCC reference genome chromosomes between UKH and UKP samples for 11,226 variants from the clinical samples set that were prepared using the iNextEra protocol and left unenriched.

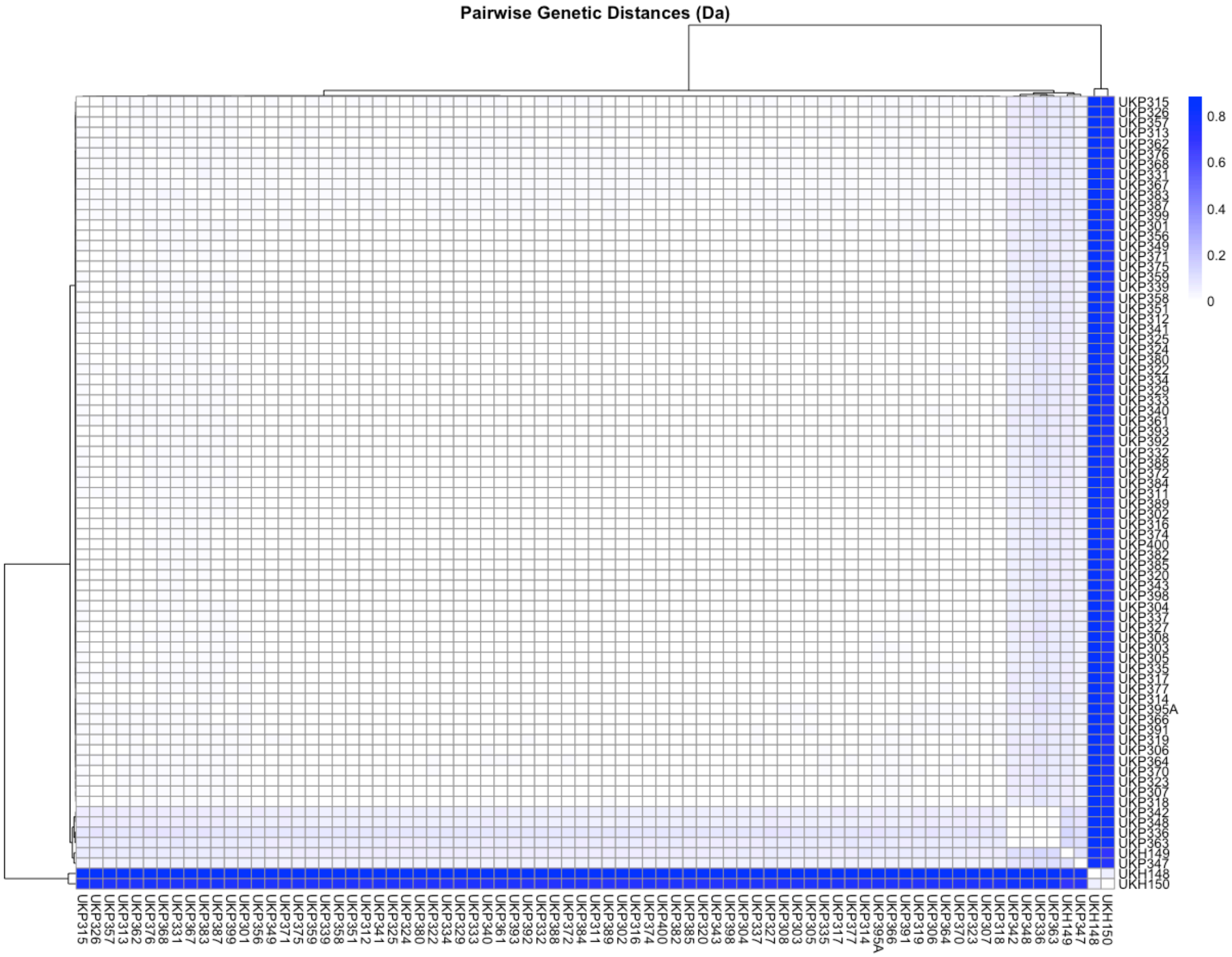


**Suppl. Fig. S74.** Heatmap of pairwise genetic distance calculated using 13,578 SNPs from the clinical samples set that were prepared using the iNextEra protocol and double-enriched with CryptoCap_100k.

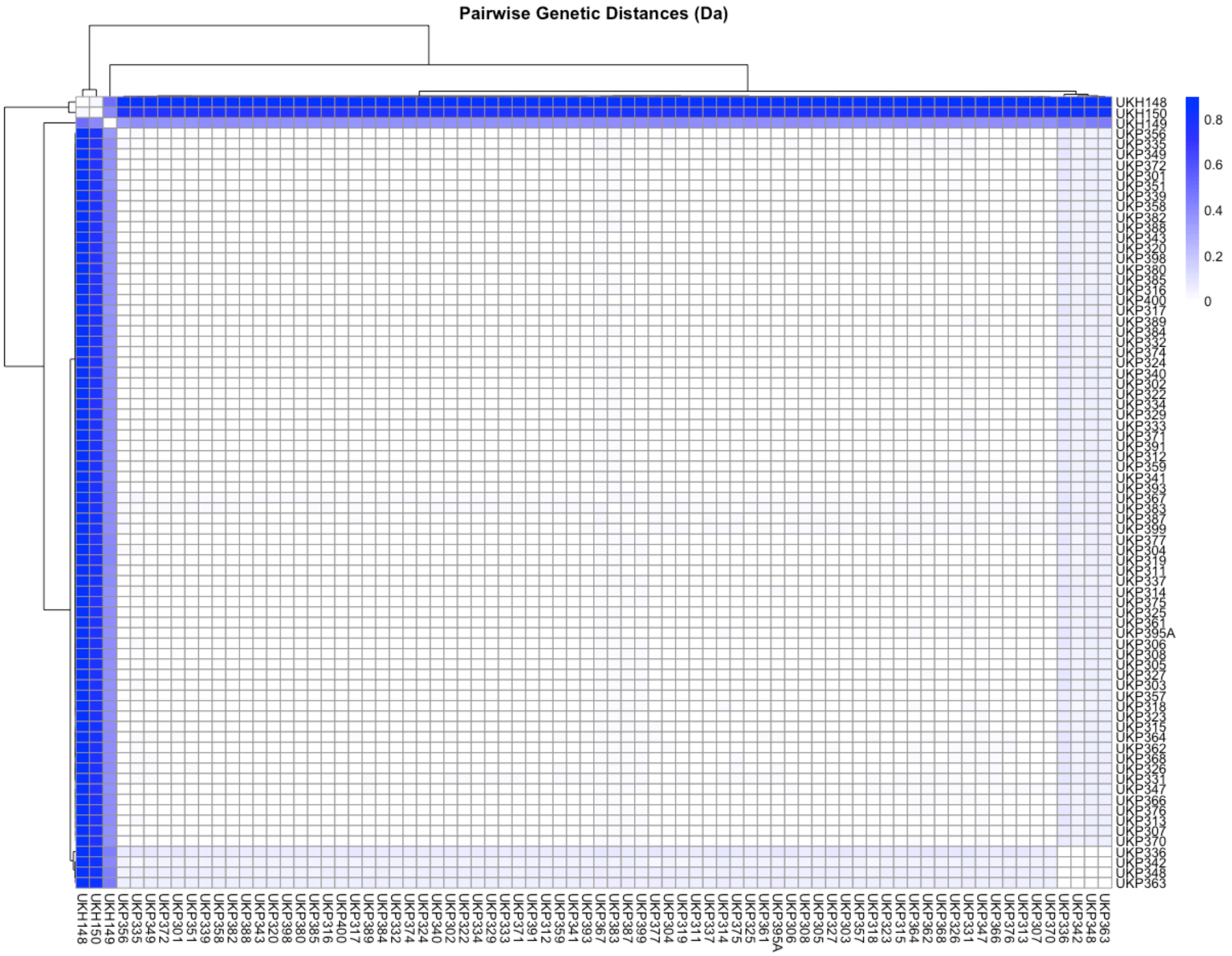


**Suppl. Fig. S75.** Heatmap of pairwise genetic distance calculated using 12,498 SNPs from the clinical samples set that were prepared using the iNextEra protocol and single-enriched with CryptoCap_100k.

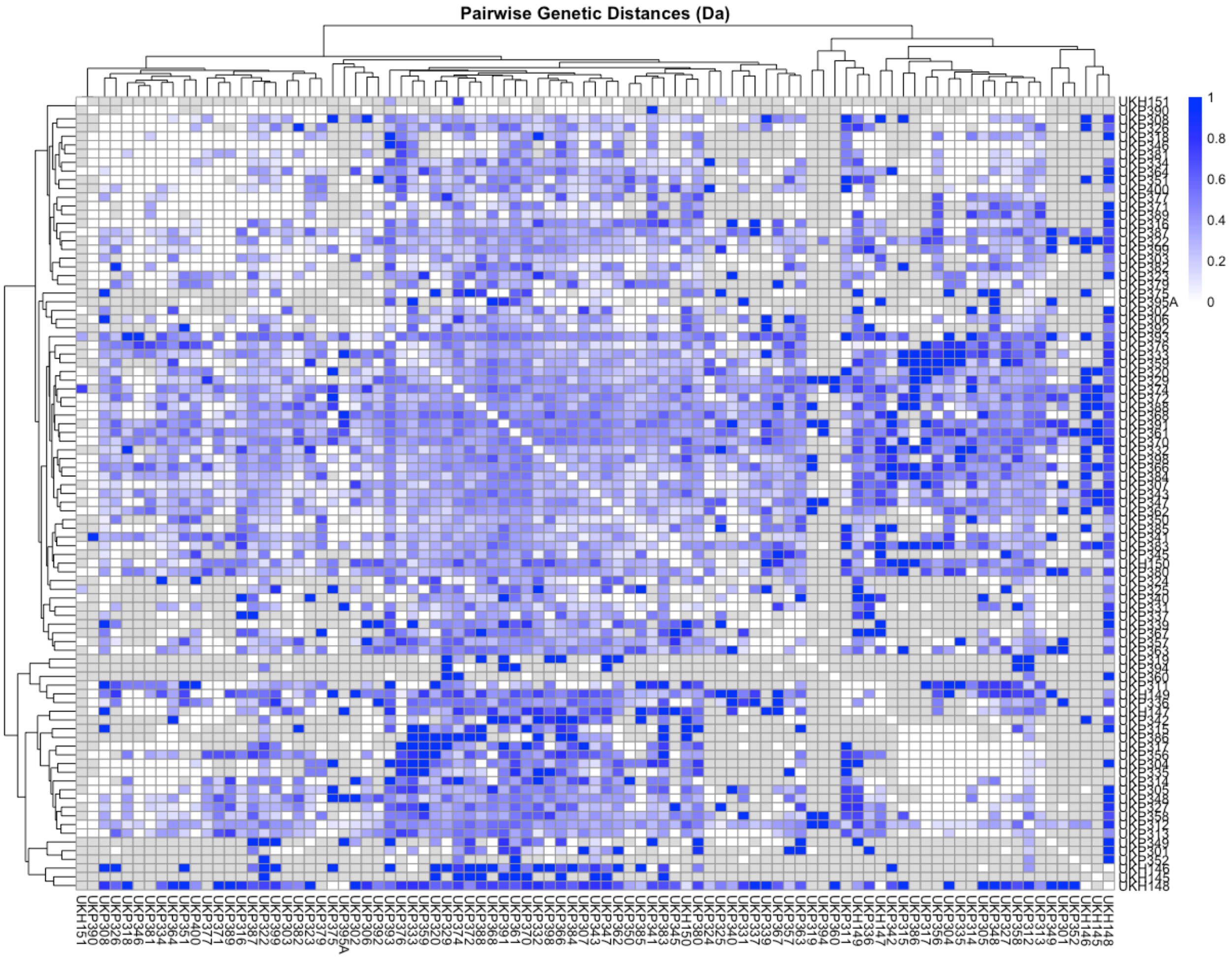


**Suppl. Fig. S76.** Heatmap of your pairwise genetic distance calculated using 13,044 SNPs from the clinical samples set that were prepared using the iNextEra protocol and left unenriched.

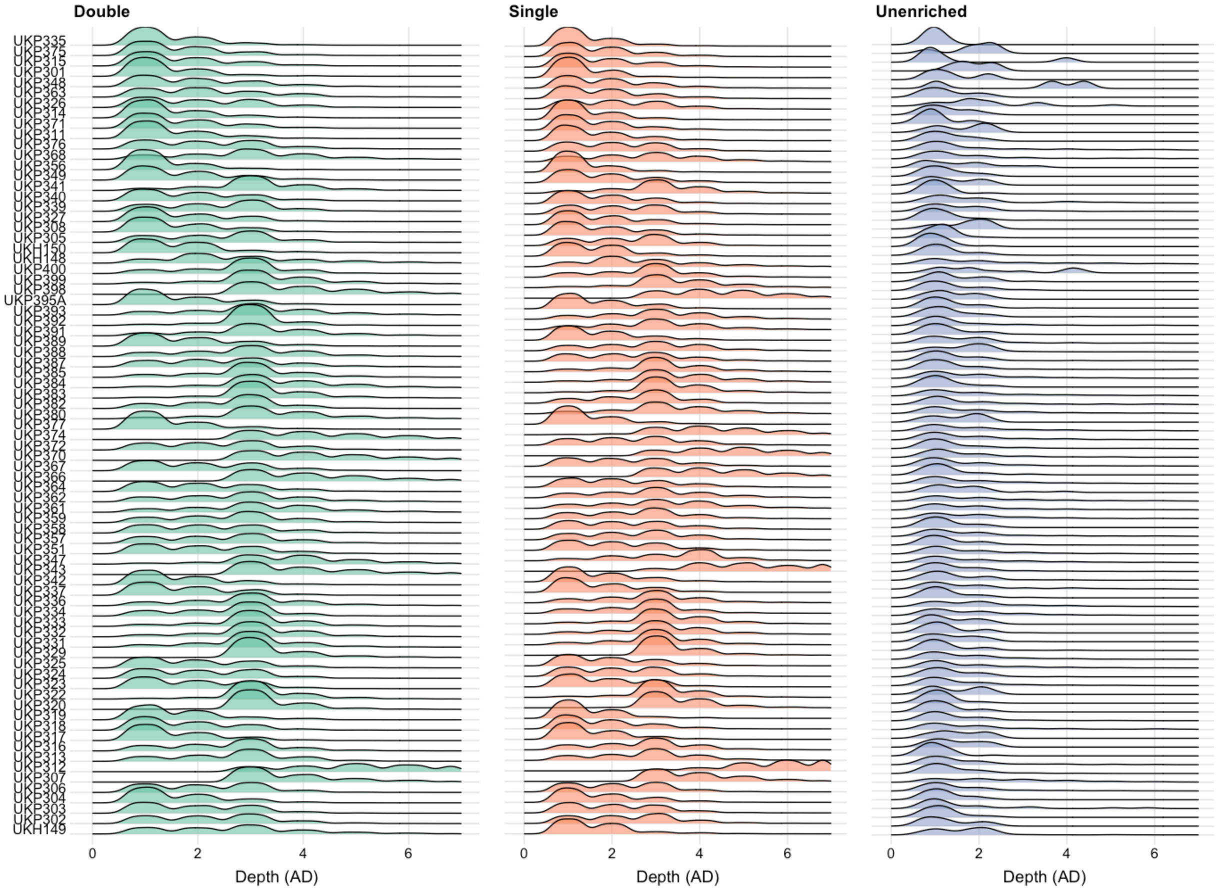


**Suppl. Fig. S77.** Ridgeline plots of the frequencies of allele depth (AD, for both reference and alternate allele, even though only one gets called) across all variants from the filtered VCFs. These were obtained for shared samples from different types of enrichments (i.e., double, single, and unenriched libraries) prepared from the clinical samples set using the iNextEra protocol.

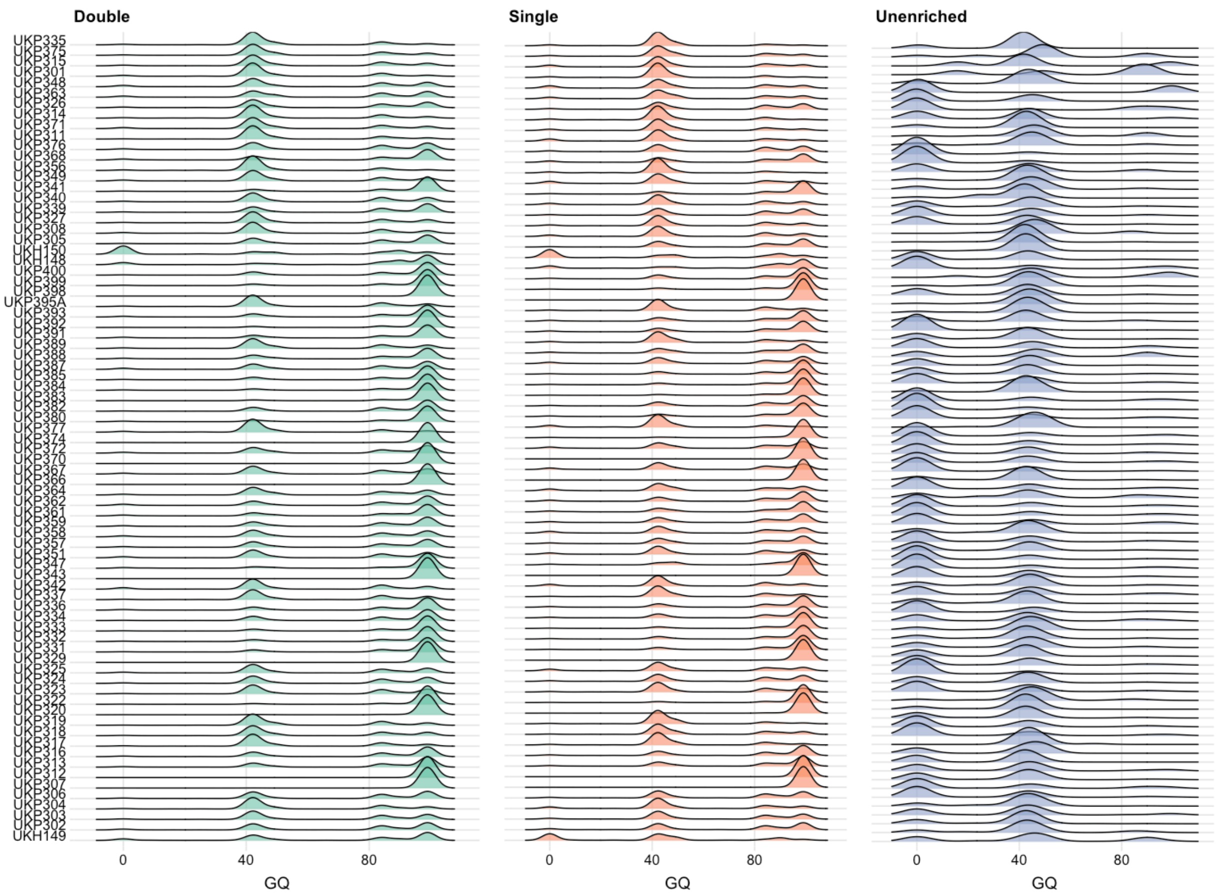


**Suppl. Fig. S78.** Ridgeline plots of the frequencies of genotype quality (QG) across all variants from the filtered VCFs. These were obtained for shared samples from different types of enrichments (i.e., double, single, and unenriched libraries) prepared from the clinical samples set using the iNextEra protocol.

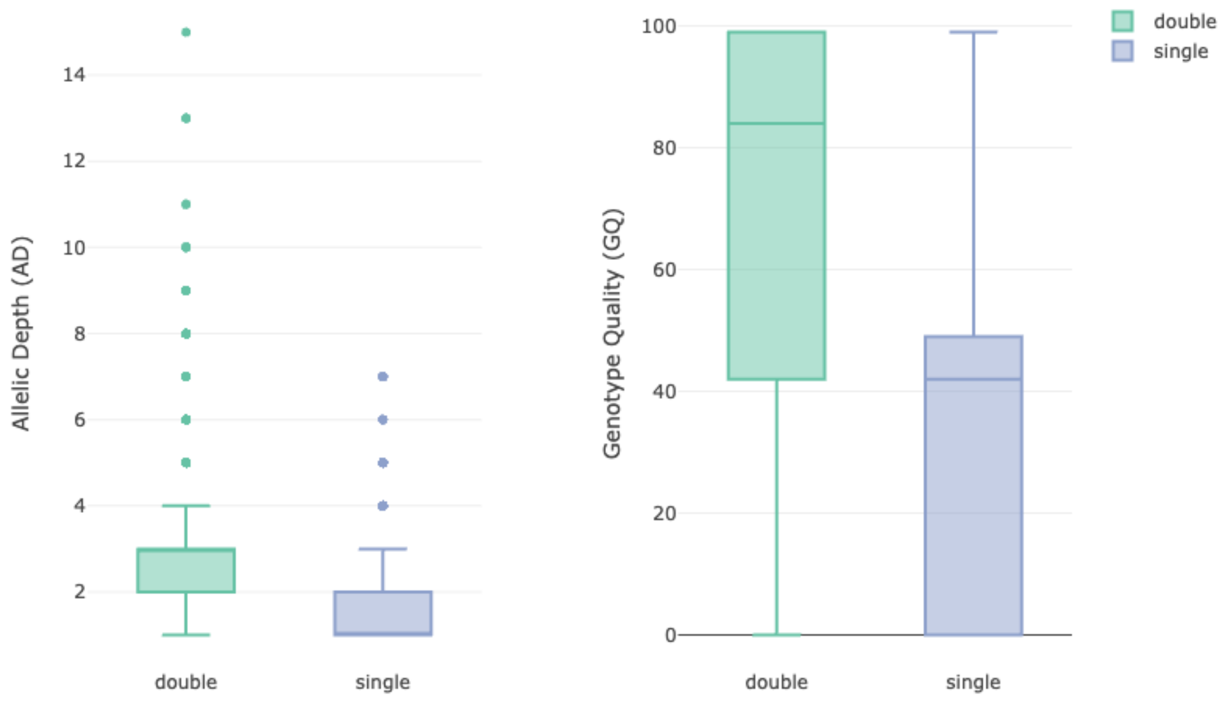


**Suppl. Fig. S79.** Left, boxplot of allelic depth (AD, for both reference and alternate allele, even though only one gets called); right, genotype quality (GQ) for all variants from double- and single-enriched libraries from the filtered VCFs for clinical sample UKH149 prepared using the iNextEra protocol. The differences between the means of both AD and GQ among the types of enrichment are statistically significant ($p < 0.0001$).

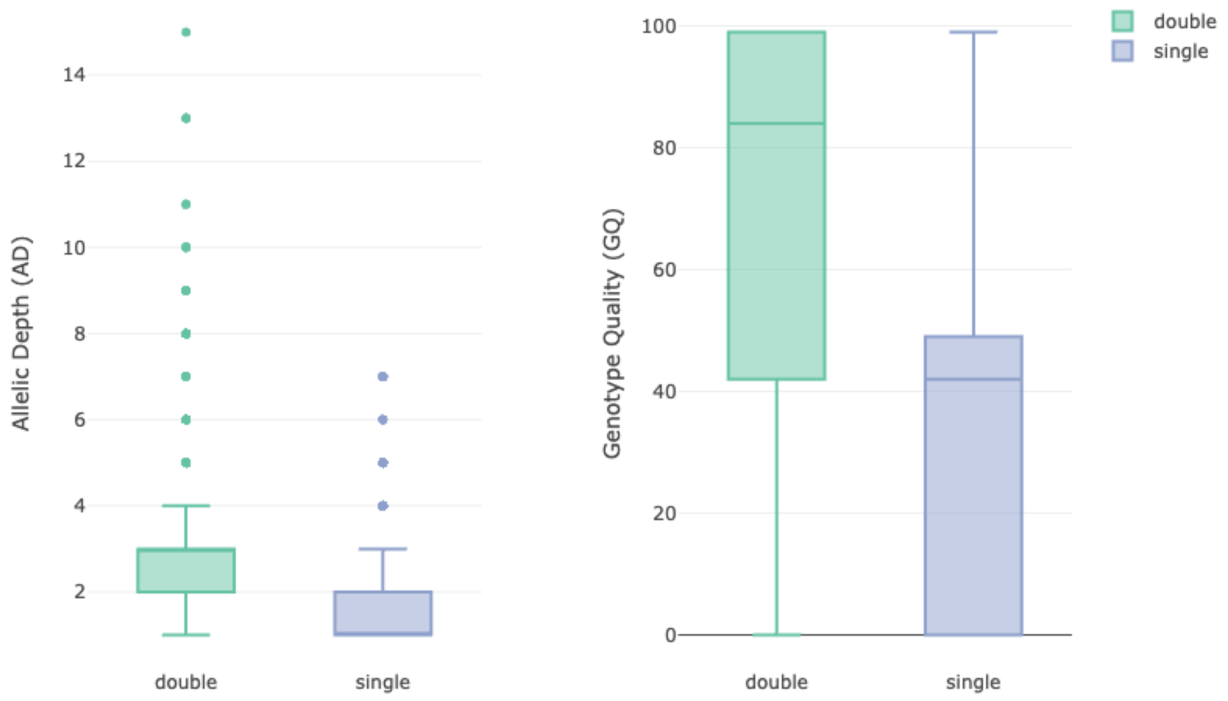


**Suppl. Fig. S80.** Left, boxplot of allelic depth (AD, for both reference and alternate allele, even though only one gets called); right, genotype quality (GQ) for all variants from double- and single-enriched libraries from the filtered VCFs for clinical sample UKP347 prepared using the iNextEra protocol. The differences between the means of both AD and GQ among the types of enrichment are statistically significant ($p < 0.0001$).

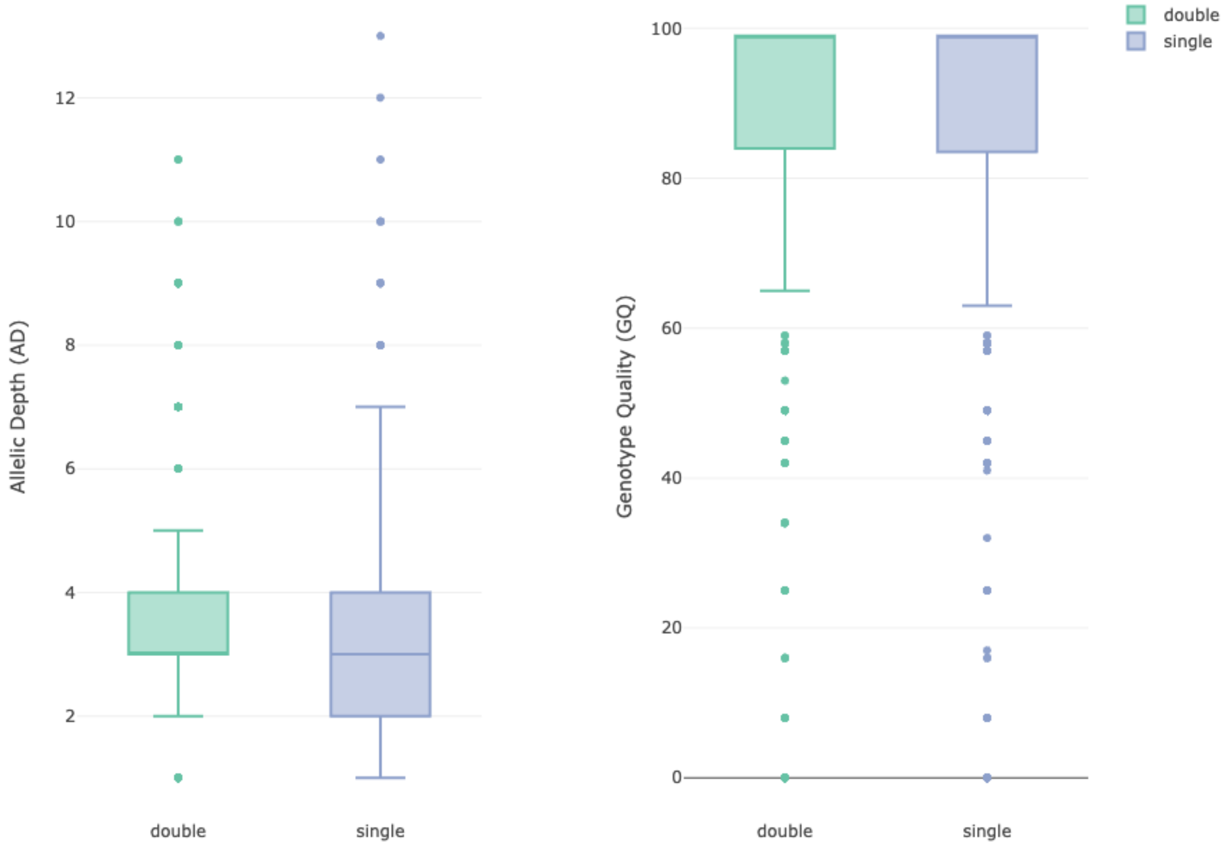


**Suppl. Fig. S81.** Left, boxplot of allelic depth (AD, for both reference and alternate allele, even though only one gets called); right, genotype quality (GQ) for all variants from double- and single-enriched libraries from the filtered VCFs for clinical sample UKP391 prepared using the iNextEra protocol. The difference between the means of AD (3.15 for double and 3.13 for single) for each type of enrichment is not statistically significant ($p = 0.64$). The difference between the means of GQ for each type of enrichment is statistically significant ($p < 0.0001$), with double enrichments having a mean of 83.8 and single enrichments having a mean of 80.4.

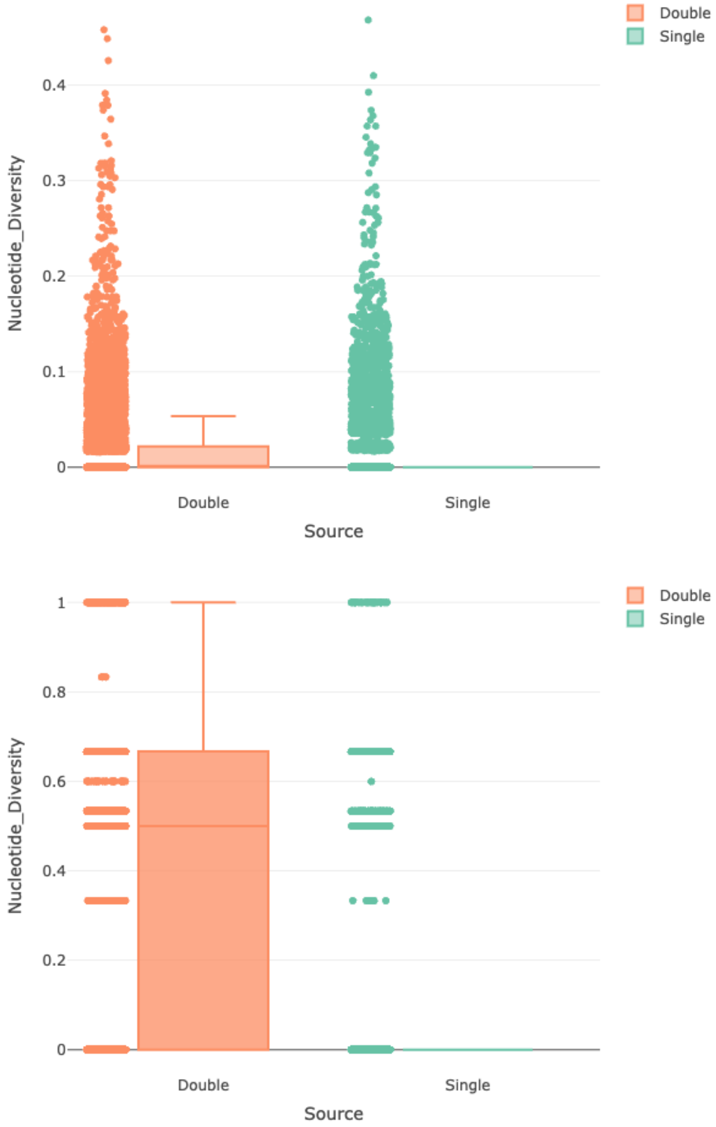


**Suppl. Fig. S82.** Top: boxplot of nucleotide diversity for UKP clinical samples for double-enriched and single-enriched libraries with CryptoCap_100k. Bottom, boxplots of nucleotide diversity for UKH clinical samples (including UKH149) for double-enriched and single-enriched libraries with CryptoCap_100k.

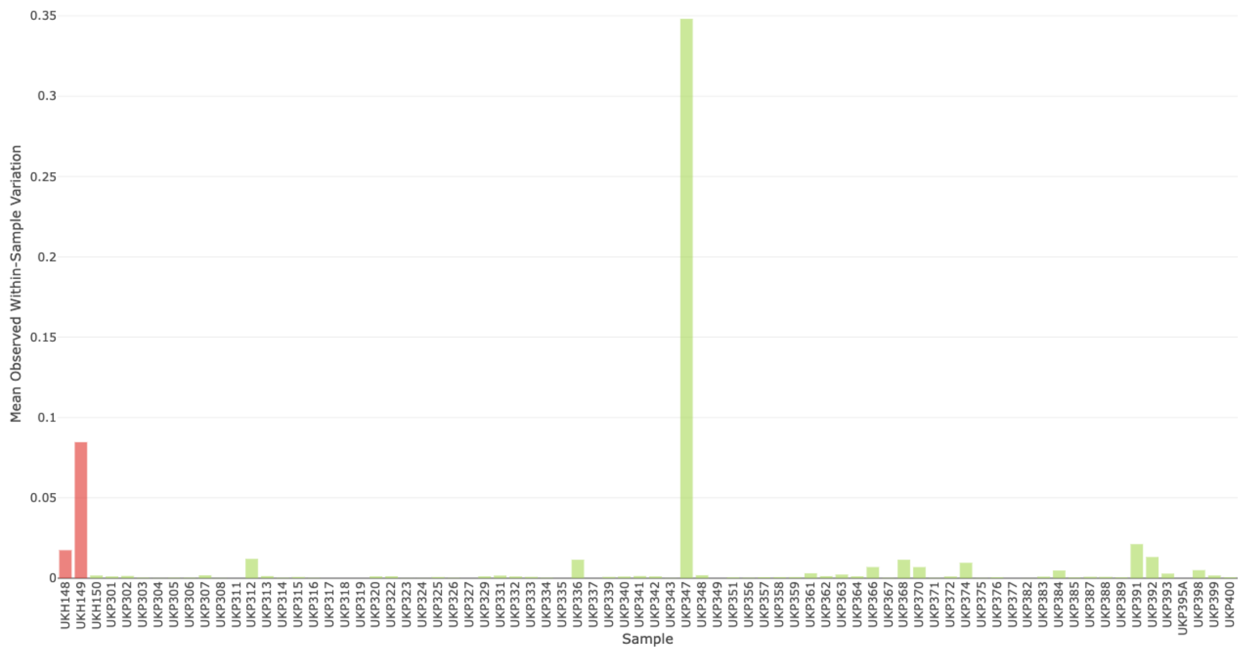


**Suppl. Fig. S83.** Mean observed within-sample allele variation calculated using 13,044 SNPs from the clinical samples set that were prepared using the iNextEra protocol and double-enriched with CryptoCap_100k.

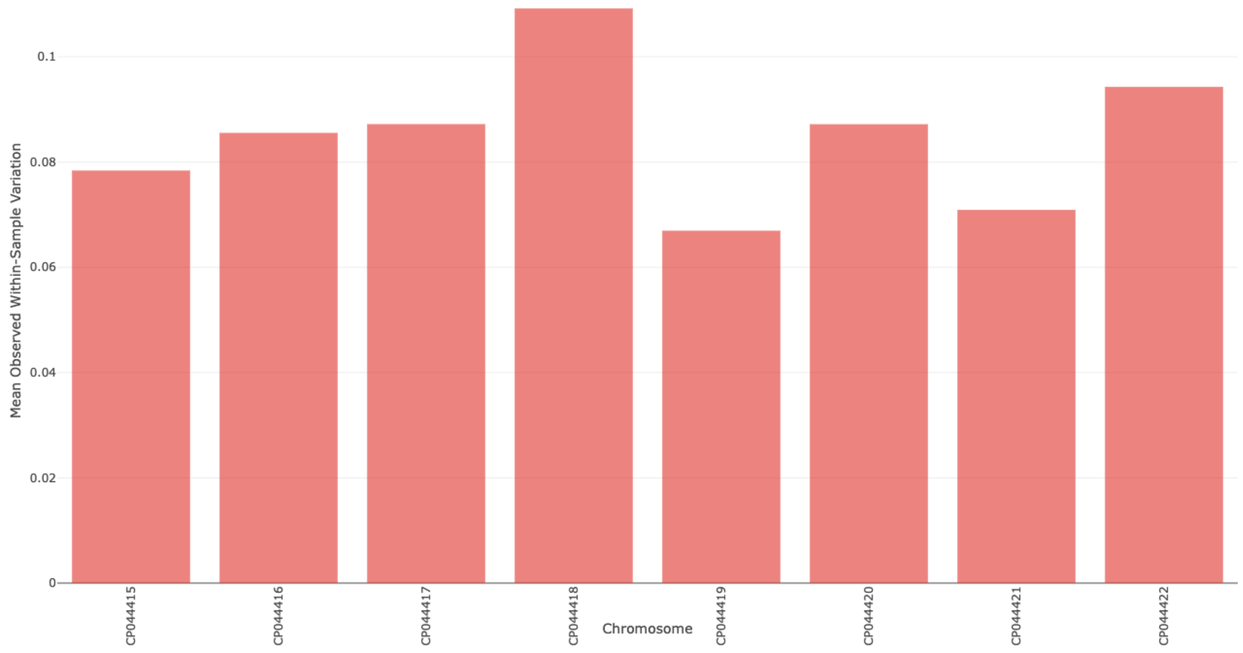


**Suppl. Fig. S84.** Mean observed within-sample allele variation per *C. parvum* IOWA-ATCC chromosome for sample UKH149 calculated using 13,044 SNPs from the clinical samples set that were prepared using the iNextEra protocol and double-enriched with CryptoCap_100k.

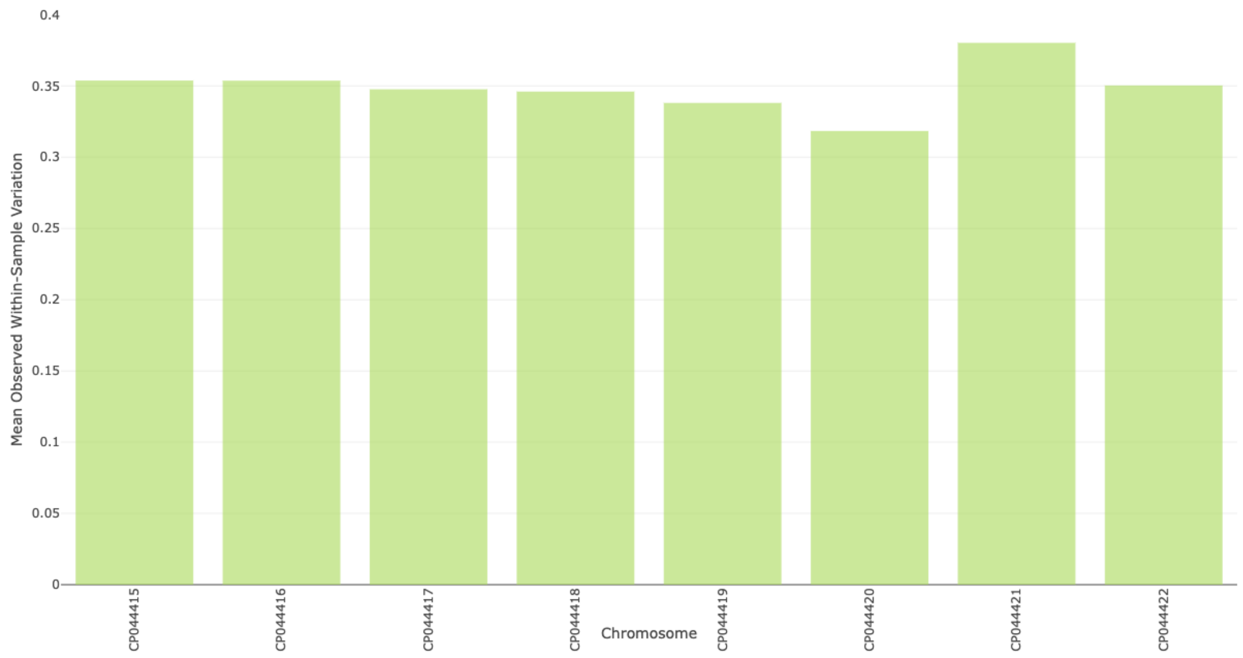


**Suppl. Fig. S85.** Mean observed within-sample allele variation per *C. parvum* IOWA-ATCC chromosome for sample UKP347 calculated using 13,044 SNPs from the clinical samples set that were prepared using the iNextEra protocol and double-enriched with CryptoCap_100k.

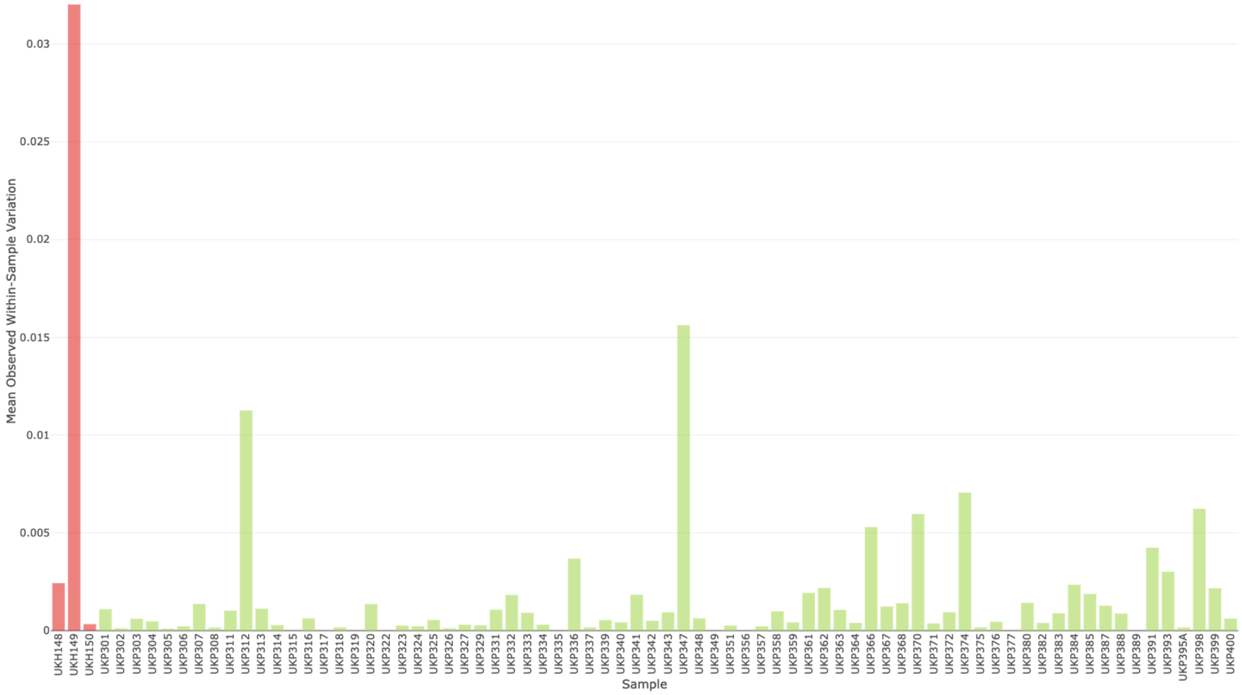


**Suppl. Fig. S86.** Mean observed within-sample allele variation calculated using 11,874 SNPs from the clinical samples set that were prepared using the iNextEra protocol and single-enriched with CryptoCap_100k.

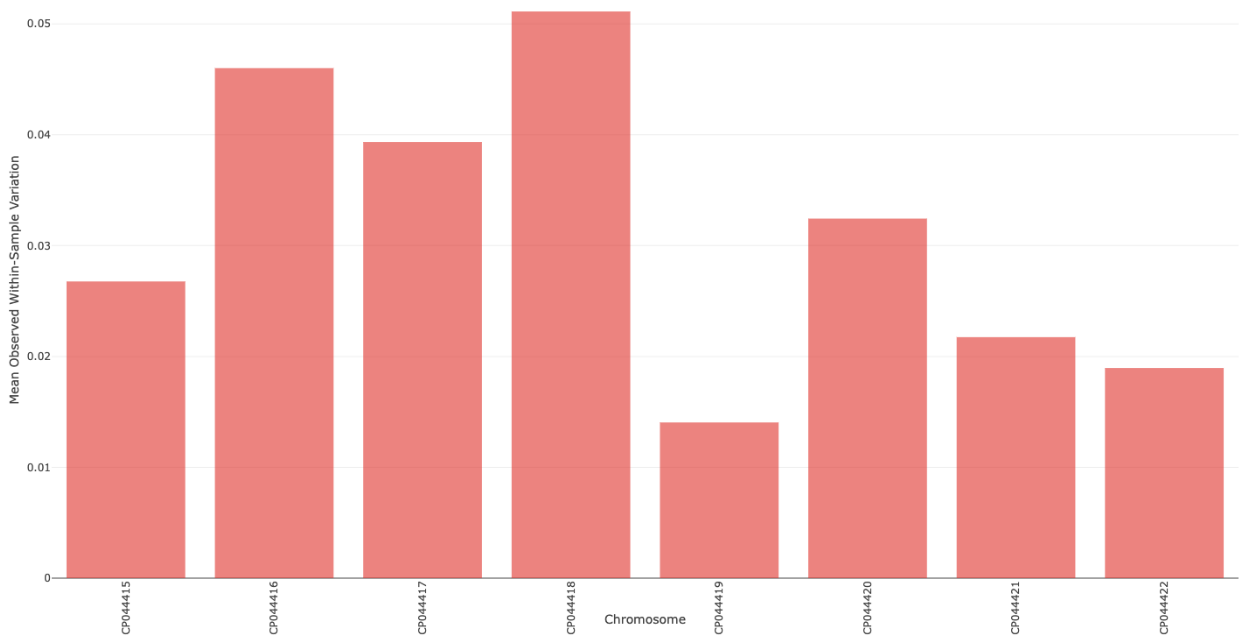


**Suppl. Fig. S87.** Mean observed within-sample allele variation per *C. parvum* IOWA-ATCC chromosome for sample UKH149 calculated using 11,874 SNPs from the clinical samples set that were prepared using the iNextEra protocol and single-enriched with CryptoCap_100k.

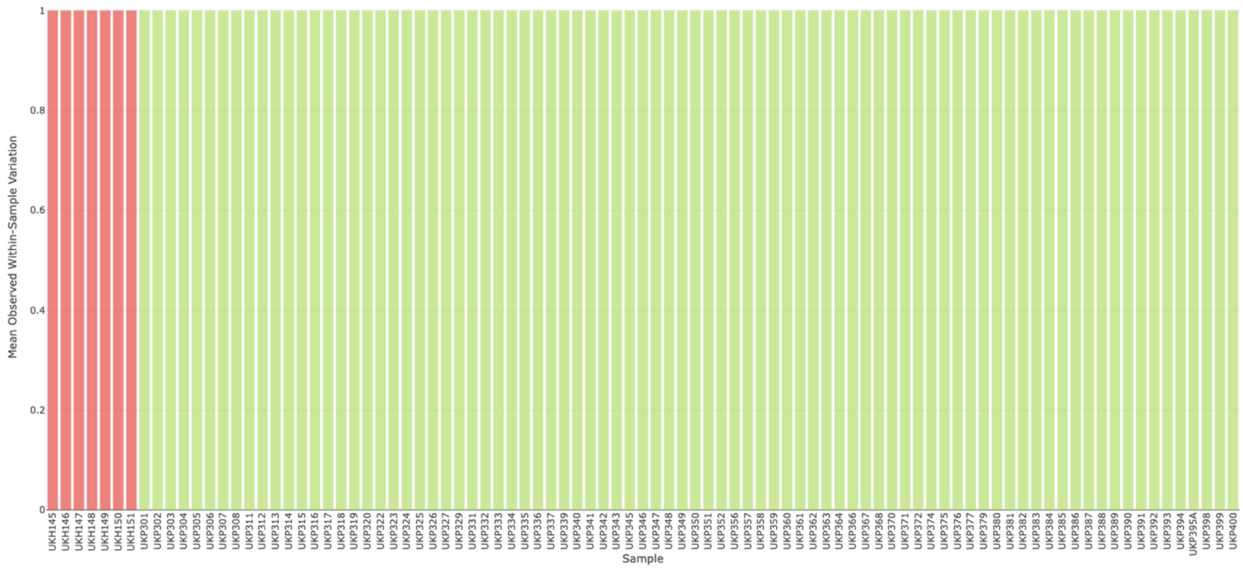


**Suppl. Fig. S88.** Mean observed within-sample allele variation calculated using 11,226 SNPs from the clinical samples set that were prepared using the iNextEra protocol and left unenriched.

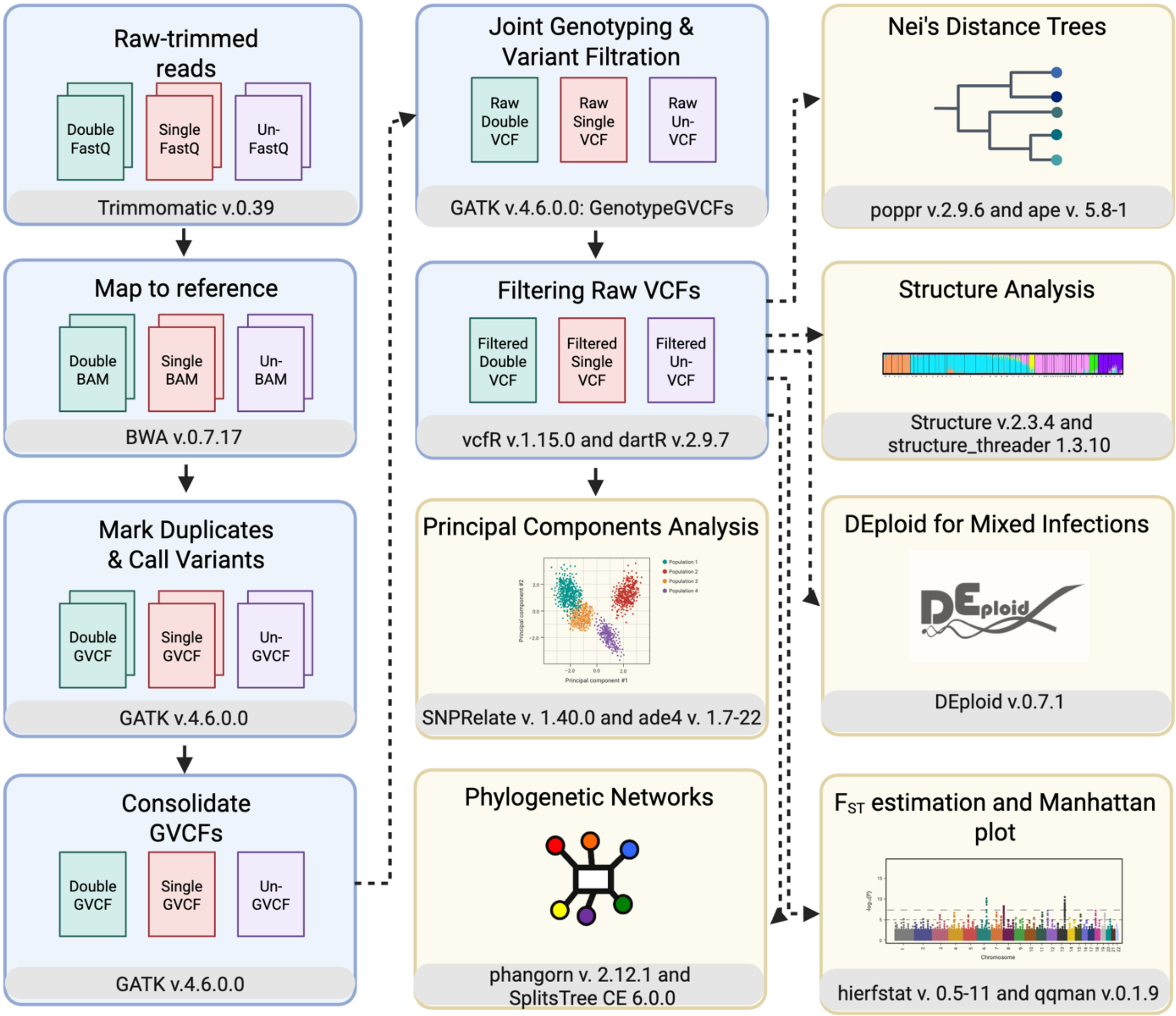


**Suppl. Fig. S89.** Workflow used for population genomics and mixed infection analyses of clinical samples set that were prepared using the iNextEra protocol. Created in BioRender. Bayona N (2025) https://BioRender.com/mrcat8s.
