## Supplementary Notes for "Genome-targeted enrichment and sequencing of human-infecting *Cryptosporidium* spp."

Supplementary Notes #4. Assessing the Validity and Accuracy of CryptoCap_100k Enrichment By Using Simulated Reads For Assigning Species Using Three Different Reference Databases 16
4.1 Mapping simulated enriched and sequenced reads to the 10-genomes reference database . 16
12.2.2.2 Sequence data mapped to 10-genomes reference database using BBMap 113

#### Supplementary Notes #1. CryptoCap_100k Baits Characterization

##### 1.1 Which genomes to use?

The following **reference** genomes were each independently used to test baits efficiency.

- *Cryptosporidium parvum*
  - *Cryptosporidium parvum* IOWA II (Reference): GCF_00000165345.1
  - *Cryptosporidium parvum* IOWA ATCC (obtained from: https://cryptodb.org/cryptodb/app/downloads/Current_Release/CparvumIOWA-ATCC/fasta/data/; 2021-11-16 10:53 8.8M)
- *Cryptosporidium hominis*
  - *Cryptosporidium hominis* 30976 (obtained from: https://cryptodb.org/cryptodb/app/downloads/Current_Release/Chominis30976/fasta/data/; 2021-11-16 10:53 8.8M)
  - *Cryptosporidium hominis* UdeA01 (obtained from: https://cryptodb.org/cryptodb/app/downloads/Current_Release/ChominisUdeA01/fasta/data/; 2021-11-16 10:53 8.8M)
- *Cryptosporidium cuniculus*
  - *Cryptosporidium cuniculus* strain UKCU2 (Reference): GCA_004337835.1
- *Cryptosporidium viatorum*
  - *Cryptosporidium viatorum* strain ABER_CVIA_1.0 (Reference): GCA_004337795.1
- *Cryptosporidium meleagridis*
  - *Cryptosporidium meleagridis* genome assembly obtained from Baptista et al. (unpublished)
- *Cryptosporidium tyzzeri*
  - *Cryptosporidium tyzzeri* isolate UGA55 (Reference): GCA_007210665.1
- *Cryptosporidium ubiquitum*
  - *Cryptosporidium ubiquitum* isolate 39726 (Reference): GCF_001865345.1
- *Cryptosporidium andersoni*
  - *Cryptosporidium andersoni* isolate 30847 (Reference): GCA_001865355.1
- *Cryptosporidium baileyi*
  - *Cryptosporidium baileyi* TAMU-09Q1 (Reference): GCA_001593455.1
- *Cryptosporidium muris*
  - *Cryptosporidium muris* RN66 (Reference): GCF_000006515.1

##### 1.2 Mapping the CryptoCap_100k bait set to each of the reference genomes

We used Burrows-Wheeler Aligner (bwa) 0.7.17 and Samtools v. 1.10 to map the CryptoCap_100k bait set to each of the genomes above, independently.

To do so we use the following code to *index* each genome, map the baits (*mem*), and convert the *.sam* files to *.bam* files, this was done independently for each genome:

```
time bwa index reference.fasta
time bwa mem -M reference.fasta baits.fa > probes.sam
samtools view -b probes.sam | samtools sort > probes.bam
samtools index probes.bam | samtools sort > probes_indexed.bam
```

##### 1.3 Plotting the mapping of the CryptoCap_100k baits against each genome

To plot each of the *.bam* files produced from mapping the baits against each genome, we first use the *bamCoverage* function from DeepTools v. 3.3.1 to generate a bigwig file (*.bw*). This tool generates a coverage track calculated as the number of reads per bin, where bins are short consecutive counting windows of a defined size (we used default: 50 bases).

```
bamCoverage -b probes.bam -o probes.bw
```

Then, this *.bw* file was visualized in the Integrative Genomics Viewer IGV-Web app version 1.10.8. igv.js version 2.11.9 and 2.4.3 by using as reference the corresponding genome.

**NOTE:** The results from these analyses are presented in **Fig. S1**.

##### 1.4 Obtaining CryptoCap_100k bait alignment statistics against each genome

To estimate the efficiency of bait alignment we used Samtools v. 1.10. and using the *depth -a* option to create a depth file.

```
samtools depth -a probes.bam -o probes.depth
```

Then, we used these instructions (https://sarahpenir.github.io/bioinformatics/awk/calculating-mapping-stats-from-a-bam-file-using-samtools-and-awk/) to use *awk* to obtain the mean read depth and proportion of baits mapped to the reference.

```
probes.depth | awk '{c++;s+=$3}END{print s/c}'
samtools flagstat probes.bam > probes.txt
```

The breadth of coverage was obtained by estimating the number of bases covered in the alignment over the length of the reference, when the depth coverage was at least 1X.

```
samtools mpileup probes.bam | awk -v X="${MIN_COVERAGE_DEPTH}" '$4>=X' | wc -l
grep -v ">" reference.fasta | wc | awk '{print $3-$1}'
```

#### Supplementary Notes #2. Comparing CryptoCap_75k and CryptoCap_100k Bait Sets

##### 2.1 Converting CryptoCap_75k *.bed* file to a FASTA file

The company that designed CryptoCap_75k bait set shared with us a bed file that contains the coordinates for their bait set, and the steps below are to use that file to obtain a fasta file with the sequences of CryptoCap_75k bait set.

Input: S3150752_Covered.bed Reference: * *Cryptosporidium parvum** IOWA II (Reference): GCF_00000165345.1

The *.bed* file was modified to match the headers of the FASTA file (they do not have chromosome number but refseq, for example, chr1 = NC_006980.1: https://www.ncbi.nlm.nih.gov/genome/?term=Cryptosporidium+parvum+Iowa+II).

New input: S3150752_Covered_modified.bed.

We used bedtools v.2.29.2 to convert the coordinates to fasta:

```
bedtools getfasta -fi C_parvum_GCF_000165345.1_ASM16534v1_genomic.fna \
-bed S3150752_Covered_modified.bed -fo agilent.fasta
```

This fasta file contains 4,538 sequences; however, these do not correspond to the baits yet. These correspond to the regions where baits have been designed for. These regions are of variable length, but all are multiples of 120 nucleotides. Thus, we split all sequences larger than 120 nucleotides into sequences of 120 nucleotides, which represent the baits. For example, if a fasta sequence is of 6,120 nucleotides. We would split it into 51 fasta sequences, each of 120 nucleotides. For that, the following bash scripts were written and run:

```
#!/bin/bash
# Input FASTA file
input_file="agilent.fasta"
# Output FASTA file
output_file="agilent_120_v4.fasta"

# Initialize a counter for subsequence numerals
subsequence_num=1

# Loop over each sequence in the input FASTA file
while read -r line
do
  # Check if the line is a header line (starts with ">")
 if [[ $line == ">"* ]]
  then
    # Write the previous sequence if it exists
    if [[ -n $sequence ]]
    then
      seq_length=${#sequence}
      # Check if the sequence is longer than 120 nucleotides
      if (( seq_length > 120 ))
      then
        # Calculate the number of sequences of 120 nucleotides
        num_sequences=$((seq_length / 120))

        # Loop over each subsequence
        for (( i=0; i<num_sequences; i++ ))
        do
          # Calculate the start and end positions of the subsequence
          start=$((i * 120))
          end=$((start + 119))

          # Extract the subsequence to the output file with the updated header
          echo "${header_line}_${subsequence_num}" >> "$output_file"
          echo "${sequence:$start:$((end - start + 1))}" >> "$output_file"
          # Increment the subsequence numeral
          subsequence_num=$((subsequence_num + 1))
        done
      else
        # Write the sequence as is to the output file with the original header
        echo "${header_line}_${subsequence_num}" >> "$output_file"
        echo "$sequence" >> "$output_file"
        subsequence_num=$((subsequence_num + 1))
      fi
```

```
    fi
    # Save the header line for the next sequence
    header_line=$line
    # Reset the sequence variable
    sequence=""
  else
    # Concatenate sequence lines
    sequence="$sequence$line"
  fi
done < "$input_file"
# Write the last sequence if it exists
if [[ -n $sequence ]]
then
  seq_length=${#sequence}
  # Check if the sequence is longer than 120 nucleotides
  if (( seq_length > 120 ))
  then
    # Calculate the number of sequences of 120 nucleotides
    num_sequences=$((seq_length / 120))
    # Loop over each subsequence
    for (( i=0; i<num_sequences; i++ ))
    do
      # Calculate the start and end positions of the subsequence
      start=$((i * 120))
      end=$((start + 119))
      # Extract the subsequence and write it to the output file with the updated header
      echo "${header_line}_${subsequence_num}" >> "$output_file"
      echo "${sequence:$start:$((end - start + 1))}" >> "$output_file"
      # Increment the subsequence numeral
      subsequence_num=$((subsequence_num + 1))
    done
  else
    # Write the sequence as is to the output file with the original header
    echo "${header_line}_${subsequence_num}" >> "$output_file"
    echo "$sequence" >> "$output_file"
  fi
fi
```

##### 2.2 Mapping the CryptoCap_75k bait set to each of the reference genomes

We used Burrows-Wheeler Aligner (bwa) 0.7.17 and Samtools v. 1.10 to map the CryptoCap_75k bait set developed by Agilent to each of the genomes above, independently, with the aim of simulating what would be an error-free, bias-free enrichment process.

To do so, we use the code:

```
time bwa index reference.fasta
time bwa mem -M reference.fasta agilent.fasta > probes.sam
samtools view -b agilent.sam | samtools sort > agilent.bam
```

##### 2.3 Obtaining CryptoCap_75k bait alignment statistics against each genome

To estimate the efficiency of CryptoCap_75k bait alignment and compare it to CryptoCap_100K, we used Samtools v.1.10. and using the *depth -a* option to create a depth file.

```
samtools depth -a probes.bam -o probes.depth
```

Then, we use these instructions (https://sarahpenir.github.io/bioinformatics/awk/calculating-mapping-stats-from-a-bam-file-using-samtools-and-awk/) to use *awk* to obtain the mean read depth and proportion of baits mapped to the reference.

```
cat probes.depth | awk '{c++;s+=$3}END{print s/c}'
samtools flagstat probes.bam > probes.txt
```

The breadth of coverage was obtained by estimating the number of bases covered in the alignment over the length of the reference, when the depth coverage was at least 1X.

```
samtools mpileup probes.bam | awk -v X="${MIN_COVERAGE_DEPTH}" '$4>=X' | wc -l
grep -v ">" reference.fasta | wc | awk '{print $3-$1}'
```

##### 2.4 Plotting the mapping of the CryptoCap_75k baits against each genome

To plot each of the *.bam* files produced from mapping the baits against each genome, we first use the *bamCoverage* function from DeepTools v. 3.3.1 to generate a bigwig file (*.bw*). This tool generates a coverage track calculated as the number of reads per bin, where bins are short consecutive counting windows of a defined size (we used default: 50 bases).

```
bamCoverage -b probes.bam -o probes.bw
```

Then, this *.bw* file was visualized in the Integrative Genomics Viewer IGV-Web app version 1.10.8, igv.js version 2.11.9, using the corresponding genome as a reference. Individual plots, as well as *.ppt* for editing figures, are in the bait_plots/ folder. NOTE: The index file is automatically created in the desktop version, and then we imported both to the online version.

**NOTE:** The results from these analyses are presented in **Fig. S2**.

We created a text document of tabulated information with the statistics of the mapping of each bait set, CryptoCap_75k and CryptoCap_100k, to the genomes.

```
file <- read.delim("~/crypto_data/2/baits_comparisons.txt",
    header = TRUE)
table <- data.frame(file)
names(table) <- c("bait_set", "species", "mean_read_depth", "percent_breadth_of_coverage",
    "percent_baits_mapped")
# table
```

##### 2.5 Mean read depth for each bait set, for each species, and overall species

```r
library(plotly)
p1 <- plot_ly(table, x = table$species, y = table$mean_read_depth,
    type = "box", color = table$bait_set) %>%
    layout(yaxis = list(title = "Mean Read Depth"), showlegend = FALSE)
p1 <- p1 %>%
    add_trace(x = table$bait_set, marker = list(color = "rgba(1,102,94,0.8)",
        line = list(width = 1, color = "rgb(255, 255, 255)")))
# p1 orca(p1, 'bait_plots/baits_sets_mean_read_depth.pdf')
```

```r
t.test(table$mean_read_depth ~ table$bait_set)
```

##### 2.5.1 Statistical test of difference of the means of mean read depth for each bait set for each species and overall species

```
Welch Two Sample t-test
```

data: table$mean_{r}ead_{d}epthbytable$bait_set t = -0.67798, df = 21.26, p-value = 0.5051 alternative hypothesis: true difference in means between group CryptoWGE_V1 and group CryptoWGE_V2 is not equal to 0 95 percent confidence interval: -0.4888236 0.2483236 sample estimates: mean in group CryptoWGE_V1 mean in group CryptoWGE_V2 0.57250 0.69275

#### 2.6 Percent breadth of coverage for each bait set for each species and overall species

```r
p2 <- plot_ly(table, x = table$species, y = table$percent_breadth_of_coverage,
    type = "box", color = table$bait_set) %>%
    layout(yaxis = list(title = "Percent Breadth Of Coverage"),
        showlegend = FALSE)
p2 <- p2 %>%
    add_trace(x = table$bait_set, marker = list(color = "rgba(1,102,94,0.8)",
        line = list(width = 1, color = "rgb(255, 255, 255)")))
```

```r
t.test(table$percent_breadth_of_coverage ~ table$bait_set)
```

##### 2.6.1 Statistical test of difference of the means of breadth of coverage for each bait set for each species and overall species

```
Welch Two Sample t-test
```

data: table$percent_{b}readth_{o}f_{c}overagebytable$bait_set t = -0.34084, df = 21.86, p-value = 0.7365 alternative hypothesis: true difference in means between group CryptoWGE_V1 and group CryptoWGE_V2 is not equal to 0 95 percent confidence interval: -40.29888 28.92604 sample estimates: mean in group CryptoWGE_V1 mean in group CryptoWGE_V2 57.06833 62.75475

##### 2.7 Percent of baits mapped from each bait set to each species and overall species

```r
p3 <- plot_ly(table, x = table$species, y = table$percent_baits_mapped,
    type = "box", color = table$bait_set) %>%
    layout(yaxis = list(title = "Percent Baits Mapped"), showlegend = FALSE)
p3 <- p3 %>%
    add_trace(x = table$bait_set, marker = list(color = "rgba(1,102,94,0.8)",
        line = list(width = 1, color = "rgb(255, 255, 255)")))
```

```r
t.test(table$percent_baits_mapped ~ table$bait_set)
```

###### 2.7.1 Statistical test of difference of the means of percent of reads mapped for each bait set for each species and overall species

```
Welch Two Sample t-test
```

data: table$percent_{b}aits_{m}appedbytable$bait_set t = 0.56144, df = 21.489, p-value = 0.5803 alternative hypothesis: true difference in means between group CryptoWGE_V1 and group CryptoWGE_V2 is not equal to 0 95 percent confidence interval: -24.66636 42.94469 sample estimates: mean in group CryptoWGE_V1 mean in group CryptoWGE_V2 64.72583 55.58667

##### 2.8 All stats summarized

```r
all_1 <- plot_ly(table, x = table$bait_set, y = table$mean_read_depth,
    type = "box", color = table$bait_set) %>%
    layout(yaxis = list(title = "Mean Read Depth"), showlegend = FALSE)
# all_1

all_2 <- plot_ly(table, x = table$bait_set, y = table$percent_breadth_of_coverage,
    type = "box", color = table$bait_set) %>%
    layout(yaxis = list(title = "Percent Breadth Of Coverage"),
        showlegend = FALSE)
# all_2

all_3 <- plot_ly(table, x = table$bait_set, y = table$percent_baits_mapped,
    type = "box", color = table$bait_set) %>%
    layout(yaxis = list(title = "Percent Baits Mapped"), showlegend = FALSE)
# all_3

all_comparisons <- subplot(list(all_1, all_2, all_3), nrows = 1,
    margin = 0.04, shareX = TRUE, titleY = TRUE)
# orca(all_comparisons, 'bait_plots/all_comparisons.pdf')
# all_comparisons
```

**NOTE:** The results from these analyses are presented in **Fig. S3**.

#### Supplementary Notes #3. Simulating the Enrichment with CryptoCap_100k Baits and Sequencing in Each of the Reference Genomes

As a first approach to validate whether we could obtain meaningful data before synthesis and experimental design with the baits, in a simple infection scenario, error-free and bias-free, we used twelve available genome sequences for ten Cryptosporidium species to simulate the enrichment and sequencing process with the CryptoCap_100k bait set. Each genome was treated as if it belonged to a sample containing only such a genome. First, we obtained the coordinates of the above-mapped baits on each of the twelve genome sequences (CryptoCap_100k Bait Design & Characterization). To simulate the hybridization process, we extended those mapping positions to several bases to the left and right, assuming that the bait was annealing to the centroid of larger fragments being captured. We performed two types of extensions to understand the effect of mean library fragment size on enrichment. We thus simulated the capture of fragments with two different size distributions. We took the first mapping positions for all baits and extended each region for large fragments, 200 bases to the left and 320 bases (120 bait +200) to the right, to simulate a library insert size of 520 bp. Also, we extended the coordinate where the bait mapped 40 bases to the left and 160 bases (120 bait + 40) to the right to simulate a library insert size of 200 bp. Then, the faidx option from Samtools was used to extract the two types of extended sequences from each genome to create a fasta file.

Importantly, BWA tolerates a hit with at most k differences (mismatches or gaps), where k is chosen such that <4% of m-long reads with 2% uniform base error rate may contain differences more than k. With this configuration, for 120 bait sequences, BWA is allowing 5 mismatches between the query and the reference (Li and Dublin, 2009). Our mapping above considered these default parameters.

##### 3.1 Obtaining the coordinates of the mapped baits on each reference genome

We use the *.sam* file from the mapping to obtain the position in each genome where the baits mapped with the aim to simulate what would be an error-free, bias-free, enrichment process.

```
cat probes.sam | grep -v "@" | cut -f 3,4 > probes_coordinates.txt
```

Coordinates that did not contain the reference name (RNAME) and therefore no mapping positions in the *.sam* file were excluded from the analyses.

The output looks something like this, but many lines per file:

> CpIOWA-ATCC-Chr1_46706 LRBS01000099.1 22131

In this example, the bait *CpIOWA-ATCC-Chr1_46706* mapped to the *LRBS01000099.1* contig from the C. andersoni genome at the leftmost position 22,131.

##### 3.2 Simulating capture of large fragments ~520 bp

In the lab, after the CryptoCap_100k baits hybridize to the *Cryptosporidium* genome(s) present in the sample, we retrieve these using a magnet that attracts the biotinylated baits. However, the fragments pulled from the sample are bigger than the 120 nucleotides of the bait's length, depending on the fragment size distribution of the library prep.

In this simulation experiment, we aim to mimic these larger fragments by taking the coordinate of the first position where the bait mapped and increasing the fragment size by adding 200 bases to the left, and 320 bases (120 of bait + 200) to the right of the mapped region in the reference. If the mapped region started within the first 200 bases, we started from base 1 of the contig to the 320th base.

To do this, the coordinates file for each genome was imported to Excel and the extra bases were added to the left and right, accordingly to the paragraph above. Using as an example the output presented above, we will extract from the *C. andersoni* genome, from the LRBS01000099.1 contig, the sequence from position 21931 to position 22451 (22131-200 — 22131+320). Then, for each genome, we created a text file called "regions_large" that contained the coordinate information of each region to extract from the reference genome (folder coordinates).

This is an example of few lines from the regions_large.txt:

LRBS01000019.1:7811-8331
LRBS01000069.1:7888-8408
LRBS01000080.1:8232-8752
LRBS01000103.1:8748-9268
LRBS01000103.1:8919-9439
LRBS01000070.1:9049-9569

Now, to extract these sequences from the reference genome we used the *faidx* option from SamTools for each species that looked like this:

```
mkdir fastas
samtools faidx reference.fna -r regions_large.txt > fastas/extracted_regions_large.fasta
```

##### 3.3 Simulating capture of short fragments ~200 bp

In this simulation experiment, we are trying to mimic shorter fragments by taking the coordinate of the first position where the bait mapped and increase the fragment size by adding 40 bases to the left, and 160 bases (120 of bait + 40) to the right of the mapped region in the reference. If the mapped region started within the first 40 bases, we started from base 1 of the contig to the 160th base.

To do this, in the same coordinates file for each genome the extra bases were added to the left and right, accordingly to the paragraph above. Using as example the output presented above, we will extract from the *C. andersoni* genome, from the LRBS01000099.1 contig, the sequence from position 22091 to position 22291 (22131-40 — 22131+160). Then, for each genome we created a text file called "regions_short" that contained the coordinate information of each region to extract from the reference genome (folder coordinates).

This is an example of few lines from the regions_short.txt:

LRBS01000019.1:7971-8171
LRBS01000069.1:8048-8248
LRBS01000080.1:8392-8592
LRBS01000103.1:8908-9108
LRBS01000103.1:9079-9279
LRBS01000070.1:9209-9409

Now, to extract these sequences from the reference genome we used the *faidx* option from SamTools for each species that looked like this:

```
mkdir fastas
samtools faidx reference.fna -r regions_short.txt > fastas/extracted_regions_short.fasta
```

##### 3.4 Simulating fastq reads from the fasta file of mapped reads

To simulate the sequencing process from these "captured" fragments, the software ART 2016.06.05 was used to reproduce ~200,000 paired-end Illumina 150 bp fastq reads from each fasta. Each set of fastq files was mapped to the respective genome sequence using BWA v 0.7.17, and we estimated depth and breadth coverage from these reads on each genome and produced bigwig files for visualization of the results.

We used ART 2016.06.05 to simulate fastq paired reads for both, short and large fragments,each read is 150 bases length, and the mean size of DNA fragments for paired-end simulations was 500 bases for large and 200 for short, we modified –fcov parameter to generate around ~200k reads for each genome, by counting the number of regions obtained in the fasta above.

```
art_illumina \
--paired \
--in fastas/extracted_regions_large.fasta \
--out reads \
--len 150 --rcount X --mflen 500 --sdev 150
```

###### 3.4.1 Simulating reads for library with large fragments

```
art_illumina \
--paired \
--in fastas/extracted_regions_short.fasta \
--out reads \
--len 150 --rcount X --mflen 200 --sdev 50
```

###### 3.4.2 Simulating reads for library with short fragments

##### 3.5 Mapping simulated fastq reads to their respective *Cryptosporidium* genome

Then, we assessed and validated the simulated data species assignment accuracy. To do this, we used three reference databases. First, we generated one fasta file containing ten Cryptosporidium genome sequences that served as a reference (10-CrypGS, from the twelve genome sequences reference, we removed C. hominis UdeA01 and C. parvum IOWA II). Second, we used a database of gp60 sequences, the most widely used genetic marker for subtyping Cryptosporidium, which contains the different subtypes identified in human cases. Third, we used a curated CryptoDB.org database of 18S rRNA, the second most characterized loci in the group.

To map each set of simulated, both large and short fragments, fastq reads against the respective genome, we used BWA (same version as above) and then converted the sam file into a bam file and indexed it:

```
time bwa index reference.fasta
time bwa mem -M 1.fq 2.fq > sim_mapped.sam
samtools view -b sim_mapped.sam | samtools sort > sim_mapped.bam
samtools index sim_mapped.bam > /sim_mapped_indexed.bam
```

**3.5.1 Visualizing the mapping of simulated fastqs across each genome** To be able to plot the bam files produced from the mapping of the simulated fastqs against each genome separately, we used again the bamCoverage function from DeepTools v. 3.3.1 to produce a bigwig file:

```
bamCoverage -b sim_mapped.bam -o sim_mapped.bw
```

Then, this *bigwig* file was visualized in the Integrative Genomics Viewer IGV-Web app version 1.10.8, igv.js version 2.11.9, by using as reference the corresponding genome.

**NOTE:** The results from these analyses are presented in **Fig. S4**.

We also obtained chromosome-level plots for: ***C. hominis* UdeA**

**NOTE:** The results from these analyses are presented in **Fig. S5**.

And we also obtained chromosome-level plots for ***C. parvum* IOWA ATCC**

**NOTE:** The results from these analyses are presented in **Fig. S6**.

Plots of breadth and depth of coverage for those genomes with chromosome-level assemblies (C. hominis UdeA01 and C. parvum IOWA-ATCC) corroborated the continuity of simulated reads from large fragments across all chromosomes (Figs. S5–S6).

**3.5.2 Obtaining simulated reads alignment statistics against each genome** To estimate the efficiency of bait alignment, we used Samtools v. 1.10. and used the *depth -a* option to create a depth file:

```
samtools depth -a probes.bam -o probes.depth
```

Then, we use these instructions (https://sarahpenir.github.io/bioinformatics/awk/calculating-mapping-stats-from-a-bam-file-using-samtools-and-awk/) to use *awk* to obtain the mean read depth and proportion of baits mapped to the reference:

```
probes.depth | awk '{c++;s+=3}END{print s/c}'
samtools flagstat probes.bam > probes.txt
samtools mpileup probes.bam | awk -v X="${MIN_COVERAGE_DEPTH}" '$4>=X' | wc -l
```

#### Supplementary Notes #4. Assessing the Validity and Accuracy of Crypto-Cap_100k Enrichment By Using Simulated Reads For Assigning Species Using Three Different Reference Databases

We used BBMap 38.90 to map fastq reads to each of the three reference files, which by default allows multi-mapping (secondary alignment) of baits to the genomes. We only considered regions where both reads were mapped to count the number of hits. We plotted the proportion of paired hits to specific Cryptosporidium species for each simulated dataset.

##### 4.1 Mapping simulated enriched and sequenced reads to the 10-genomes reference database

We mapped each set of simulated reads from each genome against the fasta containing ten genomes from different *Cryptosporidium* species, we used BBmap and as reference we use the reference file with 10 genomes. We used a maxim indel length of 20, a minid of 0.8 (read note below), we also required correct strands for paired reads (rcs), and set the mean pair size to 400, and did not rescue paired reads if avg. insert size greater than 600 (rescuedist; let's remember that the simulated size is ~520), and we sat an average pair distance *(apd)of 100 bases.

```
mkdir sim_mapped_ten
bbmap.sh in=1.fq in2=2.fq \
ref 10_crypto_genomes.fasta  \
maxindel=20 minid=0.80 rcs=t killbadpairs=t pairlen=400 \
rescuedist=600 apd=100 outu=sim_mapped_ten/ten_unmapped.sam \
outm=sim_mapped_ten/ten_mapped.sam printunmappedcount=t \
covstats=sim_mapped_twelve/twelve.cov.txt threads=12 \
-Xmx180g 2>&1 | tee sim_mapped_ten/ten.bbmap.log
```

For each species, we used the FLAG count from the third column of our SAM files to extract those sequences for which only both reads mapped (FLAG: 83 and 99):

```
cat ten_mapped.sam | grep -v "^@" |\
awk 'BEGIN{FS="\t";OFS="\t"}{if($2=="83" || $2=="99") print $0}' \
> both_mapped/ten_mapped_both.txt
```

Then, for each species, we will count each species ID hit in our mapped file. The count text has two columns, the first is the species ID, and the second is the counts:

```
cat ten_mapped_both.txt | \
awk 'BEGIN{FS="\t";OFS="\t"}{OTU[$3]+=1}END{for (i in OTU) print i,OTU[i]}' \
> crypto_species/species_ct.txt
```

This generates a file that looks like this:
> C_andersoni-LRBS01000112.1 Cryptosporidium andersoni isolate 30847 contig_19, whole genome shotgun sequence 1800

These results are summarized in **Table S2**.

###### 4.1.1 Species assignment based on mapping simulated reads from large fragment libraries ~520bp

To begin with, each file has two columns; the first one contains the reference genome information, and the second is the count of hits that the simulated reads had against target regions of each species. We split the first column into two to have in the first the "Species" name to which the reads hit, and the second is "Locus" with the rest of the information of the reference genome where the reads mapped.
Below, we import the modified files for each species and then create a fourth column, "Origin" with the name of the species for which we simulated the reads.

```
# Import Libraries not only for this chunk, but for all
# chunks
library(tidyverse)
library(extrafont)
library(ape)
library(vegan)
library(plotly)
library(lsmeans)
library(dplyr)
library(reshape2)  # to load tips data
library(tidymodels)  # for the fit() function

# Setting colors for specific species
col_c_andersoni <- c("#8dd3c7")
col_c_baileyi <- c("#ffffb3")
```

```r
col_c_cuniculus <- c("#bebada")
col_c_hominis_30976 <- c("#fb8072")
col_c_meleagridis <- c("#80b1d3")
col_c_muris <- c("#fdb462")
col_c_parvum <- c("#b3de69")
col_c_tyzzeri <- c("#fccde5")
col_c_ubiquitum <- c("#d9d9d9")
col_c_viatorum <- c("#bc80bd")
col_c_hominis_UdeA <- c("#ccebc5")
col_c_parvum_ATCC <- c("#ffed6f")
col_c_ten <- c("#8dd3c7", "#ffffb3", "#bebada", "#fb8072", "#80b1d3",
    "#fdb462", "#b3de69", "#fccde5", "#d9d9d9", "#bc80bd")

# Cryptosporidium andersoni
sim_Ca <- read.table("~/crypto_data/4/C_andersoni_largespecies_ct.txt",
    header = F, sep = "\t", stringsAsFactors = F)
colnames(sim_Ca) <- c("Species", "Locus", "Count")
# adding extra column
sim_Ca <- sim_Ca %>%
    mutate(Origin = "C_andersoni")
# Calculate Relative abundance of counts for each genome
sim_Ca <- sim_Ca %>%
    group_by(Origin) %>%
    mutate(relative_ab = Count/sum(Count))

# Cryptosporidium baileyi
sim_Cb <- read.table("~/crypto_data/4/C_baileyi_largespecies_ct.txt",
    header = F, sep = "\t", stringsAsFactors = F)
colnames(sim_Cb) <- c("Species", "Locus", "Count")
# adding extra column
sim_Cb <- sim_Cb %>%
    mutate(Origin = "C_baileyi")
# Calculate Relative abundance of counts for each genome
sim_Cb <- sim_Cb %>%
    group_by(Origin) %>%
    mutate(relative_ab = Count/sum(Count))

# Cryptosporidium cuniculus
sim_Cc <- read.table("~/crypto_data/4/C_cuniculus_largespecies_ct.txt",
    header = F, sep = "\t", stringsAsFactors = F)
colnames(sim_Cc) <- c("Species", "Locus", "Count")
# adding extra column
sim_Cc <- sim_Cc %>%
    mutate(Origin = "C_cuniculus")
# Calculate Relative abundance of counts for each genome
sim_Cc <- sim_Cc %>%
    group_by(Origin) %>%
    mutate(relative_ab = Count/sum(Count))

# Cryptosporidium hominis
sim_Ch <- read.table("~/crypto_data/4/C_hominis_30976_largespecies_ct.
    header = F, sep = "\t", stringsAsFactors = F)
colnames(sim_Ch) <- c("Species", "Locus", "Count")
```

```r
# adding extra column
sim_Ch <- sim_Ch %>%
    mutate(Origin = "C_hominis")
# Calculate Relative abundance of counts for each genome
sim_Ch <- sim_Ch %>%
    group_by(Origin) %>%
    mutate(relative_ab = Count/sum(Count))

# Cryptosporidium hominis UdeA
sim_ChUdeA <- read.table("~/crypto_data/4/C_hominis_UdeA01_largespecies_ct.txt"
    header = F, sep = "\t", stringsAsFactors = F)
colnames(sim_ChUdeA) <- c("Species", "Locus", "Count")
# adding extra column
sim_ChUdeA <- sim_ChUdeA %>%
    mutate(Origin = "C_hominis_UdeA")
# Calculate Relative abundance of counts for each genome
sim_ChUdeA <- sim_ChUdeA %>%
    group_by(Origin) %>%
    mutate(relative_ab = Count/sum(Count))

# Cryptosporidium meleagridis
sim_Cm <- read.table("~/crypto_data/4/C_meleagridis_largespecies_ct.txt"
    header = F, sep = "\t", stringsAsFactors = F)
colnames(sim_Cm) <- c("Species", "Locus", "Count")
# adding extra column
sim_Cm <- sim_Cm %>%
    mutate(Origin = "C_meleagridis")
# Calculate Relative abundance of counts for each genome
sim_Cm <- sim_Cm %>%
    group_by(Origin) %>%
    mutate(relative_ab = Count/sum(Count))

# Cryptosporidium muris
sim_Cmu <- read.table("~/crypto_data/4/C_muris_largespecies_ct.txt",
    header = F, sep = "\t", stringsAsFactors = F)
colnames(sim_Cmu) <- c("Species", "Locus", "Count")
# adding extra column
sim_Cmu <- sim_Cmu %>%
    mutate(Origin = "C_muris")
# Calculate Relative abundance of counts for each genome
sim_Cmu <- sim_Cmu %>%
    group_by(Origin) %>%
    mutate(relative_ab = Count/sum(Count))

# Cryptosporidium parvum
sim_Cp <- read.table("~/crypto_data/4/C_parvum_largespecies_ct.txt",
    header = F, sep = "\t", stringsAsFactors = F)
colnames(sim_Cp) <- c("Species", "Locus", "Count")
# adding extra column
sim_Cp <- sim_Cp %>%
    mutate(Origin = "C_parvum")
# Calculate Relative abundance of counts for each genome
sim_Cp <- sim_Cp %>%
```

```r
    group_by(Origin) %>%
    mutate(relative_ab = Count/sum(Count))

# Cryptosporidium parvum
sim_CpATCC <- read.table("~/crypto_data/4/C_parvum_IOWA-ATCC_largespecies_ct.txt"
    header = F, sep = "\t", stringsAsFactors = F)
colnames(sim_CpATCC) <- c("Species", "Locus", "Count")
# adding extra column
sim_CpATCC <- sim_CpATCC %>%
    mutate(Origin = "C_parvum_ATCC")
# Calculate Relative abundance of counts for each genome
sim_CpATCC <- sim_CpATCC %>%
    group_by(Origin) %>%
    mutate(relative_ab = Count/sum(Count))

# Cryptosporidium tyzzeri
sim_Ct <- read.table("~/crypto_data/4/C_tyzzeri_largespecies_ct.txt",
    header = F, sep = "\t", stringsAsFactors = F)
colnames(sim_Ct) <- c("Species", "Locus", "Count")
# adding extra column
sim_Ct <- sim_Ct %>%
    mutate(Origin = "C_tyzzeri")
# Calculate Relative abundance of counts for each genome
sim_Ct <- sim_Ct %>%
    group_by(Origin) %>%
    mutate(relative_ab = Count/sum(Count))

# Cryptosporidium ubiquitum
sim_Cu <- read.table("~/crypto_data/4/C_ubiquitum_largespecies_ct.txt",
    header = F, sep = "\t", stringsAsFactors = F)
colnames(sim_Cu) <- c("Species", "Locus", "Count")
# adding extra column
sim_Cu <- sim_Cu %>%
    mutate(Origin = "C_ubiquitum")
# Calculate Relative abundance of counts for each genome
sim_Cu <- sim_Cu %>%
    group_by(Origin) %>%
    mutate(relative_ab = Count/sum(Count))

# Cryptosporidium viatorum
sim_Cv <- read.table("~/crypto_data/4/C_viatorum_largespecies_ct.txt",
    header = F, sep = "\t", stringsAsFactors = F)
colnames(sim_Cv) <- c("Species", "Locus", "Count")
# adding extra column
sim_Cv <- sim_Cv %>%
    mutate(Origin = "C_viatorum")
# Calculate Relative abundance of counts for each genome
sim_Cv <- sim_Cv %>%
    group_by(Origin) %>%
    mutate(relative_ab = Count/sum(Count))

# Combine them all
sim_all_ten <- rbind(sim_Ca, sim_Cb, sim_Cc, sim_Ch, sim_ChUdeA,
```

```
    sim_Cm, sim_Cmu, sim_Cp, sim_CpATCC, sim_Ct, sim_Cu, sim_Cv)

# plot
p <- sim_all_ten %>%
    group_by(Origin) %>%
    arrange(Species) %>%
    plot_ly(x = ~Origin, y = ~relative_ab, color = ~Species,
        colors = col_c_ten, type = "bar") %>%
    layout(yaxis = list(title = "Relative Abundance"), barmode = "stack")
```

**NOTE:** The results from these analyses are presented in **Fig. S7A**.

**4.1.2 Species assignment based on mapping simulated reads from short fragment libraries ~200bp** Below, we import the modified files for each species and then create a fourth column, "Origin" with the name of the species for which we simulated the reads.

```
# Cryptosporidium andersoni
sim_Ca <- read.table("~/crypto_data/4/C_andersoni_shortspecies_ct.txt",
    header = F, sep = "\t", stringsAsFactors = F)
colnames(sim_Ca) <- c("Species", "Locus", "Count")
# adding extra column
sim_Ca <- sim_Ca %>%
    mutate(Origin = "C_andersoni")
# Calculate Relative abundance of counts for each genome
sim_Ca <- sim_Ca %>%
    group_by(Origin) %>%
    mutate(relative_ab = Count/sum(Count))

# Cryptosporidium baileyi
sim_Cb <- read.table("~/crypto_data/4/C_baileyi_shortspecies_ct.txt",
    header = F, sep = "\t", stringsAsFactors = F)
colnames(sim_Cb) <- c("Species", "Locus", "Count")
# adding extra column
sim_Cb <- sim_Cb %>%
    mutate(Origin = "C_baileyi")
# Calculate Relative abundance of counts for each genome
sim_Cb <- sim_Cb %>%
    group_by(Origin) %>%
    mutate(relative_ab = Count/sum(Count))

# Cryptosporidium cuniculus
sim_Cc <- read.table("~/crypto_data/4/C_cuniculus_shortspecies_ct.txt",
    header = F, sep = "\t", stringsAsFactors = F)
colnames(sim_Cc) <- c("Species", "Locus", "Count")
# adding extra column
sim_Cc <- sim_Cc %>%
    mutate(Origin = "C_cuniculus")
# Calculate Relative abundance of counts for each genome
sim_Cc <- sim_Cc %>%
    group_by(Origin) %>%
    mutate(relative_ab = Count/sum(Count))
```

```r
# Cryptosporidium hominis
sim_Ch <- read.table("~/crypto_data/4/C_hominis_30976_shortspecies_ct.txt"
    header = F, sep = "\t", stringsAsFactors = F)
colnames(sim_Ch) <- c("Species", "Locus", "Count")
# adding extra column
sim_Ch <- sim_Ch %>%
    mutate(Origin = "C_hominis 30976")
# Calculate Relative abundance of counts for each genome
sim_Ch <- sim_Ch %>%
    group_by(Origin) %>%
    mutate(relative_ab = Count/sum(Count))

# Cryptosporidium hominis UdeA
sim_ChUdeA <- read.table("~/crypto_data/4/C_hominis_UdeA01_shortspecies_ct.txt"
    header = F, sep = "\t", stringsAsFactors = F)
colnames(sim_ChUdeA) <- c("Species", "Locus", "Count")
# adding extra column
sim_ChUdeA <- sim_ChUdeA %>%
    mutate(Origin = "C_hominis_UdeA")
# Calculate Relative abundance of counts for each genome
sim_ChUdeA <- sim_ChUdeA %>%
    group_by(Origin) %>%
    mutate(relative_ab = Count/sum(Count))

# Cryptosporidium meleagridis
sim_Cm <- read.table("~/crypto_data/4/C_meleagridis_shortspecies_ct.txt"
    header = F, sep = "\t", stringsAsFactors = F)
colnames(sim_Cm) <- c("Species", "Locus", "Count")
# adding extra column
sim_Cm <- sim_Cm %>%
    mutate(Origin = "C_meleagridis")
# Calculate Relative abundance of counts for each genome
sim_Cm <- sim_Cm %>%
    group_by(Origin) %>%
    mutate(relative_ab = Count/sum(Count))

# Cryptosporidium muris
sim_Cmu <- read.table("~/crypto_data/4/C_muris_shortspecies_ct.txt",
    header = F, sep = "\t", stringsAsFactors = F)
colnames(sim_Cmu) <- c("Species", "Locus", "Count")
# adding extra column
sim_Cmu <- sim_Cmu %>%
    mutate(Origin = "C_muris")
# Calculate Relative abundance of counts for each genome
sim_Cmu <- sim_Cmu %>%
    group_by(Origin) %>%
    mutate(relative_ab = Count/sum(Count))

# Cryptosporidium parvum
sim_Cp <- read.table("~/crypto_data/4/C_parvum_shortspecies_ct.txt",
    header = F, sep = "\t", stringsAsFactors = F)
colnames(sim_Cp) <- c("Species", "Locus", "Count")
# adding extra column
```

```r
sim_Cp <- sim_Cp %>%
    mutate(Origin = "C_parvum")
# Calculate Relative abundance of counts for each genome
sim_Cp <- sim_Cp %>%
    group_by(Origin) %>%
    mutate(relative_ab = Count/sum(Count))

# Cryptosporidium parvum
sim_CpATCC <- read.table("~/crypto_data/4/C_parvum_IOWA-ATCC_shortspecies_ct.txt"
    header = F, sep = "\t", stringsAsFactors = F)
colnames(sim_CpATCC) <- c("Species", "Locus", "Count")
# adding extra column
sim_CpATCC <- sim_CpATCC %>%
    mutate(Origin = "C_parvum_ATCC")
# Calculate Relative abundance of counts for each genome
sim_CpATCC <- sim_CpATCC %>%
    group_by(Origin) %>%
    mutate(relative_ab = Count/sum(Count))

# Cryptosporidium tyzzeri
sim_Ct <- read.table("~/crypto_data/4/C_tyzzeri_shortspecies_ct.txt",
    header = F, sep = "\t", stringsAsFactors = F)
colnames(sim_Ct) <- c("Species", "Locus", "Count")
# adding extra column
sim_Ct <- sim_Ct %>%
    mutate(Origin = "C_tyzzeri")
# Calculate Relative abundance of counts for each genome
sim_Ct <- sim_Ct %>%
    group_by(Origin) %>%
    mutate(relative_ab = Count/sum(Count))

# Cryptosporidium ubiquitum
sim_Cu <- read.table("~/crypto_data/4/C_ubiquitum_shortspecies_ct.txt",
    header = F, sep = "\t", stringsAsFactors = F)
colnames(sim_Cu) <- c("Species", "Locus", "Count")
# adding extra column
sim_Cu <- sim_Cu %>%
    mutate(Origin = "C_ubiquitum")
# Calculate Relative abundance of counts for each genome
sim_Cu <- sim_Cu %>%
    group_by(Origin) %>%
    mutate(relative_ab = Count/sum(Count))

# Cryptosporidium viatorum
sim_Cv <- read.table("~/crypto_data/4/C_viatorum_shortspecies_ct.txt",
    header = F, sep = "\t", stringsAsFactors = F)
colnames(sim_Cv) <- c("Species", "Locus", "Count")
# adding extra column
sim_Cv <- sim_Cv %>%
    mutate(Origin = "C_viatorum")
# Calculate Relative abundance of counts for each genome
sim_Cv <- sim_Cv %>%
    group_by(Origin) %>%
```

```
    mutate(relative_ab = Count/sum(Count))

# Combine them all
sim_all_ten_short <- rbind(sim_Ca, sim_Cb, sim_Cc, sim_Ch, sim_ChUdeA, sim_Cm, sim_Cmu,
    sim_Cp, sim_CpATCC, sim_Ct, sim_Cu, sim_Cv)

# plot
p <- sim_all_ten_short %>%
    group_by(Origin) %>%
    arrange(Species) %>%
    plot_ly(x = ~Origin, y = ~relative_ab, color = ~Species, colors = col_c_ten,
        type = "bar") %>%
    layout(yaxis = list(title = "Relative Abundance"), barmode = "stack")
```

**NOTE:** The results from these analyses are presented in **Fig. S7B**.

When simulated single-infection reads were mapped to the ten-genomes database, differences between short and large fragments are only evident in the decrease in C. hominis from 82% for large fragments to 67% for short fragments of the correct assignment, given the mapping to C. cuniculus (Fig. S7A and S7B). When reads were mapped to the gp60 database, we found a small percentage of reads mapping, ranging from 0% to 0.26% (Table S3).

##### 4.2 Mapping simulated enrich and sequence reads to a gp60 database

We mapped each set of simulated reads from each genome against a fasta containing 65 gp60 sequences. We used similar parameters as above, except pairlen = 400 and maxindel = 5.

```
bbmap.sh in=1.fq in2=2.fq \
ref gp60_reference_database.fas/  \
maxindel=5 minid=0.80 rcs=t killbadpairs=t pairlen=400 \
rescuedist=600 apd=100 outu=sim_mapped/\$i\_unmapped.sam \
outm=sim_mapped/\$i\_mapped.sam printunmappedcount=t \
covstats=sim_mapped/\$i\.cov.txt threads=12 -Xmx180g 2>&1 \
| tee sim_mapped/\$i\.bbmap.log
```

These results are summarized in **Table S3**.

For each species, we used the FLAG count from the third column of our SAM files to extract those sequences for which only both reads mapped (FLAG: 83 and 99):

```
cat gp60_mapped.sam | grep -v "^@" | \
awk 'BEGIN{FS="\t";OFS="\t"}{if($2=="83" || $2=="99") print $0}' \
> both_mapped/gp60_mapped_both.txt
```

Then, for each species, we counted each species ID hit in our mapped file. The count text file has two columns, the first is the species ID, and the second is the counts:

```
cat gp60_mapped_both.txt | \
awk 'BEGIN{FS="\t";OFS="\t"}{OTU[$3]+=1}END{for (i in OTU) print i,OTU[i]}' \
> gp60_ct.txt
```

###### 4.2.1 Species assignment based on mapping simulated reads from large fragment libraries ~520bp

Before importing each file in Excel, we added an extra column with only the species name of the gp60 from the fasta header. Below, we then created a fourth column “Origin” with the name of the species for which we simulated the reads.

```
# Cryptosporidium baileyi
sim_large_gp60_Cb <- read.table("~/crypto_data/4/gp60/C_baileyi_largespecies_ct.txt"
    header = F, sep = "\t", stringsAsFactors = F)
colnames(sim_large_gp60_Cb) <- c("Species", "Locus", "Count")
# adding extra column
sim_large_gp60_Cb <- sim_large_gp60_Cb %>%
    mutate(Origin = "C_baileyi")
# Calculate Relative abundance of counts for each genome
sim_large_gp60_Cb <- sim_large_gp60_Cb %>%
    group_by(Origin) %>%
    mutate(relative_ab = Count/sum(Count))

# Cryptosporidium cuniculus
sim_large_gp60_Cc <- read.table("~/crypto_data/4/gp60/C_cuniculus_largespecies_ct.txt"
    header = F, sep = "\t", stringsAsFactors = F)
colnames(sim_large_gp60_Cc) <- c("Species", "Locus", "Count")
# adding extra column
sim_large_gp60_Cc <- sim_large_gp60_Cc %>%
    mutate(Origin = "C_cuniculus")
# Calculate Relative abundance of counts for each genome
sim_large_gp60_Cc <- sim_large_gp60_Cc %>%
    group_by(Origin) %>%
    mutate(relative_ab = Count/sum(Count))

# Cryptosporidium hominis
sim_large_gp60_Ch <- read.table("~/crypto_data/4/gp60/C_hominis_30976_largespecies_ct.txt"
    header = F, sep = "\t", stringsAsFactors = F)
colnames(sim_large_gp60_Ch) <- c("Species", "Locus", "Count")
# adding extra column
sim_large_gp60_Ch <- sim_large_gp60_Ch %>%
    mutate(Origin = "C_hominis_30976")
# Calculate Relative abundance of counts for each genome
sim_large_gp60_Ch <- sim_large_gp60_Ch %>%
    group_by(Origin) %>%
    mutate(relative_ab = Count/sum(Count))

# Cryptosporidium hominis UdeA
sim_large_gp60_ChUdeA <- read.table("~/crypto_data/4/gp60/C_hominis_UdeA01_largespecies_ct.txt"
    header = F, sep = "\t", stringsAsFactors = F)
colnames(sim_large_gp60_ChUdeA) <- c("Species", "Locus", "Count")
# adding extra column
sim_large_gp60_ChUdeA <- sim_large_gp60_ChUdeA %>%
    mutate(Origin = "C_hominis_UdeA")
# Calculate Relative abundance of counts for each genome
sim_large_gp60_ChUdeA <- sim_large_gp60_ChUdeA %>%
    group_by(Origin) %>%
    mutate(relative_ab = Count/sum(Count))

# Cryptosporidium meleagridis
```

```r
sim_large_gp60_Cm <- read.table("~/crypto_data/4/gp60/C_meleagridis_largespecies_ct.txt"
    header = F, sep = "\t", stringsAsFactors = F)
colnames(sim_large_gp60_Cm) <- c("Species", "Locus", "Count")
# adding extra column
sim_large_gp60_Cm <- sim_large_gp60_Cm %>%
    mutate(Origin = "C_meleagridis")
# Calculate Relative abundance of counts for each genome
sim_large_gp60_Cm <- sim_large_gp60_Cm %>%
    group_by(Origin) %>%
    mutate(relative_ab = Count/sum(Count))

# Cryptosporidium muris
sim_large_gp60_Cmu <- read.table("~/crypto_data/4/gp60/C_muris_largespecies_ct.txt"
    header = F, sep = "\t", stringsAsFactors = F)
colnames(sim_large_gp60_Cmu) <- c("Species", "Locus", "Count")
# adding extra column
sim_large_gp60_Cmu <- sim_large_gp60_Cmu %>%
    mutate(Origin = "C_muris")
# Calculate Relative abundance of counts for each genome
sim_large_gp60_Cmu <- sim_large_gp60_Cmu %>%
    group_by(Origin) %>%
    mutate(relative_ab = Count/sum(Count))

# Cryptosporidium parvum
sim_large_gp60_Cp <- read.table("~/crypto_data/4/gp60/C_parvum_largespecies_ct.txt"
    header = F, sep = "\t", stringsAsFactors = F)
colnames(sim_large_gp60_Cp) <- c("Species", "Locus", "Count")
# adding extra column
sim_large_gp60_Cp <- sim_large_gp60_Cp %>%
    mutate(Origin = "C_parvum")
# Calculate Relative abundance of counts for each genome
sim_large_gp60_Cp <- sim_large_gp60_Cp %>%
    group_by(Origin) %>%
    mutate(relative_ab = Count/sum(Count))

# Cryptosporidium parvum
sim_large_gp60_CpATCC <- read.table("~/crypto_data/4/gp60/C_parvum_IOWA-ATCC_largespecies_ct.txt"
    header = F, sep = "\t", stringsAsFactors = F)
colnames(sim_large_gp60_CpATCC) <- c("Species", "Locus", "Count")
# adding extra column
sim_large_gp60_CpATCC <- sim_large_gp60_CpATCC %>%
    mutate(Origin = "C_parvum_ATCC")
# Calculate Relative abundance of counts for each genome
sim_large_gp60_CpATCC <- sim_large_gp60_CpATCC %>%
    group_by(Origin) %>%
    mutate(relative_ab = Count/sum(Count))

# Cryptosporidium tyzzeri
sim_large_gp60_Ct <- read.table("~/crypto_data/4/gp60/C_tyzzeri_largespecies_ct.txt"
    header = F, sep = "\t", stringsAsFactors = F)
colnames(sim_large_gp60_Ct) <- c("Species", "Locus", "Count")
# adding extra column
sim_large_gp60_Ct <- sim_large_gp60_Ct %>%
```

```
    mutate(Origin = "C_tyzzeri")
# Calculate Relative abundance of counts for each genome
sim_large_gp60_Ct <- sim_large_gp60_Ct %>%
    group_by(Origin) %>%
    mutate(relative_ab = Count/sum(Count))

# Cryptosporidium ubiquitum
sim_large_gp60_Cu <- read.table("~/crypto_data/4/gp60/C_ubiquitum_largespecies_ct.txt"
    header = F, sep = "\t", stringsAsFactors = F)
colnames(sim_large_gp60_Cu) <- c("Species", "Locus", "Count")
# adding extra column
sim_large_gp60_Cu <- sim_large_gp60_Cu %>%
    mutate(Origin = "C_ubiquitum")
# Calculate Relative abundance of counts for each genome
sim_large_gp60_Cu <- sim_large_gp60_Cu %>%
    group_by(Origin) %>%
    mutate(relative_ab = Count/sum(Count))

# Combine them all
sim_large_all_gp60 <- rbind(sim_large_gp60_Cb, sim_large_gp60_Cc,
    sim_large_gp60_Ch, sim_large_gp60_ChUdeA, sim_large_gp60_Cm,
    sim_large_gp60_Cmu, sim_large_gp60_Cp, sim_large_gp60_CpATCC,
    sim_large_gp60_Ct, sim_large_gp60_Cu)
```

Plot relative abundance of hits from simulated reads of ~520 bp fragments against the 65 records from the GP60 database for each set of reference genomes.

```
# Setting colors for specific species
col_c_cuniculus <- c("#bebada")
col_c_erinaceai <- c("#8e0152")
col_c_fayeri <- c("#ffed6f")
col_c_hominis <- c("#fb8072")
col_c_meleagridis <- c("#80b1d3")
col_c_parvum <- c("#b3de69")
col_c_tyzzeri <- c("#fccde5")
col_c_ubiquitum <- c("#d9d9d9")
col_c_gp60 <- c("#bebada", "#8e0152", "#ffed6f", "#fb8072", "#80b1d3", "#b3de69",
    "#fccde5", "#d9d9d9")

p <- sim_large_all_gp60 %>%
    group_by(Origin) %>%
    arrange(Species) %>%
    plot_ly(x = ~Origin, y = ~relative_ab, color = ~Species, colors = col_c_gp60,
        type = "bar") %>%
    layout(yaxis = list(title = "Relative Abundance"), barmode = "stack")
```

**NOTE:** The results from these analyses are presented in **Fig. 7C**..

Estimating Relative Abundance

```
summary_gp60_asis_large <- sim_large_all_gp60 %>%
    group_by(Origin, Species) %>%
    summarise(Size_Sum = sum(relative_ab))
```

```
## `summarise()` has grouped output by 'Origin'. You can override using the
## `.groups` argument.
```

```r
# summary_gp60_asis_large
```

**NOTE:** The results from these analyses are presented in **Fig. S7C**.

###### 4.2.2 Species assignment based on mapping simulated reads from short fragment libraries ~200bp

Similarly, before importing each file, we added an extra column in Excel with only the species name of the gp60 from the FASTA header. Below, we then create a fourth column, "Origin" with the name of the species for which we simulated the reads.

```r
# Cryptosporidium baileyi
sim_short_gp60_Cb <- read.table("~/crypto_data/4/gp60/C_baileyi_shortspecies_ct.txt"
    header = F, sep = "\t", stringsAsFactors = F)
colnames(sim_short_gp60_Cb) <- c("Species", "Locus", "Count")
# adding extra column
sim_short_gp60_Cb <- sim_short_gp60_Cb %>%
    mutate(Origin = "C_baileyi")
# Calculate Relative abundance of counts for each genome
sim_short_gp60_Cb <- sim_short_gp60_Cb %>%
    group_by(Origin) %>%
    mutate(relative_ab = Count/sum(Count))

# Cryptosporidium cuniculus
sim_short_gp60_Cc <- read.table("~/crypto_data/4/gp60/C_cuniculus_shortspecies_ct.txt"
    header = F, sep = "\t", stringsAsFactors = F)
colnames(sim_short_gp60_Cc) <- c("Species", "Locus", "Count")
# adding extra column
sim_short_gp60_Cc <- sim_short_gp60_Cc %>%
    mutate(Origin = "C_cuniculus")
# Calculate Relative abundance of counts for each genome
sim_short_gp60_Cc <- sim_short_gp60_Cc %>%
    group_by(Origin) %>%
    mutate(relative_ab = Count/sum(Count))

# Cryptosporidium hominis
sim_short_gp60_Ch <- read.table("~/crypto_data/4/gp60/C_hominis_30976_shortspecies_ct.txt"
    header = F, sep = "\t", stringsAsFactors = F)
colnames(sim_short_gp60_Ch) <- c("Species", "Locus", "Count")
# adding extra column
sim_short_gp60_Ch <- sim_short_gp60_Ch %>%
    mutate(Origin = "C_hominis_30976")
# Calculate Relative abundance of counts for each genome
sim_short_gp60_Ch <- sim_short_gp60_Ch %>%
    group_by(Origin) %>%
    mutate(relative_ab = Count/sum(Count))

# Cryptosporidium hominis UdeA
sim_short_gp60_ChUdeA <- read.table("~/crypto_data/4/gp60/C_hominis_UdeA01_shortspecies_ct.txt"
    header = F, sep = "\t", stringsAsFactors = F)
colnames(sim_short_gp60_ChUdeA) <- c("Species", "Locus", "Count")
# adding extra column
```

```r
sim_short_gp60_ChUdeA <- sim_short_gp60_ChUdeA %>%
    mutate(Origin = "C_hominis_UdeA")
# Calculate Relative abundance of counts for each genome
sim_short_gp60_ChUdeA <- sim_short_gp60_ChUdeA %>%
    group_by(Origin) %>%
    mutate(relative_ab = Count/sum(Count))

# Cryptosporidium meleagridis
sim_short_gp60_Cm <- read.table("~/crypto_data/4/gp60/C_meleagridis_shortspecies_ct.txt"
    header = F, sep = "\t", stringsAsFactors = F)
colnames(sim_short_gp60_Cm) <- c("Species", "Locus", "Count")
# adding extra column
sim_short_gp60_Cm <- sim_short_gp60_Cm %>%
    mutate(Origin = "C_meleagridis")
# Calculate Relative abundance of counts for each genome
sim_short_gp60_Cm <- sim_short_gp60_Cm %>%
    group_by(Origin) %>%
    mutate(relative_ab = Count/sum(Count))

# Cryptosporidium muris
sim_short_gp60_Cmu <- read.table("~/crypto_data/4/gp60/C_muris_shortspecies_ct.txt"
    header = F, sep = "\t", stringsAsFactors = F)
colnames(sim_short_gp60_Cmu) <- c("Species", "Locus", "Count")
# adding extra column
sim_short_gp60_Cmu <- sim_short_gp60_Cmu %>%
    mutate(Origin = "C_muris")
# Calculate Relative abundance of counts for each genome
sim_short_gp60_Cmu <- sim_short_gp60_Cmu %>%
    group_by(Origin) %>%
    mutate(relative_ab = Count/sum(Count))

# Cryptosporidium parvum
sim_short_gp60_Cp <- read.table("~/crypto_data/4/gp60/C_parvum_shortspecies_ct.txt"
    header = F, sep = "\t", stringsAsFactors = F)
colnames(sim_short_gp60_Cp) <- c("Species", "Locus", "Count")
# adding extra column
sim_short_gp60_Cp <- sim_short_gp60_Cp %>%
    mutate(Origin = "C_parvum")
# Calculate Relative abundance of counts for each genome
sim_short_gp60_Cp <- sim_short_gp60_Cp %>%
    group_by(Origin) %>%
    mutate(relative_ab = Count/sum(Count))

# Cryptosporidium parvum
sim_short_gp60_CpATCC <- read.table("~/crypto_data/4/gp60/C_parvum_IOWA-ATCC_shortspecies_ct.txt"
    header = F, sep = "\t", stringsAsFactors = F)
colnames(sim_short_gp60_CpATCC) <- c("Species", "Locus", "Count")
# adding extra column
sim_short_gp60_CpATCC <- sim_short_gp60_CpATCC %>%
    mutate(Origin = "C_parvum_ATCC")
# Calculate Relative abundance of counts for each genome
sim_short_gp60_CpATCC <- sim_short_gp60_CpATCC %>%
    group_by(Origin) %>%
```

```
    mutate(relative_ab = Count/sum(Count))

# Cryptosporidium tyzzeri
sim_short_gp60_Ct <- read.table("~/crypto_data/4/gp60/C_tyzzeri_shortspecies_ct.txt"
    header = F, sep = "\t", stringsAsFactors = F)
colnames(sim_short_gp60_Ct) <- c("Species", "Locus", "Count")
# adding extra column
sim_short_gp60_Ct <- sim_short_gp60_Ct %>%
    mutate(Origin = "C_tyzzeri")
# Calculate Relative abundance of counts for each genome
sim_short_gp60_Ct <- sim_short_gp60_Ct %>%
    group_by(Origin) %>%
    mutate(relative_ab = Count/sum(Count))

# Cryptosporidium ubiquitum
sim_short_gp60_Cu <- read.table("~/crypto_data/4/gp60/C_ubiquitum_shortspecies_ct.txt"
    header = F, sep = "\t", stringsAsFactors = F)
colnames(sim_short_gp60_Cu) <- c("Species", "Locus", "Count")
# adding extra column
sim_short_gp60_Cu <- sim_short_gp60_Cu %>%
    mutate(Origin = "C_ubiquitum")
# Calculate Relative abundance of counts for each genome
sim_short_gp60_Cu <- sim_short_gp60_Cu %>%
    group_by(Origin) %>%
    mutate(relative_ab = Count/sum(Count))

# Cryptosporidium viatorum
sim_short_gp60_Cv <- read.table("~/crypto_data/4/gp60/C_viatorum_shortspecies_ct.txt"
    header = F, sep = "\t", stringsAsFactors = F)
colnames(sim_short_gp60_Cv) <- c("Species", "Locus", "Count")
# adding extra column
sim_short_gp60_Cv <- sim_short_gp60_Cv %>%
    mutate(Origin = "C_viatorum")
# Calculate Relative abundance of counts for each genome
sim_short_gp60_Cv <- sim_short_gp60_Cv %>%
    group_by(Origin) %>%
    mutate(relative_ab = Count/sum(Count))

# Combine them all
sim_short_all_gp60 <- rbind(sim_short_gp60_Cb, sim_short_gp60_Cc,
    sim_short_gp60_Ch, sim_short_gp60_ChUdeA, sim_short_gp60_Cm,
    sim_short_gp60_Cmu, sim_short_gp60_Cp, sim_short_gp60_CpATCC,
    sim_short_gp60_Ct, sim_short_gp60_Cu, sim_short_gp60_Cv)
```

Plot relative abundance of hits from simulated reads of ~200 bp fragments against the 65 records from the GP60 database for each set of reference genomes.

```
p <- sim_short_all_gp60 %>%
    group_by(Origin) %>%
    arrange(Species) %>%
    plot_ly(x = ~Origin, y = ~relative_ab, color = ~Species, colors = col_c_gp60,
        type = "bar") %>%
    layout(yaxis = list(title = "Relative Abundance"), barmode = "stack")
```

**NOTE:** The results from these analyses are presented in **Fig. S7D**.

Estimating Relative Abundance.

```
summary_gp60_asis_short <- sim_short_all_gp60 %>%
    group_by(Origin, Species) %>%
    summarise(Size_Sum = sum(relative_ab))
```

```
## `summarise()` has grouped output by 'Origin'. You can override using the
## `.groups` argument.
```

```
# summary_gp60_asis_short
```

Regarding species assignment (Fig. S7C and S7D), *C. andersoni* and *C. viatorum* reads simulated from large fragments do not map to the database. Still, short fragments of *C. viatorum* are assigned to *C. fayeri*. Also, *C. baileyi* and *C. muris* are assigned high proportions to *C. erinacei* for both fragment sizes. *C. tyzzeri* shows a proper assignment proportion of 55% when large fragments are considered, and it decreases to 34% with small fragments, these being assigned to *C. parvum* with a high proportion instead. *C. hominis*, *C. meleagridis*, and *C. parvum* show a high proportion of proper species assignment (>62%) for large fragment sizes. However, this number decreases for short fragments. *C. ubiquitum* samples show proper assignment for both fragment sizes. Overall, at least 99.1% of reads for each sample mapped to the corresponding genome, allowing high accuracy for species assignment for all *Cryptosporidium* species tested, including those not considered in the bait design (Table S2).

##### 4.3 Mapping simulated enrich and sequence reads to CryptoDB 18S rRNA database

We mapped simulated reads using similar parameters as above, except pairlen = 500 and maxindel = 5, and using as references independently a fasta file that has 57 reference sequences and was obtained from: https://cryptodb.org/cryptodb/app/downloads/Current_Release/CryptosporidiidaeReference/fasta/ (version last modified on 2022-04-13 11:11), here named as 18S_CDB.

Species identification in *Cryptosporidium* has typically been assessed by amplification and sequencing of the 18S rRNA gene. This practice is rooted in i) difficult access to whole genome data from clinical/environmental samples, ii) the presence of conserved regions amenable to universal primer design for this gene, iii) the presence of variable regions capable of distinguishing between species in this gene, and iv) in Cryptosporidium multiple copies of the gene are present, which enhances the sensitivity of the detection and aids in the resolution of closely related species (Turner et al., 2025).

```
bbmap.sh in=1.fq in2=2.fq \
ref 18S_reference.fasta  \
maxindel=5 minid=0.80 rcs=t killbadpairs=t pairlen=400 \
rescuedist=600 apd=100 outu=sim_mapped/\$i\_unmapped.sam \
outm=sim_mapped/\$i\_mapped.sam printunmappedcount=t \
covstats=sim_mapped/\$i\.cov.txt threads=12 -Xmx180g 2>&1 \
| tee sim_mapped/\$i\.bbmap.log
```

These results are summarized in **Table S4**.

For each species, we used the FLAG count from the third column of our SAM files to extract those sequences for which only both reads mapped (FLAG: 83 and 99):

```
cat 18S_mapped.sam | grep -v "^@" | \
awk 'BEGIN{FS="\t";OFS="\t"}{if($2=="83" || $2=="99") print $0}' \
> both_mapped/18S_mapped_both.txt
```

Then, for each species, we count the number of species IDs that hit our mapped file. The count text has two columns, the first is the species ID, and the second is the counts:

```
cat 18S_mapped_both.txt | \
awk 'BEGIN{FS="\t";OFS="\t"}{OTU[$3]+=1}END{for (i in OTU) print i,OTU[i]}' \
> 18S_ct.txt
```

**4.3.1 Species assignment based on mapping simulated reads from large fragment libraries ~520bp** Similarly, before importing each file, we added an extra column in Excel with only the species name of the 18S rRNA from the FASTA header. Below, we then create a fourth column, "Origin" with the name of the species for which we simulated the reads.

```
# Cryptosporidium andersoni
sim_large_18S_Ca <- read.table("~/crypto_data/4/18S_rRNA/C_andersoni_largespecies_ct.txt"
    header = F, sep = "\t", stringsAsFactors = F)
colnames(sim_large_18S_Ca) <- c("Species", "Locus", "Count")
# adding extra column
sim_large_18S_Ca <- sim_large_18S_Ca %>%
    mutate(Origin = "C_andersoni")
# Calculate Relative abundance of counts for each genome
sim_large_18S_Ca <- sim_large_18S_Ca %>%
    group_by(Origin) %>%
    mutate(relative_ab = Count/sum(Count))

# Cryptosporidium baileyi
sim_large_18S_Cb <- read.table("~/crypto_data/4/18S_rRNA/C_baileyi_largespecies_ct.txt"
    header = F, sep = "\t", stringsAsFactors = F)
colnames(sim_large_18S_Cb) <- c("Species", "Locus", "Count")
# adding extra column
sim_large_18S_Cb <- sim_large_18S_Cb %>%
    mutate(Origin = "C_baileyi")
# Calculate Relative abundance of counts for each genome
sim_large_18S_Cb <- sim_large_18S_Cb %>%
    group_by(Origin) %>%
    mutate(relative_ab = Count/sum(Count))

# Cryptosporidium cuniculus
sim_large_18S_Cc <- read.table("~/crypto_data/4/18S_rRNA/C_cuniculus_largespecies_ct.txt"
    header = F, sep = "\t", stringsAsFactors = F)
colnames(sim_large_18S_Cc) <- c("Species", "Locus", "Count")
# adding extra column
sim_large_18S_Cc <- sim_large_18S_Cc %>%
    mutate(Origin = "C_cuniculus")
# Calculate Relative abundance of counts for each genome
sim_large_18S_Cc <- sim_large_18S_Cc %>%
    group_by(Origin) %>%
    mutate(relative_ab = Count/sum(Count))
```

```r
# Cryptosporidium hominis
sim_large_18S_Ch <- read.table("~/crypto_data/4/18S_rRNA/C_hominis_30976_largespecies_ct.txt"
    header = F, sep = "\t", stringsAsFactors = F)
colnames(sim_large_18S_Ch) <- c("Species", "Locus", "Count")
# adding extra column
sim_large_18S_Ch <- sim_large_18S_Ch %>%
    mutate(Origin = "C_hominis_30976")
# Calculate Relative abundance of counts for each genome
sim_large_18S_Ch <- sim_large_18S_Ch %>%
    group_by(Origin) %>%
    mutate(relative_ab = Count/sum(Count))

# Cryptosporidium hominis UdeA
sim_large_18S_ChUdeA <- read.table("~/crypto_data/4/18S_rRNA/C_hominis_UdeA01_largespecies_ct.txt"
    header = F, sep = "\t", stringsAsFactors = F)
colnames(sim_large_18S_ChUdeA) <- c("Species", "Locus", "Count")
# adding extra column
sim_large_18S_ChUdeA <- sim_large_18S_ChUdeA %>%
    mutate(Origin = "C_hominis_UdeA")
# Calculate Relative abundance of counts for each genome
sim_large_18S_ChUdeA <- sim_large_18S_ChUdeA %>%
    group_by(Origin) %>%
    mutate(relative_ab = Count/sum(Count))

# Cryptosporidium meleagridis
sim_large_18S_Cm <- read.table("~/crypto_data/4/18S_rRNA/C_meleagridis_largespecies_ct.txt"
    header = F, sep = "\t", stringsAsFactors = F)
colnames(sim_large_18S_Cm) <- c("Species", "Locus", "Count")
# adding extra column
sim_large_18S_Cm <- sim_large_18S_Cm %>%
    mutate(Origin = "C_meleagridis")
# Calculate Relative abundance of counts for each genome
sim_large_18S_Cm <- sim_large_18S_Cm %>%
    group_by(Origin) %>%
    mutate(relative_ab = Count/sum(Count))

# Cryptosporidium muris
sim_large_18S_Cmu <- read.table("~/crypto_data/4/18S_rRNA/C_muris_largespecies_ct.txt"
    header = F, sep = "\t", stringsAsFactors = F)
colnames(sim_large_18S_Cmu) <- c("Species", "Locus", "Count")
# adding extra column
sim_large_18S_Cmu <- sim_large_18S_Cmu %>%
    mutate(Origin = "C_muris")
# Calculate Relative abundance of counts for each genome
sim_large_18S_Cmu <- sim_large_18S_Cmu %>%
    group_by(Origin) %>%
    mutate(relative_ab = Count/sum(Count))

# Cryptosporidium parvum
sim_large_18S_Cp <- read.table("~/crypto_data/4/18S_rRNA/C_parvum_largespecies_ct.txt"
    header = F, sep = "\t", stringsAsFactors = F)
colnames(sim_large_18S_Cp) <- c("Species", "Locus", "Count")
# adding extra column
```

```r
sim_large_18S_Cp <- sim_large_18S_Cp %>%
    mutate(Origin = "C_parvum")
# Calculate Relative abundance of counts for each genome
sim_large_18S_Cp <- sim_large_18S_Cp %>%
    group_by(Origin) %>%
    mutate(relative_ab = Count/sum(Count))

# Cryptosporidium parvum
sim_large_18S_CpATCC <- read.table("~/crypto_data/4/18S_rRNA/C_parvum_IOWA-ATCC_largespecies_ct.txt"
    header = F, sep = "\t", stringsAsFactors = F)
colnames(sim_large_18S_CpATCC) <- c("Species", "Locus", "Count")
# adding extra column
sim_large_18S_CpATCC <- sim_large_18S_CpATCC %>%
    mutate(Origin = "C_parvum_ATCC")
# Calculate Relative abundance of counts for each genome
sim_large_18S_CpATCC <- sim_large_18S_CpATCC %>%
    group_by(Origin) %>%
    mutate(relative_ab = Count/sum(Count))

# Cryptosporidium tyzzeri
sim_large_18S_Ct <- read.table("~/crypto_data/4/18S_rRNA/C_tyzzeri_largespecies_ct.txt"
    header = F, sep = "\t", stringsAsFactors = F)
colnames(sim_large_18S_Ct) <- c("Species", "Locus", "Count")
# adding extra column
sim_large_18S_Ct <- sim_large_18S_Ct %>%
    mutate(Origin = "C_tyzzeri")
# Calculate Relative abundance of counts for each genome
sim_large_18S_Ct <- sim_large_18S_Ct %>%
    group_by(Origin) %>%
    mutate(relative_ab = Count/sum(Count))

# Cryptosporidium ubiquitum
sim_large_18S_Cu <- read.table("~/crypto_data/4/18S_rRNA/C_ubiquitum_largespecies_ct.txt"
    header = F, sep = "\t", stringsAsFactors = F)
colnames(sim_large_18S_Cu) <- c("Species", "Locus", "Count")
# adding extra column
sim_large_18S_Cu <- sim_large_18S_Cu %>%
    mutate(Origin = "C_ubiquitum")
# Calculate Relative abundance of counts for each genome
sim_large_18S_Cu <- sim_large_18S_Cu %>%
    group_by(Origin) %>%
    mutate(relative_ab = Count/sum(Count))

# Cryptosporidium viatorum
sim_large_18S_Cv <- read.table("~/crypto_data/4/18S_rRNA/C_viatorum_largespecies_ct.txt"
    header = F, sep = "\t", stringsAsFactors = F)
colnames(sim_large_18S_Cv) <- c("Species", "Locus", "Count")
# adding extra column
sim_large_18S_Cv <- sim_large_18S_Cv %>%
    mutate(Origin = "C_viatorum")
# Calculate Relative abundance of counts for each genome
sim_large_18S_Cv <- sim_large_18S_Cv %>%
    group_by(Origin) %>%
```

```r
    mutate(relative_ab = Count/sum(Count))

# Combine them all
sim_large_all_18S <- rbind(sim_large_18S_Ca, sim_large_18S_Cb,
    sim_large_18S_Cc, sim_large_18S_Ch, sim_large_18S_ChUdeA,
    sim_large_18S_Cm, sim_large_18S_Cmu, sim_large_18S_Cp, sim_large_18S_CpATCC,
    sim_large_18S_Ct, sim_large_18S_Cu, sim_large_18S_Cv)
# sim_large_all_18S save(sim_all_18S, file
# ='simulated_18S_CDB_feature_table.Rdata')
```

Plot relative abundance of hits from simulated reads of ~520 bp fragments against the 18S reference database Crypto Database.

```r
col_c_andersoni <- c("#8dd3c7")
col_c_baileyi <- c("#ffffb3")
col_c_bovis <- c("#1f78b4")
col_c_canis <- c("#33a02c")
col_c_environmental <- c("#e31a1c")
col_c_meleagridis <- c("#80b1d3")
col_c_muris <- c("#fdb462")
col_c_parvum <- c("#b3de69")
col_c_serpentis <- c("#1a1a1a")
col_c_sp <- c("#6a3d9a")
col_c_varanii <- c("#b15928")
col_c_18S <- c("#8dd3c7", "#ffffb3", "#1f78b4", "#33a02c", "#e31a1c", "#80b1d3",
    "#fdb462", "#b3de69", "#1a1a1a", "#6a3d9a", "#b15928")

p <- sim_large_all_18S %>%
    group_by(Origin) %>%
    arrange(Species) %>%
    plot_ly(x = ~Origin, y = ~relative_ab, color = ~Species, colors = col_c_18S,
        type = "bar") %>%
    layout(yaxis = list(title = "Relative Abundance"), barmode = "stack")
```

**NOTE:** The results from these analyses are presented in **Fig. S7E**.

Estimating Relative Abundance

```r
summary_18S_CryptoDB_large <- sim_large_all_18S %>%
    group_by(Origin, Species) %>%
    summarise(Size_Sum = sum(relative_ab))
```

```
## `summarise()` has grouped output by 'Origin'. You can override using the
## `.groups` argument.
```

```r
# summary_18S_CryptoDB_large
```

**4.3.2 Species assignment based on mapping simulated reads from short fragment libraries ~200bp** Similarly, before importing each file, we added an extra column in Excel with only the species name of the 18S from the FASTA header. Below, we then create a fourth column, "Origin," with the name of the species for which we simulated the reads.

```r
# Cryptosporidium andersoni
sim_short_18S_Ca <- read.table("~/crypto_data/4/18S_rRNA/C_andersoni_shortspecies_ct.txt"
    header = F, sep = "\t", stringsAsFactors = F)
colnames(sim_short_18S_Ca) <- c("Species", "Locus", "Count")
# adding extra column
sim_short_18S_Ca <- sim_short_18S_Ca %>%
    mutate(Origin = "C_andersoni")
# Calculate Relative abundance of counts for each genome
sim_short_18S_Ca <- sim_short_18S_Ca %>%
    group_by(Origin) %>%
    mutate(relative_ab = Count/sum(Count))

# Cryptosporidium baileyi
sim_short_18S_Cb <- read.table("~/crypto_data/4/18S_rRNA/C_baileyi_shortspecies_ct.txt"
    header = F, sep = "\t", stringsAsFactors = F)
colnames(sim_short_18S_Cb) <- c("Species", "Locus", "Count")
# adding extra column
sim_short_18S_Cb <- sim_short_18S_Cb %>%
    mutate(Origin = "C_baileyi")
# Calculate Relative abundance of counts for each genome
sim_short_18S_Cb <- sim_short_18S_Cb %>%
    group_by(Origin) %>%
    mutate(relative_ab = Count/sum(Count))

# Cryptosporidium cuniculus
sim_short_18S_Cc <- read.table("~/crypto_data/4/18S_rRNA/C_cuniculus_shortspecies_ct.txt"
    header = F, sep = "\t", stringsAsFactors = F)
colnames(sim_short_18S_Cc) <- c("Species", "Locus", "Count")
# adding extra column
sim_short_18S_Cc <- sim_short_18S_Cc %>%
    mutate(Origin = "C_cuniculus")
# Calculate Relative abundance of counts for each genome
sim_short_18S_Cc <- sim_short_18S_Cc %>%
    group_by(Origin) %>%
    mutate(relative_ab = Count/sum(Count))

# Cryptosporidium hominis
sim_short_18S_Ch <- read.table("~/crypto_data/4/18S_rRNA/C_hominis_30976_shortspecies_ct.txt"
    header = F, sep = "\t", stringsAsFactors = F)
colnames(sim_short_18S_Ch) <- c("Species", "Locus", "Count")
# adding extra column
sim_short_18S_Ch <- sim_short_18S_Ch %>%
    mutate(Origin = "C_hominis_30976")
# Calculate Relative abundance of counts for each genome
sim_short_18S_Ch <- sim_short_18S_Ch %>%
    group_by(Origin) %>%
    mutate(relative_ab = Count/sum(Count))

# Cryptosporidium hominis UdeA
sim_short_18S_ChUdeA <- read.table("~/crypto_data/4/18S_rRNA/C_hominis_UdeA01_shortspecies_ct.txt"
    header = F, sep = "\t", stringsAsFactors = F)
colnames(sim_short_18S_ChUdeA) <- c("Species", "Locus", "Count")
# adding extra column
```

```r
sim_short_18S_ChUdeA <- sim_short_18S_ChUdeA %>%
    mutate(Origin = "C_hominis_UdeA")
# Calculate Relative abundance of counts for each genome
sim_short_18S_ChUdeA <- sim_short_18S_ChUdeA %>%
    group_by(Origin) %>%
    mutate(relative_ab = Count/sum(Count))

# Cryptosporidium meleagridis
sim_short_18S_Cm <- read.table("~/crypto_data/4/18S_rRNA/C_meleagridis_shortspecies_ct.txt"
    header = F, sep = "\t", stringsAsFactors = F)
colnames(sim_short_18S_Cm) <- c("Species", "Locus", "Count")
# adding extra column
sim_short_18S_Cm <- sim_short_18S_Cm %>%
    mutate(Origin = "C_meleagridis")
# Calculate Relative abundance of counts for each genome
sim_short_18S_Cm <- sim_short_18S_Cm %>%
    group_by(Origin) %>%
    mutate(relative_ab = Count/sum(Count))

# Cryptosporidium muris
sim_short_18S_Cmu <- read.table("~/crypto_data/4/18S_rRNA/C_muris_shortspecies_ct.txt"
    header = F, sep = "\t", stringsAsFactors = F)
colnames(sim_short_18S_Cmu) <- c("Species", "Locus", "Count")
# adding extra column
sim_short_18S_Cmu <- sim_short_18S_Cmu %>%
    mutate(Origin = "C_muris")
# Calculate Relative abundance of counts for each genome
sim_short_18S_Cmu <- sim_short_18S_Cmu %>%
    group_by(Origin) %>%
    mutate(relative_ab = Count/sum(Count))

# Cryptosporidium parvum
sim_short_18S_Cp <- read.table("~/crypto_data/4/18S_rRNA/C_parvum_shortspecies_ct.txt"
    header = F, sep = "\t", stringsAsFactors = F)
colnames(sim_short_18S_Cp) <- c("Species", "Locus", "Count")
# adding extra column
sim_short_18S_Cp <- sim_short_18S_Cp %>%
    mutate(Origin = "C_parvum")
# Calculate Relative abundance of counts for each genome
sim_short_18S_Cp <- sim_short_18S_Cp %>%
    group_by(Origin) %>%
    mutate(relative_ab = Count/sum(Count))

# Cryptosporidium parvum
sim_short_18S_CpATCC <- read.table("~/crypto_data/4/18S_rRNA/C_parvum_IOWA-ATCC_shortspecies_ct.txt"
    header = F, sep = "\t", stringsAsFactors = F)
colnames(sim_short_18S_CpATCC) <- c("Species", "Locus", "Count")
# adding extra column
sim_short_18S_CpATCC <- sim_short_18S_CpATCC %>%
    mutate(Origin = "C_parvum_ATCC")
# Calculate Relative abundance of counts for each genome
sim_short_18S_CpATCC <- sim_short_18S_CpATCC %>%
    group_by(Origin) %>%
```

```
    mutate(relative_ab = Count/sum(Count))

# Cryptosporidium tyzzeri
sim_short_18S_Ct <- read.table("~/crypto_data/4/18S_rRNA/C_tyzzeri_shortspecies_ct.txt"
    header = F, sep = "\t", stringsAsFactors = F)
colnames(sim_short_18S_Ct) <- c("Species", "Locus", "Count")
# adding extra column
sim_short_18S_Ct <- sim_short_18S_Ct %>%
    mutate(Origin = "C_tyzzeri")
# Calculate Relative abundance of counts for each genome
sim_short_18S_Ct <- sim_short_18S_Ct %>%
    group_by(Origin) %>%
    mutate(relative_ab = Count/sum(Count))

# Cryptosporidium ubiquitum
sim_short_18S_Cu <- read.table("~/crypto_data/4/18S_rRNA/C_ubiquitum_shortspecies_ct.txt"
    header = F, sep = "\t", stringsAsFactors = F)
colnames(sim_short_18S_Cu) <- c("Species", "Locus", "Count")
# adding extra column
sim_short_18S_Cu <- sim_short_18S_Cu %>%
    mutate(Origin = "C_ubiquitum")
# Calculate Relative abundance of counts for each genome
sim_short_18S_Cu <- sim_short_18S_Cu %>%
    group_by(Origin) %>%
    mutate(relative_ab = Count/sum(Count))

# Cryptosporidium viatorum
sim_short_18S_Cv <- read.table("~/crypto_data/4/18S_rRNA/C_viatorum_shortspecies_ct.txt"
    header = F, sep = "\t", stringsAsFactors = F)
colnames(sim_short_18S_Cv) <- c("Species", "Locus", "Count")
# adding extra column
sim_short_18S_Cv <- sim_short_18S_Cv %>%
    mutate(Origin = "C_viatorum")
# Calculate Relative abundance of counts for each genome
sim_short_18S_Cv <- sim_short_18S_Cv %>%
    group_by(Origin) %>%
    mutate(relative_ab = Count/sum(Count))

# Combine them all
sim_short_all_18S <- rbind(sim_short_18S_Ca, sim_short_18S_Cb,
    sim_short_18S_Cc, sim_short_18S_Ch, sim_short_18S_ChUdeA,
    sim_short_18S_Cm, sim_short_18S_Cmu, sim_short_18S_Cp, sim_short_18S_CpATCC,
    sim_short_18S_Ct, sim_short_18S_Cu, sim_short_18S_Cv)
```

Plot relative abundance of hits from simulated reads of ~200 bp fragments against the 18S reference database Crypto Data Base.

```
col_c_andersoni <- c("#8dd3c7")
col_c_baileyi <- c("#ffffb3")
col_c_canis <- c("#33a02c")
col_c_meleagridis <- c("#80b1d3")
col_c_muris <- c("#fdb462")
col_c_parvum <- c("#b3de69")
```

```
col_c_serpentis <- c("#1a1a1a")
col_c_sp <- c("#6a3d9a")
col_c_18S_short <- c("#8dd3c7", "#ffffb3", "#33a02c", "#80b1d3", "#fdb462", "#b3de69",
    "#1a1a1a", "#6a3d9a")

p <- sim_short_all_18S %>%
    group_by(Origin) %>%
    arrange(Species) %>%
    plot_ly(x = ~Origin, y = ~relative_ab, color = ~Species, colors = col_c_18S_short,
        type = "bar") %>%
    layout(yaxis = list(title = "Relative Abundance"), barmode = "stack")
```

**NOTE:** The results from these analyses are presented in **Fig. S7F**.

Estimating Relative Abundance.

```
summary_18S_CryptoDB_short <- sim_short_all_18S %>%
    group_by(Origin, Species) %>%
    summarise(Size_Sum = sum(relative_ab))
```

```
## `summarise()` has grouped output by 'Origin'. You can override using the
## `.groups` argument.
```

```
# summary_18S_CryptoDB_short
```

When reads are mapped to the 18S rRNA database, the percentage of reads mapping ranged from 0.1% to 6.1%, interestingly with the highest percentages for species not included in the bait design, such as *C. andersoni*, *C. baileyi*, and *C. muris*, and very low proportions for all species included in the bait design (Table S4). Regarding species assignment, *C. andersoni*, *C. baileyi*, *C. meleagridis*, *C. muris*, and *C. parvum* have high proportions of correct assignment (> 61.6%) for both types of fragments. However, all other species are incorrectly assigned to *C. parvum* with high proportions (Fig. S7E and S7F).

#### Supplementary Notes #5. Mapping CryptoCap_100k baits to Lindgreen et al. (2016) reference genomes database.

To understand the specificity of the bait set, by the sensitivity of the baits to hybridize to off-target DNA, common in environmental and clinical samples, we assessed in silico the affinity of our baits to various genome sequences, including microbial, human, and cow sequences (latter in Supplementary Notes #6). To do this, we used a fasta file with 1,008 whole genome sequences49, which includes genomes of bacteria, archaea, and eukaryote organisms, including the human genome.

To test the sensitivity of our bait set to non-target genomes such as microbial and human DNA that may be present in environmental/stool samples, we retrieved from the Supplementary Table of Lindgreen et al. (2016) (https://www.nature.com/articles/srep19233) the accession numbers of the sequences that they used to simulate their data. Using https://www.ncbi.nlm.nih.gov/sites/batchentrez we download the sequences of 1,008 of the records and save the new file as Lindgreen_genomes.fasta

Then with *BWA* 0.7.17 we mapped our CryptoCap_100k to these metagenomes:

```
time bwa index Lindgreen_genomes.fasta
time bwa mem -M Lindgreen_genomes.fasta baits.fa > L_probes.sam
samtools view -b L_probes.sam | samtools sort > L_probes.bam
```

And estimated how many baits matched to these diverse sequences:

```
samtools flagstat L_probes.bam > L_probes.txt
```

Then, we extracted hits of baits against the Lindgreen metagenome, and counted total number of hits:

```
cat L_probes.sam | grep -v "^@" | \
awk 'BEGIN{FS="\t";OFS="\t"}{if($2=="0" || $2=="16") print $0}' \
> Lindgreen_mapped_both.txt
cat Lindgreen_mapped_both.txt | \
awk 'BEGIN{FS="\t";OFS="\t"}{OTU[$3]+=1}END{for (i in OTU) print i,OTU[i]}' \
> Lindgreen_species_ct.txt
```

##### 5.1 Plotting the proportion of domains from the hits obtained from the mapping of CryptoCap_100k baits to Lindgreen et al. (2016) reference genomes

Then, we took the Lindgreen_species_ct.txt file and included and extra row that contained the name of the species, phylum, and domain of the reference sequence according to the fasta file. Below we plot the number of hits obtained.

Out of the 1,008 reference genomes present in the database our baits presented forward and/or reverse hits to 111 of these.

```
file <- read.delim("~/crypto_data/5/Lindgreen_species_ct.txt",
    header = FALSE)
table <- data.frame(file)
names(table) <- c("domain", "phylum", "species", "accession",
    "hits")
p5 <- plot_ly(table, labels = ~domain, values = ~hits, type = "pie") %>%
    layout(yaxis = list(title = "Number of hits of CryptoCap_100k against Lindgreen database"),
        showlegend = TRUE)
```

**NOTE:** The results from these analyses are presented in **Fig. S8**.

```
euk <- table %>%
    filter(domain == "Eukaryota")
```

The results from these analyses are summarized in **Table S5**.

##### 5.2 Assessing the quality of the mapping of CryptoCap_100k baits to Lindgreen et al. (2016) reference genomes database

To assess the quality of the matching between the CryptoCap_100k baits and the reference genomes, we mapped the histogram of the MAPQ field from the SAM file. MAPQ represents the mapping quality, and it is equal −10 log10 Pr{mapping position is wrong}, rounded to the nearest integer. If the probability of a correct match tends towards zero, so does the MAPQ score.

```
file <- read.delim("~/crypto_data/5/Lindgreen_mapped_both_modified.txt",
    header = FALSE)
table <- data.frame(file)
names(table) <- c("QNAME", "FLAG", "RNAME", "POS", "MAPQ", "CIGAR",
```

```
    "RNEXT", "PNEXT", "TLEN", "SEQ", "QUAL", "OTH1", "OTH2",
    "OTH3", "OTH4")

p6 <- plot_ly(data = table, x = ~MAPQ, type = "histogram")
```

**NOTE:** The results from these analyses are presented in **Fig. S9**.

We further explore which alignment pairs have high mapping quality (>=20):

```
mapq <- table %>%
    filter(MAPQ >= 20)
# write.csv(mapq, 'Lindgreen_mapped_mapq_20.csv')
```

We took this csv file and added the corresponding domain, phylum, and species name to the accession numbers to understand which groups of organisms have high mapping quality scores for our bait set. The file shows that high-quality mapping of our bait sets is present between 75 baits and 31 of the organisms included in the Lindgreen reference database.

```
file <- read.delim("~/crypto_data/5/Lindgreen_mapped_mapq_20.txt",
    header = TRUE)
table <- data.frame(file)
p7 <- plot_ly(table, y = table$MAPQ, x = table$SPECIES, type = "box",
    color = table$SPECIES) %>%
    layout(yaxis = list(title = "MAPQ"), showlegend = FALSE)
```

**NOTE:** The results from these analyses are presented in **Fig. S10**.

Only 894 of our baits (0.01%) mapped to 111 genome sequences from the 1,008 genomes database, including the human genome. In total, 84.2% of these baits mapped to eukaryotes, 13.9% to bacteria, and 1.9% to archaea (Fig. S8; Table S5). Most of the baits mapped with the lowest score (682 MAPQ == 0), and only 75 baits had mapping scores above 20 against 31 non-Cryptosporidium organisms (Fig. S9–S10).

#### Supplementary Notes #6. Mapping CryptoCap_100k baits to *Bos taurus* genome

Because we have been using calf-thymus as background media in some wet-lab tests to dilute samples for several experiments involving the CryptoCap_100k bait set, we wanted to test the sensitivity of our bait set to this non-target DNA. For this, we are using the *Bos taurus* genome available in GenBank: https://www.ncbi.nlm.nih.gov/assembly/GCF_002263795.2 (Accession number GCA_002263795.3):

```
gunzip bos_taurus/*.fna.gz
time bwa index bos_taurus/GCA_002263795.3_ARS-UCD1.3_genomic.fna
time bwa mem \
-M bos_taurus/GCA_002263795.3_ARS-UCD1.3_genomic.fna \
baits/crypto-100K-210317.fas > bos_taurus/bos_taurus.sam
samtools view -b bos_taurus/bos_taurus.sam | samtools sort \
> bos_taurus/bos_taurus.bam
samtools flagstat bos_taurus/bos_taurus.bam > bos_taurus_probes.txt
cat bos_taurus/bos_taurus.sam | grep -v "^@" | \
awk 'BEGIN{FS="\t";OFS="\t"}{if($2=="0" || $2=="16") print $0}' \
> bos_taurus/bos_taurus_mapped.txt
cat bos_taurus/bos_taurus_mapped.txt | \
```

```
awk 'BEGIN{FS="\t";OFS="\t"}{OTU[$3]+=1}END{for (i in OTU) print i,OTU[i]}' \
> bos_taurus_species_ct.txt
gzip ~/crypto_simulations/bos_taurus/*.fna
```

From this alignment, we found that out of the 100,000 baits, 856 mapped to 40 scaffolds of the *Bos taurus* genome.
Then, we plotted the mapping quality scores to see the distribution.

```
file <- read.delim("~/crypto_data/6/bos_taurus_mapped_modified.txt",
    header = FALSE)
table <- data.frame(file)
names(table) <- c("QNAME", "FLAG", "RNAME", "POS", "MAPQ", "CIGAR",
    "RNEXT", "PNEXT", "TLEN", "SEQ", "QUAL", "OTH1", "OTH2",
    "OTH3", "OTH4")
p8 <- plot_ly(data = table, x = ~MAPQ, type = "histogram", color = table$MAPQ)
```

**NOTE:** The results from these analyses are presented in **Fig. S11**.

Then, we selected only those alignments that had high MAPQ scores (>20), to understand how many out of the 856 baits that aligned to *Bos_taurus*, do so with high probability match. From this code we found that only 82 baits high a MAPQ equal or larger than 20 (probability match of 0.99). And most of the ones with highest MAPQ scores map to chromosome 11 and chromosome 27 of the *Bos taurus* genome, as well as to the 'unplaced' sequences.

```
bt_mapq <- table %>%
    filter(MAPQ >= 20)
# bt_mapq
p9 <- plot_ly(bt_mapq, y = ~MAPQ, x = ~RNAME, type = "bar", color = ~MAPQ) %>%
    layout(yaxis = list(title = "sum of MAPQ values"), xaxis = list(title = "Bos taurus chromosome"),
        showlegend = FALSE)
# orca(p9, 'bos_taurus/bostaurus.pdf')
```

**NOTE:** The results from these analyses are presented in **Fig. S12**.

Finally, because some of our in vitro experiments used calf thymus as a background community, we found, from mapping our bait set to the *Bos taurus* genome, that out of the 100,000 baits, only 856 mapped, still an amount larger than those that mapped to the human genome, but more than 600 of these hits had very low mapping quality (MAPQ ==0; Figs. S11–S12) and only 82 baits mapped with scores larger than 20 to chromosomes 11, 27, and 'unplaced' sequences.

#### Supplementary Notes #7. Simulating mixed-infection samples enrichment with CryptoCap_100k and sequence data

Finally, to explore in silico, the ability to discriminate different species within a mixed-infection sample enriched with our bait set, we created four input files to simulate different mixed-infection scenarios. We did this by concatenating the fastq files already generated for each species above and using only those designed large insert fragments. The four input types were: i) a fastq file containing reads from all twelve genomes ( *C. andersoni*, *C. baileyi*, *C. cuniculus*, *C. hominis* x 2, *C. meleagridis*, *C. muris*, *C. parvum* x 2, *C. tyzzeri*, *C. ubiquitum*, and *C. viatorum*); ii) a file containing the sequences of the ten representative species ( *C. andersoni*, *C. baileyi*, *C. cuniculus*, *C. hominis*, *C. meleagridis*, *C. muris*, *C. parvum*, *C. tyzzeri*, *C. ubiquitum*, and *C. viatorum*); iii) a file containing the cluster of five species that share high percent identity ( *C. parvum*, *C. hominis*, *C. tyzzeri*, *C. meleagridis*, and *C. ubiquitum*); and 4) a file containing the three most common species found in human infections (C. hominis, C. cuniculus, and C. parvum). Then, we mapped these mixed reads to 10-CrypGS, the gp60 database, and the 18S rRNA database following the same parameters as above.

We created four samples/files to simulate mixed infections. We achieved this by concatenating a given fastq file with the fastq files already simulated for enrichment and sequencing using CryptoCap_100k baits for each species. We used only those reads generated assuming large fragments:

1) all twelve genomes: *C. andersoni*, *C. baileyi*, *C. cuniculus*, *C. hominis* x 2, *C. meleagridis*, *C. muris*, *C. parvum* x 2, *C. tyzzeri*, *C. ubiquitum*, *C. viatorum*
2) ten species: *C. andersoni*, *C. baileyi*, *C. cuniculus*, *C. hominis*, *C. meleagridis*, *C. muris*, *C. parvum*, *C. tyzzeri*, *C. ubiquitum*, *C. viatorum*
3) five species: *C. parvum*, *C. hominis*, *C. tyzzeri*, *C. meleagridis*, and *C. ubiquitum*
4) three species: *C. hominis*, *C. cuniculus*, and *C. parvum*

##### 7.1 Map the above generated fastqs against the 10-genome reference database

We mapped the above-generated fastqs against the ten-reference genome database. These results are summarized in **Table S6**.

**7.1.1 Species assignment based on mapping simulated mixed-infection reads** We extracted those hits where both reads mapped and counted the relative abundance of species according to the number of hits.

```
# Simulated mixed infections
C_twelve_10 <- read.table("~/crypto_data/7/C_twelve_largespecies_ct.txt"
    header = F, sep = "\t", stringsAsFactors = F)
colnames(C_twelve_10) <- c("Species", "Locus", "Count")
# adding extra column
C_twelve_10 <- C_twelve_10 %>%
    mutate(Sample = "1_twelve_genomes")
# Calculate Relative abundance of counts for each genome
C_twelve_10 <- C_twelve_10 %>%
    group_by(Sample) %>%
    mutate(relative_ab = Count/sum(Count))

# Ten genomes
C_ten_10 <- read.table("~/crypto_data/7/C_ten_largespecies_ct.txt",
    header = F, sep = "\t", stringsAsFactors = F)
colnames(C_ten_10) <- c("Species", "Locus", "Count")
# adding extra column
C_ten_10 <- C_ten_10 %>%
    mutate(Sample = "2_ten_genomes")
# Calculate Relative abundance of counts for each genome
C_ten_10 <- C_ten_10 %>%
    group_by(Sample) %>%
    mutate(relative_ab = Count/sum(Count))

# Five genomes
C_five_10 <- read.table("~/crypto_data/7/C_five_largespecies_ct.txt",
    header = F, sep = "\t", stringsAsFactors = F)
```

```r
colnames(C_five_10) <- c("Species", "Locus", "Count")
# adding extra column
C_five_10 <- C_five_10 %>%
    mutate(Sample = "3_five_genomes")
# Calculate Relative abundance of counts for each genome
C_five_10 <- C_five_10 %>%
    group_by(Sample) %>%
    mutate(relative_ab = Count/sum(Count))

# Three genomes
C_three_10 <- read.table("~/crypto_data/7/C_three_largespecies_ct.txt",
    header = F, sep = "\t", stringsAsFactors = F)
colnames(C_three_10) <- c("Species", "Locus", "Count")
# adding extra column
C_three_10 <- C_three_10 %>%
    mutate(Sample = "4_three_genomes")
# Calculate Relative abundance of counts for each genome
C_three_10 <- C_three_10 %>%
    group_by(Sample) %>%
    mutate(relative_ab = Count/sum(Count))

# Combine all samples
mixed_sim_samples_10 <- rbind(C_twelve_10, C_ten_10, C_five_10,
    C_three_10)

# plot
p <- mixed_sim_samples_10 %>%
    group_by(Sample) %>%
    arrange(Species) %>%
    plot_ly(x = ~Sample, y = ~relative_ab, color = ~Species,
        colors = col_c_ten, type = "bar") %>%
    layout(yaxis = list(title = "Relative Abundance"), barmode = "stack")
```

**NOTE:** The results from these analyses are presented in **Fig. S13A**.

We used the code below to tabulate the total relative abundance for each species.

```r
totals <- aggregate(mixed_sim_samples_10$relative_ab, by = list(species = mixed_sim_samples_10$Species,
    sample = mixed_sim_samples_10$Sample), FUN = sum)
# spread(totals, 'sample', 'x') %>% remove_rownames %>%
# column_to_rownames(var='species') %>% round(digits = 2)
```

The results from these analyses are summarized in **Table S7**.

For the four data sets for which we simulated mixed infections, we obtained a high proportion of reads ( 99.88%) mapping to the ten-genome reference database (Table S7). For all datasets, all species in a sample are correctly identified, and only very slight variations regarding the expected relative abundance and the observed are shown (Fig. S13A, Table S7).

##### 7.2 Map the above-generated fastqs against the gp60 database

We mapped the above-generated fastqs against the gp60 database, extracted those hits where both reads mapped, and counted the relative abundance of species to which the reads mapped. The results from these analyses are summarized in **Table S6**.

**7.2.1 Species assignment based on mapping simulated mixed-infection reads** We extracted those hits where both reads mapped and counted the relative abundance of species according to the number of hits.

```
# Simulated mixed infections
C_twelve_10 <- read.table("~/crypto_data/7/gp60/C_twelve_largespecies_ct.txt"
    header = F, sep = "\t", stringsAsFactors = F)
colnames(C_twelve_10) <- c("Species", "Locus", "Count")
# adding extra column
C_twelve_10 <- C_twelve_10 %>%
    mutate(Sample = "1_twelve_genomes")
# Calculate Relative abundance of counts for each genome
C_twelve_10 <- C_twelve_10 %>%
    group_by(Sample) %>%
    mutate(relative_ab = Count/sum(Count))

# Ten genomes
C_ten_10 <- read.table("~/crypto_data/7/gp60/C_ten_largespecies_ct.txt",
    header = F, sep = "\t", stringsAsFactors = F)
colnames(C_ten_10) <- c("Species", "Locus", "Count")
# adding extra column
C_ten_10 <- C_ten_10 %>%
    mutate(Sample = "2_ten_genomes")
# Calculate Relative abundance of counts for each genome
C_ten_10 <- C_ten_10 %>%
    group_by(Sample) %>%
    mutate(relative_ab = Count/sum(Count))

# Five genomes
C_five_10 <- read.table("~/crypto_data/7/gp60/C_five_largespecies_ct.txt"
    header = F, sep = "\t", stringsAsFactors = F)
colnames(C_five_10) <- c("Species", "Locus", "Count")
# adding extra column
C_five_10 <- C_five_10 %>%
    mutate(Sample = "3_five_genomes")
# Calculate Relative abundance of counts for each genome
C_five_10 <- C_five_10 %>%
    group_by(Sample) %>%
    mutate(relative_ab = Count/sum(Count))

# Three genomes
C_three_10 <- read.table("~/crypto_data/7/gp60/C_three_largespecies_ct.txt"
    header = F, sep = "\t", stringsAsFactors = F)
colnames(C_three_10) <- c("Species", "Locus", "Count")
# adding extra column
C_three_10 <- C_three_10 %>%
    mutate(Sample = "4_three_genomes")
# Calculate Relative abundance of counts for each genome
C_three_10 <- C_three_10 %>%
    group_by(Sample) %>%
    mutate(relative_ab = Count/sum(Count))
```

```
# Combine all samples
mixed_sim_samples_gp60 <- rbind(C_twelve_10, C_ten_10, C_five_10,
    C_three_10)

# plot
p <- mixed_sim_samples_gp60 %>%
    group_by(Sample) %>%
    arrange(Species) %>%
    plot_ly(x = ~Sample, y = ~relative_ab, color = ~Species,
        colors = col_c_gp60, type = "bar") %>%
    layout(yaxis = list(title = "Relative Abundance"), barmode = "stack")
```

**NOTE:** The results from these analyses are presented in **Fig. S13B**.

```
totals <- aggregate(mixed_sim_samples_gp60$relative_ab, by = list(species = mixed_sim_samples_gp60$Speci
    sample = mixed_sim_samples_gp60$Sample), FUN = sum)
```

The summary of these analyses can be found in **Table S7**.

When simulated mixed infection data were mapped to the gp60 database, between 0.015 and 0.018% of reads were mapped to it (Table S6). For the twelve- and ten-genomes mixed infection data, and because of the nature of the database, not all species were assigned correctly, and relative abundances were skewed regarding what was expected, especially with *C. parvum* and *C. ubiquitum* dominating the assignments and *C. meleagridis* and *C. tyzzeri* being underrepresented (Fig. S13B).

However, for the five and three-genome mixed infection data, all species were identified, but also *C. fayeri* was incorrectly identified, although in small proportions (1-2%). The relative abundance of species for the five-genomes sample is dominated by *C. parvum*, with 36% out of the 20% expected, and *C. meleagridis* and *C. tyzzeri*, with just 8% and 12%, respectively, out of the 20% expected. The relative abundance for the three-genomes sample for the three target species was close to the expected with 0.29%, 0.34%, and 0.35% (Fig. S13B, Table S7).

##### 7.3 Map the above-generated fastqs against the CryptoDB 18S rRNA database

We map the above-generated fastqs against the ten 18S rRNA database, extract those hits where both reads are mapped, and count the relative abundance of species to which the reads map.

These results are summarized in **Table S6**.

###### 7.3.1 Species assignment based on mapping simulated mixed-infection reads

We extracted those hits were both reads mapped and count relative abundance of species according to the number of hits.

```
# Simulated mix-infections
C_twelve_10 <- read.table("~/crypto_data/7/18S_rRNA/C_twelve_largespecies_ct.txt"
    header = F, sep = "\t", stringsAsFactors = F)
colnames(C_twelve_10) <- c("Species", "Locus", "Count")
# adding extra column
C_twelve_10 <- C_twelve_10 %>%
    mutate(Sample = "1_twelve_genomes")
# Calculate Relative abundance of counts for each genome
C_twelve_10 <- C_twelve_10 %>%
```

```
    group_by(Sample) %>%
    mutate(relative_ab = Count/sum(Count))

# Ten genomes
C_ten_10 <- read.table("~/crypto_data/7/18S_rRNA/C_ten_largespecies_ct.txt"
    header = F, sep = "\t", stringsAsFactors = F)
colnames(C_ten_10) <- c("Species", "Locus", "Count")
# adding extra column
C_ten_10 <- C_ten_10 %>%
    mutate(Sample = "2_ten_genomes")
# Calculate Relative abundance of counts for each genome
C_ten_10 <- C_ten_10 %>%
    group_by(Sample) %>%
    mutate(relative_ab = Count/sum(Count))

# Five genomes
C_five_10 <- read.table("~/crypto_data/7/18S_rRNA/C_five_largespecies_ct.txt"
    header = F, sep = "\t", stringsAsFactors = F)
colnames(C_five_10) <- c("Species", "Locus", "Count")
# adding extra column
C_five_10 <- C_five_10 %>%
    mutate(Sample = "3_five_genomes")
# Calculate Relative abundance of counts for each genome
C_five_10 <- C_five_10 %>%
    group_by(Sample) %>%
    mutate(relative_ab = Count/sum(Count))

# Three genomes
C_three_10 <- read.table("~/crypto_data/7/18S_rRNA/C_three_largespecies_ct.txt"
    header = F, sep = "\t", stringsAsFactors = F)
colnames(C_three_10) <- c("Species", "Locus", "Count")
# adding extra column
C_three_10 <- C_three_10 %>%
    mutate(Sample = "4_three_genomes")
# Calculate Relative abundance of counts for each genome
C_three_10 <- C_three_10 %>%
    group_by(Sample) %>%
    mutate(relative_ab = Count/sum(Count))

# Combine all samples
mixed_sim_samples_18S <- rbind(C_twelve_10, C_ten_10, C_five_10,
    C_three_10)

# plot
col_c_andersoni <- c("#8dd3c7")
col_c_baileyi <- c("#ffffb3")
col_c_bovis <- c("#1f78b4")
col_c_canis <- c("#33a02c")
col_c_environmental <- c("#e31a1c")
col_c_meleagridis <- c("#80b1d3")
col_c_muris <- c("#fdb462")
col_c_parvum <- c("#b3de69")
```

```r
col_c_serpentis <- c("#1a1a1a")
col_c_sp <- c("#6a3d9a")
col_c_varanii <- c("#c51b7d")
col_c_wrairi <- c("#8c510a")
col_c_18S_mixed <- c("#8dd3c7", "#ffffb3", "#1f78b4", "#33a02c",
    "#e31a1c", "#80b1d3", "#fdb462", "#b3de69", "#1a1a1a", "#6a3d9a",
    "#c51b7d", "#8c510a")

P <- mixed_sim_samples_18S %>%
    group_by(Sample) %>%
    arrange(Species) %>%
    plot_ly(x = ~Sample, y = ~relative_ab, color = ~Species,
        colors = col_c_18S_mixed, type = "bar") %>%
    layout(yaxis = list(title = "Relative Abundance"), barmode = "stack")
```

**NOTE:** The results from these analyses are presented in **Fig. S13C**.

```r
totals <- aggregate(mixed_sim_samples_18S$relative_ab, by = list(species = mixed_sim_samples_18S$Specie
    sample = mixed_sim_samples_18S$Sample), FUN = sum)
```

The results from these analyses are summarized in **Table S7**.

When these mixed infection data were mapped to the 18S rRNA database, between 0.074% and 1.270% of simulated reads were mapped to it (Table S6). In this case, not all species in the sample were assigned correctly, and the abundance was skewed from the expected (Fig. S13C, Table S7). For example, *C. hominis* is not assigned to any of the samples. For the twelve and ten-genome samples, between 90-91% of the abundance is attributed to *C. andersoni*, *C. baileyi*, and *C. muris*, 4-5% of the abundance is assigned to *C. meleagridis* and *C. parvum*, and 5% to other species and genotypes. The relative abundance of *C. meleagridis* for the five-genome sample matches the expected (20%). Still, most of the remaining abundance is assigned to *C. parvum* (70%) without the other species being represented. Similarly, for the three-genome samples, most of the relative abundance is assigned to *C. parvum* (89%) (Fig. S13C, Table S7).

#### Supplementary Notes #8. In-vitro enrichment and validation with CryptoCap_100k baits in DNA from pure oocyst samples

Sequencing reads for enriched and unenriched libraries were quality-filtered using Trimmomatic v.0.39:

```bash
java -jar $EBROOTTRIMMOMATIC/trimmomatic-0.39.jar \
PE R1_001.fastq.gz R2_001.fastq.gz R1_pair_trim.fastq.gz R1_unpair_trim.fastq.gz \
R2_pair_trim.fastq.gz R2_unpair_trim.fastq.gz \
 -threads 10 -phred33 \
ILLUMINACLIP:/apps/eb/Trimmomatic/0.39-Java-1.8.0_144/adapters/TruSeq3-PE.fa\
:2:30:10:2:TRUE LEADING:3 TRAILING:3 SLIDINGWINDOW:4:15 MINLEN:75
```

Then, sequencing reads were mapped using bbmap to the their respective reference genome (C. parvum or C. meleagridis), and alignment statistics were outputted.

```bash
bbmap.sh in=R1_pair_trim.fastq.gz in2=R2_pair_trim.fastq.gz \
ref=reference_genome.fasta maxindel=20 minid=0.80 \
rcs=t killbadpairs=t pairlen=400 rescuedist=600 apd=100 outu=mapped/unmapped.sam \
outm=mapped/mapped.sam printunmappedcount=t \
```

```
covstats=cov.txt threads=12 -Xmx180g 2>&1 | tee bbmap.log;

samtools view -b sampple.sam | samtools sort > sample.bam
samtools depth -a sample.bam -o sample.depth
#Obtaining mean depth of coverage
cat sample.depth | cut -f 3 | paste -sd+ | bc
wc -l sample.depth
#Obtaining mean breadth of coverage
samtools samples.bam | awk -v X="${MIN_COVERAGE_DEPTH}" '$4>=X' | wc -l
grep -v ">" reference.fasta | wc | awk '{print $3-$1}'
```

##### 8.1 Pure oocyst DNA sequence data mapped to *C. parvum* and *C. meleagridis*

Here we import **Table S8**.

```
file = "~/crypto_data/8/pure_oocysts_summary_stats.txt" po_table = read.table(file, header
= TRUE, sep = "")
```

For the libraries made from pure oocyst DNA, we obtained an average of 926,305 reads per enriched sample and 521,121 per unenriched sample (Table S8). An average of 95.98% of reads were retained after filtering in enriched samples vs. 77.92% in unenriched samples (Table S8; p-value = 0.07).

###### 8.1.1 Testing for differences in various parameters between enriched and unenriched libraries

Then, we tested the difference in the total number of raw reads obtained between enriched and unenriched samples.

```
lsmeans(lm(data = po_table, Raw_Reads ~ Library), pairwise ~ Library)
```

```
## $lsmeans
##  Library    lsmean     SE df lower.CL upper.CL
##  Enriched   926304 165000  8   546146  1306462
##  Unenriched 521121 165000  8   140963   901278
##
## Confidence level used: 0.95
##
## $contrasts
##  contrast                estimate     SE df t.ratio p.value
##  Enriched - Unenriched     405183 233000  8   1.738  0.1204
```

We tested whether the number of reads retained after filtering differed between enriched and unenriched pure oocyst DNA libraries.

```
lsmeans(lm(data = po_table, Filtered_Reads ~ Library), pairwise ~ Library)
```

```
## $lsmeans
##  Library    lsmean     SE df lower.CL upper.CL
##  Enriched   872283 154000  8   517041  1227525
##  Unenriched 425818 154000  8    70576   781060
##
## Confidence level used: 0.95
```

```
##
## $contrasts
##  contrast                estimate     SE df t.ratio p.value
##  Enriched - Unenriched    446465 218000  8   2.049  0.0746
```

We also tested if the number of reads that mapped to *C. parvum* were different between enriched and unenriched pure oocyst DNA libraries.

```r
lsmeans(lm(data = po_table, Number_mapped_each ~ Library), pairwise ~ Library)
```

```
## $lsmeans
##  Library    lsmean     SE df lower.CL upper.CL
##  Enriched   839198 138000  8   520937  1157458
##  Unenriched 291643 138000  8   -26618   609904
##
## Confidence level used: 0.95
##
## $contrasts
##  contrast                estimate     SE df t.ratio p.value
##  Enriched - Unenriched    547555 195000  8   2.805  0.0230
```

```r
# scatter plot of number of reads obtained and number of reads mapped
fig <- plot_ly(data = po_table, x = ~Raw_Reads, y = ~Number_mapped, color = ~Library,
    symbol = ~Species, marker = list(size = 10))
```

**NOTE:** The results from these analyses are presented in **Fig. S14**.

More reads per sample mapped to their respective genomes from enriched libraries (839,198) than from unenriched libraries (291,643; Fig S14).

We also tested if the percentage of reads mapped after filtering was different between enriched and unenriched pure oocyst DNA libraries.

```r
lsmeans(lm(data = po_table, X_Both_mapped_each ~ Library), pairwise ~ Library)
```

```
## $lsmeans
##  Library    lsmean   SE df lower.CL upper.CL
##  Enriched     95.8 10.3  8     72.0    119.6
##  Unenriched   68.1 10.3  8     44.4     91.9
##
## Confidence level used: 0.95
##
## $contrasts
##  contrast                estimate   SE df t.ratio p.value
##  Enriched - Unenriched      27.7 14.6  8   1.900  0.0940
```

We also tested if the breadth of genome coverage was different between enriched and unenriched pure oocyst DNA libraries.

```r
lsmeans(lm(data = po_table, Breadth_each ~ Library), pairwise ~
    Library)
```

```
## $lsmeans
##  Library    lsmean   SE df lower.CL upper.CL
##  Enriched     85.2 7.63  8     67.6    102.8
##  Unenriched   81.6 7.63  8     64.0     99.2
##
## Confidence level used: 0.95
##
## $contrasts
##  contrast              estimate   SE df t.ratio p.value
##  Enriched - Unenriched     3.63 10.8  8   0.336  0.7456
```

```
p <- plot_ly()
p <- add_bars(p, x = c("Enriched", "Enriched0", "Unenriched"),
    y = c(85, 102, 85), opacity = 1, showlegend = F, marker = list(line = list(color = "rgba(0,0,0,0"),
        color = "rgba(0,0,0,0"), text = c("", "p-value = 0.75",
        ""), textposition = "outside", legendgroup = "1")
p <- add_lines(p, x = c("Enriched", "Enriched", "Unenriched",
    "Unenriched"), y = c(100, 101, 101, 100), showlegend = F,
    line = list(color = "black"), legendgroup = "1", hoverinfo = "none")
p <- add_boxplot(p, data = po_table, x = ~Library, y = ~Breadth_each,
    color = ~Library, type = "box", boxpoints = "all", jitter = 0.3,
    pointpos = 0, legendgroup = "1")
p <- layout(p, xaxis = list(tickmode = "array", tickvals = c("Enriched",
    "sf", "Unenriched"), ticktext = c("Enriched", "", "Unenriched")),
    yaxis = list(range = c(0, 106)))
```

**NOTE:** The results from these analyses are presented in **Fig. S15**.

A higher percentage of enriched reads mapped to the genome from enriched libraries than from unenriched libraries (p-value = 0.001).

We also tested if the mean depth was different between enriched and unenriched pure oocyst DNA samples.

```
lsmeans(lm(data = po_table, Depth_each ~ Library), pairwise ~
    Library)
```

```
## $lsmeans
##  Library    lsmean  SE df lower.CL upper.CL
##  Enriched    17.31 2.6  8    11.31     23.3
##  Unenriched   4.96 2.6  8    -1.03     11.0
##
## Confidence level used: 0.95
##
## $contrasts
##  contrast              estimate   SE df t.ratio p.value
##  Enriched - Unenriched     12.3 3.68  8   3.357  0.0100
```

```
p <- plot_ly()
p <- add_bars(p, x = c("Enriched", "Enriched0", "Unenriched"),
    y = c(17, 29, 5), opacity = 1, showlegend = F, marker = list(line = list(color = "rgba(0,0,0,0"),
        color = "rgba(0,0,0,0"), text = c("", "* p-value = 0.01",
        ""), textposition = "outside", legendgroup = "1")
p <- add_lines(p, x = c("Enriched", "Enriched", "Unenriched",
```

```
    "Unenriched"), y = c(27, 28, 28, 27), showlegend = F, line = list(color = "black"),
    legendgroup = "1", hoverinfo = "none")
p <- add_boxplot(p, data = po_table, x = ~Library, y = ~Depth_each,
    color = ~Library, type = "box", boxpoints = "all", jitter = 0.3,
    pointpos = 0, legendgroup = "1")
p <- layout(p, xaxis = list(tickmode = "array", tickvals = c("Enriched",
    "sf", "Unenriched"), ticktext = c("Enriched", "", "Unenriched")),
    yaxis = list(range = c(0, 30)))
```

**NOTE:** The results from these analyses are presented in **Fig. S16**.

The breadth of genome coverage was similar, but the depth of coverage was higher in enriched vs. unenriched libraries (p-value = 0.01).

##### 8.2 Pure oocyst DNA libraries mapped to 10-genomes reference database and species assignment

We also mapped the samples' fastqs against the ten genomes database, extracted those hits where both reads mapped, and counted the relative abundance of species to which the reads mapped.

Summary results from these analyses are presented in **Table S9**.

```
Enriched_C_parvum_MG01 <- read.table("~/crypto_data/8/Enriched_C_parvum_MG01species_ct.txt"
    header = F, sep = "\t", stringsAsFactors = F)
colnames(Enriched_C_parvum_MG01) <- c("Species", "Locus", "Count")
# adding extra column
Enriched_C_parvum_MG01 <- Enriched_C_parvum_MG01 %>%
    mutate(Sample = "Enriched_C_parvum_MG01")
# Calculate Relative abundance of counts for each genome
Enriched_C_parvum_MG01 <- Enriched_C_parvum_MG01 %>%
    group_by(Sample) %>%
    mutate(relative_ab = Count/sum(Count))

Enriched_C_parvum_MG02 <- read.table("~/crypto_data/8/Enriched_C_parvum_MG02species_ct.txt"
    header = F, sep = "\t", stringsAsFactors = F)
colnames(Enriched_C_parvum_MG02) <- c("Species", "Locus", "Count")
# adding extra column
Enriched_C_parvum_MG02 <- Enriched_C_parvum_MG02 %>%
    mutate(Sample = "Enriched_C_parvum_MG02")
# Calculate Relative abundance of counts for each genome
Enriched_C_parvum_MG02 <- Enriched_C_parvum_MG02 %>%
    group_by(Sample) %>%
    mutate(relative_ab = Count/sum(Count))

Enriched_C_parvum_Boris <- read.table("~/crypto_data/8/Enriched_C_parvum_Borisspecies_ct.txt"
    header = F, sep = "\t", stringsAsFactors = F)
colnames(Enriched_C_parvum_Boris) <- c("Species", "Locus", "Count")
# adding extra column
Enriched_C_parvum_Boris <- Enriched_C_parvum_Boris %>%
    mutate(Sample = "Enriched_C_parvum_Boris")
# Calculate Relative abundance of counts for each genome
Enriched_C_parvum_Boris <- Enriched_C_parvum_Boris %>%
    group_by(Sample) %>%
```

```
    mutate(relative_ab = Count/sum(Count))

Enriched_C_meleagridis01 <- read.table("~/crypto_data/8/Enriched_C_meleagridis01species_ct.txt"
    header = F, sep = "\t", stringsAsFactors = F)
colnames(Enriched_C_meleagridis01) <- c("Species", "Locus", "Count")
# adding extra column
Enriched_C_meleagridis01 <- Enriched_C_meleagridis01 %>%
    mutate(Sample = "Enriched_C_meleagridis01")
# Calculate Relative abundance of counts for each genome
Enriched_C_meleagridis01 <- Enriched_C_meleagridis01 %>%
    group_by(Sample) %>%
    mutate(relative_ab = Count/sum(Count))

Enriched_C_meleagridis02 <- read.table("~/crypto_data/8/Enriched_C_meleagridis02species_ct.txt"
    header = F, sep = "\t", stringsAsFactors = F)
colnames(Enriched_C_meleagridis02) <- c("Species", "Locus", "Count")
# adding extra column
Enriched_C_meleagridis02 <- Enriched_C_meleagridis02 %>%
    mutate(Sample = "Enriched_C_meleagridis02")
# Calculate Relative abundance of counts for each genome
Enriched_C_meleagridis02 <- Enriched_C_meleagridis02 %>%
    group_by(Sample) %>%
    mutate(relative_ab = Count/sum(Count))

Unenriched_C_parvum_MG01 <- read.table("~/crypto_data/8/Unenriched_C_parvum_MG01species_ct.txt"
    header = F, sep = "\t", stringsAsFactors = F)
colnames(Unenriched_C_parvum_MG01) <- c("Species", "Locus", "Count")
# adding extra column
Unenriched_C_parvum_MG01 <- Unenriched_C_parvum_MG01 %>%
    mutate(Sample = "Unenriched_C_parvum_MG01")
# Calculate Relative abundance of counts for each genome
Unenriched_C_parvum_MG01 <- Unenriched_C_parvum_MG01 %>%
    group_by(Sample) %>%
    mutate(relative_ab = Count/sum(Count))

Unenriched_C_parvum_MG02 <- read.table("~/crypto_data/8/Unenriched_C_parvum_MG02species_ct.txt"
    header = F, sep = "\t", stringsAsFactors = F)
colnames(Unenriched_C_parvum_MG02) <- c("Species", "Locus", "Count")
# adding extra column
Unenriched_C_parvum_MG02 <- Unenriched_C_parvum_MG02 %>%
    mutate(Sample = "Unenriched_C_parvum_MG02")
# Calculate Relative abundance of counts for each genome
Unenriched_C_parvum_MG02 <- Unenriched_C_parvum_MG02 %>%
    group_by(Sample) %>%
    mutate(relative_ab = Count/sum(Count))

Unenriched_C_parvum_Boris <- read.table("~/crypto_data/8/Unenriched_C_parvum_Borisspecies_ct.txt"
    header = F, sep = "\t", stringsAsFactors = F)
colnames(Unenriched_C_parvum_Boris) <- c("Species", "Count")
# adding extra column
Unenriched_C_parvum_Boris <- Unenriched_C_parvum_Boris %>%
    mutate(Sample = "Unenriched_C_parvum_Boris")
```

```r
# Calculate Relative abundance of counts for each genome
Unenriched_C_parvum_Boris <- Unenriched_C_parvum_Boris %>%
    group_by(Sample) %>%
    mutate(relative_ab = Count/sum(Count))

Unenriched_C_meleagridis01 <- read.table("~/crypto_data/8/Unenriched_C_meleagridis01species_ct.txt"
    header = F, sep = "\t", stringsAsFactors = F)
colnames(Unenriched_C_meleagridis01) <- c("Species", "Locus",
    "Count")
# adding extra column
Unenriched_C_meleagridis01 <- Unenriched_C_meleagridis01 %>%
    mutate(Sample = "Unenriched_C_meleagridis01")
# Calculate Relative abundance of counts for each genome
Unenriched_C_meleagridis01 <- Unenriched_C_meleagridis01 %>%
    group_by(Sample) %>%
    mutate(relative_ab = Count/sum(Count))

Unenriched_C_meleagridis02 <- read.table("~/crypto_data/8/Unenriched_C_meleagridis02species_ct.txt"
    header = F, sep = "\t", stringsAsFactors = F)
colnames(Unenriched_C_meleagridis02) <- c("Species", "Locus",
    "Count")
# adding extra column
Unenriched_C_meleagridis02 <- Unenriched_C_meleagridis02 %>%
    mutate(Sample = "Unenriched_C_meleagridis02")
# Calculate Relative abundance of counts for each genome
Unenriched_C_meleagridis02 <- Unenriched_C_meleagridis02 %>%
    group_by(Sample) %>%
    mutate(relative_ab = Count/sum(Count))

# Combine all samples
mixed_pure_oocyst <- rbind(Enriched_C_parvum_MG01, Enriched_C_parvum_MG02,
    Enriched_C_parvum_Boris, Enriched_C_meleagridis01, Enriched_C_meleagridis02,
    Unenriched_C_parvum_MG01, Unenriched_C_parvum_MG02, Unenriched_C_parvum_Boris,
    Unenriched_C_meleagridis01, Unenriched_C_meleagridis02)

# plot

plot <- mixed_pure_oocyst %>%
    group_by(Sample) %>%
    arrange(Species) %>%
    plot_ly(x = ~Sample, y = ~relative_ab, color = ~Species,
        colors = col_c_ten, type = "bar") %>%
    layout(yaxis = list(title = "Relative Abundance"), barmode = "stack")
```

**NOTE:** The results from these analyses are presented in **Fig. S17**.

Mapping reads to the reference database containing ten *Cryptosporidium* genome sequences showed that reads of both species were largely correctly assigned, but some *C. parvum* reads were assigned to two other species ( *C. cuniculus* and *C. hominis*), and incorrect assignments only slightly elevated in the unenriched samples (Table S9).

### Supplementary Notes #9. In Vitro Simulation and Detection of Mixed-Infection Samples

#### 9.1 Mixed-infection samples prepared with NEB protocol and iTru primers

Here, we analyze four samples: Crypt08, Crypto10, Crypto26, and Crypto28. Crypto08 and Crypto26 contain DNA from *C. parvum* for which library prep until PCR was done independently, and Crypto 10 and Crypto 28 contain DNA from *C. meleagridis* for which library prep was done independently. After PCR, these two DNA samples were pooled in equimolar ratios, one aliquot of this pool was enriched (containing 08 and 10), and one aliquot was unenriched (comprising 26 and 28). This library scheme enables us to assess whether the presence of multiple species in a sample introduces biases in the capture process, favoring one species over another.

The summary results from these analyses are depicted in **Table S10**, also imported below.

```
file = "~/crypto_data/9/mixed_infections_NEB-iTru_summary_stats.txt" mix_neb_table = read.table(file,
header = TRUE, sep = "")
```

Here we assessed the number of raw paired-reads obtained for enriched and unenriched samples.

```
lsmeans(lm(data = mix_neb_table, Raw_Paired_Reads ~ Library),
    pairwise ~ Library)
```

```
## $lsmeans
##  Library    lsmean     SE df lower.CL upper.CL
##  Enriched   558771 136000  2   -24371  1141913
##  Unenriched 498066 136000  2   -85075  1081208
##
## Confidence level used: 0.95
##
## $contrasts
##  contrast              estimate     SE df t.ratio p.value
##  Enriched - Unenriched    60704 192000  2   0.317  0.7815
```

```
col_c_mix <- c("#80b1d3", "#80b1d3", "#b3de69", "#b3de69")

p <- plot_ly(colors = col_c_mix)
p <- add_lines(p, x = c("Enriched", "Enriched0"), y = c(558771,
    558771), showlegend = F, line = list(color = "salmon"), legendgroup = "1",
    hoverinfo = "none")
p <- add_lines(p, x = c("Enriched0", "Unenriched"), y = c(498067,
    498067), showlegend = F, line = list(color = "salmon"), legendgroup = "1",
    hoverinfo = "none")
p <- add_bars(p, data = mix_neb_table, x = ~Library, y = ~Raw_Paired_Reads,
    color = ~Sample_Name)
p <- layout(p, xaxis = list(tickmode = "array", tickvals = c("Enriched",
    "sf", "Unenriched"), ticktext = c("Enriched", "", "Unenriched")),
    yaxis = list(range = c(-1, 9e+05)))
```

**NOTE:** The results from these analyses are presented in **Fig. 1A**.

**9.1.1 Sequence data mapped to 10-genomes reference database** Similarly to simulated sequencing reads, we start by mapping each sample sequenced against the fasta containing ten genomes. We used BBmap, and as a reference, we used the reference file created above. We used a maxim indel length of 20, a minid of 0.8,, we also required correct strands for paired reads (rcs), set the mean pair size to 400, and did not rescue paired reads if average insert size was greater than 600 (rescuedist; let's remember that the simulated size is 500), and we set an average pair distance *(apd)of 100 bases.

```
mkdir mixed_infections/mapped
bbmap.sh in=1.fq in2=2.fq \
ref 10_crypto_genomes.fasta  \
maxindel=20 minid=0.80 rcs=t killbadpairs=t pairlen=400 \
rescuedist=600 apd=100 outu=mixed_infections/mapped/un_sample.sam \
outm=mixed_infections/mapped/sample.sam printunmappedcount=t \
covstats=mixed_infections/mapped/sample.cov.txt threads=12 -Xmx180g 2>&1 \
| tee mixed_infections/mapped/sample.bbmap.log
```

For each sample, we used the FLAG count from the third column of our SAM files to extract those sequences for which only both reads mapped (FLAG: 83 and 99)

```
cat mixed_infections/mapped/sample.sam | grep -v "^@" | \
awk 'BEGIN{FS="\t";OFS="\t"}{if($2=="83" || $2=="99") print $0}' \
> mixed_infections/mapped/both/sample.txt
```

Then, for each sample, we counted each species ID hit in our mapped file. The count text file has two columns, the first is the species ID, and the second is the counts.

```
cat mixed_infections/mapped/both/sample.txt | \
awk 'BEGIN{FS="\t";OFS="\t"}{OTU[$3]+=1}END{for (i in OTU) print i,OTU[i]}' \
> mixed_infections/mapped/both/species_ct.txt
```

This generates a file that looks like this:

> C_andersoni-LRBS01000112.1 Cryptosporidium andersoni isolate 30847 contig_19, whole genome shotgun sequence 1800

| Label | Sample | Library | # Sequence Reads | % Pairs Mapped |
|---|---|---|---|---|
| Crypto_8 | C. parvum MG01 | Enriched | 1,426,830 | 60.33% |
| Crypto_10 | C. meleagridis01 | Enriched | 798,254 | 81.90% |
| Crypto_26 | C. parvum MG02 | Unenriched | 1,208,274 | 20.69% |
| Crypto_28 | C. meleagridis02 | Unenriched | 783,992 | 54.06% |

We tested if the percentage of reads that mapped to ten genome reference database was different between enriched and unenriched mixed-infection mocked samples.

```
lsmeans(lm(data = mix_neb_table, Per_Paired_Mapped_Reads_10_Genomes ~ Library), pairwise ~
    Library)
```

```
## $lsmeans
##  Library    lsmean    SE df lower.CL upper.CL
```

```
##  Enriched     95.7 12.5  2     42.1      149
##  Unenriched   81.4 12.5  2     27.7      135
##
## Confidence level used: 0.95
##
## $contrasts
##  contrast              estimate   SE df t.ratio p.value
##  Enriched - Unenriched     14.4 17.6  2   0.816  0.5002
```

We then assessed species assignment based on the mapping to the 10-genome database.

```
crypto_8 <- read.table("~/crypto_data/9/Crypto-8_S8species_ct.txt",
    header = F, sep = "\t", stringsAsFactors = F)
colnames(crypto_8) <- c("Species", "Locus", "Count")
# adding extra column
crypto_8 <- crypto_8 %>%
    mutate(Sample = "C_parvum_enr")
# Calculate Relative abundance of counts for each genome
crypto_8 <- crypto_8 %>%
    group_by(Sample) %>%
    mutate(relative_ab = Count/sum(Count))

# Crypto 10
crypto_10 <- read.table("~/crypto_data/9/Crypto-10_S10species_ct.txt",
    header = F, sep = "\t", stringsAsFactors = F)
colnames(crypto_10) <- c("Species", "Locus", "Count")
# adding extra column
crypto_10 <- crypto_10 %>%
    mutate(Sample = "C_meleagridis_enr")
# Calculate Relative abundance of counts for each genome
crypto_10 <- crypto_10 %>%
    group_by(Sample) %>%
    mutate(relative_ab = Count/sum(Count))

# Crypto 26
crypto_26 <- read.table("~/crypto_data/9/Crypto-26_S26species_ct.txt",
    header = F, sep = "\t", stringsAsFactors = F)
colnames(crypto_26) <- c("Species", "Locus", "Count")
# adding extra column
crypto_26 <- crypto_26 %>%
    mutate(Sample = "C_parvum_unenr")
# Calculate Relative abundance of counts for each genome
crypto_26 <- crypto_26 %>%
    group_by(Sample) %>%
    mutate(relative_ab = Count/sum(Count))

# Crypto 28
crypto_28 <- read.table("~/crypto_data/9/Crypto-28_S28species_ct.txt",
    header = F, sep = "\t", stringsAsFactors = F)
colnames(crypto_28) <- c("Species", "Locus", "Count")
# adding extra column
crypto_28 <- crypto_28 %>%
    mutate(Sample = "C_meleagridis_unenr")
# Calculate Relative abundance of counts for each genome
```

```
crypto_28 <- crypto_28 %>%
    group_by(Sample) %>%
    mutate(relative_ab = Count/sum(Count))

# Combine all samples
mixed_samples <- rbind(crypto_8, crypto_10, crypto_26, crypto_28)
# save(mixed_samples, file ='mixed_infections_feature_table_two.Rdata')

# plot
plot <- mixed_samples %>%
    group_by(Sample) %>%
    arrange(Species) %>%
    plot_ly(x = ~Sample, y = ~relative_ab, color = ~Species, colors = col_c_ten,
        type = "bar") %>%
    layout(yaxis = list(title = "Relative Abundance"), barmode = "stack")
```

**NOTE:** The results from these analyses are presented in **Fig. 1B**.

**9.1.2 Using the sequence data to map against the gp60 database** We mapped reads using similar parameters as above, except pairlen = 400 and maxindel = 5, and using as references independently one gp60 fasta file containing 65 gp60 sequences:

```
bbmap.sh in=1.fq in2=2.fq \
ref GP60new.fasta/  \
maxindel=5 minid=0.80 rcs=t killbadpairs=t pairlen=400 \
rescuedist=600 apd=100 outu=mixed_infections/mapped_gp60/\$i\_unmapped.sam \
outm=mixed_infections/mapped_gp60/\$i\_mapped.sam printunmappedcount=t \
covstats=mixed_infections/mapped_gp60/\$i\.cov.txt threads=12 -Xmx180g 2>&1 \
| tee mixed_infections/mapped_gp60/\$i\.bbmap.log
```

| Label | Sample Name | Library | # Sequence Reads | % Proper Pairs Mapped |
|---|---|---|---|---|
| Crypto_8 | C. parvum MG01 | Enriched | 1,426,830 | 0.03% |
| Crypto_10 | C. meleagridis01 | Enriched | 798,254 | 0.08% |
| Crypto_26 | C. parvum MG02 | Unenriched | 1,208,274 | 0.03% |
| Crypto_28 | C. meleagridis02 | Unenriched | 783,992 | 0.04% |

Similar to above, for each sample, we used the FLAG count from the third column of our SAM files to extract those sequences for which both reads mapped (FLAG: 83 and 99). However, samples 9, 11, 27, and 29 do not present both reads mapping the database, so they could not be considered in the plot below. These correspond to those labeled as *C. baileyi* and *C. cuniculus* samples from both enriched and unenriched libraries.

```
cat gp60_mapped.sam | grep -v "^@" | \
awk 'BEGIN{FS="\t";OFS="\t"}{if($2=="83" || $2=="99") print $0}' \
> both_mapped/gp60_mapped_both.txt
```

Then, for each species, we will count each species ID hit in our mapped file. The count text file has two columns: the first is the species ID, and the second is the counts.

```
cat gp60_mapped_both.txt | \
awk 'BEGIN{FS="\t";OFS="\t"}{OTU[$3]+=1}END{for (i in OTU) print i,OTU[i]}' \
> gp60_ct.txt
```

Similarly to above, before importing each file, in Excel, we added an extra column with only the species name of the gp60 from the fasta header. Below, we then create a fourth column, “Sample,” with the name of the sample prepared.

```
crypto_8_gp60 <- read.table("~/crypto_data/9/gp60/Crypto-8_S8_species_ct.txt"
    header = F, sep = "\t", stringsAsFactors = F)
colnames(crypto_8_gp60) <- c("Species", "Locus", "Count")
# adding extra column
crypto_8_gp60 <- crypto_8_gp60 %>%
    mutate(Sample = "C_parvum_enr")
# Calculate Relative abundance of counts for each genome
crypto_8_gp60 <- crypto_8_gp60 %>%
    group_by(Sample) %>%
    mutate(relative_ab = Count/sum(Count))

# Crypto 10
crypto_10_gp60 <- read.table("~/crypto_data/9/gp60/Crypto-10_S10_species_ct.txt"
    header = F, sep = "\t", stringsAsFactors = F)
colnames(crypto_10_gp60) <- c("Species", "Locus", "Count")
# adding extra column
crypto_10_gp60 <- crypto_10_gp60 %>%
    mutate(Sample = "C_meleagridis_enr")
# Calculate Relative abundance of counts for each genome
crypto_10_gp60 <- crypto_10_gp60 %>%
    group_by(Sample) %>%
    mutate(relative_ab = Count/sum(Count))

# Crypto 26
crypto_26_gp60 <- read.table("~/crypto_data/9/gp60/Crypto-26_S26_species_ct.txt"
    header = F, sep = "\t", stringsAsFactors = F)
colnames(crypto_26_gp60) <- c("Species", "Locus", "Count")
# adding extra column
crypto_26_gp60 <- crypto_26_gp60 %>%
    mutate(Sample = "C_parvum_unenr")
# Calculate Relative abundance of counts for each genome
crypto_26_gp60 <- crypto_26_gp60 %>%
    group_by(Sample) %>%
    mutate(relative_ab = Count/sum(Count))

# Crypto 28
crypto_28_gp60 <- read.table("~/crypto_data/9/gp60/Crypto-28_S28_species_ct.txt"
    header = F, sep = "\t", stringsAsFactors = F)
colnames(crypto_28_gp60) <- c("Species", "Locus", "Count")
# adding extra column
crypto_28_gp60 <- crypto_28_gp60 %>%
    mutate(Sample = "C_meleagridis_unenr")
# Calculate Relative abundance of counts for each genome
crypto_28_gp60 <- crypto_28_gp60 %>%
    group_by(Sample) %>%
```

```
    mutate(relative_ab = Count/sum(Count))

# Combine all samples
mixed_samples_gp60 <- rbind(crypto_8_gp60, crypto_10_gp60, crypto_26_gp60,
    crypto_28_gp60)

col_c_fayeri <- c("#ffed6f")
col_c_meleagridis <- c("#80b1d3")
col_c_parvum <- c("#b3de69")
col_c_gp60_mix <- c("#ffed6f", "#80b1d3", "#b3de69")

# plot
p <- mixed_samples_gp60 %>%
    group_by(Sample) %>%
    arrange(Species) %>%
    plot_ly(x = ~Sample, y = ~relative_ab, color = ~Species,
        colors = col_c_gp60_mix, type = "bar") %>%
    layout(yaxis = list(title = "Relative Abundance"), barmode = "stack")
```

**NOTE:** The results from these analyses are presented in **Fig. 1C**.

The reads mapped to the gp60 database performed well for the *C. parvum* sample. Still, less than half of the reads were correctly assigned for the *C. meleagridis* sample (Fig. 1C). When the mock mixed-infection reads were mapped to the gp60 database, 1.1% of retained reads mapped to the database for enriched samples and 0.07% for unenriched samples. Regarding species assignment, *C. parvum* was properly assigned with a relative abundance of 95% for the unenriched sample and 100% for the enriched sample. However, *C. meleagridis* was only properly assigned in both types of libraries, with a relative abundance of 41-47%, attributing the remaining relative abundance to *C. parvum* (Fig. 1C).

**9.1.3 Using the sequence data to map against CryptoDB 18S rRNA database** We mapped simulated reads using similar parameters as above, except pairlen = 500 and maxindel = 5, and using as references the fasta file containing 57 reference sequences from: https://cryptodb.org/cryptodb/app/downloads/Current_Release/CryptosporidiidaeReference/fasta/ (version last modified on 2022-04-13 11:11), here referred as 18S_CDB:

```
bbmap.sh in=1.fq in2=2.fq \
ref 18S_reference.fasta  \
maxindel=5 minid=0.80 rcs=t killbadpairs=t pairlen=400 \
rescuedist=600 apd=100 outu=sim_mapped/\$i\_unmapped.sam \
outm=sim_mapped/\$i\_mapped.sam printunmappedcount=t \
covstats=sim_mapped/\$i\.cov.txt threads=12 -Xmx180g 2>&1 \
| tee sim_mapped/\$i\.bbmap.log
```

| Sequence Name | Sample Name | Library | # Sequence Reads | % Proper Pairs Mapped |
|---|---|---|---|---|
| Crypto_8 | C. parvum MG | Enriched | 1,426,830 | 0.022% |
| Crypto_10 | C. meleagridis | Enriched | 798,254 | 0.047% |
| Crypto_26 | C. parvum MG | Unenriched | 1,208,274 | 0.013% |
| Crypto_28 | C. meleagridis | Unenriched | 783,992 | 0.043% |

Similar to above, for each species, we used the FLAG count from the third column of our SAM files to extract those sequences for which only both reads mapped (FLAG: 83 and 99)

```
cat 18S_mapped.sam | grep -v "^@" | \
awk 'BEGIN{FS="\t";OFS="\t"}{if($2=="83" || $2=="99") print $0}' \
> both_mapped/18S_mapped_both.txt
```

Then, for each species, we will count each species ID hit in our mapped file. The count text file has two columns: the first is the species ID, and the second is the counts.

```
cat 18S_mapped_both.txt | \
awk 'BEGIN{FS="\t";OFS="\t"}{OTU[$3]+=1}END{for (i in OTU) print i,OTU[i]}' \
> 18S_ct.txt
```

Similarly to above, before importing each file, in Excel, we added an extra column with only the species name of the 18S from the fasta header.

Below, we then create a fourth column, "Origin," with the name of the species for which we simulated the reads.

```
crypto_8_18S <- read.table("~/crypto_data/9/18S_rRNA/Crypto-8_S8species_ct.txt"
    header = F, sep = "\t", stringsAsFactors = F)
colnames(crypto_8_18S) <- c("Species", "Locus", "Count")
# adding extra column
crypto_8_18S <- crypto_8_18S %>%
    mutate(Sample = "C_parvum_enr")
# Calculate Relative abundance of counts for each genome
crypto_8_18S <- crypto_8_18S %>%
    group_by(Sample) %>%
    mutate(relative_ab = Count/sum(Count))

# Crypto 10
crypto_10_18S <- read.table("~/crypto_data/9/18S_rRNA/Crypto-10_S10species_ct.txt"
    header = F, sep = "\t", stringsAsFactors = F)
colnames(crypto_10_18S) <- c("Species", "Locus", "Count")
# adding extra column
crypto_10_18S <- crypto_10_18S %>%
    mutate(Sample = "C_meleagridis_enr")
# Calculate Relative abundance of counts for each genome
crypto_10_18S <- crypto_10_18S %>%
    group_by(Sample) %>%
    mutate(relative_ab = Count/sum(Count))

# Crypto 26
crypto_26_18S <- read.table("~/crypto_data/9/18S_rRNA/Crypto-26_S26species_ct.txt"
    header = F, sep = "\t", stringsAsFactors = F)
colnames(crypto_26_18S) <- c("Species", "Locus", "Count")
# adding extra column
crypto_26_18S <- crypto_26_18S %>%
    mutate(Sample = "C_parvum_unenr")
# Calculate Relative abundance of counts for each genome
crypto_26_18S <- crypto_26_18S %>%
    group_by(Sample) %>%
    mutate(relative_ab = Count/sum(Count))

# Crypto 28
```

```r
crypto_28_18S <- read.table("~/Documents/crypto_data/9/18S_rRNA/Crypto-28_S28species_ct.txt"
    header = F, sep = "\t", stringsAsFactors = F)
colnames(crypto_28_18S) <- c("Species", "Locus", "Count")
# adding extra column
crypto_28_18S <- crypto_28_18S %>%
    mutate(Sample = "C_meleagridis_unenr")
# Calculate Relative abundance of counts for each genome
crypto_28_18S <- crypto_28_18S %>%
    group_by(Sample) %>%
    mutate(relative_ab = Count/sum(Count))

# Combine all samples
mixed_samples_18S <- rbind(crypto_8_18S, crypto_10_18S, crypto_26_18S,
    crypto_28_18S)

# plot
col_c_andersoni <- c("#8dd3c7")
col_c_baileyi <- c("#ffffb3")
col_c_canis <- c("#33a02c")
col_c_environmental <- c("#e31a1c")
col_c_meleagridis <- c("#80b1d3")
col_c_parvum <- c("#b3de69")
col_c_serpentis <- c("#1a1a1a")
col_c_sp <- c("#6a3d9a")
col_c_varanii <- c("#c51b7d")
col_c_18S_mix2 <- c("#8dd3c7", "#ffffb3", "#33a02c", "#e31a1c",
    "#80b1d3", "#b3de69", "#1a1a1a", "#6a3d9a", "#c51b7d")

plot <- mixed_samples_18S %>%
    group_by(Sample) %>%
    arrange(Species) %>%
    plot_ly(x = ~Sample, y = ~relative_ab, color = ~Species,
        colors = col_c_18S_mix2, type = "bar") %>%
    layout(yaxis = list(title = "Relative Abundance"), barmode = "stack")
```

**NOTE:** The results from these analyses are presented in **Fig. 1D**.

Most reads mapped to the correct species in the 18S rRNA database. Still, percentages were substantially misassigned (Fig. 1D). When the mock mixed-infection reads were mapped to the 18S rRNA database, 0.07% of retained reads mapped to the database for enriched libraries and 0.06% for unenriched. Regarding species assignment, *C. parvum* was properly assigned with a relative abundance of 75% for the unenriched sample and 70% for the enriched sample. *C. meleagridis* was properly assigned with a relative abundance of 66% for the unenriched sample and 73% for the enriched sample (Fig. 1D).

#### Supplementary Notes #10. Limits Of Detection: Minimum Genomic Input of *Cryptosporidium*

##### 10.1 Limits Of Detection: Minimum genomic input of *Cryptosporidium* with iTru libraries

We tested different levels of input DNA of C. parvum with serial dilutions to determine the limit of detection of target genomic DNA using both NEB-iTru and iNextEra libraries.

For this experiment, a 45-ng aliquot of calf thymus was pipetted into five tubes. In the first tube, 5 ng of *C. parvum* genomic DNA was added. Following this, a series of three 1:10 serial dilutions was performed using the first tube as a starter. The fifth tube corresponded to the negative control, which contained only calf thymus DNA. Summary results of these analyses can be found in **Table S11**, also imported below.

For NEB-iTru libraries, we obtained for the category with the highest input to the lowest input on average of 4,644,856, 1,422,483, 627,794, 481,962, 561,110, 616,115 paired reads per library, the negative control made from only complex background had on average 477,291 paired reads per library (Table S11). The number of paired reads was only statistically different between all the comparisons with the highest concentration (p-values $< 0.0001$). Similarly, the number of reads retained after trimming and quality filtering differed only between the highest concentration and the rest (p-values $< 0.0001$; Table S11).

```
file = "~/crypto_data/10/C_parvum_bait_dilution_NEB.txt" dil_neb = read.table(file, header =
TRUE, sep = "")
```

We assessed the number of raw paired reads obtained for each *C. parvum* dilution level.

```
lsmeans(lm(data = dil_neb, Paired_Raw_Reads ~ Category), pairwise ~
    Category)
```

```
## $lsmeans
##  Category  lsmean     SE df  lower.CL upper.CL
##  A        4644856 339000 35  3957214  5332498
##  B        1422483 339000 35   734841  2110125
##  C         627794 339000 35   -59848  1315436
##  D         481962 339000 35  -205680  1169604
##  E         561110 339000 35  -126533  1248752
##  F         616115 339000 35   -71527  1303757
##  G         477291 339000 35  -210351  1164934
##
## Confidence level used: 0.95
##
## $contrasts
##  contrast estimate     SE df t.ratio p.value
##  A - B     3222373 479000 35   6.727  <.0001
##  A - C     4017062 479000 35   8.386  <.0001
##  A - D     4162894 479000 35   8.690  <.0001
##  A - E     4083746 479000 35   8.525  <.0001
##  A - F     4028741 479000 35   8.410  <.0001
##  A - G     4167565 479000 35   8.700  <.0001
##  B - C      794689 479000 35   1.659  0.6465
##  B - D      940521 479000 35   1.963  0.4547
##  B - E      861373 479000 35   1.798  0.5580
##  B - F      806368 479000 35   1.683  0.6311
##  B - G      945192 479000 35   1.973  0.4488
##  C - D      145832 479000 35   0.304  0.9999
##  C - E       66684 479000 35   0.139  1.0000
##  C - F       11679 479000 35   0.024  1.0000
##  C - G      150503 479000 35   0.314  0.9999
##  D - E      -79148 479000 35  -0.165  1.0000
##  D - F     -134153 479000 35  -0.280  1.0000
##  D - G        4671 479000 35   0.010  1.0000
##  E - F      -55005 479000 35  -0.115  1.0000
##  E - G       83818 479000 35   0.175  1.0000
##  F - G      138824 479000 35   0.290  0.9999
##
```

```
## P value adjustment: tukey method for comparing a family of 7 estimates
```

```r
p <- plot_ly()
p <- add_boxplot(p, data = dil_neb, x = ~Category, y = ~Paired_Raw_Reads,
    color = I("#c51b7d"), type = "box", boxpoints = "all", jitter = 0.3,
    pointpos = 0, legendgroup = "1")
p <- layout(p, xaxis = list(tickmode = "array", tickvals = c("A",
    "sf", "B", "sf", "C", "sf", "D", "sf", "E", "sf", "F", "sf",
    "G"), ticktext = c("A", "", "B", "", "C", "", "D", "", "E",
    "", "F", "", "G")), yaxis = list(range = c(-1, 6e+06)))
```

Then, we assessed the number of retained reads retained after trimming for each dilution level of *C. parvum*

```r
lsmeans(lm(data = dil_neb, Paired_Trim_Reads ~ Category), pairwise ~ Category)
```

```
## $lsmeans
##  Category  lsmean     SE df lower.CL upper.CL
##  A        4395152 314000 35  3757799  5032505
##  B        1322603 314000 35   685251  1959956
##  C         558295 314000 35   -79058  1195648
##  D         417093 314000 35  -220260  1054445
##  E         486334 314000 35  -151019  1123686
##  F         533271 314000 35  -104082  1170623
##  G         411513 314000 35  -225839  1048866
##
## Confidence level used: 0.95
##
## $contrasts
##  contrast estimate     SE df t.ratio p.value
##  A - B     3072549 444000 35   6.920  <.0001
##  A - C     3836857 444000 35   8.642  <.0001
##  A - D     3978059 444000 35   8.960  <.0001
##  A - E     3908818 444000 35   8.804  <.0001
##  A - F     3861881 444000 35   8.698  <.0001
##  A - G     3983639 444000 35   8.972  <.0001
##  B - C      764308 444000 35   1.721  0.6069
##  B - D      905511 444000 35   2.039  0.4094
##  B - E      836270 444000 35   1.884  0.5040
##  B - F      789333 444000 35   1.778  0.5710
##  B - G      911090 444000 35   2.052  0.4021
##  C - D      141202 444000 35   0.318  0.9999
##  C - E       71961 444000 35   0.162  1.0000
##  C - F       25024 444000 35   0.056  1.0000
##  C - G      146782 444000 35   0.331  0.9999
##  D - E      -69241 444000 35  -0.156  1.0000
##  D - F     -116178 444000 35  -0.262  1.0000
##  D - G        5579 444000 35   0.013  1.0000
##  E - F      -46937 444000 35  -0.106  1.0000
##  E - G       74821 444000 35   0.169  1.0000
##  F - G      121757 444000 35   0.274  1.0000
##
## P value adjustment: tukey method for comparing a family of 7 estimates
```

We tested how the depth and breadth of genome coverage varied for the different dilutions.

```
lsmeans(lm(data = dil_neb, Per_Mapped_Cparvum ~ Category), pairwise ~
    Category)
```

```
## $lsmeans
##  Category lsmean   SE df lower.CL upper.CL
##  A         81.55 8.48 35     64.3     98.8
##  B         71.16 8.48 35     53.9     88.4
##  C         32.59 8.48 35     15.4     49.8
##  D          6.42 8.48 35    -10.8     23.6
##  E          1.79 8.48 35    -15.4     19.0
##  F          1.28 8.48 35    -15.9     18.5
##  G          1.19 8.48 35    -16.0     18.4
##
## Confidence level used: 0.95
##
## $contrasts
##  contrast estimate SE df t.ratio p.value
##  A - B     10.3917 12 35   0.867  0.9752
##  A - C     48.9583 12 35   4.084  0.0042
##  A - D     75.1250 12 35   6.266  <.0001
##  A - E     79.7533 12 35   6.652  <.0001
##  A - F     80.2650 12 35   6.695  <.0001
##  A - G     80.3617 12 35   6.703  <.0001
##  B - C     38.5667 12 35   3.217  0.0403
##  B - D     64.7333 12 35   5.399  0.0001
##  B - E     69.3617 12 35   5.785  <.0001
##  B - F     69.8733 12 35   5.828  <.0001
##  B - G     69.9700 12 35   5.836  <.0001
##  C - D     26.1667 12 35   2.183  0.3303
##  C - E     30.7950 12 35   2.569  0.1669
##  C - F     31.3067 12 35   2.611  0.1535
##  C - G     31.4033 12 35   2.619  0.1510
##  D - E      4.6283 12 35   0.386  0.9997
##  D - F      5.1400 12 35   0.429  0.9995
##  D - G      5.2367 12 35   0.437  0.9994
##  E - F      0.5117 12 35   0.043  1.0000
##  E - G      0.6083 12 35   0.051  1.0000
##  F - G      0.0967 12 35   0.008  1.0000
##
## P value adjustment: tukey method for comparing a family of 7 estimates
```

```
p <- plot_ly()
p <- add_boxplot(p, data = dil_neb, x = ~Category, y = ~Per_Mapped_Cparvum,
    color = I("#c51b7d"), type = "box", boxpoints = "all", jitter = 0.3,
    pointpos = 0, legendgroup = "1")
p <- layout(p, xaxis = list(tickmode = "array", tickvals = c("A",
    "sf", "B", "sf", "C", "sf", "D", "sf", "E", "sf", "F"), ticktext = c("A",
    "", "B", "", "C", "", "D", "", "E", "", "F")), yaxis = list(range = c(-1,
    100)))
```

**NOTE:** The results from these analyses are presented in **Fig. S18**.

The percentage of reads mapped per sample to *C. parvum* was the highest for the high 1 ng input, with an average of 81.6%. The next dilution of 0.1 ng had an average of 71.2% of reads mapping, the following dilution of 0.01 ng had a 32.6% average, and the following dilution of 0.001 ng had a 6.4% average. The two lowest inputs of 0.0001 ng and 0.00001 ng had below 2% of the reads mapped to *C. parvum* and the negative control. Statistically significant differences in the percentage of reads mapped were found between 1 ng and all the values of 0.01 ng and below (p-values  0.004) and 0.1 ng and all 0.01 ng and below (p-values  0.040, Fig. S18).

```
lsmeans(lm(data = dil_neb, Depth_Cparvum ~ Category), pairwise ~
    Category)
```

```
## $lsmeans
##  Category   lsmean   SE df lower.CL upper.CL
##  A        126.5950 10.1 35   106.14    147.0
##  B         29.2400 10.1 35     8.79     49.7
##  C          3.7333 10.1 35   -16.72     24.2
##  D          0.4517 10.1 35   -20.00     20.9
##  E          0.1050 10.1 35   -20.35     20.6
##  F          0.0667 10.1 35   -20.39     20.5
##  G          0.0433 10.1 35   -20.41     20.5
##
## Confidence level used: 0.95
##
## $contrasts
##  contrast estimate   SE df t.ratio p.value
##  A - B     97.3550 14.2 35   6.833  <.0001
##  A - C    122.8617 14.2 35   8.624  <.0001
##  A - D    126.1433 14.2 35   8.854  <.0001
##  A - E    126.4900 14.2 35   8.878  <.0001
##  A - F    126.5283 14.2 35   8.881  <.0001
##  A - G    126.5517 14.2 35   8.883  <.0001
##  B - C     25.5067 14.2 35   1.790  0.5630
##  B - D     28.7883 14.2 35   2.021  0.4204
##  B - E     29.1350 14.2 35   2.045  0.4062
##  B - F     29.1733 14.2 35   2.048  0.4046
##  B - G     29.1967 14.2 35   2.049  0.4037
##  C - D      3.2817 14.2 35   0.230  1.0000
##  C - E      3.6283 14.2 35   0.255  1.0000
##  C - F      3.6667 14.2 35   0.257  1.0000
##  C - G      3.6900 14.2 35   0.259  1.0000
##  D - E      0.3467 14.2 35   0.024  1.0000
##  D - F      0.3850 14.2 35   0.027  1.0000
##  D - G      0.4083 14.2 35   0.029  1.0000
##  E - F      0.0383 14.2 35   0.003  1.0000
##  E - G      0.0617 14.2 35   0.004  1.0000
##  F - G      0.0233 14.2 35   0.002  1.0000
##
## P value adjustment: tukey method for comparing a family of 7 estimates
```

```
depth <- plot_ly()
depth <- add_boxplot(depth, data = dil_neb, x = ~Category, y = ~Depth_Cparvum,
    color = I("#c51b7d"), type = "box", boxpoints = "all", jitter = 0.3,
    pointpos = 0, legendgroup = "1")
```

```r
depth <- layout(depth, xaxis = list(tickmode = "array", tickvals = c("A",
    "sf", "B", "sf", "C", "sf", "D", "sf", "E", "sf", "F", "sf",
    "G"), ticktext = c("A", "", "B", "", "C", "", "D", "", "E",
    "", "F", "", "G")), yaxis = list(range = c(-1, 180)))
lsmeans(lm(data = dil_neb, Breadth_Cparvum ~ Category), pairwise ~
    Category)
```

```
## $lsmeans
##  Category lsmean   SE df lower.CL upper.CL
##  A        89.088 8.34 35    72.15    106.0
##  B        81.362 8.34 35    64.42     98.3
##  C        54.830 8.34 35    37.89     71.8
##  D        13.758 8.34 35    -3.18     30.7
##  E         2.248 8.34 35   -14.69     19.2
##  F         0.550 8.34 35   -16.39     17.5
##  G         0.283 8.34 35   -16.66     17.2
##
## Confidence level used: 0.95
##
## $contrasts
##  contrast estimate   SE df t.ratio p.value
##  A - B       7.727 11.8 35   0.655  0.9942
##  A - C      34.258 11.8 35   2.903  0.0831
##  A - D      75.330 11.8 35   6.384  <.0001
##  A - E      86.840 11.8 35   7.360  <.0001
##  A - F      88.538 11.8 35   7.503  <.0001
##  A - G      88.805 11.8 35   7.526  <.0001
##  B - C      26.532 11.8 35   2.249  0.2970
##  B - D      67.603 11.8 35   5.729  <.0001
##  B - E      79.113 11.8 35   6.705  <.0001
##  B - F      80.812 11.8 35   6.849  <.0001
##  B - G      81.078 11.8 35   6.871  <.0001
##  C - D      41.072 11.8 35   3.481  0.0209
##  C - E      52.582 11.8 35   4.456  0.0015
##  C - F      54.280 11.8 35   4.600  0.0010
##  C - G      54.547 11.8 35   4.623  0.0009
##  D - E      11.510 11.8 35   0.975  0.9560
##  D - F      13.208 11.8 35   1.119  0.9178
##  D - G      13.475 11.8 35   1.142  0.9104
##  E - F       1.698 11.8 35   0.144  1.0000
##  E - G       1.965 11.8 35   0.167  1.0000
##  F - G       0.267 11.8 35   0.023  1.0000
##
## P value adjustment: tukey method for comparing a family of 7 estimates
```

```r
breadth <- plot_ly()
breadth <- add_boxplot(breadth, data = dil_neb, x = ~Category,
    y = ~Breadth_Cparvum, color = I("#c51b7d"), type = "box",
    boxpoints = "all", jitter = 0.3, pointpos = 0, legendgroup = "1")
breadth <- layout(breadth, xaxis = list(tickmode = "array", tickvals = c("A",
    "sf", "B", "sf", "C", "sf", "D", "sf", "E", "sf", "F", "sf",
    "G"), ticktext = c("A", "", "B", "", "C", "", "D", "", "E",
    "", "F", "", "G")), yaxis = list(range = c(-1, 110)))
```

```r
# creating subplot
fig <- subplot(depth, breadth, nrows = 1, titleY = TRUE, titleX = TRUE,
    margin = 0.1)
```

**NOTE:** The results from these analyses are presented in **Fig. S19**.

We tested how the depth and breadth of genome coverage varied for the different dilutions once we normalized for the number of sequences used in the mapping.

```r
lsmeans(lm(data = dil_neb, Normalized_Depth_Cparvum ~ Category),
    pairwise ~ Category)
```

```
## $lsmeans
##  Category lsmean    SE df lower.CL upper.CL
##  A        2.5250 0.262 35    1.993    3.057
##  B        2.1950 0.262 35    1.663    2.727
##  C        0.9983 0.262 35    0.467    1.530
##  D        0.1850 0.262 35   -0.347    0.717
##  E        0.0400 0.262 35   -0.492    0.572
##  F        0.0250 0.262 35   -0.507    0.557
##  G        0.0233 0.262 35   -0.508    0.555
##
## Confidence level used: 0.95
##
## $contrasts
##  contrast estimate   SE df t.ratio p.value
##  A - B     0.33000 0.37 35   0.891  0.9716
##  A - C     1.52667 0.37 35   4.121  0.0038
##  A - D     2.34000 0.37 35   6.317  <.0001
##  A - E     2.48500 0.37 35   6.708  <.0001
##  A - F     2.50000 0.37 35   6.749  <.0001
##  A - G     2.50167 0.37 35   6.753  <.0001
##  B - C     1.19667 0.37 35   3.230  0.0390
##  B - D     2.01000 0.37 35   5.426  0.0001
##  B - E     2.15500 0.37 35   5.817  <.0001
##  B - F     2.17000 0.37 35   5.858  <.0001
##  B - G     2.17167 0.37 35   5.862  <.0001
##  C - D     0.81333 0.37 35   2.196  0.3236
##  C - E     0.95833 0.37 35   2.587  0.1610
##  C - F     0.97333 0.37 35   2.627  0.1486
##  C - G     0.97500 0.37 35   2.632  0.1473
##  D - E     0.14500 0.37 35   0.391  0.9997
##  D - F     0.16000 0.37 35   0.432  0.9994
##  D - G     0.16167 0.37 35   0.436  0.9994
##  E - F     0.01500 0.37 35   0.040  1.0000
##  E - G     0.01667 0.37 35   0.045  1.0000
##  F - G     0.00167 0.37 35   0.004  1.0000
##
## P value adjustment: tukey method for comparing a family of 7 estimates
```

```r
depth <- plot_ly()
depth <- add_boxplot(depth, data = dil_neb, x = ~Category, y = ~Normalized_Depth_Cparvum,
    color = I("#c51b7d"), type = "box", boxpoints = "all", jitter = 0.3,
```

```
    pointpos = 0, legendgroup = "1")
depth <- layout(depth, xaxis = list(tickmode = "array", tickvals = c("A",
    "sf", "B", "sf", "C", "sf", "D", "sf", "E", "sf", "F", "sf",
    "G"), ticktext = c("A", "", "B", "", "C", "", "D", "", "E",
    "", "F", "", "G")), yaxis = list(range = c(-1, 4)))
lsmeans(lm(data = dil_neb, Normalized_Breadth ~ Category), pairwise ~
    Category)
```

```
## $lsmeans
##  Category lsmean   SE df lower.CL upper.CL
##  A         2.158 1.31 35    -0.51     4.83
##  B         6.223 1.31 35     3.55     8.89
##  C        14.670 1.31 35    12.00    17.34
##  D         5.535 1.31 35     2.87     8.20
##  E         0.877 1.31 35    -1.79     3.55
##  F         0.203 1.31 35    -2.47     2.87
##  G         0.150 1.31 35    -2.52     2.82
##
## Confidence level used: 0.95
##
## $contrasts
##  contrast estimate   SE df t.ratio p.value
##  A - B     -4.0650 1.86 35  -2.186  0.3283
##  A - C    -12.5117 1.86 35  -6.730  <.0001
##  A - D     -3.3767 1.86 35  -1.816  0.5465
##  A - E      1.2817 1.86 35   0.689  0.9924
##  A - F      1.9550 1.86 35   1.052  0.9378
##  A - G      2.0083 1.86 35   1.080  0.9298
##  B - C     -8.4467 1.86 35  -4.543  0.0011
##  B - D      0.6883 1.86 35   0.370  0.9998
##  B - E      5.3467 1.86 35   2.876  0.0883
##  B - F      6.0200 1.86 35   3.238  0.0383
##  B - G      6.0733 1.86 35   3.267  0.0357
##  C - D      9.1350 1.86 35   4.914  0.0004
##  C - E     13.7933 1.86 35   7.419  <.0001
##  C - F     14.4667 1.86 35   7.781  <.0001
##  C - G     14.5200 1.86 35   7.810  <.0001
##  D - E      4.6583 1.86 35   2.506  0.1883
##  D - F      5.3317 1.86 35   2.868  0.0899
##  D - G      5.3850 1.86 35   2.897  0.0844
##  E - F      0.6733 1.86 35   0.362  0.9998
##  E - G      0.7267 1.86 35   0.391  0.9997
##  F - G      0.0533 1.86 35   0.029  1.0000
##
## P value adjustment: tukey method for comparing a family of 7 estimates
```

```
breadth <- plot_ly()
breadth <- add_boxplot(breadth, data = dil_neb, x = ~Category,
    y = ~Normalized_Breadth, color = I("#c51b7d"), type = "box",
    boxpoints = "all", jitter = 0.3, pointpos = 0, legendgroup = "1")
breadth <- layout(breadth, xaxis = list(tickmode = "array", tickvals = c("A",
    "sf", "B", "sf", "C", "sf", "D", "sf", "E", "sf", "F", "sf",
    "G"), ticktext = c("A", "", "B", "", "C", "", "D", "", "E",
```

```
    "", "F", "", "G")), yaxis = list(range = c(-1, 20)))
# creating subplot
fig <- subplot(depth, breadth, nrows = 1, titleY = TRUE, titleX = TRUE,
    margin = 0.1)
```

**NOTE:** The results from these analyses are presented in **Fig. 2A**.

Then, we evaluated species assignments from the mapping to the ten-genomes database.

```
# Loop through the list of file names
data_c_parvum_list <- list()
cpar_list <- gsub(" ", "", dil_neb$Sample)
for (i in cpar_list) {
    # Construct the file path
    file_path <- paste("~/Documents/crypto_data/10/iTru_species_renamed/",
        i, "_species.txt", sep = "")

    # Check if the file exists and has data (not empty)
    if (file.exists(file_path) && file.info(file_path)$size >
        0) {
        # Read the file into a data frame
        data <- read.table(file_path, header = FALSE, sep = "\t",
            stringsAsFactors = FALSE)

        # Remove columns 3 to 5
        data <- data[, -c(3:5)]

        # Rename columns
        colnames(data) <- c("Species", "Count")

        # Calculate Relative abundance of counts for each
        # genome
        data <- data %>%
            mutate(relative_ab = Count/sum(Count))

        # Add the processed data frame to the list with the
        # file name as the list element name
        data_c_parvum_list[[i]] <- data
    }
}

# Combine the data frames in data_list into a single data
# frame
combined_data_cparvum <- bind_rows(data_c_parvum_list, .id = "Sample")

# Sort the data by Sample and Species
combined_data_cparvum <- combined_data_cparvum %>%
    arrange(Sample, Species)

# Create the plot
plot <- combined_data_cparvum %>%
    group_by(Sample) %>%
    arrange(Species) %>%
    plot_ly(x = ~Sample, y = ~relative_ab, color = ~Species,
```

```
        colors = col_c_ten, type = "bar") %>%
    layout(yaxis = list(title = "Relative Abundance"), barmode = "stack")
```

**NOTE:** The results from these analyses are presented in **Fig. 2B**.

The depth of genome coverage per 100,000 reads was below 1X for 0.01 ng, 2.2X for 0.1 ng, and 2.5X for 1 ng, with significant differences between 1 ng and 0.1 ng and all other dilutions (p-values < 0.0001; Fig. 2A; non-normalized values Fig. S19). The breadth of genome coverage, normalized per 100,000 reads, did not correlate with target input (Fig. 2A), with significant differences observed only between 0.01 ng and all other dilutions (non-normalized values Fig. S19). For unenriched libraries, accurate species assignment is observed only for the 1 ng and 0.1 ng inputs (Fig. 2B). For single enrichments, *C. parvum* is accurately detected in inputs 0.001 ng (Fig. 2B).

##### 10.2 Limits of Detection: Minimum genomic input of *C. parvum* with iNextEra libraries

The summary results of these analyses can be found in **Table S12**, and are imported below.

For the *Cryptosporidium* dilution experiment performed with iNextEra libraries, we observed that the number of raw reads obtained varies depending on whether the experiment was unenriched, single-captured, or double-captured. We observed a trend where double enrichment treatments obtained fewer reads, on average 1128,803, followed by unenriched treatments with 2,196,810 paired reads per library on average, and single enrichments with 6,115,285 reads on average (Table S12).

```
file = "~/crypto_data/10/C_par_dilution_iNextEra.txt" dil_inext = read.table(file, header
= TRUE, sep = "")
```

We assessed the number of raw paired reads obtained for each dilution level of *C. parvum*

```
lsmeans(lm(data = dil_inext, Raw_Read_Pairs ~ Category), pairwise ~
    Category)
```

```
## $lsmeans
##  Category  lsmean      SE df  lower.CL upper.CL
##  A        5457696 1830000 28  1709805  9205588
##  B        4831591 1490000 28  1771450  7891731
##  C        3130268 1490000 28    70127  6190408
##  D        3025721 1490000 28   -34420  6085861
##  E        3251745 1490000 28   191604  6311886
##  G        3715136 1490000 28   654995  6775277
##
## Confidence level used: 0.95
##
## $contrasts
##  contrast estimate      SE df t.ratio p.value
##  A - B      626106 2360000 28   0.265  0.9998
##  A - C     2327429 2360000 28   0.985  0.9188
##  A - D     2431976 2360000 28   1.030  0.9039
##  A - E     2205951 2360000 28   0.934  0.9342
##  A - G     1742560 2360000 28   0.738  0.9754
##  B - C     1701323 2110000 28   0.805  0.9642
##  B - D     1805870 2110000 28   0.855  0.9540
##  B - E     1579846 2110000 28   0.748  0.9739
```

```
## B - G 1116454 2110000 28 0.528 0.9945
## C - D 104547 2110000 28 0.049 1.0000
## C - E -121478 2110000 28 -0.057 1.0000
## C - G -584868 2110000 28 -0.277 0.9998
## D - E -226024 2110000 28 -0.107 1.0000
## D - G -689416 2110000 28 -0.326 0.9995
## E - G -463391 2110000 28 -0.219 0.9999
##
## P value adjustment: tukey method for comparing a family of 6 estimates
```

```r
lsmeans(lm(data = dil_inext, Raw_Read_Pairs ~ Library), pairwise ~
    Library)
```

```
## $lsmeans
##  Library     lsmean      SE df lower.CL upper.CL
##  Double     1128803  795000 31  -492199  2749804
##  Single     6115285  639000 31  4811352  7419218
##  Unenriched 2196810 1080000 31     1963  4391656
##
## Confidence level used: 0.95
##
## $contrasts
##  contrast            estimate      SE df t.ratio p.value
##  Double - Single     -4986482 1020000 31  -4.889  0.0001
##  Double - Unenriched -1068007 1340000 31  -0.798  0.7068
##  Single - Unenriched  3918476 1250000 31   3.130  0.0103
##
## P value adjustment: tukey method for comparing a family of 3 estimates
```

```r
my_palette2 <- c("#fdae61", "#1b9e77", "#7570b3")
fig <- plot_ly(colors = my_palette2)
fig <- add_boxplot(fig, data = dil_inext, x = ~Category, y = ~Raw_Read_Pairs,
    color = ~Library, type = "box")
fig <- fig %>%
    layout(boxmode = "group")
```

Then, we assessed the similarity between the percentage of retained reads that mapped to the *C. parvum* genome for each dilution level.

```r
lsmeans(lm(data = dil_inext, Per_Map_Paired_Reads_Cparvum ~ Category),
    pairwise ~ Category)
```

```
## $lsmeans
##  Category lsmean    SE df lower.CL upper.CL
##  A        55.392 10.00 28    34.89     75.9
##  B        35.625  8.17 28    18.89     52.4
##  C         9.425  8.17 28    -7.31     26.2
##  D         1.497  8.17 28   -15.24     18.2
##  E         0.397  8.17 28   -16.34     17.1
##  G         0.030  8.17 28   -16.71     16.8
##
## Confidence level used: 0.95
```

```
##
## $contrasts
##  contrast estimate   SE df t.ratio p.value
##  A - B      19.767 12.9 28   1.530  0.6488
##  A - C      45.968 12.9 28   3.558  0.0154
##  A - D      53.896 12.9 28   4.171  0.0033
##  A - E      54.996 12.9 28   4.256  0.0026
##  A - G      55.362 12.9 28   4.285  0.0024
##  B - C      26.200 11.6 28   2.267  0.2408
##  B - D      34.128 11.6 28   2.953  0.0627
##  B - E      35.228 11.6 28   3.048  0.0509
##  B - G      35.595 11.6 28   3.080  0.0474
##  C - D       7.928 11.6 28   0.686  0.9821
##  C - E       9.028 11.6 28   0.781  0.9685
##  C - G       9.395 11.6 28   0.813  0.9627
##  D - E       1.100 11.6 28   0.095  1.0000
##  D - G       1.467 11.6 28   0.127  1.0000
##  E - G       0.367 11.6 28   0.032  1.0000
##
## P value adjustment: tukey method for comparing a family of 6 estimates
```

```
lsmeans(lm(data = dil_inext, Per_Map_Paired_Reads_Cparvum ~ Library),
    pairwise ~ Library)
```

```
## $lsmeans
##  Library    lsmean    SE df lower.CL upper.CL
##  Double      27.55  7.84 31    11.56     43.5
##  Single      11.75  6.31 31    -1.11     24.6
##  Unenriched   0.11 10.60 31   -21.54     21.8
##
## Confidence level used: 0.95
##
## $contrasts
##  contrast            estimate   SE df t.ratio p.value
##  Double - Single         15.8 10.1 31   1.570  0.2735
##  Double - Unenriched     27.4 13.2 31   2.079  0.1108
##  Single - Unenriched     11.6 12.3 31   0.943  0.6179
##
## P value adjustment: tukey method for comparing a family of 3 estimates
```

```
fig <- plot_ly(colors = my_palette2)
fig <- add_boxplot(fig, data = dil_inext, x = ~Category, y = ~Per_Map_Paired_Reads_Cparvum,
    color = ~Library, type = "box")
fig <- fig %>%
    layout(boxmode = "group")
```

**NOTE:** The results from these analyses are presented in **Fig. S20**.

We then investigated the depth and breadth of genome coverage for *C. parvum*.

```
lsmeans(lm(data = dil_inext, Average_Depth_Cparvum ~ Category),
    pairwise ~ Category)
```

```
## $lsmeans
##  Category  lsmean   SE df lower.CL upper.CL
##  A        72.2943 20.6 28   30.051    114.5
##  B        34.7965 16.8 28    0.305     69.3
##  C         3.7023 16.8 28  -30.789     38.2
##  D         0.4273 16.8 28  -34.064     34.9
##  E         0.1325 16.8 28  -34.359     34.6
##  G         0.0575 16.8 28  -34.434     34.5
##
## Confidence level used: 0.95
##
## $contrasts
##  contrast estimate   SE df t.ratio p.value
##  A - B      37.498 26.6 28   1.408  0.7218
##  A - C      68.592 26.6 28   2.576  0.1368
##  A - D      71.867 26.6 28   2.699  0.1071
##  A - E      72.162 26.6 28   2.710  0.1047
##  A - G      72.237 26.6 28   2.713  0.1041
##  B - C      31.094 23.8 28   1.306  0.7794
##  B - D      34.369 23.8 28   1.443  0.7012
##  B - E      34.664 23.8 28   1.456  0.6938
##  B - G      34.739 23.8 28   1.459  0.6919
##  C - D       3.275 23.8 28   0.138  1.0000
##  C - E       3.570 23.8 28   0.150  1.0000
##  C - G       3.645 23.8 28   0.153  1.0000
##  D - E       0.295 23.8 28   0.012  1.0000
##  D - G       0.370 23.8 28   0.016  1.0000
##  E - G       0.075 23.8 28   0.003  1.0000
##
## P value adjustment: tukey method for comparing a family of 6 estimates
```

```
depth <- plot_ly(colors = my_palette2)
depth <- add_boxplot(depth, data = dil_inext, x = ~Category,
    y = ~Average_Depth_Cparvum, color = I("#fdb462"), type = "box",
    boxpoints = "all", jitter = 0.3, pointpos = 0, legendgroup = "1")
depth <- layout(depth, xaxis = list(tickmode = "array", tickvals = c("A",
    "sf", "B", "sf", "C", "sf", "D", "sf", "E", "sf", "G"), ticktext = c("A",
    "", "B", "", "C", "", "D", "", "E", "", "G")), yaxis = list(range = c(-1,
    250)))
lsmeans(lm(data = dil_inext, Average_Breadth_Cparvum ~ Category),
    pairwise ~ Category)
```

```
## $lsmeans
##  Category lsmean    SE df lower.CL upper.CL
##  A        56.968 11.90 28     32.5     81.4
##  B        58.233  9.75 28     38.3     78.2
##  C        18.372  9.75 28     -1.6     38.3
##  D         2.785  9.75 28    -17.2     22.8
##  E         1.405  9.75 28    -18.6     21.4
##  G         0.143  9.75 28    -19.8     20.1
##
## Confidence level used: 0.95
##
## $contrasts
```

```
##  contrast estimate   SE df t.ratio p.value
##  A - B       -1.27 15.4 28  -0.082  1.0000
##  A - C       38.60 15.4 28   2.503  0.1574
##  A - D       54.18 15.4 28   3.514  0.0171
##  A - E       55.56 15.4 28   3.604  0.0138
##  A - G       56.82 15.4 28   3.685  0.0113
##  B - C       39.86 13.8 28   2.890  0.0719
##  B - D       55.45 13.8 28   4.021  0.0048
##  B - E       56.83 13.8 28   4.121  0.0037
##  B - G       58.09 13.8 28   4.212  0.0029
##  C - D       15.59 13.8 28   1.130  0.8646
##  C - E       16.97 13.8 28   1.230  0.8184
##  C - G       18.23 13.8 28   1.322  0.7708
##  D - E        1.38 13.8 28   0.100  1.0000
##  D - G        2.64 13.8 28   0.192  1.0000
##  E - G        1.26 13.8 28   0.091  1.0000
##
## P value adjustment: tukey method for comparing a family of 6 estimates
```

```
breadth <- plot_ly()
breadth <- add_boxplot(breadth, data = dil_inext, x = ~Category,
    y = ~Average_Breadth_Cparvum, color = I("#fdb462"), type = "box",
    boxpoints = "all", jitter = 0.3, pointpos = 0, legendgroup = "1")
breadth <- layout(breadth, xaxis = list(tickmode = "array", tickvals = c("A",
    "sf", "B", "sf", "C", "sf", "D", "sf", "E", "sf", "G"), ticktext = c("A",
    "", "B", "", "C", "", "D", "", "E", "", "G")), yaxis = list(range = c(-1,
    110)))
# creating subplot
fig <- subplot(depth, breadth, nrows = 1, titleY = TRUE, titleX = TRUE,
    margin = 0.1)
```

**NOTE:** The results from these analyses are presented in **Fig. 2C**.

We then investigated the depth and breadth of genome coverage for *C. parvum* normalized by the number of retained reads.

```
lsmeans(lm(data = dil_inext, Normalized_Depth_Cparvum ~ Category),
    pairwise ~ Category)
```

```
## $lsmeans
##  Category  lsmean    SE df lower.CL upper.CL
##  A        1.47700 0.341 28    0.779    2.175
##  B        1.04667 0.278 28    0.477    1.617
##  C        0.29517 0.278 28   -0.275    0.865
##  D        0.04817 0.278 28   -0.522    0.618
##  E        0.01383 0.278 28   -0.556    0.584
##  G        0.00167 0.278 28   -0.568    0.572
##
## Confidence level used: 0.95
##
## $contrasts
##  contrast estimate    SE df t.ratio p.value
##  A - B      0.4303 0.440 28   0.978  0.9212
```

```
##   A - C      1.1818 0.440 28   2.686  0.1101
##   A - D      1.4288 0.440 28   3.247  0.0323
##   A - E      1.4632 0.440 28   3.325  0.0270
##   A - G      1.4753 0.440 28   3.352  0.0252
##   B - C      0.7515 0.394 28   1.909  0.4177
##   B - D      0.9985 0.394 28   2.537  0.1476
##   B - E      1.0328 0.394 28   2.624  0.1246
##   B - G      1.0450 0.394 28   2.655  0.1171
##   C - D      0.2470 0.394 28   0.628  0.9880
##   C - E      0.2813 0.394 28   0.715  0.9785
##   C - G      0.2935 0.394 28   0.746  0.9742
##   D - E      0.0343 0.394 28   0.087  1.0000
##   D - G      0.0465 0.394 28   0.118  1.0000
##   E - G      0.0122 0.394 28   0.031  1.0000
##
## P value adjustment: tukey method for comparing a family of 6 estimates
```

```
depth <- plot_ly(colors = my_palette2)
depth <- add_boxplot(depth, data = dil_inext, x = ~Category,
    y = ~Normalized_Depth_Cparvum, color = I("#fdb462"), type = "box",
    boxpoints = "all", jitter = 0.3, pointpos = 0, legendgroup = "1")
depth <- layout(depth, xaxis = list(tickmode = "array", tickvals = c("A",
    "sf", "B", "sf", "C", "sf", "D", "sf", "E", "sf", "G"), ticktext = c("A",
    "", "B", "", "C", "", "D", "", "E", "", "G")), yaxis = list(range = c(-1,
    4)))
lsmeans(lm(data = dil_inext, Normalized_Breadth ~ Category),
    pairwise ~ Category)
```

```
## $lsmeans
##   Category  lsmean    SE df lower.CL upper.CL
##   A        2.44375 0.389 28    1.647    3.240
##   B        1.51250 0.318 28    0.862    2.163
##   C        0.82350 0.318 28    0.173    1.474
##   D        0.12250 0.318 28   -0.528    0.773
##   E        0.23767 0.318 28   -0.413    0.888
##   G        0.00767 0.318 28   -0.643    0.658
##
## Confidence level used: 0.95
##
## $contrasts
##   contrast estimate    SE df t.ratio p.value
##   A - B       0.931 0.502 28   1.855  0.4493
##   A - C       1.620 0.502 28   3.227  0.0339
##   A - D       2.321 0.502 28   4.623  0.0010
##   A - E       2.206 0.502 28   4.394  0.0018
##   A - G       2.436 0.502 28   4.852  0.0005
##   B - C       0.689 0.449 28   1.534  0.6461
##   B - D       1.390 0.449 28   3.095  0.0458
##   B - E       1.275 0.449 28   2.839  0.0802
##   B - G       1.505 0.449 28   3.351  0.0254
##   C - D       0.701 0.449 28   1.561  0.6296
##   C - E       0.586 0.449 28   1.304  0.7802
##   C - G       0.816 0.449 28   1.817  0.4719
##   D - E      -0.115 0.449 28  -0.256  0.9998
```

```
##   D - G       0.115 0.449 28   0.256  0.9998
##   E - G       0.230 0.449 28   0.512  0.9953
##
## P value adjustment: tukey method for comparing a family of 6 estimates
```

```
breadth <- plot_ly()
breadth <- add_boxplot(breadth, data = dil_inext, x = ~Category,
    y = ~Normalized_Breadth, color = I("#fdb462"), type = "box",
    boxpoints = "all", jitter = 0.3, pointpos = 0, legendgroup = "1")
breadth <- layout(breadth, xaxis = list(tickmode = "array", tickvals = c("A",
    "sf", "B", "sf", "C", "sf", "D", "sf", "E", "sf", "G"), ticktext = c("A",
    "", "B", "", "C", "", "D", "", "E", "", "G")), yaxis = list(range = c(-1,
    5)))
# creating subplot
fig <- subplot(depth, breadth, nrows = 1, titleY = TRUE, titleX = TRUE,
    margin = 0.1)
```

**NOTE:** The results from these analyses are presented in **Fig. S21**.

We estimated the percentage of reads that mapped to *C. parvum* reference genome. We found that the average percentage for negative control libraries was 0.03%, and it increased with the different input levels, from 0.40% for 0.0001 ng, 1.50% for 0.001 ng, 9.42% for 0.01 ng, 35.63% for 0.1 ng, and 55.39% for 1 ng on average per library (Fig S20). These percentages resulted in statistical differences in the comparisons of 1 ng with all other inputs, except with 0.1 ng (p-values  0.015). Other marginally significant differences were found between the 0.1 ng input and 0.001 ng and 0.0001 ng (p-values = 0.05). These percentage values were not affected by the type of library, nor the bait dilution used (p-values  0.111). When normalizing by 100,000 retained reads, depth and breadth are significantly higher in 1 ng inputs compared to < 0.01 ng inputs (p-values  0.034, Fig. S21), and significantly higher for 0.1 ng compared to 0.001 ng and the negative control (p-values  0.046).

Then, we evaluated species assignment from the mapping to the ten-genomes database.

```
# Create an empty list to store the data frames
data_c_parvum_list <- list()
cpar_list <- gsub(" ", "", dil_inext$New_label)
# Loop through the list of file names in 'elist'
for (i in cpar_list) {
    # Construct the file path
    file_path <- paste("~/crypto_data/10/iNextEra_species_renamed/",
        i, "_species.txt", sep = "")

    # Check if the file exists and has data (not empty)
    if (file.exists(file_path) && file.info(file_path)$size >
        0) {
        # Read the file into a data frame
        data <- read.table(file_path, header = FALSE, sep = "\t",
            stringsAsFactors = FALSE)

        # Remove columns 3 to 5
        data <- data[, -c(3:5)]

        # Rename columns
        colnames(data) <- c("Species", "Count")
```

```r
        # Calculate Relative abundance of counts for each
        # genome
        data <- data %>%
            mutate(relative_ab = Count/sum(Count))

        # Add the processed data frame to the list with the
        # file name as the list element name
        data_c_parvum_list[[i]] <- data
    }
}

# Combine the data frames in data_list into a single data
# frame
combined_data_cparvum <- bind_rows(data_c_parvum_list, .id = "Sample")

# Sort the data by Sample and Species
combined_data_cparvum <- combined_data_cparvum %>%
    arrange(Sample, Species)

# Create the plot
plot <- combined_data_cparvum %>%
    group_by(Sample) %>%
    arrange(Species) %>%
    plot_ly(x = ~Sample, y = ~relative_ab, color = ~Species,
        colors = col_c_ten, type = "bar") %>%
    layout(yaxis = list(title = "Relative Abundance"), barmode = "stack")
```

**NOTE:** The results from these analyses are presented in **Fig. 2D**.

We also assessed the percentage of retained reads that mapped to the *Bos taurus* genome for each dilution level.

```r
lsmeans(lm(data = dil_inext, Per_Map_Paired_Reads_Bos_taurus ~
    Category), pairwise ~ Category)
```

```
## $lsmeans
##  Category lsmean   SE df lower.CL upper.CL
##  A          35.2 9.39 28     16.0     54.5
##  B          54.5 7.67 28     38.8     70.2
##  C          76.9 7.67 28     61.2     92.6
##  D          83.7 7.67 28     68.0     99.4
##  E          84.3 7.67 28     68.6    100.0
##  G          85.1 7.67 28     69.4    100.8
##
## Confidence level used: 0.95
##
## $contrasts
##  contrast estimate   SE df t.ratio p.value
##  A - B     -19.242 12.1 28  -1.587  0.6133
##  A - C     -41.646 12.1 28  -3.435  0.0207
##  A - D     -48.461 12.1 28  -3.997  0.0051
##  A - E     -49.064 12.1 28  -4.047  0.0045
##  A - G     -49.874 12.1 28  -4.114  0.0038
```

```
##   B - C     -22.403 10.8 28  -2.066  0.3331
##   B - D     -29.218 10.8 28  -2.694  0.1082
##   B - E     -29.822 10.8 28  -2.750  0.0966
##   B - G     -30.632 10.8 28  -2.825  0.0826
##   C - D      -6.815 10.8 28  -0.628  0.9879
##   C - E      -7.418 10.8 28  -0.684  0.9823
##   C - G      -8.228 10.8 28  -0.759  0.9722
##   D - E      -0.603 10.8 28  -0.056  1.0000
##   D - G      -1.413 10.8 28  -0.130  1.0000
##   E - G      -0.810 10.8 28  -0.075  1.0000
##
## P value adjustment: tukey method for comparing a family of 6 estimates
```

```
lsmeans(lm(data = dil_inext, Per_Map_Paired_Reads_Bos_taurus ~
    Library), pairwise ~ Library)
```

```
## $lsmeans
##   Library    lsmean   SE df lower.CL upper.CL
##   Double       59.8 6.92 31     45.7     73.9
##   Single       73.4 5.57 31     62.0     84.7
##   Unenriched   90.4 9.37 31     71.3    109.5
##
## Confidence level used: 0.95
##
## $contrasts
##   contrast            estimate    SE df t.ratio p.value
##   Double - Single        -13.6  8.88 31  -1.535  0.2889
##   Double - Unenriched    -30.6 11.60 31  -2.628  0.0344
##   Single - Unenriched    -17.0 10.90 31  -1.558  0.2788
##
## P value adjustment: tukey method for comparing a family of 3 estimates
```

```
fig <- plot_ly(colors = my_palette2)
fig <- add_boxplot(fig, data = dil_inext, x = ~Category, y = ~Per_Map_Paired_Reads_Bos_taurus,
    color = ~Library, type = "box")
fig <- fig %>%
    layout(boxmode = "group")
```

**NOTE:** The results from these analyses are presented in **Fig. S22**.

The average percentage of reads mapped to the *Bos taurus* genome sequence decreased as the *C. parvum* input increased in enriched libraries, given that *C. parvum* DNA was diluted in calf thymus DNA. In unenriched libraries, percentages of mapping were overall similarly high across all dilutions, with an average of 90.5% of sequencing reads mapping to *Bos taurus* (Table S12, Fig. S22). For enriched libraries, the percentage of reads mapping to *Bos taurus* was, on average, 35.2% for samples with 1 ng of *C. parvum*, 54.5% for samples with 0.1 ng of *C. parvum*, 76.9% for samples with 0.01 ng of *C. parvum*, 83.7% for samples with 0.001 ng of *C. parvum*, 84.7% for samples with 0.0001 ng of *C. parvum*, and 85.1% for the negative controls (Fig. S22). The percentage of mapped reads was only significantly lower for inputs of 1 ng vs. all the other input values (0.004   p-values   0.021).

### Supplementary Notes #11. Limits of Detection: Minimum CryptoCap_100k 2.0 Bait Set Input

#### 11.1 Limits of Detection: Minimum CryptoCap_100k bait set input with NEB libraries with iTru primers

```r
lsmeans(lm(data = dil_neb, Paired_Raw_Reads ~ Bait_Dilution),
    pairwise ~ Bait_Dilution)
```

```
## $lsmeans
##  Bait_Dilution  lsmean     SE df lower.CL upper.CL
##  01_Full       1324530 649000 36     9000  2640061
##  02_Half       1258041 649000 36   -57490  2573571
##  03_Quarter    1161069 649000 36  -154462  2476599
##  04_Eighth     1089351 649000 36  -226179  2404882
##  05_Sixteenth  1027866 649000 36  -287664  2343397
##  06_Unenriched 1709095 649000 36   393565  3024626
##
## Confidence level used: 0.95
##
## $contrasts
##  contrast                     estimate     SE df t.ratio p.value
##  01_Full - 02_Half               66489 917000 36   0.072  1.0000
##  01_Full - 03_Quarter           163461 917000 36   0.178  1.0000
##  01_Full - 04_Eighth            235179 917000 36   0.256  0.9998
##  01_Full - 05_Sixteenth         296664 917000 36   0.323  0.9995
##  01_Full - 06_Unenriched       -384565 917000 36  -0.419  0.9982
##  02_Half - 03_Quarter            96972 917000 36   0.106  1.0000
##  02_Half - 04_Eighth            168690 917000 36   0.184  1.0000
##  02_Half - 05_Sixteenth         230175 917000 36   0.251  0.9999
##  02_Half - 06_Unenriched       -451054 917000 36  -0.492  0.9962
##  03_Quarter - 04_Eighth          71718 917000 36   0.078  1.0000
##  03_Quarter - 05_Sixteenth      133202 917000 36   0.145  1.0000
##  03_Quarter - 06_Unenriched    -548026 917000 36  -0.597  0.9906
##  04_Eighth - 05_Sixteenth        61485 917000 36   0.067  1.0000
##  04_Eighth - 06_Unenriched     -619744 917000 36  -0.676  0.9836
##  05_Sixteenth - 06_Unenriched  -681229 917000 36  -0.743  0.9751
##
## P value adjustment: tukey method for comparing a family of 6 estimates
```

```r
p <- plot_ly()
p <- add_boxplot(p, data = dil_neb, x = ~Bait_Dilution, y = ~Paired_Raw_Reads,
    color = I("#c51b7d"), type = "box", boxpoints = "all", jitter = 0.3,
    pointpos = 0, legendgroup = "1")
p <- layout(p, xaxis = list(tickmode = "array", tickvals = c("01_Full",
    "sf", "02_Half", "sf", "03_Quarter", "sf", "04_Eighth", "sf",
    "05_Sixteenth", "sf", "06_Unenriched"), ticktext = c("01_Full",
    "", "02_Half", "", "03_Quarter", "", "04_Eighth", "", "05_Sixteenth",
    "", "06_Unenriched")), yaxis = list(range = c(-1, 6e+06)))
```

**NOTE:** The results from these analyses are presented in **Fig. S23**.

We assessed the percentage of retained reads that mapped to the *C. parvum* genome for each bait dilution level.

```
lsmeans(lm(data = dil_neb, Per_Mapped_Cparvum ~ Bait_Dilution),
    pairwise ~ Bait_Dilution)
```

```
## $lsmeans
##  Bait_Dilution lsmean   SE df lower.CL upper.CL
##  01_Full       33.521 14.4 36     4.22     62.8
##  02_Half       34.324 14.4 36     5.02     63.6
##  03_Quarter    34.406 14.4 36     5.10     63.7
##  04_Eighth     30.864 14.4 36     1.56     60.2
##  05_Sixteenth  34.519 14.4 36     5.21     63.8
##  06_Unenriched  0.341 14.4 36   -28.96     29.6
##
## Confidence level used: 0.95
##
## $contrasts
##  contrast                     estimate   SE df t.ratio p.value
##  01_Full - 02_Half             -0.8029 20.4 36  -0.039  1.0000
##  01_Full - 03_Quarter          -0.8843 20.4 36  -0.043  1.0000
##  01_Full - 04_Eighth            2.6571 20.4 36   0.130  1.0000
##  01_Full - 05_Sixteenth        -0.9971 20.4 36  -0.049  1.0000
##  01_Full - 06_Unenriched       33.1800 20.4 36   1.624  0.5889
##  02_Half - 03_Quarter          -0.0814 20.4 36  -0.004  1.0000
##  02_Half - 04_Eighth            3.4600 20.4 36   0.169  1.0000
##  02_Half - 05_Sixteenth        -0.1943 20.4 36  -0.010  1.0000
##  02_Half - 06_Unenriched       33.9829 20.4 36   1.663  0.5640
##  03_Quarter - 04_Eighth         3.5414 20.4 36   0.173  1.0000
##  03_Quarter - 05_Sixteenth     -0.1129 20.4 36  -0.006  1.0000
##  03_Quarter - 06_Unenriched    34.0643 20.4 36   1.667  0.5615
##  04_Eighth - 05_Sixteenth      -3.6543 20.4 36  -0.179  1.0000
##  04_Eighth - 06_Unenriched     30.5229 20.4 36   1.494  0.6704
##  05_Sixteenth - 06_Unenriched  34.1771 20.4 36   1.672  0.5580
##
## P value adjustment: tukey method for comparing a family of 6 estimates
```

```
p <- plot_ly()
p <- add_boxplot(p, data = dil_neb, x = ~Bait_Dilution, y = ~Per_Mapped_Cparvum,
    color = I("#c51b7d"), type = "box", boxpoints = "all", jitter = 0.3,
    pointpos = 0, legendgroup = "1")
p <- layout(p, xaxis = list(tickmode = "array", tickvals = c("01_Full",
    "sf", "02_Half", "sf", "03_Quarter", "sf", "04_Eighth", "sf",
    "05_Sixteenth", "sf", "06_Unenriched"), ticktext = c("01_Full",
    "", "02_Half", "", "03_Quarter", "", "04_Eighth", "", "05_Sixteenth",
    "", "06_Unenriched")), yaxis = list(range = c(-1, 100)))
```

**NOTE:** The results from these analyses are presented in **Fig. S24**.

We assessed depth of genome coverage at each bait dilution level.

```
lsmeans(lm(data = dil_neb, Depth_Cparvum ~ Bait_Dilution), pairwise ~
    Bait_Dilution)
```

```
## $lsmeans
##  Bait_Dilution lsmean   SE df lower.CL upper.CL
##  01_Full       30.849 19.5 36    -8.75     70.4
##  02_Half       29.817 19.5 36    -9.78     69.4
##  03_Quarter    27.884 19.5 36   -11.71     67.5
##  04_Eighth     23.503 19.5 36   -16.09     63.1
##  05_Sixteenth  25.170 19.5 36   -14.43     64.8
##  06_Unenriched  0.121 19.5 36   -39.47     39.7
##
## Confidence level used: 0.95
##
## $contrasts
##  contrast                    estimate   SE df t.ratio p.value
##  01_Full - 02_Half               1.03 27.6 36   0.037  1.0000
##  01_Full - 03_Quarter            2.96 27.6 36   0.107  1.0000
##  01_Full - 04_Eighth             7.35 27.6 36   0.266  0.9998
##  01_Full - 05_Sixteenth          5.68 27.6 36   0.206  0.9999
##  01_Full - 06_Unenriched        30.73 27.6 36   1.113  0.8728
##  02_Half - 03_Quarter            1.93 27.6 36   0.070  1.0000
##  02_Half - 04_Eighth             6.31 27.6 36   0.229  0.9999
##  02_Half - 05_Sixteenth          4.65 27.6 36   0.168  1.0000
##  02_Half - 06_Unenriched        29.70 27.6 36   1.076  0.8878
##  03_Quarter - 04_Eighth          4.38 27.6 36   0.159  1.0000
##  03_Quarter - 05_Sixteenth       2.71 27.6 36   0.098  1.0000
##  03_Quarter - 06_Unenriched     27.76 27.6 36   1.006  0.9131
##  04_Eighth - 05_Sixteenth       -1.67 27.6 36  -0.060  1.0000
##  04_Eighth - 06_Unenriched      23.38 27.6 36   0.847  0.9564
##  05_Sixteenth - 06_Unenriched   25.05 27.6 36   0.907  0.9421
##
## P value adjustment: tukey method for comparing a family of 6 estimates
```

```r
depth <- plot_ly(colors = my_palette2)
depth <- add_boxplot(depth, data = dil_neb, x = ~Bait_Dilution,
    y = ~Depth_Cparvum, color = I("#c51b7d"), type = "box", boxpoints = "all",
    jitter = 0.3, pointpos = 0, legendgroup = "1")
depth <- layout(depth, xaxis = list(tickmode = "array", tickvals = c("01_Full",
    "sf", "02_Half", "sf", "03_Quarter", "sf", "04_Eighth", "sf",
    "05_Sixteenth", "sf", "06_Unenriched"), ticktext = c("01_Full",
    "", "02_Half", "", "03_Quarter", "", "04_Eighth", "", "05_Sixteenth",
    "", "06_Unenriched", "", "G")), yaxis = list(range = c(-1,
    180)))
lsmeans(lm(data = dil_neb, Breadth_Cparvum ~ Bait_Dilution),
    pairwise ~ Bait_Dilution)
```

```
## $lsmeans
##  Bait_Dilution lsmean   SE df lower.CL upper.CL
##  01_Full        41.88 15.9 36     9.60     74.2
##  02_Half        42.53 15.9 36    10.25     74.8
##  03_Quarter     40.78 15.9 36     8.49     73.1
##  04_Eighth      39.22 15.9 36     6.94     71.5
##  05_Sixteenth   37.24 15.9 36     4.95     69.5
##  06_Unenriched   5.88 15.9 36   -26.41     38.2
##
## Confidence level used: 0.95
```

```
##
## $contrasts
##  contrast                     estimate   SE df t.ratio p.value
##  01_Full - 02_Half               -0.65 22.5 36  -0.029  1.0000
##  01_Full - 03_Quarter             1.11 22.5 36   0.049  1.0000
##  01_Full - 04_Eighth              2.66 22.5 36   0.118  1.0000
##  01_Full - 05_Sixteenth           4.65 22.5 36   0.206  0.9999
##  01_Full - 06_Unenriched         36.01 22.5 36   1.599  0.6043
##  02_Half - 03_Quarter             1.76 22.5 36   0.078  1.0000
##  02_Half - 04_Eighth              3.31 22.5 36   0.147  1.0000
##  02_Half - 05_Sixteenth           5.30 22.5 36   0.235  0.9999
##  02_Half - 06_Unenriched         36.66 22.5 36   1.628  0.5860
##  03_Quarter - 04_Eighth           1.55 22.5 36   0.069  1.0000
##  03_Quarter - 05_Sixteenth        3.54 22.5 36   0.157  1.0000
##  03_Quarter - 06_Unenriched      34.90 22.5 36   1.550  0.6353
##  04_Eighth - 05_Sixteenth         1.98 22.5 36   0.088  1.0000
##  04_Eighth - 06_Unenriched       33.34 22.5 36   1.481  0.6781
##  05_Sixteenth - 06_Unenriched    31.36 22.5 36   1.393  0.7308
##
## P value adjustment: tukey method for comparing a family of 6 estimates
```

```
breadth <- plot_ly(colors = my_palette2)
breadth <- add_boxplot(breadth, data = dil_neb, x = ~Bait_Dilution,
    y = ~Breadth_Cparvum, color = I("#c51b7d"), type = "box",
    boxpoints = "all", jitter = 0.3, pointpos = 0, legendgroup = "1")
breadth <- layout(breadth, xaxis = list(tickmode = "array", tickvals = c("01_Full",
    "sf", "02_Half", "sf", "03_Quarter", "sf", "04_Eighth", "sf",
    "05_Sixteenth", "sf", "06_Unenriched"), ticktext = c("01_Full",
    "", "02_Half", "", "03_Quarter", "", "04_Eighth", "", "05_Sixteenth",
    "", "06_Unenriched", "", "G")), yaxis = list(range = c(-1,
    110)))
# creating subplot
fig <- subplot(depth, breadth, nrows = 1, titleY = TRUE, titleX = TRUE,
    margin = 0.1)
```

**NOTE:** The results from these analyses are presented in **Fig. S25**.

##### 11.2 Limits of Detection: Minimum CryptoCap_100k bait set input with iNextEra libraries and iNext primers

```
lsmeans(lm(data = dil_inext, Raw_Read_Pairs ~ Bait_Dilution),
    pairwise ~ Bait_Dilution)
```

```
## $lsmeans
##  Bait_Dilution  lsmean      SE df lower.CL upper.CL
##  01_Quarter    6035566 1380000 31  3219899  8851234
##  02_Eighth     3643786  721000 31  2173351  5114220
##  03_Unenriched 2196810 1380000 31  -618858  5012478
##
## Confidence level used: 0.95
##
```

```
## $contrasts
##  contrast                   estimate      SE df t.ratio p.value
##  01_Quarter - 02_Eighth      2391781 1560000 31   1.536  0.2886
##  01_Quarter - 03_Unenriched  3838756 1950000 31   1.966  0.1376
##  02_Eighth - 03_Unenriched   1446976 1560000 31   0.929  0.6263
##
## P value adjustment: tukey method for comparing a family of 3 estimates
```

```
lsmeans(lm(data = dil_inext, Raw_Read_Pairs ~ Library.Bait.dilution),
    pairwise ~ Library.Bait.dilution)
```

```
## $lsmeans
##  Library.Bait.dilution  lsmean      SE df lower.CL upper.CL
##  Double-One-eighth     1128803  808000 30  -520997  2778603
##  Single-One-eighth     6158768  808000 30  4508968  7808569
##  Single-One-quarter    6035566 1090000 30  3801726  8269406
##  Unenriched            2196810 1090000 30   -37030  4430650
##
## Confidence level used: 0.95
##
## $contrasts
##  contrast                                  estimate      SE df t.ratio p.value
##  (Double-One-eighth) - (Single-One-eighth)  -5029966 1140000 30  -4.403  0.0007
##  (Double-One-eighth) - (Single-One-quarter) -4906763 1360000 30  -3.609  0.0058
##  (Double-One-eighth) - Unenriched           -1068007 1360000 30  -0.785  0.8605
##  (Single-One-eighth) - (Single-One-quarter)   123202 1360000 30   0.091  0.9997
##  (Single-One-eighth) - Unenriched            3961959 1360000 30   2.914  0.0321
##  (Single-One-quarter) - Unenriched           3838756 1550000 30   2.482  0.0835
##
## P value adjustment: tukey method for comparing a family of 4 estimates
```

```
fig <- plot_ly()
fig <- add_boxplot(fig, data = dil_inext, x = ~Bait_Dilution,
    y = ~Raw_Read_Pairs, color = I("#fdb462"), type = "box")
fig <- fig %>%
    layout(boxmode = "group")
```

**NOTE:** The results from these analyses are presented in **Fig. S26**.

We also investigated the percentage of reads mapped:

```
lsmeans(lm(data = dil_inext, Per_Map_Paired_Reads_Cparvum ~ Bait_Dilution),
    pairwise ~ Bait_Dilution)
```

```
## $lsmeans
##  Bait_Dilution lsmean    SE df lower.CL upper.CL
##  01_Quarter     18.04 11.00 31    -4.45     40.5
##  02_Eighth      17.93  5.76 31     6.18     29.7
##  03_Unenriched   0.11 11.00 31   -22.39     22.6
##
## Confidence level used: 0.95
##
## $contrasts
```

```
##   contrast                     estimate   SE df t.ratio p.value
##   01_Quarter - 02_Eighth          0.11 12.4 31   0.009  1.0000
##   01_Quarter - 03_Unenriched     17.93 15.6 31   1.150  0.4916
##   02_Eighth - 03_Unenriched      17.82 12.4 31   1.432  0.3372
##
## P value adjustment: tukey method for comparing a family of 3 estimates
```

```
p <- plot_ly()
p <- add_boxplot(p, data = dil_inext, x = ~Bait_Dilution, y = ~Per_Map_Paired_Reads_Cparvum,
    color = I("#fdb462"), type = "box", boxpoints = "all", jitter = 0.3,
    pointpos = 0, legendgroup = "1")
```

**NOTE:** The results from these analyses are presented in **Fig. S27**.

We then investigated the depth and breadth of genome coverage for *C. parvum*.

```
lsmeans(lm(data = dil_inext, Average_Depth_Cparvum ~ Bait_Dilution),
    pairwise ~ Bait_Dilution)
```

```
## $lsmeans
##   Bait_Dilution lsmean    SE df lower.CL upper.CL
##   01_Quarter    43.868 18.10 31     6.86     80.9
##   02_Eighth     11.840  9.48 31    -7.49     31.2
##   03_Unenriched  0.029 18.10 31   -36.98     37.0
##
## Confidence level used: 0.95
##
## $contrasts
##   contrast                     estimate   SE df t.ratio p.value
##   01_Quarter - 02_Eighth           32.0 20.5 31   1.564  0.2759
##   01_Quarter - 03_Unenriched       43.8 25.7 31   1.708  0.2182
##   02_Eighth - 03_Unenriched        11.8 20.5 31   0.577  0.8334
##
## P value adjustment: tukey method for comparing a family of 3 estimates
```

```
depth <- plot_ly(colors = my_palette2)
depth <- add_boxplot(depth, data = dil_inext, x = ~Bait_Dilution,
    y = ~Average_Depth_Cparvum, color = I("#fdb462"), type = "box",
    boxpoints = "all", jitter = 0.3, pointpos = 0, legendgroup = "1")
depth <- layout(depth, xaxis = list(tickmode = "array", tickvals = c("01_Quarter",
    "sf", "02_Eighth", "sf", "03_Unenriched"), ticktext = c("01_Quarter",
    "", "02_Eighth", "", "03_Unenriched")), yaxis = list(range = c(-1,
    250)))
lsmeans(lm(data = dil_inext, Average_Breadth_Cparvum ~ Bait_Dilution),
    pairwise ~ Bait_Dilution)
```

```
## $lsmeans
##   Bait_Dilution lsmean   SE df lower.CL upper.CL
##   01_Quarter     41.67 13.0 31    15.10     68.2
##   02_Eighth      20.64  6.8 31     6.77     34.5
##   03_Unenriched   1.55 13.0 31   -25.02     28.1
##
## Confidence level used: 0.95
```

```
##
## $contrasts
##  contrast                    estimate   SE df t.ratio p.value
##  01_Quarter - 02_Eighth          21.0 14.7 31   1.431  0.3379
##  01_Quarter - 03_Unenriched      40.1 18.4 31   2.177  0.0911
##  02_Eighth - 03_Unenriched       19.1 14.7 31   1.299  0.4065
##
## P value adjustment: tukey method for comparing a family of 3 estimates
```

```
breadth <- plot_ly(colors = my_palette2)
breadth <- add_boxplot(breadth, data = dil_inext, x = ~Bait_Dilution,
    y = ~Average_Breadth_Cparvum, color = I("#fdb462"), type = "box",
    boxpoints = "all", jitter = 0.3, pointpos = 0, legendgroup = "1")
breadth <- layout(breadth, xaxis = list(tickmode = "array", tickvals = c("01_Quarter",
    "sf", "02_Eighth", "sf", "03_Unenriched"), ticktext = c("01_Quarter",
    "", "02_Eighth", "", "03_Unenriched")), yaxis = list(range = c(-1,
    110)))
# creating subplot
fig <- subplot(depth, breadth, nrows = 1, titleY = TRUE, titleX = TRUE,
    margin = 0.1)
```

**NOTE:** The results from these analyses are presented in **Fig. S28**.

We then investigated the depth and breadth of genome coverage for *C. parvum* when normalized by retained reads.

```
lsmeans(lm(data = dil_inext, Normalized_Depth_Cparvum ~ Bait_Dilution),
    pairwise ~ Bait_Dilution)
```

```
## $lsmeans
##  Bait_Dilution  lsmean    SE df lower.CL upper.CL
##  01_Quarter    0.41050 0.340 31   -0.282    1.103
##  02_Eighth     0.53891 0.177 31    0.177    0.901
##  03_Unenriched 0.00367 0.340 31   -0.689    0.696
##
## Confidence level used: 0.95
##
## $contrasts
##  contrast                    estimate    SE df t.ratio p.value
##  01_Quarter - 02_Eighth        -0.128 0.383 31  -0.335  0.9401
##  01_Quarter - 03_Unenriched     0.407 0.480 31   0.847  0.6770
##  02_Eighth - 03_Unenriched      0.535 0.383 31   1.397  0.3547
##
## P value adjustment: tukey method for comparing a family of 3 estimates
```

```
depth <- plot_ly(colors = my_palette2)
depth <- add_boxplot(depth, data = dil_inext, x = ~Bait_Dilution,
    y = ~Normalized_Depth_Cparvum, color = I("#fdb462"), type = "box",
    boxpoints = "all", jitter = 0.3, pointpos = 0, legendgroup = "1")
depth <- layout(depth, xaxis = list(tickmode = "array", tickvals = c("01_Quarter",
    "sf", "02_Eighth", "sf", "03_Unenriched"), ticktext = c("01_Quarter",
    "", "02_Eighth", "", "03_Unenriched")), yaxis = list(range = c(-1,
    4)))
```

```r
lsmeans(lm(data = dil_inext, Normalized_Breadth ~ Bait_Dilution),
    pairwise ~ Bait_Dilution)
```

```
## $lsmeans
##  Bait_Dilution lsmean    SE df lower.CL upper.CL
##  01_Quarter     0.983 0.447 31   0.0717     1.89
##  02_Eighth      0.845 0.233 31   0.3689     1.32
##  03_Unenriched  0.251 0.447 31  -0.6605     1.16
##
## Confidence level used: 0.95
##
## $contrasts
##  contrast                   estimate    SE df t.ratio p.value
##  01_Quarter - 02_Eighth        0.138 0.504 31   0.274  0.9594
##  01_Quarter - 03_Unenriched    0.732 0.632 31   1.158  0.4865
##  02_Eighth - 03_Unenriched     0.594 0.504 31   1.178  0.4751
##
## P value adjustment: tukey method for comparing a family of 3 estimates
```

```r
breadth <- plot_ly(colors = my_palette2)
breadth <- add_boxplot(breadth, data = dil_inext, x = ~Bait_Dilution,
    y = ~Normalized_Breadth, color = I("#fdb462"), type = "box",
    boxpoints = "all", jitter = 0.3, pointpos = 0, legendgroup = "1")
breadth <- layout(breadth, xaxis = list(tickmode = "array", tickvals = c("01_Quarter",
    "sf", "02_Eighth", "sf", "03_Unenriched"), ticktext = c("01_Quarter",
    "", "02_Eighth", "", "03_Unenriched")), yaxis = list(range = c(-1,
    6)))
# creating subplot
fig <- subplot(depth, breadth, nrows = 1, titleY = TRUE, titleX = TRUE,
    margin = 0.1)
```

**NOTE:** The results from these analyses are presented in **Fig. S29**.

Raw reads obtained were similar for both bait dilutions and unenriched. The percentage of mapped reads was correlated with the bait input amount, averaging ~18% for the enriched libraries and 0.11% for unenriched libraries. The depth and breadth of *C. parvum* genome coverage were, on average, 43.9X and 41.67% for libraries enriched with quarter dilutions of the baits, 11.8X, and 20.64% for libraries enriched with an eighth dilution of the baits, vs. 0.03X and 1.55% for unenriched libraries.

#### Supplementary Notes #12. In Vitro Enrichment and Validation with CryptoCap_100k Baits in Clinical Samples

##### 12.1 Set of 7 clinical samples

Similarly to simulated sequencing reads, we started by mapping each sample sequenced against the fasta containing ten genomes. We used BBmap and as reference we used the reference file created above. We used a maxim indel length of 20, a minid of 0.8 (read note below), we also required correct strands for paired reads (rcs), set the mean pair size to 400, and did not rescue paired reads if avg. insert size greater than 600 (rescuedist; let's remember that the simulated size is 500), and we set an average pair distance *(apd)of 100 bases.

Then, we obtained the hits per sample. For each, we used the FLAG count from the third column of our SAM files to extract those sequences for which both reads mapped (FLAG: 83 and 99). Then, for each sample, we counted each species ID hit in our mapped file. The count text has two columns: the first is the species ID, and the second is the counts.

Before importing each file from each sample, we separated the first column in Excel, delimiting by "-", separating the added species name. To begin with, each file has two columns; the first one contains the reference genome information, and the second is the count of hits that the simulated reads had against target regions of each species. We split the first column into two to have in the first the "Species" name to which the reads hit, and the second is "Locus" with the rest of the information of the reference genome where the reads mapped. Summary results of these analyses are found in **Table S13**, also imported below.

```
file = "~/crypto_data/12/seven_clinical_samples_summary_stats.txt" cl_seven_table = read.table(file,
header = TRUE, sep = "")
```

We tested whether the number of raw paired reads obtained for the seven clinical samples from enriched CryptoCap_100k and unenriched samples was different.

```
lsmeans(lm(data = cl_seven_table, RawReadPairs ~ Library), pairwise ~
    Library)
```

```
## $lsmeans
##  Library    lsmean    SE df lower.CL upper.CL
##  Enriched   116757 50200 12     7416   226098
##  Unenriched 221769 50200 12   112428   331110
##
## Confidence level used: 0.95
##
## $contrasts
##  contrast              estimate    SE df t.ratio p.value
##  Enriched - Unenriched  -105012 71000 12  -1.480  0.1647
```

```
p <- plot_ly()
p <- add_bars(p, x = c("Unenriched", "Unenriched0", "Enriched"),
    y = c(5e+05, 570000, 5e+05), opacity = 1, showlegend = F,
    marker = list(line = list(color = "rgba(0,0,0,0"), color = "rgba(0,0,0,0"),
    text = c("", "p-value = 0.165", ""), textposition = "outside",
    legendgroup = "1")
p <- add_lines(p, x = c("Unenriched", "Unenriched", "Enriched",
    "Enriched"), y = c(550000, 540000, 540000, 550000), showlegend = F,
    line = list(color = "black"), legendgroup = "1", hoverinfo = "none")
p <- add_boxplot(p, data = cl_seven_table, x = ~Library, y = ~RawReadPairs,
    color = ~Library, colormodel = c("rgba(197,27,125"), type = "box",
    boxpoints = "all", jitter = 0.3, pointpos = 0, legendgroup = "1")
p <- layout(p, xaxis = list(tickmode = "array", tickvals = c("Unenriched",
    "sf", "Enriched"), ticktext = c("Unenriched", "", "Enriched")),
    yaxis = list(range = c(-1, 6e+05)))
```

**NOTE:** The results from these analyses are presented in **Fig. S30**.

We tested if the average percentage of reads retained after filtering was different between single and double captures with CryptoCap_100k.

```
lsmeans(lm(data = cl_seven_table, Trimmed_Reads ~ Library), pairwise ~ Library)
```

```
## $lsmeans
##  Library    lsmean    SE df lower.CL upper.CL
##  Enriched   111719 46700 12     9997   213441
##  Unenriched 201337 46700 12    99615   303059
##
## Confidence level used: 0.95
##
## $contrasts
##  contrast              estimate    SE df t.ratio p.value
##  Enriched - Unenriched   -89618 66000 12  -1.357  0.1997
```

For libraries from DNA obtained from seven clinical patients' fecal samples, a similar average number of reads was obtained for unenriched and enriched libraries (Table S13, Fig. S30).

**12.1.1 Sequence data mapped to the 10-genomes reference database** We also tested if the number of reads that mapped to the ten-genome reference after filtering were different between single and double captures.

```r
lsmeans(lm(data = cl_seven_table, MappedPairedReads10RefGenomes ~ Library), pairwise ~
    Library)
```

```
## $lsmeans
##  Library    lsmean   SE df lower.CL upper.CL
##  Enriched   5571.7 1440 12     2426     8718
##  Unenriched   67.7 1440 12    -3078     3214
##
## Confidence level used: 0.95
##
## $contrasts
##  contrast              estimate   SE df t.ratio p.value
##  Enriched - Unenriched     5504 2040 12   2.696  0.0195
```

We also tested if the percentage of reads mapped to *C.parvum* were different between single and double captures.

```r
lsmeans(lm(data = cl_seven_table, PerMapPairedReads10RefGenomes ~
    Library), pairwise ~ Library)
```

```
## $lsmeans
##  Library    lsmean   SE df lower.CL upper.CL
##  Enriched   7.1471 1.66 12     3.53    10.76
##  Unenriched 0.0471 1.66 12    -3.57     3.66
##
## Confidence level used: 0.95
##
## $contrasts
##  contrast              estimate   SE df t.ratio p.value
##  Enriched - Unenriched      7.1 2.35 12   3.026  0.0105
```

```
p <- plot_ly()
p <- add_boxplot(p, data = cl_seven_table, x = ~Sample, y = ~PerMapPairedReads10RefGenomes,
    color = ~Library, boxpoints = "all", jitter = 0.3, pointpos = 0,
    legendgroup = "1")
p <- layout(p, xaxis = list(tickmode = "array", tickvals = c("Single",
    "sf", "Double"), ticktext = c("Single", "", "Double")), yaxis = list(range = c(-1,
    20)), barmode = "stack")
```

**NOTE:** The results from these analyses are presented in **Fig. S31**.

When mapped to the ten-genome reference, the number and percentage of trimmed reads from enriched samples exceeded those from unenriched samples. Enriched samples had, on average, 5,572 paired reads, representing an average of 7.15% of trimmed reads mapped to the reference. In contrast, unenriched samples had an average of 68 paired reads mapping, representing 0.05% of trimmed reads mapping to the reference. Both values were statistically different for both library types (number of mapped reads p-value = 0.019, percentage of reads mapped p-value = 0.011; Figs. S30-S31).

Below, we imported the modified files for each species and then created a fourth column "Sample" with the name of the species for which we simulated the reads.

```
# Crypto 12
crypto_12 <- read.table("~/crypto_data/12/Crypto-12_species.txt",
    header = F, sep = "\t", stringsAsFactors = F)
colnames(crypto_12) <- c("Species", "Count")
# adding extra column
crypto_12 <- crypto_12 %>%
    mutate(Sample = "enriched DOK1")
# Calculate Relative abundance of counts for each genome
crypto_12 <- crypto_12 %>%
    group_by(Sample) %>%
    mutate(relative_ab = Count/sum(Count))

# Crypto 13
crypto_13 <- read.table("~/crypto_data/12/Crypto-13_species.txt",
    header = F, sep = "\t", stringsAsFactors = F)
colnames(crypto_13) <- c("Species", "Count")
# adding extra column
crypto_13 <- crypto_13 %>%
    mutate(Sample = "enriched DOK2")
# Calculate Relative abundance of counts for each genome
crypto_13 <- crypto_13 %>%
    group_by(Sample) %>%
    mutate(relative_ab = Count/sum(Count))

# Crypto 14
crypto_14 <- read.table("~/crypto_data/12/Crypto-14_species.txt",
    header = F, sep = "\t", stringsAsFactors = F)
colnames(crypto_14) <- c("Species", "Count")
# adding extra column
crypto_14 <- crypto_14 %>%
    mutate(Sample = "enriched DOK3")
# Calculate Relative abundance of counts for each genome
crypto_14 <- crypto_14 %>%
    group_by(Sample) %>%
```

```
    mutate(relative_ab = Count/sum(Count))

# Crypto 15
crypto_15 <- read.table("~/crypto_data/12/Crypto-15_species.txt",
    header = F, sep = "\t", stringsAsFactors = F)
colnames(crypto_15) <- c("Species", "Count")
# adding extra column
crypto_15 <- crypto_15 %>%
    mutate(Sample = "enriched DOK4")
# Calculate Relative abundance of counts for each genome
crypto_15 <- crypto_15 %>%
    group_by(Sample) %>%
    mutate(relative_ab = Count/sum(Count))

# Crypto 16
crypto_16 <- read.table("~/crypto_data/12/Crypto-16_species.txt",
    header = F, sep = "\t", stringsAsFactors = F)
colnames(crypto_16) <- c("Species", "Count")
# adding extra column
crypto_16 <- crypto_16 %>%
    mutate(Sample = "enriched UK101")
# Calculate Relative abundance of counts for each genome
crypto_16 <- crypto_16 %>%
    group_by(Sample) %>%
    mutate(relative_ab = Count/sum(Count))

# Crypto 17
crypto_17 <- read.table("~/crypto_data/12/Crypto-17_species.txt",
    header = F, sep = "\t", stringsAsFactors = F)
colnames(crypto_17) <- c("Species", "Count")
# adding extra column
crypto_17 <- crypto_17 %>%
    mutate(Sample = "enriched UK196")
# Calculate Relative abundance of counts for each genome
crypto_17 <- crypto_17 %>%
    group_by(Sample) %>%
    mutate(relative_ab = Count/sum(Count))

# Crypto 18
crypto_18 <- read.table("~/crypto_data/12/Crypto-18_species.txt",
    header = F, sep = "\t", stringsAsFactors = F)
colnames(crypto_18) <- c("Species", "Count")
# adding extra column
crypto_18 <- crypto_18 %>%
    mutate(Sample = "enriched EC4")
# Calculate Relative abundance of counts for each genome
crypto_18 <- crypto_18 %>%
    group_by(Sample) %>%
    mutate(relative_ab = Count/sum(Count))

# Crypto 30
crypto_30 <- read.table("~/crypto_data/12/Crypto-30_species.txt",
    header = F, sep = "\t", stringsAsFactors = F)
```

```r
colnames(crypto_30) <- c("Species", "Count")
# adding extra column
crypto_30 <- crypto_30 %>%
    mutate(Sample = "unenriched DOK1")
# Calculate Relative abundance of counts for each genome
crypto_30 <- crypto_30 %>%
    group_by(Sample) %>%
    mutate(relative_ab = Count/sum(Count))

# Crypto 31
crypto_31 <- read.table("~/crypto_data/12/Crypto-31_species.txt",
    header = F, sep = "\t", stringsAsFactors = F)
colnames(crypto_31) <- c("Species", "Count")
# adding extra column
crypto_31 <- crypto_31 %>%
    mutate(Sample = "unenriched DOK2")
# Calculate Relative abundance of counts for each genome
crypto_31 <- crypto_31 %>%
    group_by(Sample) %>%
    mutate(relative_ab = Count/sum(Count))

# Crypto 32
crypto_32 <- read.table("~/crypto_data/12/Crypto-32_species.txt",
    header = F, sep = "\t", stringsAsFactors = F)
colnames(crypto_32) <- c("Species", "Count")
# adding extra column
crypto_32 <- crypto_32 %>%
    mutate(Sample = "unenriched DOK3")
# Calculate Relative abundance of counts for each genome
crypto_32 <- crypto_32 %>%
    group_by(Sample) %>%
    mutate(relative_ab = Count/sum(Count))

# Crypto 33
crypto_33 <- read.table("~/crypto_data/12/Crypto-33_species.txt",
    header = F, sep = "\t", stringsAsFactors = F)
colnames(crypto_33) <- c("Species", "Count")
# adding extra column
crypto_33 <- crypto_33 %>%
    mutate(Sample = "unenriched DOK4")
# Calculate Relative abundance of counts for each genome
crypto_33 <- crypto_33 %>%
    group_by(Sample) %>%
    mutate(relative_ab = Count/sum(Count))

# Crypto 34
crypto_34 <- read.table("~/crypto_data/12/Crypto-34_species.txt",
    header = F, sep = "\t", stringsAsFactors = F)
colnames(crypto_34) <- c("Species", "Count")
# adding extra column
crypto_34 <- crypto_34 %>%
    mutate(Sample = "unenriched UK101")
# Calculate Relative abundance of counts for each genome
```

```r
crypto_34 <- crypto_34 %>%
    group_by(Sample) %>%
    mutate(relative_ab = Count/sum(Count))

# Crypto 35
crypto_35 <- read.table("~/crypto_data/12/Crypto-35_species.txt",
    header = F, sep = "\t", stringsAsFactors = F)
colnames(crypto_35) <- c("Species", "Count")
# adding extra column
crypto_35 <- crypto_35 %>%
    mutate(Sample = "unenriched UK196")
# Calculate Relative abundance of counts for each genome
crypto_35 <- crypto_35 %>%
    group_by(Sample) %>%
    mutate(relative_ab = Count/sum(Count))

# Crypto 36
crypto_36 <- read.table("~/crypto_data/12/Crypto-36_species.txt",
    header = F, sep = "\t", stringsAsFactors = F)
colnames(crypto_36) <- c("Species", "Count")
# adding extra column
crypto_36 <- crypto_36 %>%
    mutate(Sample = "unenriched EC4")
# Calculate Relative abundance of counts for each genome
crypto_36 <- crypto_36 %>%
    group_by(Sample) %>%
    mutate(relative_ab = Count/sum(Count))

# Combine all samples
clinical_samples <- rbind(crypto_12, crypto_13, crypto_14, crypto_15,
    crypto_16, crypto_17, crypto_18, crypto_30, crypto_31, crypto_32,
    crypto_33, crypto_34, crypto_35, crypto_36)
clinical_samples
```

```
## # A tibble: 519 x 4
## # Groups:   Sample [14]
##    Species         Count Sample        relative_ab
##    <chr>           <int> <chr>               <dbl>
##  1 C_hominis_30976  2779 enriched DOK1    0.681
##  2 C_baileyi           5 enriched DOK1    0.00122
##  3 C_ubiquitum        17 enriched DOK1    0.00416
##  4 C_cuniculus         4 enriched DOK1    0.000980
##  5 C_muris            40 enriched DOK1    0.00980
##  6 C_muris            10 enriched DOK1    0.00245
##  7 C_hominis_30976    27 enriched DOK1    0.00661
##  8 C_hominis_30976    65 enriched DOK1    0.0159
##  9 C_parvum           22 enriched DOK1    0.00539
## 10 C_muris            32 enriched DOK1    0.00784
## # i 509 more rows
```

```r
plot <- clinical_samples %>%
    group_by(Sample) %>%
    arrange(Species) %>%
```

```
    plot_ly(x = ~Sample, y = ~relative_ab, color = ~Species,
        colors = col_c_ten, type = "bar") %>%
    layout(yaxis = list(title = "Relative Abundance"), barmode = "stack")
```

**NOTE:** The results from these analyses are presented in **Fig. S32**.

Using the ten-genomes reference, the species assignment shows a drastic differential between enriched and unenriched samples (Fig. S32). Species assignment of all clinical samples from unenriched libraries shows high abundances of *C. muris* (Fig. S32, right). The assignments for the enriched libraries for the same samples show that most samples contain in their majority *C. hominis* followed by *C. cuniculus*, except for samples EC4 and UKP196, which contain high proportions of *C. parvum* (Fig. S32, left).

**12.1.2 Sequence data mapped to the gp60 reference database** We mapped reads using similar parameters as above, except pairlen = 400 and maxindel = 5, and using as references independently one gp60 fasta file containing 65 gp60 record:

```
bbmap.sh in=1.fq in2=2.fq \
ref GP60new.fasta/  \
maxindel=5 minid=0.80 rcs=t killbadpairs=t pairlen=400 \
rescuedist=600 apd=100 outu=mixed_infections/mapped_gp60/\$i\_unmapped.sam \
outm=mixed_infections/mapped_gp60/\$i\_mapped.sam printunmappedcount=t \
covstats=mixed_infections/mapped_gp60/\$i\.cov.txt threads=12 -Xmx180g 2>&1 \
| tee mixed_infections/mapped_gp60/\$i\.bbmap.log
```

None of the sequencing reads from the 14 samples analyzed here had hits against the gp60 reference database.

**12.1.3 Sequence data mapped to the CryptoDB 18S rRNA database.** We mapped reads using similar parameters as above, except pairlen = 500 and maxindel = 5, and using as references the fasta file containing 57 reference sequences from: https://cryptodb.org/cryptodb/app/downloads/Current_Release/CryptosporidiidaeReference/fasta/ (version last modified on 2022-04-13 11:11), here referred to as 18S_CDB:

```
bbmap.sh in=1.fq in2=2.fq \
ref 18S_reference.fasta  \
maxindel=5 minid=0.80 rcs=t killbadpairs=t pairlen=400 \
rescuedist=600 apd=100 outu=sim_mapped/\$i\_unmapped.sam \
outm=sim_mapped/\$i\_mapped.sam printunmappedcount=t \
covstats=sim_mapped/\$i\.cov.txt threads=12 -Xmx180g 2>&1 \
| tee sim_mapped/\$i\.bbmap.log
```

We can observe that only three enriched samples mapped to the database, and none of the unenriched samples.

We imported species count files to plot species assignments based on the 18S rRNA database.

```
# Crypto 15
crypto_15_18S <- read.table("~/crypto_data/12/18S_rRNA/Crypto-15_S15species_ct.txt"
    header = F, sep = "\t", stringsAsFactors = F)
colnames(crypto_15_18S) <- c("Species", "Locus", "Count")
# adding extra column
crypto_15_18S <- crypto_15_18S %>%
    mutate(Sample = "enriched DOK4")
# Calculate Relative abundance of counts for each genome
```

```r
crypto_15_18S <- crypto_15_18S %>%
    group_by(Sample) %>%
    mutate(relative_ab = Count/sum(Count))

# Crypto 16
crypto_16_18S <- read.table("~/crypto_data/12/18S_rRNA/Crypto-16_S16species_ct.txt"
    header = F, sep = "\t", stringsAsFactors = F)
colnames(crypto_16_18S) <- c("Species", "Locus", "Count")
# adding extra column
crypto_16_18S <- crypto_16_18S %>%
    mutate(Sample = "enriched UK101")
# Calculate Relative abundance of counts for each genome
crypto_16_18S <- crypto_16_18S %>%
    group_by(Sample) %>%
    mutate(relative_ab = Count/sum(Count))

# Crypto 17
crypto_17_18S <- read.table("~/crypto_data/12/18S_rRNA/Crypto-17_S17species_ct.txt"
    header = F, sep = "\t", stringsAsFactors = F)
colnames(crypto_17_18S) <- c("Species", "Locus", "Count")
# adding extra column
crypto_17_18S <- crypto_17_18S %>%
    mutate(Sample = "enriched UK196")
# Calculate Relative abundance of counts for each genome
crypto_17_18S <- crypto_17_18S %>%
    group_by(Sample) %>%
    mutate(relative_ab = Count/sum(Count))

# Combine all samples
clinical_samples_18S <- rbind(crypto_15_18S, crypto_16_18S, crypto_17_18S)
clinical_samples_18S
```

```
## # A tibble: 5 x 5
## # Groups:   Sample [3]
##   Species                    Locus                          Count Sample relative_ab
##   <chr>                      <chr>                          <int> <chr>        <dbl>
## 1 Cryptosporidium_sp         AY504515 | organism=Crypto~     8 enric~       1
## 2 Cryptosporidium_sp         DQ295013 | organism=Crypto~     1 enric~       1
## 3 Cryptosporidium_andersoni AB089285 | organism=Crypto~     5 enric~       0.263
## 4 Cryptosporidium_canis      AB210854 | organism=Crypto~    12 enric~       0.632
## 5 Cryptosporidium_serpentis AF093502 | organism=Crypto~     2 enric~       0.105
```

```r
# plot
col_c_andersoni <- c("#8dd3c7")
col_c_canis <- c("#33a02c")
col_c_serpentis <- c("#1a1a1a")
col_c_sp <- c("#6a3d9a")
col_c_18S_mix3 <- c("#8dd3c7", "#33a02c", "#1a1a1a", "#6a3d9a")

plot <- clinical_samples_18S %>%
    group_by(Sample) %>%
    arrange(Species) %>%
    plot_ly(x = ~Sample, y = ~relative_ab, color = ~Species,
```

```
        colors = col_c_18S_mix3, type = "bar") %>%
    layout(yaxis = list(title = "Relative Abundance"), barmode = "stack")
```

**NOTE:** The results from these analyses are presented in **Fig. S33**.

Similarly, in mapping the 18S rRNA database, only three enriched libraries were mapped to the reference, and none of the unenriched libraries were mapped (Table S13). The ones that mapped showed that the percentage of reads mapping was between 0.001 and 0.021%. The species assignment for the three samples shows a drastic difference between enriched and unenriched samples (Fig. S33).

##### 12.2 Set of 100 clinical samples

A real-time qPCR assay was used to quantify *Cryptosporidium* in this set of samples, using primers for the 18S rRNA gene and probes slightly modified from Stroup et al (2006). We used as forward primer 5' -GGGTTGTATTTATTAGATAAAGAACCA- 3', reverse primer 5'-AGGCCAATACCCTACCGTCT-3', and an internal probe 5'-TGACATATCATTCAAGTTTCTGAC-3'. Amplifications were carried out on a final volume of 20 µL, with 5 µL of 4x TaqPathTM 1-step RT-qPCR Master Mix (Thermo Fisher Scientific, MD, USA), 1 µL of 10 µM forward primer, 1 µL of 10 µM reverse primer, 2 µL of 2.5 µM fluorescent probe, 5 µL of 1:5 diluted DNA, and 6 µL of water.

In the results below, it is notable that, particularly in unenriched samples, many reads map to all genomes, introducing a bias towards *C. baileyi* and *C. muris*. We suspect this may be due to i) likely contamination of the C. baileyi and C. muris genome sequences with likely bacterial reads, and ii) a high proportion of bacterial reads after library prep that carry over particularly when libraries are not enriched with CryptoCap_100K, which map to these portions of the genomes.

**12.2.1 Enriched and unenriched clinical samples from NEB libraries with iTru primers** Twenty-six samples required more than one attempt and an increased number of PCR cycles (from 14 to 18) at library preparation to produce a library that passed QC. We obtained an average of 2.6 million paired reads per enriched library for the 100 samples, and 4.1 million paired reads for 73 samples from unenriched libraries. After trimming and quality filtering the clinical samples prepared with the NEB protocol, 83.9% and 61.5% of reads were retained for the enriched and unenriched samples, respectively (Table S14).

**12.2.1.1 Sequence data mapped to *C. parvum*** This section analyzes data from 100 enriched and unenriched clinical samples using the NEB protocol with iTru primers. Summary results from these analyses can be found in **Table S14**, which is imported below.

```
file = "~/crypto_data/12/clinical_samples_iTru_summary_stats.txt" clneb_table = read.table(file,
header = TRUE, sep = "")
```

We tested whether the raw number of sequencing reads was different between enriched and unenriched clinical samples.

```
lsmeans(lm(data = clneb_table, Raw_Paired_Reads ~ Library), pairwise ~ Library)
```

```
## $lsmeans
##  Library     lsmean     SE  df lower.CL upper.CL
##  Enriched   2603198 355000 171  1902580  3303815
##  Unenriched 4078984 415000 171  3258973  4898994
##
## Confidence level used: 0.95
```

```
##
## $contrasts
##  contrast                estimate     SE  df t.ratio p.value
##  Enriched - Unenriched -1475786 546000 171  -2.701  0.0076
```

```
p <- plot_ly()
p <- add_bars(p, x = c("Enriched", "Enriched0", "Unenriched"), y = c(1.3e+07, 2.8e+07,
    1.3e+07), opacity = 1, showlegend = F, marker = list(line = list(color = "rgba(0,0,0,0"),
    color = "rgba(0,0,0,0"), text = c("", "p-value = 0.008**", ""), textposition = "outside",
    legendgroup = "1")
p <- add_lines(p, x = c("Enriched", "Enriched", "Unenriched", "Unenriched"), y = c(26500000,
    26800000, 26800000, 26500000), showlegend = F, line = list(color = "black"),
    legendgroup = "1", hoverinfo = "none")
p <- add_boxplot(p, data = clneb_table, x = ~Library, y = ~Raw_Paired_Reads, color = ~Library,
    type = "box", boxpoints = "all", jitter = 0.3, pointpos = 0, legendgroup = "1")
p <- layout(p, xaxis = list(tickmode = "array", tickvals = c("Enriched", "sf", "Unenriched"),
    ticktext = c("Enriched", "", "Unenriched")), yaxis = list(range = c(-1, 3e+07)))
```

**NOTE:** The results from these analyses are presented in **Fig. S34**.

We also tested if the number of raw reads that mapped to *C. parvum* was different between enriched and unenriched clinical samples.

```
lsmeans(lm(data = clneb_table, Mapped_Paired_Reads_Cparvum ~ Library), pairwise ~
    Library)
```

```
## $lsmeans
##  Library    lsmean     SE  df lower.CL upper.CL
##  Enriched   914219 169000 171   580030  1248409
##  Unenriched   1374 198000 171  -389765   392513
##
## Confidence level used: 0.95
##
## $contrasts
##  contrast                estimate     SE  df t.ratio p.value
##  Enriched - Unenriched   912845 261000 171   3.502  0.0006
```

```
p <- plot_ly()
p <- add_bars(p, x = c("Enriched", "Enriched0", "Unenriched"), y = c(5e+05, 2e+07,
    5e+05), opacity = 1, showlegend = F, marker = list(line = list(color = "rgba(0,0,0,0"),
    color = "rgba(0,0,0,0"), text = c("", "***p-value = 0.0006", ""), textposition = "outside",
    legendgroup = "1")
p <- add_lines(p, x = c("Enriched", "Enriched", "Unenriched", "Unenriched"), y = c(1.9e+07,
    2e+07, 2e+07, 1.9e+07), showlegend = F, line = list(color = "black"), legendgroup = "1",
    hoverinfo = "none")
p <- add_boxplot(p, data = clneb_table, x = ~Library, y = ~Mapped_Paired_Reads_Cparvum,
    color = ~Library, type = "box", boxpoints = "all", jitter = 0.3, pointpos = 0,
    legendgroup = "1")
p <- layout(p, xaxis = list(tickmode = "array", tickvals = c("Enriched", "sf", "Unenriched"),
    ticktext = c("Enriched", "", "Unenriched")), yaxis = list(range = c(-1, 2.1e+07)))
```

**NOTE:** The results from these analyses are presented in **Fig. S35**.

We also tested if the percentage of reads that mapped to *C. parvum* was different between enriched and unenriched clinical samples

```
lsmeans(lm(data = clneb_table, Per_Map_Paired_Reads_Cparvum ~ Library), pairwise ~
    Library)
```

```
## $lsmeans
##  Library     lsmean   SE  df lower.CL upper.CL
##  Enriched   35.6176 1.72 151    32.22    39.02
##  Unenriched  0.0803 2.36 151    -4.59     4.75
##
## Confidence level used: 0.95
##
## $contrasts
##  contrast              estimate   SE  df t.ratio p.value
##  Enriched - Unenriched     35.5 2.92 151  12.154  <.0001
```

```
p <- plot_ly()
p <- add_boxplot(p, data = clneb_table, x = ~Sample, y = ~Per_Map_Paired_Reads_Cparvum,
    color = ~Library, type = "box", boxpoints = "all", jitter = 0.3, pointpos = 0,
    legendgroup = "1")
p <- add_bars(p, data = clneb_table, x = ~Sample, y = ~Per_Map_Paired_Reads_Cparvum,
    color = ~Library)
p <- layout(p, xaxis = list(tickmode = "array", tickvals = c("Enriched", "sf", "Unenriched"),
    ticktext = c("Enriched", "", "Unenriched")), yaxis = list(range = c(-1, 80)))
```

**NOTE:** The results from these analyses are presented in **Fig. S35**.

We tested if the depth of genome coverage obtained by mapping sequence reads to *C. parvum* was different between enriched and unenriched clinical samples.

```
lsmeans(lm(data = clneb_table, Depth_Cparvum ~ Library), pairwise ~ Library)
```

```
## $lsmeans
##  Library     lsmean   SE  df lower.CL upper.CL
##  Enriched   28.1050 5.36 171     17.5     38.7
##  Unenriched  0.0397 6.28 171    -12.4     12.4
##
## Confidence level used: 0.95
##
## $contrasts
##  contrast              estimate   SE  df t.ratio p.value
##  Enriched - Unenriched     28.1 8.26 171   3.399  0.0008
```

```
boxplot_depth <- plot_ly()
boxplot_depth <- add_bars(boxplot_depth, x = c("Enriched", "Enriched0", "Unenriched"),
    y = c(565, 570, 565), opacity = 1, showlegend = F, marker = list(line = list(color = "rgba(0,0,0,0"
        color = "rgba(0,0,0,0"), text = c("", "***p-value = 0.0008", ""), textposition = "outside",
    legendgroup = "1")
boxplot_depth <- add_lines(boxplot_depth, x = c("Enriched", "Enriched", "Unenriched",
    "Unenriched"), y = c(565, 570, 570, 565), showlegend = F, line = list(color = "black"),
    legendgroup = "1", hoverinfo = "none")
```

```r
boxplot_depth <- add_boxplot(boxplot_depth, data = clneb_table, x = ~Library, y = ~Depth_Cparvum,
    color = ~Library, type = "box", boxpoints = "all", jitter = 0.3, pointpos = 0,
    legendgroup = "1")
boxplot_depth <- layout(boxplot_depth, xaxis = list(tickmode = "array", tickvals = c("Enriched",
    "sf", "Unenriched"), ticktext = c("Enriched", "", "Unenriched")), yaxis = list(range = c(-1,
    600), title = "Depth (X)"))
```

We also tested whether the breadth of genome coverage from mapping sequence reads to *C. parvum* was different between enriched and unenriched clinical samples with the NEB protocol.

```r
lsmeans(lm(data = clneb_table, Breadth_Cparvum ~ Library), pairwise ~ Library)
```

```
## $lsmeans
##  Library     lsmean   SE  df lower.CL upper.CL
##  Enriched   31.3031 2.02 171    27.31     35.3
##  Unenriched  0.0221 2.37 171    -4.66      4.7
##
## Confidence level used: 0.95
##
## $contrasts
##  contrast              estimate   SE  df t.ratio p.value
##  Enriched - Unenriched     31.3 3.12 171  10.037  <.0001
```

```r
boxplot_breadth <- plot_ly()
boxplot_breadth <- add_bars(boxplot_breadth, x = c("Enriched", "Enriched0", "Unenriched"),
    y = c(80, 106, 80), opacity = 1, showlegend = F, marker = list(line = list(color = "rgba(0,0,0,0"),
        color = "rgba(0,0,0,0"), text = c("", "***p-value < 0.0001", ""), textposition = "outside",
    legendgroup = "1")
boxplot_breadth <- add_lines(boxplot_breadth, x = c("Enriched", "Enriched", "Unenriched",
    "Unenriched"), y = c(102, 105, 105, 102), showlegend = F, line = list(color = "black"),
    legendgroup = "1", hoverinfo = "none")
boxplot_breadth <- add_boxplot(boxplot_breadth, data = clneb_table, x = ~Library,
    y = ~Breadth_Cparvum, color = ~Library, type = "box", boxpoints = "all", jitter = 0.3,
    pointpos = 0, legendgroup = "1")
boxplot_breadth <- layout(boxplot_breadth, xaxis = list(tickmode = "array", tickvals = c("Enriched",
    "sf", "Unenriched"), ticktext = c("Enriched", "", "Unenriched")), yaxis = list(range = c(-1,
    113), title = "Breadth (%)"))

# creating subplot
fig <- subplot(boxplot_depth, boxplot_breadth, nrows = 1, titleY = TRUE, titleX = TRUE,
    margin = 0.1)
```

**NOTE:** The results from these analyses are presented in **Fig. S36**.

Then, we tested if these patterns are maintained when normalizing by the number of retained reads.

```r
lsmeans(lm(data = clneb_table, Normalized_Depth_Cparvum ~ Library),
    pairwise ~ Library)
```

```
## $lsmeans
##  Library      lsmean     SE  df lower.CL upper.CL
##  Enriched   1.119600 0.0598 169     1.00     1.24
```

```
##  Unenriched 0.000423 0.0709 169    -0.14     0.14
##
## Confidence level used: 0.95
##
## $contrasts
##  contrast              estimate     SE  df t.ratio p.value
##  Enriched - Unenriched     1.12 0.0928 169  12.063  <.0001
```

```
boxplot_depth <- plot_ly()
boxplot_depth <- add_bars(boxplot_depth, x = c("Enriched", "Enriched0",
    "Unenriched"), y = c(4, 5, 4), opacity = 1, showlegend = F,
    marker = list(line = list(color = "rgba(0,0,0,0"), color = "rgba(0,0,0,0"),
    text = c("", "***p-value < 0.0001", ""), textposition = "outside",
    legendgroup = "1")
boxplot_depth <- add_lines(boxplot_depth, x = c("Enriched", "Enriched",
    "Unenriched", "Unenriched"), y = c(4.7, 5, 5, 4.7), showlegend = F,
    line = list(color = "black"), legendgroup = "1", hoverinfo = "none")
boxplot_depth <- add_boxplot(boxplot_depth, data = clneb_table,
    x = ~Library, y = ~Normalized_Depth_Cparvum, color = ~Library,
    type = "box", boxpoints = "all", jitter = 0.3, pointpos = 0,
    legendgroup = "1")
boxplot_depth <- layout(boxplot_depth, xaxis = list(tickmode = "array",
    tickvals = c("Enriched", "sf", "Unenriched"), ticktext = c("Enriched",
        "", "Unenriched")), yaxis = list(range = c(0, 6), title = "Depth (X)"))
lsmeans(lm(data = clneb_table, Normalized_Breadth_Cparvum ~ Library),
    pairwise ~ Library)
```

```
## $lsmeans
##  Library      lsmean    SE  df lower.CL upper.CL
##  Enriched   2.244900 0.126 169    1.995    2.494
##  Unenriched 0.000845 0.150 169   -0.295    0.297
##
## Confidence level used: 0.95
##
## $contrasts
##  contrast              estimate    SE  df t.ratio p.value
##  Enriched - Unenriched     2.24 0.196 169  11.442  <.0001
```

```
boxplot_breadth <- plot_ly()
boxplot_breadth <- add_bars(boxplot_breadth, x = c("Enriched",
    "Enriched0", "Unenriched"), y = c(8.5, 9, 8.5), opacity = 1,
    showlegend = F, marker = list(line = list(color = "rgba(0,0,0,0"),
        color = "rgba(0,0,0,0"), text = c("", "***p-value < 0.0001",
        ""), textposition = "outside", legendgroup = "1")
boxplot_breadth <- add_lines(boxplot_breadth, x = c("Enriched",
    "Enriched", "Unenriched", "Unenriched"), y = c(8.3, 8.6,
    8.6, 8.3), showlegend = F, line = list(color = "black"),
    legendgroup = "1", hoverinfo = "none")
boxplot_breadth <- add_boxplot(boxplot_breadth, data = clneb_table,
    x = ~Library, y = ~Normalized_Breadth_Cparvum, color = ~Library,
    type = "box", boxpoints = "all", jitter = 0.3, pointpos = 0,
    legendgroup = "1")
boxplot_breadth <- layout(boxplot_breadth, xaxis = list(tickmode = "array",
```

```
    tickvals = c("Enriched", "sf", "Unenriched"), ticktext = c("Enriched",
        "", "Unenriched")), yaxis = list(range = c(-1, 10), title = "Breadth (%)"))

# creating subplot
fig <- subplot(boxplot_depth, boxplot_breadth, nrows = 1, titleY = TRUE,
    titleX = TRUE, margin = 0.1)
```

**NOTE:** The results from these analyses are presented in **Fig. S36**.

For fecal DNA obtained from 100 patients when using the NEB-iTru protocol (Table S14; Fig. S34), on average, 35.6% of enriched vs. 0.08% of unenriched retained reads mapped to the C. parvum genome sequence (p-value < 0.0001; Table S14; Fig. S35). Libraries were enriched with C. parvum by an average factor of 2,095X (7.20–5,097X; Table S14). The depth and breadth of genome coverage were significantly higher for enriched samples, with average depth of 18X and breadth of 31% vs. 0.04X and 0.02% in unenriched samples (Fig. S36; normalized parameters per 100,000 reads can also be found in Fig. S36).

The depth and breadth of genome coverage were significantly higher for enriched samples with an average depth of 28.1X and breadth of 31.3% for enriched vs. an average depth of 0.04X and breadth of 0.02% for the unenriched libraries (depth p-value = 0.0008, breadth p-value < 0.0001; Fig. S35; Table S14). This remains true with normalized parameters per 100,000 reads, obtaining 1.12X and breadth of 2.25% for enriched vs. an average depth of 0.0004X and breadth of 0.0008% for the unenriched libraries (p-values < 0.0001; Fig. S36; Table S14).

**12.2.1.2 Sequence data mapped to 10-genomes reference database** Similarly to simulated sequencing reads, we mapped each sample sequenced against the fasta containing ten genomes, we used BBmap, and as a reference, we used the reference file created above. We used a maxim indel length of 20, a minid of 0.8 (read note below), we also required correct strands for paired reads (rcs), and set a the mean pair size to 400, and did not rescue paired reads if avg. insert size greater than 600 (rescuedist; let's remember that the simulated size is 500), and we sat an average pair distance *(apd)of 100 bases:

```
mkdir clinical_samples/mapped
bbmap.sh in=1.fq in2=2.fq \
ref 10_crypto_genomes.fasta  \
maxindel=20 minid=0.80 rcs=t killbadpairs=t pairlen=400 \
rescuedist=600 apd=100 outu=clinical_samples/mapped/un_sample.sam \
outm=clinical_samples/mapped/sample.sam printunmappedcount=t \
covstats=clinical_samples/mapped/sample.cov.txt threads=12 -Xmx180g 2>&1 \
| tee clinical_samples/mapped/sample.bbmap.log
```

A 5.6% increase in paired reads mapping to 10-CrypGS vs. the C. parvum genome was observed for enriched libraries and 0.05% for unenriched (Table S14). Of the 73 unenriched clinical samples with sequence data, only 53 had hits to 10-CrypGS (Table S14), with an average of 2,705 mapped reads per sample.

12.2.1.2.1 Assessing associations of qPCR Ct values and percentage of reads mapped to reference

We also tested the association between Ct values and the raw number of reads obtained via a linear regression model, using only unenriched libraries.

```
# scatter plot of number of reads obtained and number of reads mapped
per_mapped_unenriched <- clneb_table[c(101:200), c(3, 14)]

y <- per_mapped_unenriched$Per_Map_Paired_Reads_10_genomes
X <- per_mapped_unenriched$qPCR_Ct
```

```r
lm_model <- linear_reg() %>%
    set_engine("lm") %>%
    set_mode("regression") %>%
    fit(Per_Map_Paired_Reads_10_genomes ~ qPCR_Ct, data = per_mapped_unenriched)
lm_model
```

```
## parsnip model object
##
##
## Call:
## stats::lm(formula = Per_Map_Paired_Reads_10_genomes ~ qPCR_Ct,
##     data = data)
##
## Coefficients:
## (Intercept)      qPCR_Ct
##    -0.20602      0.01233
```

```r
summary(lm(Per_Map_Paired_Reads_10_genomes ~ qPCR_Ct, data = per_mapped_unenriched))
```

```
##
## Call:
## lm(formula = Per_Map_Paired_Reads_10_genomes ~ qPCR_Ct, data = per_mapped_unenriched)
##
## Residuals:
##      Min       1Q   Median       3Q      Max
## -0.23630 -0.10630 -0.07601 -0.03394  2.19140
##
## Coefficients:
##             Estimate Std. Error t value Pr(>|t|)
## (Intercept) -0.20602    0.34381  -0.599    0.551
## qPCR_Ct      0.01233    0.01333   0.926    0.358
##
## Residual standard error: 0.3231 on 71 degrees of freedom
##   (27 observations deleted due to missingness)
## Multiple R-squared:  0.01192,    Adjusted R-squared:  -0.001994
## F-statistic: 0.8567 on 1 and 71 DF,  p-value: 0.3578
```

```r
x_range <- seq(min(X), max(X), length.out = 100)
x_range <- matrix(x_range, nrow = 100, ncol = 1)
xdf <- data.frame(x_range)
colnames(xdf) <- c("qPCR_Ct")

ydf <- lm_model %>%
    predict(xdf)

colnames(ydf) <- c("Per_Map_Paired_Reads_10_genomes")
xy <- data.frame(xdf, ydf)

fig <- plot_ly(data = per_mapped_unenriched, x = ~qPCR_Ct, y = ~Per_Map_Paired_Reads_10_genomes,
    type = "scatter", alpha = 0.65, mode = "markers", name = "% Reads Mapped to 10-genomes Reference")
fig <- fig %>%
    add_trace(data = xy, x = ~qPCR_Ct, y = ~Per_Map_Paired_Reads_10_genomes, name = "prediction",
        mode = "lines", alpha = 1)
```

**NOTE:** The results from these analyses are presented in **Fig. S37**.

We tested the association between Ct values and the raw number of reads obtained via a linear regression model, using only enriched libraries.

```
# scatter plot of number of reads obtained and number of reads mapped
per_mapped_enriched <- clneb_table[c(1:100), c(3, 14)]

y <- per_mapped_enriched$Per_Map_Paired_Reads_10_genomes
X <- per_mapped_enriched$qPCR_Ct

lm_model <- linear_reg() %>%
    set_engine("lm") %>%
    set_mode("regression") %>%
    fit(Per_Map_Paired_Reads_10_genomes ~ qPCR_Ct, data = per_mapped_enriched)
lm_model
```

```
## parsnip model object
##
##
## Call:
## stats::lm(formula = Per_Map_Paired_Reads_10_genomes ~ qPCR_Ct,
##     data = data)
##
## Coefficients:
## (Intercept)      qPCR_Ct
##     112.580       -2.808
```

```
summary(lm(Per_Map_Paired_Reads_10_genomes ~ qPCR_Ct, data = per_mapped_enriched))
```

```
##
## Call:
## lm(formula = Per_Map_Paired_Reads_10_genomes ~ qPCR_Ct, data = per_mapped_enriched)
##
## Residuals:
##     Min      1Q  Median      3Q     Max
## -39.440 -12.167   0.332  10.889  35.819
##
## Coefficients:
##             Estimate Std. Error t value Pr(>|t|)
## (Intercept) 112.5804    16.9328   6.649 1.69e-09 ***
## qPCR_Ct      -2.8084     0.6627  -4.238 5.11e-05 ***
## ---
## Signif. codes:  0 '***' 0.001 '**' 0.01 '*' 0.05 '.' 0.1 ' ' 1
##
## Residual standard error: 17.98 on 98 degrees of freedom
## Multiple R-squared:  0.1549, Adjusted R-squared:  0.1463
## F-statistic: 17.96 on 1 and 98 DF,  p-value: 5.108e-05
```

```
x_range <- seq(min(X), max(X), length.out = 100)
x_range <- matrix(x_range, nrow = 100, ncol = 1)
xdf <- data.frame(x_range)
colnames(xdf) <- c("qPCR_Ct")
```

```
ydf <- lm_model %>%
    predict(xdf)

colnames(ydf) <- c("Per_Map_Paired_Reads_10_genomes")
xy <- data.frame(xdf, ydf)

fig <- plot_ly(data = per_mapped_enriched, x = ~qPCR_Ct, y = ~Per_Map_Paired_Reads_10_genomes,
    type = "scatter", alpha = 0.65, mode = "markers", name = "% Reads Mapped to 10-genomes Reference")
fig <- fig %>%
    add_trace(data = xy, x = ~qPCR_Ct, y = ~Per_Map_Paired_Reads_10_genomes, name = "prediction",
        mode = "lines", alpha = 1)
```

**NOTE:** The results from these analyses are presented in **Fig. S37**.

For unenriched libraries, qPCR CT values were not a good predictor of the percentage of reads mapping (R2 = 0.012, p-value = 0.36; Fig. S37). In enriched libraries the qPCR CT values were good predictor of the 15.5% of the variation in the percentage of mapping reads with a statistically significant effect showing that for every increase of one qPCR CT unit, the percentage decreased by 2.81% (R2 = 0.155, p-value = 5.11e-5; ±18%; Fig. S37; Table S14).

12.2.1.2.2 Counting and plotting the proportion of hits to different species' genomes

For each sample, we used the FLAG count from the third column of our SAM files to extract those sequences for which both reads mapped (FLAG: 83 and 99):

```
cat clinical_samples/mapped/sample.sam | grep -v "^@" | \
awk 'BEGIN{FS="\t";OFS="\t"}{if($2=="83" || $2=="99") print $0}' \
> clinical_samples/mapped/both/sample.txt
```

Then, for each sample, we will count each species ID hit in our mapped file. The count text file has two columns, the first is the species ID, and the second is the counts:

```
cat clinical_samples/mapped/both/sample.txt | \
awk 'BEGIN{FS="\t";OFS="\t"}{OTU[$3]+=1}END{for (i in OTU) print i,OTU[i]}' \
> clinical_samples/mapped/both/species_ct.txt
```

First, we analyzed unenriched libraries. Before importing each file from each sample, we delimited the first column by "-", separating the added species name.

```
awk -F '\t' '{gsub(/-.*/, "", $1); print}' sample_species_ct.txt > sample_species_ct_edited.txt
```

Below, we imported the edited files for each species and then created a fourth column, "Sample," with the name of the species for which we simulated the reads.

```
# Create an empty list to store the data frames
data_list_unenriched <- list()
# Loop through the list of file names in 'elist'
unlist <- gsub(" ", "", paste(clneb_table$Sample, "_U"))
for (i in unlist) {
    # Construct the file path
    file_path <- paste("~/crypto_data/12/iTru-NEB/species_edited/",
        i, "_species_ct.txt", sep = "")
```

```
    # Check if the file exists and has data (not empty)
    if (file.exists(file_path) && file.info(file_path)$size > 0) {
        # Read the file into a data frame
        data <- read.table(file_path, header = FALSE, sep = "\t", stringsAsFactors = FALSE)

        # Remove columns 3 to 5
        data <- data[, -c(3:5)]

        # Rename columns
        colnames(data) <- c("Species", "Count")

        # Calculate Relative abundance of counts for each genome
        data <- data %>%
            mutate(relative_ab = Count/sum(Count))

        # Add the processed data frame to the list with the file name as the
        # list element name
        data_list_unenriched[[i]] <- data
    }
}

# Combine the data frames in data_list into a single data frame
combined_data_unenriched <- bind_rows(data_list_unenriched, .id = "Sample")

# Sort the data by Sample and Species
combined_data_unenriched <- combined_data_unenriched %>%
    arrange(Sample, Species)

# Create the plot
p <- plot_ly(data = combined_data_unenriched, x = ~Sample, y = ~relative_ab, color = ~Species,
    colors = col_c_ten, type = "bar") %>%
    layout(yaxis = list(title = "Relative Abundance"), barmode = "stack")
```

**NOTE:** The results from these analyses are presented in **Fig. S38**.

Species assignments for unenriched libraries resemble the complex background from LOD experiments (compare to Fig. 2B). Samples with hits to species known to infect humans with high likelihood include nine samples with high proportion assignments (>50%) to *C. parvum*, seven samples with high proportion (>50%) of *C. hominis* (UKP323, UKP328, UKP332, UKP345, UKP350, UKP376, UKP392), and one containing >25% of *C. cuniculus*. There were higher abundances of *C. muris* hits (>50% in 13 samples).

And then, we followed the same procedure for the enriched libraries that got hits against the 10-genomes reference.

```
# Create an empty list to store the data frames
data_list_enriched <- list()
# Loop through the list of file names in 'elist'
elist <- gsub(" ", "", paste(clneb_table$Sample, "_E"))
for (i in elist) {
    # Read the file into a data frame
    file_path <- paste("~/crypto_data/12/iTru-NEB/species_edited/",
        i, "_species_ct.txt", sep = "")
    data <- read.table(file_path, header = FALSE, sep = "\t", stringsAsFactors = FALSE)
    # Remove columns 3 to 5
```

```
    data <- data[, -c(3:5)]
    # Rename columns
    colnames(data) <- c("Species", "Count")
    # Calculate Relative abundance of counts for each genome
    data <- data %>%
        mutate(relative_ab = Count/sum(Count))

    # Add the processed data frame to the list with the file name as the list
    # element name
    data_list_enriched[[i]] <- data
}
# Combine the data frames in data_list into a single data frame
combined_data_enriched <- bind_rows(data_list_enriched, .id = "Sample")

# Sort the data by Sample and Species
combined_data_enriched <- combined_data_enriched %>%
    arrange(Sample, Species)

# Create the plot
p <- plot_ly(data = combined_data_enriched, x = ~Sample, y = ~relative_ab, color = ~Species,
    colors = col_c_ten, type = "bar") %>%
    layout(yaxis = list(title = "Relative Abundance"), barmode = "stack")
```

**NOTE:** The results from these analyses are presented in **Fig. S38**.

Following enrichment, an average of 1.03 million reads hit the database. The relative abundance of species for the single enriched libraries shows that 63 out of the 100 samples' reads are assigned in the majority (>50%) to *C. parvum*. The other 37 samples have other species assignments in high proportions. For example, five samples (UKH146, UKH147, UKH148, UKH150, and UKH151) contain a high proportion (>50%) of *C. hominis*, and all of these, except UKH151, also show moderate proportions of *C. cuniculus* (between 12% and 35%). Notice that these samples with *C. hominis* do not correspond to those samples with high levels of *C. hominis* in the unenriched. Some samples also show moderate proportions of *C. baileyi*, *C. meleagridis*, *C. tyzzeri*, and *C. viatorum*, but consistently below 25% each. The other species, *C. andersoni*, *C. muris*, and *C. ubiquitum* were assigned to most samples but always in percentages lower than 10%.

##### 12.2.2 Enriched and unenriched clinical samples from iNextEra libraries with iNext primers

This section analyzes data from 100 unenriched, single-enriched, and double-enriched clinical samples using the iNextEra protocol. The results from these analyses are summarized in **Table S15**, also imported below.

We obtained an average of 3.7 million paired reads per unenriched library, 5.3 million paired reads for single-enriched libraries, and 3.6 million for double-enriched libraries.

```
file = "~/crypto_data/12/clinical_samples_iNext_summary_stats.txt" clinext_table = read.table(file,
header = TRUE, sep = "")
```

We tested if the raw number of sequencing reads was different between unenriched, single-enriched, and double-enriched clinical samples.

```
lsmeans(lm(data = clinext_table, RawReadPairs ~ Library), pairwise ~
    Library)
```

```
## $lsmeans
```

```
##  Library          lsmean     SE  df lower.CL upper.CL
##  Double-enriched 3600743 646000 270  2328101  4873385
##  Single-enriched 5298444 646000 270  4025802  6571086
##  Unenriched      3720999 646000 270  2448358  4993641
##
## Confidence level used: 0.95
##
## $contrasts
##  contrast                              estimate     SE  df t.ratio p.value
##  (Double-enriched) - (Single-enriched) -1697701 914000 270  -1.857  0.1533
##  (Double-enriched) - Unenriched         -120256 914000 270  -0.132  0.9905
##  (Single-enriched) - Unenriched         1577445 914000 270   1.726  0.1975
##
## P value adjustment: tukey method for comparing a family of 3 estimates
```

```r
p <- plot_ly()
p <- add_bars(p, x = c("Double-enriched", "Double-enriched0",
    "Single-enriched", "Single-enriched0", "Unenriched"), y = c(6.9e+07,
    6.9e+07, 7.3e+07, 6.9e+07, 6.9e+07), opacity = 1, showlegend = F,
    marker = list(line = list(color = "rgba(0,0,0,0"), color = "rgba(0,0,0,0"),
    text = c("", "p-value = 0.153", "p-value = 0.991", "p-value = 0.198",
        ""), textposition = "outside", legendgroup = "1")

p <- add_lines(p, x = c("Double-enriched", "Double-enriched",
    "Single-enriched", "Single-enriched", "Single-enriched",
    "Single-enriched", "Unenriched", "Unenriched"), y = c(6.7e+07,
    68500000, 68500000, 6.7e+07, 6.7e+07, 68500000, 68500000,
    6.7e+07), showlegend = F, line = list(color = "black"), legendgroup = "1",
    hoverinfo = "none")

p <- add_lines(p, x = c("Double-enriched", "Double-enriched",
    "Unenriched", "Unenriched"), y = c(7.2e+07, 72500000, 72500000,
    7.2e+07), showlegend = F, line = list(color = "black"), legendgroup = "1",
    hoverinfo = "none")
p <- add_boxplot(p, data = clinext_table, x = ~Library, y = ~RawReadPairs,
    color = ~Library, type = "box", boxpoints = "all", jitter = 0.3,
    pointpos = 0, legendgroup = "1")
p <- layout(p, xaxis = list(tickmode = "array", tickvals = c("Double-enriched",
    "sf", "Single-enriched", "sf", "Unenriched"), ticktext = c("Double-enriched",
    "", "Single-enriched", "", "Unenriched")), yaxis = list(range = c(-1,
    8e+07)))
```

**NOTE:** The results from these analyses are presented in **Fig. S39**.

The depth and breadth of genome coverage were significantly higher for double-enriched samples than for single-enriched libraries and unenriched libraries when normalized per 100,000 reads, obtaining 0.001X and 0.06% breadth for unenriched, 0.56X, and 1.79% breadth for single-enriched, and 2.70X and 5.99% breadth for double-enriched (Fig. S39, Table S15).

**12.2.2.1 Sequence data mapped to *C. parvum* genome** We also tested if the number of raw reads that mapped to *C. parvum* was different between enriched and unenriched clinical samples.

```r
lsmeans(lm(data = clinext_table, MappedPairedReadsCparvum ~ Library),
    pairwise ~ Library)
```

```
## $lsmeans
##  Library          lsmean     SE  df lower.CL upper.CL
##  Double-enriched 2406965 352000 270  1713601  3100330
##  Single-enriched  941467 352000 270   248102  1634831
##  Unenriched         1424 352000 270  -691941   694788
##
## Confidence level used: 0.95
##
## $contrasts
##  contrast                              estimate     SE  df t.ratio p.value
##  (Double-enriched) - (Single-enriched)  1465499 498000 270   2.942  0.0099
##  (Double-enriched) - Unenriched         2405542 498000 270   4.830  <.0001
##  (Single-enriched) - Unenriched          940043 498000 270   1.887  0.1443
##
## P value adjustment: tukey method for comparing a family of 3 estimates
```

```r
p <- plot_ly()
p <- add_bars(p, x = c("Double-enriched", "Double-enriched0",
    "Single-enriched", "Single-enriched0", "Unenriched"), y = c(3.4e+07,
    3e+07, 37500000, 3e+07, 3.4e+07), opacity = 1, showlegend = F,
    marker = list(line = list(color = "rgba(0,0,0,0"), color = "rgba(0,0,0,0"),
    text = c("", "*p-value = 0.010", "***p-value < 0.0001", "p-value = 0.1443",
        ""), textposition = "outside", legendgroup = "1")

p <- add_lines(p, x = c("Double-enriched", "Double-enriched",
    "Single-enriched", "Single-enriched", "Single-enriched",
    "Single-enriched", "Unenriched", "Unenriched"), y = c(3.4e+07,
    3.5e+07, 3.5e+07, 3.4e+07, 3.4e+07, 3.5e+07, 3.5e+07, 3.4e+07),
    showlegend = F, line = list(color = "black"), legendgroup = "1",
    hoverinfo = "none")

p <- add_lines(p, x = c("Double-enriched", "Double-enriched",
    "Unenriched", "Unenriched"), y = c(3.6e+07, 3.7e+07, 3.7e+07,
    3.6e+07), showlegend = F, line = list(color = "black"), legendgroup = "1",
    hoverinfo = "none")
p <- add_boxplot(p, data = clinext_table, x = ~Library, y = ~MappedPairedReadsCparvum,
    color = ~Library, type = "box", boxpoints = "all", jitter = 0.3,
    pointpos = 0, legendgroup = "1")
p <- layout(p, xaxis = list(tickmode = "array", tickvals = c("Double-enriched",
    "sf", "Single-enriched", "sf", "Unenriched"), ticktext = c("Double-enriched",
    "", "Single-enriched", "", "Unenriched")), yaxis = list(range = c(-1,
    4e+07)))
```

**NOTE:** The results from these analyses are presented in **Fig. S40**.

We also tested if the percentage of reads that mapped to *C. parvum* was different between enriched and unenriched clinical samples.

```r
lsmeans(lm(data = clinext_table, PerMapPairedReadsCparvum ~ Library),
    pairwise ~ Library)
```

```
## $lsmeans
##  Library          lsmean   SE  df lower.CL upper.CL
##  Double-enriched 64.9843 1.35 270    62.34    67.63
##  Single-enriched 15.8511 1.35 270    13.20    18.50
##  Unenriched       0.0331 1.35 270    -2.62     2.68
##
## Confidence level used: 0.95
##
## $contrasts
##  contrast                              estimate  SE  df t.ratio p.value
##  (Double-enriched) - (Single-enriched)     49.1 1.9 270  25.828  <.0001
##  (Double-enriched) - Unenriched            65.0 1.9 270  34.143  <.0001
##  (Single-enriched) - Unenriched            15.8 1.9 270   8.315  <.0001
##
## P value adjustment: tukey method for comparing a family of 3 estimates
```

```r
p <- plot_ly()
p <- add_bars(p, x = c("Double-enriched", "Double-enriched0",
    "Single-enriched", "Single-enriched0", "Unenriched"), y = c(109,
    109, 116, 109, 109), opacity = 1, showlegend = F, marker = list(line = list(color = "rgba(0,0,0,0")
    color = "rgba(0,0,0,0"), text = c("", "***p-value < 0.0001",
    "***p-value < 0.0001", "***p-value < 0.0001", ""), textposition = "outside",
    legendgroup = "1")

p <- add_lines(p, x = c("Double-enriched", "Double-enriched",
    "Single-enriched", "Single-enriched", "Single-enriched",
    "Single-enriched", "Unenriched", "Unenriched"), y = c(105,
    107, 107, 105, 105, 107, 107, 105), showlegend = F, line = list(color = "black"),
    legendgroup = "1", hoverinfo = "none")

p <- add_lines(p, x = c("Double-enriched", "Double-enriched",
    "Unenriched", "Unenriched"), y = c(112, 115, 115, 112), showlegend = F,
    line = list(color = "black"), legendgroup = "1", hoverinfo = "none")
p <- add_boxplot(p, data = clinext_table, x = ~Library, y = ~PerMapPairedReadsCparvum,
    color = ~Library, type = "box", boxpoints = "all", jitter = 0.3,
    pointpos = 0, legendgroup = "1")
p <- layout(p, xaxis = list(tickmode = "array", tickvals = c("Double-enriched",
    "sf", "Single-enriched", "sf", "Unenriched"), ticktext = c("Double-enriched",
    "", "Single-enriched", "", "Unenriched")), yaxis = list(range = c(-1,
    120)))
```

**NOTE:** The results from these analyses are presented in **Fig. 3A**.

We tested if the depth of genome coverage obtained by mapping sequence reads to *C. parvum* was different between enriched and unenriched clinical samples.

```r
lsmeans(lm(data = clinext_table, AverageDepthCparvum ~ Library),
    pairwise ~ Library)
```

```
## $lsmeans
```

```
##  Library          lsmean   SE  df lower.CL upper.CL
##  Double-enriched 95.4994 14.2 270     67.5    123.5
##  Single-enriched 32.1179 14.2 270      4.1     60.1
##  Unenriched       0.0503 14.2 270    -28.0     28.1
##
## Confidence level used: 0.95
##
## $contrasts
##  contrast                              estimate   SE  df t.ratio p.value
##  (Double-enriched) - (Single-enriched)     63.4 20.1 270   3.149  0.0052
##  (Double-enriched) - Unenriched            95.4 20.1 270   4.743  <.0001
##  (Single-enriched) - Unenriched            32.1 20.1 270   1.593  0.2502
##
## P value adjustment: tukey method for comparing a family of 3 estimates
```

```
boxplot_depth <- plot_ly()
boxplot_depth <- add_bars(boxplot_depth, x = c("Double-enriched",
    "Double-enriched0", "Single-enriched", "Single-enriched0",
    "Unenriched"), y = c(1709, 1709, 1920, 1709, 1709), opacity = 1,
    showlegend = F, marker = list(line = list(color = "rgba(0,0,0,0"),
        color = "rgba(0,0,0,0"), text = c("", "**p-value = 0.005",
        "***p-value < 0.0001", "p-value = 0.250", ""), textposition = "outside",
    legendgroup = "1")

boxplot_depth <- add_lines(boxplot_depth, x = c("Double-enriched",
    "Double-enriched", "Single-enriched", "Single-enriched",
    "Single-enriched", "Single-enriched", "Unenriched", "Unenriched"),
    y = c(1605, 1677, 1677, 1605, 1605, 1677, 1677, 1605), showlegend = F,
    line = list(color = "black"), legendgroup = "1", hoverinfo = "none")

boxplot_depth <- add_lines(boxplot_depth, x = c("Double-enriched",
    "Double-enriched", "Unenriched", "Unenriched"), y = c(1812,
    1915, 1915, 1812), showlegend = F, line = list(color = "black"),
    legendgroup = "1", hoverinfo = "none")
boxplot_depth <- add_boxplot(boxplot_depth, data = clinext_table,
    x = ~Library, y = ~AverageDepthCparvum, color = ~Library,
    type = "box", boxpoints = "all", jitter = 0.3, pointpos = 0,
    legendgroup = "1")
boxplot_depth <- layout(boxplot_depth, xaxis = list(tickmode = "array",
    tickvals = c("Double-enriched", "sf", "Single-enriched",
        "sf", "Unenriched"), ticktext = c("Double-enriched",
        "", "Single-enriched", "", "Unenriched")), yaxis = list(range = c(-1,
    2000), title = "Depth (X)"))
```

We also tested if the breadth of genome coverage from the mapping of sequence reads to *C. parvum* was different between enriched and unenriched clinical samples.

```
lsmeans(lm(data = clinext_table, AverageBreadthCparvum ~ Library),
    pairwise ~ Library)
```

```
## $lsmeans
##  Library         lsmean   SE  df lower.CL upper.CL
##  Double-enriched  48.09 2.62 270    42.93    53.24
```

```
##   Single-enriched  54.38 2.62 270    49.22    59.54
##   Unenriched        2.56 2.62 270    -2.59     7.72
##
## Confidence level used: 0.95
##
## $contrasts
##   contrast                              estimate   SE  df t.ratio p.value
##   (Double-enriched) - (Single-enriched)    -6.29 3.7 270  -1.699  0.2073
##   (Double-enriched) - Unenriched           45.52 3.7 270  12.292  <.0001
##   (Single-enriched) - Unenriched           51.82 3.7 270  13.991  <.0001
##
## P value adjustment: tukey method for comparing a family of 3 estimates
```

```
boxplot_breadth <- plot_ly()
boxplot_breadth <- add_bars(boxplot_breadth, x = c("Double-enriched",
    "Double-enriched0", "Single-enriched", "Single-enriched0",
    "Unenriched"), y = c(109, 109, 116, 109, 109), opacity = 1,
    showlegend = F, marker = list(line = list(color = "rgba(0,0,0,0"),
        color = "rgba(0,0,0,0"), text = c("", "p-value = 0.2073",
        "***p-value < 0.0001", "***p-value < 0.0001", ""), textposition = "outside",
    legendgroup = "1")

boxplot_breadth <- add_lines(boxplot_breadth, x = c("Double-enriched",
    "Double-enriched", "Single-enriched", "Single-enriched",
    "Single-enriched", "Single-enriched", "Unenriched", "Unenriched"),
    y = c(105, 107, 107, 105, 105, 107, 107, 105), showlegend = F,
    line = list(color = "black"), legendgroup = "1", hoverinfo = "none")
boxplot_breadth <- add_lines(boxplot_breadth, x = c("Double-enriched",
    "Double-enriched", "Unenriched", "Unenriched"), y = c(112,
    115, 115, 112), showlegend = F, line = list(color = "black"),
    legendgroup = "1", hoverinfo = "none")
boxplot_breadth <- add_boxplot(boxplot_breadth, data = clinext_table,
    x = ~Library, y = ~AverageBreadthCparvum, color = ~Library,
    type = "box", boxpoints = "all", jitter = 0.3, pointpos = 0,
    legendgroup = "1")
boxplot_breadth <- layout(boxplot_breadth, xaxis = list(tickmode = "array",
    tickvals = c("Double-enriched", "sf", "Single-enriched",
        "sf", "Unenriched"), ticktext = c("Double-enriched",
        "", "Single-enriched", "", "Unenriched")), yaxis = list(range = c(-1,
    123), title = "Breadth (%)"))
# creating subplot
fig <- subplot(boxplot_depth, boxplot_breadth, nrows = 1, titleY = TRUE,
    titleX = TRUE, margin = 0.1)
```

**NOTE:** The results from these analyses are presented in **Fig. 3B**.

We tested how the depth and breadth of genome coverage varied for the library types once we normalized for the number of sequenced used in the mapping

```
lsmeans(lm(data = clinext_table, Normalized_Depth_Cparvum ~ Library),
    pairwise ~ Library)
```

```
## $lsmeans
```

```
##   Library          lsmean   SE  df lower.CL upper.CL
##   Double-enriched 2.70221 0.13 270    2.446    2.959
##   Single-enriched 0.56343 0.13 270    0.307    0.820
##   Unenriched      0.00115 0.13 270   -0.255    0.258
##
## Confidence level used: 0.95
##
## $contrasts
##   contrast                              estimate    SE  df t.ratio p.value
##   (Double-enriched) - (Single-enriched)    2.139 0.184 270  11.602  <.0001
##   (Double-enriched) - Unenriched           2.701 0.184 270  14.652  <.0001
##   (Single-enriched) - Unenriched           0.562 0.184 270   3.050  0.0071
##
## P value adjustment: tukey method for comparing a family of 3 estimates
```

```
boxplot_depth <- plot_ly()
boxplot_depth <- add_bars(boxplot_depth, x = c("Double-enriched",
    "Double-enriched0", "Single-enriched", "Single-enriched0",
    "Unenriched"), y = c(24, 24, 27, 24, 24), opacity = 1, showlegend = F,
    marker = list(line = list(color = "rgba(0,0,0,0"), color = "rgba(0,0,0,0"),
    text = c("", "***p-value < 0.0001", "***p-value < 0.0001",
        "**p-value = 0.007", ""), textposition = "outside", legendgroup = "1")

boxplot_depth <- add_lines(boxplot_depth, x = c("Double-enriched",
    "Double-enriched", "Single-enriched", "Single-enriched",
    "Single-enriched", "Single-enriched", "Unenriched", "Unenriched"),
    y = c(23, 24, 24, 23, 23, 24, 24, 23), showlegend = F, line = list(color = "black"),
    legendgroup = "1", hoverinfo = "none")

boxplot_depth <- add_lines(boxplot_depth, x = c("Double-enriched",
    "Double-enriched", "Unenriched", "Unenriched"), y = c(26,
    27, 27, 26), showlegend = F, line = list(color = "black"),
    legendgroup = "1", hoverinfo = "none")
boxplot_depth <- add_boxplot(boxplot_depth, data = clinext_table,
    x = ~Library, y = ~Normalized_Depth_Cparvum, color = ~Library,
    type = "box", boxpoints = "all", jitter = 0.3, pointpos = 0,
    legendgroup = "1")
boxplot_depth <- layout(boxplot_depth, xaxis = list(tickmode = "array",
    tickvals = c("Double-enriched", "sf", "Single-enriched",
        "sf", "Unenriched"), ticktext = c("Double-enriched",
        "", "Single-enriched", "", "Unenriched")), yaxis = list(range = c(-1,
    30), title = "Depth (X)"))
```

We also tested if the breadth of genome coverage from the mapping of sequence reads to *C. parvum* was different between enriched and unenriched clinical samples.

```
lsmeans(lm(data = clinext_table, Normalized_Breadth_Cparvum ~
    Library), pairwise ~ Library)
```

```
## $lsmeans
##   Library         lsmean   SE  df lower.CL upper.CL
##   Double-enriched 5.9926 0.27 270    5.462    6.524
##   Single-enriched 1.7902 0.27 270    1.259    2.321
```

```
##  Unenriched      0.0602 0.27 270   -0.471    0.591
##
## Confidence level used: 0.95
##
## $contrasts
##  contrast                               estimate    SE  df t.ratio p.value
##  (Double-enriched) - (Single-enriched)     4.20 0.381 270  11.019  <.0001
##  (Double-enriched) - Unenriched            5.93 0.381 270  15.555  <.0001
##  (Single-enriched) - Unenriched            1.73 0.381 270   4.536  <.0001
##
## P value adjustment: tukey method for comparing a family of 3 estimates
```

```
boxplot_breadth <- plot_ly()
boxplot_breadth <- add_bars(boxplot_breadth, x = c("Double-enriched",
    "Double-enriched0", "Single-enriched", "Single-enriched0",
    "Unenriched"), y = c(23, 23, 26, 23, 23), opacity = 1, showlegend = F,
    marker = list(line = list(color = "rgba(0,0,0,0"), color = "rgba(0,0,0,0"),
    text = c("", "p-value = 0.2073", "***p-value < 0.0001", "***p-value < 0.0001",
        ""), textposition = "outside", legendgroup = "1")

boxplot_breadth <- add_lines(boxplot_breadth, x = c("Double-enriched",
    "Double-enriched", "Single-enriched", "Single-enriched",
    "Single-enriched", "Single-enriched", "Unenriched", "Unenriched"),
    y = c(22, 23, 23, 22, 22, 23, 23, 22), showlegend = F, line = list(color = "black"),
    legendgroup = "1", hoverinfo = "none")
boxplot_breadth <- add_lines(boxplot_breadth, x = c("Double-enriched",
    "Double-enriched", "Unenriched", "Unenriched"), y = c(25,
    26, 26, 25), showlegend = F, line = list(color = "black"),
    legendgroup = "1", hoverinfo = "none")
boxplot_breadth <- add_boxplot(boxplot_breadth, data = clinext_table,
    x = ~Library, y = ~Normalized_Breadth_Cparvum, color = ~Library,
    type = "box", boxpoints = "all", jitter = 0.3, pointpos = 0,
    legendgroup = "1")
boxplot_breadth <- layout(boxplot_breadth, xaxis = list(tickmode = "array",
    tickvals = c("Double-enriched", "sf", "Single-enriched",
        "sf", "Unenriched"), ticktext = c("Double-enriched",
        "", "Single-enriched", "", "Unenriched")), yaxis = list(range = c(-1,
    28), title = "Breadth (%)"))
# creating subplot
fig <- subplot(boxplot_depth, boxplot_breadth, nrows = 1, titleY = TRUE,
    titleX = TRUE, margin = 0.1)
```

**NOTE:** The results from these analyses are presented in **Fig. S41**.

**12.2.2.2 Sequence data mapped to 10-genomes reference database using BBMap** Similarly to NEB libraries, we mapped each sample sequenced against the fasta containing ten genomes, we used BBmap, and as a reference, we used the reference file created above. We used a maxim indel length of 20, a minid of 0.8 (read note below), we also required correct strands for paired reads (rcs), and set a the mean pair size to 400, and did not rescue paired reads if avg. insert size greater than 600 (rescuedist; let's remember that the simulated size is 500), and we sat an average pair distance *(apd)of 100 bases.

**NOTE:** The results from these analyses are presented in **Fig. 3B**.

12.2.2.2.1 Visualizing read mapping across the reference

From the mapped reads, bigwig files were generated and visualized for each sample and library type. For this, we used the Integrative Genomics Viewer IGV-Web app version 2.1.2 and igv.js version 3.1.2.

**NOTE:** The results from these analyses are presented in **Fig. S42**.

12.2.2.2.2 Assessing associations of qPCR Ct values and percentage of reads mapped to reference

Then, we tested the association between Ct values and the raw number of reads obtained via a linear regression model, using only unenriched libraries.

```
# scatter plot of number of reads obtained and number of
# reads mapped
per_mapped_unenriched <- clinext_table[c(1:100), c(3, 15)]

y <- per_mapped_unenriched$PerMapPairedReads10RefGenomes
X <- per_mapped_unenriched$Ct_qPCR

lm_model <- linear_reg() %>%
    set_engine("lm") %>%
    set_mode("regression") %>%
    fit(PerMapPairedReads10RefGenomes ~ Ct_qPCR, data = per_mapped_unenriched)
lm_model
```

```
## parsnip model object
##
##
## Call:
## stats::lm(formula = PerMapPairedReads10RefGenomes ~ Ct_qPCR,
##     data = data)
##
## Coefficients:
## (Intercept)      Ct_qPCR
##     0.33778     -0.00939
```

```
summary(lm(PerMapPairedReads10RefGenomes ~ Ct_qPCR, data = per_mapped_unenriched))
```

```
##
## Call:
## lm(formula = PerMapPairedReads10RefGenomes ~ Ct_qPCR, data = per_mapped_unenriched)
##
## Residuals:
##      Min       1Q   Median       3Q      Max
## -0.12896 -0.07954 -0.05543 -0.00933  0.89785
##
## Coefficients:
##              Estimate Std. Error t value Pr(>|t|)
## (Intercept)  0.337784   0.161488   2.092   0.0393 *
## Ct_qPCR     -0.009390   0.006337  -1.482   0.1419
## ---
## Signif. codes:  0 '***' 0.001 '**' 0.01 '*' 0.05 '.' 0.1 ' ' 1
##
## Residual standard error: 0.1667 on 89 degrees of freedom
##   (9 observations deleted due to missingness)
## Multiple R-squared:  0.02408,    Adjusted R-squared:  0.01311
## F-statistic: 2.196 on 1 and 89 DF,  p-value: 0.1419
```

```
x_range <- seq(min(X), max(X), length.out = 100)
x_range <- matrix(x_range, nrow = 100, ncol = 1)
xdf <- data.frame(x_range)
colnames(xdf) <- c("Ct_qPCR")

ydf <- lm_model %>%
    predict(xdf)

colnames(ydf) <- c("PerMapPairedReads10RefGenomes")
xy <- data.frame(xdf, ydf)

fig <- plot_ly(data = per_mapped_unenriched, x = ~Ct_qPCR, y = ~PerMapPairedReads10RefGenomes,
    type = "scatter", alpha = 0.65, mode = "markers", name = "% Reads Mapped to 10-genomes Reference")
fig <- fig %>%
    add_trace(data = xy, x = ~Ct_qPCR, y = ~PerMapPairedReads10RefGenomes,
        name = "prediction", mode = "lines", alpha = 1)
```

**NOTE:** The results from these analyses are presented in **Fig. 3C**.

We also tested the association between Ct values and the raw number of reads obtained via a linear regression model, using only single-enriched libraries.

```
# scatter plot of number of reads obtained and number of
# reads mapped
per_mapped_s_enriched <- clinext_table[c(101:200), c(3, 15)]

y <- per_mapped_s_enriched$PerMapPairedReads10RefGenomes
X <- per_mapped_s_enriched$Ct_qPCR

lm_model <- linear_reg() %>%
    set_engine("lm") %>%
    set_mode("regression") %>%
    fit(PerMapPairedReads10RefGenomes ~ Ct_qPCR, data = per_mapped_s_enriched)
lm_model
```

```
## parsnip model object
##
##
## Call:
## stats::lm(formula = PerMapPairedReads10RefGenomes ~ Ct_qPCR,
##     data = data)
##
## Coefficients:
## (Intercept)      Ct_qPCR
##      50.555       -1.289
```

```
summary(lm(PerMapPairedReads10RefGenomes ~ Ct_qPCR, data = per_mapped_s_enriched))
```

```
##
## Call:
## lm(formula = PerMapPairedReads10RefGenomes ~ Ct_qPCR, data = per_mapped_s_enriched)
##
```

```
## Residuals:
##     Min      1Q  Median      3Q     Max
## -17.795 -11.006  -3.508   6.201  49.415
##
## Coefficients:
##             Estimate Std. Error t value Pr(>|t|)
## (Intercept)  50.5555    13.3871   3.776 0.000287 ***
## Ct_qPCR      -1.2888     0.5253  -2.454 0.016094 *
## ---
## Signif. codes:  0 '***' 0.001 '**' 0.01 '*' 0.05 '.' 0.1 ' ' 1
##
## Residual standard error: 13.82 on 89 degrees of freedom
##   (9 observations deleted due to missingness)
## Multiple R-squared:  0.06335,    Adjusted R-squared:  0.05283
## F-statistic:  6.02 on 1 and 89 DF,  p-value: 0.01609
```

```
x_range <- seq(min(X), max(X), length.out = 100)
x_range <- matrix(x_range, nrow = 100, ncol = 1)
xdf <- data.frame(x_range)
colnames(xdf) <- c("Ct_qPCR")

ydf <- lm_model %>%
    predict(xdf)

colnames(ydf) <- c("PerMapPairedReads10RefGenomes")
xy <- data.frame(xdf, ydf)

fig <- plot_ly(data = per_mapped_s_enriched, x = ~Ct_qPCR, y = ~PerMapPairedReads10RefGenomes,
    type = "scatter", alpha = 0.65, mode = "markers", name = "% Reads Mapped to 10-genomes Reference")
fig <- fig %>%
    add_trace(data = xy, x = ~Ct_qPCR, y = ~PerMapPairedReads10RefGenomes,
        name = "prediction", mode = "lines", alpha = 1)
```

**NOTE:** The results from these analyses are presented in **Fig. 3D**.

We also tested the association between Ct values and the raw number of reads obtained via a linear regression model, using only double-enriched libraries.

```
# scatter plot of number of reads obtained and number of
# reads mapped
per_mapped_d_enriched <- clinext_table[c(201:300), c(3, 15)]

y <- per_mapped_d_enriched$PerMapPairedReads10RefGenomes
X <- per_mapped_d_enriched$Ct_qPCR

lm_model <- linear_reg() %>%
    set_engine("lm") %>%
    set_mode("regression") %>%
    fit(PerMapPairedReads10RefGenomes ~ Ct_qPCR, data = per_mapped_d_enriched)
lm_model
```

```
## parsnip model object
##
##
```

```
## Call:
## stats::lm(formula = PerMapPairedReads10RefGenomes ~ Ct_qPCR,
##     data = data)
##
## Coefficients:
## (Intercept)      Ct_qPCR
##     90.2645      -0.9663
```

```
summary(lm(PerMapPairedReads10RefGenomes ~ Ct_qPCR, data = per_mapped_d_enriched))
```

```
##
## Call:
## lm(formula = PerMapPairedReads10RefGenomes ~ Ct_qPCR, data = per_mapped_d_enriched)
##
## Residuals:
##     Min      1Q  Median      3Q     Max
## -51.342  -6.713   0.586  11.110  28.881
##
## Coefficients:
##             Estimate Std. Error t value Pr(>|t|)
## (Intercept)  90.2645    16.1260   5.597 2.38e-07 ***
## Ct_qPCR      -0.9663     0.6328  -1.527     0.13
## ---
## Signif. codes:  0 '***' 0.001 '**' 0.01 '*' 0.05 '.' 0.1 ' ' 1
##
## Residual standard error: 16.64 on 89 degrees of freedom
##   (9 observations deleted due to missingness)
## Multiple R-squared:  0.02553,    Adjusted R-squared:  0.01458
## F-statistic: 2.332 on 1 and 89 DF,  p-value: 0.1303
```

```
x_range <- seq(min(X), max(X), length.out = 100)
x_range <- matrix(x_range, nrow = 100, ncol = 1)
xdf <- data.frame(x_range)
colnames(xdf) <- c("Ct_qPCR")

ydf <- lm_model %>%
    predict(xdf)

colnames(ydf) <- c("PerMapPairedReads10RefGenomes")
xy <- data.frame(xdf, ydf)

fig <- plot_ly(data = per_mapped_d_enriched, x = ~Ct_qPCR, y = ~PerMapPairedReads10RefGenomes,
    type = "scatter", alpha = 0.65, mode = "markers", name = "% Reads Mapped to 10-genomes Reference")
fig <- fig %>%
    add_trace(data = xy, x = ~Ct_qPCR, y = ~PerMapPairedReads10RefGenomes,
        name = "prediction", mode = "lines", alpha = 1)
```

**NOTE:** The results from these analyses are presented in **Fig. 3E**.

12.2.2.2.3 Counting and plotting the proportion of hits to different species' genomes

Before importing each file from each sample, we delimited the first column by "-", separating the added species name.

```
awk -F '\t' '{gsub(/-.*/, "", $1); print}' sample_species_ct.txt > sample_species_ct_edited.txt
```

Below, we imported the edited files for each species and then created a fourth column, "Sample," with the name of the species for the uneriched libraries.

```
# Create an empty list to store the data frames
data_list_unenriched <- list()
# Loop through the list of file names in 'elist'
unlist <- gsub(" ", "", paste(clneb_table$Sample, "_U"))
for (i in unlist) {
    # Construct the file path
    file_path <- paste("~/Documents/crypto_data/12/iNextEra/species_edited/",
        i, "_species.txt", sep = "")

    # Check if the file exists and has data (not empty)
    if (file.exists(file_path) && file.info(file_path)$size > 0) {
        # Read the file into a data frame
        data <- read.table(file_path, header = FALSE, sep = "\t", stringsAsFactors = FALSE)

        # Remove columns 3 to 5
        data <- data[, -c(3:5)]

        # Rename columns
        colnames(data) <- c("Species", "Count")

        # Calculate Relative abundance of counts for each genome
        data <- data %>%
            mutate(relative_ab = Count/sum(Count))

        # Add the processed data frame to the list with the file name as the
        # element name
        data_list_unenriched[[i]] <- data
    }
}

# Combine the data frames in data_list into a single data frame
combined_data_unenriched <- bind_rows(data_list_unenriched, .id = "Sample")

# Sort the data by Sample and Species
combined_data_unenriched <- combined_data_unenriched %>%
    arrange(Sample, Species)

# Create the plot
p <- plot_ly(data = combined_data_unenriched, x = ~Sample, y = ~relative_ab, color = ~Species,
    colors = col_c_ten, type = "bar") %>%
    layout(yaxis = list(title = "Relative Abundance"), barmode = "stack")
```

**NOTE:** The results from these analyses are presented in **Fig. 3F**.

We then plotted species assignments for single-enriched reads.

```
# Create an empty list to store the data frames
data_list_enriched <- list()
```

```r
# Loop through the list of file names in 'elist'
for (i in elist) {
    # Construct the file path
    file_path <- paste("~/crypto_data/12/iNextEra/species_edited/",
        i, "_species.txt", sep = "")

    # Check if the file exists and has data (not empty)
    if (file.exists(file_path) && file.info(file_path)$size >
        0) {
        # Read the file into a data frame
        data <- read.table(file_path, header = FALSE, sep = "\t",
            stringsAsFactors = FALSE)

        # Remove columns 3 to 5
        data <- data[, -c(3:5)]

        # Rename columns
        colnames(data) <- c("Species", "Count")

        # Calculate Relative abundance of counts for each
        # genome
        data <- data %>%
            mutate(relative_ab = Count/sum(Count))

        # Add the processed data frame to the list with the
        # file name as the list element name
        data_list_enriched[[i]] <- data
    }
}

# Combine the data frames in data_list into a single data
# frame
combined_data_enriched <- bind_rows(data_list_enriched, .id = "Sample")

# Sort the data by Sample and Species
combined_data_enriched <- combined_data_enriched %>%
    arrange(Sample, Species)

# Create the plot
p <- plot_ly(data = combined_data_enriched, x = ~Sample, y = ~relative_ab,
    color = ~Species, colors = col_c_ten, type = "bar") %>%
    layout(yaxis = list(title = "Relative Abundance"), barmode = "stack")
```

**NOTE:** The results from these analyses are presented in **Fig. 3G**.

And then, we followed the same procedure for the double-enriched libraries that got hits against the 10-genomes reference.

```r
# Create an empty list to store the data frames
data_list_enriched <- list()
delist <- gsub(" ", "", paste(clneb_table$Sample, "_DE"))
# Loop through the list of file names in 'elist'
for (i in delist) {
    # Construct the file path
```

```r
    file_path <- paste("~/crypto_data/12/iNextEra/double_species_edited/",
        i, "_species.txt", sep = "")

    # Check if the file exists and has data (not empty)
    if (file.exists(file_path) && file.info(file_path)$size >
        0) {
        # Read the file into a data frame
        data <- read.table(file_path, header = FALSE, sep = "\t",
            stringsAsFactors = FALSE)

        # Remove columns 3 to 5
        data <- data[, -c(3:5)]

        # Rename columns
        colnames(data) <- c("Species", "Count")

        # Calculate Relative abundance of counts for each
        # genome
        data <- data %>%
            mutate(relative_ab = Count/sum(Count))

        # Add the processed data frame to the list with the
        # file name as the list element name
        data_list_enriched[[i]] <- data
    }
}

# Combine the data frames in data_list into a single data
# frame
combined_data_double_enriched <- bind_rows(data_list_enriched,
    .id = "Sample")

# Sort the data by Sample and Species
combined_data_double_enriched <- combined_data_double_enriched %>%
    arrange(Sample, Species)

# Create the plot
p <- plot_ly(data = combined_data_double_enriched, x = ~Sample,
    y = ~relative_ab, color = ~Species, colors = col_c_ten, type = "bar") %>%
    layout(yaxis = list(title = "Relative Abundance"), barmode = "stack")
```

**NOTE:** The results from these analyses are presented in **Fig. 3H**.

When mapping the clinical samples prepared with the iNextEra protocol to the ten-genomes reference database. Sixty-six of 100 samples had a majority (>50%) of reads mapping to *C. parvum*. Similarly, five samples (UKH145, UKH146, UKH147, UKH148, and UKH150), the last four overlapping with NEB-iTru results, contain a high proportion (>50%) of *C. hominis* (Fig. 3F) and show moderate proportions of *C. cuniculus*. Of the 91 unenriched iNextEra samples with sequence data, 89 mapped to the ten-genomes database, a higher proportion than NEB (Table S15). Similarly to the NEB-iTru libraries, the analysis shows a drastic differential assignment of species between single-enriched and unenriched samples (Fig. 3F vs. 3G). However, the actual assignments differ: 10 samples had >50% of *C. parvum* reads, and 23 samples contained >50% of *C. hominis* reads (Fig. 3G). A majority of *C. hominis* samples were detected in the unenriched, showing a mismatch in species assignments from those in the enriched sets. The relative abundance of species for the double enriched libraries shows that 86 of the 93 for which we got data are assigned in the majority

(>50%) to *C. parvum*, even with higher proportions than those observed from single enrichments. Here, six samples (UKH145, UKH146, UKH147, UKH148, UKH149, and UKH150) contain a high proportion (>50%) of *C. hominis* (Fig. 3G), which also show moderate proportions of *C. cuniculus* (14-25%). These patterns are similar to those from single enrichments with iNextEra, except that five and not six samples were detected in single enrichments with *C. hominis* (all except UKH149). All seven other *Cryptosporidium* species had relative abundances of less than 1.3%.

###### 12.2.3 Comparison between single-enriched clinical samples with iNextEra and NEB libraries

The summary statistics from these results can be found in **Table S16**, also imported below.

```
file = "~/crypto_data/12/clinical_samples_iTruvsiNext_summary_stats.txt" cl_next_table = read.table(file
header = TRUE, sep = "")
```

We tested if the number of raw paired reads obtained for clinical samples using two protocols, NEB-iTru and iNextEra-iNext, and then enriched with CryptoCap_100k, were different.

```
lsmeans(lm(data = cl_next_table, RawReadPairs ~ Protocol), pairwise ~
    Protocol)
```

```
## $lsmeans
##  Protocol  lsmean     SE  df lower.CL upper.CL
##  iNextEra 5298444 518000 189  4277591  6319297
##  NEB-iTru 2603198 494000 189  1629366  3577029
##
## Confidence level used: 0.95
##
## $contrasts
##  contrast              estimate     SE  df t.ratio p.value
##  iNextEra - (NEB-iTru)  2695246 715000 189   3.768  0.0002
```

```
my_palette <- c("#fdb462", "#c51b7d")
p <- plot_ly(colors = my_palette)
p <- add_bars(p, x = c("iNextEra", "iNextEra0", "NEB-iTru"),
    y = c(3e+07, 3.7e+07, 3e+07), opacity = 1, showlegend = F,
    marker = list(line = list(color = "rgba(0,0,0,0"), color = "rgba(0,0,0,0"),
    text = c("", "*** p-value = 0.0002", ""), textposition = "outside",
    legendgroup = "1")
p <- add_lines(p, x = c("iNextEra", "iNextEra", "NEB-iTru", "NEB-iTru"),
    y = c(3.5e+07, 3.4e+07, 3.4e+07, 3.5e+07), showlegend = F,
    line = list(color = "black"), legendgroup = "1", hoverinfo = "none")
p <- add_boxplot(p, data = cl_next_table, x = ~Protocol, y = ~RawReadPairs,
    color = ~Protocol, colormodel = c("rgba(197,27,125"), type = "box",
    boxpoints = "all", jitter = 0.3, pointpos = 0, legendgroup = "1")
p <- layout(p, xaxis = list(tickmode = "array", tickvals = c("iNextEra",
    "sf", "NEB-iTru"), ticktext = c("iNextEra", "", "NEB-iTru")),
    yaxis = list(range = c(-1, 4e+07)))
```

**NOTE:** The results from these analyses are presented in **Fig. S43**.

We tested if the average percentage of reads retained after filtering was different between the two protocols NEB-iTru and iNextEra-iNext.

```r
lsmeans(lm(data = cl_next_table, PerReadsRetained ~ Protocol), pairwise ~ Protocol)
```

```
## $lsmeans
##  Protocol lsmean    SE  df lower.CL upper.CL
##  iNextEra  0.908 0.428 189    0.063     1.75
##  NEB-iTru 83.873 0.409 189   83.067    84.68
##
## Confidence level used: 0.95
##
## $contrasts
##  contrast              estimate    SE  df  t.ratio p.value
##  iNextEra - (NEB-iTru)      -83 0.592 189 -140.127  <.0001
```

We also tested if the number of retained reads after filtering was different between the two protocols, NEB-iTru and iNextEra-iNext.

```r
lsmeans(lm(data = cl_next_table, TrimmedReadPairs ~ Protocol), pairwise ~ Protocol)
```

```
## $lsmeans
##  Protocol  lsmean     SE  df lower.CL upper.CL
##  iNextEra 4725889 441000 189  3856782  5594997
##  NEB-iTru 2151983 420000 189  1322907  2981060
##
## Confidence level used: 0.95
##
## $contrasts
##  contrast              estimate     SE  df t.ratio p.value
##  iNextEra - (NEB-iTru)  2573906 609000 189   4.227  <.0001
```

We also tested if the percentage of reads mapped to *C.parvum* was different between the two protocols NEB-iTru and iNextEra-iNext.

```r
lsmeans(lm(data = cl_next_table, PerMapPairedReadsCparvum ~ Protocol),
    pairwise ~ Protocol)
```

```
## $lsmeans
##  Protocol lsmean   SE  df lower.CL upper.CL
##  iNextEra   15.9 1.89 189     12.1     19.6
##  NEB-iTru   35.6 1.81 189     32.1     39.2
##
## Confidence level used: 0.95
##
## $contrasts
##  contrast              estimate   SE  df t.ratio p.value
##  iNextEra - (NEB-iTru)    -19.8 2.62 189  -7.553  <.0001
```

```r
p <- plot_ly(colors = my_palette)
p <- add_boxplot(p, data = cl_next_table, x = ~Sample, y = ~PerMapPairedReadsCparvum,
    color = ~Protocol, boxpoints = "all", jitter = 0.3, pointpos = 0,
    legendgroup = "1")
p <- layout(p, xaxis = list(tickmode = "array", tickvals = c("iNextEra",
    "sf", "NEB-iTru"), ticktext = c("iNextEra", "", "NEB-iTru")),
    yaxis = list(range = c(-1, 100)), barmode = "stack")
```

**NOTE:** The results from these analyses are presented in **Fig. S44**.

We tested if the depth of genome coverage obtained by the mapping of sequence reads to *C. parvum* was different between enriched and unenriched clinical samples.

```
lsmeans(lm(data = cl_next_table, AverageDepthCparvum ~ Protocol),
    pairwise ~ Protocol)
```

```
## $lsmeans
##  Protocol lsmean   SE  df lower.CL upper.CL
##  iNextEra   32.1 7.26 189     17.8     46.4
##  NEB-iTru   28.1 6.93 189     14.4     41.8
##
## Confidence level used: 0.95
##
## $contrasts
##  contrast              estimate SE  df t.ratio p.value
##  iNextEra - (NEB-iTru)     4.01 10 189   0.400  0.6896
```

```
p <- plot_ly(colors = my_palette)
p <- add_bars(p, x = c("iNextEra", "iNextEra0", "NEB-iTru"),
    y = c(690, 695, 690), opacity = 1, showlegend = F, marker = list(line = list(color = "rgba(0,0,0,0"
        color = "rgba(0,0,0,0"), text = c("", "p-value = 0.690",
        ""), textposition = "outside", legendgroup = "1")
p <- add_lines(p, x = c("iNextEra", "iNextEra", "NEB-iTru", "NEB-iTru"),
    y = c(689, 685, 685, 689), showlegend = F, line = list(color = "black"),
    legendgroup = "1", hoverinfo = "none")
p <- add_boxplot(p, data = cl_next_table, x = ~Protocol, y = ~AverageDepthCparvum,
    color = ~Protocol, colormodel = c("rgba(197,27,125"), type = "box",
    boxpoints = "all", jitter = 0.3, pointpos = 0, legendgroup = "1")
p <- layout(p, xaxis = list(tickmode = "array", tickvals = c("iNextEra",
    "sf", "NEB-iTru"), ticktext = c("iNextEra", "", "NEB-iTru")),
    yaxis = list(range = c(-1, 730)))
```

**NOTE:** The results from these analyses are presented in **Fig. S45**.

We also tested whether the breadth of genome coverage from mapping sequence reads to *C. parvum* was different between enriched and unenriched clinical samples.

```
lsmeans(lm(data = cl_next_table, AverageBreadthCparvum ~ Protocol),
    pairwise ~ Protocol)
```

```
## $lsmeans
##  Protocol lsmean   SE  df lower.CL upper.CL
##  iNextEra   54.4 3.02 189     48.4     60.3
##  NEB-iTru   31.3 2.88 189     25.6     37.0
##
## Confidence level used: 0.95
##
## $contrasts
##  contrast              estimate   SE  df t.ratio p.value
##  iNextEra - (NEB-iTru)     23.1 4.17 189   5.537  <.0001
```

```
p <- plot_ly(colors = my_palette)
p <- add_bars(p, x = c("iNextEra", "iNextEra0", "NEB-iTru"),
    y = c(110, 115, 110), opacity = 1, showlegend = F, marker = list(line = list(color = "rgba(0,0,0,0"
        color = "rgba(0,0,0,0"), text = c("", "***p-value < 0.0001",
        ""), textposition = "outside", legendgroup = "1")
p <- add_lines(p, x = c("iNextEra", "iNextEra", "NEB-iTru", "NEB-iTru"),
    y = c(109, 105, 105, 109), showlegend = F, line = list(color = "black"),
    legendgroup = "1", hoverinfo = "none")
p <- add_boxplot(p, data = cl_next_table, x = ~Protocol, y = ~AverageBreadthCparvum,
    color = ~Protocol, colormodel = c("rgba(197,27,125"), type = "box",
    boxpoints = "all", jitter = 0.3, pointpos = 0, legendgroup = "1")
p <- layout(p, xaxis = list(tickmode = "array", tickvals = c("iNextEra",
    "sf", "NEB-iTru"), ticktext = c("iNextEra", "", "NEB-iTru")),
    yaxis = list(range = c(-1, 125)))
```

**NOTE:** The results from these analyses are presented in **Fig. S47**.

We tested how the depth and breadth of genome coverage varied for the library types once we normalized for the number of sequences used in the mapping.

```
lsmeans(lm(data = cl_next_table, Normalized_Depth_Cparvum ~ Protocol),
    pairwise ~ Protocol)
```

```
## $lsmeans
##  Protocol lsmean     SE  df lower.CL upper.CL
##  iNextEra  0.564 0.0804 189    0.405    0.722
##  NEB-iTru  0.596 0.0767 189    0.445    0.747
##
## Confidence level used: 0.95
##
## $contrasts
##  contrast              estimate    SE  df t.ratio p.value
##  iNextEra - (NEB-iTru)  -0.0323 0.111 189  -0.291  0.7718
```

```
depth <- plot_ly(colors = my_palette)
depth <- add_bars(depth, x = c("iNextEra", "iNextEra0", "NEB-iTru"),
    y = c(9, 9.5, 9), opacity = 1, showlegend = F, marker = list(line = list(color = "rgba(0,0,0,0"),
        color = "rgba(0,0,0,0"), text = c("", "p-value = 0.772",
        ""), textposition = "outside", legendgroup = "1")
depth <- add_lines(depth, x = c("iNextEra", "iNextEra", "NEB-iTru",
    "NEB-iTru"), y = c(9, 8.5, 8.5, 9), showlegend = F, line = list(color = "black"),
    legendgroup = "1", hoverinfo = "none")
depth <- add_boxplot(depth, data = cl_next_table, x = ~Protocol,
    y = ~Normalized_Depth_Cparvum, color = ~Protocol, colormodel = c("rgba(197,27,125"),
    type = "box", boxpoints = "all", jitter = 0.3, pointpos = 0,
    legendgroup = "1")
depth <- layout(depth, xaxis = list(tickmode = "array", tickvals = c("iNextEra",
    "sf", "NEB-iTru"), ticktext = c("iNextEra", "", "NEB-iTru")),
    yaxis = list(range = c(-1, 11)))
```

We also tested whether the breadth of genome coverage from mapping sequence reads to *C. parvum* was different between enriched and unenriched clinical samples.

```r
lsmeans(lm(data = cl_next_table, Normalized_Breadth_Cparvum ~
    Protocol), pairwise ~ Protocol)
```

```
## $lsmeans
##  Protocol lsmean    SE  df lower.CL upper.CL
##  iNextEra   1.79 0.110 189    1.574    2.007
##  NEB-iTru   0.00 0.105 189   -0.207    0.207
##
## Confidence level used: 0.95
##
## $contrasts
##  contrast              estimate    SE  df t.ratio p.value
##  iNextEra - (NEB-iTru)     1.79 0.152 189  11.795  <.0001
```

```r
breadth <- plot_ly(colors = my_palette)
breadth <- add_bars(breadth, x = c("iNextEra", "iNextEra0", "NEB-iTru"),
    y = c(9, 9.5, 9), opacity = 1, showlegend = F, marker = list(line = list(color = "rgba(0,0,0,0"),
        color = "rgba(0,0,0,0"), text = c("", "***p-value < 0.0001",
        ""), textposition = "outside", legendgroup = "1")
breadth <- add_lines(breadth, x = c("iNextEra", "iNextEra", "NEB-iTru",
    "NEB-iTru"), y = c(9, 8.5, 8.5, 9), showlegend = F, line = list(color = "black"),
    legendgroup = "1", hoverinfo = "none")
breadth <- add_boxplot(breadth, data = cl_next_table, x = ~Protocol,
    y = ~Normalized_Breadth_Cparvum, color = ~Protocol, colormodel = c("rgba(197,27,125"),
    type = "box", boxpoints = "all", jitter = 0.3, pointpos = 0,
    legendgroup = "1")
breadth <- layout(breadth, xaxis = list(tickmode = "array", tickvals = c("iNextEra",
    "sf", "NEB-iTru"), ticktext = c("iNextEra", "", "NEB-iTru")),
    yaxis = list(range = c(-1, 11)))
breadth
```

```r
# creating subplot
fig <- subplot(depth, breadth, nrows = 1, titleY = TRUE, titleX = TRUE,
    margin = 0.1)
```

**NOTE:** The results from these analyses are presented in **Fig. S46**.

iNextEra and NEB-iTru libraries performed similarly (Table S16) but with noteworthy differences. For single-enriched libraries, we obtained an average of 2.04X more reads and retained more reads following quality filtering for iNextEra than NEB-iTru libraries (Table S16; Fig. S43). However, a higher percentage of reads mapped to the *C. parvum* genome sequence in samples from NEB-iTru vs. iNextEra libraries (35.6% vs. 15.9%, respectively; p-value $< 0.0001$; Fig. S44), which resulted in NEB-iTru libraries yielding an average enrichment factor about twice as large as iNextEra libraries (Table S16). The average depth of coverage (based on total reads obtained) of *C. parvum* was not significantly different between the two protocols (p-value $= 0.690$). The normalized depth of coverage is very similar for both protocols (0.5X, p-value $=$ 0.7718; non-normalized values in Fig. S45, normalized values in Fig. S46). However, the total breadth of genome coverage was higher for iNextEra samples (p-value $< 0.0001$); the same as the normalized breadth of coverage was significantly higher for the iNextEra libraries (p-value $< 0.0001$; non-normalized values in Fig. S47, normalized values in Fig. S46).

#### Supplementary Notes #13. Population Genomic Analyses of Clinical Samples

##### 13.1 Filtering and read mapping against *C. parvum* genome

To further examine the genetic variation obtained from the CrytoCap_100K approach in natural populations of *Cryptosporidium* from clinical samples in the UK, the evolutionary relationships of these samples were studied. For these sets of analyses, samples from the iNextEra protocol were batched by library type (i.e., unenriched, single-enriched, and double-enriched). All samples were mapped individually to the *Cryptosporidium parvum* genome (ATCC) using BWA v.0.7.17.

> time bwa mem -M $ref_genome.fasta *inputs*/i_$x_R1_pair_trim.fastq.gz *inputs*/i_*x_R2pairtrim.fastq.gz* > ./*bwaaligned*/i_$x.sam

##### 13.2 Raw depth and breadth of coverage

From the mapped reads, *bigwig* files were obtained using SAMTools v.1.16.1 and deepTools v.3.5.2 and visualized for each sample, for each library type. For this, we used the Integrative Genomics Viewer (IGV) Web app version 2.1.2 and igv.js version 3.1.2.

> samtools view -b ./bwa_aligned/*i*_x.sam | samtools sort > ./bwa_aligned/*i*_x.bam samtools index ./bwa_aligned/*i*_x.bam bamCoverage -b ./bwa_aligned/*i*_x.bam -o ./bwa_aligned/*i*_x.bw

##### 13.3 SNP calling and filtering

We followed the best practices pipeline from GATK (https://gatk.broadinstitute.org/hc/en-us) for variant calling.
From the .bam files obtained from the mapping of clinical samples against *C. parvum* IOWA ATCC reference genome, we move to GATK v4.6.0.0 to:

**13.3.1 From raw reads to analysis-ready reads** Illumina reads mapping information from each bam file was assigned to read groups using their sample ID:

> gatk –java-options "-Xmx20g" AddOrReplaceReadGroups –INPUT *.bam –OUTPUT* .rg.bam –RGLB lib1 –RGPL illumina –RGPU unit1 –RGSM 20

Then, these assigned bam files were used to flag (do not remove) duplicates using MarkDuplicatesSpark:

> gatk –java-options "-Xmx20g" MarkDuplicatesSpark –spark-runner LOCAL –input *BAMdudup*/i_$x.rg.bam –output *BAMdudup*/i_$x.md.bam –conf 'spark.executor.cores=4'

Then, these bam files were used to calculate the mismatch base count (NM), mismatch bases (MD), and the unaligned quality (UQ) tags by comparing with the reference.

> gatk –java-options "-Xmx20g" SetNmMdAndUqTags –INPUT *BAMdudup*/i_$x.md.bam –OUTPUT *BAMdudup*/i_$x.md.fx.bam –REFERENCE_SEQUENCE $REF

Finally, these bam files were indexed. > gatk –java-options "-Xmx20g" BuildBamIndex –INPUT *BAMdudup*/i_$x.md.fx.bam

**13.3.2 Calling variants per-sample and consolidation of all samples** For variant calling per sample, HaplotypeCaller was used for local re-assembly of haplotypes. We ran this software by setting the minimum phred-scaled confidence threshold at which variants should be called at 30, ploidy of 1 (given that *Cryptosporidium* has haploid genomes), and using both the VCGF and the BP_Resolution modes for emitting reference confidence scores.

```
gatk –java-options “-Xmx100G -Djava.io.tmpdir=$tempdir” HaplotypeCaller
-R $REF
-I BAMdudup/i_$x.md.fx.bam
-O vcf_gvcf_mode/i_$x.raw.snps.indels.g.vcf
–emit-ref-confidence GVCF
-stand-call-conf 30.0
–native-pair-hmm-threads 10
–sample-ploidy 1

gatk –java-options “-Xmx100G -Djava.io.tmpdir=$tempdir” HaplotypeCaller
-R $REF
-I BAMdudup/i_$x.md.fx.bam
-O vcf_bp_mode/i_$x.raw.snps.indels.vcf
-ERC BP_RESOLUTION
-stand-call-conf 30.0
–native-pair-hmm-threads 10
–sample-ploidy 1
```

To merge all single-sample GVCFs into one database, we converted our reference file to a .bed coordinates file and then used GenomicsDBImport for each batch of library type:

```
awk ‘BEGIN {FS=“”̂}; {print $1 FS”0” FS $2}’ C_parvum.fasta.fai > C_parvum.bed

gatk –java-options “-Xmx10g” GenomicsDBImport
–genomicsdb-workspace-path my_database_x
–sample-name-map gvcfs-for-db-import.library_type_sample_map_mg
–tmp-dir ./tmp
–batch-size 32
-L C_parvum.bed
```

**13.3.3 Joint genotyping and variant selection and filtering** Finally, to perform joint genotyping on the group of samples pre-called with HaplotypeCaller for each library type we used GenotypeGVCFs:

```
gatk –java-options “-Xmx20g”
GenotypeGVCFs
-R C_parvum.fasta
-V gendb://my_database_x
-O workingdir/ref_output_x_mg.vcf.gz
```

We then selected SNP variants from the above VCF file and flagged variants based on filters for QualByDepth (QD), FisherStrand (FS), RMSMappingQuality (MQ), MQRankSum, and ReadPosRankSumTest, following the lenient hard-filtering recommendations from GATK documentation:

```
gatk –java-options “-Xmx100G -Djava.io.tmpdir=$tempdir” SelectVariants
-R REF
-V workingdir/ref_output_x.vcf.gz
–select-type-to-include SNP
-O workingdir/raw_snps_gatk_all_x.g.vcf
```

gatk –java-options “-Xmx100G -Djava.io.tmpdir=$tempdir” VariantFiltration
-R REF
-V workingdir/raw_snps_gatk_all_x.g.vcf
–filter-name “filter_QD” –filter-expression “QD < 2.0”
–filter-name “filter_FS” –filter-expression “FS > 60.0”
–filter-name “filter_MQ” –filter-expression “MQ < 40.0”
–filter-name “filter_MQRankSum” –filter-expression “MQRankSum < -12.5”
–filter-name “filter_ReadPosRankSum” –filter-expression “ReadPosRankSum < -8.0”
-O workingdir/filtered_snps_gatk_all_x.g.vcf

##### 13.4 Exploring and filtering raw VCF

**13.4.1 Exploring raw VCFs** Raw VCF files contained 91 samples and 94,783 SNPs for double enrichments (45.2% missing data, Figs. S48–S49), 127,102 SNPs for single enrichments (42% missing data, Figs. S50–S51), and 15,430 SNPs for unenriched libraries (91.8% missing data, Figs. S52–S53). The read depth (DP) used for variant calling of genomic positions differed widely between enriched and unenriched raw VCF files of enriched and unenriched libraries, being on average 151.6 for double enrichments, 184.6 for single enrichments, and 13 for unenriched (Fig. S54).

**13.4.1.1 Exploring double enrichment VCF** The VCF from GATK for double enrichments:

```
library(vcfR)
```

```
##
##    *****       ***   vcfR   ***       *****
##    This is vcfR 1.15.0
##      browseVignettes('vcfR') # Documentation
##      citation('vcfR') # Citation
##    *****       *****      *****       *****
```

```
library(ggplot2)
library(plotly)

# Importing VCF containing 94,783 variants and 45.18% missing data
vcf_double <- read.vcfR("~/crypto_data/13/filtered_snps_gatk_all_D.g.vcf"
```

Scanning file to determine attributes. File attributes: meta lines: 49 header_line: 50 variant count: 94783 column count: 100 Meta line 49 read in. All meta lines processed. gt matrix initialized. Character matrix gt created. Character matrix gt rows: 94783 Character matrix gt cols: 100 skip: 0 nrows: 94783 row_num: 0 Processed variant 1000Processed variant 2000Processed variant 3000Processed variant 4000Processed variant 5000Processed variant 6000Processed variant 7000Processed variant 8000Processed variant 9000Processed variant 10000Processed variant 11000Processed variant 12000Processed variant 13000Processed variant 14000Processed variant 15000Processed variant 16000Processed variant 17000Processed variant 18000Processed variant 19000Processed variant 20000Processed variant 21000Processed variant 22000Processed variant 23000Processed variant 24000Processed variant 25000Processed variant 26000Processed variant 27000Processed variant 28000Processed variant 29000Processed variant 30000Processed variant 31000Processed variant 32000Processed variant 33000Processed variant 34000Processed variant 35000Processed variant 36000Processed variant 37000Processed variant 38000Processed variant 39000Processed variant 40000Processed variant 41000Processed variant 42000Processed variant 43000Processed variant 44000Processed variant 45000Processed variant 46000Processed variant 47000Processed variant 48000Processed variant 49000Processed variant 50000Processed variant 51000Processed variant 52000Processed variant 53000Processed variant 54000Processed variant 55000Processed variant 56000Processed variant 57000Processed variant 58000Processed variant

59000Processed variant 60000Processed variant 61000Processed variant 62000Processed variant 63000Processed variant 64000Processed variant 65000Processed variant 66000Processed variant 67000Processed variant 68000Processed variant 69000Processed variant 70000Processed variant 71000Processed variant 72000Processed variant 73000Processed variant 74000Processed variant 75000Processed variant 76000Processed variant 77000Processed variant 78000Processed variant 79000Processed variant 80000Processed variant 81000Processed variant 82000Processed variant 83000Processed variant 84000Processed variant 85000Processed variant 86000Processed variant 87000Processed variant 88000Processed variant 89000Processed variant 90000Processed variant 91000Processed variant 92000Processed variant 93000Processed variant 94000Processed variant: 94783 All variants processed

```
## Extract the FILTER column and subset the VCF for variants with 'PASS' in the
## FILTER column
filter_column <- vcf_double@fix[, "FILTER"]
# This filtered VCF contains 94,422 variants and 45.21% missing data
vcf_pass_double <- vcf_double[filter_column == "PASS", ]

# Extract the DP (Depth of Coverage) values from the INFO column
dp_info_double <- extract.info(vcf_pass_double, element = "DP")
dp_info_double_numeric <- as.numeric(dp_info_double)

# Extract the DP values from the FORMAT column for each individual
dp_format_double <- extract.gt(vcf_pass_double, element = "DP", as.numeric = TRUE)

coverage_double_df <- as.data.frame(dp_format_double)
coverage_double_df <- reshape2::melt(coverage_double_df, variable.name = "Individual",
    value.name = "Depth")
```

```
## No id variables; using all as measure variables
```

```
# Remove NA values from the data
coverage_double_df <- na.omit(coverage_double_df)

# Create the boxplot using Plotly
boxplot_coverage_double <- plot_ly(data = coverage_double_df, x = ~Individual, y = ~Depth,
    type = "box", marker = list(color = "#66C2A5")) %>%
    layout(xaxis = list(title = "Sample", tickangle = 45, tickfont = list(size = 10)),
        yaxis = list(title = "Depth of Coverage"), showlegend = FALSE)
```

**NOTE:** The results from these analyses are presented in **Fig. S48**.

```
# Extract chromosome and position info
chrom_double <- vcf_pass_double@fix[, "CHROM"]

# Aggregate mean depth per chromosome
mean_depth_per_chrom_double <- tapply(dp_info_double_numeric, chrom_double, mean,
    na.rm = TRUE)
coverage_chrom_double <- data.frame(Chromosome = names(mean_depth_per_chrom_double),
    MeanCoverage = mean_depth_per_chrom_double)

# Barplot of mean depth per chromosome
plot_coverage_chrom_double <- plot_ly(data = coverage_chrom_double, x = ~Chromosome,
    y = ~MeanCoverage, type = "bar", marker = list(color = "#66C2A5")) %>%
    layout(xaxis = list(title = "Chromosome", tickangle = 45, tickfont = list(size = 10)),
        yaxis = list(title = "Mean Depth"), bargap = 0.2)
```

**NOTE:** The results from these analyses are presented in **Fig. S49**.

**13.4.1.2 Exploring single enrichment VCF** The VCF from GATK for single enrichments:

```
# Importing VCF containing 127,102 variants and 41.92% missing data
vcf_single <- read.vcfR("~/crypto_data/13/filtered_snps_gatk_all_S.g.vcf"
```

Scanning file to determine attributes. File attributes: meta lines: 49 header_line: 50 variant count: 127102 column count: 100 Meta line 49 read in. All meta lines processed. gt matrix initialized. Character matrix gt created. Character matrix gt rows: 127102 Character matrix gt cols: 100 skip: 0 nrows: 127102 row_num: 0 Processed variant 1000Processed variant 2000Processed variant 3000Processed variant 4000Processed variant 5000Processed variant 6000Processed variant 7000Processed variant 8000Processed variant 9000Processed variant 10000Processed variant 11000Processed variant 12000Processed variant 13000Processed variant 14000Processed variant 15000Processed variant 16000Processed variant 17000Processed variant 18000Processed variant 19000Processed variant 20000Processed variant 21000Processed variant 22000Processed variant 23000Processed variant 24000Processed variant 25000Processed variant 26000Processed variant 27000Processed variant 28000Processed variant 29000Processed variant 30000Processed variant 31000Processed variant 32000Processed variant 33000Processed variant 34000Processed variant 35000Processed variant 36000Processed variant 37000Processed variant 38000Processed variant 39000Processed variant 40000Processed variant 41000Processed variant 42000Processed variant 43000Processed variant 44000Processed variant 45000Processed variant 46000Processed variant 47000Processed variant 48000Processed variant 49000Processed variant 50000Processed variant 51000Processed variant 52000Processed variant 53000Processed variant 54000Processed variant 55000Processed variant 56000Processed variant 57000Processed variant 58000Processed variant 59000Processed variant 60000Processed variant 61000Processed variant 62000Processed variant 63000Processed variant 64000Processed variant 65000Processed variant 66000Processed variant 67000Processed variant 68000Processed variant 69000Processed variant 70000Processed variant 71000Processed variant 72000Processed variant 73000Processed variant 74000Processed variant 75000Processed variant 76000Processed variant 77000Processed variant 78000Processed variant 79000Processed variant 80000Processed variant 81000Processed variant 82000Processed variant 83000Processed variant 84000Processed variant 85000Processed variant 86000Processed variant 87000Processed variant 88000Processed variant 89000Processed variant 90000Processed variant 91000Processed variant 92000Processed variant 93000Processed variant 94000Processed variant 95000Processed variant 96000Processed variant 97000Processed variant 98000Processed variant 99000Processed variant 100000Processed variant 101000Processed variant 102000Processed variant 103000Processed variant 104000Processed variant 105000Processed variant 106000Processed variant 107000Processed variant 108000Processed variant 109000Processed variant 110000Processed variant 111000Processed variant 112000Processed variant 113000Processed variant 114000Processed variant 115000Processed variant 116000Processed variant 117000Processed variant 118000Processed variant 119000Processed variant 120000Processed variant 121000Processed variant 122000Processed variant 123000Processed variant 124000Processed variant 125000Processed variant 126000Processed variant 127000Processed variant: 127102 All variants processed

```
## Extract the FILTER column and subset the VCF for variants with 'PASS' in the
## FILTER column
filter_column <- vcf_single@fix[, "FILTER"]
# This filtered VCF contains 126,598 variants and 41.96% missing data
vcf_pass_single <- vcf_single[filter_column == "PASS", ]

# Extract the DP (Depth of Coverage) values from the INFO column
dp_info_single <- extract.info(vcf_pass_single, element = "DP")
dp_info_single_numeric <- as.numeric(dp_info_single)

# Extract the DP values from the FORMAT column for each individual
dp_format_single <- extract.gt(vcf_pass_single, element = "DP", as.numeric = TRUE)
```

```r
coverage_single_df <- as.data.frame(dp_format_single)
coverage_single_df <- reshape2::melt(coverage_single_df, variable.name = "Individual",
    value.name = "Depth")
```

```
## No id variables; using all as measure variables
```

```r
# Remove NA values from the data
coverage_single_df <- na.omit(coverage_single_df)

# Create the boxplot using Plotly
boxplot_coverage_single <- plot_ly(data = coverage_single_df, x = ~Individual, y = ~Depth,
    type = "box", marker = list(color = "#FC8D62")) %>%
    layout(xaxis = list(title = "Sample", tickangle = 45, tickfont = list(size = 10)),
        yaxis = list(title = "Depth of Coverage"), showlegend = FALSE)
```

**NOTE:** The results from these analyses are presented in **Fig. S50**.

```r
# Extract chromosome and position info
chrom_single <- vcf_pass_single@fix[, "CHROM"]

# Aggregate mean depth per chromosome
mean_depth_per_chrom_single <- tapply(dp_info_single_numeric, chrom_single, mean,
    na.rm = TRUE)
coverage_chrom_single <- data.frame(Chromosome = names(mean_depth_per_chrom_single),
    MeanCoverage = mean_depth_per_chrom_single)

# Barplot of mean depth per chromosome
plot_coverage_chrom_single <- plot_ly(data = coverage_chrom_single, x = ~Chromosome,
    y = ~MeanCoverage, type = "bar", marker = list(color = "#FC8D62")) %>%
    layout(xaxis = list(title = "Chromosome", tickangle = 45, tickfont = list(size = 10)),
        yaxis = list(title = "Mean Depth"), bargap = 0.2)
```

**NOTE:** The results from these analyses are presented in **Fig. S51**.

**13.4.1.3 Exploring unenrichment VCF** The VCF from GATK from unenriched libraries:

```r
# Importing VCF containing 15,430 variants and 91.77% missing data
vcf_unenriched <- read.vcfR("~/crypto_data/13/filtered_snps_gatk_all_U.g.vcf"
```

Scanning file to determine attributes. File attributes: meta lines: 49 header_line: 50 variant count: 15430 column count: 100 Meta line 49 read in. All meta lines processed. gt matrix initialized. Character matrix gt created. Character matrix gt rows: 15430 Character matrix gt cols: 100 skip: 0 nrows: 15430 row_num: 0 Processed variant 1000Processed variant 2000Processed variant 3000Processed variant 4000Processed variant 5000Processed variant 6000Processed variant 7000Processed variant 8000Processed variant 9000Processed variant 10000Processed variant 11000Processed variant 12000Processed variant 13000Processed variant 14000Processed variant 15000Processed variant: 15430 All variants processed

```r
## Extract the FILTER column and subset the VCF for variants with 'PASS' in the
## FILTER column
filter_column <- vcf_unenriched@fix[, "FILTER"]
```

```r
# This filtered VCF contains 15,198 variants and 91.83% missing data
vcf_pass_unenriched <- vcf_unenriched[filter_column == "PASS", ]

# Extract the DP (Depth of Coverage) values from the INFO column
dp_info_unenriched <- extract.info(vcf_pass_unenriched, element = "DP")
dp_info_unenriched_numeric <- as.numeric(dp_info_unenriched)

# Extract the DP values from the FORMAT column for each individual
dp_format_unenriched <- extract.gt(vcf_pass_unenriched, element = "DP", as.numeric = TRUE)

coverage_unenriched_df <- as.data.frame(dp_format_unenriched)
coverage_unenriched_df <- reshape2::melt(coverage_unenriched_df, variable.name = "Individual",
    value.name = "Depth")
```

```
## No id variables; using all as measure variables
```

```r
# Remove NA values from the data
coverage_unenriched_df <- na.omit(coverage_unenriched_df)

# Create the boxplot using Plotly
boxplot_coverage_unenriched <- plot_ly(data = coverage_unenriched_df, x = ~Individual,
    y = ~Depth, type = "box", marker = list(color = "#8DA0CB")) %>%
    layout(xaxis = list(title = "Sample", tickangle = 45, tickfont = list(size = 10)),
        yaxis = list(title = "Depth of Coverage"), showlegend = FALSE)
```

**NOTE:** The results from these analyses are presented in **Fig. S52**.

```r
# Extract chromosome and position info
chrom_unenriched <- vcf_pass_unenriched@fix[, "CHROM"]

# Aggregate mean depth per chromosome
mean_depth_per_chrom_unenriched <- tapply(dp_info_unenriched_numeric, chrom_unenriched,
    mean, na.rm = TRUE)
coverage_chrom_unenriched <- data.frame(Chromosome = names(mean_depth_per_chrom_unenriched),
    MeanCoverage = mean_depth_per_chrom_unenriched)

# Barplot of mean depth per chromosome
plot_coverage_chrom_unenriched <- plot_ly(data = coverage_chrom_unenriched, x = ~Chromosome,
    y = ~MeanCoverage, type = "bar", marker = list(color = "#8DA0CB")) %>%
    layout(xaxis = list(title = "Chromosome", tickangle = 45, tickfont = list(size = 10)),
        yaxis = list(title = "Mean Depth"), bargap = 0.2)
```

**NOTE:** The results from these analyses are presented in **Fig. S53**.

```r
# Double enrichments Extract DP from INFO field (e.g., total depth per site
# across all samples)
dp_info_double <- extract.info(vcf_pass_double, element = "DP")
dp_info_double_numeric <- as.numeric(dp_info_double)
```

```
# Convert to data frame
dp_info_df <- data.frame(Depth = dp_info_double_numeric)

# Basic histogram
DP_double <- ggplot(dp_info_df, aes(x = Depth)) + geom_histogram(binwidth = 5, fill = "#66C2A5",
    color = "black") + theme_minimal() + labs(title = "Frequency Distribution of Site Depth (INFO field)
    x = "Total Site Depth (DP from INFO)", y = "Number of Variants") + xlim(0, quantile(dp_info_df$Depth
    0.99, na.rm = TRUE))

# Single enrichments Extract DP from INFO field (e.g., total depth per site
# across all samples)
dp_info_unenriched <- extract.info(vcf_pass_single, element = "DP")
dp_info_single_numeric <- as.numeric(dp_info_single)

# Convert to data frame
dp_info_df <- data.frame(Depth = dp_info_single_numeric)

# Basic histogram
DP_single <- ggplot(dp_info_df, aes(x = Depth)) + geom_histogram(binwidth = 5, fill = "#FC8D62",
    color = "black") + theme_minimal() + labs(title = "Frequency Distribution of Site Depth (INFO field)
    x = "Total Site Depth (DP from INFO)", y = "Number of Variants") + xlim(0, quantile(dp_info_df$Depth
    0.99, na.rm = TRUE))

# Unenriched samples Extract DP from INFO field (e.g., total depth per site
# across all samples)
dp_info_unenriched <- extract.info(vcf_pass_unenriched, element = "DP")
dp_info_unenriched_numeric <- as.numeric(dp_info_unenriched)

# Convert to data frame
dp_info_df <- data.frame(Depth = dp_info_unenriched_numeric)

# Basic histogram
DP_unenriched <- ggplot(dp_info_df, aes(x = Depth)) + geom_histogram(binwidth = 5,
    fill = "#8DA0CB", color = "black") + theme_minimal() + labs(title = "Frequency Distribution of Site
    x = "Total Site Depth (DP from INFO)", y = "Number of Variants") + xlim(0, quantile(dp_info_df$Depth
    0.99, na.rm = TRUE))

# Load patchwork
library(patchwork)
dp_all <- DP_double/DP_single/DP_unenriched + plot_layout(ncol = 1)  # 1 column = vertical layout
```

**13.4.1.4 Comparing overall depth (DP) in raw datasets among library types** **NOTE:** The results from these analyses are presented in **Fig. S54**.

**13.4.2 Filtering variants** Filtered VCF files contained, for double enrichments: 77 samples and 13,578 SNPs (24.1% missing data, Figs. S55–S56); for single enrichments: 76 samples and 12,498 SNPs (23.5% of missing data, Figs S57–S58); and for unenriched libraries missing data filtering would remove all variants and all samples, except for three, thus only filtering based on read depth and biallelic variants could be performed yielding a database with 91 samples, 11,226 SNPs (91% missing data; Figs. S59–S60).

```r
library(dartR)

# FILTER BASED ON MEAN DEPTH Filter variants with DP within reasonable range
# (>=5 to 200)
vcf_double_filtered <- vcf_pass_double[dp_info_double_numeric >= 5 & dp_info_double_numeric <=
    200, ]

# FILTER OUT VARIANTS WITH MORE THAN TWO ALLELES
is_biallelic <- function(vcf) {
    alt_alleles <- strsplit(vcf@fix[, "ALT"], ",")
    sapply(alt_alleles, length) == 1  # True if only 1 ALT allele
}
biallelic_variants <- is_biallelic(vcf_double_filtered)
vcf_biallelic_double <- vcf_double_filtered[biallelic_variants, ]
vcf_biallelic_double
```

###### 13.4.2.1 Filtering double-enriched VCF

```
## ***** Object of Class vcfR *****
## 91 samples
## 8 CHROMs
## 78,683 variants
## Object size: 78 Mb
## 48.2 percent missing data
## *****        *****         *****
```

```r
# FILTER LOCI BASED ON MISSING DATA
genotype_data_double <- extract.gt(vcf_biallelic_double, element = "GT", as.numeric = FALSE)
missing_data_proportion_double <- rowMeans(is.na(genotype_data_double))
# Filter out loci with more than 40% of missing
loci_to_keep_double <- which(missing_data_proportion_double <= 0.4)
vcf_filtered_loci_double <- vcf_biallelic_double[loci_to_keep_double, ]
vcf_filtered_loci_double
```

```
## ***** Object of Class vcfR *****
## 91 samples
## 8 CHROMs
## 13,578 variants
## Object size: 13.7 Mb
## 33.64 percent missing data
## *****        *****         *****
```

```r
# FILTER SAMPLES BASED ON MISSING DATA Extract the DP (Depth of Coverage)
# values from the INFO column
gt_matrix_double <- extract.gt(vcf_filtered_loci_double, element = "GT", as.numeric = FALSE)
missing_data_per_sample_double <- colMeans(is.na(gt_matrix_double))
# Filter out samples with more than 80% of missing
samples_to_keep_double <- which(missing_data_per_sample_double <= 0.8)
vcf_filtered_samples_double <- vcf_filtered_loci_double
vcf_filtered_samples_double@gt <- vcf_filtered_samples_double@gt[, c(1, samples_to_keep_double +
    1)]
vcf_filtered_samples_double
```

```
## ***** Object of Class vcfR *****
## 77 samples
## 8 CHROMs
## 13,578 variants
## Object size: 12.3 Mb
## 24.11 percent missing data
## *****        *****         *****
```

```r
# EXLPORING FILTERED VCF Extract the DP (Depth of Coverage) values from the
# INFO column
dp_info_double_target <- extract.info(vcf_filtered_samples_double, element = "DP")
dp_info_double_target_numeric <- as.numeric(dp_info_double_target)
# Extract the DP values from the FORMAT column for each individual
dp_format_double_target <- extract.gt(vcf_filtered_samples_double, element = "DP",
    as.numeric = TRUE)

coverage_double_df_target <- as.data.frame(dp_format_double_target)
coverage_double_df_target <- reshape2::melt(coverage_double_df_target, variable.name = "Individual",
    value.name = "Depth")
coverage_double_df_target <- na.omit(coverage_double_df_target)

# EXPLORE IF MONOMORPHIC LOCI ARE PRESENT AFTER FILTERING Transforming to
# genind
double_genind <- vcfR2genind(vcf_filtered_samples_double, ploidy = 1)
gl_double <- gi2gl(double_genind, parallel = FALSE, verbose = NULL)
```

```
## Starting gi2gl
## Starting gl.compliance.check
##   Processing genlight object with SNP data
##   The slot loc.all, which stores allele name for each locus, is empty.
## Creating a dummy variable (A/C) to insert in this slot.
##   Checking coding of SNPs
##     SNP data scored NA, 0, 1 or 2 confirmed
##   Checking locus metrics and flags
##   Recalculating locus metrics
##   Checking for monomorphic loci
##     No monomorphic loci detected
##   Checking for loci with all missing data
##     No loci with all missing data detected
##   Checking whether individual names are unique.
##   Checking for individual metrics
##   Warning: Creating a slot for individual metrics
##   Checking for population assignments
##   Population assignments not detected, individuals assigned
##                     to a single population labelled 'pop1'
##   Spelling of coordinates checked and changed if necessary to
##               lat/lon
## Completed: gl.compliance.check
## Completed: gi2gl
```

```r
# Create the boxplot using Plotly
boxplot_coverage_target_double <- plot_ly(data = coverage_double_df_target, x = ~Individual,
    y = ~Depth, type = "box", marker = list(color = "#66C2A5")) %>%
```

```
    layout(xaxis = list(title = "Sample", tickangle = 45, tickfont = list(size = 10)),
        yaxis = list(title = "Depth of Coverage"), showlegend = FALSE)
```

**NOTE:** The results from these analyses are presented in **Fig. S55**.

```
# Extract chromosome and position info
chrom_double <- vcf_filtered_samples_double@fix[, "CHROM"]
pos_double <- vcf_filtered_samples_double@fix[, "POS"]

# Aggregate mean depth per chromosome
mean_depth_per_chrom_double <- tapply(dp_info_double_target_numeric, chrom_double,
    mean, na.rm = TRUE)

coverage_chrom_double_fil <- data.frame(Chromosome = names(mean_depth_per_chrom_double),
    MeanCoverage = mean_depth_per_chrom_double)

# Barplot of mean depth per chromosome
plot_coverage_chrom_fil_double <- plot_ly(data = coverage_chrom_double_fil, x = ~Chromosome,
    y = ~MeanCoverage, type = "bar", marker = list(color = "#66C2A5")) %>%
    layout(xaxis = list(title = "Chromosome", tickangle = 45, tickfont = list(size = 10)),
        yaxis = list(title = "Mean Depth"), bargap = 0.2)
```

**NOTE:** The results from these analyses are presented in **Fig. S56**.

```
# FILTER BASED ON MEAN DEPTH Filter variants with DP within reasonable range
# (>=5 to 200)
vcf_single_filtered <- vcf_pass_single[dp_info_single_numeric >= 5 & dp_info_single_numeric <=
    200, ]

# FILTER OUT VARIANTS WITH MORE THAN TWO ALLELES
biallelic_variants <- is_biallelic(vcf_single_filtered)
vcf_biallelic_single <- vcf_single_filtered[biallelic_variants, ]
vcf_biallelic_single
```

###### 13.4.2.2 Filtering single-enriched VCF

```
## ***** Object of Class vcfR *****
## 91 samples
## 8 CHROMs
## 99,350 variants
## Object size: 97.4 Mb
## 44.77 percent missing data
## *****        *****         *****
```

```
# FILTER LOCI BASED ON MISSING DATA
genotype_data_single <- extract.gt(vcf_biallelic_single, element = "GT", as.numeric = FALSE)
missing_data_proportion_single <- rowMeans(is.na(genotype_data_single))
# Filter out loci with more than 40% of missing
loci_to_keep_single <- which(missing_data_proportion_single <= 0.4)
vcf_filtered_loci_single <- vcf_biallelic_single[loci_to_keep_single, ]
vcf_filtered_loci_single
```

```
## ***** Object of Class vcfR *****
## 91 samples
## 8 CHROMs
## 12,498 variants
## Object size: 12.5 Mb
## 33.96 percent missing data
## *****        *****         *****
```

```
# FILTER SAMPLES BASED ON MISSING DATA Extract the DP (Depth of Coverage)
# values from the INFO column
gt_matrix_single <- extract.gt(vcf_filtered_loci_single, element = "GT", as.numeric = FALSE)
missing_data_per_sample_single <- colMeans(is.na(gt_matrix_single))
# Filter out samples with more than 80% of missing
samples_to_keep_single <- which(missing_data_per_sample_single <= 0.8)
vcf_filtered_samples_single <- vcf_filtered_loci_single
vcf_filtered_samples_single@gt <- vcf_filtered_samples_single@gt[, c(1, samples_to_keep_single +
    1)]
vcf_filtered_samples_single
```

```
## ***** Object of Class vcfR *****
## 76 samples
## 8 CHROMs
## 12,498 variants
## Object size: 11.1 Mb
## 23.47 percent missing data
## *****        *****         *****
```

```
# EXLPORING FILTERED VCF Extract the DP (Depth of Coverage) values from the
# INFO column
dp_info_single_target <- extract.info(vcf_filtered_samples_single, element = "DP")
dp_info_single_target_numeric <- as.numeric(dp_info_single_target)
# Extract the DP values from the FORMAT column for each individual
dp_format_single_target <- extract.gt(vcf_filtered_samples_single, element = "DP",
    as.numeric = TRUE)

coverage_single_df_target <- as.data.frame(dp_format_single_target)
coverage_single_df_target <- reshape2::melt(coverage_single_df_target, variable.name = "Individual",
    value.name = "Depth")
```

```
## No id variables; using all as measure variables
```

```
coverage_single_df_target <- na.omit(coverage_single_df_target)

# EXPLORE IF MONOMORPHIC LOCI ARE PRESENT AFTER FILTERING Transforming to
# genind
single_genind <- vcfR2genind(vcf_filtered_samples_single, ploidy = 1)
gl_single <- gi2gl(single_genind, parallel = FALSE, verbose = NULL)
```

```
## Starting gi2gl
## Starting gl.compliance.check
##   Processing genlight object with SNP data
##   The slot loc.all, which stores allele name for each locus, is empty.
```

```
## Creating a dummy variable (A/C) to insert in this slot.
##   Checking coding of SNPs
##     SNP data scored NA, 0, 1 or 2 confirmed
##   Checking locus metrics and flags
##   Recalculating locus metrics
##   Checking for monomorphic loci
##     No monomorphic loci detected
##   Checking for loci with all missing data
##     No loci with all missing data detected
##   Checking whether individual names are unique.
##   Checking for individual metrics
##   Warning: Creating a slot for individual metrics
##   Checking for population assignments
##   Population assignments not detected, individuals assigned
##                     to a single population labelled 'pop1'
##   Spelling of coordinates checked and changed if necessary to
##             lat/lon
## Completed: gl.compliance.check
## Completed: gi2gl
```

```
# Create the boxplot using Plotly
boxplot_coverage_target_single <- plot_ly(data = coverage_single_df_target, x = ~Individual,
    y = ~Depth, type = "box", marker = list(color = "#FC8D62")) %>%
    layout(xaxis = list(title = "Sample", tickangle = 45, tickfont = list(size = 10)),
        yaxis = list(title = "Depth of Coverage"), showlegend = FALSE)
```

**NOTE:** The results from these analyses are presented in **Fig. S57**.

```
# Extract chromosome and position info
chrom_single <- vcf_filtered_samples_single@fix[, "CHROM"]
pos_single <- vcf_filtered_samples_single@fix[, "POS"]

# Aggregate mean depth per chromosome
mean_depth_per_chrom_single <- tapply(dp_info_single_target_numeric, chrom_single,
    mean, na.rm = TRUE)

coverage_chrom_single_fil <- data.frame(Chromosome = names(mean_depth_per_chrom_single),
    MeanCoverage = mean_depth_per_chrom_single)

# Barplot of mean depth per chromosome
plot_coverage_chrom_fil_single <- plot_ly(data = coverage_chrom_single_fil, x = ~Chromosome,
    y = ~MeanCoverage, type = "bar", marker = list(color = "#FC8D62")) %>%
    layout(xaxis = list(title = "Chromosome", tickangle = 45, tickfont = list(size = 10)),
        yaxis = list(title = "Mean Depth"), bargap = 0.2)
```

**NOTE:** The results from these analyses are presented in **Fig. S58**.

```

# FILTER BASED ON MEAN DEPTH Filter variants with DP within
# reasonable range (>=5 to 100)
vcf_unenriched_filtered <- vcf_pass_unenriched[dp_info_unenriched_numeric >=
```

```r
    5 & dp_info_unenriched_numeric <= 200, ]

# FILTER OUT VARIANTS WITH MORE THAN TWO ALLELES
biallelic_variants <- is_biallelic(vcf_unenriched_filtered)
vcf_biallelic_unenriched <- vcf_unenriched_filtered[biallelic_variants,
    ]
vcf_biallelic_unenriched
```

###### 13.4.2.3 Filtering unenriched VCF

```
## ***** Object of Class vcfR *****
## 91 samples
## 8 CHROMs
## 11,226 variants
## Object size: 11.1 Mb
## 91.01 percent missing data
## *****        *****         *****
```

```r
# FILTER LOCI BASED ON MISSING DATA
genotype_data_unenriched <- extract.gt(vcf_biallelic_unenriched,
    element = "GT", as.numeric = FALSE)
missing_data_proportion_unenriched <- rowMeans(is.na(genotype_data_unenriched))
# Filter out loci with more than 40% of missing, removes
# all loci!
loci_to_keep_unenriched <- which(missing_data_proportion_unenriched <=
    0.4)

# FILTER SAMPLES BASED ON MISSING DATA Extract the DP
# (Depth of Coverage) values from the INFO column
gt_matrix_unenriched <- extract.gt(vcf_biallelic_unenriched,
    element = "GT", as.numeric = FALSE)
missing_data_per_sample_unenriched <- colMeans(is.na(gt_matrix_unenriched))
# Filter out samples with more than 80% of missing, removes
# all except 3 samples!
samples_to_keep_unenriched <- which(missing_data_per_sample_unenriched <=
    0.9)

# EXLPORING FILTERED VCF Extract the DP (Depth of Coverage)
# values from the INFO column
dp_info_unenriched_target <- extract.info(vcf_biallelic_unenriched,
    element = "DP")
dp_info_unenriched_target_numeric <- as.numeric(dp_info_unenriched_target)
# Extract the DP values from the FORMAT column for each
# individual
dp_format_unenriched_target <- extract.gt(vcf_biallelic_unenriched,
    element = "DP", as.numeric = TRUE)

coverage_unenriched_df_target <- as.data.frame(dp_format_unenriched_target)
coverage_unenriched_df_target <- reshape2::melt(coverage_unenriched_df_target,
    variable.name = "Individual", value.name = "Depth")
```

```
## No id variables; using all as measure variables
```

```r
coverage_unenriched_df_target <- na.omit(coverage_unenriched_df_target)

# EXPLORE IF MONOMORPHIC LOCI ARE PRESENT AFTER FILTERING
# Transforming to genind
unenriched_genind <- vcfR2genind(vcf_biallelic_unenriched, ploidy = 1)
gl_unenriched <- gi2gl(unenriched_genind, parallel = FALSE, verbose = NULL)
```

```
## Starting gi2gl
## Starting gl.compliance.check
##   Processing genlight object with SNP data
##   The slot loc.all, which stores allele name for each locus, is empty.
## Creating a dummy variable (A/C) to insert in this slot.
##   Checking coding of SNPs
##     SNP data scored NA, 0, 1 or 2 confirmed
##   Checking locus metrics and flags
##   Recalculating locus metrics
##   Checking for monomorphic loci
##     No monomorphic loci detected
##   Checking for loci with all missing data
##     No loci with all missing data detected
##   Checking whether individual names are unique.
##   Checking for individual metrics
##   Warning: Creating a slot for individual metrics
##   Checking for population assignments
##   Population assignments not detected, individuals assigned
##                      to a single population labelled 'pop1'
##   Spelling of coordinates checked and changed if necessary to
##              lat/lon
## Completed: gl.compliance.check
## Completed: gi2gl
```

```r
# Create the boxplot using Plotly
boxplot_coverage_target_unenriched <- plot_ly(data = coverage_unenriched_df_target,
    x = ~Individual, y = ~Depth, type = "box", marker = list(color = "#8DA0CB")) %>%
    layout(xaxis = list(title = "Sample", tickangle = 45, tickfont = list(size = 10)),
        yaxis = list(title = "Depth of Coverage"), showlegend = FALSE)
```

**NOTE:** The results from these analyses are presented in **Fig. S59**.

```r
# Extract chromosome and position info
chrom_unenriched <- vcf_biallelic_unenriched@fix[, "CHROM"]
pos_unenriched <- vcf_biallelic_unenriched@fix[, "POS"]

# Aggregate mean depth per chromosome
mean_depth_per_chrom_unenriched <- tapply(dp_info_unenriched_target_numeric, chrom_unenriched,
    mean, na.rm = TRUE)

coverage_chrom_unenriched_fil <- data.frame(Chromosome = names(mean_depth_per_chrom_unenriched),
    MeanCoverage = mean_depth_per_chrom_unenriched)

# Barplot of mean depth per chromosome
plot_coverage_chrom_fil_unenriched <- plot_ly(data = coverage_chrom_unenriched_fil,
```

```
    x = ~Chromosome, y = ~MeanCoverage, type = "bar", marker = list(color = "#8DA0CB")) %>%
    layout(xaxis = list(title = "Chromosome", tickangle = 45, tickfont = list(size = 10)),
        yaxis = list(title = "Mean Depth"), bargap = 0.2)
```

**NOTE:** The results from these analyses are presented in **Fig. S60**.

##### 13.5 Genetic variation and distances

###### 13.5.1 Principal Components Analysis

```
library(SNPRelate)
```

###### 13.5.1.1 Double-enriched samples

```
## Loading required package: gdsfmt
```

```
# write.vcf(vcf_filtered_samples_double, file =
# 'vcf_filtered_samples_double.vcf')
double.fn <- "~/crypto_data/13/vcf_filtered_samples_double.vcf" snpgdsVCF2GDS(double.fn,
"double_enrichments.gds", method = "biallelic.only")
```

```
## Start file conversion from VCF to SNP GDS ...
## Method: extracting biallelic SNPs
## Number of samples: 77
## Parsing "~/crypto_data/13/vcf_filtered_samples_double.vcf" ... ##  import 13578 variants.
## + genotype   { Bit2 77x13578, 255.3K } *
## Optimize the access efficiency ...
## Clean up the fragments of GDS file:

##     open the file 'double_enrichments.gds' (313.9K)
##     # of fragments: 47
##     save to 'double_enrichments.gds.tmp'
##     rename 'double_enrichments.gds.tmp' (313.5K, reduced: 324B)
##     # of fragments: 20
```

```
snpgdsSummary("double_enrichments.gds")
```

```
## The file name: ~/crypto_miscellaneous/Simulations/v9/double_enrichments.gds ## The total number of sa
## The total number of SNPs: 13578
## SNP genotypes are stored in SNP-major mode (Sample X SNP).
```

```
genofile_double <- snpgdsOpen("double_enrichments.gds", readonly = FALSE)
head(read.gdsn(index.gdsn(genofile_double, "snp.id")))
```

```
## [1] 1 2 3 4 5 6
```

```r
head(read.gdsn(index.gdsn(genofile_double, "snp.rs.id")))
```

```
## [1] "" "" "" "" "" ""
```

```r
snpgdsSampMissRate(genofile_double)
```

```
##  [1] 0.2057740463 0.2294888791 0.7820739431 0.7902489321 0.1795551628
##  [6] 0.2856090735 0.4720871999 0.1779348947 0.2029017528 0.0015466195
## [11] 0.4684784210 0.6064958020 0.0000000000 0.1182059213 0.7129916041
## [16] 0.4708351745 0.1119457947 0.5409485933 0.4299602298 0.7365591398
## [21] 0.0038297246 0.0212107821 0.3212549713 0.2329503609 0.3147738990
## [26] 0.2764766534 0.4709824716 0.0008837826 0.0413168361 0.0482397997
## [31] 0.0343202239 0.1159964649 0.7250699661 0.0895566357 0.4561054647
## [36] 0.1816173221 0.3867285314 0.0506702018 0.5088378259 0.0000000000
## [41] 0.0475769627 0.4460156135 0.4815878627 0.3136691707 0.7347915746
## [46] 0.1935483871 0.2493003388 0.1425835911 0.1187951097 0.1507585801
## [51] 0.5180438945 0.3074826926 0.0012520253 0.2947414936 0.1334511710
## [56] 0.0004418913 0.5640742377 0.2304463102 0.0058918839 0.5204742967
## [61] 0.2403152158 0.7206510532 0.0505965532 0.0899248785 0.0171601119
## [66] 0.0312269848 0.0545735749 0.2288996907 0.1375018412 0.5362350862
## [71] 0.0849904257 0.0191486228 0.1073795846 0.4700250405 0.0001472971
## [76] 0.0346148181 0.0758580056
```

```r
# Add sample annotation that has been corrected
samp.annot <- data.frame(pop.group = c("UKH", "UKH", "UKH", "UKP", "UKP", "UKP",
    "UKP", "UKP", "UKP", "UKP", "UKP", "UKP", "UKP", "UKP", "UKP", "UKP", "UKP",
    "UKP", "UKP", "UKP", "UKP", "UKP", "UKP", "UKP", "UKP", "UKP", "UKP", "UKP",
    "UKP", "UKP", "UKP", "UKP", "UKP", "UKP", "UKP", "UKP", "UKP", "UKP", "UKP",
    "UKP", "UKP", "UKP", "UKP", "UKP", "UKP", "UKP", "UKP", "UKP", "UKP", "UKP",
    "UKP", "UKP", "UKP", "UKP", "UKP", "UKP", "UKP", "UKP", "UKP", "UKP", "UKP",
    "UKP", "UKP", "UKP", "UKP", "UKP", "UKP", "UKP", "UKP", "UKP", "UKP", "UKP",
    "UKP", "UKP", "UKP", "UKP", "UKP"))
add.gdsn(genofile_double, "sample.annot", samp.annot, replace = TRUE)
# Get sample id
sample.id <- read.gdsn(index.gdsn(genofile_double, "sample.id"))
# Get population information
pop_code_double <- read.gdsn(index.gdsn(genofile_double, "sample.annot/pop.group"))
pop_code_double <- as.factor(pop_code_double)
# color
myColHP <- c(rgb(222, 45, 38, maxColorValue = 255, alpha = 150), rgb(166, 216, 84,
    maxColorValue = 255, alpha = 150))
```

```r
# PCA
x.double <- tab(double_genind, freq = TRUE, NA.method = "mean")
pca_double_two <- dudi.pca(x.double, cent = TRUE, scale = FALSE, scannf = FALSE,
    nf = 4)
barplot(pca_double_two$eig[1:8], main = "PCA eigenvalues", col = heat.colors(8))

# Adjust the margins and aspect ratio if needed par(mar = c(3, 4, 4, 4) + 0.1)
# # Adjust margins as needed Set the axis limits to ensure the plot is properly
# scaled
xlim <- range(pca_double_two$li[, 1]) + c(-0.1, 0.1) * diff(range(pca_double_two$li[,
```

```
    1]))
ylim <- range(pca_double_two$li[, 2]) + c(-0.1, 0.1) * diff(range(pca_double_two$li[,
    2]))
# Plot
levels(pop_code_double)[levels(pop_code_double) == "UKP"] <- ""
plot(pca_double_two$li[, 1:2], type = "n", xlim = xlim, ylim = ylim, xlab = "PC1 56.43%",
    ylab = "PC2 11.52%")
s.class(pca_double_two$li[, 1:2], fac = (pop_code_double), col = transp(myColHP,
    0.6), axesel = FALSE, cstar = 0, cpoint = 5, add.plot = TRUE)
sample_name <- c("UKH148", "UKH149", "UKH150")
coords <- pca_double_two$li[rownames(pca_double_two$li) == sample_name, ]
text(coords[, 1], coords[, 2], labels = sample_name, cex = 2, pos = 3, col = "black")
abline(v = 0, h = 0, col = "grey", lty = 2)

# first four PCs variance
eig <- pca_double_two$eig
var_explained_two <- eig/sum(eig) * 100
lbls <- paste("PC", 1:4, "\n", format(var_explained_two[1:4], digits = 2), "%", sep = "")
pca_data <- pca_double_two$li[, 1:4]
pairs(pca_data, col = myColHP[pop_code_double], labels = lbls)
```

**NOTE:** The results from these analyses are presented in **Fig. 4A**.

```

# write.vcf(vcf_filtered_samples_single, file =
# 'vcf_filtered_samples_single.vcf')
single.fn <- "~/crypto_data/13/vcf_filtered_samples_single.vcf" snpgdsVCF2GDS(single.fn,
"single_enrichments.gds", method = "biallelic.only")
```

###### 13.5.1.2 Single-enriched samples

```
## Start file conversion from VCF to SNP GDS ...
## Method: extracting biallelic SNPs
## Number of samples: 76
## Parsing "~/crypto_data/13/vcf_filtered_samples_single.vcf" ... ##  import 12498 variants.
## + genotype   { Bit2 76x12498, 231.9K } *
## Optimize the access efficiency ...
## Clean up the fragments of GDS file:

##     open the file 'single_enrichments.gds' (287.0K)
##     # of fragments: 47
##     save to 'single_enrichments.gds.tmp'
##     rename 'single_enrichments.gds.tmp' (286.7K, reduced: 324B)
##     # of fragments: 20
```

```
snpgdsSummary("single_enrichments.gds")
```

```
## The file name:   ~/crypto_miscellaneous/Simulations/v9/single_enrichments.gds ## The total number of
## The total number of SNPs: 12498
## SNP genotypes are stored in SNP-major mode (Sample X SNP).
```

```r
genofile_single <- snpgdsOpen("single_enrichments.gds", readonly = FALSE)
head(read.gdsn(index.gdsn(genofile_single, "snp.id")))
```

```
## [1] 1 2 3 4 5 6
```

```r
head(read.gdsn(index.gdsn(genofile_single, "snp.rs.id")))
```

```
## [1] "" "" "" "" "" ""
```

```r
snpgdsSampMissRate(genofile_single)
```

```
##  [1] 0.2293967035 0.7799647944 0.7634821571 0.7660425668 0.1725076012
##  [6] 0.2738838214 0.4461513842 0.1517042727 0.1899503921 0.0014402304
## [11] 0.4162265963 0.5772923668 0.0000000000 0.0999359898 0.6933909426
## [16] 0.4694351096 0.1030564890 0.5336853897 0.4655144823 0.7239558329
## [21] 0.0030404865 0.0189630341 0.3356537046 0.2065930549 0.3474155865
## [26] 0.2204352696 0.4260681709 0.0011201792 0.0385661706 0.0506481037
## [31] 0.0237638022 0.1073771803 0.7390782525 0.0900144023 0.4647943671
## [36] 0.1857097136 0.3787806049 0.0564890382 0.4824771964 0.0000000000
## [41] 0.1058569371 0.4369499120 0.4817570811 0.3038886222 0.7547607617
## [46] 0.1585853737 0.2447591615 0.1320211234 0.1206593055 0.1274603937
## [51] 0.5142422788 0.3034085454 0.0020803329 0.2523603777 0.1579452712
## [56] 0.0007201152 0.5330452872 0.2090734518 0.0178428549 0.4629540727
## [61] 0.2217154745 0.6905104817 0.0552088334 0.0899343895 0.0160825732
## [66] 0.0312850056 0.0554488718 0.2043526964 0.1378620579 0.5118418947
## [71] 0.1037766043 0.1091374620 0.4452712434 0.0000800128 0.0217634822
## [76] 0.1044967195
```

```r
# Add sample annotation that has been corrected
samp.annot <- data.frame(pop.group = c("UKH", "UKH", "UKH", "UKP",
    "UKP", "UKP", "UKP", "UKP", "UKP", "UKP", "UKP", "UKP", "UKP",
    "UKP", "UKP", "UKP", "UKP", "UKP", "UKP", "UKP", "UKP", "UKP",
    "UKP", "UKP", "UKP", "UKP", "UKP", "UKP", "UKP", "UKP", "UKP",
    "UKP", "UKP", "UKP", "UKP", "UKP", "UKP", "UKP", "UKP", "UKP",
    "UKP", "UKP", "UKP", "UKP", "UKP", "UKP", "UKP", "UKP", "UKP",
    "UKP", "UKP", "UKP", "UKP", "UKP", "UKP", "UKP", "UKP", "UKP",
    "UKP", "UKP", "UKP", "UKP", "UKP", "UKP", "UKP", "UKP", "UKP",
    "UKP", "UKP", "UKP", "UKP", "UKP", "UKP", "UKP", "UKP", "UKP"))
add.gdsn(genofile_single, "sample.annot", samp.annot, replace = TRUE)
# Get sample id
sample.id <- read.gdsn(index.gdsn(genofile_single, "sample.id"))
# Get population information
pop_code_single <- read.gdsn(index.gdsn(genofile_single, "sample.annot/pop.group"))
pop_code_single <- as.factor(pop_code_single)
```

```r
# PCA
x.single <- tab(single_genind, freq = TRUE, NA.method = "mean")
pca_single_two <- dudi.pca(x.single, cent = TRUE, scale = FALSE, scannf = FALSE,
    nf = 4)
barplot(pca_single_two$eig[1:8], main = "PCA eigenvalues", col = heat.colors(8))
# Adjust the margins and aspect ratio if needed par(mar = c(5, 4, 4, 4) + 0.1)
```

```
# # Adjust margins as needed Set the axis limits to ensure the plot is properly
# scaled
xlim <- range(pca_single_two$li[, 1]) + c(-0.1, 0.1) * diff(range(pca_single_two$li[,
    1]))
ylim <- range(pca_single_two$li[, 2]) + c(-0.1, 0.1) * diff(range(pca_single_two$li[,
    2]))
# plot
levels(pop_code_single)[levels(pop_code_single) == "UKP"] <- ""
plot(pca_single_two$li[, 1:2], type = "n", xlim = xlim, ylim = ylim, xlab = "PC1 57.2%",
    ylab = "PC2 13.0%")
s.class(pca_single_two$li[, 1:2], fac = (pop_code_single), col = transp(myColHP,
    0.6), axesel = FALSE, cstar = 0, cpoint = 5, add.plot = TRUE)
sample_name <- c("UKH148", "UKH149", "UKH150")
coords <- pca_single_two$li[rownames(pca_single_two$li) == sample_name, ]
text(coords[, 1], coords[, 2], labels = sample_name, cex = 2, pos = 3, col = "black")
abline(v = 0, h = 0, col = "grey", lty = 2)

# first four PCs variance
eig_single <- pca_single_two$eig
var_single_two <- eig_single/sum(eig_single) * 100
lbls <- paste("PC", 1:4, "\n", format(var_single_two[1:4], digits = 2), "%", sep = "")
pca_data_single <- pca_single_two$li[, 1:4]
pairs(pca_data_single, col = myColHP[pop_code_single], labels = lbls)
```

**NOTE:** The results from these analyses are presented in **Fig. 4B**.

```
# write.vcf(vcf_biallelic_unenriched, file =
# 'vcf_biallelic_unenriched.vcf')
unenriched.fn <- "~/crypto_data/13/vcf_biallelic_unenriched.vcf" snpgdsVCF2GDS(unenriched.fn,
"unenriched_enrichments.gds", method = "biallelic.only")
```

###### 13.5.1.3 Unenriched samples

```
## Start file conversion from VCF to SNP GDS ...
## Method: extracting biallelic SNPs
## Number of samples: 91
## Parsing "~/crypto_data/13/vcf_biallelic_unenriched.vcf" ...##  import 11226 variants.
## + genotype   { Bit2 91x11226, 249.4K } *
## Optimize the access efficiency ...
## Clean up the fragments of GDS file:

##     open the file 'unenriched_enrichments.gds' (293.9K)
##     # of fragments: 47
##     save to 'unenriched_enrichments.gds.tmp'
##     rename 'unenriched_enrichments.gds.tmp' (293.5K, reduced: 324B)
##     # of fragments: 20
```

```
snpgdsSummary("unenriched_enrichments.gds")
```

```
## The file name: ~/crypto_miscellaneous/Simulations/v9/unenriched_enrichments.gds
```

```
## The total number of samples: 91
## The total number of SNPs: 11226
## SNP genotypes are stored in SNP-major mode (Sample X SNP).
```

```r
genofile_unenriched <- snpgdsOpen("unenriched_enrichments.gds",
    readonly = FALSE)
head(read.gdsn(index.gdsn(genofile_unenriched, "snp.id")))
```

```
## [1] 1 2 3 4 5 6
```

```r
head(read.gdsn(index.gdsn(genofile_unenriched, "snp.rs.id")))
```

```
## [1] "" "" "" "" "" ""
```

```r
snpgdsSampMissRate(genofile_unenriched)
```

```
##  [1] 0.9991092 0.9950116 0.9929628 0.9325673 0.9698022 0.9756815 0.9995546
##  [8] 0.9961696 0.9917157 0.9925174 0.9966150 0.9869945 0.9933191 0.9301621
## [15] 0.9763941 0.9977730 0.5664529 0.9757705 0.9975949 0.9965259 0.9832532
## [22] 0.9984857 0.9978621 0.9980403 0.9120791 0.9541244 0.9934972 0.9934972
## [29] 0.9922501 0.9913593 0.9937645 0.9437912 0.9840549 0.9156423 0.9586674
## [36] 0.9853020 0.9986638 0.9741671 0.9919829 0.9918047 0.9945662 0.9768395
## [43] 0.9967041 0.6857296 0.9837876 0.9992874 0.8543560 0.9810262 0.9965259
## [50] 0.9822733 0.9902013 0.9994655 0.9989311 0.9678425 0.9795118 0.9666845
## [57] 0.9995546 0.8803670 0.9429004 0.9615179 0.9778193 0.8904329 0.9887761
## [64] 0.8955104 0.8363620 0.9954570 0.9203634 0.7874577 0.9959024 0.9858365
## [71] 0.9967041 0.9983075 0.9337253 0.9993764 0.9862818 0.9603599 0.9600926
## [78] 0.9727418 0.9995546 0.9572421 0.9107429 0.9958133 0.9999109 0.9091395
## [85] 0.9859255 0.8743987 0.9994655 0.9976839 0.8881169 0.9750579 0.9819170
```

```r
# Add sample annotation that has been corrected
samp.annot <- data.frame(pop.group = c("UKH", "UKH", "UKH", "UKH",
    "UKH", "UKP", "UKP", "UKP", "UKP", "UKP", "UKP", "UKP", "UKP",
    "UKP", "UKP", "UKP", "UKP", "UKP", "UKP", "UKP", "UKP", "UKP",
    "UKP", "UKP", "UKP", "UKP", "UKP", "UKP", "UKP", "UKP", "UKP",
    "UKP", "UKP", "UKP", "UKP", "UKP", "UKP", "UKP", "UKP", "UKP",
    "UKP", "UKP", "UKP", "UKP", "UKP", "UKP", "UKP", "UKP", "UKP",
    "UKP", "UKP", "UKP", "UKP", "UKP", "UKP", "UKP", "UKP", "UKP",
    "UKP", "UKP", "UKP", "UKP", "UKP", "UKP", "UKP", "UKP", "UKP",
    "UKP", "UKP", "UKP", "UKP", "UKP", "UKP", "UKP", "UKP", "UKP",
    "UKP", "UKP", "UKP", "UKP", "UKP", "UKP", "UKP", "UKP", "UKP",
    "UKP", "UKP", "UKP", "UKP", "UKP", "UKP"))
add.gdsn(genofile_unenriched, "sample.annot", samp.annot, replace = TRUE)
# Run PCA
pca_unenriched_one <- snpgdsPCA(genofile_unenriched, autosome.only = FALSE,
    maf = 0.05, num.thread = 1)
```

```
## Principal Component Analysis (PCA) on genotypes:
## Excluding 1,272 SNPs (monomorphic: TRUE, MAF: 0.05, missing rate: NaN)
##     # of samples: 91
##     # of SNPs: 9,954
```

```
##     using 1 thread
##     # of principal components: 32
## PCA:    the sum of all selected genotypes (0,1,2) = 26534
## CPU capabilities:
## Thu Apr 23 19:55:56 2026    (internal increment: 5400)
## [..................................................]  0%, ETC: ---        [==========================
## Thu Apr 23 19:55:56 2026    Begin (eigenvalues and eigenvectors)
## Thu Apr 23 19:55:56 2026    Done.
```

```
# variance proportion (%)
pc.percent_one <- pca_unenriched_one$varprop * 100
head(round(pc.percent_one, 2))
```

```
## [1] 12.57  7.18  6.39  5.45  4.75  4.34
```

```
# Get sample id
sample.id <- read.gdsn(index.gdsn(genofile_unenriched, "sample.id"))
# Get population information
pop_code_unenriched <- read.gdsn(index.gdsn(genofile_unenriched,
    "sample.annot/pop.group"))
pop_code_unenriched <- as.factor(pop_code_unenriched)
```

```
# PCA
x.unenriched <- tab(unenriched_genind, freq = TRUE, NA.method = "mean")
pca_unenriched_two <- dudi.pca(x.unenriched, cent = TRUE, scale = FALSE, scannf = FALSE,
    nf = 4)
barplot(pca_unenriched_two$eig[1:8], main = "PCA eigenvalues", col = heat.colors(8))
# Adjust the margins and aspect ratio if needed par(mar = c(5, 4, 4, 4) + 0.1)
# # Adjust margins as needed Set the axis limits to ensure the plot is properly
# scaled
xlim <- range(pca_unenriched_two$li[, 1]) + c(-0.1, 0.1) * diff(range(pca_unenriched_two$li[,
    1]))
ylim <- range(pca_unenriched_two$li[, 2]) + c(-0.1, 0.1) * diff(range(pca_unenriched_two$li[,
    2]))
# plot
levels(pop_code_unenriched)[levels(pop_code_unenriched) == "UKP"] <- ""
plot(pca_unenriched_two$li[, 1:2], type = "n", xlim = xlim, ylim = ylim, xlab = "PC1 12.1%",
    ylab = "PC2 6.9%")
s.class(pca_unenriched_two$li[, 1:2], fac = (pop_code_unenriched), col = transp(myColHP,
    0.6), axesel = FALSE, cstar = 0, cpoint = 5, add.plot = TRUE)
sample_name <- c("UKP312", "UKP343")
coords <- pca_unenriched_two$li[, 1:2][rownames(pca_unenriched_two$li[, 1:2]) %in%
    sample_name, ]
text(coords[, 1], coords[, 2], labels = sample_name, cex = 2, pos = 3, col = "black")
abline(v = 0, h = 0, col = "grey", lty = 2)

# first four PCs variance
eig_unenriched <- pca_unenriched_two$eig
var_unenriched_two <- eig_unenriched/sum(eig_unenriched) * 100
lbls <- paste("PC", 1:4, "\n", format(var_unenriched_two[1:4], digits = 2), "%",
    sep = "")
pca_data_unenriched <- pca_unenriched_two$li[, 1:4]
pairs(pca_data_unenriched, col = myColHP[pop_code_unenriched], labels = lbls)
```

**NOTE:** The results from these analyses are presented in **Fig. 4C**.

##### 13.5.2 Exploring phylogenetic networks

```r
library(phangorn)
library(gridExtra)
# Before using vcf2phylip to convert to phylip, I had to: mv
# ./vcf_filtered_samples_double.vcf
# ./SplitsTree/vcf_filtered_samples_double.vcf.gz gunzip
# ./SplitsTree/vcf_filtered_samples_double.vcf.gz Then, using vcf2phylip 2.9
# ran: python vcf2phylip.py -i ../vcf_filtered_samples_double.vcf
# --output-folder ../double-phylip -b -n -f

# Read the PHYLIP file into R
phy_double <- read.phyDat("~/crypto_data/13/SplitsTree/double-phylip/vcf_filtered_samples_double.min4.phy"

dist_matrix <- dist.hamming(phy_double, exclude = "pairwise")
nnet_double <- neighborNet(dist_matrix)
# write.nexus.networx(file
# ='~/crypto_data/13/SplitsTree/double-phylip/double_network',
# nnet_double)
```

**13.5.2.1 Double-enriched samples** **NOTE:** The results from these analyses were visualized in SplitsTree 6.4.11 and presented in **Fig. 4A**.

```r
# Before using vcf2phylip to convert to phylip, I had to: mv
# ./Population_analyses/single_ref/vcf_filtered_samples_single.vcf
# ./SplitsTree/vcf_filtered_samples_single.vcf.gz gunzip
# ./SplitsTree/vcf_filtered_samples_single.vcf.gz Then, using vcf2phylip 2.9
# ran: python vcf2phylip.py -i ../vcf_filtered_samples_single.vcf
# --output-folder ../single-phylip -b -n -f

# Read the PHYLIP file into R
phy_single <- read.phyDat("~/crypto_data/13/SplitsTree/single-phylip/vcf_filtered_samples_single.min4.phy"

dist_matrix_single <- dist.hamming(phy_single, exclude = "pairwise")
nnet_single <- neighborNet(dist_matrix_single)
# write.nexus.networx(file
# ='~/crypto_data/13/SplitsTree/single-phylip/single_network',
# nnet_single)
```

**13.5.2.2 Single-enriched samples** **NOTE:** The results from these analyses were visualized in SplitsTree 6.4.11 and presented in **Fig. 4B**.

```
# Before using vcf2phylip to convert to phylip, I had to: mv
# ./Population_analyses/single_ref/vcf_biallelic_unenriched.vcf
# ./SplitsTree/vcf_biallelic_unenriched.vcf.gz gunzip
# ./SplitsTree/vcf_biallelic_unenriched.vcf.gz Then, using vcf2phylip 2.9 ran:
# python vcf2phylip.py -i ../vcf_biallelic_unenriched.vcf --output-folder
# ../unenriched-phylip -b -n -f

# Read the PHYLIP file into R
phy_unenriched <- read.phyDat("~/crypto_data/13/SplitsTree/unenriched-phylip/

vcf_biallelic_unenriched.min4.phy"

dist_matrix_unenriched <- dist.hamming(phy_unenriched, exclude = "pairwise")

# Because high presence of missing data, I need to handle these values:
dist_matrix_unenriched[is.na(dist_matrix_unenriched)] <- 0

# Calculate network
nnet_unenriched <- neighborNet(dist_matrix_unenriched)
# write.nexus.networx(file
# ='~/crypto_data/13/SplitsTree/unenriched-phylip/unenriched_network', # nnet_unenriched)
```

**13.5.2.3 Unenriched samples** **NOTE:** The results from these analyses were visualized in SplitsTree 6.4.11 and presented in **Fig. 4C**.

**13.5.3 Estimate and plot a Nei's genetic distances tree** We use the *aboot* function from *poppr* to calculate a UPGMA tree with the estimated Nei's genetic distances, and plot it.

```
library(poppr)
```

###### 13.5.3.1 Double-enriched samples

```
## This is poppr version 2.9.6. To get started, type package?poppr
## OMP parallel support: unavailable
```

```
library(ape)
double_tree <- aboot(double_genind, tree = "upgma", distance = nei.dist, sample = 1000,
    showtree = F, cutoff = 50, quiet = T)
```

```
# Adjust the margins to make more space for the legend
par(mar = c(5, 4, 4, 8) + 0.1)  # increase the right margin
# Plot the tree
plot.phylo(double_tree, cex = 0.4, adj = 0, tip.color = myColHP[pop_code_double],
    xpd = TRUE)
nodelabels(double_tree$node.label, adj = c(1.3, -1), frame = "n", cex = 0.5, font = 1,
    xpd = TRUE)
# Add the legend, allowing it to be outside the plot area
legend("topleft", legend = c("UKH", "UKP"), fill = myColHP, border = FALSE, bty = "n",
```

```
    cex = 0.7, xpd = TRUE, inset = c(-0.1, 0))
# Add the axis and title
axis(side = 1)
title(xlab = "Nei's genetic distance")
```

**NOTE:** The results from these analyses are presented in **Fig. 4A**.

```
single_tree <- aboot(single_genind, tree = "upgma", distance = nei.dist, sample = 1000,
    showtree = F, cutoff = 50, quiet = T)
```

```
# Adjust the margins to make more space for the legend
par(mar = c(5, 4, 4, 8) + 0.1)  # increase the right margin
# Plot the tree
plot.phylo(single_tree, cex = 0.4, adj = 0, tip.color = myColHP[pop_code_single],
    xpd = TRUE)
nodelabels(single_tree$node.label, adj = c(1.3, -1), frame = "n", cex = 0.5, font = 1,
    xpd = TRUE)
# Add the legend, allowing it to be outside the plot area
legend("topleft", legend = c("UKH", "UKP"), fill = myColHP, border = FALSE, bty = "n",
    cex = 0.7, xpd = TRUE, inset = c(-0.1, 0))
# Add the axis and title
axis(side = 1)
title(xlab = "Nei's genetic distance")
```

**13.5.3.2 Single-enriched samples** **NOTE:** The results from these analyses are presented in **Fig. 4B**.

```
unenriched_tree <- aboot(unenriched_genind, tree = "upgma", distance = nei.dist,
    sample = 1000, showtree = F, cutoff = 50, quiet = T)
```

```
# Adjust the margins to make more space for the legend
par(mar = c(5, 4, 4, 8) + 0.1)  # increase the right margin
# Plot the tree
plot.phylo(unenriched_tree, cex = 0.4, adj = 0, tip.color = myColHP[pop_code_unenriched],
    xpd = TRUE)
nodelabels(unenriched_tree$node.label, adj = c(1.3, -1), frame = "n", cex = 0.5,
    font = 1, xpd = TRUE)
# Add the legend, allowing it to be outside the plot area
legend("topleft", legend = c("UKH", "UKP"), fill = myColHP, border = FALSE, bty = "n",
    cex = 0.7, xpd = TRUE, inset = c(-0.1, 0))
# Add the axis and title
axis(side = 1)
title(xlab = "Nei's genetic distance")
```

**13.5.3.3 Unenriched samples** **NOTE:** The results from these analyses are presented in **Fig. 4C**.

**13.5.4 Exploring Clustering with Structure** To run Structure 2.3.4, we first converted each filtered VCF file into structure format using PGDSpider 3.0.0.0, setting the ploidy to 1, and left all other options as default. To run Structure, we used structure_threader 1.3.10. We ran for each VCF K = 1-5 with 3 replicates each. We used 10,000 chains for burn-in and 100,000 chains of MCMC. Each mainparams file was created with the appropriate number of loci, individuals, and ploidy of 1; all other options were left as default. The software also runs Structure Harvester v0.6.94 to assess the best value from Evanno's method. We ran Structure header:

> ml structure_threader/1.3.10-foss-2022a module load Structure/2.3.4-GCC-11.3.0
>
> structure_threader run
> -i librarytype.structure
> –params structure/params/params_library_type/mainparams
> -o structure/library_type_results
> -K 5 -R 3 -t4
> -st /apps/eb/Structure/2.3.4-GCC-11.3.0/bin/structure

```
library(readr)
# Read summary.txt for meanLnP(K)
#This file was manually edited to remove comments and include simpler column titles
summary_data_double <- read_tsv("~/crypto_data/13/Structure/double_results/bestK/summary_edited.txt"
```

###### 13.5.4.1 Double-enriched samples

```
## Rows: 15 Columns: 5
## -- Column specification --------------------------------------------------------
## Delimiter: "\t"
## chr (1): File_name
## dbl (4): K, LnP, MeanLnP, VarianceLnP
##
## i Use `spec()` to retrieve the full column specification for this data.
## i Specify the column types or set `show_col_types = FALSE` to quiet this message.
```

```
# Read evanno.txt for deltaK
evanno_data_double <- read_tsv("~/crypto_data/13/Structure/double_results/bestK/evanno.txt"
  comment = "#",
  col_names = c("K", "Reps", "Mean_LnP_K", "Stdev_LnP_K", "LnPrime", "AbsLnDoublePrime", "DeltaK"),
  col_types = "iddddd")

meanLnP_double <- plot_ly(
  data = summary_data_double,
  x = ~K,
  y = ~MeanLnP,
  type = "box",
  boxpoints = "all",
  jitter = 0.3,
  pointpos = -1.8,
  marker = list(color = 'rgba(7,40,89,0.5)'),
  line = list(color = 'rgba(7,40,89,1)')
```

```r
) %>%
  layout(
    xaxis = list(title = "K Value"),
    yaxis = list(title = "MeanLnP(K)")
  )

evanno_double <- plot_ly(
  data = evanno_data_double,
  x = ~K,
  y = ~DeltaK,
  type = 'scatter',
  mode = 'lines+markers',
  line = list(color = 'blue', width = 2),
  marker = list(color = 'red', size = 8)
) %>%
  layout(
    xaxis = list(title = "K Value"),
    yaxis = list(title = "DeltaK")
  )

combined_double <- subplot(
  meanLnP_double,      # First plot
  evanno_double,     # Second plot
  nrows = 1,        # Arrange plots in a single row
  shareY = FALSE,   # Do not share the Y-axis
  titleX = TRUE,    # Keep X-axis titles
  titleY = TRUE     # Keep Y-axis titles
)
```

**NOTE:** The results from these analyses are presented in **Fig. S61**.

```r
# Load data.
library("starmie")
library(stringr)

double_ktwo <- read.csv(file = "~/crypto_data/13/Structure/double_results/str_K2_rep2_f.csv"
    header = TRUE)

# Samples names
double_ktwo_names <- double_ktwo$Label
double_ktwo_names
```

```
##  [1] "UKH148"  "UKH149"  "UKH150"  "UKP301"  "UKP302"  "UKP303"  "UKP304"
##  [8] "UKP305"  "UKP306"  "UKP307"  "UKP308"  "UKP311"  "UKP312"  "UKP313"
## [15] "UKP314"  "UKP315"  "UKP316"  "UKP317"  "UKP318"  "UKP319"  "UKP320"
## [22] "UKP322"  "UKP323"  "UKP324"  "UKP325"  "UKP326"  "UKP327"  "UKP329"
## [29] "UKP331"  "UKP332"  "UKP333"  "UKP334"  "UKP335"  "UKP336"  "UKP337"
## [36] "UKP339"  "UKP340"  "UKP341"  "UKP342"  "UKP343"  "UKP347"  "UKP348"
## [43] "UKP349"  "UKP351"  "UKP356"  "UKP357"  "UKP358"  "UKP359"  "UKP361"
## [50] "UKP362"  "UKP363"  "UKP364"  "UKP366"  "UKP367"  "UKP368"  "UKP370"
## [57] "UKP371"  "UKP372"  "UKP374"  "UKP375"  "UKP376"  "UKP377"  "UKP380"
## [64] "UKP382"  "UKP383"  "UKP384"  "UKP385"  "UKP387"  "UKP388"  "UKP389"
## [71] "UKP391"  "UKP392"  "UKP393"  "UKP395A" "UKP398"  "UKP399"  "UKP400"
```

```r
# Membership probability to K=2.
double_ktwo_anc <- double_ktwo[, 5:6]
double_ktwo_anc
```

```
##    Inferred clusters
## 1     0.000    1.000
## 2     0.930    0.070
## 3     0.000    1.000
## 4     1.000    0.000
## 5     1.000    0.000
## 6     1.000    0.000
## 7     1.000    0.000
## 8     1.000    0.000
## 9     1.000    0.000
## 10    1.000    0.000
## 11    1.000    0.000
## 12    1.000    0.000
## 13    1.000    0.000
## 14    1.000    0.000
## 15    1.000    0.000
## 16    1.000    0.000
## 17    1.000    0.000
## 18    1.000    0.000
## 19    1.000    0.000
## 20    1.000    0.000
## 21    1.000    0.000
## 22    1.000    0.000
## 23    1.000    0.000
## 24    1.000    0.000
## 25    1.000    0.000
## 26    1.000    0.000
## 27    1.000    0.000
## 28    1.000    0.000
## 29    1.000    0.000
## 30    1.000    0.000
## 31    1.000    0.000
## 32    1.000    0.000
## 33    1.000    0.000
## 34    0.933    0.067
## 35    1.000    0.000
## 36    1.000    0.000
## 37    1.000    0.000
## 38    1.000    0.000
## 39    0.973    0.027
## 40    1.000    0.000
## 41    0.964    0.036
## 42    0.976    0.024
## 43    1.000    0.000
## 44    1.000    0.000
## 45    1.000    0.000
## 46    1.000    0.000
## 47    1.000    0.000
## 48    1.000    0.000
```

```
## 49    1.000    0.000
## 50    1.000    0.000
## 51    0.956    0.044
## 52    1.000    0.000
## 53    1.000    0.000
## 54    1.000    0.000
## 55    1.000    0.000
## 56    1.000    0.000
## 57    1.000    0.000
## 58    1.000    0.000
## 59    1.000    0.000
## 60    1.000    0.000
## 61    1.000    0.000
## 62    1.000    0.000
## 63    1.000    0.000
## 64    1.000    0.000
## 65    1.000    0.000
## 66    1.000    0.000
## 67    1.000    0.000
## 68    1.000    0.000
## 69    1.000    0.000
## 70    1.000    0.000
## 71    1.000    0.000
## 72    1.000    0.000
## 73    1.000    0.000
## 74    1.000    0.000
## 75    1.000    0.000
## 76    1.000    0.000
## 77    1.000    0.000
```

```
# Make spacing
double_ktwo_space <- rep(8/length(double_ktwo_names) * 0.1, length(double_ktwo_names))
double_ktwo_space
```

```
##  [1] 0.01038961 0.01038961 0.01038961 0.01038961 0.01038961 0.01038961
##  [7] 0.01038961 0.01038961 0.01038961 0.01038961 0.01038961 0.01038961
## [13] 0.01038961 0.01038961 0.01038961 0.01038961 0.01038961 0.01038961
## [19] 0.01038961 0.01038961 0.01038961 0.01038961 0.01038961 0.01038961
## [25] 0.01038961 0.01038961 0.01038961 0.01038961 0.01038961 0.01038961
## [31] 0.01038961 0.01038961 0.01038961 0.01038961 0.01038961 0.01038961
## [37] 0.01038961 0.01038961 0.01038961 0.01038961 0.01038961 0.01038961
## [43] 0.01038961 0.01038961 0.01038961 0.01038961 0.01038961 0.01038961
## [49] 0.01038961 0.01038961 0.01038961 0.01038961 0.01038961 0.01038961
## [55] 0.01038961 0.01038961 0.01038961 0.01038961 0.01038961 0.01038961
## [61] 0.01038961 0.01038961 0.01038961 0.01038961 0.01038961 0.01038961
## [67] 0.01038961 0.01038961 0.01038961 0.01038961 0.01038961 0.01038961
## [73] 0.01038961 0.01038961 0.01038961 0.01038961 0.01038961
```

```
# Make alternative spacings.
double_ktwo_space[3] <- 8/length(double_ktwo_names) * 0.5

# Create labels.
double_ktwo_labels <- c("UKH", "UKP")
```

```r
# and x coords for those labels
double_ktwo_yx <- c(30, 75, 120)

# Plot
structure_d <- barplot(t(as.matrix(double_ktwo_anc[, 1:2])),
    col = c("#A6D85496", "#DE2D2696"), ylab = "", border = NA,
    main = "", space = double_ktwo_space, names.arg = double_ktwo_names,
    cex.names = 0.5, las = 2)
```

**NOTE:** The results from these analyses are presented in **Fig. 4A**.

###### 13.5.4.1.1 *C. hominis* from double-enriched samples

```r
#This files was manually edited to remove comments and include simpler column titles
ch_summary_data_double <- read_tsv("~/crypto_data/13/Structure/C_hominis/C_hominis_double_results/bestK
summary_edited.txt"
```

```
## Rows: 15 Columns: 5
## -- Column specification --------------------------------------------------------
## Delimiter: "\t"
## chr (1): File_name
## dbl (4): K, LnP, MeanLnP, VarianceLnP
##
## i Use `spec()` to retrieve the full column specification for this data.
## i Specify the column types or set `show_col_types = FALSE` to quiet this message.
```

```r
# Read evanno.txt for deltaK
ch_evanno_data_double <- read_tsv("~/crypto_data/13/Structure/C_hominis/C_hominis_single_results/
bestK/evanno.txt"

comment = "#",
col_names = c("K", "Reps", "Mean_LnP_K", "Stdev_LnP_K", "LnPrime", "AbsLnDoublePrime", "DeltaK"),
col_types = "iddddd")
ch_meanLnP_double <- plot_ly(
  data = ch_summary_data_double,
  x = ~K,
  y = ~MeanLnP,
  type = "box",
  boxpoints = "all",
  jitter = 0.3,
  pointpos = -1.8,
  marker = list(color = 'rgba(7,40,89,0.5)'),
  line = list(color = 'rgba(7,40,89,1)')
) %>%
  layout(
    xaxis = list(title = "K Value"),
    yaxis = list(title = "MeanLnP(K)")
  )

ch_evanno_double <- plot_ly(
  data = ch_evanno_data_double,
  x = ~K,
  y = ~DeltaK,
  type = 'scatter',
```

```r
  mode = 'lines+markers',
  line = list(color = 'blue', width = 2),
  marker = list(color = 'red', size = 8)
) %>%
  layout(
    xaxis = list(title = "K Value"),
    yaxis = list(title = "DeltaK")
  )

ch_combined_double <- subplot(
  ch_meanLnP_double,      # First plot
  ch_evanno_double,     # Second plot
  nrows = 1,        # Arrange plots in a single row
  shareY = FALSE,   # Do not share the Y-axis
  titleX = TRUE,    # Keep X-axis titles
  titleY = TRUE     # Keep Y-axis titles
)
ch_combined_double
```

**NOTE:** The results from these analyses are presented in **Fig. S62**.

```r
# Load data.
double_ktwo_ch <- read.csv(file = "~/crypto_data/13/Structure/C_hominis/C_hominis_double_results/
str_K2_rep2_f.csv" header = TRUE)
```

```
## Warning in read.table(file = file, header = header, sep = sep, quote = quote, :
## incomplete final line found by readTableHeader on
## '~/crypto_data/13/Structure/C_hominis/C_hominis_double_results/str_K2_rep2_f.csv'
```

```r
# Samples names
double_ktwo_names_ch <- double_ktwo_ch$Label
double_ktwo_names_ch
```

```
## [1] "UKH148" "UKH149" "UKH150"
```

```r
# Membership probability to K=2.
double_ktwo_anc_ch <- double_ktwo_ch[, 5:6]
double_ktwo_anc_ch
```

```
##   Inferred clusters
## 1        0        1
## 2        1        0
## 3        0        1
```

```r
# Make spacing
double_ktwo_space_ch <- rep(0.2, length(double_ktwo_names_ch))
double_ktwo_space_ch
```

```
## [1] 0.2 0.2 0.2
```

```r
# and x coords for those labels
double_ktwo_yx_ch <- c(30, 75, 120)

# Plot
ch_structure_d <- barplot(t(as.matrix(double_ktwo_anc_ch[, 1:2])),
    col = c("#A6D85496", "#DE2D2696"), ylab = "", border = NA,
    main = "", space = double_ktwo_space_ch, names.arg = double_ktwo_names_ch,
    cex.names = 1.2, las = 2)
```

**NOTE:** The results from these analyses are presented in **Fig. S63**.

**13.5.4.1.2 *C. parvum* from double-enriched samples**

```r
#This files was manually edited to remove comments and include simpler column titles
cp_summary_data_double <- read_tsv("~/crypto_data/13/Structure/C_parvum/C_parvum_double_results/bestK/
summary_edited.txt"
```

```
## Rows: 15 Columns: 5
## -- Column specification --------------------------------------------------------
## Delimiter: "\t"
## chr (1): File_name
## dbl (4): K, LnP, MeanLnP, VarianceLnP
##
## i Use `spec()` to retrieve the full column specification for this data.
## i Specify the column types or set `show_col_types = FALSE` to quiet this message.
```

```r
# Read evanno.txt for deltaK
cp_evanno_data_double <- read_tsv("~/crypto_data/13/Structure/C_parvum/C_parvum_single_results/bestK/
evanno.txt"

comment = "#",
col_names = c("K", "Reps", "Mean_LnP_K", "Stdev_LnP_K", "LnPrime", "AbsLnDoublePrime", "DeltaK"),
col_types = "iddddd")

cp_meanLnP_double <- plot_ly(
  data = cp_summary_data_double,
  x = ~K,
  y = ~MeanLnP,
  type = "box",
  boxpoints = "all",
  jitter = 0.3,
  pointpos = -1.8,
  marker = list(color = 'rgba(7,40,89,0.5)'),
  line = list(color = 'rgba(7,40,89,1)')
) %>%
  layout(
    xaxis = list(title = "K Value"),
    yaxis = list(title = "MeanLnP(K)")
  )

cp_evanno_double <- plot_ly(
  data = cp_evanno_data_double,
  x = ~K,
  y = ~DeltaK,
  type = 'scatter',
```

```r
  mode = 'lines+markers',
  line = list(color = 'blue', width = 2),
  marker = list(color = 'red', size = 8)
) %>%
  layout(
    xaxis = list(title = "K Value"),
    yaxis = list(title = "DeltaK")
  )

cp_combined_double <- subplot(
  cp_meanLnP_double,     # First plot
  cp_evanno_double,     # Second plot
  nrows = 1,         # Arrange plots in a single row
  shareY = FALSE,   # Do not share the Y-axis
  titleX = TRUE,    # Keep X-axis titles
  titleY = TRUE     # Keep Y-axis titles
)
cp_combined_double
```

**NOTE:** The results from these analyses are presented in **Fig. S64**.

```r
# Load data.
double_ktwo_cp <- read.csv(file = "~/crypto_data/13/Structure/C_parvum/C_parvum_double_results/
str_K2_rep2_f.csv"
    header = TRUE)

# Samples names
double_ktwo_names_cp <- double_ktwo_cp$Label
double_ktwo_names_cp
```

```
##  [1] "UKP301"  "UKP302"  "UKP303"  "UKP304"  "UKP305"  "UKP306"  "UKP307"
##  [8] "UKP308"  "UKP311"  "UKP312"  "UKP313"  "UKP314"  "UKP315"  "UKP316"
## [15] "UKP317"  "UKP318"  "UKP319"  "UKP320"  "UKP322"  "UKP323"  "UKP324"
## [22] "UKP325"  "UKP326"  "UKP327"  "UKP329"  "UKP331"  "UKP332"  "UKP333"
## [29] "UKP334"  "UKP335"  "UKP336"  "UKP337"  "UKP339"  "UKP340"  "UKP341"
## [36] "UKP342"  "UKP343"  "UKP347"  "UKP348"  "UKP349"  "UKP351"  "UKP356"
## [43] "UKP357"  "UKP358"  "UKP359"  "UKP361"  "UKP362"  "UKP363"  "UKP364"
## [50] "UKP366"  "UKP367"  "UKP368"  "UKP370"  "UKP371"  "UKP372"  "UKP374"
## [57] "UKP375"  "UKP376"  "UKP377"  "UKP380"  "UKP382"  "UKP383"  "UKP384"
## [64] "UKP385"  "UKP387"  "UKP388"  "UKP389"  "UKP391"  "UKP392"  "UKP393"
## [71] "UKP395A" "UKP398"  "UKP399"  "UKP400"
```

```r
# Membership probability to K=2.
double_ktwo_anc_cp <- double_ktwo_cp[, 5:6]
double_ktwo_anc_cp
```

```
##    Inferred clusters
## 1     1.000    0.000
## 2     1.000    0.000
## 3     1.000    0.000
## 4     1.000    0.000
## 5     1.000    0.000
## 6     1.000    0.000
```

```
## 7     1.000   0.000
## 8     1.000   0.000
## 9     1.000   0.000
## 10    1.000   0.000
## 11    1.000   0.000
## 12    1.000   0.000
## 13    1.000   0.000
## 14    1.000   0.000
## 15    1.000   0.000
## 16    1.000   0.000
## 17    1.000   0.000
## 18    1.000   0.000
## 19    1.000   0.000
## 20    1.000   0.000
## 21    1.000   0.000
## 22    1.000   0.000
## 23    1.000   0.000
## 24    1.000   0.000
## 25    1.000   0.000
## 26    1.000   0.000
## 27    1.000   0.000
## 28    1.000   0.000
## 29    1.000   0.000
## 30    1.000   0.000
## 31    0.116   0.884
## 32    1.000   0.000
## 33    1.000   0.000
## 34    1.000   0.000
## 35    1.000   0.000
## 36    0.244   0.756
## 37    1.000   0.000
## 38    0.708   0.292
## 39    0.285   0.715
## 40    1.000   0.000
## 41    1.000   0.000
## 42    1.000   0.000
## 43    1.000   0.000
## 44    1.000   0.000
## 45    1.000   0.000
## 46    1.000   0.000
## 47    1.000   0.000
## 48    0.109   0.891
## 49    1.000   0.000
## 50    1.000   0.000
## 51    1.000   0.000
## 52    0.991   0.009
## 53    1.000   0.000
## 54    1.000   0.000
## 55    1.000   0.000
## 56    1.000   0.000
## 57    1.000   0.000
## 58    1.000   0.000
## 59    1.000   0.000
## 60    1.000   0.000
```

```
## 61    1.000    0.000
## 62    0.998    0.002
## 63    1.000    0.000
## 64    1.000    0.000
## 65    1.000    0.000
## 66    1.000    0.000
## 67    1.000    0.000
## 68    1.000    0.000
## 69    1.000    0.000
## 70    1.000    0.000
## 71    1.000    0.000
## 72    1.000    0.000
## 73    1.000    0.000
## 74    1.000    0.000
```

```
# Make spacing
double_ktwo_space_cp <- rep(0.2, length(double_ktwo_names_cp))
double_ktwo_space_cp
```

```
##  [1] 0.2 0.2 0.2 0.2 0.2 0.2 0.2 0.2 0.2 0.2 0.2 0.2 0.2 0.2 0.2 0.2 0.2 0.2 0.2
## [20] 0.2 0.2 0.2 0.2 0.2 0.2 0.2 0.2 0.2 0.2 0.2 0.2 0.2 0.2 0.2 0.2 0.2 0.2 0.2
## [39] 0.2 0.2 0.2 0.2 0.2 0.2 0.2 0.2 0.2 0.2 0.2 0.2 0.2 0.2 0.2 0.2 0.2 0.2 0.2
## [58] 0.2 0.2 0.2 0.2 0.2 0.2 0.2 0.2 0.2 0.2 0.2 0.2 0.2 0.2 0.2 0.2 0.2
```

```
# and x coords for those labels
double_ktwo_yx_cp <- c(30, 75, 120)

# Plot
cp_structure_d <- barplot(t(as.matrix(double_ktwo_anc_cp[, 1:2])),
    col = c("#A6D85496", "#00441b"), ylab = "", border = NA,
    main = "", space = double_ktwo_space_cp, names.arg = double_ktwo_names_cp,
    cex.names = 0.5, las = 2)
```

**NOTE:** The results from these analyses are presented in **Fig. S65**.

```
# Read summary.txt for meanLnP(K)
#These file was manually edited to remove comments and include simpler column titles
summary_data_single <- read_tsv("~/crypto_data/13/Structure/single_results/bestK/summary_edited.txt"
```

###### 13.5.4.2 Single-enriched samples

```
## Rows: 15 Columns: 5
## -- Column specification --------------------------------------------------------
## Delimiter: "\t"
## chr (1): File_name
## dbl (4): K, LnP, MeanLnP, VarianceLnP
##
## i Use `spec()` to retrieve the full column specification for this data.
## i Specify the column types or set `show_col_types = FALSE` to quiet this message.
```

```r
# Read evanno.txt for deltaK
evanno_data_single <- read_tsv("~/crypto_data/13/Structure/single_results/bestK/evanno.txt"
  comment = "#",
  col_names = c("K", "Reps", "Mean_LnP_K", "Stdev_LnP_K", "LnPrime", "AbsLnDoublePrime", "DeltaK"),
  col_types = "iddddd")

meanLnP_single <- plot_ly(
  data = summary_data_single,
  x = ~K,
  y = ~MeanLnP,
  type = "box",
  boxpoints = "all",
  jitter = 0.3,
  pointpos = -1.8,
  marker = list(color = 'rgba(7,40,89,0.5)'),
  line = list(color = 'rgba(7,40,89,1)')
) %>%
  layout(
    xaxis = list(title = "K Value"),
    yaxis = list(title = "MeanLnP(K)")
  )

evanno_single <- plot_ly(
  data = evanno_data_single,
  x = ~K,
  y = ~DeltaK,
  type = 'scatter',
  mode = 'lines+markers',
  line = list(color = 'blue', width = 2),
  marker = list(color = 'red', size = 8)
) %>%
  layout(
    xaxis = list(title = "K Value"),
    yaxis = list(title = "DeltaK")
  )

combined_single <- subplot(
  meanLnP_single,     # First plot
  evanno_single,     # Second plot
  nrows = 1,        # Arrange plots in a single row
  shareY = FALSE,   # Do not share the Y-axis
  titleX = TRUE,    # Keep X-axis titles
  titleY = TRUE     # Keep Y-axis titles
)
```

**NOTE:** The results from these analyses are presented in **Fig. S66**.

```r
# Load data.
single_ktwo <- read.csv(file = "~/crypto_data/13/Structure/single_results/str_K2_rep2_f.csv"
    header = TRUE)

# Samples names
```

```r
single_ktwo_names <- single_ktwo$Label
single_ktwo_names
```

```
##  [1] "UKH148"  "UKH149"  "UKH150"  "UKP301"  "UKP302"  "UKP303"  "UKP304"
##  [8] "UKP305"  "UKP306"  "UKP307"  "UKP308"  "UKP311"  "UKP312"  "UKP313"
## [15] "UKP314"  "UKP315"  "UKP316"  "UKP317"  "UKP318"  "UKP319"  "UKP320"
## [22] "UKP322"  "UKP323"  "UKP324"  "UKP325"  "UKP326"  "UKP327"  "UKP329"
## [29] "UKP331"  "UKP332"  "UKP333"  "UKP334"  "UKP335"  "UKP336"  "UKP337"
## [36] "UKP339"  "UKP340"  "UKP341"  "UKP342"  "UKP343"  "UKP347"  "UKP348"
## [43] "UKP349"  "UKP351"  "UKP356"  "UKP357"  "UKP358"  "UKP359"  "UKP361"
## [50] "UKP362"  "UKP363"  "UKP364"  "UKP366"  "UKP367"  "UKP368"  "UKP370"
## [57] "UKP371"  "UKP372"  "UKP374"  "UKP375"  "UKP376"  "UKP377"  "UKP380"
## [64] "UKP382"  "UKP383"  "UKP384"  "UKP385"  "UKP387"  "UKP388"  "UKP389"
## [71] "UKP391"  "UKP393"  "UKP395A" "UKP398"  "UKP399"  "UKP400"
```

```r
# Membership probability to K=2.
single_ktwo_anc <- single_ktwo[, 5:6]
single_ktwo_anc
```

```
##    Inferred clusters
## 1     0.000    1.000
## 2     0.337    0.663
## 3     0.000    1.000
## 4     1.000    0.000
## 5     1.000    0.000
## 6     1.000    0.000
## 7     1.000    0.000
## 8     1.000    0.000
## 9     1.000    0.000
## 10    1.000    0.000
## 11    1.000    0.000
## 12    1.000    0.000
## 13    1.000    0.000
## 14    1.000    0.000
## 15    1.000    0.000
## 16    1.000    0.000
## 17    1.000    0.000
## 18    1.000    0.000
## 19    1.000    0.000
## 20    1.000    0.000
## 21    1.000    0.000
## 22    1.000    0.000
## 23    1.000    0.000
## 24    1.000    0.000
## 25    1.000    0.000
## 26    1.000    0.000
## 27    1.000    0.000
## 28    1.000    0.000
## 29    1.000    0.000
## 30    1.000    0.000
## 31    1.000    0.000
## 32    1.000    0.000
```

```
## 33    1.000    0.000
## 34    0.959    0.041
## 35    1.000    0.000
## 36    1.000    0.000
## 37    1.000    0.000
## 38    1.000    0.000
## 39    1.000    0.000
## 40    1.000    0.000
## 41    1.000    0.000
## 42    0.999    0.001
## 43    1.000    0.000
## 44    1.000    0.000
## 45    1.000    0.000
## 46    1.000    0.000
## 47    1.000    0.000
## 48    1.000    0.000
## 49    1.000    0.000
## 50    1.000    0.000
## 51    0.993    0.007
## 52    1.000    0.000
## 53    1.000    0.000
## 54    1.000    0.000
## 55    1.000    0.000
## 56    1.000    0.000
## 57    1.000    0.000
## 58    1.000    0.000
## 59    1.000    0.000
## 60    1.000    0.000
## 61    1.000    0.000
## 62    1.000    0.000
## 63    1.000    0.000
## 64    1.000    0.000
## 65    1.000    0.000
## 66    1.000    0.000
## 67    1.000    0.000
## 68    1.000    0.000
## 69    1.000    0.000
## 70    1.000    0.000
## 71    1.000    0.000
## 72    1.000    0.000
## 73    1.000    0.000
## 74    1.000    0.000
## 75    1.000    0.000
## 76    1.000    0.000
```

```r
# Make spacing
single_ktwo_space <- rep(8/length(single_ktwo_names) * 0.1, length(single_ktwo_names))
single_ktwo_space
```

```
##  [1] 0.01052632 0.01052632 0.01052632 0.01052632 0.01052632 0.01052632
##  [7] 0.01052632 0.01052632 0.01052632 0.01052632 0.01052632 0.01052632
## [13] 0.01052632 0.01052632 0.01052632 0.01052632 0.01052632 0.01052632
## [19] 0.01052632 0.01052632 0.01052632 0.01052632 0.01052632 0.01052632
## [25] 0.01052632 0.01052632 0.01052632 0.01052632 0.01052632 0.01052632
```

```
## [31] 0.01052632 0.01052632 0.01052632 0.01052632 0.01052632 0.01052632
## [37] 0.01052632 0.01052632 0.01052632 0.01052632 0.01052632 0.01052632
## [43] 0.01052632 0.01052632 0.01052632 0.01052632 0.01052632 0.01052632
## [49] 0.01052632 0.01052632 0.01052632 0.01052632 0.01052632 0.01052632
## [55] 0.01052632 0.01052632 0.01052632 0.01052632 0.01052632 0.01052632
## [61] 0.01052632 0.01052632 0.01052632 0.01052632 0.01052632 0.01052632
## [67] 0.01052632 0.01052632 0.01052632 0.01052632 0.01052632 0.01052632
## [73] 0.01052632 0.01052632 0.01052632 0.01052632
```

```
# Make alternative spacings.
single_ktwo_space[3] <- 8/length(single_ktwo_names) * 0.5

# Create labels.
single_ktwo_labels <- c("UKH", "UKP")

# and x coords for those labels
single_ktwo_yx <- c(30, 75, 120)

# Plot
structure_s <- barplot(t(as.matrix(single_ktwo_anc[, 1:2])),
    col = c("#A6D85496", "#DE2D2696"), ylab = "", border = NA,
    main = "", space = single_ktwo_space, names.arg = single_ktwo_names,
    cex.names = 0.5, las = 2)
```

**NOTE:** The results from these analyses are presented in **Fig. 4B**.

###### 13.5.4.2.1 *C. hominis* from single-enriched samples

```
#This files was manually edited to remove comments and include simpler column titles
ch_summary_data_single <- read_tsv("~/crypto_data/13/Structure/C_hominis/C_hominis_single_results/bestK
summary_edited.txt"
```

```
## Rows: 15 Columns: 5
## -- Column specification --------------------------------------------------------
## Delimiter: "\t"
## chr (1): File_name
## dbl (4): K, LnP, MeanLnP, VarianceLnP
##
## i Use `spec()` to retrieve the full column specification for this data.
## i Specify the column types or set `show_col_types = FALSE` to quiet this message.
```

```
# Read evanno.txt for deltaK
ch_evanno_data_single <- read_tsv("~/crypto_data/13/Structure/C_hominis/C_hominis_single_results/bestK/
evanno.txt"

comment = "#",
col_names = c("K", "Reps", "Mean_LnP_K", "Stdev_LnP_K", "LnPrime", "AbsLnDoublePrime", "DeltaK"),
col_types = "iddddd")

ch_meanLnP_single <- plot_ly(
  data = ch_summary_data_single,
  x = ~K,
  y = ~MeanLnP,
  type = "box",
  boxpoints = "all",
```

```
  jitter = 0.3,
  pointpos = -1.8,
  marker = list(color = 'rgba(7,40,89,0.5)'),
  line = list(color = 'rgba(7,40,89,1)')
) %>%
  layout(
    xaxis = list(title = "K Value"),
    yaxis = list(title = "MeanLnP(K)")
  )

ch_evanno_single <- plot_ly(
  data = ch_evanno_data_single,
  x = ~K,
  y = ~DeltaK,
  type = 'scatter',
  mode = 'lines+markers',
  line = list(color = 'blue', width = 2),
  marker = list(color = 'red', size = 8)
) %>%
  layout(
    xaxis = list(title = "K Value"),
    yaxis = list(title = "DeltaK")
  )

ch_combined_single <- subplot(
  ch_meanLnP_single,     # First plot
  ch_evanno_single,     # Second plot
  nrows = 1,        # Arrange plots in a single row
  shareY = FALSE,   # Do not share the Y-axis
  titleX = TRUE,    # Keep X-axis titles
  titleY = TRUE     # Keep Y-axis titles
)
ch_combined_single
```

**NOTE:** The results from these analyses are presented in **Fig. S67**.

```
# Load data.
single_ktwo_ch <- read.csv(file = "~/crypto_data/13/Structure/C_hominis/C_hominis_single_results/
str_K2_rep2_f.csv"  header = TRUE)
```

```
## Warning in read.table(file = file, header = header, sep = sep, quote = quote, :
## incomplete final line found by readTableHeader on
## '~/crypto_data/13/Structure/C_hominis/C_hominis_single_results/str_K2_rep2_f.csv'
```

```
# Samples names
single_ktwo_names_ch <- single_ktwo_ch$Label
single_ktwo_names_ch
```

```
## [1] "UKH148" "UKH149" "UKH150"
```

```r
# Membership probability to K=2.
single_ktwo_anc_ch <- single_ktwo_ch[, 5:6]
single_ktwo_anc_ch
```

```
##   Inferred clusters
## 1     0.000    1.000
## 2     1.000    0.000
## 3     0.001    0.999
```

```r
# Make spacing
single_ktwo_space_ch <- rep(0.2, length(single_ktwo_names_ch))
single_ktwo_space_ch
```

```
## [1] 0.2 0.2 0.2
```

```r
# and x coords for those labels
single_ktwo_yx_ch <- c(30, 75, 120)

# Plot
ch_structure_s <- barplot(t(as.matrix(single_ktwo_anc_ch[, 1:2])),
    col = c("#A6D85496", "#DE2D2696"), ylab = "", border = NA,
    main = "", space = single_ktwo_space_ch, names.arg = single_ktwo_names_ch,
    cex.names = 1.2, las = 2)
```

**NOTE:** The results from these analyses are presented in **Fig. S68**.

###### 13.5.4.2.2 *C. parvum* from single-enriched samples

```r
#This files was manually edited to remove comments and include simpler column titles
cp_summary_data_single <- read_tsv("~/crypto_data/13/Structure/C_parvum/C_parvum_single_results/bestK/
summary_edited.txt"
```

```
## Rows: 15 Columns: 5
## -- Column specification --------------------------------------------------------
## Delimiter: "\t"
## chr (1): File_name
## dbl (4): K, LnP, MeanLnP, VarianceLnP
##
## i Use `spec()` to retrieve the full column specification for this data.
## i Specify the column types or set `show_col_types = FALSE` to quiet this message.
```

```r
# Read evanno.txt for deltaK
cp_evanno_data_single <- read_tsv("~/crypto_data/13/Structure/C_parvum/C_parvum_single_results/bestK/
evanno.txt"
comment = "#",
col_names = c("K", "Reps", "Mean_LnP_K", "Stdev_LnP_K", "LnPrime", "AbsLnsinglePrime", "DeltaK"),
col_types = "iddddd")

cp_meanLnP_single <- plot_ly(
  data = cp_summary_data_single,
  x = ~K,
  y = ~MeanLnP,
```

```
  type = "box",
  boxpoints = "all",
  jitter = 0.3,
  pointpos = -1.8,
  marker = list(color = 'rgba(7,40,89,0.5)'),
  line = list(color = 'rgba(7,40,89,1)')
) %>%
  layout(
    xaxis = list(title = "K Value"),
    yaxis = list(title = "MeanLnP(K)")
  )

cp_evanno_single <- plot_ly(
  data = cp_evanno_data_single,
  x = ~K,
  y = ~DeltaK,
  type = 'scatter',
  mode = 'lines+markers',
  line = list(color = 'blue', width = 2),
  marker = list(color = 'red', size = 8)
) %>%
  layout(
    xaxis = list(title = "K Value"),
    yaxis = list(title = "DeltaK")
  )

cp_combined_single <- subplot(
  cp_meanLnP_single,      # First plot
  cp_evanno_single,     # Second plot
  nrows = 1,        # Arrange plots in a single row
  shareY = FALSE,   # Do not share the Y-axis
  titleX = TRUE,    # Keep X-axis titles
  titleY = TRUE     # Keep Y-axis titles
)
cp_combined_single
```

**NOTE:** The results from these analyses are presented in **Fig. S69**.

```
# Load data.
single_ktwo_cp <- read.csv(file = "~/crypto_data/13/Structure/C_parvum/C_parvum_single_results/
str_K2_rep2_f.csv"
    header = TRUE)

# Samples names
single_ktwo_names_cp <- single_ktwo_cp$Label
single_ktwo_names_cp
```

```
##  [1] "UKP301"  "UKP302"  "UKP303"  "UKP304"  "UKP305"  "UKP306"  "UKP307"
##  [8] "UKP308"  "UKP311"  "UKP312"  "UKP313"  "UKP314"  "UKP315"  "UKP316"
## [15] "UKP317"  "UKP318"  "UKP319"  "UKP320"  "UKP322"  "UKP323"  "UKP324"
## [22] "UKP325"  "UKP326"  "UKP327"  "UKP329"  "UKP331"  "UKP332"  "UKP333"
## [29] "UKP334"  "UKP335"  "UKP336"  "UKP337"  "UKP339"  "UKP340"  "UKP341"
## [36] "UKP342"  "UKP343"  "UKP347"  "UKP348"  "UKP349"  "UKP351"  "UKP356"
## [43] "UKP357"  "UKP358"  "UKP359"  "UKP361"  "UKP362"  "UKP363"  "UKP364"
```

```
## [50] "UKP366"  "UKP367"  "UKP368"  "UKP370"  "UKP371"  "UKP372"  "UKP374"
## [57] "UKP375"  "UKP376"  "UKP377"  "UKP380"  "UKP382"  "UKP383"  "UKP384"
## [64] "UKP385"  "UKP387"  "UKP388"  "UKP389"  "UKP391"  "UKP393"  "UKP395A"
## [71] "UKP398"  "UKP399"  "UKP400"  "UKP400"
```

```
# Membership probability to K=2.
single_ktwo_anc_cp <- single_ktwo_cp[, 5:6]
single_ktwo_anc_cp
```

```
##    Inferred clusters
## 1     1.000    0.000
## 2     1.000    0.000
## 3     1.000    0.000
## 4     1.000    0.000
## 5     1.000    0.000
## 6     1.000    0.000
## 7     1.000    0.000
## 8     1.000    0.000
## 9     1.000    0.000
## 10    1.000    0.000
## 11    1.000    0.000
## 12    1.000    0.000
## 13    1.000    0.000
## 14    1.000    0.000
## 15    1.000    0.000
## 16    1.000    0.000
## 17    1.000    0.000
## 18    1.000    0.000
## 19    1.000    0.000
## 20    1.000    0.000
## 21    1.000    0.000
## 22    1.000    0.000
## 23    1.000    0.000
## 24    1.000    0.000
## 25    1.000    0.000
## 26    1.000    0.000
## 27    1.000    0.000
## 28    1.000    0.000
## 29    1.000    0.000
## 30    1.000    0.000
## 31    0.000    1.000
## 32    1.000    0.000
## 33    1.000    0.000
## 34    1.000    0.000
## 35    1.000    0.000
## 36    0.002    0.998
## 37    1.000    0.000
## 38    1.000    0.000
## 39    0.000    1.000
## 40    1.000    0.000
## 41    1.000    0.000
## 42    1.000    0.000
## 43    1.000    0.000
## 44    1.000    0.000
```

```
## 45    1.000    0.000
## 46    1.000    0.000
## 47    1.000    0.000
## 48    0.000    1.000
## 49    1.000    0.000
## 50    1.000    0.000
## 51    0.995    0.005
## 52    1.000    0.000
## 53    1.000    0.000
## 54    1.000    0.000
## 55    1.000    0.000
## 56    1.000    0.000
## 57    1.000    0.000
## 58    1.000    0.000
## 59    1.000    0.000
## 60    1.000    0.000
## 61    1.000    0.000
## 62    0.967    0.033
## 63    1.000    0.000
## 64    1.000    0.000
## 65    1.000    0.000
## 66    1.000    0.000
## 67    1.000    0.000
## 68    1.000    0.000
## 69    1.000    0.000
## 70    1.000    0.000
## 71    1.000    0.000
## 72    1.000    0.000
## 73    1.000    0.000
## 74    1.000    0.000
```

```
# Make spacing
single_ktwo_space_cp <- rep(0.2, length(single_ktwo_names_cp))
single_ktwo_space_cp
```

```
##  [1] 0.2 0.2 0.2 0.2 0.2 0.2 0.2 0.2 0.2 0.2 0.2 0.2 0.2 0.2 0.2 0.2 0.2 0.2 0.2
## [20] 0.2 0.2 0.2 0.2 0.2 0.2 0.2 0.2 0.2 0.2 0.2 0.2 0.2 0.2 0.2 0.2 0.2 0.2 0.2
## [39] 0.2 0.2 0.2 0.2 0.2 0.2 0.2 0.2 0.2 0.2 0.2 0.2 0.2 0.2 0.2 0.2 0.2 0.2 0.2
## [58] 0.2 0.2 0.2 0.2 0.2 0.2 0.2 0.2 0.2 0.2 0.2 0.2 0.2 0.2 0.2 0.2 0.2
```

```
# and x coords for those labels
single_ktwo_yx_cp <- c(30, 75, 120)

# Plot
cp_structure_s <- barplot(t(as.matrix(single_ktwo_anc_cp[, 1:2])),
    col = c("#A6D85496", "#00441b"), ylab = "", border = NA,
    main = "", space = single_ktwo_space_cp, names.arg = single_ktwo_names_cp,
    cex.names = 0.5, las = 2)
```

**NOTE:** The results from these analyses are presented in **Fig. S70**.

**13.5.5 Fst estimation between UKH and UKP** Pairwise Fst estimation between *C. parvum* and *C. hominis* by the method of Weir & Cockerham (1984), using the diploid = FALSE to account for haploidity, along the seven chromosomes of *C. parvum* (without considering that some samples harbor mixed infections).

```r
library(hierfstat)
library(qqman)

double_genind <- vcfR2genind(vcf_filtered_samples_double, ploidy = 1)
pop_code_double <- read.gdsn(index.gdsn(genofile_double, "sample.annot/pop.group"))
pop_code_double <- as.factor(pop_code_double)
double_genind$pop <- pop_code_double
Fst_double_hierfstat <- genet.dist(double_genind, diploid = FALSE,
    method = "WC84")
basic_double_hierfstat <- basic.stats(double_genind, diploid = FALSE,
    digits = 2)

# Create dataframe
chrom <- vcf_filtered_samples_double@fix[, "CHROM"]
pos <- vcf_filtered_samples_double@fix[, "POS"]
double_fst_chr <- cbind(chrom, pos, basic_double_hierfstat$perloc[,
    7:8])  #fst and fst prime
dim(double_fst_chr)
```

###### 13.5.5.1 Double-enriched samples

```
## [1] 13578     4
```

```r
colnames(double_fst_chr) <- c("CHROM", "POS", "Fst")
double_fst_chr <- as.data.frame(double_fst_chr)
# Create a factor from the CHROM column, which
# automatically assigns numeric levels
double_fst_chr$CHROM <- as.factor(double_fst_chr$CHROM)
# Convert the factor to numeric (this will map each unique
# chromosome to a numeric value)
double_fst_chr$CHROM <- as.numeric(double_fst_chr$CHROM)
double_fst_chr$POS <- as.numeric(as.character(double_fst_chr$POS))
double_fst_chr$Fst <- as.numeric(as.character(double_fst_chr$Fst))

# Check the transformation
head(double_fst_chr)
```

```
##                 CHROM  POS   Fst    NA
## CP044415_4027     1 4027 -0.32 -0.96
## CP044415_7031     1 7031  0.19  0.32
## CP044415_7040     1 7040  0.19  0.32
## CP044415_7069     1 7069  0.19  0.33
## CP044415_7082     1 7082  0.04  0.07
## CP044415_7102     1 7102  0.19  0.32
```

```r
# plot Manhattan
plot <- manhattan(double_fst_chr, chr = "CHROM", bp = "POS", p = "Fst", snp = "POS",
    logp = FALSE, ylim = c(0, 1), xlab = "Chr", ylab = "Fst", col = c("#A6D854",
        "black"))
```

**NOTE:** The results from these analyses are presented in **Fig. S71**.

```r
single_genind <- vcfR2genind(vcf_filtered_samples_single, ploidy = 1)
pop_code_single <- read.gdsn(index.gdsn(genofile_single, "sample.annot/pop.group"))
pop_code_single <- as.factor(pop_code_single)

single_genind$pop <- pop_code_single
Fst_single_hierfstat <- genet.dist(single_genind, diploid = FALSE, method = "WC84")
basic_single_hierfstat <- basic.stats(single_genind, diploid = FALSE, digits = 2)

# Create dataframe
chrom <- vcf_filtered_samples_single@fix[, "CHROM"]
pos <- vcf_filtered_samples_single@fix[, "POS"]
single_fst_chr <- cbind(chrom, pos, basic_single_hierfstat$perloc[, 7:8])  #fst and fst prime

colnames(single_fst_chr) <- c("CHROM", "POS", "Fst")
single_fst_chr <- as.data.frame(single_fst_chr)
# Create a factor from the CHROM column, which automatically assigns numeric
# levels
single_fst_chr$CHROM <- as.factor(single_fst_chr$CHROM)
# Convert the factor to numeric (this will map each unique chromosome to a
# numeric value)
single_fst_chr$CHROM <- as.numeric(single_fst_chr$CHROM)
single_fst_chr$POS <- as.numeric(as.character(single_fst_chr$POS))
single_fst_chr$Fst <- as.numeric(as.character(single_fst_chr$Fst))

# Check the transformation
head(single_fst_chr)
```

###### 13.5.5.2 Single-enriched samples

```
##                CHROM  POS   Fst    NA
## CP044415_962       1  962 -0.14 -0.32
## CP044415_963       1  963 -0.14 -0.32
## CP044415_970       1  970  1.00  1.00
## CP044415_976       1  976 -0.14 -0.32
## CP044415_1120      1 1120 -0.32 -0.96
## CP044415_4925      1 4925 -0.13 -0.30
```

```r
# plot Manhattan
plot <- manhattan(single_fst_chr, chr = "CHROM", bp = "POS", p = "Fst", snp = "POS",
    logp = FALSE, ylim = c(0, 1), xlab = "Chr", ylab = "Fst", col = c("#A6D854",
        "black"))
```

**NOTE:** The results from these analyses are presented in **Fig. S72**.

```r
unenriched_genind <- vcfR2genind(vcf_biallelic_unenriched, ploidy = 1)
pop_code_unenriched <- read.gdsn(index.gdsn(genofile_unenriched, "sample.annot/pop.group"))
pop_code_unenriched <- as.factor(pop_code_unenriched)
```

```
unenriched_genind$pop <- pop_code_unenriched
Fst_unenriched_hierfstat <- genet.dist(unenriched_genind, diploid = FALSE, method = "WC84")
basic_unenriched_hierfstat <- basic.stats(unenriched_genind, diploid = FALSE, digits = 2)

# Create dataframe
chrom <- vcf_biallelic_unenriched@fix[, "CHROM"]
pos <- vcf_biallelic_unenriched@fix[, "POS"]
unenriched_fst_chr <- cbind(chrom, pos, basic_unenriched_hierfstat$perloc[, 7:8])  # getting fst and fs
dim(unenriched_fst_chr)
```

###### 13.5.5.3 Unenriched samples.

```
## [1] 11226     4
```

```
colnames(unenriched_fst_chr) <- c("CHROM", "POS", "Fst")
unenriched_fst_chr <- as.data.frame(unenriched_fst_chr)
# Create a factor from the CHROM column, which automatically assigns numeric
# levels
unenriched_fst_chr$CHROM <- as.factor(unenriched_fst_chr$CHROM)
# Convert the factor to numeric (this will map each unique chromosome to a
# numeric value)
unenriched_fst_chr$CHROM <- as.numeric(unenriched_fst_chr$CHROM)
unenriched_fst_chr$POS <- as.numeric(as.character(unenriched_fst_chr$POS))
unenriched_fst_chr$Fst <- as.numeric(as.character(unenriched_fst_chr$Fst))

# Check the transformation
head(unenriched_fst_chr)
```

```
##                CHROM  POS Fst  NA
## CP044415_905       1  905   0 NaN
## CP044415_1326      1 1326   0 NaN
## CP044415_3859      1 3859   0 NaN
## CP044415_4413      1 4413   0 NaN
## CP044415_4415      1 4415   0 NaN
## CP044415_4417      1 4417   0 NaN
```

```
# plot Manhattan
plot <- manhattan(unenriched_fst_chr, chr = "CHROM", bp = "POS", p = "Fst", snp = "POS",
    logp = FALSE, ylim = c(0, 1), xlab = "Chr", ylab = "Fst", col = c("#A6D854",
        "black"))
```

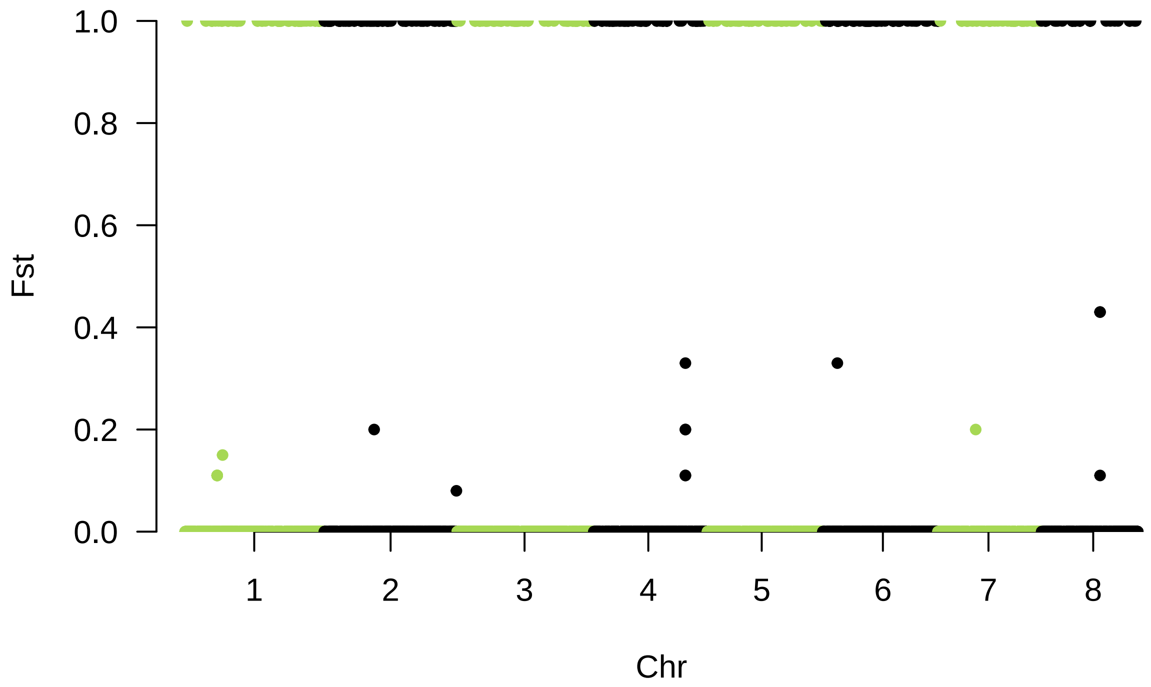

**NOTE:** The results from these analyses are presented in **Fig. S73**.

**13.5.6 Estimation of Nei's genetic distances between and within UKH and UKP** Estimation of Nei's genetic distances, diploid=FALSE, between *C. parvum* and *C. hominis* and all samples. Pairwise genetic distances among samples of *C. parvum* reveal three clusters in double-enriched libraries (UKP347 + [UKP363+UKP336+UKP348+UKP342] + the rest of UKP; Fig. S74), and two clusters in single-enriched libraries ([UKP363+UKP336+UKP348+UKP342] + the rest of UKP; Fig. S75). For *C. hominis*, both capture types reveal that the three samples are highly distinct from each other, even more than the differences shown between clusters of *C. parvum* (Figs. S74–S75). Pairwise distances using unenriched libraries fail to reflect any biologically meaningful pattern (Fig. S76).

```
double_dist <- genet.dist(double_genind, diploid = FALSE, method = "Da")
sample.id <- read.gdsn(index.gdsn(genofile_double, "sample.id"))
samp.annot <- data.frame(pop.group = c("UKH148", "UKH149", "UKH150",
    "UKP301", "UKP302", "UKP303", "UKP304", "UKP305", "UKP306",
    "UKP307", "UKP308", "UKP311", "UKP312", "UKP313", "UKP314",
    "UKP315", "UKP316", "UKP317", "UKP318", "UKP319", "UKP320",
    "UKP322", "UKP323", "UKP324", "UKP325", "UKP326", "UKP327",
    "UKP329", "UKP331", "UKP332", "UKP333", "UKP334", "UKP335",
    "UKP336", "UKP337", "UKP339", "UKP340", "UKP341", "UKP342",
    "UKP343", "UKP347", "UKP348", "UKP349", "UKP351", "UKP356",
    "UKP357", "UKP358", "UKP359", "UKP361", "UKP362", "UKP363",
    "UKP364", "UKP366", "UKP367", "UKP368", "UKP370", "UKP371",
```

```
    "UKP372", "UKP374", "UKP375", "UKP376", "UKP377", "UKP380",
    "UKP382", "UKP383", "UKP384", "UKP385", "UKP387", "UKP388",
    "UKP389", "UKP391", "UKP392", "UKP393", "UKP395A", "UKP398",
    "UKP399", "UKP400"))
add.gdsn(genofile_double, "sample.annot", samp.annot, replace = TRUE)
ind_code_double <- read.gdsn(index.gdsn(genofile_double, "sample.annot/pop.group"))
ind_code_double <- as.factor(ind_code_double)
double_genind <- vcfR2genind(vcf_filtered_samples_double, ploidy = 1)
double_genind@pop <- as.factor(ind_code_double)
double_dist_ind <- genet.dist(double_genind, diploid = FALSE,
    method = "Da")
double_dist_matrix <- as.matrix(double_dist_ind)
```

```
library(pheatmap)
pheatmap(double_dist_matrix, clustering_distance_rows = "euclidean", clustering_distance_cols = "euclid
    clustering_method = "complete", color = colorRampPalette(c("white", "blue"))(100),
    main = "Pairwise Genetic Distances (Da)")
```

**13.5.6.1 Double-enriched samples** **NOTE:** The results from these analyses are presented in **Fig. S74**.

```
single_dist <- genet.dist(single_genind, diploid = FALSE, method = "Da")
sample.id <- read.gdsn(index.gdsn(genofile_single, "sample.id"))
samp.annot <- data.frame(pop.group = c("UKH148", "UKH149", "UKH150", "UKP301", "UKP302",
    "UKP303", "UKP304", "UKP305", "UKP306", "UKP307", "UKP308", "UKP311", "UKP312",
    "UKP313", "UKP314", "UKP315", "UKP316", "UKP317", "UKP318", "UKP319", "UKP320",
    "UKP322", "UKP323", "UKP324", "UKP325", "UKP326", "UKP327", "UKP329", "UKP331",
    "UKP332", "UKP333", "UKP334", "UKP335", "UKP336", "UKP337", "UKP339", "UKP340",
    "UKP341", "UKP342", "UKP343", "UKP347", "UKP348", "UKP349", "UKP351", "UKP356",
    "UKP357", "UKP358", "UKP359", "UKP361", "UKP362", "UKP363", "UKP364", "UKP366",
    "UKP367", "UKP368", "UKP370", "UKP371", "UKP372", "UKP374", "UKP375", "UKP376",
    "UKP377", "UKP380", "UKP382", "UKP383", "UKP384", "UKP385", "UKP387", "UKP388",
    "UKP389", "UKP391", "UKP393", "UKP395A", "UKP398", "UKP399", "UKP400"))
add.gdsn(genofile_single, "sample.annot", samp.annot, replace = TRUE)
ind_code_single <- read.gdsn(index.gdsn(genofile_single, "sample.annot/pop.group"))
ind_code_single <- as.factor(ind_code_single)
single_genind <- vcfR2genind(vcf_filtered_samples_single, ploidy = 1)
single_genind@pop <- as.factor(ind_code_single)
single_dist_ind <- genet.dist(single_genind, diploid = FALSE, method = "Da")
single_dist_matrix <- as.matrix(single_dist_ind)
```

```
pheatmap(single_dist_matrix, clustering_distance_rows = "euclidean", clustering_distance_cols = "euclid
    clustering_method = "complete", color = colorRampPalette(c("white", "blue"))(100),
    main = "Pairwise Genetic Distances (Da)")
```

**13.5.6.2 Single-enriched samples** **NOTE:** The results from these analyses are presented in **Fig. S75**.

```
unenriched_dist <- genet.dist(unenriched_genind, diploid = FALSE, method = "Da")
sample.id <- read.gdsn(index.gdsn(genofile_unenriched, "sample.id"))
samp.annot <- data.frame(pop.group = c("UKH145", "UKH146", "UKH147", "UKH148", "UKH149",
    "UKH150", "UKH151", "UKP301", "UKP302", "UKP303", "UKP304", "UKP305", "UKP306",
    "UKP307", "UKP308", "UKP311", "UKP312", "UKP313", "UKP314", "UKP315", "UKP316",
    "UKP317", "UKP318", "UKP319", "UKP320", "UKP322", "UKP323", "UKP324", "UKP325",
    "UKP326", "UKP327", "UKP329", "UKP331", "UKP332", "UKP333", "UKP334", "UKP335",
    "UKP336", "UKP337", "UKP339", "UKP340", "UKP341", "UKP342", "UKP343", "UKP345",
    "UKP346", "UKP347", "UKP348", "UKP349", "UKP350", "UKP351", "UKP352", "UKP356",
    "UKP357", "UKP358", "UKP359", "UKP360", "UKP361", "UKP362", "UKP363", "UKP364",
    "UKP366", "UKP367", "UKP368", "UKP370", "UKP371", "UKP372", "UKP374", "UKP375",
    "UKP376", "UKP377", "UKP379", "UKP380", "UKP381", "UKP382", "UKP383", "UKP384",
    "UKP385", "UKP386", "UKP387", "UKP388", "UKP389", "UKP390", "UKP391", "UKP392",
    "UKP393", "UKP394", "UKP395A", "UKP398", "UKP399", "UKP400"))
add.gdsn(genofile_unenriched, "sample.annot", samp.annot, replace = TRUE)
ind_code_unenriched <- read.gdsn(index.gdsn(genofile_unenriched, "sample.annot/pop.group"))
ind_code_unenriched <- as.factor(ind_code_unenriched)
unenriched_genind <- vcfR2genind(vcf_biallelic_unenriched, ploidy = 1)
unenriched_genind@pop <- as.factor(ind_code_unenriched)
unenriched_dist_ind <- genet.dist(unenriched_genind, diploid = FALSE, method = "Da")
unenriched_dist_matrix <- as.matrix(unenriched_dist_ind)
```

```
pheatmap(unenriched_dist_matrix, clustering_distance_rows = "euclidean", clustering_distance_cols = "eu
    clustering_method = "complete", color = colorRampPalette(c("white", "blue"))(100),
    main = "Pairwise Genetic Distances (Da)")
```

**13.5.6.3 Unenriched samples** **NOTE:** The results from these analyses are presented in **Fig. S76**.

##### 13.6 Assessment of heterogeneity within samples

**13.6.1 Deconvoluting samples with DEploid v0.7.1** Deconvolution analyses revealed different numbers of inferred strains (inferred K) for double-, single-enriched, and unenriched libraries (Figs. 4A–4C). For double enrichments, 72 samples were detected as having only one strain, and five were detected as having two different strains according to the inferred K; however, the effective K of just three of these samples, UKH149, UKP347, and UKP391, was well above 1.2 (Fig. 4A), and their haplotype proportions for the less dominant haplotype were above 5% (Table S17). For single enrichments, 75 samples were detected to have only one strain. One, UKH149, was detected as having two different strains, but with an effective K of 1.02 (Fig. 4B) and a less dominant haplotype proportion of 1.3% (Table S17). Finally, in unenriched libraries, 61 samples were detected as having only one strain, and 30 were detected as having two different strains, with effective Ks ranging up to 1.96 (Fig. 4C, Table S17). Sample UKH149 was detected as a mixed-infection sample in both of our capture methods, but not with the unenriched libraries.

**13.6.1.1 Generating VCF with a group of samples** We used VCFtools to filter the original VCF, given that exporting from vcfR removes relevant info. We obtained the list of samples and SNPs to keep from corresponding genind objects. Then, imported filtered VCF files here.

```
vcftools_double <- read.vcfR("~/crypto_data/13/DEploid/crypto_double_info.recode.vcf"
```

```
## Scanning file to determine attributes.
## File attributes:
##   meta lines: 49
##   header_line: 50
##   variant count: 13578
##   column count: 86
## Meta line 49 read in.
## All meta lines processed.
## gt matrix initialized.
## Character matrix gt created.
##   Character matrix gt rows: 13578
##   Character matrix gt cols: 86
##   skip: 0
##   nrows: 13578
##   row_num: 0
## Processed variant 1000Processed variant 2000Processed variant 3000Processed variant 4000Processed va
## All variants processed
```

```
vcftools_single <- read.vcfR("~/crypto_data/13/DEploid/crypto_single_info.recode.vcf"
```

```
## Scanning file to determine attributes.
## File attributes:
##   meta lines: 49
##   header_line: 50
##   variant count: 12498
##   column count: 85
## Meta line 49 read in.
## All meta lines processed.
## gt matrix initialized.
## Character matrix gt created.
##   Character matrix gt rows: 12498
##   Character matrix gt cols: 85
##   skip: 0
##   nrows: 12498
##   row_num: 0
## Processed variant 1000Processed variant 2000Processed variant 3000Processed variant 4000Processed va
## All variants processed
```

```
vcftools_unenriched <- read.vcfR("~/crypto_data/13/DEploid/crypto_unenriched_info.recode.vcf"
```

```
## Scanning file to determine attributes.
## File attributes:
##   meta lines: 49
##   header_line: 50
##   variant count: 11226
##   column count: 100
## Meta line 49 read in.
## All meta lines processed.
## gt matrix initialized.
## Character matrix gt created.
##   Character matrix gt rows: 11226
##   Character matrix gt cols: 100
##   skip: 0
```

```
##   nrows: 11226
##   row_num: 0
## Processed variant 1000Processed variant 2000Processed variant 3000Processed variant 4000Processed va
## All variants processed
```

```r
## Add a VQSLOD column to the INFO field
vcftools_double@fix[, "INFO"] <- paste0(vcftools_double@fix[,
    "INFO"], ";VQSLOD=0")
vcftools_single@fix[, "INFO"] <- paste0(vcftools_single@fix[,
    "INFO"], ";VQSLOD=0")
vcftools_unenriched@fix[, "INFO"] <- paste0(vcftools_unenriched@fix[,
    "INFO"], ";VQSLOD=0")

## Because DEploid expects two values in the AD field even
## for variants with missing data. I will:

# Apply to each VCF and write fixed versions
vcf_paths <- list(double = vcftools_double, single = vcftools_single,
    unenriched = vcftools_unenriched)
output_dir <- "~/crypto_data/13/DEploid/testing_haploid/"

fix_missing_ad <- function(vcf_obj) {
    gt_matrix <- vcf_obj@gt
    format_fields <- strsplit(gt_matrix[, "FORMAT"], ":")[[1]]
    ad_index <- which(format_fields == "AD")

    for (i in 2:ncol(gt_matrix)) {
        splitted <- strsplit(gt_matrix[, i], ":")
        for (j in seq_along(splitted)) {
            if (length(splitted[[j]]) >= ad_index && splitted[[j]][ad_index] ==
                ".") {
                splitted[[j]][ad_index] <- "0,0"
            }
        }
        gt_matrix[, i] <- sapply(splitted, paste, collapse = ":")
    }

    vcf_obj@gt <- gt_matrix
    return(vcf_obj)
}

for (name in names(vcf_paths)) {
    vcf_fixed <- fix_missing_ad(vcf_paths[[name]])
    assign(paste0("vcftools_", name, "_edited"), vcf_fixed)  # Save in environment
    output_file <- file.path(output_dir, paste0(name, "_edited.vcf"))
    write.vcf(vcf_fixed, file = output_file)
}
```

**13.6.1.2 Generating VCFs for each sample** Given that the current implementation of DEploid only takes the first sample in the VCF file. I have to create VCFs for each sample.

```r
# Double
sample_names <- colnames(vcftools_double@gt)[-1]  # Exclude FORMAT column
output_dir <- "~/crypto_data/13/DEploid/testing_haploid/double_individual_vcfs" for (sample in sample_na

    # Subset: FORMAT column and one sample
    gt_sub <- vcftools_double_edited@gt[, c("FORMAT", sample),
        drop = FALSE]
    # Create new vcfR object
    vcf_individual <- new("vcfR", meta = vcftools_double@meta,
        fix = vcftools_double_edited@fix, gt = gt_sub)
    # Write to file
    output_file <- file.path(output_dir, paste0(sample, ".vcf"))
    write.vcf(vcf_individual, file = output_file)
}

# Single
sample_names <- colnames(vcftools_single@gt)[-1]  # Exclude FORMAT column
output_dir <- "~/crypto_data/13/DEploid/testing_haploid/single_individual_vcfs" for (sample in sample_na

    # Subset: FORMAT column and one sample
    gt_sub <- vcftools_single_edited@gt[, c("FORMAT", sample),
        drop = FALSE]
    # Create new vcfR object
    vcf_individual <- new("vcfR", meta = vcftools_single_edited@meta,
        fix = vcftools_single_edited@fix, gt = gt_sub)
    # Write to file
    output_file <- file.path(output_dir, paste0(sample, ".vcf"))
    write.vcf(vcf_individual, file = output_file)
}

# Unenriched
sample_names <- colnames(vcftools_unenriched@gt)[-1]  # Exclude FORMAT column
output_dir <- "~/crypto_data/13/DEploid/testing_haploid/unenriched_individual_vcfs" for (sample in sampl

    # Subset: FORMAT column and one sample
    gt_sub <- vcftools_unenriched_edited@gt[, c("FORMAT", sample),
        drop = FALSE]
    # Create new vcfR object
    vcf_individual <- new("vcfR", meta = vcftools_unenriched@meta,
        fix = vcftools_unenriched_edited@fix, gt = gt_sub)
    # Write to file
    output_file <- file.path(output_dir, paste0(sample, ".vcf"))
    write.vcf(vcf_individual, file = output_file)
}
```

**13.6.1.3 Generating Plaf files** Here we estimate allele frequencies from the VCF for each type of library and generate the plaf files for DEploid.

```r
# Extract genotype matrix
geno_double <- extract.gt(vcftools_double, element = "GT")
geno_single <- extract.gt(vcftools_single, element = "GT")
geno_unenriched <- extract.gt(vcftools_unenriched, element = "GT")
```

```r
# Function to compute allele frequencies
compute_AF <- function(gt_matrix) {
    # Create an empty matrix to store allele frequencies
    freq_matrix <- matrix(0, nrow = nrow(gt_matrix), ncol = 2)

    # Loop over each SNP (row) to calculate allele
    # frequencies
    for (i in 1:nrow(gt_matrix)) {
        # Count the number of alleles 0 and 1 for each
        # individual
        allele_counts <- table(unlist(strsplit(gt_matrix[i, ],
            "/")))
        # Store the allele frequencies (percentage of 0s
        # and 1s)
        freq_matrix[i, 1] <- allele_counts["0"]/length(gt_matrix[i,
            ])
        freq_matrix[i, 2] <- allele_counts["1"]/length(gt_matrix[i,
            ])
    }

    return(freq_matrix)
}

# Get allele frequencies
allele_frequencies_double <- compute_AF(geno_double)
allele_frequencies_single <- compute_AF(geno_single)
allele_frequencies_unenriched <- compute_AF(geno_unenriched)

# Extract CHROM and POS information from the VCF
chromosomes_double <- vcftools_double@fix[, "CHROM"]
positions_double <- vcftools_double@fix[, "POS"]
chromosomes_single <- vcftools_single@fix[, "CHROM"]
positions_single <- vcftools_single@fix[, "POS"]
chromosomes_unenriched <- vcftools_unenriched@fix[, "CHROM"]
positions_unenriched <- vcftools_unenriched@fix[, "POS"]

# Create a PLAF file (three columns: CHROM, POS, PLAF)
plaf_double <- data.frame(CHROM = chromosomes_double, POS = positions_double,
    PLAF = allele_frequencies_double[, 2]  # Use the frequency of allele 1
)

plaf_single <- data.frame(CHROM = chromosomes_single, POS = positions_single,
    PLAF = allele_frequencies_single[, 2]  # Use the frequency of allele 1
)

plaf_unenriched <- data.frame(CHROM = chromosomes_unenriched,
    POS = positions_unenriched, PLAF = allele_frequencies_unenriched[,
        2]  # Use the frequency of allele 1
)

# Ensure format
plaf_double$CHROM <- as.character(plaf_double$CHROM)
plaf_double$POS <- as.integer(plaf_double$POS)  # Make sure it's integer
```

```r
plaf_double$PLAF <- as.numeric(plaf_double$PLAF)

plaf_single$CHROM <- as.character(plaf_single$CHROM)
plaf_single$POS <- as.integer(plaf_single$POS)  # Make sure it's integer
plaf_single$PLAF <- as.numeric(plaf_single$PLAF)

plaf_unenriched$CHROM <- as.character(plaf_unenriched$CHROM)
plaf_unenriched$POS <- as.integer(plaf_unenriched$POS)  # Make sure it's integer
plaf_unenriched$PLAF <- as.numeric(plaf_unenriched$PLAF)

# Write the PLAF file to a CSV (you can change the file
# extension if needed) write.csv(plaf_double, file =
# '~/crypto_data/13/DEploid/testing_haploid/plaf_file_double.csv',
# row.names = FALSE)

# write.csv(plaf_single, file =
# '~/crypto_data/13/DEploid/testing_haploid/plaf_file_single.csv',
# row.names = FALSE)

# write.csv(plaf_unenriched, file =
# '~/crypto_data/13/DEploid/testing_haploid/plaf_file_unenriched.csv', # row.names = FALSE)
```

**13.6.1.4 Running DEploid** We used locally the following script using DEploid v0.7.1-beta:

!/bin/bash cd DEploid

Double Enrichments for vcf in double_individual_vcfs/*.vcf; do sample=($basename$"vcf" .vcf) docker run –platform linux/amd64 -v $PWD:/tmp/-w/tmp/ shajoezhu/deploid -$
$vcfdouble_individual_vcfs/${sample}.vcf
-plaf plaf_file_double.csv
-o double_outputs/deploid_${sample}
-noPanel done

Single Enrichments for vcf in single_individual_vcfs/*.vcf; do sample=($basename$"vcf" .vcf) docker run –platform linux/amd64 -v $PWD:/tmp/-w/tmp/ shajoezhu/deploid -$
$vcfsingle_individual_vcfs/${sample}.vcf
-plaf plaf_file_single.csv
-o single_outputs/deploid_${sample}
-noPanel done

Unenriched Samples for vcf in unenriched_individual_vcfs/*.vcf; do sample=($basename$"vcf" .vcf) docker run –platform linux/amd64 -v $PWD:/tmp/-w/tmp/ shajoezhu/deploid -$
$vcfunenriched_individual_vcfs/${sample}.vcf
-plaf plaf_file_unenriched.csv
-o unenriched_outputs/deploid_${sample}
-noPanel done

**13.6.1.5 Graphing effective number of K from DEploid** 13.6.1.5.1 Double enrichments

```r
library(tidyverse)
# Directory where final haplotype files are located
double_log_dir <- "~/crypto_data/13/DEploid/testing_haploid/DEploid_outputs/double_outputs/"

double_log_files <- list.files(double_log_dir, pattern = "^[^.]+\\.log$",
    full.names = TRUE)

extract_k_values <- function(log_file) {
    lines <- readLines(log_file)

    eff_k_line <- grep("Effective_K:", lines, value = TRUE)
    inf_k_line <- grep("Inferred_K:", lines, value = TRUE)

    eff_k <- as.numeric(str_extract(eff_k_line, "[0-9.]+"))
    inf_k <- as.numeric(str_extract(inf_k_line, "[0-9.]+"))

    prop_string <- NA_character_
    prop_idx <- grep("^Proportions:", lines)
    if (length(prop_idx) == 1 && prop_idx < length(lines)) {
        prop_line <- trimws(lines[prop_idx + 1])

        if (nzchar(prop_line)) {
            proportions <- suppressWarnings(as.numeric(str_split(prop_line,
                "\\s+")[[1]]))
            proportions <- proportions[!is.na(proportions) &
                proportions > 0.01]

            if (length(proportions) > 0) {
                prop_string <- paste0(round(proportions, 3),
                  collapse = ", ")
            }
        }
    }

    sample_name <- basename(log_file) %>%
        str_remove("^deploid_") %>%
        str_remove("\\.log$")

    tibble(Sample = sample_name, Effective_K = eff_k, Inferred_K = inf_k,
        Proportions = prop_string)
}

double_k_data <- map_dfr(double_log_files, extract_k_values)
# write.csv(double_k_data, 'double_DEploid_data.csv',
# row.names = FALSE)

double_k_long <- double_k_data %>%
    pivot_longer(cols = c(Effective_K, Inferred_K), names_to = "Type",
        values_to = "K") %>%
    mutate(Symbol = ifelse(Type == "Effective_K", "circle", "x"))

double_k_plot <- plot_ly(double_k_long, x = ~Sample, y = ~K,
    type = "scatter", mode = "markers", symbol = ~Symbol, symbols = c("circle",
```

```
        "x"), color = ~Type, marker = list(size = 10)) %>%
    layout(xaxis = list(title = "Sample", tickangle = -90), yaxis = list(title = "K"),
        legend = list(title = list(text = "K Type")))
```

**NOTE:** The results from these analyses are presented in **Fig. 4A** and **Table S17**.

13.6.1.5.2 Single enrichments

```
# Directory where final haplotype files are located
single_log_dir <- "~/crypto_data/13/DEploid/testing_haploid/DEploid_outputs/single_outputs/"

single_log_files <- list.files(single_log_dir, pattern = "^[^.]+\\.log$", full.names = TRUE)

single_k_data <- map_dfr(single_log_files, extract_k_values)
# write.csv(single_k_data, 'single_DEploid_data.csv', row.names = FALSE)

single_k_long <- single_k_data %>%
    pivot_longer(cols = c(Effective_K, Inferred_K), names_to = "Type", values_to = "K") %>%
    mutate(Symbol = ifelse(Type == "Effective_K", "circle", "x"))

single_k_plot <- plot_ly(single_k_long, x = ~Sample, y = ~K, type = "scatter", mode = "markers",
    symbol = ~Symbol, symbols = c("circle", "x"), color = ~Type, marker = list(size = 10)) %>%
    layout(xaxis = list(title = "Sample", tickangle = -90), yaxis = list(title = "K"),
        legend = list(title = list(text = "K Type")))
```

**NOTE:** The results from these analyses are presented in **Fig. 4B** and **Table S17**.

13.6.1.5.3 Unenriched samples

```
unenriched_log_dir <- "~/crypto_data/13/DEploid/testing_haploid/DEploid_outputs/unenriched_outputs/"

unenriched_log_files <- list.files(unenriched_log_dir, pattern = "^[^.]+\\.log$", full.names = TRUE)

unenriched_k_data <- map_dfr(unenriched_log_files, extract_k_values)
#write.csv(unenriched_k_data, "unenriched_DEploid_data.csv", row.names = FALSE)

unenriched_k_long <- unenriched_k_data %>%
  pivot_longer(cols = c(Effective_K, Inferred_K), names_to = "Type", values_to = "K") %>%
  mutate(Symbol = ifelse(Type == "Effective_K", "circle", "x"))

unenriched_k_plot <- plot_ly(unenriched_k_long, x = ~Sample, y = ~K, type = "scatter", mode = "markers"
        symbol = ~Symbol, symbols = c("circle", "x"),
        color = ~Type, marker = list(size = 10)) %>%#
  layout(xaxis = list(title = "Sample", tickangle = -90),
         yaxis = list(title = "K"),
         legend = list(title = list(text = "K Type")))
```

**NOTE:** The results from these analyses are presented in **Fig. 4C** and **Table S17**.

**13.6.1.6 Combining graphs** 13.6.1.6.1 Double enriched samples

```r
library(ggtree)
```

```
## ggtree v3.14.0 Learn more at https://yulab-smu.top/contribution-tree-data/
##
## Please cite:
##
## Guangchuang Yu. Using ggtree to visualize data on tree-like structures.
## Current Protocols in Bioinformatics. 2020, 69:e96. doi:10.1002/cpbi.96
```

```
##
## Attaching package: 'ggtree'
```

```
## The following object is masked from 'package:ape':
##
##     rotate
```

```
## The following object is masked from 'package:tidyr':
##
##     expand
```

```r
library(ape)
library(tidyverse)
library(ggtreeExtra)
```

```
## ggtreeExtra v1.19.0 For help: https://yulab-smu.top/treedata-book/
##
## If you use the ggtree package suite in published research, please cite
## the appropriate paper(s):
##
## S Xu, Z Dai, P Guo, X Fu, S Liu, L Zhou, W Tang, T Feng, M Chen, L
## Zhan, T Wu, E Hu, Y Jiang, X Bo, G Yu. ggtreeExtra: Compact
## visualization of richly annotated phylogenetic data. Molecular Biology
## and Evolution. 2021, 38(9):4039-4042. doi: 10.1093/molbev/msab166
##
## Guangchuang Yu.  Data Integration, Manipulation and Visualization of
## Phylogenetic Trees (1st edition). Chapman and Hall/CRC. 2022,
## doi:10.1201/9781003279242
##
## Shuangbin Xu, Lin Li, Xiao Luo, Meijun Chen, Wenli Tang, Li Zhan, Zehan
## Dai, Tommy T. Lam, Yi Guan, Guangchuang Yu. Ggtree: A serialized data
## object for visualization of a phylogenetic tree and annotation data.
## iMeta 2022, 1(4):e56. doi:10.1002/imt2.56
```

```r
# convert tree to ggtree
tree <- double_tree

tree_plot <- ggtree(tree, layout = "rectangular")
```

```
## Warning: `aes_()` was deprecated in ggplot2 3.0.0.
## i Please use tidy evaluation idioms with `aes()`
## i The deprecated feature was likely used in the ggtree package.
```

```
##   Please report the issue at <https://github.com/YuLab-SMU/ggtree/issues>.
## This warning is displayed once every 8 hours.
## Call `lifecycle::last_lifecycle_warnings()` to see where this warning was
## generated.

## Warning in fortify(data, ...): Arguments in `...` must be used.
## x Problematic arguments:
## * as.Date = as.Date
## * yscale_mapping = yscale_mapping
## * hang = hang
## i Did you misspell an argument name?
```

```
# Get the vertical tip order (top to bottom in plot)
tree_tips <- tree_plot$data %>%
    filter(isTip) %>%
    arrange(y) %>%
    pull(label)

# convert structure to dataframe
structure_df <- double_ktwo %>%
    select(Label, Anc1 = 5, Anc2 = 6) %>%
    pivot_longer(cols = starts_with("Anc"), names_to = "Cluster",
        values_to = "Proportion")
structure_df$Cluster <- factor(structure_df$Cluster, levels = c("Anc1",
    "Anc2"))
structure_df <- structure_df %>%
    rename(label = Label)
structure_df$label <- factor(structure_df$label, levels = tree_tips)

# Format Deploid K data
k_data_wide <- double_k_data %>%
    rename(Sample = Sample, EffK = Effective_K, InfK = Inferred_K)
k_data_wide <- k_data_wide %>%
    rename(label = Sample)
k_data_wide$label <- factor(k_data_wide$label, levels = tree_tips)
# Effective K bar data
eff_k_df <- k_data_wide %>%
    select(label, EffK) %>%
    mutate(panel = "Effective K")
eff_k_df$label <- factor(eff_k_df$label, levels = tree_tips)
# Inferred K point data
inf_k_df <- k_data_wide %>%
    select(label, InfK) %>%
    mutate(panel = "Inferred K")
inf_k_df$label <- factor(inf_k_df$label, levels = tree_tips)

pop_code_double <- read.gdsn(index.gdsn(genofile_double, "sample.annot/pop.group"))
pop_code_double <- as.factor(pop_code_double)
myColHP <- c(UKH = "#DE2D2696", UKP = "#A6D85496")
tip_colors <- myColHP[as.character(pop_code_double)]
names(tip_colors) <- names(pop_code_double)

pop_df <- data.frame(label = tree$tip.label, pop = pop_code_double)
```

```r
# Plots
p_tree <- ggtree(tree) %<+% pop_df + geom_tippoint(aes(color = pop),
    size = 4) + geom_tiplab(size = 4, hjust = -0.2) + scale_color_manual(values = myColHP) +
    theme_tree2() + theme(legend.position = "none")
```

```
## Warning in fortify(data, ...): Arguments in `...` must be used.
## x Problematic arguments:
## * as.Date = as.Date
## * yscale_mapping = yscale_mapping
## * hang = hang
## i Did you misspell an argument name?
```

```
## Warning: `aes_string()` was deprecated in ggplot2 3.0.0.
## i Please use tidy evaluation idioms with `aes()`.
## i See also `vignette("ggplot2-in-packages")` for more information.
## i The deprecated feature was likely used in the ggtree package.
##   Please report the issue at <https://github.com/YuLab-SMU/ggtree/issues>.
## This warning is displayed once every 8 hours.
## Call `lifecycle::last_lifecycle_warnings()` to see where this warning was
## generated.
```

```r
p_structure <- ggplot(structure_df, aes(x = Proportion, y = label,
    fill = Cluster)) + geom_col(position = "stack", width = 0.8) +
    scale_fill_manual(values = c("#A6D854", "#DE2D26")) + theme_minimal() +
    theme(axis.text.y = element_blank(), axis.title.y = element_blank(),
        axis.ticks.y = element_blank(), axis.title.x = element_text(size = 14),
        axis.text.x = element_text(size = 12), legend.position = "none") +
    labs(x = "Structure K=2")

p_k <- ggplot(k_data_wide, aes(y = label)) + geom_col(aes(x = EffK),
    fill = "gray", width = 0.6) + geom_point(aes(x = InfK), shape = 19,
    size = 5, color = "#2b8cbe") + theme_minimal() + theme(axis.text.y = element_blank(),
    axis.title.y = element_blank(), axis.ticks.y = element_blank(),
    axis.title.x = element_text(size = 14), axis.text.x = element_text(size = 12),
    legend.position = "none") + labs(x = "DEploid K")

library(patchwork)
# All together
double_fig <- p_tree + p_structure + p_k + plot_layout(ncol = 3,
    widths = c(2.5, 0.2, 0.2))
```

**NOTE:** The results from these analyses are presented in **Fig. 4A**.

13.6.1.6.2 Single enriched samples

```r
# convert tree to ggtree
tree <- single_tree

tree_plot <- ggtree(tree, layout = "rectangular")
```

```
## Warning in fortify(data, ...): Arguments in `...` must be used.
## x Problematic arguments:
```

```
## * as.Date = as.Date
## * yscale_mapping = yscale_mapping
## * hang = hang
## i Did you misspell an argument name?
```

```r
# Get the vertical tip order (top to bottom in plot)
tree_tips <- tree_plot$data %>%
    filter(isTip) %>%
    arrange(y) %>%
    pull(label)

# convert structure to dataframe
structure_df <- single_ktwo %>%
    select(Label, Anc1 = 5, Anc2 = 6) %>%
    pivot_longer(cols = starts_with("Anc"), names_to = "Cluster",
        values_to = "Proportion")
structure_df$Cluster <- factor(structure_df$Cluster, levels = c("Anc1",
    "Anc2"))
structure_df <- structure_df %>%
    rename(label = Label)
structure_df$label <- factor(structure_df$label, levels = tree_tips)

# Format Deploid K data
k_data_wide <- single_k_data %>%
    rename(Sample = Sample, EffK = Effective_K, InfK = Inferred_K)
k_data_wide <- k_data_wide %>%
    rename(label = Sample)
k_data_wide$label <- factor(k_data_wide$label, levels = tree_tips)
# Effective K bar data
eff_k_df <- k_data_wide %>%
    select(label, EffK) %>%
    mutate(panel = "Effective K")
eff_k_df$label <- factor(eff_k_df$label, levels = tree_tips)
# Inferred K point data
inf_k_df <- k_data_wide %>%
    select(label, InfK) %>%
    mutate(panel = "Inferred K")
inf_k_df$label <- factor(inf_k_df$label, levels = tree_tips)

pop_code_single <- read.gdsn(index.gdsn(genofile_single, "sample.annot/pop.group"))
pop_code_single <- as.factor(pop_code_single)
myColHP <- c(UKH = "#DE2D2696", UKP = "#A6D85496")
tip_colors <- myColHP[as.character(pop_code_single)]
names(tip_colors) <- names(pop_code_single)

pop_df <- data.frame(label = tree$tip.label, pop = pop_code_single)

# Plots
p_tree <- ggtree(tree) %<+% pop_df + geom_tippoint(aes(color = pop),
    size = 4) + geom_tiplab(size = 4, hjust = -0.2) + scale_color_manual(values = myColHP) +
    theme_tree2() + theme(legend.position = "none")
```

```
## Warning in fortify(data, ...): Arguments in `...` must be used.
## x Problematic arguments:
## * as.Date = as.Date
## * yscale_mapping = yscale_mapping
## * hang = hang
## i Did you misspell an argument name?
```

```
p_structure <- ggplot(structure_df, aes(x = Proportion, y = label,
    fill = Cluster)) + geom_col(position = "stack", width = 0.8) +
    scale_fill_manual(values = c("#A6D854", "#DE2D26")) + theme_minimal() +
    theme(axis.text.y = element_blank(), axis.title.y = element_blank(),
        axis.ticks.y = element_blank(), axis.title.x = element_text(size = 14),
        axis.text.x = element_text(size = 12), legend.position = "none") +
    labs(x = "Structure K=2")

p_k <- ggplot(k_data_wide, aes(y = label)) + geom_col(aes(x = EffK),
    fill = "gray", width = 0.6) + geom_point(aes(x = InfK), shape = 19,
    size = 5, color = "#2b8cbe") + theme_minimal() + theme(axis.text.y = element_blank(),
    axis.title.y = element_blank(), axis.ticks.y = element_blank(),
    axis.title.x = element_text(size = 14), axis.text.x = element_text(size = 12),
    legend.position = "none") + labs(x = "DEploid K")

# All together
single_fig <- p_tree + p_structure + p_k + plot_layout(ncol = 3,
    widths = c(2.5, 0.2, 0.2))
```

**NOTE:** The results from these analyses are presented in **Fig. 4B**.

13.6.1.6.3 Unenriched samples

```
# convert tree to ggtree
tree <- unenriched_tree

tree_plot <- ggtree(tree, layout = "rectangular")
```

```
## Warning in fortify(data, ...): Arguments in `...` must be used.
## x Problematic arguments:
## * as.Date = as.Date
## * yscale_mapping = yscale_mapping
## * hang = hang
## i Did you misspell an argument name?
```

```
# Get the vertical tip order (top to bottom in plot)
tree_tips <- tree_plot$data %>%
    filter(isTip) %>%
    arrange(y) %>%
    pull(label)

# Format Deploid K data
k_data_wide <- unenriched_k_data %>%
    rename(Sample = Sample, EffK = Effective_K, InfK = Inferred_K)
k_data_wide <- k_data_wide %>%
    rename(label = Sample)
```

```r
k_data_wide$label <- factor(k_data_wide$label, levels = tree_tips)
# Effective K bar data
eff_k_df <- k_data_wide %>%
    select(label, EffK) %>%
    mutate(panel = "Effective K")
eff_k_df$label <- factor(eff_k_df$label, levels = tree_tips)
# Inferred K point data
inf_k_df <- k_data_wide %>%
    select(label, InfK) %>%
    mutate(panel = "Inferred K")
inf_k_df$label <- factor(inf_k_df$label, levels = tree_tips)

pop_code_unenriched <- read.gdsn(index.gdsn(genofile_unenriched,
    "sample.annot/pop.group"))
pop_code_unenriched <- as.factor(pop_code_unenriched)
myColHP <- c(UKH = "#DE2D2696", UKP = "#A6D85496")
tip_colors <- myColHP[as.character(pop_code_unenriched)]
names(tip_colors) <- names(pop_code_unenriched)

pop_df <- data.frame(label = tree$tip.label, pop = pop_code_unenriched)

# Plots
p_tree <- ggtree(tree) %<+% pop_df + geom_tippoint(aes(color = pop),
    size = 4) + geom_tiplab(size = 4, hjust = -0.2) + scale_color_manual(values = myColHP) +
    theme_tree2() + theme(legend.position = "none")
```

```
## Warning in fortify(data, ...): Arguments in `...` must be used.
## x Problematic arguments:
## * as.Date = as.Date
## * yscale_mapping = yscale_mapping
## * hang = hang
## i Did you misspell an argument name?
```

```r
p_k <- ggplot(k_data_wide, aes(y = label)) + geom_col(aes(x = EffK),
    fill = "gray", width = 0.6) + geom_point(aes(x = InfK), shape = 19,
    size = 5, color = "#2b8cbe") + theme_minimal() + theme(axis.text.y = element_blank(),
    axis.title.y = element_blank(), axis.ticks.y = element_blank(),
    axis.title.x = element_text(size = 14), axis.text.x = element_text(size = 12),
    legend.position = "none") + labs(x = "DEploid K")

# All together
unenriched_fig <- p_tree + p_k + plot_layout(ncol = 4, widths = c(2.5,
    0.1, 0.5, 0.5))
```

**NOTE:** The results from these analyses are presented in **Fig. 4C**.

##### 13.6.2 Assessing sample depth among libraries

```
library(ggridges)

# Function to sum both allele depths
sum_ad <- function(x) {
    sapply(strsplit(x, ","), function(y) {
        nums <- suppressWarnings(as.numeric(y))
        if (length(nums) == 2 && !any(is.na(nums)))
            sum(nums) else NA
    })
}

# Extract raw AD strings
double_gt_ad_raw <- extract.gt(vcf_filtered_samples_double, element = "AD",
    as.numeric = FALSE)

# Apply to matrix
double_gt_ad_total <- apply(double_gt_ad_raw, 2, sum_ad)

# Convert to long format
double_ad_long <- as.data.frame(double_gt_ad_total) %>%
    rownames_to_column("Locus") %>%
    pivot_longer(-Locus, names_to = "label", values_to = "AD") %>%
    filter(!is.na(AD))

double_ad_long$AD <- as.numeric(double_ad_long$AD)

# Remove big valleys of intermediate values
double_ad_long$jittered_AD <- jitter(double_ad_long$AD, amount = 0.4)  # add random noise ±0.4

# plot
double_AD_ridges <- ggplot(double_ad_long, aes(x = jittered_AD,
    y = label)) + ggridges::geom_density_ridges(scale = 2, fill = "#66C2A5",
    color = "black", alpha = 0.6) + scale_x_continuous(limits = c(0,
    7)) + theme_ridges() + labs(x = "Depth (AD)", y = NULL) +
    theme(axis.title.x = element_text(hjust = 0.5, size = 14)  # hjust = 0.5 centers it
)
```

###### 13.6.2.1 Double enriched samples

```
# Extract raw AD strings
single_gt_ad_raw <- extract.gt(vcf_filtered_samples_single, element = "AD",
    as.numeric = FALSE)

# Apply to matrix
single_gt_ad_total <- apply(single_gt_ad_raw, 2, sum_ad)

# Convert to long format
single_ad_long <- as.data.frame(single_gt_ad_total) %>%
    rownames_to_column("Locus") %>%
```

```r
    pivot_longer(-Locus, names_to = "label", values_to = "AD") %>%
    filter(!is.na(AD))

single_ad_long$AD <- as.numeric(single_ad_long$AD)

# Remove big valleys of intermediate values
single_ad_long$jittered_AD <- jitter(single_ad_long$AD, amount = 0.4)  # add random noise ±0.4

# plot
single_AD_ridges <- ggplot(single_ad_long, aes(x = jittered_AD,
    y = label)) + ggridges::geom_density_ridges(scale = 2, fill = "#FC8D62",
    color = "black", alpha = 0.6) + scale_x_continuous(limits = c(0,
    7)) + theme_ridges() + labs(x = "Depth (AD)", y = NULL) +
    theme(axis.title.x = element_text(hjust = 0.5, size = 14)  # hjust = 0.5 centers it
)
```

###### 13.6.2.2 Single enriched samples

```r
# Extract raw AD strings
unenriched_gt_ad_raw <- extract.gt(vcf_biallelic_unenriched,
    element = "AD", as.numeric = FALSE)

# Apply to matrix
unenriched_gt_ad_total <- apply(unenriched_gt_ad_raw, 2, sum_ad)

# Convert to long format
unenriched_ad_long <- as.data.frame(unenriched_gt_ad_total) %>%
    rownames_to_column("Locus") %>%
    pivot_longer(-Locus, names_to = "label", values_to = "AD") %>%
    filter(!is.na(AD))
unenriched_ad_long$AD <- as.numeric(unenriched_ad_long$AD)

# Remove big valleys of intermediate values
unenriched_ad_long$jittered_AD <- jitter(unenriched_ad_long$AD,
    amount = 0.4)  # add random noise ±0.4

# plot
unenriched_AD_ridges <- ggplot(unenriched_ad_long, aes(x = jittered_AD,
    y = label)) + ggridges::geom_density_ridges(scale = 2, fill = "#8DA0CB",
    color = "black", alpha = 0.6) + scale_x_continuous(limits = c(0,
    7)) + theme_ridges() + labs(x = "Depth (AD)", y = NULL) +
    theme(axis.title.x = element_text(hjust = 0.5, size = 14)  # hjust = 0.5 centers it
)
```

###### 13.6.2.3 Unenriched samples

```r
all_labels <- union(union(unique(double_ad_long$label), unique(single_ad_long$label)),
    unique(unenriched_ad_long$label))
```

```
double_ad_long$label <- factor(double_ad_long$label, levels = all_labels)
single_ad_long$label <- factor(single_ad_long$label, levels = all_labels)
unenriched_ad_long$label <- factor(unenriched_ad_long$label,
    levels = all_labels)

double_AD_ridges <- ggplot(double_ad_long, aes(x = jittered_AD,
    y = label)) + geom_density_ridges(scale = 2, fill = "#66C2A5",
    color = "black", alpha = 0.6) + scale_x_continuous(limits = c(0,
    7)) + theme_ridges() + labs(title = "Double", x = "Depth (AD)",
    y = NULL) + theme(axis.title.x = element_text(hjust = 0.5,
    size = 14))

single_AD_ridges <- ggplot(single_ad_long, aes(x = jittered_AD,
    y = label)) + geom_density_ridges(scale = 2, fill = "#FC8D62",
    color = "black", alpha = 0.6) + scale_x_continuous(limits = c(0,
    7)) + theme_ridges() + labs(title = "Single", x = "Depth (AD)",
    y = NULL) + theme(axis.title.x = element_text(hjust = 0.5,
    size = 14), axis.text.y = element_blank())  # hide y-axis to avoid repetition

unenriched_AD_ridges <- ggplot(unenriched_ad_long, aes(x = jittered_AD,
    y = label)) + geom_density_ridges(scale = 2, fill = "#8DA0CB",
    color = "black", alpha = 0.6) + scale_x_continuous(limits = c(0,
    7)) + theme_ridges() + labs(title = "Unenriched", x = "Depth (AD)",
    y = NULL) + theme(axis.title.x = element_text(hjust = 0.5,
    size = 14), axis.text.y = element_blank())  # only show y-axis on first panel

library(patchwork)
combined_ridges <- double_AD_ridges + single_AD_ridges + unenriched_AD_ridges +
    plot_layout(ncol = 3, widths = c(1.2, 1, 1))
```

###### 13.6.2.4 All library types

**NOTE:** The results from these analyses are presented in **Fig. S77**.

##### 13.6.3 Assessing sample genotype quality among libraries

```
# Extract raw GQ strings
double_gt_gq_raw <- extract.gt(vcf_filtered_samples_double, element = "GQ",
    as.numeric = FALSE)

# Convert to long format
double_gq_long <- as.data.frame(double_gt_gq_raw) %>%
    rownames_to_column("Locus") %>%
    pivot_longer(-Locus, names_to = "label", values_to = "GQ") %>%
    filter(!is.na(GQ))

double_gq_long$GQ <- as.numeric(double_gq_long$GQ)

# Remove big valleys of intermediate values
double_gq_long$jittered_GQ <- jitter(double_gq_long$GQ, amount = 0.9)  # add random noise ±0.4
```

```r
# plot
double_GQ_ridges <- ggplot(double_gq_long, aes(x = jittered_GQ,
    y = label)) + ggridges::geom_density_ridges(scale = 2, fill = "#66C2A5",
    color = "black", alpha = 0.6) + scale_x_continuous(limits = c(-10,
    110)) + theme_ridges() + labs(x = "Genotype Quality (GQ)",
    y = NULL) + theme(axis.title.x = element_text(hjust = 0.5,
    size = 14)  # hjust = 0.5 centers it
)
```

###### 13.6.3.1 Double enriched samples

```r
# Extract raw GQ strings
single_gt_gq_raw <- extract.gt(vcf_filtered_samples_single, element = "GQ",
    as.numeric = FALSE)

# Convert to long format
single_gq_long <- as.data.frame(single_gt_gq_raw) %>%
    rownames_to_column("Locus") %>%
    pivot_longer(-Locus, names_to = "label", values_to = "GQ") %>%
    filter(!is.na(GQ))

single_gq_long$GQ <- as.numeric(single_gq_long$GQ)

# Remove big valleys of intermediate values
single_gq_long$jittered_GQ <- jitter(single_gq_long$GQ, amount = 0.9)  # add random noise ±0.4

# plot
single_GQ_ridges <- ggplot(single_gq_long, aes(x = jittered_GQ,
    y = label)) + ggridges::geom_density_ridges(scale = 2, fill = "#FC8D62",
    color = "black", alpha = 0.6) + scale_x_continuous(limits = c(-10,
    110)) + theme_ridges() + labs(x = "Genotype Quality (GQ)",
    y = NULL) + theme(axis.title.x = element_text(hjust = 0.5,
    size = 14)  # hjust = 0.5 centers it
)
```

###### 13.6.3.2 Single enriched samples

```r
# Extract raw GQ strings
unenriched_gt_gq_raw <- extract.gt(vcf_biallelic_unenriched,
    element = "GQ", as.numeric = FALSE)

# Convert to long format
unenriched_gq_long <- as.data.frame(unenriched_gt_gq_raw) %>%
    rownames_to_column("Locus") %>%
    pivot_longer(-Locus, names_to = "label", values_to = "GQ") %>%
    filter(!is.na(GQ))
```

```
unenriched_gq_long$GQ <- as.numeric(unenriched_gq_long$GQ)

# Remove big valleys of intermediate values
unenriched_gq_long$jittered_GQ <- jitter(unenriched_gq_long$GQ,
    amount = 0.9)  # add random noise ±0.4

# plot
unenriched_GQ_ridges <- ggplot(unenriched_gq_long, aes(x = jittered_GQ,
    y = label)) + ggridges::geom_density_ridges(scale = 2, fill = "#8DA0CB",
    color = "black", alpha = 0.6) + scale_x_continuous(limits = c(-10,
    110)) + theme_ridges() + labs(x = "Genotype Quality (GQ)",
    y = NULL) + theme(axis.title.x = element_text(hjust = 0.5,
    size = 14)  # hjust = 0.5 centers it
)
```

###### 13.6.3.3 Unenriched samples

```
all_labels_gq <- union(union(unique(double_gq_long$label), unique(single_gq_long$label)),
    unique(unenriched_gq_long$label))

double_gq_long$label <- factor(double_gq_long$label, levels = all_labels_gq)
single_gq_long$label <- factor(single_gq_long$label, levels = all_labels_gq)
unenriched_gq_long$label <- factor(unenriched_gq_long$label,
    levels = all_labels_gq)

double_GQ_ridges <- ggplot(double_gq_long, aes(x = jittered_GQ,
    y = label)) + geom_density_ridges(scale = 2, fill = "#66C2A5",
    color = "black", alpha = 0.6) + scale_x_continuous(limits = c(-10,
    110)) + theme_ridges() + labs(title = "Double", x = "GQ",
    y = NULL) + theme(axis.title.x = element_text(hjust = 0.5,
    size = 14))

single_GQ_ridges <- ggplot(single_gq_long, aes(x = jittered_GQ,
    y = label)) + geom_density_ridges(scale = 2, fill = "#FC8D62",
    color = "black", alpha = 0.6) + scale_x_continuous(limits = c(-10,
    110)) + theme_ridges() + labs(title = "Single", x = "GQ",
    y = NULL) + theme(axis.title.x = element_text(hjust = 0.5,
    size = 14), axis.text.y = element_blank())  # hide y-axis to avoid repetition

unenriched_GQ_ridges <- ggplot(unenriched_gq_long, aes(x = jittered_GQ,
    y = label)) + geom_density_ridges(scale = 2, fill = "#8DA0CB",
    color = "black", alpha = 0.6) + scale_x_continuous(limits = c(-10,
    110)) + theme_ridges() + labs(title = "Unenriched", x = "GQ",
    y = NULL) + theme(axis.title.x = element_text(hjust = 0.5,
    size = 14), axis.text.y = element_blank())  # only show y-axis on first panel

combined_ridges_GQ <- double_GQ_ridges + single_GQ_ridges + unenriched_GQ_ridges +
    plot_layout(ncol = 3, widths = c(1.2, 1, 1))
```

**13.6.3.4 All library types** **NOTE:** The results from these analyses are presented in **Fig. S78**.

```r
# Extract chromosome and position
loci_double <- as.data.frame(vcf_filtered_samples_double@fix[,
    c("CHROM", "POS")])
loci_single <- as.data.frame(vcf_filtered_samples_single@fix[,
    c("CHROM", "POS")])

# Now create the unique locus ID
loci_double$Locus <- paste(loci_double$CHROM, loci_double$POS,
    sep = "_")
loci_single$Locus <- paste(loci_single$CHROM, loci_single$POS,
    sep = "_")

# Extract data from FORMAT field for UKH149
gt_double <- extract.gt(vcf_filtered_samples_double, element = "GT",
    as.numeric = FALSE)[, "UKH149"]
ad_double <- extract.gt(vcf_filtered_samples_double, element = "AD",
    as.numeric = FALSE)[, "UKH149"]
gq_double <- extract.gt(vcf_filtered_samples_double, element = "GQ",
    as.numeric = TRUE)[, "UKH149"]

gt_single <- extract.gt(vcf_filtered_samples_single, element = "GT",
    as.numeric = FALSE)[, "UKH149"]
ad_single <- extract.gt(vcf_filtered_samples_single, element = "AD",
    as.numeric = FALSE)[, "UKH149"]
gq_single <- extract.gt(vcf_filtered_samples_single, element = "GQ",
    as.numeric = TRUE)[, "UKH149"]

# Create data frames
df_double <- data.frame(Locus = loci_double$Locus, CHROM = loci_double$CHROM,
    POS = loci_double$POS, GT_double = gt_double, AD_double = ad_double,
    GQ_double = gq_double)

df_single <- data.frame(Locus = loci_single$Locus, CHROM = loci_single$CHROM,
    POS = loci_single$POS, GT_single = gt_single, AD_single = ad_single,
    GQ_single = gq_single)

# Merge based on common loci
comparison_df <- merge(df_double, df_single, by = "Locus", all = TRUE)

# Remove rows where both GTs are NA
comparison_df <- comparison_df %>%
    filter(!(is.na(GT_double) & is.na(GT_single)))

# View where genotype or depth or quality differ
comparison_df_diff <- comparison_df %>%
    filter(GT_double != GT_single | AD_double != AD_single |
        GQ_double != GQ_single)

# Or print a summary
summary(comparison_df_diff)
```

###### 13.6.3.5 Assessing differences in UKH149 for both enrichment types

```
##     Locus              CHROM.x            POS.x             GT_double
##  Length:1092        Length:1092        Length:1092        Length:1092
##  Class :character   Class :character   Class :character   Class :character
##  Mode  :character   Mode  :character   Mode  :character   Mode  :character
##
##
##
##   AD_double            GQ_double        CHROM.y             POS.y
##  Length:1092        Min.   : 0.00   Length:1092        Length:1092
##  Class :character   1st Qu.:45.00   Class :character   Class :character
##  Mode  :character   Median :90.00   Mode  :character   Mode  :character
##                     Mean   :74.41
##                     3rd Qu.:99.00
##                     Max.   :99.00
##   GT_single          AD_single            GQ_single
##  Length:1092        Length:1092        Min.   : 0.00
##  Class :character   Class :character   1st Qu.: 0.00
##  Mode  :character   Mode  :character   Median :42.00
##                                        Mean   :38.42
##                                        3rd Qu.:49.00
##                                        Max.   :99.00
```

```r
# write.csv(comparison_df_diff,
# 'UKH149_double_vs_single_comparison.csv', row.names =
# FALSE)

# Count number of genotype changes Focus only on GT
# differences
gt_diff_df <- comparison_df_diff %>%
    select(Locus, CHROM.x, POS.x, GT_double, GT_single) %>%
    rename(CHROM = CHROM.x, POS = POS.x) %>%
    mutate(GT_double = ifelse(is.na(GT_double), ".", GT_double),
        GT_single = ifelse(is.na(GT_single), ".", GT_single),
        Change = paste(GT_double, "→", GT_single))

# Count all change types from double to single enrichments
gt_change_summary <- gt_diff_df %>%
    count(Change, sort = TRUE)
gt_change_summary
```

```
##   Change   n
## 1  0 → 0 480
## 2  0 → . 290
## 3  0 → 1 121
## 4  1 → 1 103
## 5  1 → .  54
## 6  . → 1  27
## 7  . → 0  12
## 8  1 → 0   5
```

**NOTE:** The results from these analyses are presented in **Table S18**.

```r
# Plot shared SNPs Add CHROM and POS for both VCFs
loci_double <- data.frame(Locus = paste(vcf_filtered_samples_double@fix[,
    "CHROM"], vcf_filtered_samples_double@fix[, "POS"], sep = "_"),
    AD = ad_double, GQ = gq_double)

loci_single <- data.frame(Locus = paste(vcf_filtered_samples_single@fix[,
    "CHROM"], vcf_filtered_samples_single@fix[, "POS"], sep = "_"),
    AD = ad_single, GQ = gq_single)

loci_double$Source <- "Double-enriched"
loci_single$Source <- "Single-enriched"

# Merge datasets on Locus
merged_ad_gq <- inner_join(loci_double, loci_single, by = "Locus",
    suffix = c("_double", "_single"))

# I need to sum AD data: Convert AD to numeric: sum the
# comma-separated values (e.g., '1,0' => 1)
merged_ad_gq <- merged_ad_gq %>%
    mutate(AD_double = as.numeric(sapply(strsplit(AD_double,
        ","), function(x) sum(as.numeric(x)))), AD_single = as.numeric(sapply(strsplit(AD_single,
        ","), function(x) sum(as.numeric(x)))))

# Reshape to long format
ad_gq_df <- merged_ad_gq %>%
    select(Locus, AD_double, GQ_double, AD_single, GQ_single) %>%
    pivot_longer(cols = -Locus, names_to = c("Metric", "Source"),
        names_sep = "_", values_to = "Value") %>%
    pivot_wider(names_from = Metric, values_from = Value)

lsmeans(lm(data = ad_gq_df, AD ~ Source), pairwise ~ Source)
```

```
## $lsmeans
##  Source lsmean     SE   df lower.CL upper.CL
##  double   2.72 0.0221 5291     2.68     2.76
##  single   1.68 0.0352 5291     1.61     1.75
##
## Confidence level used: 0.95
##
## $contrasts
##  contrast        estimate     SE   df t.ratio p.value
##  double - single     1.04 0.0415 5291  25.082  <.0001
```

```r
lsmeans(lm(data = ad_gq_df, GQ ~ Source), pairwise ~ Source)
```

```
## $lsmeans
##  Source lsmean    SE   df lower.CL upper.CL
##  double   68.2 0.518 5393     67.2     69.3
##  single   36.1 0.814 5393     34.5     37.7
##
## Confidence level used: 0.95
```

```
##
## $contrasts
##  contrast        estimate    SE   df t.ratio p.value
##  double - single     32.2 0.965 5393  33.310  <.0001
```

```
# AD boxplot
AD_UKH149 <- plot_ly(ad_gq_df, x = ~Source, y = ~AD, color = ~Source,
    type = "box") %>%
    layout(yaxis = list(title = "Allelic Depth (AD)"))

# GQ boxplot
GQ_UKH149 <- plot_ly(ad_gq_df, x = ~Source, y = ~GQ, color = ~Source,
    type = "box") %>%
    layout(yaxis = list(title = "Genotype Quality (GQ)"))

fig_UKH149 <- subplot(AD_UKH149, GQ_UKH149, nrows = 1, titleY = TRUE,
    titleX = FALSE, margin = 0.1)
```

```
## Warning: Ignoring 4239 observations

## Warning: Ignoring 4137 observations
```

**NOTE:** The results from these analyses are presented in **Fig. S79**.

```
# Extract chromosome and position
loci_double <- as.data.frame(vcf_filtered_samples_double@fix[,
    c("CHROM", "POS")])
loci_single <- as.data.frame(vcf_filtered_samples_single@fix[,
    c("CHROM", "POS")])

# Now create the unique locus ID
loci_double$Locus <- paste(loci_double$CHROM, loci_double$POS,
    sep = "_")
loci_single$Locus <- paste(loci_single$CHROM, loci_single$POS,
    sep = "_")

# Extract data from FORMAT field for UKP347
gt_double <- extract.gt(vcf_filtered_samples_double, element = "GT",
    as.numeric = FALSE)[, "UKP347"]
ad_double <- extract.gt(vcf_filtered_samples_double, element = "AD",
    as.numeric = FALSE)[, "UKP347"]
gq_double <- extract.gt(vcf_filtered_samples_double, element = "GQ",
    as.numeric = TRUE)[, "UKP347"]

gt_single <- extract.gt(vcf_filtered_samples_single, element = "GT",
    as.numeric = FALSE)[, "UKP347"]
ad_single <- extract.gt(vcf_filtered_samples_single, element = "AD",
    as.numeric = FALSE)[, "UKP347"]
gq_single <- extract.gt(vcf_filtered_samples_single, element = "GQ",
```

```r
    as.numeric = TRUE)[, "UKP347"]

# Create data frames
df_double <- data.frame(Locus = loci_double$Locus, CHROM = loci_double$CHROM,
    POS = loci_double$POS, GT_double = gt_double, AD_double = ad_double,
    GQ_double = gq_double)

df_single <- data.frame(Locus = loci_single$Locus, CHROM = loci_single$CHROM,
    POS = loci_single$POS, GT_single = gt_single, AD_single = ad_single,
    GQ_single = gq_single)

# Merge based on common loci
comparison_df <- merge(df_double, df_single, by = "Locus", all = TRUE)

# Remove rows where both GTs are NA
comparison_df <- comparison_df %>%
    filter(!(is.na(GT_double) & is.na(GT_single)))

# View where genotype or depth or quality differ
comparison_df_diff_347 <- comparison_df %>%
    filter(GT_double != GT_single | AD_double != AD_single |
        GQ_double != GQ_single)

# Or print a summary
summary(comparison_df_diff_347)
```

###### 13.6.3.6 Assessing differences in UKP347 for both enrichment types

```
##     Locus              CHROM.x             POS.x             GT_double        
##  Length:3467        Length:3467        Length:3467        Length:3467       
##  Class :character   Class :character   Class :character   Class :character  
##  Mode  :character   Mode  :character   Mode  :character   Mode  :character  
##                                                                             
##                                                                             
##                                                                             
##   AD_double           GQ_double       CHROM.y              POS.y          
##  Length:3467        Min.   : 0.00   Length:3467        Length:3467       
##  Class :character   1st Qu.:84.00   Class :character   Class :character  
##  Mode  :character   Median :99.00   Mode  :character   Mode  :character  
##                     Mean   :84.31                                        
##                     3rd Qu.:99.00                                        
##                     Max.   :99.00                                        
##   GT_single          AD_single            GQ_single   
##  Length:3467        Length:3467        Min.   : 0.0  
##  Class :character   Class :character   1st Qu.:49.0  
##  Mode  :character   Mode  :character   Median :99.0  
##                                        Mean   :78.3  
##                                        3rd Qu.:99.0  
##                                        Max.   :99.0
```

```r
# write.csv(comparison_df_diff_347,
# 'UKP347_double_vs_single_comparison.csv', row.names =
# FALSE)
```

```r
# Count number of genotype changes Focus only on GT
# differences
gt_diff_df_347 <- comparison_df_diff_347 %>%
    select(Locus, CHROM.x, POS.x, GT_double, GT_single) %>%
    rename(CHROM = CHROM.x, POS = POS.x) %>%
    mutate(GT_double = ifelse(is.na(GT_double), ".", GT_double),
        GT_single = ifelse(is.na(GT_single), ".", GT_single),
        Change = paste(GT_double, "→", GT_single))

# Count all change types from double to single enrichments
gt_change_summary_347 <- gt_diff_df_347 %>%
    count(Change, sort = TRUE)
gt_change_summary_347
```

```
##   Change    n
## 1  0 → 0 3103
## 2  0 → .  131
## 3  . → 0  115
## 4  1 → 0   94
## 5  1 → .   13
## 6  0 → 1    8
## 7  1 → 1    2
## 8  . → 1    1
```

**NOTE:** The results from these analyses are presented in **Table S19**.

```r
# Plot shared SNPs Add CHROM and POS for both VCFs
loci_double <- data.frame(Locus = paste(vcf_filtered_samples_double@fix[,
    "CHROM"], vcf_filtered_samples_double@fix[, "POS"], sep = "_"),
    AD = ad_double, GQ = gq_double)

loci_single <- data.frame(Locus = paste(vcf_filtered_samples_single@fix[,
    "CHROM"], vcf_filtered_samples_single@fix[, "POS"], sep = "_"),
    AD = ad_single, GQ = gq_single)

loci_double$Source <- "Double-enriched"
loci_single$Source <- "Single-enriched"

# Merge datasets on Locus
merged_ad_gq <- inner_join(loci_double, loci_single, by = "Locus",
    suffix = c("_double", "_single"))

# I need to sum AD data: Convert AD to numeric: sum the
# comma-separated values (e.g., '1,0' => 1)
merged_ad_gq <- merged_ad_gq %>%
    mutate(AD_double = as.numeric(sapply(strsplit(AD_double,
        ","), function(x) sum(as.numeric(x)))), AD_single = as.numeric(sapply(strsplit(AD_single,
        ","), function(x) sum(as.numeric(x)))))

# Reshape to long format
ad_gq_df <- merged_ad_gq %>%
```

```r
    select(Locus, AD_double, GQ_double, AD_single, GQ_single) %>%
    pivot_longer(cols = -Locus, names_to = c("Metric", "Source"),
        names_sep = "_", values_to = "Value") %>%
    pivot_wider(names_from = Metric, values_from = Value)

lsmeans(lm(data = ad_gq_df, AD ~ Source), pairwise ~ Source)
```

```
## $lsmeans
##  Source lsmean     SE   df lower.CL upper.CL
##  double   4.15 0.0253 8954     4.10      4.2
##  single   3.44 0.0261 8954     3.39      3.5
##
## Confidence level used: 0.95
##
## $contrasts
##  contrast        estimate     SE   df t.ratio p.value
##  double - single    0.703 0.0363 8954  19.350  <.0001
```

```r
lsmeans(lm(data = ad_gq_df, GQ ~ Source), pairwise ~ Source)
```

```
## $lsmeans
##  Source lsmean    SE   df lower.CL upper.CL
##  double   84.1 0.409 9064     83.3     84.9
##  single   80.5 0.422 9064     79.7     81.3
##
## Confidence level used: 0.95
##
## $contrasts
##  contrast        estimate    SE   df t.ratio p.value
##  double - single     3.58 0.588 9064   6.088  <.0001
```

```r
# AD boxplot
AD_UKP347 <- plot_ly(ad_gq_df, x = ~Source, y = ~AD, color = ~Source,
    type = "box") %>%
    layout(yaxis = list(title = "Allelic Depth (AD)"))

# GQ boxplot
GQ_UKP347 <- plot_ly(ad_gq_df, x = ~Source, y = ~GQ, color = ~Source,
    type = "box") %>%
    layout(yaxis = list(title = "Genotype Quality (GQ)"))

fig_UKP347 <- subplot(AD_UKP347, GQ_UKP347, nrows = 1, titleY = TRUE,
    titleX = FALSE, margin = 0.1)
```

```
## Warning: Ignoring 576 observations

## Warning: Ignoring 466 observations
```

**NOTE:** The results from these analyses are presented in **Fig. S80**.

```r
# Extract chromosome and position
loci_double <- as.data.frame(vcf_filtered_samples_double@fix[,
    c("CHROM", "POS")])
loci_single <- as.data.frame(vcf_filtered_samples_single@fix[,
    c("CHROM", "POS")])

# Now create the unique locus ID
loci_double$Locus <- paste(loci_double$CHROM, loci_double$POS,
    sep = "_")
loci_single$Locus <- paste(loci_single$CHROM, loci_single$POS,
    sep = "_")

# Extract data from FORMAT field for UKP347
gt_double <- extract.gt(vcf_filtered_samples_double, element = "GT",
    as.numeric = FALSE)[, "UKP391"]
ad_double <- extract.gt(vcf_filtered_samples_double, element = "AD",
    as.numeric = FALSE)[, "UKP391"]
gq_double <- extract.gt(vcf_filtered_samples_double, element = "GQ",
    as.numeric = TRUE)[, "UKP391"]

gt_single <- extract.gt(vcf_filtered_samples_single, element = "GT",
    as.numeric = FALSE)[, "UKP391"]
ad_single <- extract.gt(vcf_filtered_samples_single, element = "AD",
    as.numeric = FALSE)[, "UKP391"]
gq_single <- extract.gt(vcf_filtered_samples_single, element = "GQ",
    as.numeric = TRUE)[, "UKP391"]

# Create data frames
df_double <- data.frame(Locus = loci_double$Locus, CHROM = loci_double$CHROM,
    POS = loci_double$POS, GT_double = gt_double, AD_double = ad_double,
    GQ_double = gq_double)

df_single <- data.frame(Locus = loci_single$Locus, CHROM = loci_single$CHROM,
    POS = loci_single$POS, GT_single = gt_single, AD_single = ad_single,
    GQ_single = gq_single)

# Merge based on common loci
comparison_df <- merge(df_double, df_single, by = "Locus", all = TRUE)

# Remove rows where both GTs are NA
comparison_df <- comparison_df %>%
    filter(!(is.na(GT_double) & is.na(GT_single)))

# View where genotype or depth or quality differ
comparison_df_diff_391 <- comparison_df %>%
    filter(GT_double != GT_single | AD_double != AD_single |
        GQ_double != GQ_single)

# Or print a summary
summary(comparison_df_diff_391)
```

###### 13.6.3.7 Assessing differences in UKP391 for both enrichment types

```
##     Locus              CHROM.x            POS.x             GT_double
##  Length:2489        Length:2489        Length:2489        Length:2489
##  Class :character   Class :character   Class :character   Class :character
##  Mode  :character   Mode  :character   Mode  :character   Mode  :character
##
##
##
##   AD_double            GQ_double      CHROM.y             POS.y
##  Length:2489        Min.   : 0.0   Length:2489        Length:2489
##  Class :character   1st Qu.:84.0   Class :character   Class :character
##  Mode  :character   Median :99.0   Mode  :character   Mode  :character
##                     Mean   :86.2
##                     3rd Qu.:99.0
##                     Max.   :99.0
##   GT_single           AD_single            GQ_single
##  Length:2489        Length:2489        Min.   : 0.00
##  Class :character   Class :character   1st Qu.:66.00
##  Mode  :character   Mode  :character   Median :91.00
##                                        Mean   :79.89
##                                        3rd Qu.:99.00
##                                        Max.   :99.00
```

```
write.csv(comparison_df_diff_391, "UKP391_double_vs_single_comparison.csv",
    row.names = FALSE)

# Count number of genotype changes Focus only on GT
# differences
gt_diff_df_391 <- comparison_df_diff_391 %>%
    select(Locus, CHROM.x, POS.x, GT_double, GT_single) %>%
    rename(CHROM = CHROM.x, POS = POS.x) %>%
    mutate(GT_double = ifelse(is.na(GT_double), ".", GT_double),
        GT_single = ifelse(is.na(GT_single), ".", GT_single),
        Change = paste(GT_double, "→", GT_single))

# Count all change types from double to single enrichments
gt_change_summary_391 <- gt_diff_df_391 %>%
    count(Change, sort = TRUE)
gt_change_summary_391
```

```
##   Change    n
## 1  0 → 0 2335
## 2  0 → .   86
## 3  . → 0   59
## 4  1 → .    3
## 5  1 → 0    3
## 6  1 → 1    2
## 7  . → 1    1
```

**NOTE:** The results from these analyses are presented in **Table S20**.

```r
# Plot shared SNPs Add CHROM and POS for both VCFs
loci_double <- data.frame(Locus = paste(vcf_filtered_samples_double@fix[,
    "CHROM"], vcf_filtered_samples_double@fix[, "POS"], sep = "_"),
    AD = ad_double, GQ = gq_double)

loci_single <- data.frame(Locus = paste(vcf_filtered_samples_single@fix[,
    "CHROM"], vcf_filtered_samples_single@fix[, "POS"], sep = "_"),
    AD = ad_single, GQ = gq_single)

loci_double$Source <- "Double-enriched"
loci_single$Source <- "Single-enriched"

# Merge datasets on Locus
merged_ad_gq <- inner_join(loci_double, loci_single, by = "Locus",
    suffix = c("_double", "_single"))

# I need to sum AD data: Convert AD to numeric: sum the
# comma-separated values (e.g., '1,0' => 1)
merged_ad_gq <- merged_ad_gq %>%
    mutate(AD_double = as.numeric(sapply(strsplit(AD_double,
        ","), function(x) sum(as.numeric(x)))), AD_single = as.numeric(sapply(strsplit(AD_single,
        ","), function(x) sum(as.numeric(x)))))

# Reshape to long format
ad_gq_df <- merged_ad_gq %>%
    select(Locus, AD_double, GQ_double, AD_single, GQ_single) %>%
    pivot_longer(cols = -Locus, names_to = c("Metric", "Source"),
        names_sep = "_", values_to = "Value") %>%
    pivot_wider(names_from = Metric, values_from = Value)

lsmeans(lm(data = ad_gq_df, AD ~ Source), pairwise ~ Source)
```

```
## $lsmeans
##  Source lsmean     SE   df lower.CL upper.CL
##  double   3.15 0.0193 8865     3.11     3.18
##  single   3.13 0.0195 8865     3.09     3.17
##
## Confidence level used: 0.95
##
## $contrasts
##  contrast        estimate     SE   df t.ratio p.value
##  double - single   0.0129 0.0274 8865   0.469  0.6390
```

```r
lsmeans(lm(data = ad_gq_df, GQ ~ Source), pairwise ~ Source)
```

```
## $lsmeans
##  Source lsmean    SE   df lower.CL upper.CL
##  double   83.8 0.402 8946     83.0     84.6
##  single   80.4 0.405 8946     79.6     81.2
##
## Confidence level used: 0.95
```

```
##
## $contrasts
##  contrast        estimate    SE   df t.ratio p.value
##  double - single     3.41 0.571 8946   5.962  <.0001
```

```
# AD boxplot
AD_UKP391 <- plot_ly(ad_gq_df, x = ~Source, y = ~AD, color = ~Source,
    type = "box") %>%
    layout(yaxis = list(title = "Allelic Depth (AD)"))

# GQ boxplot
GQ_UKP391 <- plot_ly(ad_gq_df, x = ~Source, y = ~GQ, color = ~Source,
    type = "box") %>%
    layout(yaxis = list(title = "Genotype Quality (GQ)"))

fig_UKP391 <- subplot(AD_UKP391, GQ_UKP391, nrows = 1, titleY = TRUE,
    titleX = FALSE, margin = 0.1)
```

```
## Warning: Ignoring 665 observations
```

```
## Warning: Ignoring 584 observations
```

**NOTE:** The results from these analyses are presented in **Fig. S81**.

###### 13.6.4 Assessing levels of nucleotidic diversity among samples

Here, we we run VCFtools 0.1.16 on VCFs filtered to contain only UKH *C. hominis* or UKP *C. parvum* samples. Here, we are using the VCFs originated from GATK, with parameters optimized for diploid organisms filtered for samples and SNPs that fit our filtered database. The samples and SNPs here considered are fewer than in the previous (polidy =1) filtered database, but all present in the previous database.

```
# check vcfs used
pi_vcf_CH_double <- read.vcfR("~/crypto_data/13/nucleotide_diversity/C_hominis_double.recode.vcf"
```

###### 13.6.4.1 Double and single enriched samples

```
## Scanning file to determine attributes.
## File attributes:
##   meta lines: 52
##   header_line: 53
##   variant count: 13444
##   column count: 12
## Meta line 52 read in.
## All meta lines processed.
## gt matrix initialized.
## Character matrix gt created.
##   Character matrix gt rows: 13444
##   Character matrix gt cols: 12
##   skip: 0
##   nrows: 13444
```

```
##   row_num: 0
## Processed variant 1000Processed variant 2000Processed variant 3000Processed variant 4000Processed va
## All variants processed
```

```r
pi_vcf_CH_double
```

```
## ***** Object of Class vcfR *****
## 3 samples
## 8 CHROMs
## 13,444 variants
## Object size: 2.9 Mb
## 0 percent missing data
## *****        *****         *****
```

```r
pi_vcf_CH_single <- read.vcfR("~/crypto_data/13/nucleotide_diversity/C_hominis_single.recode.vcf"
```

```
## Scanning file to determine attributes.
## File attributes:
##   meta lines: 52
##   header_line: 53
##   variant count: 12402
##   column count: 12
## Meta line 52 read in.
## All meta lines processed.
## gt matrix initialized.
## Character matrix gt created.
##   Character matrix gt rows: 12402
##   Character matrix gt cols: 12
##   skip: 0
##   nrows: 12402
##   row_num: 0
## Processed variant 1000Processed variant 2000Processed variant 3000Processed variant 4000Processed va
## All variants processed
```

```r
pi_vcf_CH_single
```

```
## ***** Object of Class vcfR *****
## 3 samples
## 8 CHROMs
## 12,402 variants
## Object size: 2.6 Mb
## 0 percent missing data
## *****        *****         *****
```

```r
pi_vcf_CP_double <- read.vcfR("~/Documents/crypto_data/13/nucleotide_diversity/
C_parvum_double.recode.vcf"
```

```
## Scanning file to determine attributes.
## File attributes:
##   meta lines: 52
##   header_line: 53
##   variant count: 13444
```

```
##   column count: 82
## Meta line 52 read in.
## All meta lines processed.
## gt matrix initialized.
## Character matrix gt created.
##   Character matrix gt rows: 13444
##   Character matrix gt cols: 82
##   skip: 0
##   nrows: 13444
##   row_num: 0
## Processed variant 1000Processed variant 2000Processed variant 3000Processed variant 4000Processed va
## All variants processed
```

```
pi_vcf_CP_double
```

```
## ***** Object of Class vcfR *****
## 73 samples
## 8 CHROMs
## 13,444 variants
## Object size: 11.7 Mb
## 0 percent missing data
## *****        *****         *****
```

```
pi_vcf_CP_single <- read.vcfR("~/crypto_data/13/nucleotide_diversity/C_parvum_single.recode.vcf"
```

```
## Scanning file to determine attributes.
## File attributes:
##   meta lines: 52
##   header_line: 53
##   variant count: 12402
##   column count: 82
## Meta line 52 read in.
## All meta lines processed.
## gt matrix initialized.
## Character matrix gt created.
##   Character matrix gt rows: 12402
##   Character matrix gt cols: 82
##   skip: 0
##   nrows: 12402
##   row_num: 0
## Processed variant 1000Processed variant 2000Processed variant 3000Processed variant 4000Processed va
## All variants processed
```

```
pi_vcf_CP_single
```

```
## ***** Object of Class vcfR *****
## 73 samples
## 8 CHROMs
## 12,402 variants
## Object size: 10.5 Mb
## 0 percent missing data
## *****        *****         *****
```

```r
# Obtain data
CH_pi_double <- read.table("~/crypto_data/13/nucleotide_diversity/pi_results_maf_CH_double.sites.pi"
    header = TRUE)
CP_pi_double <- read.table("~/crypto_data/13/nucleotide_diversity/pi_results_maf_CP_double.sites.pi"
    header = TRUE)
CH_pi_single <- read.table("~/crypto_data/13/nucleotide_diversity/pi_results_maf_CH_single.sites.pi"
    header = TRUE)
CP_pi_single <- read.table("~/crypto_data/13/nucleotide_diversity/pi_results_maf_CP_single.sites.pi"
    header = TRUE)

# Create data frames for each category
pi_CH_double <- data.frame(Nucleotide_Diversity = CH_pi_double$PI, Source = "Double",
    Species = "C_hominis")
pi_CP_double <- data.frame(Nucleotide_Diversity = CP_pi_double$PI, Source = "Double",
    Species = "C_parvum")
pi_CH_single <- data.frame(Nucleotide_Diversity = CH_pi_single$PI, Source = "Single",
    Species = "C_hominis")
pi_CP_single <- data.frame(Nucleotide_Diversity = CP_pi_single$PI, Source = "Single",
    Species = "C_parvum")

# Combine the data frames using rbind
all_pi <- rbind(pi_CH_double, pi_CP_double, pi_CH_single, pi_CP_single)
double_pi <- rbind(pi_CH_double, pi_CP_double)
single_pi <- rbind(pi_CH_single, pi_CP_single)
CH_pi <- rbind(pi_CH_single, pi_CH_double)
CP_pi <- rbind(pi_CP_single, pi_CP_double)

# Check the structure of the combined data frame to ensure it is correct
str(all_pi)
```

```
## 'data.frame':    51691 obs. of  3 variables:
##  $ Nucleotide_Diversity: num  0 0.667 0.667 0.667 0 ...
##  $ Source              : chr  "Double" "Double" "Double" "Double" ...
##  $ Species             : chr  "C_hominis" "C_hominis" "C_hominis" "C_hominis" ...
```

```r
lsmeans(lm(data = all_pi, Nucleotide_Diversity ~ Source), pairwise ~ Source)
```

```
## $lsmeans
##  Source lsmean      SE    df lower.CL upper.CL
##  Double 0.1734 0.00138 49855   0.1707   0.1761
##  Single 0.0379 0.00146 49855   0.0351   0.0408
##
## Confidence level used: 0.95
##
## $contrasts
##  contrast        estimate      SE    df t.ratio p.value
##  Double - Single    0.135 0.00201 49855  67.459  <.0001
```

```r
lsmeans(lm(data = all_pi, Nucleotide_Diversity ~ Species), pairwise ~ Species)
```

```
## $lsmeans
```

```
##   Species   lsmean      SE    df lower.CL upper.CL
##   C_hominis 0.2102 0.00137 49855    0.207   0.2129
##   C_parvum  0.0156 0.00132 49855    0.013   0.0182
##
## Confidence level used: 0.95
##
## $contrasts
##   contrast              estimate      SE    df t.ratio p.value
##   C_hominis - C_parvum     0.195 0.00191 49855 102.077  <.0001
```

```
lsmeans(lm(data = CH_pi, Nucleotide_Diversity ~ Source), pairwise ~ Source)
```

```
## $lsmeans
##   Source lsmean      SE    df lower.CL upper.CL
##   Double 0.3355 0.00241 24010   0.3308   0.3402
##   Single 0.0664 0.00258 24010   0.0613   0.0715
##
## Confidence level used: 0.95
##
## $contrasts
##   contrast        estimate      SE    df t.ratio p.value
##   Double - Single    0.269 0.00354 24010  76.116  <.0001
```

```
lsmeans(lm(data = CP_pi, Nucleotide_Diversity ~ Source), pairwise ~ Source)
```

```
## $lsmeans
##   Source lsmean       SE    df lower.CL upper.CL
##   Double 0.0187 0.000287 25843   0.0181   0.0192
##   Single 0.0122 0.000299 25843   0.0117   0.0128
##
## Confidence level used: 0.95
##
## $contrasts
##   contrast        estimate       SE    df t.ratio p.value
##   Double - Single  0.00642 0.000415 25843  15.460  <.0001
```

```
lsmeans(lm(data = double_pi, Nucleotide_Diversity ~ Species), pairwise ~ Species)
```

```
## $lsmeans
##   Species   lsmean      SE    df lower.CL upper.CL
##   C_hominis 0.3355 0.00202 26269   0.3315   0.3395
##   C_parvum  0.0187 0.00197 26269   0.0148   0.0225
##
## Confidence level used: 0.95
##
## $contrasts
##   contrast              estimate      SE    df t.ratio p.value
##   C_hominis - C_parvum     0.317 0.00282 26269 112.273  <.0001
```

```
lsmeans(lm(data = single_pi, Nucleotide_Diversity ~ Species), pairwise ~ Species)
```

```
## $lsmeans
##  Species   lsmean      SE    df lower.CL upper.CL
##  C_hominis 0.0664 0.00130 23584  0.06384   0.0690
##  C_parvum  0.0122 0.00124 23584  0.00982   0.0147
##
## Confidence level used: 0.95
##
## $contrasts
##  contrast             estimate     SE    df t.ratio p.value
##  C_hominis - C_parvum   0.0541 0.0018 23584  30.107  <.0001
```

```
# Create boxplot for double
double_pi_plot <- plotly::plot_ly()
double_pi_plot <- plotly::add_boxplot(double_pi_plot, data = all_pi, x = ~Species,
    y = ~Nucleotide_Diversity, type = "box", color = ~Species, colors = c("#DE2D2696",
        "#A6D85496"), boxpoints = "all")

# Create boxplot for single
single_pi_plot <- plotly::plot_ly()
single_pi_plot <- plotly::add_boxplot(single_pi_plot, data = all_pi, x = ~Species,
    y = ~Nucleotide_Diversity, type = "box", color = ~Species, colors = c("#DE2D2696",
        "#A6D85496"), boxpoints = "all")

# Create boxplot for CH
CH_pi_plot <- plotly::plot_ly()
CH_pi_plot <- plotly::add_boxplot(CH_pi_plot, data = CH_pi, x = ~Source, y = ~Nucleotide_Diversity,
    type = "box", color = ~Source, colors = c("#FC8D62", "#66C2A5"), boxpoints = "all")

# Create boxplot for CP
CP_pi_plot <- plotly::plot_ly()
CP_pi_plot <- plotly::add_boxplot(CP_pi_plot, data = CP_pi, x = ~Source, y = ~Nucleotide_Diversity,
    type = "box", color = ~Source, colors = c("#FC8D62", "#66C2A5"), boxpoints = "all")

fig_pi <- subplot(CH_pi_plot, CP_pi_plot, nrows = 1, titleY = TRUE, titleX = FALSE,
    margin = 0.1)
```

```
## Warning: Ignoring 1834 observations
```

```
fig_pi
```

```
## Assessing UKH without UKH149
pi_vcf_CH_double_noUKH149 <- read.vcfR("~/crypto_data/13/nucleotide_diversity/
C_hominis_double_noUKH149.recode.vcf"
```

```
## Scanning file to determine attributes.
## File attributes:
##   meta lines: 52
##   header_line: 53
##   variant count: 13444
##   column count: 11
## Meta line 52 read in.
## All meta lines processed.
## gt matrix initialized.
```

```
## Character matrix gt created.
##   Character matrix gt rows: 13444
##   Character matrix gt cols: 11
##   skip: 0
##   nrows: 13444
##   row_num: 0
## Processed variant 1000Processed variant 2000Processed variant 3000Processed variant 4000Processed va
## All variants processed
```

```
pi_vcf_CH_double_noUKH149
```

```
## ***** Object of Class vcfR *****
## 2 samples
## 8 CHROMs
## 13,444 variants
## Object size: 2.7 Mb
## 0 percent missing data
## *****        *****         *****
```

```
pi_vcf_CH_single_noUKH149 <- read.vcfR("~/crypto_data/13/nucleotide_diversity/
C_hominis_single_noUKH149.recode.vcf"
```

```
## Scanning file to determine attributes.
## File attributes:
##   meta lines: 52
##   header_line: 53
##   variant count: 12402
##   column count: 11
## Meta line 52 read in.
## All meta lines processed.
## gt matrix initialized.
## Character matrix gt created.
##   Character matrix gt rows: 12402
##   Character matrix gt cols: 11
##   skip: 0
##   nrows: 12402
##   row_num: 0
## Processed variant 1000Processed variant 2000Processed variant 3000Processed variant 4000Processed va
## All variants processed
```

```
pi_vcf_CH_single_noUKH149
```

```
## ***** Object of Class vcfR *****
## 2 samples
## 8 CHROMs
## 12,402 variants
## Object size: 2.4 Mb
## 0 percent missing data
## *****        *****         *****
```

```
# Obtain data
CH_pi_double_noUKH149 <- read.table("~/crypto_data/13/nucleotide_diversity/
pi_results_maf_CH_double_noUKH149.sites.pi"
```

```r
    header = TRUE)
CH_pi_single_noUKH149 <- read.table("~/crypto_data/13/nucleotide_diversity/
pi_results_maf_CH_single_noUKH149.sites.pi"
    header = TRUE)

# Create data frames for each category
pi_CH_double_noUKH149 <- data.frame(Nucleotide_Diversity = CH_pi_double_noUKH149$PI,
    Source = "Double", Species = "C_hominis")
pi_CH_single_noUKH149 <- data.frame(Nucleotide_Diversity = CH_pi_single_noUKH149$PI,
Source = "Single", Species = "C_hominis")
# Combine the data frames using rbind
CH_pi_noUKH149 <- rbind(pi_CH_single_noUKH149, pi_CH_double_noUKH149)

lsmeans(lm(data = CH_pi_noUKH149, Nucleotide_Diversity ~ Source), pairwise ~ Source)
```

```
## $lsmeans
##  Source  lsmean       SE    df lower.CL upper.CL
##  Double 0.02037 0.000976 22220  0.01846  0.02229
##  Single 0.00499 0.001020 22220  0.00299  0.00699
##
## Confidence level used: 0.95
##
## $contrasts
##  contrast        estimate      SE    df t.ratio p.value
##  Double - Single   0.0154 0.00141 22220  10.895  <.0001
```

```r
# Create boxplot for CH
CH_pi_plot_noUKH149 <- plotly::plot_ly()
CH_pi_plot_noUKH149 <- plotly::add_boxplot(CH_pi_plot_noUKH149, data = CH_pi_noUKH149,
    x = ~Source, y = ~Nucleotide_Diversity, type = "box", color = ~Source, colors = c("#FC8D62",
        "#66C2A5"), boxpoints = "all")

all_pi_noUKH149 <- rbind(pi_CH_double_noUKH149, pi_CP_double, pi_CH_single_noUKH149,
    pi_CP_single)
lsmeans(lm(data = all_pi_noUKH149, Nucleotide_Diversity ~ Species), pairwise ~ Species)
```

```
## $lsmeans
##  Species   lsmean       SE    df lower.CL upper.CL
##  C_hominis 0.0130 0.000508 48065   0.0120   0.0140
##  C_parvum  0.0156 0.000471 48065   0.0147   0.0165
##
## Confidence level used: 0.95
##
## $contrasts
##  contrast            estimate       SE    df t.ratio p.value
##  C_hominis - C_parvum -0.00256 0.000693 48065  -3.690  0.0002
```

**NOTE:** The results from these analyses are presented in **Fig. S82**.

**13.6.5 Assessing levels of within samples variation** Here, we are using the VCFs originated from GATK, with parameters optimized for diploid organisms, allowing us to estimate levels of variation per sample.

```r
vcf_double_diploid <- read.vcfR("~/crypto_data/13/nucleotide_diversity/
crypto_new_double.recode.vcf"
```

###### 13.6.5.1 Transforming double-enriched VCF

```
## Scanning file to determine attributes.
## File attributes:
##   meta lines: 52
##   header_line: 53
##   variant count: 13444
##   column count: 85
## Meta line 52 read in.
## All meta lines processed.
## gt matrix initialized.
## Character matrix gt created.
##   Character matrix gt rows: 13444
##   Character matrix gt cols: 85
##   skip: 0
##   nrows: 13444
##   row_num: 0
## Processed variant 1000Processed variant 2000Processed variant 3000Processed variant 4000Processed va
## All variants processed
```

```r
# Transforming to genind
double_genind <- vcfR2genind(vcf_double_diploid, ploidy = 2)
gl_double <- gi2gl(double_genind, parallel = FALSE, verbose = NULL)
```

```
## Starting gi2gl
## Starting gl.compliance.check
##   Processing genlight object with SNP data
##   The slot loc.all, which stores allele name for each locus, is empty.
## Creating a dummy variable (A/C) to insert in this slot.
##   Checking coding of SNPs
##     SNP data scored NA, 0, 1 or 2 confirmed
##   Checking locus metrics and flags
##   Recalculating locus metrics
##   Checking for monomorphic loci
##     Dataset contains monomorphic loci
##   Checking for loci with all missing data
##     No loci with all missing data detected
##   Checking whether individual names are unique.
##   Checking for individual metrics
##   Warning: Creating a slot for individual metrics
##   Checking for population assignments
##   Population assignments not detected, individuals assigned
##                   to a single population labelled 'pop1'
##   Spelling of coordinates checked and changed if necessary to
##           lat/lon
## Completed: gl.compliance.check
## Completed: gi2gl
```

```r
# removing monomorphic SNPs, 400
gl_double <- gl.filter.monomorphs(gl_double, verbose = 3)
```

```
## Starting gl.filter.monomorphs
##   Processing genlight object with SNP data
##   Identifying monomorphic loci
##   Removing monomorphic loci and loci with all missing
##                          data
##     Original No. of loci: 13444
##     Monomorphic loci: 400
##     Loci scored all NA: 0
##     No. of loci deleted: 400
##     No. of loci retained: 13044
##     No. of individuals: 76
##     No. of populations: 1
## Completed: gl.filter.monomorphs
```

```r
double_genind <- gl2gi(gl_double, verbose = NULL)
```

```
## Starting gl2gi
##   Processing genlight object with SNP data
## Matrix converted.. Prepare genind object...
## Completed: gl2gi
```

```r
indNames(double_genind)
```

```
##  [1] "UKH148"  "UKH149"  "UKH150"  "UKP301"  "UKP302"  "UKP303"  "UKP304"
##  [8] "UKP305"  "UKP306"  "UKP307"  "UKP308"  "UKP311"  "UKP312"  "UKP313"
## [15] "UKP314"  "UKP315"  "UKP316"  "UKP317"  "UKP318"  "UKP319"  "UKP320"
## [22] "UKP322"  "UKP323"  "UKP324"  "UKP325"  "UKP326"  "UKP327"  "UKP329"
## [29] "UKP331"  "UKP332"  "UKP333"  "UKP334"  "UKP335"  "UKP336"  "UKP337"
## [36] "UKP339"  "UKP340"  "UKP341"  "UKP342"  "UKP343"  "UKP347"  "UKP348"
## [43] "UKP349"  "UKP351"  "UKP356"  "UKP357"  "UKP358"  "UKP359"  "UKP361"
## [50] "UKP362"  "UKP363"  "UKP364"  "UKP366"  "UKP367"  "UKP368"  "UKP370"
## [57] "UKP371"  "UKP372"  "UKP374"  "UKP375"  "UKP376"  "UKP377"  "UKP382"
## [64] "UKP383"  "UKP384"  "UKP385"  "UKP387"  "UKP388"  "UKP389"  "UKP391"
## [71] "UKP392"  "UKP393"  "UKP395A" "UKP398"  "UKP399"  "UKP400"
```

```r
vcf_single_diploid <- read.vcfR("~/crypto_data/13/nucleotide_diversity/
crypto_new_single.recode.vcf"
```

###### 13.6.5.2 Transforming single-enriched VCF

```
## Scanning file to determine attributes.
## File attributes:
##   meta lines: 52
##   header_line: 53
##   variant count: 12402
##   column count: 85
```

```
## Meta line 52 read in.
## All meta lines processed.
## gt matrix initialized.
## Character matrix gt created.
##   Character matrix gt rows: 12402
##   Character matrix gt cols: 85
##   skip: 0
##   nrows: 12402
##   row_num: 0
## Processed variant 1000Processed variant 2000Processed variant 3000Processed variant 4000Processed va
## All variants processed
```

```
# Transforming to genind
single_genind <- vcfR2genind(vcf_single_diploid, ploidy = 2)
gl_single <- gi2gl(single_genind, parallel = FALSE, verbose = NULL)
```

```
## Starting gi2gl
## Starting gl.compliance.check
##   Processing genlight object with SNP data
##   The slot loc.all, which stores allele name for each locus, is empty.
## Creating a dummy variable (A/C) to insert in this slot.
##   Checking coding of SNPs
##     SNP data scored NA, 0, 1 or 2 confirmed
##   Checking locus metrics and flags
##   Recalculating locus metrics
##   Checking for monomorphic loci
##     Dataset contains monomorphic loci
##   Checking for loci with all missing data
##     No loci with all missing data detected
##   Checking whether individual names are unique.
##   Checking for individual metrics
##   Warning: Creating a slot for individual metrics
##   Checking for population assignments
##   Population assignments not detected, individuals assigned
##                     to a single population labelled 'pop1'
##   Spelling of coordinates checked and changed if necessary to
##             lat/lon
## Completed: gl.compliance.check
## Completed: gi2gl
```

```
# removing monomorphic SNPs, 528
gl_single <- gl.filter.monomorphs(gl_single, verbose = 3)
```

```
## Starting gl.filter.monomorphs
##   Processing genlight object with SNP data
##   Identifying monomorphic loci
##   Removing monomorphic loci and loci with all missing
##                        data
##     Original No. of loci: 12402
##     Monomorphic loci: 528
##     Loci scored all NA: 0
##     No. of loci deleted: 528
##     No. of loci retained: 11874
```

```
##     No. of individuals: 76
##     No. of populations: 1
## Completed: gl.filter.monomorphs
```

```
single_genind <- gl2gi(gl_single, verbose = NULL)
```

```
## Starting gl2gi
##   Processing genlight object with SNP data
## Matrix converted.. Prepare genind object...
## Completed: gl2gi
```

```
indNames(single_genind)
```

```
##  [1] "UKH148"  "UKH149"  "UKH150"  "UKP301"  "UKP302"  "UKP303"  "UKP304"
##  [8] "UKP305"  "UKP306"  "UKP307"  "UKP308"  "UKP311"  "UKP312"  "UKP313"
## [15] "UKP314"  "UKP315"  "UKP316"  "UKP317"  "UKP318"  "UKP319"  "UKP320"
## [22] "UKP322"  "UKP323"  "UKP324"  "UKP325"  "UKP326"  "UKP327"  "UKP329"
## [29] "UKP331"  "UKP332"  "UKP333"  "UKP334"  "UKP335"  "UKP336"  "UKP337"
## [36] "UKP339"  "UKP340"  "UKP341"  "UKP342"  "UKP343"  "UKP347"  "UKP348"
## [43] "UKP349"  "UKP351"  "UKP356"  "UKP357"  "UKP358"  "UKP359"  "UKP361"
## [50] "UKP362"  "UKP363"  "UKP364"  "UKP366"  "UKP367"  "UKP368"  "UKP370"
## [57] "UKP371"  "UKP372"  "UKP374"  "UKP375"  "UKP376"  "UKP377"  "UKP380"
## [64] "UKP382"  "UKP383"  "UKP384"  "UKP385"  "UKP387"  "UKP388"  "UKP389"
## [71] "UKP391"  "UKP393"  "UKP395A" "UKP398"  "UKP399"  "UKP400"
```

```
# FILTER BASED ON MEAN DEPTH Filter variants with DP within
# reasonable range (>=5 to 100)
vcf_unenriched_filtered <- vcf_pass_unenriched[dp_info_unenriched_numeric >=
    5 & dp_info_unenriched_numeric <= 200, ]

# FILTER OUT VARIANTS WITH MORE THAN TWO ALLELES
biallelic_variants <- is_biallelic(vcf_unenriched_filtered)
vcf_biallelic_unenriched <- vcf_unenriched_filtered[biallelic_variants,
    ]
vcf_biallelic_unenriched
```

###### 13.6.5.3 Importing and filtering unenriched VCF

```
## ***** Object of Class vcfR *****
## 91 samples
## 8 CHROMs
## 11,226 variants
## Object size: 11.1 Mb
## 91.01 percent missing data
## *****        *****         *****
```

```
# FILTER LOCI BASED ON MISSING DATA
genotype_data_unenriched <- extract.gt(vcf_biallelic_unenriched,
    element = "GT", as.numeric = FALSE)
```

```r
missing_data_proportion_unenriched <- rowMeans(is.na(genotype_data_unenriched))
# Filter out loci with more than 60% of missing, removes
# all loci!
loci_to_keep_unenriched <- which(missing_data_proportion_unenriched <=
    0.6)

# FILTER SAMPLES BASED ON MISSING DATA Extract the DP
# (Depth of Coverage) values from the INFO column
gt_matrix_unenriched <- extract.gt(vcf_biallelic_unenriched,
    element = "GT", as.numeric = FALSE)
missing_data_per_sample_unenriched <- colMeans(is.na(gt_matrix_unenriched))
# Filter out samples with more than 80% of missing, removes
# all except 2 samples!
samples_to_keep_unenriched <- which(missing_data_per_sample_unenriched <=
    0.8)

# EXLPORING FILTERED VCF Extract the DP (Depth of Coverage)
# values from the INFO column
dp_info_unenriched_target <- extract.info(vcf_biallelic_unenriched,
    element = "DP")
dp_info_unenriched_target_numeric <- as.numeric(dp_info_unenriched_target)
# Extract the DP values from the FORMAT column for each
# individual
dp_format_unenriched_target <- extract.gt(vcf_biallelic_unenriched,
    element = "DP", as.numeric = TRUE)

coverage_unenriched_df_target <- as.data.frame(dp_format_unenriched_target)
coverage_unenriched_df_target <- reshape2::melt(coverage_unenriched_df_target,
    variable.name = "Individual", value.name = "Depth")
```

```
## No id variables; using all as measure variables
```

```r
coverage_unenriched_df_target <- na.omit(coverage_unenriched_df_target)

# EXPLORE IF MONOMORPHIC LOCI ARE PRESENT AFTER FILTERING
# Transforming to genind
unenriched_genind <- vcfR2genind(vcf_biallelic_unenriched, ploidy = 2)
gl_unenriched <- gi2gl(unenriched_genind, parallel = FALSE, verbose = NULL)
```

```
## Starting gi2gl
## Starting gl.compliance.check
##   Processing genlight object with SNP data
##   The slot loc.all, which stores allele name for each locus, is empty.
## Creating a dummy variable (A/C) to insert in this slot.
##   Checking coding of SNPs
##     SNP data scored NA, 0, 1 or 2 confirmed
##   Checking locus metrics and flags
##   Recalculating locus metrics
##   Checking for monomorphic loci
##     No monomorphic loci detected
##   Checking for loci with all missing data
##     No loci with all missing data detected
```

```
##   Checking whether individual names are unique.
##   Checking for individual metrics
##   Warning: Creating a slot for individual metrics
##   Checking for population assignments
##   Population assignments not detected, individuals assigned
##                     to a single population labelled 'pop1'
##   Spelling of coordinates checked and changed if necessary to
##            lat/lon
## Completed: gl.compliance.check
## Completed: gi2gl
```

```
indNames(unenriched_genind)
```

```
##  [1] "UKH145"  "UKH146"  "UKH147"  "UKH148"  "UKH149"  "UKH150"  "UKH151"
##  [8] "UKP301"  "UKP302"  "UKP303"  "UKP304"  "UKP305"  "UKP306"  "UKP307"
## [15] "UKP308"  "UKP311"  "UKP312"  "UKP313"  "UKP314"  "UKP315"  "UKP316"
## [22] "UKP317"  "UKP318"  "UKP319"  "UKP320"  "UKP322"  "UKP323"  "UKP324"
## [29] "UKP325"  "UKP326"  "UKP327"  "UKP329"  "UKP331"  "UKP332"  "UKP333"
## [36] "UKP334"  "UKP335"  "UKP336"  "UKP337"  "UKP339"  "UKP340"  "UKP341"
## [43] "UKP342"  "UKP343"  "UKP345"  "UKP346"  "UKP347"  "UKP348"  "UKP349"
## [50] "UKP350"  "UKP351"  "UKP352"  "UKP356"  "UKP357"  "UKP358"  "UKP359"
## [57] "UKP360"  "UKP361"  "UKP362"  "UKP363"  "UKP364"  "UKP366"  "UKP367"
## [64] "UKP368"  "UKP370"  "UKP371"  "UKP372"  "UKP374"  "UKP375"  "UKP376"
## [71] "UKP377"  "UKP379"  "UKP380"  "UKP381"  "UKP382"  "UKP383"  "UKP384"
## [78] "UKP385"  "UKP386"  "UKP387"  "UKP388"  "UKP389"  "UKP390"  "UKP391"
## [85] "UKP392"  "UKP393"  "UKP394"  "UKP395A" "UKP398"  "UKP399"  "UKP400"
```

```
#Importing strata
double_strata <- read.table("~/crypto_data/13/Heterozygosity/double_ #Add strata to obj
strata(double_genind) <- double_strata

# Summarizing diversity for the whole dataset
## Summarizing expected and observed "heterozygosity"
setPop(double_genind) <- ~Enrichment
double.smry <- summary(double_genind)
mean(double.smry$Hobs)
```

###### 13.6.5.4 Within-sample variation in double enrichments

```
## [1] 0.008738537
```

```
## Summarizing allelic richness
library(PopGenReport)
double_ar <- allel.rich(double_genind) #allele richness
double_ar$mean.richness
```

```
##   Double
## 1.833386
```

```r
double_ar$sum.richness
```

```
##   Double
## 23914.69
```

```r
## Summarizing Fis
library(hierfstat)
double_basicstat<- basic.stats(double_genind, diploid = TRUE, digits = 2)
mean(double_basicstat$Fis[,c(1)], na.rm = TRUE) #Global
```

```
## [1] 0.8346619
```

```r
#Summarizing Diversity for Samples
setPop(double_genind) <- ~Sample
double_genind$pop
```

```
##  [1] UKH148  UKH149  UKH150  UKP301  UKP302  UKP303  UKP304  UKP305  UKP306
## [10] UKP307  UKP308  UKP311  UKP312  UKP313  UKP314  UKP315  UKP316  UKP317
## [19] UKP318  UKP319  UKP320  UKP322  UKP323  UKP324  UKP325  UKP326  UKP327
## [28] UKP329  UKP331  UKP332  UKP333  UKP334  UKP335  UKP336  UKP337  UKP339
## [37] UKP340  UKP341  UKP342  UKP343  UKP347  UKP348  UKP349  UKP351  UKP356
## [46] UKP357  UKP358  UKP359  UKP361  UKP362  UKP363  UKP364  UKP366  UKP367
## [55] UKP368  UKP370  UKP371  UKP372  UKP374  UKP375  UKP376  UKP377  UKP382
## [64] UKP383  UKP384  UKP385  UKP387  UKP388  UKP389  UKP391  UKP392  UKP393
## [73] UKP395A UKP398  UKP399  UKP400
## 76 Levels: UKH148 UKH149 UKH150 UKP301 UKP302 UKP303 UKP304 UKP305 ... UKP400
```

```r
pop.double <- seppop(double_genind)
summary.by.pop = lapply(pop.double, summary)

double_obs_het_df <- data.frame(
  Sample = names(summary.by.pop),
  Ho = sapply(summary.by.pop, function(x) mean(x$Hobs, na.rm = TRUE))
)

double_obs_het_df$color <- ifelse(double_obs_het_df$Sample %in% c("UKH148", "UKH149"),
                           "#DE2D2696",  # highlight color
                           "#A6D85496")  # default color

het_double_plot <- plot_ly(
  data = double_obs_het_df,
  x = ~Sample,
  y = ~Ho,
  type = 'bar',
  marker = list(color = double_obs_het_df$color)
) %>%
  layout(xaxis = list(title = "Sample", tickangle = -90),
    yaxis = list(title = "Mean Observed Within-Sample Variation"),
    margin = list(b = 100),
    showlegend = FALSE
  )
```

**NOTE:** The results from these analyses are presented in **Fig. S83**.

13.6.5.4.1 Detailing within-sample variation of sample UKH149

```
Ho_149 <- summary.by.pop$UKH149$Hobs
loci_149 <- locNames(pop.double$UKH149)

het_df <- data.frame(Locus = loci_149, Ho = Ho_149)

het_df$Chromosome <- sub("_.*", "", het_df$Locus)

mean_Ho_by_chr <- het_df %>%
    group_by(Chromosome) %>%
    summarise(mean_Ho = mean(Ho, na.rm = TRUE)) %>%
    ungroup()

het_double_UKH149 <- plot_ly(data = mean_Ho_by_chr, x = ~Chromosome, y = ~mean_Ho,
    type = "bar", marker = list(color = "#DE2D2696")) %>%
    layout(xaxis = list(title = "Chromosome", tickangle = -90), yaxis = list(title = "Mean Observed Wit
        margin = list(b = 100), showlegend = FALSE)
```

**NOTE:** The results from these analyses are presented in **Fig. S84**.

13.6.5.4.2 Detailing within-sample variation of sample UKP347

```
Ho_347 <- summary.by.pop$UKP347$Hobs
loci_347 <- locNames(pop.double$UKP347)

het_df <- data.frame(Locus = loci_347, Ho = Ho_347)

het_df$Chromosome <- sub("_.*", "", het_df$Locus)

mean_Ho_by_chr <- het_df %>%
    group_by(Chromosome) %>%
    summarise(mean_Ho = mean(Ho, na.rm = TRUE)) %>%
    ungroup()

het_double_UKP347 <- plot_ly(data = mean_Ho_by_chr, x = ~Chromosome, y = ~mean_Ho,
    type = "bar", marker = list(color = "#A6D85496")) %>%
    layout(xaxis = list(title = "Chromosome", tickangle = -90), yaxis = list(title = "Mean Observed Wit
        margin = list(b = 100), showlegend = FALSE)
```

**NOTE:** The results from these analyses are presented in **Fig. S85**.

```
#Importing strata
single_strata <- read.table("~/crypto_data/13/Heterozygosity/single_strata.csv" #Add strata to obj
strata(single_genind) <- single_strata

# Summarizing diversity for the whole dataset
## Summarizing expected and observed "heterozygosity"
```

```r
setPop(single_genind) <- ~Enrichment
single.smry <- summary(single_genind)
mean(single.smry$Hobs)
```

###### 13.6.5.5 Within-sample variation in single enrichments

```
## [1] 0.001782951
```

```r
## Summarizing allelic richness
single_ar <- allel.rich(single_genind) #allele richness
single_ar$mean.richness
```

```
##   Single
## 1.827664
```

```r
single_ar$sum.richness
```

```
##   Single
## 21701.68
```

```r
## Summarizing Fis
library(hierfstat)
single_basicstat<- basic.stats(single_genind, diploid = TRUE, digits = 2)
mean(single_basicstat$Fis[,c(1)], na.rm = TRUE) #Global
```

```
## [1] 0.9662616
```

```r
#Summarizing Diversity for Samples
setPop(single_genind) <- ~Sample
single_genind$pop
```

```
##  [1] UKH148  UKH149  UKH150  UKP301  UKP302  UKP303  UKP304  UKP305  UKP306
## [10] UKP307  UKP308  UKP311  UKP312  UKP313  UKP314  UKP315  UKP316  UKP317
## [19] UKP318  UKP319  UKP320  UKP322  UKP323  UKP324  UKP325  UKP326  UKP327
## [28] UKP329  UKP331  UKP332  UKP333  UKP334  UKP335  UKP336  UKP337  UKP339
## [37] UKP340  UKP341  UKP342  UKP343  UKP347  UKP348  UKP349  UKP351  UKP356
## [46] UKP357  UKP358  UKP359  UKP361  UKP362  UKP363  UKP364  UKP366  UKP367
## [55] UKP368  UKP370  UKP371  UKP372  UKP374  UKP375  UKP376  UKP377  UKP380
## [64] UKP382  UKP383  UKP384  UKP385  UKP387  UKP388  UKP389  UKP391  UKP393
## [73] UKP395A UKP398  UKP399  UKP400
## 76 Levels: UKH148 UKH149 UKH150 UKP301 UKP302 UKP303 UKP304 UKP305 ... UKP400
```

```r
pop.single <- seppop(single_genind)
single_summary.by.pop = lapply(pop.single, summary)

single_obs_het_df <- data.frame(
  Sample = names(single_summary.by.pop),
  Ho = sapply(single_summary.by.pop, function(x) mean(x$Hobs, na.rm = TRUE))
)
```

```r
single_obs_het_df$color <- ifelse(single_obs_het_df$Sample %in% c("UKH148","UKH149","UKH150"),
                            "#DE2D2696",  # highlight color
                            "#A6D85496")  # default color

het_single_plot <- plot_ly(
  data = single_obs_het_df,
  x = ~Sample,
  y = ~Ho,
  type = 'bar',
  marker = list(color = single_obs_het_df$color)
) %>%
  layout(
    xaxis = list(title = "Sample", tickangle = -90),
    yaxis = list(title = "Mean Observed Within-Sample Variation"),
    margin = list(b = 100),
    showlegend = FALSE
  )
```

**NOTE:** The results from these analyses are presented in **Fig. S86**.

13.6.5.5.1 Detailing within-sample variation of sample UKH149

```r
single_Ho_149 <- single_summary.by.pop$UKH149$Hobs
single_loci_149 <- locNames(pop.single$UKH149)

single_het_df <- data.frame(Locus = single_loci_149, Ho = single_Ho_149)

single_het_df$Chromosome <- sub("_.*", "", single_het_df$Locus)

single_mean_Ho_by_chr <- single_het_df %>%
    group_by(Chromosome) %>%
    summarise(mean_Ho = mean(Ho, na.rm = TRUE)) %>%
    ungroup()

het_single_UKH149 <- plot_ly(data = single_mean_Ho_by_chr, x = ~Chromosome, y = ~mean_Ho,
    type = "bar", marker = list(color = "#DE2D2696")) %>%
    layout(xaxis = list(title = "Chromosome", tickangle = -90), yaxis = list(title = "Mean Observed Wit
        margin = list(b = 100), showlegend = FALSE)
```

**NOTE:** The results from these analyses are presented in **Fig. S87**.

```r
#Importing strata
unenriched_strata <- read.table("~/crypto_data/13/Heterozygosity/unenriched_strata.csv" #Add strata to d
strata(unenriched_genind) <- unenriched_strata

# Summarizing diversity for the whole dataset
## Summarizing expected and observed "heterozygosity"
setPop(unenriched_genind) <- ~Enrichment
unenriched.smry <- summary(unenriched_genind)
mean(unenriched.smry$Hobs)
```

###### 13.6.5.6 Within-sample variation in unenriched libraries

```
## [1] 1
```

```
## Summarizing allelic richness
unenriched_ar <- allel.rich(unenriched_genind) #allele richness
unenriched_ar$mean.richness
```

```
## Unenriched
##   1.443103
```

```
unenriched_ar$sum.richness
```

```
## Unenriched
##   16200.27
```

```
## Summarizing Fis
library(hierfstat)
unenriched_basicstat<- basic.stats(unenriched_genind, diploid = TRUE, digits = 2)
mean(unenriched_basicstat$Fis[,c(1)], na.rm = TRUE) #Global
```

```
## [1] -0.03419984
```

```
#Summarizing Diversity for Samples
setPop(unenriched_genind) <- ~Sample
unenriched_genind$pop
```

```
##  [1] UKH145  UKH146  UKH147  UKH148  UKH149  UKH150  UKH151  UKP301  UKP302
## [10] UKP303  UKP304  UKP305  UKP306  UKP307  UKP308  UKP311  UKP312  UKP313
## [19] UKP314  UKP315  UKP316  UKP317  UKP318  UKP319  UKP320  UKP322  UKP323
## [28] UKP324  UKP325  UKP326  UKP327  UKP329  UKP331  UKP332  UKP333  UKP334
## [37] UKP335  UKP336  UKP337  UKP339  UKP340  UKP341  UKP342  UKP343  UKP345
## [46] UKP346  UKP347  UKP348  UKP349  UKP350  UKP351  UKP352  UKP356  UKP357
## [55] UKP358  UKP359  UKP360  UKP361  UKP362  UKP363  UKP364  UKP366  UKP367
## [64] UKP368  UKP370  UKP371  UKP372  UKP374  UKP375  UKP376  UKP377  UKP379
## [73] UKP380  UKP381  UKP382  UKP383  UKP384  UKP385  UKP386  UKP387  UKP388
## [82] UKP389  UKP390  UKP391  UKP392  UKP393  UKP394  UKP395A UKP398  UKP399
## [91] UKP400
## 91 Levels: UKH145 UKH146 UKH147 UKH148 UKH149 UKH150 UKH151 UKP301 ... UKP400
```

```
pop.unenriched <- seppop(unenriched_genind)
unenriched_summary.by.pop = lapply(pop.unenriched, summary)

unenriched_obs_het_df <- data.frame(
  Sample = names(unenriched_summary.by.pop),
  Ho = sapply(unenriched_summary.by.pop, function(x) mean(x$Hobs, na.rm = TRUE))
)

unenriched_obs_het_df$color <- ifelse(unenriched_obs_het_df$Sample %in% c("UKH145","UKH146","UKH147","U
                           "#DE2D2696",  # highlight color
                           "#A6D85496")  # default color
```

```r
het_unenriched_plot <- plot_ly(
  data = unenriched_obs_het_df,
  x = ~Sample,
  y = ~Ho,
  type = 'bar',
  marker = list(color = unenriched_obs_het_df$color)
) %>%
  layout(
    xaxis = list(title = "Sample", tickangle = -90),
    yaxis = list(title = "Mean Observed Within-Sample Variation"),
    margin = list(b = 100),
    showlegend = FALSE
  )
```

**NOTE:** The results from these analyses are presented in **Fig. S88**.

#### Supplementary Notes #14. Cost Reduction in Cryptosporidium Genomics by Enrichment.

Our assay can be implemented in various scenarios, and the cost savings will vary depending on the abundance of *Cryptosporidium* DNA in the sample. If pure *Cryptosporidium* DNA is available, the approximate cost of sequencing to 20X coverage is ~$1 USD (Table S17). However, if the same *Cryptosporidium* DNA is present in a fecal sample with a typical abundance observed here (~0.1%), sequencing costs increase 1000-fold to approximately $1,000 USD. However, if the fecal DNA from the second scenario is enriched to just 10% on-target *Cryptosporidium* DNA reads, the sequencing costs drop to approximately $10 USD, considering costs through sample pooling and bait dilution while still obtaining a large number of sequences. If we add the cost of enrichment, approximately $10 USD, then the total assay cost is approximately $20 USD. This example also clarifies that implementing our assay is economically desirable for any sample where *Cryptosporidium* DNA constitutes  5% of the total DNA, and no human clinical fecal samples we tested had *Cryptosporidium* DNA at an abundance that high. However, this assay requires significant sample volumes to be processed, thereby spreading out the buy-in costs, and our calculations do not account for staff time. However, the enrichment process for a set of samples should take ~ half a day of hands-on time per enrichment. Generally, one technician can process up to 16 pools with 8-12 samples per pool, but most commonly, 8-12 pools (from 96 samples) are processed simultaneously. Thus, double-enrichment would require ~1 day of hands-on technician time, which should be divided by the number of samples in the batch to arrive at the per-sample cost (most commonly, the cost of one technician day/96; at $25/hr, this is ~$2/sample; Table S21).
